# Mouse primary T cell phosphotyrosine proteomics enabled by BOOST

**DOI:** 10.1101/2022.05.13.491817

**Authors:** Xien Yu Chua, Kenneth P. Callahan, Alijah A. Griffith, Tobias Hildebrandt, Guoping Fu, Mengzhou Hu, Renren Wen, Arthur R. Salomon

**Affiliations:** Department of Molecular Pharmacology, Physiology & Biotechnology, Brown University, Providence, RI, 02912; Department of Molecular Biology, Cell Biology & Biochemistry, Brown University, Providence, RI, 02912; Blood Research Institute, Blood Center of Wisconsin, Milwaukee, WI, 53226

**Keywords:** Mice, T cell, BOOST, TMT, SH2 superbinder, Phosphotyrosine proteomics, Phase-constrained spectrum deconvolution method

## Abstract

The Broad Spectrum Optimization of Selective Triggering (BOOST) approach was recently developed to increase the quantitative depth of the tyrosine phosphoproteome by mass spectrometry-based proteomics. While BOOST has been demonstrated in the Jurkat T cell line, it has not been demonstrated in scarce mice primary T cells. Here, we show the first phosphotyrosine proteomics experiment performed in mice primary T cells using BOOST. We identify and precisely quantify more than 2,000 unique pTyr sites from more than 3,000 unique pTyr peptide PSMs using only 1 mg of protein from T cell receptor-stimulated primary T cells from mice. We further reveal the importance of the phase-constrained spectrum deconvolution method (ΦSDM) parameter on Orbitrap instruments that, when disabled, enhances quantitation depth, accuracy, and precision in low-abundance samples. Using samples with contrived ratios, we find that disabling ΦSDM allows for up to a two-fold increase in the number of statistically significant intensity ratios detected while enabling ΦSDM degrades quantitation, especially in low-abundance samples.

**TOC Graphic:** 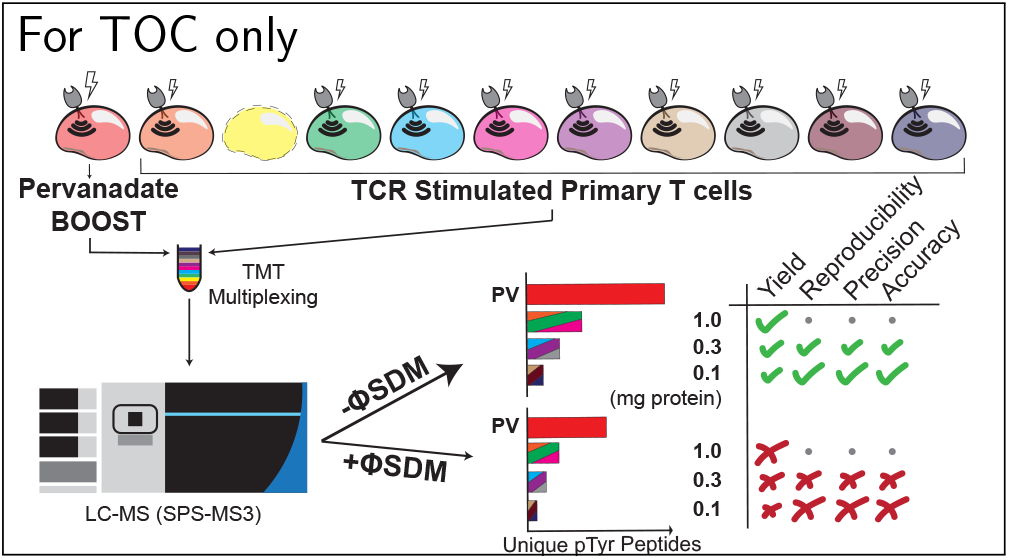

## Introduction

Kinase signaling cascades regulate key cellular processes including growth, differentiation, and transcriptional regulation. In T cells, T cell receptor (TCR) engagement by peptide major histocompatibility complexes on antigen-presenting cells initiates early tyrosine kinase-mediated signaling, leading to serine/threonine kinase activation that regulates transcriptional activation.^1^ Signal initiation from the TCR begins via recruitment of the Src family tyrosine kinases Lck and Fyn, which phosphorylate tyrosine residues in immunotyrosine activation motifs (ITAMs) on the intracellular domain of TCR complex subunits TCR*ζ*, CD3*ϵ*, CD3*δ*, and CD3*γ*.^1,2^ Next, the Syk-family tyrosine kinase Zap70 binds to phosphorylated ITAMs, is itself phosphorylated by TCR-proximal Lck, and then phosphorylates the adaptor proteins linker for activation of T cells (LAT) and SH2 domain containing protein of 76 kDa (SLP76).^3^ Zap70 in combination with the Lck-activated Tec-family tyrosine kinase Itk phosphorylate and activate many LAT-associated proteins, culminating in serine/threonine kinase activation upstream of cytokine expression and actin cytoskeletal regulation. ^1,4^ Despite the importance of tyrosine phosphorylation in the early stages of TCR signaling, tyrosine phosphorylation is scarce, accounting for less than 1% of all phosphorylation events.^5,6^

Due to the scarce nature of tyrosine phosphorylation, large-scale phosphotyrosine (pTyr) proteomic studies of TCR signaling in mice primary T cells are often impaired by low yield. Additionally, the low abundance of tyrosine phosphorylation requires researchers to use a large number of cells per replicate (usually 100 million or more), and therefore a large amount of protein input (around 5 to 10 milligrams), to reproducibly characterize the tyrosine phosphoproteome. Recently, Locard-Paulet et al. performed a pTyr proteomic study of TCR signaling in CD4^+^ T cells by treating 100 million T cells per replicate with *α*-CD3 and *α*-CD4 antibodies before phophotyrosine enrichment using the pTyr-1000 enrichment kit. Using this approach, Locard-Paulet et al. identified a total of 254 unique pTyr sites from 786 unique pTyr peptides, which is comparable to phosphoproteomic studies of TCR and chimeric antigen receptor signaling in primary human T cells. ^8–10^ Phosphotyrosine-specific enrichment methods provide improved pTyr sequencing compared to the more commonly used global phosphopeptide enrichment strategies like immobilized metal affinity chromatography (IMAC) or titanium dioxide (TiO_2_).^11,12^ One phosphoproteomic study of TCR signaling in mice using 20 million cells per replicate and IMAC phosphopeptide enrichment identified only 77 unique pTyr peptides,^13^ whereas other studies report 1-2% (about 250 to 700) of their total yield as unique pTyr peptides using IMAC or TiO_2_ with 30 million to 100 million cells per replicate.^11,14^ As demonstrated by Iwai et al., decreasing the number of cells and therefore the amount of protein input can severely limit pTyr quantitation depth in primary T cells from mice.

To increase the accuracy and precision of pTyr quantitation in cases of limited protein availability, both experimental and computational approaches are being developed. For example, recent improvements in pTyr enrichment reagents, namely the superbinder SH2 enrichment method,^15–20^ have improved quantitation depth of the pTyr proteome.^21–23^ Additionally, the use of isobaric labeling reagents like tandem mass tags (TMT) have allowed for accurate phosphopeptide quantitation in multiplexed samples with a higher probability of identifying unique peptides compared to label-free quantitation. ^24–29^ To improve the spectral quality and speed of acquiring TMT samples on Fourier transform-mass spectrometers (FT-MS), instrument settings like the phase-constrained spectrum deconvolution method (ΦSDM) are available. By applying the ΦSDM, FT spectra are deconvolved into frequency distributions, allowing for efficient extraction of the harmonic components of oscillating ions and ultimately achieving higher mass accuracy and resolution in shorter times.^30^

Recently, we combined the multiplexing capability of TMT, the selectivity of superbinder SH2, and a carrier channel using broad inhibition of tyrosine phosphatases to develop the broad-spectrum optimization of selective triggering (BOOST) method to increase pTyr quantitation depth in proteomics experiments.^21^ During the development of BOOST we used Jurkat T cells, a model system for studying TCR signaling. ^31^ However the BOOST method has not yet been demonstrated in the more biologically relevant primary T cells from mice. Here, we report the first pTyr proteomics study in primary T cells from mice using the BOOST method. By using predetermined protein input amounts, we show that BOOST increases the sequencing depth of low abundance samples, yielding more than 3,000 unique pTyr peptides and more than 2,000 unique pTyr sites in experimental samples. We also show that acquiring samples using the ΦSDM degrades quantitation in low-abundance samples. By using samples with contrived ratios, samples acquired with the ΦSDM disabled have higher replicate reproducibility, are more accurate, and are more precise than equivalent samples acquired with the ΦSDM enabled.

## Materials and Methods

### Stimulation of mice primary T cells

Splenic CD8^+^ T cells from B6 mice were isolated using the EasySep Mouse CD8^+^ T cell Isolation Kit (StemCell, cat. # 19853), primed with plate-bound 1 *µ*g/mL of *α*-CD3 (eBioscience, clone 145-2C11, cat. # 14-0031-86) and 0.5 *µ*g/mL *α*-CD28 (eBioscience, clone 37.51, cat. # 553294) for two days, then expanded using 50 U/mL of IL-2 for 4 days. Cells were rested in 1% BSA T cell media for 2 hours at 2 *×* 10^6^ cells/ml prior to stimulation. To initiate T cell stimulation, 25 *µ*g/mL of biotin-labeled *α*-CD3 antibody(eBioscience, clone 145-2C11, cat. # 14-0031-86) and 25 *µ*g/mL streptavidin were added to the cells resuspended at 1 *×* 10^8^ cells/ml for 5 minutes at 37°C. After 5 minutes of stimulation, cells were lysed with 2% (w/v) sodium dodecyl sulfate (SDS) in 100 mM Tris-HCl (pH 7.6). Pervanadate treatment for the carrier channel sample was performed by incubating cells with 500 *µ*M PV (prepared by mixing equal volume of 1 mM sodium orthovanadate and 1 mM hydrogen peroxide) for 20 minutes at 37°C.

### Sample processing

Lysates were processed using the QIAshredder Mini Spin Column by centrifugation at 20,000*×*g at 37 °C for 5 minutes. The concentration of protein in each sample was determined using the Pierce BCA Protein Assay (Thermo Fisher Scientific, 23225) before treatment with 100 mM dithiothreitol at room temperature for 30 minutes. Lysate was subsequently processed and digested using the filter-aided sample preparation (FASP) method^32^ as described previously. Digested peptides were collected and acidified by trifluoroacetic acid and desalted using Sep-Pak C18 Cartridge (Waters WAT020515) as described previously.^33^ Desalted peptides were labeled using a Tandem Mass Tag as described previously.^21^ TMT-labeled peptides were mixed and purified for phosphotyrosine peptides using the Src SH2 domain superbinder as described previously.^21^

### Liquid chromatography tandem mass spectrometry

Offline basic (pH 10) fractionation of peptides was performed on a 100 mm *×* 1.0 mm Acquity BEH C18 column (Waters) using an UltiMate 3000 UHPLC system (ThermoFisher Scientific) with a 40-minute gradient from 1% to 40% Buffer B_basic_ into 36 fractions. The 36 fractions were subsequently consolidated into 12 super-fractions (Buffer A_basic_ = 10 mM ammonium hydroxide in 99.5% (v/v) HPLC-grade water, 0.5% (v/v) HPLC-grade acetonitrile; Buffer B_basic_ = 10 mM ammonium hydroxide in 100% HPLC-grade acetonitrile), which were further separated on an in-line 150 mm *×* 75 *µ*m reversed phase analytical column packed in-house with XSelect CSH C18 2.5 *µ*m resin (Waters) using an UltiMate 3000 RSLCnano system (ThermoFisher Scientific) at a flow rate of 300 nL/min. A 65 minute gradient from 5% to 30% Buffer B_acidic_ followed by a 6-minute gradient 30% to 90% Buffer B_acidic_ (Buffer A_acidic_ = 0.1% (v/v) formic acid in 99.4% (v/v) HPLC-grade water, 0.5% (v/v) HPLC-grade acetonitrile; Buffer B_basic_ = 0.1% (v/v) formic acid in 99.9% (v/v) HPLC-grade acetonitrile) was used to separate peptides. Data were acquired in data-dependent acquisition (DDA) mode on a Orbitrap Eclipse Tribrid mass spectrometer (ThermoFisher Scientific) with a positive spray voltage of 2.25 kV, multinotch TMT-MS3 settings, ^34^ and a cycle time was set at 2.5 seconds. For MS1 level scans, precursor ions with charge states from 2 to 5 were acquired on the Orbitrap detector with a scan range of 400 − 1600 *m/z*, 120,000 resolution, maximum injection time of 50 ms, automatic gain control (AGC) target of 800,000, and a dynamic exclusion time of 15 seconds. An isolation window of 0.7 *m/z* was used to isolate MS1 precursor ions on the quadrupole for MS2 scans. The MS2 scans were acquired in centroid mode on the ion trap detector using a scan range of 400 − 1400 *m/z*. Precursor ions were fractured via higher-energy dissociation (HCD, 33% energy) activation with an AGC target of 5,000 and a maximum injection time of 75 ms. Using synchronous precursor selection (SPS), ^34^ 10 notches were further isolated on the quadrupole using an MS2 isolation window of 3 *m/z* for MS3 scans, which are acquired on the Orbitrap detector on a scan range of 100 − 500 *m/z* in a mass resolution of 50,000 via HCD activation (55% energy) with a AGC target of 250,000 and maximum injection time of 150 ms in centroid mode.

### Database search parameters and acceptance criteria for identifications

Raw files were processed using MaxQuant^35^ version 1.6.17.0 with the integrated peptide search engine Andromeda. ^36^ MS/MS spectra were searched against a mouse UniProt database (*Mus musculus*, last modified 12/01/2019) comprised of 55,412 forward protein sequences. The false discovery rate (FDR) for peptide spectrum matches was determined using a reverse decoy database approach and peptides with FDR*<* 0.01 were retained for further analysis. For database searching, the following parameters were set: carbamidomethylation (cysteine) was set as fixed modification; oxidation (methionine) was set as a variable modification; acetylation (protein N-termini) was set as a variable modification; phosphorylation (serine, threonine, tyrosine) was set as a variable modification. The specificity of Trypsin was used with up to two missed cleavages. The peptide tolerance for the main search was set to 5 ppm, whereas the FTMS tolerance was 20 ppm and ITMS MS/MS tolerance was 0.5 Da. For MS3 quantitation, the reporter ion mass tolerance was set to 3 mDa using the isotopic correction factors provided by the manufacturer (Lot UK291564, Lot UH283151). The search parameter file (mqpar.xml) and all MaxQuant output files are provided in Supporting Folder 1.

### Data analysis & code availability

All analysis and data visualization were preformed on Ubuntu 20.04 LTS in the Windows Subsystem for Linux version 2 using Python 3.8.10 with the packages “Matplotlib” (Version 3.3.2), “SciPy” (Version 1.7.3), “pandas” (Version 1.2.3), “NumPy” (Version 1.19.2), “Biopython” (version 1.78), and “matplotlib-venn” (version 0.11.6) and is available in Supporting Folder 1. Analysis of unique PSMs was performed using the MaxQuant output file “evidence.txt” (Supporting Folder 2). Unique PSMs were defined by a non-redundant amino acid sequence (including posttranslational modifications), the charge state of the peptide, and the least number of missing values across all TMT channels. In the cases where redundancy was still present, we kept the peptide with the highest median reporter intensity. For assigning flanking sequences to each peptide and generating Supporting Figure 1, the MaxQuant output file “Phospho (STY)Sites.txt” (Supplementary Folder 2) was used. For determining previously annotated pTyr sites we used the PhosphoSitePlus^®^ (www.phosphosite.org)^37^ posttranslational modification database file “Phosphorylation_site_dataset.txt” (Supporting Folder 1). Before analysis, all peptides at a 1% FDR flagged as “potential contaminants” or “reverse hits” were removed, and reporter ions from the PV-treated sample (TMT126) and the Blank channel (TMT127N) were excluded from further analysis unless otherwise stated. Statistical significance between the mean corrected reporter intensities for 1.0 mg, 0.3 mg, and 0.1 mg protein input samples was determined using unpaired Student’s T-tests to calculate *p*-values before correcting for multiple hypotheses (generating *q*-values) using the method of Benjamini & Hochberg. ^38^ For all comparisons, statistical significance was only attained for peptides where reporter intensity values were present for all three replicates. In line with the previous literature, we did not impute or interpolate missing values at any point during data analysis.^21,22^ To evaluate replicate reproducibility, we performed least squares linear regression^39^ on pairwise comparisons between replicates for each protein input amount in a given experiment, removing peptides for which one or both replicates contained missing values. For all volcano plots, we plotted − log_10_(*q*-value) as a function of log_10_(Ratio of Mean pTyr Intensities) for comparisons between 1.0 mg and 0.1 mg, 1.0 mg and 0.3 mg, and 0.3 mg and 0.1 mg of protein input for each TMT mix. For cases where volcano plots were constructed for each portion of a Venn diagram, separate volcano plots for each overlapping portion were constructed using the reporter intensity data from each experiment. For BOOST Factor plots, only pTyr peptides with at least one reporter ion value were used, and we used the following equation to calculate BOOST Factor for each pTyr peptide:

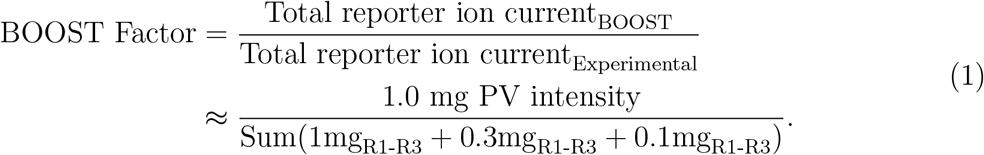

The results of all analysis are provided in Supporting Tables 1-6, including references to the identification number(s) for each peptide in the original MaxQuant output file(s). All Python scripts are also available on [GitHub](https://github.com/kendeegz/2022_chua_callahan_BOOST_code).

## Results

### Experimental design & rationale

In this study, we sought to determine the performance of the recently developed BOOST method for pTyr proteomics in primary T cells from mice. In short, BOOST is a method used to increase the precursor ion triggering and fragmentation of pTyr-containing peptides using a pervanadate (PV) treated sample as a carrier channel in multiplexed TMT experiments, thus increasing quantitation depth of low-abundance posttranslational modifications.^21^ Our design is similar to that of our previous BOOST studies, ^21,22^ using one 1.0 mg PV treated sample (or 1.0 mg protein control) and predetermined protein sample amounts (1.0 mg, 0.3 mg, and 0.1 mg) from 5-minute CD3-stimulated primary mouse T cells to define the accuracy and precision of the BOOST method in primary T cells (Figure 1A,B). The protein amounts used in our experiments range from 1.0 mg to 0.1 mg, which is about 5- to 10-fold less at a maximum and 50- to 100-fold less at a minimum than what is typically used for in pTyr proteomics studies.^7,9–11,14,23^ In PV BOOST experiments, there is potential for reporter ion leakage between nearby channels, which can negatively impact quantitation and increase the potential for false positive discoveries. ^40^ In our previous study we found evidence of reporter ion leakage from channel 126 (+PV) to 127C, however we found no evidence of leakage from 126 to 127N.^22^ Therefore, we included a “Blank” channel (127C) to catch potential reporter ion leakage from the 1.0 mg PV-treated sample (126; Figure 1B). To enrich for pTyr-containing peptides, we used the superbinder SH2 method ^19–21,23^ prior to acquisition and analysis by LC-MS and MaxQuant, respectively. To understand how the ΦSDM affects pTyr quantitation in BOOST experiments, our BOOST and control TMT mixes were acquired with (+ΦSDM) and without the ΦSDM on an Orbitrap Eclipse Tribrid mass spectrometer (Figure 1A). From all experiments (BOOST and control, with and without the ΦSDM), the majority of identified phosphorylation sites were localized to tyrosines (70%) with 94.1% of pTyr sites being assigned with probability *>* 0.75 (Supporting Figure 1).

**Figure 1:**
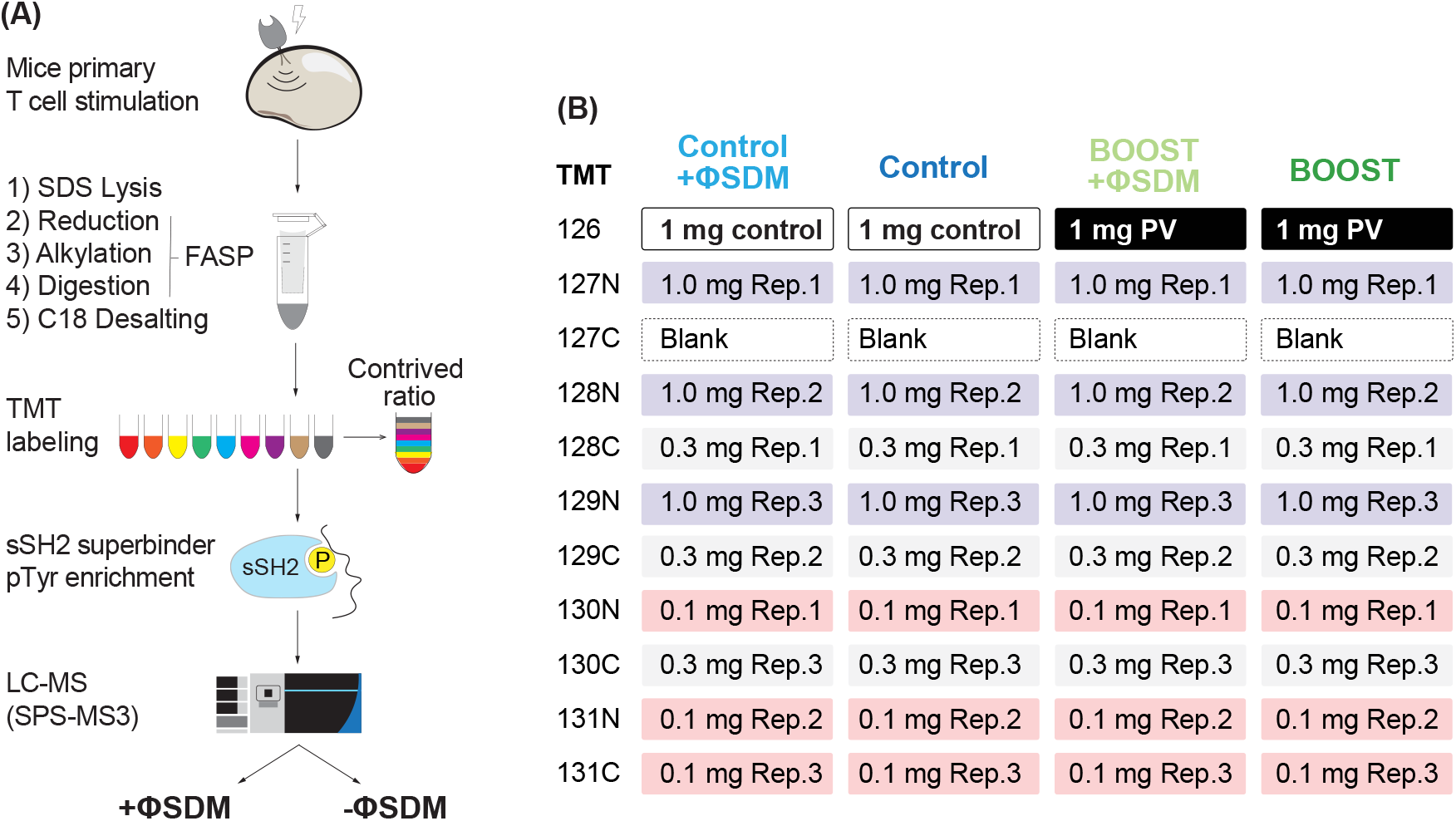
Schematic of (A) experimental workflow and (B) TMT mix and channel design.

### Disabling the ΦSDM and BOOST increases pTyr quantitation depth in BOOST experiments with minimal ratio compression or reporter ion interference

To our surprise, disabling the ΦSDM increased the number of pTyr peptide PSMs with quantifiable reporters in both the BOOST and control TMT mixes. In the 1.0 mg PV-treated channels, we observed 5,741 unique pTyr peptide PSMs with the ΦSDM disabled and only 4,839 with the ΦSDM enabled. On average, 1.0 mg protein input samples using BOOST yielded 2,425 quantifiable pTyr peptide PSMs with the ΦSDM disabled compared to 1,066 when the ΦSDM is enabled, a 2.3 fold increase. We observed improvement when disabling the ΦSDM in both 0.3 and 0.1 mg protein input samples in BOOST, with an average of 1,019 and 369 quantifiable pTyr peptides compared to 640 and 142 with the ΦSDM enabled, respectively (Figure 2C,D). The increased quantitation depth also came with fewer missing values. The average percentage of missing data for 1.0 mg, 0.3 mg, and 0.1 mg samples using BOOST dropped from 78.4, 87.0, and 97.1 to 59.0, 82.8, and 93.8 when the ΦSDM was disabled, respectively (Supporting Figure 2C,D). While the control samples did benefit from disabling the ΦSDM, the magnitude of improvement was smaller (1.3-fold for 1.0 mg, 1.3-fold for 0.3 mg, and 1.7-fold for 0.1 mg; Figure 2A,B) and the percentage of missing values between replicates were similar (Supporting Figure 2A,B). Using the BOOST method without the ΦSDM led to a 7.6-fold, 3.6-fold, and 3.1-fold increase in the number of unique pTyr peptide PSMs quantified in the 1.0 mg, 0.3 mg, and 0.1 mg samples, respectively, when compared to the control samples acquired without the ΦSDM. These data suggest that the ΦSDM negatively effects pTyr quantitation in both control and BOOST experiments, and that using the BOOST method substantially increases pTyr quantitation depth.

**Figure 2:**
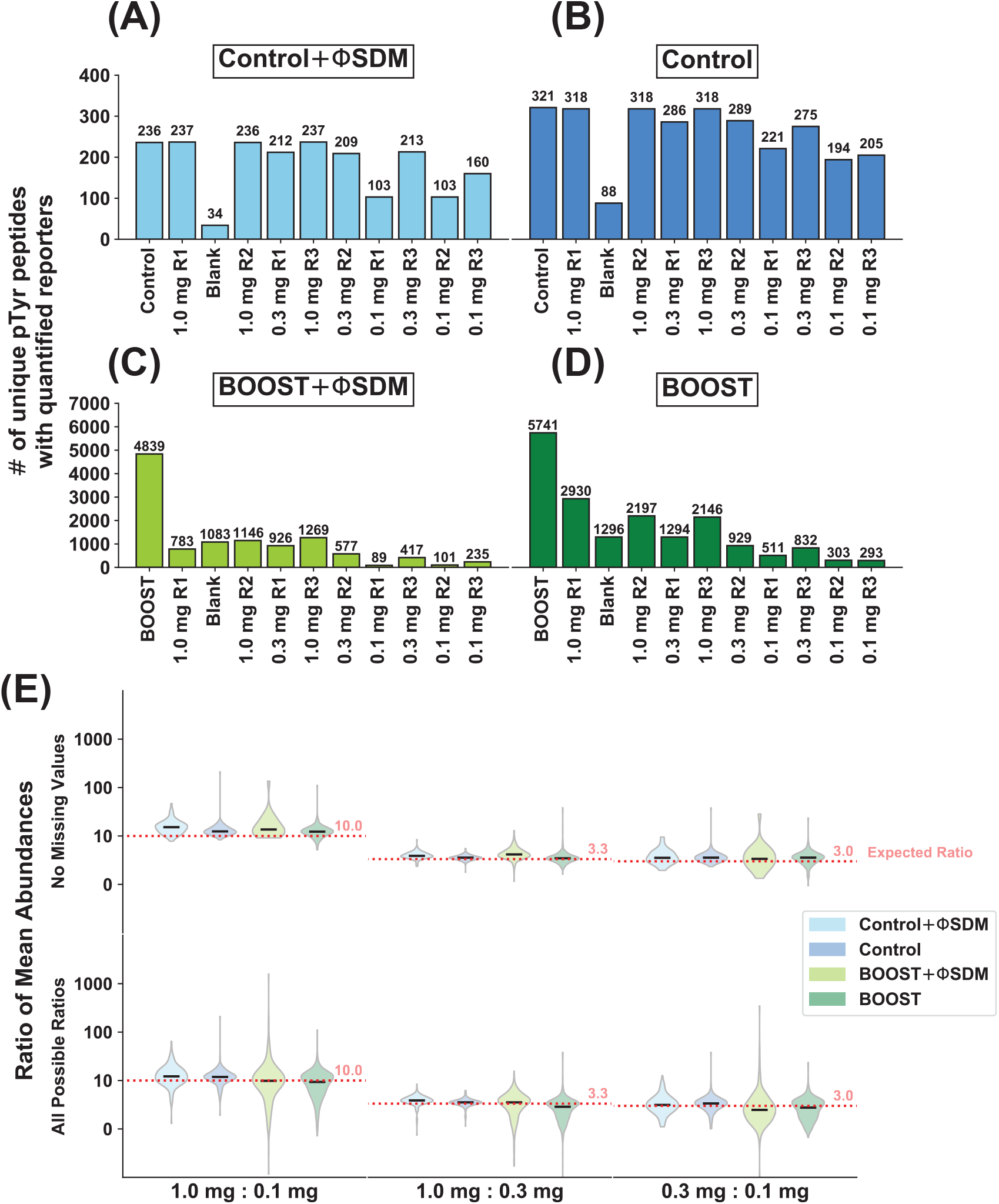
Quantitation depth is improved in BOOST and 1.0 mg Control experiments when the ΦSDM is disabled without detectable ratio suppression. The number of unique pTyr peptides identified in each TMT channel for the (A) 1.0 mg Control experiment with the ΦSDM enabled, (B) 1.0 mg Control experiment with the ΦSDM disabled, (C) BOOST experiment with ΦSDM enabled, (D) BOOST experiment with the ΦSDM disabled. The exact number of unique pTyr peptides is indicated above each bar. (E) Ratio of mean intensities for each unique pTyr peptide observed. Top row depicts ratios with no missing values (*n* = 3 intensiy values in each group). Bottom row depicts all possible ratios (*n ≥* 1 intensity values in each group).

Previous literature evaluating the BOOST method suggested that a PV BOOST channel can promote ratio compression and increase reporter ion interference.^40^ Therefore, we evaluated whether the PV BOOST channel influenced quantitation in neighboring channels in our experiments. In both control and BOOST samples with or without the ΦSDM enabled, we quantified peptides in the blank channels (Figure 2A-D) and the distribution of reporter ion intensities in the blank channels were markedly lower than all other channels (Supporting Figure 3) as observed previously.^22^ The distribution of intensity values in experimental channels were consistent for each replicate in each condition (Supporting Figure 3). To determine whether ratio compression was present, we plotted the ratio of all pTyr peptide PSMs with no missing values (Figure 2E, upper row) or with at least one value (Figure 2E, lower row) in each group. For pTyr peptide PSMs with no missing values and all possible ratios, the median ratios between all protein input amounts were similar to the controls with or without the ΦSDM enabled and aligned well with the expected ratios (Figure 2E, upper row).

Next, we evaluated whether reporter ion leakage was prevalent in any individual experimental channel by evaluating deviation of individual replicates from the mean intensity value for each pTyr peptide PSM present in all replicates in each condition (Supporting Figures 4-6). The mean intensity values for 1.0 mg replicates predicted each individual replicate well in all experiments with *r*^2^ values greater than 0.94 in all comparisons, while the counts of pTyr peptide PSMs were greatly elevated when comparing BOOST to control. We also observed high *r*^2^ values (*>* 0.9) in the 0.3 mg and 0.1 mg replicates in all experiments where the ΦSDM was disabled, however experiments with ΦSDM enabled had noticably lower *r*^2^ values and fewer repeating pTyr peptide PSMs for low abundance samples (Supporting Figures 5,6). Together, these data support the conclusion that ratio compression and reporter ion leakage were minimal while yield of pTyr peptide PSMs were greatly elevated with BOOST.

### Disabling the ΦSDM increases precision while BOOST increases the yield of of pTyr quantitation in low abundance samples

Interestingly, we observed a substantial increase in replicate reproducibility after disabling the ΦSDM in both BOOST and control experiments, especially in low abundance samples. We assessed replicate reproducibility by performing simple least squares regression in a pairwise manner on replicates for 1.0 mg, 0.3 mg, and 0.1 mg protein inputs for BOOST and control experiments acquired with and without the ΦSDM (Figure 3A, Supporting Figures 7-10). When the ΦSDM was disabled, we observed higher average values for the coefficient of determination (*r*^2^), a measure of the linear relationship between data, in all conditions. This effect was clearest in the low abundance samples, where the average *r*^2^ for BOOST experiments with 0.1 mg of protein increased from 0.527 to 0.775 by disabling the ΦSDM (Figure 3A, Supporting Figures 7, 8). We observed similar results in the control samples, where disabling the ΦSDM increased the *r*^2^ from 0.566 to 0.863 (Figure 3A, Supporting Figures 9, 10). When comparing the BOOST and control samples, we observed a modest decrease in replicate reproducibility when using BOOST for all protein input amounts (Figure 3A). However, the average number of peptides with quantifiable reporter ions increased 2 to 7-fold using BOOST (Supporting Figures 7, 9), leading to an average 1.6-fold increase in the number of significantly changing peptides (Figure 3B). Our data reveal that the ΦSDM generally reduces replicate reproducibility and that using the BOOST method increases the number of significantly changing pTyr peptide PSMs observed.

**Figure 3:**
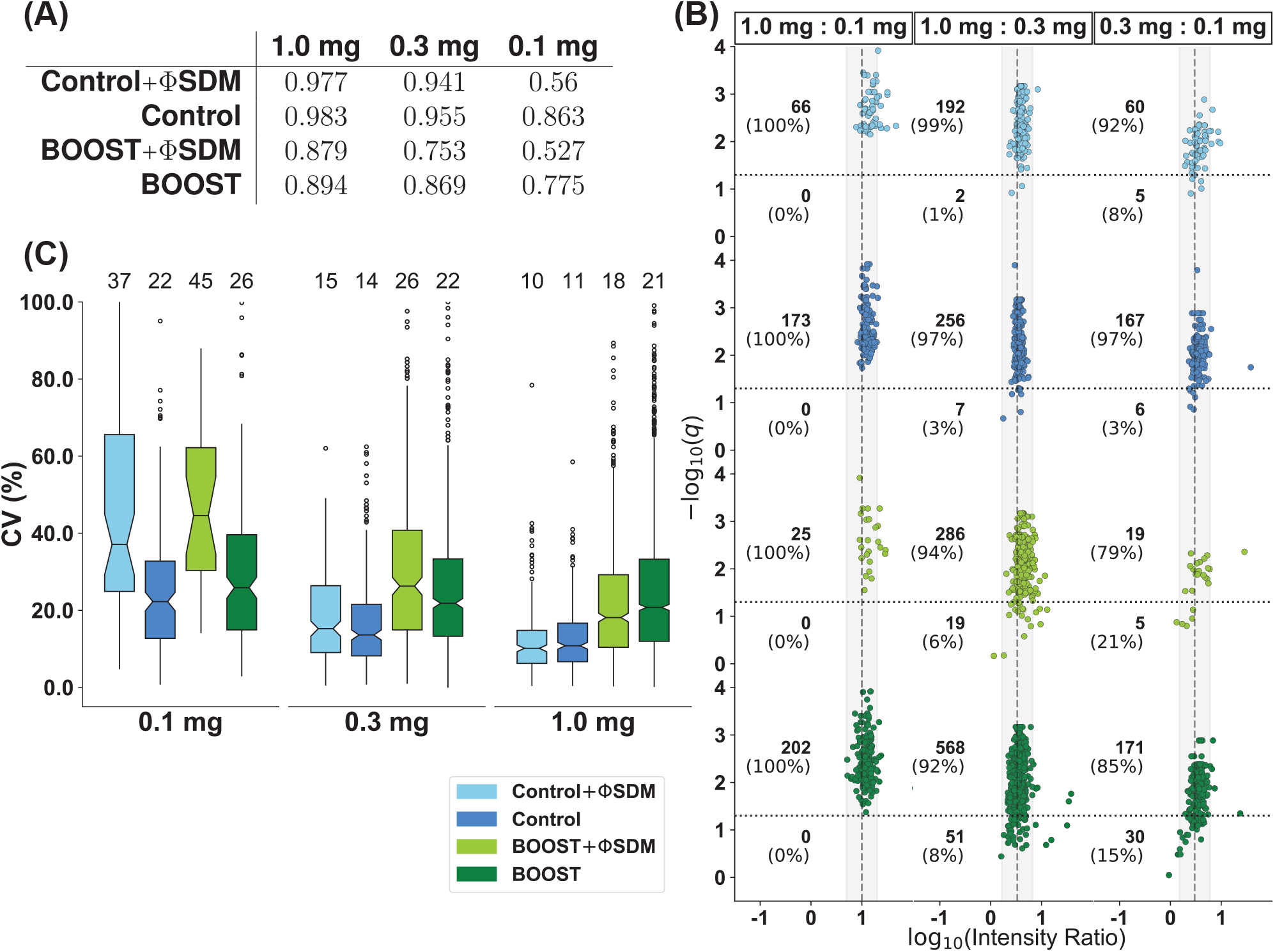
ΦSDM decreases both replicate reproducibility and detection of significant changes with or without BOOST, while BOOST permits quantification of more significantly changing PSMs at the cost of elevated replicate variance. (A) Table showing the average coefficient of determination (*r*^2^) from least squares linear regression preformed on replicates. (B) Volcano plot showing contrived ratios for each TMT mix as indicated. The numbers and proportions [in percentages] of ratios above and below a *q*-value of 0.05 [horizontal black, dotted lines] are indicated. The grey, dashed line indicates the theoretically expected ratio, and the grey shaded area represents 2-fold above and below theoretically expected ratios. (C) Box-and-whisker plots showing the percentage coefficient of variation of the triplicate intensities for each protein input as indicated. The median coefficient of variation percentages are shown above each boxplot. Color labels apply to (B) and (C).

In addition to increasing reproducibility, disabling the ΦSDM led to a modest incresase the accuracy of pTyr quantitaiton. We assessed accuracy by observing clustering around the theoretically expected peptide intensity ratios in violin plots (Figure 2E) and in volcano plots (Figure 3B), which also show the number of significantly changing pTyr peptide PSMs in each comparison. In both the control and BOOST experiments with the ΦSDM disabled, we observed tight clustering of values around theoretical truth, especially in the 1.0 mg to 0.1 mg comparison (Figure 2E, upper row). In contrast, enabling the ΦSDM decreased both clustering around the theoretical truth (Figure 2E) and the number of peptides with a statistically significant difference in mean reporter intensity (Figure 3B). Disabling the ΦSDM lead to a 2.8-fold increase in statistically significant ratios between the 0.3 mg and 0.1 mg protein input conditions for control experiments, and a 9.0-fold increase for BOOST experiments (Figure 3B). The increased number of statistically significant ratios with the ΦSDM disabled was coupled with an increase in quantitative precision in low abundance samples. For 0.1 mg protein input samples, disabling the ΦSDM decreased the median coefficient of variation (CV) from 37% to 22% in control experiments and 45% to 26% in BOOST experiments, while the CV% for 0.3 mg and 1.0 mg samples remained similar in the presence or absence of the ΦSDM for control and BOOST experiments (Figure 3C). Although we observed an increase in CV% for 0.3 mg and 1.0 mg when comparing the control and BOOST experiments without the ΦSDM, the increase in CV% did not impact the number of statistically significant pTyr peptide PSMs identified (Figure 3B,C). Together, our data show that disabling the ΦSDM for multiplexed TMT experiments with low protein input substantially increases the quality of pTyr quantitation, and the BOOST method improves detection of significantly changed pTyr peptide PSMs.

### The magnitude of pTyr quantitation in BOOST experiments is improved when ΦSDM is disabled

Because the goal of BOOST is to improve quantitation of low abundance peptides, we chose to examine the magnitude of improvement with the ΦSDM disabled. We first determined the populations of pTyr peptide PSMs unique to BOOST (“BOOST-Gained”), unique to the control (“Control-Only”), or found in both experiments (“Overlap”) when the ΦSDM was disabled or enabled (Figure 4A,B, Supporting Tables 1-6). We identified 1.94-fold more BOOST-Gained pTyr peptide PSMs with the ΦSDM disabled than with the ΦSDM enabled (Figure 4A,B). The accuracy of reporter intensity measurements was almost identical in overlapping peptides identified in the BOOST and control experiments with the ΦSDM disabled, with a large increase in the number of significant BOOST-Gained peptides in all contrived ratios (Supporting Figure 11). In contrast, the precision and yield of significant overlapping peptides were severely lowered in the BOOST experiment compared to the control with the ΦSDM enabled (Supporting Figure 12). Disabling the ΦSDM led to a large increase in BOOST-Gained pTyr peptide PSMs, whereas the gain in control pTyr peptide PSMs was minimal (Supporting Figure 12). Our data suggest that the ΦSDM degrades the precision of measurements in control-overlapping pTyr peptide PSMs and limits the potential to identify unique pTyr peptides PSMs.

**Figure 4:**
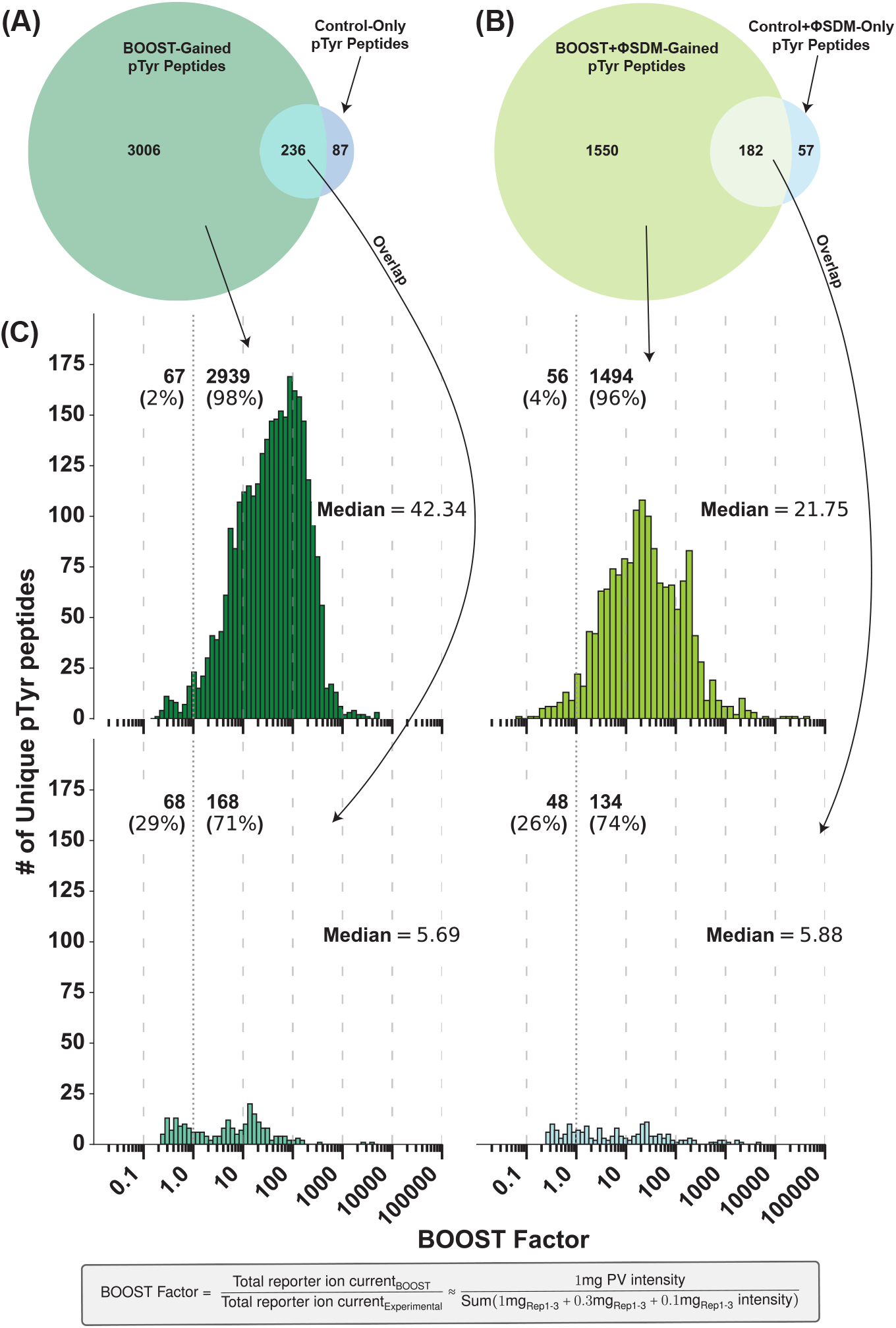
The number of peptides and the magnitude of BOOST factor are increased when the ΦSDM is disabled. (A) Venn diagram showing the number of unique pTyr peptides quantified in the PV carrier channel and observed in at least one experimental channel when the ΦSDM is disabled in the BOOST, the 1.0 mg Control, and in both the BOOST and 1.0 mg Control TMT mixes. (B) Venn diagram showing the number of unique pTyr peptides observed when the ΦSDM is enabled in the BOOST, the 1.0 mg Control, and in both the BOOST and 1.0 mg Control TMT mixes. (C) Histogram showing the distribution of BOOST factor values with or without the ΦSDM. Top row show BOOST-gained pTyr peptide PSMs. Bottom row show pTyr peptide PSMs observed in each BOOST experiment and the corresponding control experiment.

In our paper describing the BOOST method, we determined the magnitude of quantitative improvement in BOOST experiments using “BOOST factors”, defined as the ratio of the reporter intensity from the PV-treated sample to the sum of reporter intensities from experimental channels for a specific pTyr peptide PSM (Figure 4C, bottom). ^21^ A pTyr peptide PSM with a BOOST factor exceeding 1 occurs when the summed reporter ion current of the PV-treated sample is greater than the reporter ion current of the experimental channels, indicating the peptide is generally scarce in the experimental samples. ^21^ Overall, the majority of BOOST-Gained peptide PSMs had BOOST factors greater than 1 regardless of whether the ΦSDM was enabled or disabled. BOOST-Gained pTyr peptide PSMs had a median BOOST factor of 7.4-fold higher than Control-overlapping pTyr peptide PSMs, while this effect was muted when the ΦSDM was enabled. To account for BOOST-Gained pTyr peptide PSMs with missing values, we filtered the data to contain pTyr peptide PSMs where intensity ratios and statistical significance could be attained and plotted their cumulative distributions (Supporting Figure 14). For ratios including the low abundance 0.1 mg samples (0.3 mg to 0.1 mg, *n* = 92 and 1.0 mg to 0.1 mg, *n* = 115) acquired without the ΦSDM, 90% of the significantly changing pTyr peptide PSMs had a BOOST factor less than 5, which shifted to about 18 in the higher abundance ratio (1.0 mg to 0.3 mg). The BOOST factor distribution for the 1.0 mg to 0.3 mg ratio with the ΦSDM enabled was similar to BOOST without the ΦSDM, however too few significantly changing ratios were detected for this analysis to be useful for ratios including the low abundance 0.1 mg sample (Supporting Figure 14). When we included ratios where at least one replicate value was identified for each protein input sample, the distribution of BOOST factors were almost identical for ratios including the 0.1 mg samples, with 90% of pTyr peptide PSMs having BOOST factors less than 20 (Supporting Figure 14). A BOOST factor of 15− to 20− fold is in agreement with the current recommendations for carrier channel use in single cell proteomics, where a carrier channel boost of *∼* 20 is optimal.^41^ These data suggest that disabling the ΦSDM increases the quantity and quality of significantly changing pTyr peptide PSMs in low abundance 0.1 mg samples observed using the BOOST method in primary T cells from mice.

### BOOST reveals pTyr sites critical for T cell receptor signaling in primary T cells

To show the efficacy of BOOST pTyr proteomics in primary T cells we used *α*-CD3 antibodies to stimulate TCR signaling (Figure 1). Therefore, we expected to observe many unique pTyr sites on TCR signaling proteins. In accordance with our expectations, we observed a total of 113 unique pTyr sites on proteins in the Kyoto Encyclopedia of Genes and Genomes (KEGG) T cell receptor signaling pathway (Figure 5, Supporting Figure 15). To determine the profiling range of unique pTyr site sequencing in primary T cells using BOOST, we compared our BOOST experiment with the ΦSDM disabled to our previously published BOOST experiment in Jurkat T cells which used an equivalent amount of cellular protein and set of contrived ratios (1.0:0.3:0.1 mg total protein).^21^ Our Jurkat BOOST experiment was acquired using an Orbitrap Fusion Lumos Tribrid mass spectrometer without the ΦSDM and had a similar BOOST-Gained yield to the present primary T cell BOOST data, ^21,42^ and is therefore suitable for comparison. Within the KEGG TCR annotated proteins, we observed 60 pTyr sites in both mouse and Jurkat BOOST experiments (Figure 5 inset, Supporting Figure 15), with the majority of these sites on ITAMs (TCR*ζ*, CD3*δ*/*ϵ*/*γ*; 12), the tyrosine kinase Zap70 (10), and the canonical activation sites on mitogen activated protein kinases (Erk1/2, p38*α*/*β*, Jnk1*/*2*/*3; 7). We found 53 unique pTyr sites exclusive to BOOST in primary T cells from mice, compared to 32 unique pTyr sites exclusive to BOOST in Jurkat T cells (Figure 5 inset, Supporting Figure 15). Notably, using the BOOST method in primary T cells from mice uncovered 8 pTyr sites on CD45, a phosphatase critical for the activation of Lck,^43^ and 4 pTyr sites on Tec, a non receptor tyrosine kinase with overlapping function with Itk in TCR signaling. ^44,45^ Of the 4 unique pTyr sites observed on Tec, the activation site (Y519) was quantified in all three replicates in the 1.0, 0.3, and 0.1 mg samples of the BOOST experiment, whereas it was quantified in some but not all replicates of all conditions of the 1.0 mg Control experiment. Similar results were found for SHP-1^Y536^, Itk^Y512^, Zap70^Y290^, and Zap70^Y492^ from our BOOST experiment in primary T cells from mice (Supporting Table 2). These results suggest that using BOOST in primary T cells from mice increases pTyr profiling depth similarly to what we observed previously using Jurkat T cells. ^21,22^

**Figure 5:**
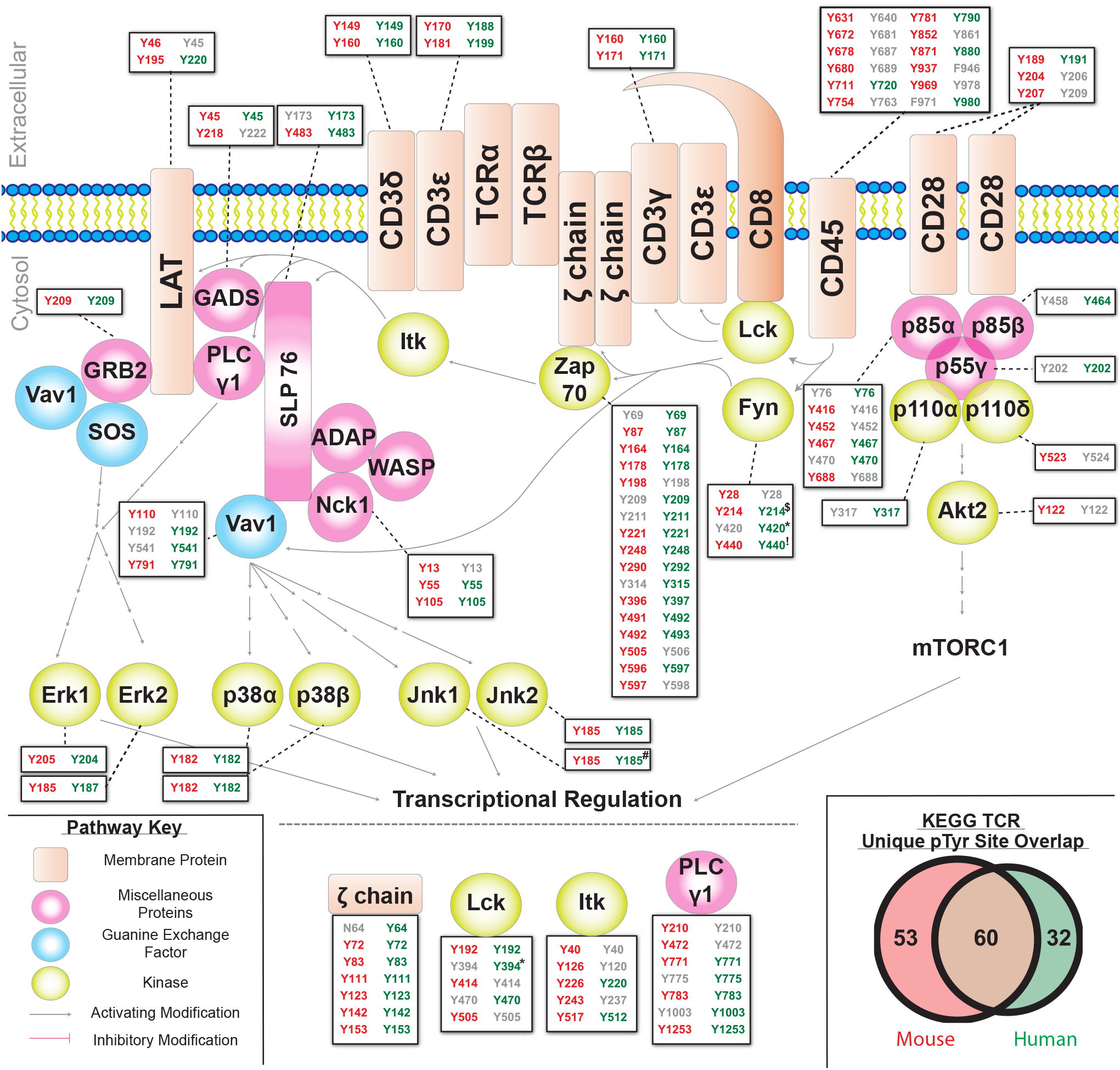
BOOST pTyr proteomics reveals a comparable number of unique TCR-related pTyr sites in primary T cells from mice and Jurkat T cells. Unique pTyr sites identified in our BOOST experiment with the ΦSDM disabled are colored in red. Unique pTyr sites identified by Chua et al. 2020 in the Jurkat T cell model system for studying TCR signaling are colored in green. Inset shows the overlap between unique pTyr sites in the Kyoto Encyclopedia of Genes and Genomes T cell receptor signaling pathway between our BOOST experiment and the BOOST experiment performed by Chua et al. 2020. Special characters next to site numbers indicate the presence of PSMs that arise from multiple proteins: ^$^Fyn/Yes1 | **\*** Fyn/Yes1/Src/Lck | ^**!**^Fyn/Yes1/Src | ^**#**^Jnk1/Jnk3.

## Discussion

To improve our understanding of the critical tyrosine phosphorylation events involved in TCR signaling and other cellular processes, accurate methods to perform deep profiling of the pTyr proteome are required. Although the accuracy of LC-MS/MS techniques are desirable for pTyr proteomics, the low abundance of tyrosine phosphorylation events in the proteome hinder the frequently used global phosphoenrichment methods like TiO_2_ or IMAC.^11,12,46–48^ Although recent developments in pTyr-specific enrichment techniques like p-Tyr-1000 and the superbinder SH2 method have improved pTyr proteomics,^15–21^ the issue of low pTyr abundance is apparent when using samples that are difficult or expensive to collect, such as primary cells from humans or mice. ^7,9,10^ Increasing quantitative yield in low abundance samples has been achieved in multiplexed TMT experiments using a carrier proteome sample,^40,41,49^ and we developed the BOOST method using a PV treated sample to increase quantitative yield of pTyr peptides.^21,22^

Here, we demonstrate the first use of the BOOST method in primary T cells from mice, defining the accuracy, precision, and profiling depth of the mouse T cell pTyr proteome in low abundance samples. Our multiplexed TMT experiments were designed to minimize reporter ion interference from the PV channel by including a “Blank” (127C) channel where maximal reporter ion interference has been observed previously (Supporting Figures 3-6, Figure 2). ^22,40^ In addition, our maximal protein amount of 1.0 mg is 5- to 10-fold less and our minimal protein amount is 50- to 100-fold less than what most pTyr proteomics studies typically use. Using BOOST, we were able to quantify more than 2,000, 900, and 300 unqiue pTyr peptide PSMs in 1.0 mg, 0.3 mg, and 0.1 mg protein samples, respectively (Figure 2D), while maintaining accuracy and precision (Figure 3, Supporting Figure 7). Using BOOST allowed for 3,006 BOOST-gained pTyr peptide PSMs to be quantified with 2,939 pTyr peptides that were low abundance in the samples (Figure 4C). This included 113 unique pTyr sites on proteins involved in the T cell receptor signaling pathway, with 53 of these sites being uniquely identified in mice when compared to our the pTyr sites we originally identified in Jurkat T cells (Figure 5, Supporting Figure 15). ^21^ Of the TCR signaling proteins identified using BOOST, many of the unique pTyr sites were found in all replicates of the 0.3 and/or 0.1 mg samples in the BOOST experiment and not in the 1.0 mg Control experiment. Together, our data suggest that including a PV BOOST channel increases quantitative depth of low abundance peptides in higher abundance samples and overall quantitation in low abundance samples without large compromises to accuracy or precision while increasing the number of significant changes observed in the tyrosine phosphoproteome.

We also examined the influence of the acquisition parameter ΦSDM, a computational method that increases acquisition rate of FT-MS by efficient and noise tolerant deconvolution of FT spectra,^30^ on our BOOST and 1.0 mg Control samples. Although previous research has shown that using the ΦSDM on long gradients or scarce samples may reduce the efficiency of the algorithm due to low ion currents, ^30,42,50^ the influence of the ΦSDM on TMT mixes with carrier proteome channels has yet to be evaluated. Our data are in agreement with previous literature suggesting that enabling the ΦSDM degrades low abundance samples. We observed a decrease of about 75 to 100 unique pTyr peptides across our 1.0 mg Control samples with the ΦSDM enabled, with the largest loss in the 0.1 mg R1 sample (221 to 104 unique pTyr peptides; Figure 2A,B). Surprisingly, enabling the ΦSDM degraded the quality of data from BOOST experiments. We observed a large reduction of unique pTyr peptide yield in experimental channels, with the largest difference being 1.0 mg R1 dropping from 2,931 unique pTyr peptides to 784 unique pTyr peptides with the ΦSDM enabled (Figure 2C,D). We also observed a reduction in accuracy (Figure 3B), precision (Figure 3C), and replicate reproducibility (Supporting Figures 4, 5) with the ΦSDM enabled. Our study indicates that disabling the ΦSDM, or “Turbo-TMT” on the method editor on Orbitrap instruments, substantially improves the quantitation depth of low abundance posttranslational modification samples, especially when a BOOST channel is present.

With increased interest in using proteomics to study rare or specific posttranslational modifications^20,51,52^ and the proteomes of single-cells,^41,49,53^ reliable methods to increase multiplexing capabilities, ^54^ posttranslational modification selection, ^55^ and quantitation^21,22,56^ are desired. These experimental techniques will come with a wave of computational methods to further improve quantitation,^30,57,58^ which will require stringent testing for both experimental-computational method compatibility and to understand the range of biological processes that these methods can work with. In this study, we displayed both of these features for the BOOST method by showing that BOOST and the ΦSDM were incompatible and that BOOST can increase the yield of pTyr peptides in primary T cells from mice, which are notoriously refractory for wide scale analysis.^7^ By using this study as a benchmark for the BOOST method in primary T cells from mice, future research into the pTyr proteome of primary T cells from mice is possible using far less sample than is conventionally used.

## Conclusion

Our study defines the accuracy, precision, and profiling depth of multiplexed TMT experiments using a pervandate BOOST channel to increase detection of significantly changed pTyr peptides in stimulated primary T cells from mice. We found that including the BOOST channel increases the quantitative yield of unique pTyr peptide PSMs without jeopardizing the accuracy in low abundance samples. Importantly, we found no evidence of reporter ion leakage between the PV BOOST channel and the experimental channels or ratio compression when using BOOST and SPS-MS3 quantitation. While we did observe an increase in the median coefficient of variation when using BOOST, detection of significant changes was greatly increased. For example, important pTyr sites on the tyrosine kinases Tec, Itk, and Zap70, as well as pTyr sites on the phosphatase SHP-1 would not have been detected without the use of BOOST. The majority of the BOOST-Gained pTyr peptide PSMs were scarce, revealing that BOOST increases identification of rare pTyr peptide PSMs. Interestingly, we found that enabling the ΦSDM degrades the quality of data in BOOST experiments, almost halving the unique pTyr peptide PSM yield and reducing accuracy and precision in low abundance samples. Together, our study shows that multiplexed TMT experiments using a PV BOOST channel increase quantitative yield of meaningful unique pTyr peptide PSMs in primary T cells from mice and that the ΦSDM should be used cautiously for experiment employing a carrier channel.

## Supporting information

Supporting Folder 2 Data Analysis

Supporting Folder 1 MaxQuant Results

## Supporting Information

The Supporting Information is available free of charge at https://pubs.acs.org/. Supporting Information includes:

- All tables generated after MaxQuant analysis of .raw files (“summary.txt”, “evidence.txt”, “peptides.txt”, “modificationSpecificPeptides.txt”, “Oxidation (M)Sites.txt”, “Phospho (STY)Sites.txt”, “proteinGroups.txt”, “allPeptides.txt”, “msScans.txt”, “mzRange.txt”, “msmsScans.txt”, and “msms.txt”) (.ZIP)
- All Python3 code used to perform data analysis and representation, including statistical analyses, replicate reproducibility assessments, BOOST factor analysis, and comparisons between TMT experiments (.ZIP)
- All output from statistical analyses performed and selected MaxQuant output required for statistical analysis (.XLSX)

## Acknowledgements

The authors wish to thank Dr. Ricky Edmondson and Dr. Samuel G. Mackintosh from University of Arkansas for Medical Sciences (UAMS) for collecting the proteomic data. We acknowledge financial support from NIH grants R01AI083636, P20GM121293, and R24GM137786. Kenneth P. Callahan was supported by T32GM136566 and the Sidney E. Frank Fellowship.

## Data Availability

The mass spectrometry proteomic data have been deposited to the ProteomeXchange Consortium (http://proteomecentral.proteomexchange.org) via the PRIDE partner repository^59^ with the dataset identifier PXD025853 (Username: reviewer _pxd025853ebi.ac.uk, Password: RDtiS7iG).

