## Supporting Folder 2 Data Analysis for "Mouse primary T cell phosphotyrosine proteomics enabled by BOOST": data_analysis.py

```
"""
============================================================================================================================================================
Mouse primary T cell phosphotyrosine proteomics enabled by BOOST
Xien Yu Chua(1), Kenneth P. Callahan(2), Alijah A. Griffith(2), Tobias Hildebrandt(1), Guoping Fu(3), Mengzhou Hu(1), Renren Wen(3), Arthur R. Salomon(2,*)
(1)Department of Molecular Pharmocology, Physiology & Biotechnology, Brown University, Providence, RI, 02912
(2)Department of Molecular Biology, Cell Biology & Biochemistry, Brown University, Providence, RI, 02912
(3)Blood Research Institute, Blood Center of Wisconsin, Milwaukee, WI, 53226
* Corresponding Author,
============================================================================================================================================================
Kenneth P. Callahan
19 February 2022
Salomon Laboratory
Post-MaxQuant Data Analysis

Python 3.8.10

Dependencies:

helpers: A folder of common functions created by Kenneth P. Callahan for data analysis
         Put this folder in the same directory as this file and include '__init__.py' in this
         this directory as well

NumPy Version 1.19.2
SciPy Version 1.7.2
Pandas Version 1.2.3
matplotlib Version 3.3.2
BioPython Version 1.78
matplotlib-venn Version 0.11.6

============================================================================================================================================================
This script was used to process the MaxQuant output files 'evidence.txt' and 'Phospho (STY)Sites.txt'
and produce the data-based figures in the manuscript. These include:

- Coefficient of Variation box-and-whisker plots (Figure 3C)
- Correlation plots and simple least squares regression fits (Supporting Figures 3-6)
- Contrived ratio Volcano Plots (Figure 3B, Supporting Figures 7 & 8)
- Unique pTyr Peptide Counts & Missing Values (Figure 2)
- Reporter Intensity Boxplots (Supporting Figure 2)
- pTyr Peptide Venn Diagrams (Figure 4A & B, Supporting Figures 7-9)
- BOOST Factor Histograms (Figure 4C)
- BOOST Factor Empirical Distributions (Supporting Figure 10)
- Localisation Probability Histogram (Supporting Figure 1)

The data are available in Supporting Tables 1-6.
============================================================================================================================================================
"""

###################################################################################################
#
#  Importables

# Check for the PSP Dataset
import os
# For managing system-level processes
import sys
#
if not os.path.exists("database/phosphositeplus_ptm_data.txt") and not os.path.exists("database/Phosphorylation_site_dataset"):
    print("The PhosphoSitePlus PTM database was not found.")
    print("Please visit phosphosite.org and download the 'Phosphorylation_site_dataset.gz' file,")
    print("unzip the file, and place it in the database folder.")
    sys.exit(1)
elif os.path.exists("database/phosphositeplus_ptm_data.txt"):
    psp_rename = True
else:
    psp_rename = False

# Homemade helper modules for statistics, plotting, and genral data manipulation
from helpers import stats_helpers as sh
from helpers import mpl_plotting_helpers as mph
from helpers import general_helpers as gh


# For avoiding manipulation of original data structures
import copy
# For plotting
import matplotlib.pyplot as plt
# For log histogram plotting
import numpy as np

# For grabbing gene names from Entrez IDs
from Bio import Entrez
# *Always* tell NCBI who you are
Entrez.email = ""
# For transforming data
from math import log2, log10
# For colouring the pairwise correlation plots
from scipy.stats import gaussian_kde
# For easy Venn Diagram plotting
from matplotlib_venn import venn2
# For creating legend labels
from matplotlib.patches import Patch
# For manipulating plot sizes
from matplotlib.gridspec import GridSpec

#
#
###################################################################################################
#
#  Functions: Data Management

def keep_first(a_matrix, id_col):
    keep = {}
    for item in a_matrix:
        if item[id_col] not in list(keep.keys()):
            keep[item[id_col]] = item
    return sorted([value for key, value in keep.items()], key = lambda x: x[id_col])

def ratio(m,n):
    if m != m or n != n:
        return float("nan")
    else:
        return m/n

def index_ps(pvals):
    indices = [[i,pvals[i]] for i in range(len(pvals))]
    ps = [p for p in indices if p[1] == p[1]]
    return list(zip(*ps))

def fill_nans(indices, qvals, total_len):
    returner = []
    for i in range(total_len):
        if i in indices:
            returner.append(float(qvals.pop(0)))
        else:
            returner.append(float("nan"))
    return returner

def remove_nanpairs(d1,d2):
    pairs = list(zip(d1,d2))
    pairs = [list(pair) for pair in pairs if pair[0] == pair[0] and pair[1] == pair[1]]
    return list(zip(*pairs))
    
def filter_nanpairs(list_1, list_2):
    """
    redundant to remove_nanpairs, except it returns both lists separately rather
    than zipped
    """
    newlist = []
    for item in zip(list_1, list_2):
        if item[0] == item[0] and item[1] == item[1]:
            newlist.append(item)
    return ([item[0] for item in newlist], [item[1] for item in newlist])

def boost_factor(experiment_matrix,
                 boost_ind, exp_inds,
                 head_ind = 0):
    """
    """
    factors = []
    heads = experiment_matrix[head_ind]
    for row in experiment_matrix:
        if row == heads:
            continue
        else:
            newrow = [row[boost_ind]] + [row[i] for i in exp_inds if row[i] == row[i]]
            if len(newrow) == 1:
                factors.append(float("nan"))
            else:
                factors.append(newrow[0] / sum(newrow[1:]))
    return factors

#
#
###################################################################################################
#
#  Functions: Entrez Annotation Management

def retrieve_annotation(id_list, verbose = True):

    """Annotates Entrez Gene IDs using Bio.Entrez, in particular epost (to
    submit the data to NCBI) and esummary to retrieve the information.
    Returns a list of dictionaries with the annotations."""

    request = Entrez.epost("gene", id=",".join(id_list))
    try:
        result = Entrez.read(request)
    except RuntimeError as e:
        # FIXME: How generate NAs instead of causing an error with invalid IDs?
        print("An error occurred while retrieving the annotations.")
        print(f"The error returned was {e}")
        sys.exit(-1)

    webEnv = result["WebEnv"]
    queryKey = result["QueryKey"]
    data = Entrez.esummary(db="gene", webenv=webEnv, query_key=queryKey)
    annotations = Entrez.read(data)["DocumentSummarySet"]["DocumentSummary"]
    if verbose:
        print(f"Retrieved {len(annotations)} annotations for {len(id_list)} genes\n")
    return annotations

def make_id_name_dict(annotations):
    return {gene_data.attributes["uid"] : gene_data["Name"] for gene_data in annotations}

def get_gene_names(id_list, annotations):
    id_name_dict = make_id_name_dict(annotations)
    return [id_name_dict[gene_id] for gene_id in id_list]

# Functions to handle reading the GMT databases from WikiPathway
def get_database(filename, delim = "\t"):
    return gh.read_file(filename, delim = "\t")

def update_entrez_ids(database, verbose = True):
    with_genes = []
    organism = False
    for row in database:
        if not organism:
            organism = row[0].split("%")[-1]
            if verbose:
                print(f"{organism:^30}\n")
        newrow = [row[0].split("%")[0], row[1]]
        if verbose:
            print(newrow[0])
        annotations = retrieve_annotation(row[2:], verbose = verbose)
        gene_names = get_gene_names(row[2:], annotations)
        newrow += gene_names
        with_genes.append(newrow)
    return organism, with_genes

def write_database(database,
                   newfilename,
                   delim = "\t"):
    newdb = [gh.list_to_str(row, delimiter = "\t",
                            newline = True) for row in database]
    gh.write_outfile(newdb, newfilename, writestyle = "w")
    print(f"{newfilename} written\n")
    
# Functions to flip the database
def get_all_genes(database):
    return list(set(gh.unpack_list([row[2:] for row in database])))

def make_genedb(database):
    genes = get_all_genes(database)
    genedict = {gene.upper() : [] for gene in genes}
    for row in database:
        for gene in row[2:]:
            genedict[gene.upper()].append(row[0])
    genedict = {gene : gh.list_to_str(annots,
                                      delimiter = ";",
                                      newline = False) for gene, annots in genedict.items()}
    return genedict

# Function to add database annotation to current matrix
def annotate_matrix(a_matrix, genedb,
                    newhead = "WikiPathway Annotation",
                    gene_col = 0):
    heads = a_matrix[0] + [newhead]
    newmatrix = [heads]
    for row in a_matrix[1:]:
        try:
            genedb[row[gene_col].upper()]
        except:
            newmatrix.append(row + ["None"])
        else:
            newmatrix.append(row + [genedb[row[gene_col].upper()]])
    return newmatrix

def wikipath_annotation(database_filename, a_matrix,
                        database_delim = "\t", 
                        write_genedb = False,
                        gene_col = 0,
                        verbose = True):
    database = get_database(database_filename)
    org,database = update_entrez_ids(database, verbose = verbose)
    if write_genedb:
        newfilename = database_filename.split(".")[0]+"_genes.gmt"
        write_databse(database, newfilename)
    genedb = make_genedb(database)
    return annotate_matrix(a_matrix, genedb,
                           gene_col = gene_col,
                           newhead = f"WikiPathway Annotation ({org})")

#
#
###################################################################################################
#
#  Functions: Plotting-related

def scatter_gausskde(d1,d2, ax, scatterargs = {"s" : 20,
                                               "alpha" : 0.5,
                                               "marker" : "s"},
                    llsr = True,
                    update_ticks = True,
                    xlim = [0,25],
                    ylim = [0,25],
                    colour = None):
    if llsr and xlim != None and ylim != None:
        data1, data2 = remove_nanpairs(d1,d2)
        a, b, r2= sh.least_squares_fit(data1, data2)
        xs = [xlim[0], ylim[1]]
        # Finish by getting y values and plotting a line
        ys = [a+b*xs[0], a+b*xs[1]]
        ax.plot(xs, ys, color = "black", linestyle = ":",
                label = f"$y={a:.2f}+{b:.2f}x$\n$r^2 = {r2:.3f}$")
    else:
        data1, data2 = remove_nanpairs(d1,d2)
        a, b, r2= sh.least_squares_fit(data1, data2)
        xs = [min(data1), max(data1)]
        # Finish by getting y values and plotting a line
        ys = [a+b*xs[0], a+b*xs[1]]
        ax.plot(xs, ys, color = "black", linestyle = ":",
                label = f"$y={a:.2f}+{b:.2f}x$\n$r^2 = {r2:.3f}$")
    if colour == None:
        xy=np.vstack(remove_nanpairs(d1,d2))
        z = gaussian_kde(xy)(xy)
        ax.scatter(*remove_nanpairs(d1,d2), c=z, label = f"$n={len(data1)}$",
                **scatterargs)
    else:
        ax.scatter(*remove_nanpairs(d1,d2), c=colour, label = f"$n={len(data1)}$",
                **scatterargs)
    if xlim != None:
        ax.set_xlim(*xlim)
    if ylim != None:
        ax.set_ylim(*ylim)
    if update_ticks:
        ax.set_xticks(ax.get_xticks())
        xlabs = []
        for tick in ax.get_xticks():
            if int(tick) == float(tick):
                xlabs.append(str(int(tick)))
            else:
                xlabs.append(str(tick))
        ax.set_xticklabels(xlabs,
                           fontfamily ="sans-serif", font = "Arial",
                           fontweight = "bold")
        ax.set_yticks(ax.get_yticks())
        ylabs = []
        for tick in ax.get_yticks():
            if int(tick) == float(tick):
                ylabs.append(str(int(tick)))
            else:
                ylabs.append(str(tick))
        ax.set_yticklabels(ylabs,
                           fontfamily ="sans-serif", font = "Arial",
                           fontweight = "bold")
    return ax

def get_above_below(a_list, value = 1):
    total = len(a_list)
    counts = {"below" : 0,
              "above" : 0}
    for item in a_list:
        if item <= value:
            counts["below"] += 1
        else:
            counts["above"] += 1
    counts["below"] = f'{counts["below"]}\n(${(counts["below"]/total)*100:.0f}\%$)'
    counts["above"] = f'{counts["above"]}\n(${(counts["above"]/total)*100:.0f}\%$)'
    return [counts["below"], counts["above"]]

def cdf(data, bins, ax, colour, label = "",
        plot_90 = False):
    values, base = np.histogram(data, bins = bins)
    logbase = np.logspace(log10(base[0]),log10(base[-1]),len(base))
    cumulative = np.cumsum(values)
    counts, bins, bars = ax.hist(data, cumulative = True,bins = logbase[:-1],
                                color = "white", histtype = "barstacked", density = True,
                                 alpha = 0)
    ax.scatter(bins[:-1], counts, color = colour, s = 10)
    ax.plot(bins[:-1], counts, color = colour,
            label = label)
    ax.set_xscale("log")
    return ax

#
#
###################################################################################################
#
#  Functions: Flanking Sequences & PSP Database Comparisons

def _make_flankhelper(loc_prob_seq,
                          highest_locprob):
    if float(highest_locprob) == int(float(highest_locprob)):
        seqs = [gh.remove_chars(string) for string in loc_prob_seq.split(f"Y({int(float(highest_locprob))})")]
    else:
        seqs = [gh.remove_chars(string) for string in loc_prob_seq.split(f"Y({highest_locprob})")]
    if len(seqs[0]) <= 2 and len(seqs[1]) <= 2:
        return seqs[0], seqs[1]
    elif len(seqs[0]) == 1:
        return seqs[0], seqs[1][:4]
    elif len(seqs[0]) == 2:
        return seqs[0], seqs[1][:3]
    elif len(seqs[1]) == 1:
        return seqs[0][-4:], seqs[1]
    elif len(seqs[1]) == 2:
        return seqs[0][-3:], seqs[1]
    else:
        return seqs[0][-3:], seqs[1][:2]
    
    
def grab_loc_prob(loc_prob_seq, amino_acid = "Y"):
    """
    """
    # Split the string on Ys that have probabilities
    split = loc_prob_seq.split("Y(")
    probs = []
    for seq in split:
        if ")" in seq:
            # The probability is everything up to the next parenthesee
            # So add it to the probs
            try:
                probs.append(float(seq.split(")")[0]))
            except:
                pass
        # Otherwsie just pass
        else:
            pass
    # Return the maximum localization probability
    # and all of them as a string, in order
    return max(probs), gh.list_to_str(probs, delimiter = ";", newline = False)

def make_loc_probs(a_matrix, 
                   loc_prob_col, 
                   head_col = 0,
                   newcol_names = ["Highest (Y) Localisation Probability",
                                   "All (Y) Localisation Probabilities"]):
    locs = gh.grab_col(a_matrix, loc_prob_col)[1:]
    loc_tups = [grab_loc_prob(seq) for seq in locs]
    loc_probs = list(zip(*loc_tups))
    loc_probs= [list(row) for row in loc_tups]
    new_matrix = [a_matrix[head_col] + newcol_names] + [a_matrix[i+1] + loc_probs[i] for i in range(len(loc_probs))]
    return new_matrix

def add_sequence_window(evidence_matrix,
                        sty_matrix,
                        loc_prob = "Highest (Y) Localisation Probability",
                        loc_prob_seq = "Localisation Probability (Sequence)",
                        name = "Phospho (STY) site IDs",
                        aa = "Y"):
    """
    """
    seq_wind = gh.grab_col(sty_matrix, "Sequence window")
    ids = gh.grab_col(sty_matrix, "id")
    prob_col = evidence_matrix[0].index(name)
    newrows = [evidence_matrix[0] + ["Sequence Window", "Flanking Sequence"]]
    loc_prob_id = evidence_matrix[0].index(loc_prob)
    loc_prob_seq_id = evidence_matrix[0].index(loc_prob_seq)
    # Loop over the rows
    for row in evidence_matrix[1:]:
        # Get the potential IDs for Phospho (STY)Sites
        if ";" in str(row[prob_col]):
            prob_ids = [int(item) for item in row[prob_col].split(";")]
        elif row[prob_col] == "":
            prob_ids = float("nan")
        else:
            prob_ids = [int(row[prob_col])]
        # Check the IDs
        if prob_ids != prob_ids:
            newrows.append(row + ["", ""])
            continue
        else:
            found = 0
            for p_id in prob_ids:
                if found == 1:
                    continue
                else:
                    index = ids.index(p_id)
                    windows = seq_wind[index].split(";")
                    for wind in windows:
                    # If the center of the sequence is a tyrosine
                        if wind[len(wind)//2] == aa:
                            flank1,flank2 = _make_flankhelper(row[loc_prob_seq_id],
                                                             row[loc_prob_id])
                            break_seq = wind[:len(wind)//2], wind[len(wind)//2+1:] 
                            if break_seq[0][-len(flank1):] == flank1 and break_seq[1][:len(flank2)] == flank2:
                                newrows.append(row + [wind, wind[8:23]])
                                found = 1
                            elif flank1 == "" and break_seq[1][:len(flank2)] == flank2:
                                newrows.append(row + [wind, wind[8:23]])
                                found = 1
                            elif break_seq[0][-len(flank1):] == flank1 and flank2 == "":
                                newrows.append(row + [wind, wind[8:23]])
                                found = 1
                            else:
                                continue
                        else:
                            continue
            if found == 0:
                newrows.append(row + ["", ""])
    return newrows

def _merge_database_seqs(matrix, matrix_gene_col,
                         matrix_flank_col, psp_genes,
                         psp_flanks, psp_sites,
                         psp_groupids, psp_orgs):
    # Get all of the indices for these columns
    m_gene_ind = matrix[0].index(matrix_gene_col)
    m_flank_ind = matrix[0].index(matrix_flank_col)
    #
    new_matrix = [matrix[0] + [f"Modified Site",
                               "PSP Site Group ID",
                               "Organism (Origin)"]]
    #
    i=0
    for row in matrix[1:]:
        i += 1
        if row[m_flank_ind] in psp_flanks:
            occurs = [i for i in range(len(psp_flanks)) if psp_flanks[i] == row[m_flank_ind]]
            found = 0
            for o in occurs:
                if psp_genes[o] in row[m_gene_ind]:
                    new_matrix.append(row + [psp_sites[o],
                                             psp_groupids[o],
                                             psp_orgs[o]])
                    found = 1
                    break
                else:
                    continue
            if found == 0:
                new_matrix.append(row + ["", "", ""])
        else:
            new_matrix.append(row + ["", "", ""])
    if i == 77:
        return None
    return new_matrix

def compare_seqs_to_database(psp_database, matrices,
                             matrix_gene_col = "Gene names",
                             matrix_flank_col = "Flanking Sequence",
                             psp_flank_col = "SITE_+/-7_AA",
                             psp_gene_col = "GENE",
                             psp_site_col = "MOD_RSD", 
                             psp_groupid_col = "SITE_GRP_ID",
                             psp_org_col = "ORGANISM"):
    """
    """
    
    #
    psp_genes = gh.grab_col(psp_database, psp_gene_col)
    psp_flanks = gh.grab_col(psp_database, psp_flank_col)
    psp_flanks = [flank.upper() for flank in psp_flanks]
    psp_sites = gh.grab_col(psp_database, psp_site_col)
    psp_groupids = gh.grab_col(psp_database, psp_groupid_col)
    psp_orgs = gh.grab_col(psp_database, psp_org_col)
    #
    return [_merge_database_seqs(matrix, matrix_gene_col,
                         matrix_flank_col, psp_genes,
                         psp_flanks, psp_sites,
                         psp_groupids, psp_orgs) for matrix in matrices]
    

def _merge_other_species_sites(matrix, matrix_groupid_col,
                               psp_groupids, psp_sites,
                               psp_flanks, organism = "human"):
    m_groupid_ind = matrix[0].index(matrix_groupid_col)
    new_matrix = [matrix[0] + [f"PSP Modified Site (Equivalent in {organism.upper()})",
                               f"PSP Flanking Sequence (Equivalent in {organism.upper()})"]]
    for row in matrix[1:]:
        if row[m_groupid_ind] in psp_groupids:
            ind = psp_groupids.index(row[m_groupid_ind])
            new_matrix.append(row + [psp_sites[ind],
                                      psp_flanks[ind]])
        else:
            new_matrix.append(row + ["", ""])
    return new_matrix

def get_other_species_sites(psp_database, matrices,
                            organism = "human",
                            matrix_groupid_col = "PSP Site Group ID",
                            psp_groupid_col = "SITE_GRP_ID",
                            psp_site_col = "MOD_RSD",
                            psp_flank_col = "SITE_+/-7_AA"):
    """
    """
    #m_groupid_ind = matrices[0][0].index(matrix_groupid_col)
    psp_flanks = gh.grab_col(psp_database, psp_flank_col)
    psp_flanks = [flank.upper() for flank in psp_flanks]
    psp_sites = gh.grab_col(psp_database, psp_site_col)
    psp_groupids = gh.grab_col(psp_database, psp_groupid_col)
    return [_merge_other_species_sites(matrix, matrix_groupid_col,
                                       psp_groupids, psp_sites,
                                       psp_flanks,
                                       organism = organism) for matrix in matrices]

#
#
###################################################################################################
#
#  Colours, labels, headers to rename, etc.

# These are the colours for the plots, in the following order:
    # [0] Control$+\Phi$SDM
    # [1] Control
    # [2] BOOST$+\Phi$SDM
    # [3] BOOST
colours = ["skyblue", "dodgerblue", "yellowgreen", "green"]
labels = [r"Control$+\Phi$SDM", "Control", "BOOST$+\Phi$SDM", "BOOST"]

ten_x = [1,3,5]
three_x = [4,6,8]
one_x = [7,9,10]

# Dictionary with new names for each column of evidence.txt that
# we care about
rename_cols = {"Reporter intensity corrected 1" : "PV Boost",
               "Reporter intensity corrected 2" : "1.0 mg R1",
               "Reporter intensity corrected 3" : "Blank",
               "Reporter intensity corrected 4" : "1.0 mg R2",
               "Reporter intensity corrected 5" : "0.3 mg R1",
               "Reporter intensity corrected 6" : "1.0 mg R3",
               "Reporter intensity corrected 7" : "0.3 mg R2",
               "Reporter intensity corrected 8" : "0.1 mg R1",
               "Reporter intensity corrected 9" : "0.3 mg R3",
               "Reporter intensity corrected 10" : "0.1 mg R2",
               "Reporter intensity corrected 11" : "0.1 mg R3",
               "Phospho (STY) Probabilities" : "Localisation Probability (Sequence)",
               "Gene names" : "Gene names",
               "Protein names" : "Protein names",
               "WikiPathway Annotation (Homo sapiens)" : "WikiPathway Annotation (Homo sapiens)",
               "WikiPathway Annotation (Mus musculus)" : "WikiPathway Annotation (Mus musculus)",
               "id" : "id",
               "Phospho (STY) site IDs" : "Phospho (STY) site IDs",
               "Modified sequence" : "Modified sequence",
               "Charge" : "Charge",
               "Experiment" : "Experiment"}

# General characteristics of fonts in plots
textdict = {"fontfamily" : "sans-serif",
            "font" : "Arial",
            "fontsize" : "16"}
            
ylabels = [r"$\log_2$(R2)", r"$\log_2$(R3)", r"$\log_2$(R3)"]
xlabels = [r"$\log_2$(R1)", r"$\log_2$(R1)", r"$\log_2$(R2)"]
rows = ["1.0 mg", "0.3 mg", "0.1 mg"]

control_labs = ["Control", "1.0 mg R1", "Blank", "1.0 mg R2", "0.3 mg R1", 
               "1.0 mg R3", "0.3 mg R2", "0.1 mg R1", "0.3 mg R3",
                "0.1 mg R2", "0.1 mg R3"]

boost_labs = ["PV BOOST", "1.0 mg R1", "Blank", "1.0 mg R2", "0.3 mg R1", 
               "1.0 mg R3", "0.3 mg R2", "0.1 mg R1", "0.3 mg R3",
                "0.1 mg R2", "0.1 mg R3"]

plot_heads = [[labels[0], labels[1]],
              [labels[2], labels[3]]]

cdf_colours = ["brown",  "grey","hotpink"]
cdf_labs = [f"1.0:0.1 mg\n",
            f"1.0:0.3 mg\n",
            f"0.3:0.1 mg\n"]

overlap_bc_colours = [colours[3], "mediumaquamarine", "mediumaquamarine", colours[1]]
overlap_bscs_colours = [colours[2], "paleturquoise", "paleturquoise", colours[0]]

keep_boost_cols = {"Gene names" : "Gene names",
               "Protein names" : "Protein names",
                "Modified sequence" : "Modified sequence",
               "Localisation Probability (Sequence)" : "Localisation Probability (Sequence)",
               "Modified Site" : "Modified Site",
               "PSP Modified Site (Equivalent in HUMAN)" : "PSP Modified Site (Equivalent in HUMAN)",
               "Highest (Y) Localisation Probability" : "Highest (Y) Localisation Probability",
               "All (Y) Localisation Probabilities":"All (Y) Localisation Probabilities",
               "Sequence Window" : "Sequence Window",
               "Flanking Sequence" : "Flanking Sequence",
               "PSP Flanking Sequence (Equivalent in HUMAN)" : "PSP Flanking Sequence (Equivalent in HUMAN)",
               "Charge" : "Charge",
               "Experiment" : "Experiment",
               "PV Boost" : "PV Boost or 1.0 mg Control",
               "1.0 mg R1" : "1.0 mg R1",
               "Blank"     : "Blank",
               "1.0 mg R2" : "1.0 mg R2",
               "0.3 mg R1" : "0.3 mg R1",
               "1.0 mg R3" : "1.0 mg R3",
               "0.3 mg R2" : "0.3 mg R2",
               "0.1 mg R1" : "0.1 mg R1",
               "0.3 mg R3" : "0.3 mg R3",
               "0.1 mg R2" : "0.1 mg R2",
               "0.1 mg R3" : "0.1 mg R3",
               "Median"   : "Row Median",
               "Missing values"  : "Missing Values",
               "10X mean" : "1.0 mg Mean Intensity", 
               "3X mean" : "0.3 mg Mean Intensity", 
               "1X mean" : "0.1 mg Mean Intensity",
               "10X/3X" : "1.0 mg Mean / 0.3 mg Mean", 
               "10X/1X" : "1.0 mg Mean / 0.1 mg Mean", 
               "3X/1X"  : "0.3 mg Mean / 0.1 mg Mean",  
               "10X SD" : "1.0 mg Standard Deviation",
               "3X SD"  : "0.3 mg Standard Deviation",
               "1X SD"  : "0.1 mg Standard Deviation",  
               "10X CV" : "1.0 mg Coefficient of Variation",
               "3X CV"  : "0.3 mg Coefficient of Variation",
               "1X CV"  : "0.1 mg Coefficient of Variation",  
               "10X/3X P" : "1.0 mg - 0.3 mg p value",
               "10X/1X P" : "1.0 mg - 0.1 mg p value",
               "3X/1X P"  : "0.3 mg - 0.1 mg p value",
               "10X/3X Q" : "1.0 mg - 0.3 mg q value",
               "10X/1X Q" : "1.0 mg - 0.1 mg q value",
               "3X/1X Q"  : "0.3 mg - 0.1 mg q value",
               "Boost Factor" : "BOOST Factor",
               "Unique ID" : "Unique ID",
               "WikiPathway Annotation (Homo sapiens)" : "WikiPathway Annotation (Homo sapiens)",
               "WikiPathway Annotation (Mus musculus)" : "WikiPathway Annotation (Mus musculus)",
               "PSP Site Group ID" : "PSP Site Group ID",
               "id" : "evidence.txt ID",
               "Phospho (STY) site IDs" : "Phospho (STY)Sites.txt ID"}

keep_cont_cols = {"Gene names" : "Gene names",
               "Protein names" : "Protein names",
                  "Modified sequence" : "Modified sequence",
               "Localisation Probability (Sequence)" : "Localisation Probability (Sequence)",
               "Modified Site" : "Modified Site",
               "PSP Modified Site (Equivalent in HUMAN)" : "PSP Modified Site (Equivalent in HUMAN)",
               "Highest (Y) Localisation Probability" : "Highest (Y) Localisation Probability",
               "All (Y) Localisation Probabilities":"All (Y) Localisation Probabilities",
               "Sequence Window" : "Sequence Window",
               "Flanking Sequence" : "Flanking Sequence",
               "PSP Flanking Sequence (Equivalent in HUMAN)" : "PSP Flanking Sequence (Equivalent in HUMAN)",
               "Charge" : "Charge",
               "Experiment" : "Experiment",
               "PV Boost" : "PV Boost or 1.0 mg Control",
               "1.0 mg R1" : "1.0 mg R1",
               "Blank"     : "Blank",
               "1.0 mg R2" : "1.0 mg R2",
               "0.3 mg R1" : "0.3 mg R1",
               "1.0 mg R3" : "1.0 mg R3",
               "0.3 mg R2" : "0.3 mg R2",
               "0.1 mg R1" : "0.1 mg R1",
               "0.3 mg R3" : "0.3 mg R3",
               "0.1 mg R2" : "0.1 mg R2",
               "0.1 mg R3" : "0.1 mg R3",
               "Median"   : "Row Median",
               "Missing values"  : "Missing Values",
               "10X mean" : "1.0 mg Mean Intensity", 
               "3X mean" : "0.3 mg Mean Intensity", 
               "1X mean" : "0.1 mg Mean Intensity",
               "10X/3X" : "1.0 mg Mean / 0.3 mg Mean", 
               "10X/1X" : "1.0 mg Mean / 0.1 mg Mean", 
               "3X/1X"  : "0.3 mg Mean / 0.1 mg Mean",  
               "10X SD" : "1.0 mg Standard Deviation",
               "3X SD"  : "0.3 mg Standard Deviation",
               "1X SD"  : "0.1 mg Standard Deviation",  
               "10X CV" : "1.0 mg Coefficient of Variation",
               "3X CV"  : "0.3 mg Coefficient of Variation",
               "1X CV"  : "0.1 mg Coefficient of Variation",  
               "10X/3X P" : "1.0 mg - 0.3 mg p value",
               "10X/1X P" : "1.0 mg - 0.1 mg p value",
               "3X/1X P"  : "0.3 mg - 0.1 mg p value",
               "10X/3X Q" : "1.0 mg - 0.3 mg q value",
               "10X/1X Q" : "1.0 mg - 0.1 mg q value",
               "3X/1X Q"  : "0.3 mg - 0.1 mg q value",
               "Unique ID" : "Unique ID",
               "WikiPathway Annotation (Homo sapiens)" : "WikiPathway Annotation (Homo sapiens)",
               "WikiPathway Annotation (Mus musculus)" : "WikiPathway Annotation (Mus musculus)",
               "PSP Site Group ID" : "PSP Site Group ID",
               "id" : "evidence.txt ID",
               "Phospho (STY) site IDs" : "Phospho (STY)Sites.txt ID"}

#
#
###################################################################################################
#
#  main() function

def main():
    
    print("STAGE 1: Loading Data and Performing Calculations ")
    
    print("\tReading the file 'maxquant_results/evidence.txt'")
    evidence = gh.read_file("maxquant_results/evidence.txt", delim = "\t")
    print(f"\tTotal rows in evidence: {len(evidence)-1}\n")

    print(f"\tFiltering reverse and potential contaminants")
    ev_clean = gh.filter_matrix(gh.filter_matrix(evidence, "Reverse", "+", False),
                                "Potential contaminant", "+", False)
    print(f"\t{len(ev_clean)-1} PSMs remaining\n")

    print(f"\tAnnotating the rows using WikiPathways Human Database\n")
    ev_clean = wikipath_annotation("database/wikipathways-20220410-gmt-Homo_sapiens.gmt",
                                   ev_clean, gene_col = ev_clean[0].index("Gene names"))
    print("\tDone\n")
    print(f"\tAnnotating the rows using WikiPathways Mouse Database")
    ev_clean = wikipath_annotation("database/wikipathways-20220410-gmt-Mus_musculus.gmt",
                                   ev_clean, gene_col = ev_clean[0].index("Gene names"))
    print("\tDone\n")
    print("\tGrabbing the columns relevant for data analysis\n")
    print("\tand renaming them:")
    for key, value in rename_cols.items():
        print(f"\t\t{key:^40} -> {value:^40}")
    ev_clean = gh.select_cols(ev_clean, rename_cols)
    print("\tDone\n")

    # Save the current index of the modified sequence column
    mod_seq_col = ev_clean[0].index("Modified sequence")

    print(f"\tTransforming all number strings to floats")
    ev_clean = [gh.transform_values(item) for item in ev_clean]
    print("\tDone\n")

    print(f"\tReplacing all 0 values with nan")
    ev_clean = [gh.replace_value(item, 0, float("nan")) for item in ev_clean]
    print("\tDone\n")

    print(f"\tPopulating 'Unique ID' column: '<Modified sequence><Charge><Experiment>'")
    ev_clean = [item + [gh.list_to_str(item[-3:], delimiter = "", newline = False)] for item in ev_clean]
    ev_clean[0][-1] = "Unique ID"
    print("\tDone\n")

    print(f"\tCalculating missing values in experimental channels")
    ev_clean = [item + [sum([1 for _ in [item[1]]+item[3:11] if _ != _])] for item in ev_clean]
    ev_clean[0][-1] = "Missing values"
    print("\tDone\n")

    print(f"\tCalculating median intensity for each row")
    ev_clean = [item + [sh.median([r for r in [item[1]]+item[3:11]])] for item in ev_clean]
    ev_clean[0][-1] = "Median"
    print("\tDone\n")

    print("\tSort the rows by : Unique ID (ascending), Missing Values (ascending), and Median (descending).")
    ev_clean = [ev_clean[0]] + sorted(ev_clean[1:], key = lambda x: (x[-3],x[-2],-x[-1]))
    print("\tDone\n")

    print(f"\tRemoving duplicate PSMs. Keeping least missing values and \nhighest median intensity")
    ev_clean = [ev_clean[0]] + keep_first(ev_clean[1:], -3)
    print(f"\t{len(ev_clean)-1} remaining\n")

    print(f"\tAdding identifier for Phosphotyrosine containing")
    ev_clean = [item + ["Y(Phospho" in item[mod_seq_col]] for item in ev_clean]
    ev_clean[0][-1] = "ispY"
    print("\tDone\n")

    print("\tUpdating experiment column to be human-readable:")
    print("\t\t1.0 == Control$+\Phi$SDM")
    print("\t\t2.0 == Control")
    print("\t\t3.0 == BOOST$+\Phi$SDM")
    print("\t\t4.0 == BOOST")
    experiment_ind = ev_clean[0].index("Experiment")
    for item in ev_clean:
        if item[experiment_ind] == 1.0:
            item[experiment_ind] = r"Control$+\Phi$SDM"
        elif item[experiment_ind] == 2.0:
            item[experiment_ind] = r"Control"
        elif item[experiment_ind] == 3.0:
            item[experiment_ind] = r"BOOST$+\Phi$SDM"
        elif item[experiment_ind] == 4.0:
            item[experiment_ind] = r"BOOST"
    print("\tDone\n")

    print(f"\tRemove all non-phosphotyrosine peptides")
    ev_clean_py = [item for item in ev_clean if item[-1]]
    print(f"\t{len(ev_clean_py)-1} remaining\n")
    
    # New columns we will add to the matrix                        # Indices
    heads = ev_clean_py[0] + ["10X mean", "3X mean", "1X mean",    # -18,-17,-16
                              "10X/3X", "10X/1X", "3X/1X",         # -15,-14,-13
                              "10X SD", "3X SD", "1X SD",          # -12,-11,-10
                              "10X CV", "3X CV", "1X CV",          # -9,-8,-7
                              "10X/3X P", "10X/1X P", "3X/1X P",   # -6,-5,-4
                              "10X/3X Q", "10X/1X Q", "3X/1X Q"]   # -3,-2,-1
    
    print("\tCalculating means (at least 1 reporter intensity in all replicates)")
    # Take the mean of each condition and make a new column for each
    # Note that the headers are removed in the first calculations
    ev_clean_py = [item + [sh.mean([item[x] for x in ten_x])] for item in ev_clean_py[1:]]
    ev_clean_py = [item + [sh.mean([item[x] for x in three_x])] for item in ev_clean_py]
    ev_clean_py = [item + [sh.mean([item[x] for x in one_x])] for item in ev_clean_py]
    print("\tDone\n")
    
    print("\tCalculating ratios of conditions (at least 1 reporter intensity in all replicates)")
    # Take the ratio of each condition
    ev_clean_py = [item + [ratio(item[-3],item[-2])] for item in ev_clean_py]
    ev_clean_py = [item + [ratio(item[-4],item[-2])] for item in ev_clean_py]
    ev_clean_py = [item + [ratio(item[-4],item[-3])] for item in ev_clean_py]
    print("\tDone\n")

    print("\tCalcuting standard deviations (all 3 reporter intensities must be present)")
    # Calculate the SD for each condition
    ev_clean_py = [item + [sh.standard_deviation([item[x] for x in ten_x], threshold = 3)] for item in ev_clean_py]
    ev_clean_py = [item + [sh.standard_deviation([item[x] for x in three_x], threshold = 3)] for item in ev_clean_py]
    ev_clean_py = [item + [sh.standard_deviation([item[x] for x in one_x], threshold = 3)] for item in ev_clean_py]
    print("\tDone\n")

    print("\tCalculating CV percentages (all 3 reporter intensities must be present)")
    # Calculate CV (as a percentage). This is easy because adding a column
    # shifts the index by one :)
    ev_clean_py = [item + [(item[-3]/item[-9]) * 100] for item in ev_clean_py]
    ev_clean_py = [item + [(item[-3]/item[-9]) * 100] for item in ev_clean_py]
    ev_clean_py = [item + [(item[-3]/item[-9]) * 100] for item in ev_clean_py]
    print("\tDone\n")

    print("\tPerforming Pairwise Welch's TTests (all 3 reporter intensities must be present)")
    # Do Welch's TTests
    ev_clean_py = [item + sh.TTest([item[x] for x in ten_x],
                                   [item[x] for x in three_x],
                                   test_type = "w",
                                   threshold = 3,
                                   labels = False).output[0]["pvalue"] for item in ev_clean_py]
    ev_clean_py = [item + sh.TTest([item[x] for x in ten_x],
                                   [item[x] for x in one_x],
                                   test_type = "w",
                                   threshold = 3,
                                   labels = False).output[0]["pvalue"] for item in ev_clean_py]
    ev_clean_py = [item + sh.TTest([item[x] for x in three_x],
                                   [item[x] for x in one_x],
                                   test_type = "w", 
                                   threshold = 3,
                                   labels = False).output[0]["pvalue"] for item in ev_clean_py]
    print("\tDone\n")

    print("\tPerforming Benjamini & Hochberg FDR Correction")
    # And correct for FDR using Benjamini & Hochberg. Again, each time
    # we add this to the list, the index increases by one
    inds, ps = index_ps([item[-3] for item in ev_clean_py])
    qs = list(sh.storey(list(ps), pi0=1)["qvalue"])
    qs = fill_nans(inds, qs, len(ev_clean_py))
    ev_clean_py = [ev_clean_py[i] + [qs[i]] for i in range(len(qs))]

    inds, ps = index_ps([item[-3] for item in ev_clean_py])
    qs = list(sh.storey(list(ps), pi0=1)["qvalue"])
    qs = fill_nans(inds, qs, len(ev_clean_py))
    ev_clean_py = [ev_clean_py[i] + [qs[i]] for i in range(len(qs))]

    inds, ps = index_ps([item[-3] for item in ev_clean_py])
    qs = list(sh.storey(list(ps),pi0=1)["qvalue"])
    qs = fill_nans(inds, qs, len(ev_clean_py))
    ev_clean_py = [ev_clean_py[i] + [qs[i]] for i in range(len(qs))]
    print("\tDone\n")
    
    print("\tParsing the data by experiment")
    experiments = gh.bin_by_col([heads] + ev_clean_py, experiment_ind)
    for key in list(experiments.keys()):
        print(f"\t\t{key:^20}")
    print("\tDone\n")
    
    print("\tLog2 Transforming Replicates for Correlation Plots")
    
    print("\t\tControl$+\Phi$SDM 1.0 mg")
    csten_x_r1 = [log2(item[1]) for item in experiments["Control$+\Phi$SDM"][1:]]
    csten_x_r2 = [log2(item[3]) for item in experiments["Control$+\Phi$SDM"][1:]]
    csten_x_r3 = [log2(item[5]) for item in experiments["Control$+\Phi$SDM"][1:]]
    
    print("\t\tControl$+\Phi$SDM 0.3 mg")
    csthree_x_r1 = [log2(item[4]) for item in experiments["Control$+\Phi$SDM"][1:]]
    csthree_x_r2 = [log2(item[6]) for item in experiments["Control$+\Phi$SDM"][1:]]
    csthree_x_r3 = [log2(item[8]) for item in experiments["Control$+\Phi$SDM"][1:]]
    
    print("\t\tControl$+\Phi$SDM 0.1\n")
    csone_x_r1 = [log2(item[7]) for item in experiments["Control$+\Phi$SDM"][1:]]
    csone_x_r2 = [log2(item[9]) for item in experiments["Control$+\Phi$SDM"][1:]]
    csone_x_r3 = [log2(item[10]) for item in experiments["Control$+\Phi$SDM"][1:]]

    print("\t\tControl 1.0 mg")
    cten_x_r1 = [log2(item[1]) for item in experiments["Control"][1:]]
    cten_x_r2 = [log2(item[3]) for item in experiments["Control"][1:]]
    cten_x_r3 = [log2(item[5]) for item in experiments["Control"][1:]]
    
    print("\t\tControl 0.3 mg")
    cthree_x_r1 = [log2(item[4]) for item in experiments["Control"][1:]]
    cthree_x_r2 = [log2(item[6]) for item in experiments["Control"][1:]]
    cthree_x_r3 = [log2(item[8]) for item in experiments["Control"][1:]]
    
    print("\t\tControl 0.1 mg")
    cone_x_r1 = [log2(item[7]) for item in experiments["Control"][1:]]
    cone_x_r2 = [log2(item[9]) for item in experiments["Control"][1:]]
    cone_x_r3 = [log2(item[10]) for item in experiments["Control"][1:]]
    
    print("\t\tBOOST 1.0 mg")
    bsten_x_r1 = [log2(item[1]) for item in experiments["BOOST$+\Phi$SDM"][1:]]
    bsten_x_r2 = [log2(item[3]) for item in experiments["BOOST$+\Phi$SDM"][1:]]
    bsten_x_r3 = [log2(item[5]) for item in experiments["BOOST$+\Phi$SDM"][1:]]
    
    print("\t\tBOOST 0.3 mg")
    bsthree_x_r1 = [log2(item[4]) for item in experiments["BOOST$+\Phi$SDM"][1:]]
    bsthree_x_r2 = [log2(item[6]) for item in experiments["BOOST$+\Phi$SDM"][1:]]
    bsthree_x_r3 = [log2(item[8]) for item in experiments["BOOST$+\Phi$SDM"][1:]]
    
    print("\t\tBOOST 0.1 mg")
    bsone_x_r1 = [log2(item[7]) for item in experiments["BOOST$+\Phi$SDM"][1:]]
    bsone_x_r2 = [log2(item[9]) for item in experiments["BOOST$+\Phi$SDM"][1:]]
    bsone_x_r3 = [log2(item[10]) for item in experiments["BOOST$+\Phi$SDM"][1:]]
    
    print("\t\tBOOST$+\Phi$SDM 1.0 mg")
    bten_x_r1 = [log2(item[1]) for item in experiments["BOOST"][1:]]
    bten_x_r2 = [log2(item[3]) for item in experiments["BOOST"][1:]]
    bten_x_r3 = [log2(item[5]) for item in experiments["BOOST"][1:]]
    
    print("\t\tBOOST$+\Phi$SDM 0.3 mg")
    bthree_x_r1 = [log2(item[4]) for item in experiments["BOOST"][1:]]
    bthree_x_r2 = [log2(item[6]) for item in experiments["BOOST"][1:]]
    bthree_x_r3 = [log2(item[8]) for item in experiments["BOOST"][1:]]
    
    print("\t\tBOOST$+\Phi$SDM 0.1 mg")
    bone_x_r1 = [log2(item[7]) for item in experiments["BOOST"][1:]]
    bone_x_r2 = [log2(item[9]) for item in experiments["BOOST"][1:]]
    bone_x_r3 = [log2(item[10]) for item in experiments["BOOST"][1:]]
    
    print("\tLog10 Transforming q-values & Ratios for Volcano Plots")
    
    print("\t\tControl$+\Phi$SDM q-values")
    cs_103_q = [-log10(q[-3]) for q in experiments["Control$+\Phi$SDM"][1:]]
    cs_101_q = [-log10(q[-2]) for q in experiments["Control$+\Phi$SDM"][1:]]
    cs_31_q = [-log10(q[-1]) for q in experiments["Control$+\Phi$SDM"][1:]]

    print("\t\tControl$+\Phi$SDM ratios")
    cs_103_r = [log10(q[-15]) for q in experiments["Control$+\Phi$SDM"][1:]]
    cs_101_r = [log10(q[-14]) for q in experiments["Control$+\Phi$SDM"][1:]]
    cs_31_r = [log10(q[-13]) for q in experiments["Control$+\Phi$SDM"][1:]]

    print("\t\tControl$+\Phi$SDM Removing nanpairs and counting q > 0.05")
    cs_103_rq = remove_nanpairs(cs_103_r, cs_103_q)
    cs_103_qcount = sum([1 for _ in cs_103_rq[1] if _ > -log10(0.05)])
    cs_101_rq = remove_nanpairs(cs_101_r, cs_101_q)
    cs_101_qcount = sum([1 for _ in cs_101_rq[1] if _ > -log10(0.05)])
    cs_31_rq = remove_nanpairs(cs_31_r, cs_31_q)
    cs_31_qcount = sum([1 for _ in cs_31_rq[1] if _ > -log10(0.05)])

    cs_rq = [cs_101_rq, cs_103_rq, cs_31_rq]
    cs_counts = [cs_101_qcount, cs_103_qcount, cs_31_qcount]
    
    print("\t\tControl q-values")
    c_103_q = [-log10(q[-3]) for q in experiments["Control"][1:]]
    c_101_q = [-log10(q[-2]) for q in experiments["Control"][1:]]
    c_31_q = [-log10(q[-1]) for q in experiments["Control"][1:]]

    print("\t\tControl ratios")
    c_103_r = [log10(q[-15]) for q in experiments["Control"][1:]]
    c_101_r = [log10(q[-14]) for q in experiments["Control"][1:]]
    c_31_r = [log10(q[-13]) for q in experiments["Control"][1:]]

    print("\t\tControl Removing nanpairs and counting q > 0.05")
    c_103_rq = remove_nanpairs(c_103_r, c_103_q)
    c_103_qcount = sum([1 for _ in c_103_rq[1] if _ > -log10(0.05)])
    c_101_rq = remove_nanpairs(c_101_r, c_101_q)
    c_101_qcount = sum([1 for _ in c_101_rq[1] if _ > -log10(0.05)])
    c_31_rq = remove_nanpairs(c_31_r, c_31_q)
    c_31_qcount = sum([1 for _ in c_31_rq[1] if _ > -log10(0.05)])

    c_rq = [c_101_rq, c_103_rq, c_31_rq]
    c_counts = [c_101_qcount, c_103_qcount, c_31_qcount]

    print("\t\tBOOST$+\Phi$SDM q-values")
    bs_103_q = [-log10(q[-3]) for q in experiments["BOOST$+\Phi$SDM"][1:]]
    bs_101_q = [-log10(q[-2]) for q in experiments["BOOST$+\Phi$SDM"][1:]]
    bs_31_q = [-log10(q[-1]) for q in experiments["BOOST$+\Phi$SDM"][1:]]

    print("\t\tBOOST$+\Phi$SDM ratios")
    bs_103_r = [log10(q[-15]) for q in experiments["BOOST$+\Phi$SDM"][1:]]
    bs_101_r = [log10(q[-14]) for q in experiments["BOOST$+\Phi$SDM"][1:]]
    bs_31_r = [log10(q[-13]) for q in experiments["BOOST$+\Phi$SDM"][1:]]

    print("\t\tBOOST$+\Phi$SDM Removing nanpairs and counting q > 0.05")
    bs_103_rq = remove_nanpairs(bs_103_r, bs_103_q)
    bs_103_qcount = sum([1 for _ in bs_103_rq[1] if _ > -log10(0.05)])
    bs_101_rq = remove_nanpairs(bs_101_r, bs_101_q)
    bs_101_qcount = sum([1 for _ in bs_101_rq[1] if _ > -log10(0.05)])
    bs_31_rq = remove_nanpairs(bs_31_r, bs_31_q)
    bs_31_qcount = sum([1 for _ in bs_31_rq[1] if _ > -log10(0.05)])

    bs_rq = [bs_101_rq, bs_103_rq, bs_31_rq]
    bs_counts = [bs_101_qcount, bs_103_qcount, bs_31_qcount]

    print("\t\tBOOST q-values")
    b_103_q = [-log10(q[-3]) for q in experiments["BOOST"][1:]]
    b_101_q = [-log10(q[-2]) for q in experiments["BOOST"][1:]]
    b_31_q = [-log10(q[-1]) for q in experiments["BOOST"][1:]]

    print("\t\tBOOST ratios")
    b_103_r = [log10(q[-15]) for q in experiments["BOOST"][1:]]
    b_101_r = [log10(q[-14]) for q in experiments["BOOST"][1:]]
    b_31_r = [log10(q[-13]) for q in experiments["BOOST"][1:]]

    print("\t\tBOOST Removing nanpairs and counting q > 0.05")
    b_103_rq = remove_nanpairs(b_103_r, b_103_q)
    b_103_qcount = sum([1 for _ in b_103_rq[1] if _ > -log10(0.05)])
    b_101_rq = remove_nanpairs(b_101_r, b_101_q)
    b_101_qcount = sum([1 for _ in b_101_rq[1] if _ > -log10(0.05)])
    b_31_rq = remove_nanpairs(b_31_r, b_31_q)
    b_31_qcount = sum([1 for _ in b_31_rq[1] if _ > -log10(0.05)])

    b_rq = [b_101_rq, b_103_rq, b_31_rq]
    b_counts = [b_101_qcount, b_103_qcount, b_31_qcount]

    qr_data = [cs_rq, c_rq, bs_rq, b_rq]
    # Grab the data for the ratio suppression graph
    ratios = gh.transpose(*[[item[0] for item in group] for group in qr_data])
    without_filt = gh.transpose(*[[cs_101_r, cs_103_r, cs_31_r],
                [c_101_r, c_103_r, c_31_r],
               [bs_101_r, bs_103_r, bs_31_r],
               [b_101_r, b_103_r, b_31_r]])
    without_filt = [[[d for d in col if d == d] for col in group] for group in without_filt]
    loggedmean = [ratios, without_filt]
    ratiomeds = [[[10**sh.median(col) for col in group] for group in ratios],
             [[10**sh.median(col) for col in group] for group in without_filt]]
    #
    qcounts = [cs_counts, c_counts, bs_counts, b_counts]
    
    print("\tCounting the number of unique pTyr peptides with quantified reporters")
    print("\t\tControl$+\Phi$SDM")
    cs_report_count = [sum([1 for j in range(1,len(experiments["Control$+\Phi$SDM"])) if experiments["Control$+\Phi$SDM"][j][i] == experiments["Control$+\Phi$SDM"][j][i]])
                       for i in range(11)]
    print("\t\tControl")
    c_report_count = [sum([1 for j in range(1,len(experiments["Control"])) if experiments["Control"][j][i] == experiments["Control"][j][i]])
                       for i in range(11)]
    print("\t\tBOOST$+\Phi$SDM")
    bs_report_count = [sum([1 for j in range(1,len(experiments["BOOST$+\Phi$SDM"])) if experiments["BOOST$+\Phi$SDM"][j][i] == experiments["BOOST$+\Phi$SDM"][j][i]])
                       for i in range(11)]
    print("\t\tBOOST")
    b_report_count = [sum([1 for j in range(1,len(experiments["BOOST"])) if experiments["BOOST"][j][i] == experiments["BOOST"][j][i]])
                       for i in range(11)]
    report_counts = [[cs_report_count, c_report_count],[bs_report_count, b_report_count]]
    
    print("\tComputing the percentage of unique pTyr peptides with missing reporter intensities")
    print("\t\tControl$+\Phi$SDM")
    cs_report_perc = [sum([1 for j in range(1,len(experiments["Control$+\Phi$SDM"])) if experiments["Control$+\Phi$SDM"][j][i] != experiments["Control$+\Phi$SDM"][j][i]])/len(experiments["Control$+\Phi$SDM"][1:])*100
                       for i in range(11)]
    print("\t\tControl")
    c_report_perc = [sum([1 for j in range(1,len(experiments["Control"])) if experiments["Control"][j][i] != experiments["Control"][j][i]])/len(experiments["Control"][1:])*100 
                       for i in range(11)]
    print("\t\tBOOST$+\Phi$SDM")
    bs_report_perc = [sum([1 for j in range(1,len(experiments["BOOST$+\Phi$SDM"])) if experiments["BOOST$+\Phi$SDM"][j][i] != experiments["BOOST$+\Phi$SDM"][j][i]])/len(experiments["BOOST$+\Phi$SDM"][1:])*100
                       for i in range(11)]
    print("\t\tBOOST")
    b_report_perc = [sum([1 for j in range(1,len(experiments["BOOST"])) if experiments["BOOST"][j][i] != experiments["BOOST"][j][i]])/len(experiments["BOOST"][1:])*100
                       for i in range(11)]

    report_percs = [[cs_report_perc, c_report_perc], [bs_report_perc, b_report_perc]]
    
    print("\tCounting the number of unique pTyr peptides with missing reporter intensities")
    print("\t\tControl$+\Phi$SDM")
    cs_miss_count = [sum([1 for j in range(1,len(experiments["Control$+\Phi$SDM"])) if experiments["Control$+\Phi$SDM"][j][i] != experiments["Control$+\Phi$SDM"][j][i]])
                       for i in range(11)]
    print("\t\tControl")
    c_miss_count = [sum([1 for j in range(1,len(experiments["Control"])) if experiments["Control"][j][i] != experiments["Control"][j][i]])
                       for i in range(11)]
    print("\t\tBOOST$+\Phi$SDM")
    bs_miss_count = [sum([1 for j in range(1,len(experiments["BOOST$+\Phi$SDM"])) if experiments["BOOST$+\Phi$SDM"][j][i] != experiments["BOOST$+\Phi$SDM"][j][i]])
                       for i in range(11)]
    print("\t\tBOOST")
    b_miss_count = [sum([1 for j in range(1,len(experiments["BOOST"])) if experiments["BOOST"][j][i] != experiments["BOOST"][j][i]])
                       for i in range(11)]

    miss_counts = [[cs_miss_count, c_miss_count],[bs_miss_count, b_miss_count]]
    
    print("\tGather the reporter intensities for each replicate and Log10 transform them")
    print("\t\tControl$+\Phi$SDM")
                         # Matrix position[row][column]         for all columns except header                           if the value is not nan
    cs_report_signal = [[experiments["Control$+\Phi$SDM"][j][i] for j in range(1,len(experiments["Control$+\Phi$SDM"])) if experiments["Control$+\Phi$SDM"][j][i] == experiments["Control$+\Phi$SDM"][j][i]]
                        for i in range(11)]  # For columns 0-10, which are the reporter channels
    cs_meds = [sh.median(channel) for channel in cs_report_signal]
    cs_report_signal = [[log10(data) for data in channel] for channel in cs_report_signal]
    print("\t\tControl")
    c_report_signal = [[experiments["Control"][j][i] for j in range(1,len(experiments["Control"])) if experiments["Control"][j][i] == experiments["Control"][j][i]]
                       for i in range(11)]
    c_meds = [sh.median(channel) for channel in c_report_signal]
    c_report_signal = [[log10(data) for data in channel] for channel in c_report_signal]
    print("\t\tBOOST$+\Phi$SDM")
    bs_report_signal = [[experiments["BOOST$+\Phi$SDM"][j][i] for j in range(1,len(experiments["BOOST$+\Phi$SDM"])) if experiments["BOOST$+\Phi$SDM"][j][i] == experiments["BOOST$+\Phi$SDM"][j][i]]
                       for i in range(11)]
    bs_meds = [sh.median(channel) for channel in bs_report_signal]
    bs_report_signal = [[log10(data) for data in channel] for channel in bs_report_signal]
    print("\t\tBOOST")
    b_report_signal = [[experiments["BOOST"][j][i] for j in range(1,len(experiments["BOOST"])) if experiments["BOOST"][j][i] == experiments["BOOST"][j][i]]
                       for i in range(11)]
    b_meds = [sh.median(channel) for channel in b_report_signal]
    b_report_signal = [[log10(data) for data in channel] for channel in b_report_signal]
    meds_list = [cs_meds, c_meds, bs_meds, b_meds]
    sigsnals = [cs_report_signal, c_report_signal, bs_report_signal, b_report_signal]
    
    print("\tPrepare data for making Venn Diagrams")
    print("\t\tGetting peptides observed in experimental channels of Control$+\Phi$SDM")
    controlsdm_peps = [experiments["Control$+\Phi$SDM"][0]] + [row for row in experiments["Control$+\Phi$SDM"][1:] if any([data == data for data in [row[1]] + row[3:11]])]
    print(f"\t\tTotal peptides identified in the Control: {len(controlsdm_peps)}\n")
    u_id = experiments["Control"][0].index("Unique ID")
    controlsdm_peps_f = [item[u_id][:-3] for item in controlsdm_peps[1:]]
    print("\t\tGetting peptides observed in experimental channels of Control")
    control_peps =  [experiments["Control"][0]] + [row for row in experiments["Control"][1:] if any([data == data for data in [row[1]] + row[3:11]])]
    print(f"\t\tTotal peptides identified in the Control: {len(control_peps)}\n")
    control_peps_f = [row[u_id][:-3] for row in control_peps[1:]]
    
    print("\t\tGetting peptides observed in experimental channels of BOOST$+\Phi$SDM")
    boostsdm_peps = gh.filter_matrix(experiments["BOOST$+\Phi$SDM"], "PV Boost",
                                  0, False)
    print(f"\t\tNumber of peptides observed in BOOST$+\Phi$SDM: {len(boostsdm_peps)}\n")
    boostsdm_peps = [row for row in boostsdm_peps if any([data == data for data in [row[1]] + row[3:11]])]
    print(f"\t\tSubset whose peptides were also seen in an experimental channel: {len(boostsdm_peps)}\n")
    boostsdm_peps_f = [item[u_id][:-3] for item in boostsdm_peps[1:]]
    
    print("\t\tGetting peptides observed in experimental channels of BOOST")
    boost_peps = gh.filter_matrix(experiments["BOOST"], "PV Boost",
                               0, False)
    print(f"\t\tNumber of peptides observed in BOOST: {len(boost_peps)}\n")
    boost_peps =  [row for row in boost_peps if any([data == data for data in [row[1]] + row[3:11]])]
    print(f"\t\tSubset whose peptides were also seen in an experimental channel: {len(boost_peps)}\n")
    boost_peps_f = [item[u_id][:-3] for item in boost_peps[1:]]
    
    print("\tPrepare data for making BOOST Factor Histograms")
    boost_peps = [boost_peps[0] + ["ID no exp"]] + [row + [row[u_id][:-3]] for row in boost_peps[1:]]
    boostsdm_peps = [boostsdm_peps[0] + ["ID no exp"]] + [row + [row[u_id][:-3]] for row in boostsdm_peps[1:]]
    control_peps = [control_peps[0] + ["ID no exp"]] + [row + [row[u_id][:-3]] for row in control_peps[1:]]
    controlsdm_peps = [controlsdm_peps[0] + ["ID no exp"]] + [row + [row[u_id][:-3]] for row in controlsdm_peps[1:]]
    print("\t\tUsing UniqueIDs without Experiment numbers to determine peptide overlap")
    control_mod_seq = list(set(gh.grab_col(control_peps, "ID no exp")[1:]))
    boost_gained_peps = gh.filter_matrix(boost_peps, "ID no exp",
                                      control_mod_seq, True, compare = "not in")
    print(f"\t\tBoost gained peptides: {len(boost_gained_peps)}")
    boost_control_peps = gh.filter_matrix(boost_peps, "ID no exp",
                                      control_mod_seq, True, compare = "in")
    print(f"\t\tBoost-Control overlap peptides: {len(boost_control_peps)}")
    print(f"\t\tApplying the BOOST Factor function to BOOST Gained Peptides")
    boost_gained_fact = boost_factor(boost_gained_peps, 0, [1,3,4,5,6,7,8,9,10])
    boost_gained_peps = [boost_gained_peps[0] + ["Boost Factor"]] + [boost_gained_peps[i+1] + [boost_gained_fact[i]] for i in range(len(boost_gained_fact))]
    print(f"\t\tApplying the BOOST Factor function to BOOST-Control Peptides")
    boost_control_fact = boost_factor(boost_control_peps, 0, [1,3,4,5,6,7,8,9,10])
    boost_control_peps = [boost_control_peps[0] + ["Boost Factor"]] + [boost_control_peps[i+1] + [boost_control_fact[i]] for i in range(len(boost_control_fact))]
    controlsdm_mod_seq = list(set(gh.grab_col(controlsdm_peps, "ID no exp")[1:]))
    boostsdm_gained_peps = gh.filter_matrix(boostsdm_peps, "ID no exp",
                                      controlsdm_mod_seq, True, compare = "not in")
    print(f"\t\tBoost+SDM gained peptides: {len(boostsdm_gained_peps)}\n")
    boostsdm_controlsdm_peps = gh.filter_matrix(boostsdm_peps, "ID no exp",
                                      controlsdm_mod_seq, True, compare = "in")
    print(f"\t\tBoost+SDM-Control+SDM overlap peptides: {len(boostsdm_controlsdm_peps)}\n")
    print(f"\t\tApplying the BOOST Factor function to BOOST$+\Phi$SDM Gained Peptides")
    boostsdm_gained_fact = boost_factor(boostsdm_gained_peps, 0, [1,3,4,5,6,7,8,9,10])
    boostsdm_gained_peps = [boostsdm_gained_peps[0] + ["Boost Factor"]] + [boostsdm_gained_peps[i+1] + [boostsdm_gained_fact[i]] for i in range(len(boostsdm_gained_fact))]
    print(f"\t\tApplying the BOOST Factor function to BOOST$+\Phi$SDM-Control$+\Phi$SDM Peptides\n")
    boostsdm_controlsdm_fact = boost_factor(boostsdm_controlsdm_peps, 0, [1,3,4,5,6,7,8,9,10])
    boostsdm_controlsdm_peps = [boostsdm_controlsdm_peps[0] + ["Boost Factor"]] + [boostsdm_controlsdm_peps[i+1] + [boostsdm_controlsdm_fact[i]] for i in range(len(boostsdm_controlsdm_fact))]
    
    facts = [[boost_gained_fact, boostsdm_gained_fact],
             [boost_control_fact, boostsdm_controlsdm_fact]]
    
    meds = [[sh.median(facts[i][j]) for j in range(2)] for i in range(2)]
    
    overunder = [[get_above_below(facts[i][j]) for j in range(2)] for i in range(2)]


    print("\tPreparing Data for BOOST Factor CDFs")
    boost_ind = boost_gained_peps[0].index("Boost Factor")
    print("\t\tCreating BOOST Gained Matrix with 1.0 mg - 0.1 mg q-value < 0.05")
    b_ten_one_q = gh.filter_matrix(boost_gained_peps, "10X/1X Q", float("nan"), keep = False)
    b_ten_one_q = gh.filter_matrix(boost_gained_peps, "10X/1X Q", 0.05, keep = True, compare = "<")
    b_ten_one_q = [row[boost_ind] for row in b_ten_one_q[1:] if row[boost_ind] == row[-1]]
    print("\t\tCreating BOOST Gained Matrix with 1.0 mg - 0.3 mg q-value < 0.05")
    b_ten_three_q = gh.filter_matrix(boost_gained_peps, "10X/3X Q", float("nan"), keep = False)
    b_ten_three_q = gh.filter_matrix(boost_gained_peps, "10X/3X Q", 0.05, keep = True, compare = "<")
    b_ten_three_q = [row[boost_ind] for row in b_ten_three_q[1:] if row[boost_ind] == row[boost_ind]]
    print("\t\tCreating BOOST Gained Matrix with 0.3 mg - 0.1 mg q-value < 0.05")
    b_three_one_q = gh.filter_matrix(boost_gained_peps, "3X/1X Q", float("nan"), keep = False)
    b_three_one_q = gh.filter_matrix(boost_gained_peps, "3X/1X Q", 0.05, keep = True, compare = "<")
    b_three_one_q = [row[boost_ind] for row in b_three_one_q[1:] if row[boost_ind] == row[boost_ind]]
    print("\t\tCreating BOOST$+\Phi$SDM Gained Matrix with 1.0 mg - 0.1 mg q-value < 0.05")
    bs_ten_one_q = gh.filter_matrix(boostsdm_gained_peps, "10X/1X Q", float("nan"), keep = False)
    bs_ten_one_q = gh.filter_matrix(boostsdm_gained_peps, "10X/1X Q", 0.05, keep = True, compare = "<")
    bs_ten_one_q = [row[boost_ind] for row in bs_ten_one_q[1:] if row[boost_ind] == row[-1]]
    print("\t\tCreating BOOST$+\Phi$SDM Gained Matrix with 1.0 mg - 0.3 mg q-value < 0.05")
    bs_ten_three_q = gh.filter_matrix(boostsdm_gained_peps, "10X/3X Q", float("nan"), keep = False)
    bs_ten_three_q = gh.filter_matrix(boostsdm_gained_peps, "10X/3X Q", 0.05, keep = True, compare = "<")
    bs_ten_three_q = [row[boost_ind] for row in bs_ten_three_q[1:] if row[boost_ind] == row[boost_ind]]
    print("\t\tCreating BOOST$+\Phi$SDM Gained Matrix with 0.3 mg - 0.1 mg q-value < 0.05")
    bs_three_one_q = gh.filter_matrix(boostsdm_gained_peps, "3X/1X Q", float("nan"), keep = False)
    bs_three_one_q = gh.filter_matrix(boostsdm_gained_peps, "3X/1X Q", 0.05, keep = True, compare = "<")
    bs_three_one_q = [row[boost_ind] for row in bs_three_one_q[1:] if row[boost_ind] == row[boost_ind]]
    print("\t\tCreating BOOST Gained Matrix with 1.0 mg - 0.1 mg ratios")
    b_ten_one_r = gh.filter_matrix(boost_gained_peps, "10X/1X", float("nan"), keep = False)
    b_ten_one_r = gh.filter_matrix(boost_gained_peps, "10X/1X", 0, keep = True, compare = ">=")
    b_ten_one_r = [row[boost_ind] for row in b_ten_one_r[1:] if row[boost_ind] == row[-1]]
    print("\t\tCreating BOOST Gained Matrix with 1.0 mg - 0.3 mg ratios")
    b_ten_three_r = gh.filter_matrix(boost_gained_peps, "10X/3X", float("nan"), keep = False)
    b_ten_three_r = gh.filter_matrix(boost_gained_peps, "10X/3X", 0, keep = True, compare = ">=")
    b_ten_three_r = [row[boost_ind] for row in b_ten_three_r[1:] if row[boost_ind] == row[boost_ind]]
    print("\t\tCreating BOOST Gained Matrix with 0.3 mg - 0.1 mg ratios")
    b_three_one_r = gh.filter_matrix(boost_gained_peps, "3X/1X", float("nan"), keep = False)
    b_three_one_r = gh.filter_matrix(boost_gained_peps, "3X/1X", 0, keep = True, compare = ">=")
    b_three_one_r = [row[boost_ind] for row in b_three_one_r[1:] if row[boost_ind] == row[boost_ind]]
    print("\t\tCreating BOOST$+\Phi$SDM Gained Matrix with 1.0 mg - 0.1 mg ratios")
    bs_ten_one_r = gh.filter_matrix(boostsdm_gained_peps, "10X/1X", float("nan"), keep = False)
    bs_ten_one_r = gh.filter_matrix(boostsdm_gained_peps, "10X/1X", 0, keep = True, compare = ">=")
    bs_ten_one_r = [row[boost_ind] for row in bs_ten_one_r[1:] if row[boost_ind] == row[-1]]
    print("\t\tCreating BOOST$+\Phi$SDM Gained Matrix with 1.0 mg - 0.3 mg ratios")
    bs_ten_three_r = gh.filter_matrix(boostsdm_gained_peps, "10X/3X", float("nan"), keep = False)
    bs_ten_three_r = gh.filter_matrix(boostsdm_gained_peps, "10X/3X", 0, keep = True, compare = ">=")
    bs_ten_three_r = [row[boost_ind] for row in bs_ten_three_r[1:] if row[boost_ind] == row[boost_ind]]
    print("\t\tCreating BOOST$+\Phi$SDM Gained Matrix with 0.3 mg - 0.1 mg ratios\n")
    bs_three_one_r = gh.filter_matrix(boostsdm_gained_peps, "3X/1X", float("nan"), keep = False)
    bs_three_one_r = gh.filter_matrix(boostsdm_gained_peps, "3X/1X", 0, keep = True, compare = ">=")
    bs_three_one_r = [row[boost_ind] for row in bs_three_one_r[1:] if row[boost_ind] == row[boost_ind]]

    cdf_data = [[[b_ten_one_q, b_ten_three_q, b_three_one_q], 
                 [bs_ten_one_q, bs_ten_three_q, bs_three_one_q]],
                [[b_ten_one_r, b_ten_three_r, b_three_one_r], 
                 [bs_ten_one_r, bs_ten_three_r, bs_three_one_r]]]
    print("\tGrabbing q-values and ratios for BOOST-Control Overlap Volcano Plots")
    print("\t\tCreating BOOST Gained list with 1.0 mg - 0.1 mg q-values")
    b_only_101_q = gh.grab_col(boost_gained_peps, "10X/1X Q")[1:]
    print("\t\tCreating BOOST Gained list with 1.0 mg - 0.1 mg ratios")
    b_only_101_r = gh.grab_col(boost_gained_peps, "10X/1X")[1:]
    print("\t\tCombining q-values and ratios for 1.0 mg - 0.1 mg BOOST")
    b_only_101_q, b_only_101_r = filter_nanpairs(b_only_101_q, b_only_101_r)
    print("\t\tCreating BOOST Gained list with 1.0 mg - 0.3 mg q-values")
    b_only_103_q = gh.grab_col(boost_gained_peps, "10X/3X Q")[1:]
    print("\t\tCreating BOOST Gained list with 1.0 mg - 0.3 mg ratios")
    b_only_103_r = gh.grab_col(boost_gained_peps, "10X/3X")[1:]
    print("\t\tCombining q-values and ratios for 1.0 mg - 0.3 mg BOOST")
    b_only_103_q, b_only_103_r = filter_nanpairs(b_only_103_q, b_only_103_r)
    print("\t\tCreating BOOST Gained list with 0.3 mg - 0.1 mg q-values")
    b_only_031_q = gh.grab_col(boost_gained_peps, "3X/1X Q")[1:]
    print("\t\tCreating BOOST Gained list with 0.3 mg - 0.1 mg ratios")
    b_only_031_r = gh.grab_col(boost_gained_peps, "3X/1X")[1:]
    print("\t\tCombining q-values and ratios for 0.3 mg - 0.1 mg BOOST")
    b_only_031_q, b_only_031_r = filter_nanpairs(b_only_031_q, b_only_031_r)
    print("\t\tCreating BOOST-Control list with 1.0 mg - 0.1 mg q-values")
    bc_only_101_q = gh.grab_col(boost_control_peps, "10X/1X Q")[1:]
    print("\t\tCreating BOOST-Control Gained list with 1.0 mg - 0.1 mg ratios")
    bc_only_101_r = gh.grab_col(boost_control_peps, "10X/1X")[1:]
    print("\t\tCombining q-values and ratios for 1.0 mg - 0.1 mg BOOST-Control")
    bc_only_101_q, bc_only_101_r = filter_nanpairs(bc_only_101_q, bc_only_101_r)
    print("\t\tCreating BOOST-Control list with 1.0 mg - 0.3 mg q-values")
    bc_only_103_q = gh.grab_col(boost_control_peps, "10X/3X Q")[1:]
    print("\t\tCreating BOOST-Control Gained list with 1.0 mg - 0.3 mg ratios")
    bc_only_103_r = gh.grab_col(boost_control_peps, "10X/3X")[1:]
    print("\t\tCombining q-values and ratios for 1.0 mg - 0.1 mg BOOST-Control")
    bc_only_103_q, bc_only_103_r = filter_nanpairs(bc_only_103_q, bc_only_103_r)
    print("\t\tCreating BOOST-Control list with 0.3 mg - 0.1 mg q-values")
    bc_only_031_q = gh.grab_col(boost_control_peps, "3X/1X Q")[1:]
    print("\t\tCreating BOOST-Control Gained list with 0.3 mg - 0.1 mg ratios")
    bc_only_031_r = gh.grab_col(boost_control_peps, "3X/1X")[1:]
    print("\t\tCombining q-values and ratios for 0.3 mg - 0.1 mg BOOST-Control")
    bc_only_031_q, bc_only_031_r = filter_nanpairs(bc_only_031_q, bc_only_031_r)
    boost_mod_seq = list(set(gh.grab_col(boost_peps, "ID no exp")[1:]))
    control_only_peps = gh.filter_matrix(control_peps, "ID no exp",
                                      boost_mod_seq, True, compare = "not in")
    control_boost_peps = gh.filter_matrix(control_peps, "ID no exp",
                                      boost_mod_seq, True, compare = "in")
    print("\t\tCreating Control Only list with 1.0 mg - 0.1 mg q-values")
    c_only_101_q = gh.grab_col(control_only_peps, "10X/1X Q")[1:]
    print("\t\tCreating Control Only list with 1.0 mg - 0.1 mg ratios")
    c_only_101_r = gh.grab_col(control_only_peps, "10X/1X")[1:]
    print("\t\tCombining q-values and ratios for 1.0 mg - 0.1 mg Control")
    c_only_101_q, c_only_101_r = filter_nanpairs(c_only_101_q, c_only_101_r)
    print("\t\tCreating Control Only list with 1.0 mg - 0.3 mg q-values")
    c_only_103_q = gh.grab_col(control_only_peps, "10X/3X Q")[1:]
    print("\t\tCreating Control Only list with 1.0 mg - 0.3 mg ratios")
    c_only_103_r = gh.grab_col(control_only_peps, "10X/3X")[1:]
    print("\t\tCombining q-values and ratios for 1.0 mg - 0.3 mg Control")
    c_only_103_q, c_only_103_r = filter_nanpairs(c_only_103_q, c_only_103_r)
    print("\t\tCreating Control Only list with 0.3 mg - 0.1 mg q-values")
    c_only_031_q = gh.grab_col(control_only_peps, "3X/1X Q")[1:]
    print("\t\tCreating Control Only list with 0.3 mg - 0.1 mg ratios")
    c_only_031_r = gh.grab_col(control_only_peps, "3X/1X")[1:]
    print("\t\tCombining q-values and ratios for 0.3 mg - 0.1 mg Control")
    c_only_031_q, c_only_031_r = filter_nanpairs(c_only_031_q, c_only_031_r)
    print("\t\tCreating Control-BOOST list with 1.0 mg - 0.1 mg q-values")
    cb_only_101_q = gh.grab_col(control_boost_peps, "10X/1X Q")[1:]
    print("\t\tCreating Control-BOOST list with 1.0 mg - 0.1 mg ratios")
    cb_only_101_r = gh.grab_col(control_boost_peps, "10X/1X")[1:]
    print("\t\tCombining q-values and ratios for 1.0 mg - 0.1 mg Control-BOOST")
    cb_only_101_q, cb_only_101_r = filter_nanpairs(cb_only_101_q, cb_only_101_r)
    print("\t\tCreating Control-BOOST list with 1.0 mg - 0.3 mg q-values")
    cb_only_103_q = gh.grab_col(control_boost_peps, "10X/3X Q")[1:]
    print("\t\tCreating Control-BOOST list with 1.0 mg - 0.3 mg ratios")
    cb_only_103_r = gh.grab_col(control_boost_peps, "10X/3X")[1:]
    print("\t\tCombining q-values and ratios for 1.0 mg - 0.3 mg Control-BOOST")
    cb_only_103_q, cb_only_103_r = filter_nanpairs(cb_only_103_q, cb_only_103_r)
    print("\t\tCreating Control-BOOST list with 0.3 mg - 0.1 mg q-values")
    cb_only_031_q = gh.grab_col(control_boost_peps, "3X/1X Q")[1:]
    print("\t\tCreating Control-BOOST list with 0.3 mg - 0.1 mg ratios")
    cb_only_031_r = gh.grab_col(control_boost_peps, "3X/1X")[1:]
    print("\t\tCreating Control-BOOST list with 0.3 mg - 0.1 mg ratios\n")
    cb_only_031_q, cb_only_031_r = filter_nanpairs(cb_only_031_q, cb_only_031_r)
    print("\t\tCreating BOOST$+\Phi$SDM Gained list with 1.0 mg - 0.1 mg q-values")
    bs_only_101_q = gh.grab_col(boostsdm_gained_peps, "10X/1X Q")[1:]
    print("\t\tCreating BOOST$+\Phi$SDM Gained list with 1.0 mg - 0.1 mg ratios")
    bs_only_101_r = gh.grab_col(boostsdm_gained_peps, "10X/1X")[1:]
    print("\t\tCombining q-values and ratios for 1.0 mg - 0.1 mg BOOST$+\Phi$SDM")
    bs_only_101_q, bs_only_101_r = filter_nanpairs(bs_only_101_q, bs_only_101_r)
    print("\t\tCreating BOOST$+\Phi$SDM Gained list with 1.0 mg - 0.3 mg q-values")
    bs_only_103_q = gh.grab_col(boostsdm_gained_peps, "10X/3X Q")[1:]
    print("\t\tCreating BOOST$+\Phi$SDM Gained list with 1.0 mg - 0.3 mg ratios")
    bs_only_103_r = gh.grab_col(boostsdm_gained_peps, "10X/3X")[1:]
    print("\t\tCombining q-values and ratios for 1.0 mg - 0.3 mg BOOST$+\Phi$SDM")
    bs_only_103_q, bs_only_103_r = filter_nanpairs(bs_only_103_q, bs_only_103_r)
    print("\t\tCreating BOOST$+\Phi$SDM Gained list with 0.3 mg - 0.1 mg q-value")
    bs_only_031_q = gh.grab_col(boostsdm_gained_peps, "3X/1X Q")[1:]
    print("\t\tCreating BOOST$+\Phi$SDM Gained list with 0.3 mg - 0.1 mg ratios")
    bs_only_031_r = gh.grab_col(boostsdm_gained_peps, "3X/1X")[1:]
    print("\t\tCombining q-values and ratios for 0.3 mg - 0.1 mg BOOST$+\Phi$SDM\n")
    bs_only_031_q, bs_only_031_r = filter_nanpairs(bs_only_031_q, bs_only_031_r)
    print("\tGrabbing q-values and ratios for BOOST$+\Phi$SDM-Control$+\Phi$SDM Overlap Volcano Plots")
    print("\t\tCreating BOOST$+\Phi$SDM-Control$+\Phi$SDM list with 1.0 mg - 0.1 mg q-values")
    bscs_only_101_q = gh.grab_col(boostsdm_controlsdm_peps, "10X/1X Q")[1:]
    print("\t\tCreating BOOST$+\Phi$SDM-Control$+\Phi$SDM list with 1.0 mg - 0.1 mg ratios")
    bscs_only_101_r = gh.grab_col(boostsdm_controlsdm_peps, "10X/1X")[1:]
    print("\t\tCombining q-values and ratios for 1.0 mg - 0.1 mg BOOST$+\Phi$SDM-Control$+\Phi$SDM")
    bscs_only_101_q, bscs_only_101_r = filter_nanpairs(bscs_only_101_q, bscs_only_101_r)
    print("\t\tCreating BOOST$+\Phi$SDM-Control$+\Phi$SDM list with 1.0 mg - 0.3 mg q-values")
    bscs_only_103_q = gh.grab_col(boostsdm_controlsdm_peps, "10X/3X Q")[1:]
    print("\t\tCreating BOOST$+\Phi$SDM-Control$+\Phi$SDM list with 1.0 mg - 0.3 mg ratios")
    bscs_only_103_r = gh.grab_col(boostsdm_controlsdm_peps, "10X/3X")[1:]
    print("\t\tCombining q-values and ratios for 1.0 mg - 0.3 mg BOOST$+\Phi$SDM-Control$+\Phi$SDM")
    bscs_only_103_q, bscs_only_103_r = filter_nanpairs(bscs_only_103_q, bscs_only_103_r)
    print("\t\tCreating BOOST$+\Phi$SDM-Control$+\Phi$SDM list with 0.3 mg - 0.1 mg q-values")
    bscs_only_031_q = gh.grab_col(boostsdm_controlsdm_peps, "3X/1X Q")[1:]
    print("\t\tCreating BOOST$+\Phi$SDM-Control$+\Phi$SDM list with 0.3 mg - 0.1 mg ratios")
    bscs_only_031_r = gh.grab_col(boostsdm_controlsdm_peps, "3X/1X")[1:]
    print("\t\tCombining q-values and ratios for 0.3 mg - 0.1 mg BOOST$+\Phi$SDM-Control$+\Phi$SDM")
    bscs_only_031_q, bscs_only_031_r = filter_nanpairs(bscs_only_031_q, bscs_only_031_r)
    
    boostsdm_mod_seq = list(set(gh.grab_col(boostsdm_peps, "ID no exp")[1:]))
    controlsdm_only_peps = gh.filter_matrix(controlsdm_peps, "ID no exp",
                                      boostsdm_mod_seq, True, compare = "not in")
    controlsdm_boostsdm_peps = gh.filter_matrix(controlsdm_peps, "ID no exp",
                                      boostsdm_mod_seq, True, compare = "in")
    print("\t\tCreating Control$+\Phi$SDM Only list with 1.0 mg - 0.1 mg q-values")
    cs_only_101_q = gh.grab_col(controlsdm_only_peps, "10X/1X Q")[1:]
    print("\t\tCreating Control$+\Phi$SDM Only list with 1.0 mg - 0.1 mg ratios")
    cs_only_101_r = gh.grab_col(controlsdm_only_peps, "10X/1X")[1:]
    print("\t\tCombining q-values and ratios for 1.0 mg - 0.1 mg Control$+\Phi$SDM Only")
    cs_only_101_q, cs_only_101_r = filter_nanpairs(cs_only_101_q, cs_only_101_r)
    print("\t\tCreating Control$+\Phi$SDM Only list with 1.0 mg - 0.3 mg q-values")
    cs_only_103_q = gh.grab_col(controlsdm_only_peps, "10X/3X Q")[1:]
    print("\t\tCreating Control$+\Phi$SDM Only list with 1.0 mg - 0.3 mg ratios")
    cs_only_103_r = gh.grab_col(controlsdm_only_peps, "10X/3X")[1:]
    print("\t\tCombining q-values and ratios for 1.0 mg - 0.3 mg Control$+\Phi$SDM Only")
    cs_only_103_q, cs_only_103_r = filter_nanpairs(cs_only_103_q, cs_only_103_r)
    print("\t\tCreating Control$+\Phi$SDM Only list with 0.3 mg - 0.1 mg q-values")
    cs_only_031_q = gh.grab_col(controlsdm_only_peps, "3X/1X Q")[1:]
    print("\t\tCreating Control$+\Phi$SDM Only list with 0.3 mg - 0.1 mg ratios")
    cs_only_031_r = gh.grab_col(controlsdm_only_peps, "3X/1X")[1:]
    print("\t\tCombining q-values and ratios for 0.3 mg - 0.1 mg Control$+\Phi$SDM Only")
    cs_only_031_q, cs_only_031_r = filter_nanpairs(cs_only_031_q, cs_only_031_r)
    print("\t\tCreating Control$+\Phi$SDM-BOOST$+\Phi$SDM list with 1.0 mg - 0.1 mg q-values")
    csbs_only_101_q = gh.grab_col(controlsdm_boostsdm_peps, "10X/1X Q")[1:]
    print("\t\tCreating Control$+\Phi$SDM-BOOST$+\Phi$SDM list with 1.0 mg - 0.1 mg ratios")
    csbs_only_101_r = gh.grab_col(controlsdm_boostsdm_peps, "10X/1X")[1:]
    print("\t\tCombining q-values and ratios for 1.0 mg - 0.1 mg Control$+\Phi$SDM-BOOST$+\Phi$SDM")
    csbs_only_101_q, csbs_only_101_r = filter_nanpairs(csbs_only_101_q, csbs_only_101_r)
    print("\t\tCreating Control$+\Phi$SDM-BOOST$+\Phi$SDM list with 1.0 mg - 0.3 mg q-values")
    csbs_only_103_q = gh.grab_col(controlsdm_boostsdm_peps, "10X/3X Q")[1:]
    print("\t\tCreating Control$+\Phi$SDM-BOOST$+\Phi$SDM list with 1.0 mg - 0.3 mg ratios")
    csbs_only_103_r = gh.grab_col(controlsdm_boostsdm_peps, "10X/3X")[1:]
    print("\t\tCombining q-values and ratios for 1.0 mg - 0.3 mg Control$+\Phi$SDM-BOOST$+\Phi$SDM")
    csbs_only_103_q, csbs_only_103_r = filter_nanpairs(csbs_only_103_q, csbs_only_103_r)
    print("\t\tCreating Control$+\Phi$SDM-BOOST$+\Phi$SDM list with 0.3 mg - 0.1 mg q-values")
    csbs_only_031_q = gh.grab_col(controlsdm_boostsdm_peps, "3X/1X Q")[1:]
    print("\t\tCreating Control$+\Phi$SDM-BOOST$+\Phi$SDM list with 0.3 mg - 0.1 mg ratios")
    csbs_only_031_r = gh.grab_col(controlsdm_boostsdm_peps, "3X/1X")[1:]
    print("\t\tCombining q-values and ratios for 0.3 mg - 0.1 mg Control$+\Phi$SDM-BOOST$+\Phi$SDM")
    csbs_only_031_q, csbs_only_031_r = filter_nanpairs(csbs_only_031_q, csbs_only_031_r)
    # Aggregate all the data for the first overlap volcano plot
    boost_control_qs = [[b_only_101_q, b_only_103_q, b_only_031_q],    # BOOST Gained
                        [bc_only_101_q, bc_only_103_q, bc_only_031_q], # BOOST \cap Control
                        [cb_only_101_q, cb_only_103_q, cb_only_031_q], # Control \cap BOOST
                        [c_only_101_q, c_only_103_q, c_only_031_q]]    # Control
    boost_control_rs = [[b_only_101_r, b_only_103_r, b_only_031_r],
                        [bc_only_101_r, bc_only_103_r, bc_only_031_r],
                        [cb_only_101_r, cb_only_103_r, cb_only_031_r],
                        [c_only_101_r, c_only_103_r, c_only_031_r]]
    # And add the counts for q < 0.05
    boost_control_qcounts = [[sum([1 for _ in q_list if _ < 0.05])
                         for q_list in groups]
                         for groups in boost_control_qs]
    # Then log10 transform the q-values and ratios
    boost_control_qs = [[[-log10(q) for q in q_list] 
                         for q_list in group] 
                        for group in boost_control_qs]
    boost_control_rs = [[[log10(r) for r in r_list] 
                         for r_list in group] 
                        for group in boost_control_rs]
    # Aggregate all the data for the second volcano plot
    boostsdm_controlsdm_qs = [[bs_only_101_q, bs_only_103_q, bs_only_031_q],
                              [bscs_only_101_q, bscs_only_103_q, bscs_only_031_q],
                              [csbs_only_101_q, csbs_only_103_q, csbs_only_031_q],
                              [cs_only_101_q, cs_only_103_q, cs_only_031_q]]
    boostsdm_controlsdm_rs = [[bs_only_101_r, bs_only_103_r, bs_only_031_r],
                              [bscs_only_101_r, bscs_only_103_r, bscs_only_031_r],
                              [csbs_only_101_r, csbs_only_103_r, csbs_only_031_r],
                              [cs_only_101_r, cs_only_103_r, cs_only_031_r]]
    # And add the counts for q < 0.05
    boostsdm_controlsdm_qcounts = [[sum([1 for _ in q_list if _ < 0.05])
                                   for q_list in groups]
                                   for groups in boostsdm_controlsdm_qs]
    # Then log10 transform the q-values and ratios
    boostsdm_controlsdm_qs = [[[-log10(q) for q in q_list] 
                               for q_list in group] 
                               for group in boostsdm_controlsdm_qs]
    boostsdm_controlsdm_rs = [[[log10(r) for r in r_list] 
                                for r_list in group] 
                                for group in boostsdm_controlsdm_rs]
    print("\tReading the 'maxquant_results/Phospho (STY)Sites.txt' file")
    sty = gh.read_file("maxquant_results/Phospho (STY)Sites.txt", delim = "\t")
    print("\t\tTransforming values and replacing empty strings by nans")
    sty = [gh.transform_values(row) for row in sty]
    sty = [gh.replace_value(row, "", float("nan")) for row in sty]
    print("\t\tDone\n")
    sty_split = gh.bin_by_col( sty[1:], 31)
    for key, value in sty_split.items():
        sty_split[key] = [sty[0]] + value
    sty_split = sorted(sty_split.items(), key = lambda x: len(x[1]),
                       reverse = True)
    if psp_rename:
        print("\tReading the database/phosphositeplus_ptm_data.txt' file\n")
        psp = gh.read_file("database/phosphositeplus_ptm_data.txt")[3:]
    else:
        print("\tReading the database/Phosphorylation_site_dataset' file\n")
        psp = gh.read_file("database/Phosphorylation_site_dataset")[3:]
    psp_database = gh.bin_by_col(psp, psp[0].index("ORGANISM"))
    #####################################################################################################
    print("STAGE 2: Writing the output files \n")
    # Grab the index of the localisation probability column
    loc_col = boost_gained_peps[0].index("Localisation Probability (Sequence)")
    # Aggregate the data we want for the new sheets
    new_sheets = [copy.copy(boost_gained_peps),
                  copy.copy(boost_control_peps),
                  copy.copy(control_boost_peps),
                  copy.copy(control_only_peps),
                  copy.copy(boostsdm_gained_peps),
                  copy.copy(boostsdm_controlsdm_peps),
                  copy.copy(controlsdm_boostsdm_peps),
                  copy.copy(controlsdm_only_peps)]
    print("\tAdding localisation probabilities column\n")
    new_sheets = [make_loc_probs(matrix, "Localisation Probability (Sequence)") for matrix in new_sheets]
    print("\tDone\n")
    print("\tAdding sequence windows and flanking sequences")
    for i in range(len(new_sheets)):
        if i in [0,1,4,5]:
            new_sheets[i] = add_sequence_window(new_sheets[i], sty)
        else:
            new_sheets[i] = add_sequence_window(new_sheets[i], sty)
    print("\tDone\n")
    print("\tAdding site numbers using PSP database")
    new_sheets = compare_seqs_to_database(psp_database["mouse"], new_sheets)
    print("\tDone\n")
    print("\tAdding Human equivalent sites")
    new_sheets = get_other_species_sites(psp_database["human"], new_sheets)
    print("\tDone\n")
    print("\tReordering the columns for writing")
    newsheets = []
    for i in range(len(new_sheets)):
        if i in [0,1,4,5]:
            newsheets.append(gh.select_cols(new_sheets[i], keep_boost_cols))
        else:
            contsheet = gh.select_cols(new_sheets[i], keep_cont_cols)
            contsheet = gh.add_col(contsheet, "BOOST Factor", newhead_pos = -7)
            newsheets.append(contsheet)
    print("\tSorting the rows by Gene Name, PSP phosphorylation site, and localisation probability.")
    newsheets = [[gh.transform_values(row) for row in sheet] for sheet in newsheets]
    newsheets = [[matrix[0]] + sorted(matrix[1:], key = lambda x: (x[0],
                                                                   x[4],
                                                                   -x[6])) for matrix in newsheets]
    print("\tMaking the lists into tab separated strings")
    newsheets = [[gh.list_to_str(row,
                                  delimiter = "\t",
                                  newline = True) for row in matrix]
                  for matrix in newsheets]
    names = ["curated_results/boost_gained_peptides.txt",
             "curated_results/boost_control_peptides.txt",
             "curated_results/control_boost_peptides.txt",
             "curated_results/control_only_peptides.txt",
             "curated_results/boostsdm_gained_peptides.txt",
             "curated_results/boostsdm_controlsdm_peptides.txt",
             "curated_results/controlsdm_boostsdm_peptides.txt",
             "curated_results/controlsdm_only_peptides.txt"]
    print("\tWriting the output files")
    i=0
    for sheet in newsheets:
        print(f"Writing {names[i]}\n")
        gh.write_outfile(sheet, filename = names[i], writestyle = "w")
        i+=1
    print("")
    ####################################################################################################
    print("STAGE 3: Making the plots")
    print("\tGenerating Figure 2E: Ratios")
    ratio_labs = ["1.0 mg : 0.1 mg",
                  "1.0 mg : 0.3 mg",
                  "0.3 mg : 0.1 mg",
                  ""]
    labels = [r"Control$+\Phi$SDM", "Control", "BOOST$+\Phi$SDM", "BOOST"]
    xpos = []
    i=1
    for _ in range(4):
        xpos.append(i)
        i+= 0.3
    figg, axx = plt.subplots(2,4,figsize=(12,6),
                             sharex = True,
                             gridspec_kw = {"hspace" : 0.01,
                                              "wspace" :0.01})
    axx[0][3].spines["top"].set_visible(False)
    axx[0][3].spines["right"].set_visible(False)
    axx[0][3].spines["left"].set_visible(False)
    axx[0][3].spines["bottom"].set_visible(False)
    axx[0][3].set_xticks([])
    axx[0][3].set_yticks([])
    axx[0][3].set_xticklabels([])
    axx[0][3].set_yticklabels([])
    axx[1][3].spines["top"].set_visible(False)
    axx[1][3].spines["right"].set_visible(False)
    axx[1][3].spines["left"].set_visible(False)
    axx[1][3].spines["bottom"].set_visible(False)
    axx[1][3].set_xticks([])
    axx[1][3].set_yticks([])
    axx[1][3].set_xticklabels([])
    axx[1][3].set_yticklabels([])
    for h in range(2):
        for i in range(len(ratios)):
            axx[h][i].set_ylim(-1,4)
            if i == 0:
                axx[h][i].text(2.18, log10(10)+0.05, f"{10**log10(10):.1f}", alpha = 0.5,
                               ha = "right", va = "bottom",
                       fontfamily = "sans-serif", font = "Arial",
                                         fontweight = "bold", fontsize = 10, color = "red")
                axx[h][i].plot([0.5,2.5], [log10(10), log10(10)], linestyle = ":", color = "red")
                axx[h][i].spines["top"].set_visible(False)
                axx[h][i].spines["right"].set_visible(False)
                axx[h][i].set_yticks([0,1,2,3])
                axx[h][i].set_yticklabels(["0", "10","100","1000"],fontfamily = "sans-serif", font = "Arial",
                                         fontweight = "bold", fontsize = 12 )
            elif i == 1:
                axx[h][i].text(2.18, log10(3+1/3)+0.05, f"{10**log10(3+1/3):.1f}", alpha = 0.5,
                               ha = "right", va = "bottom",
                               fontfamily = "sans-serif", font = "Arial",
                               fontweight = "bold", fontsize = 10, color = "red")
                axx[h][i].plot([0.5,2.5], [log10(3 + 1/3), log10(3 + 1/3)], linestyle = ":", color = "red")
                axx[h][i].set_yticks([])
                axx[h][i].set_yticklabels([])
                axx[h][i].spines["top"].set_visible(False)
                axx[h][i].spines["right"].set_visible(False)
                axx[h][i].spines["left"].set_visible(False)
            elif i == 2:
                axx[h][i].plot([0.5,2.5], [log10(3), log10(3)], linestyle = ":", color = "red")
                axx[h][i].set_yticks([])
                axx[h][i].set_yticklabels([])
                axx[h][i].spines["top"].set_visible(False)
                axx[h][i].spines["right"].set_visible(False)
                axx[h][i].spines["left"].set_visible(False)
                axx[h][i].text(2.18, log10(3)+0.05, f"{10**log10(3):.1f}", alpha = 0.5,
                               ha = "right", va = "bottom",
                       fontfamily = "sans-serif", font = "Arial",
                                         fontweight = "bold", fontsize = 10, color = "red")
            violin_parts = axx[h][i].violinplot(loggedmean[h][i], showmedians = True,
                        positions = xpos, widths = 0.2, points = 1000,
                                 showextrema = False)
            axx[h][i].set_xlim(0.7,2.2)
            for j in range(4):
                #axx[h][i].text(xpos[j], 3.5, f"{ratiomeds[h][i][j]:.2f}",
                #                va = "bottom", fontfamily = "sans-serif",
                #                   font = "Arial", fontweight = "bold", ha = "center", 
                #                fontsize = 10)
                violin_parts["bodies"][j].set_facecolor(colours[j])
                violin_parts["bodies"][j].set_edgecolor("black")
            violin_parts["cmedians"].set_color("black")
    #        print([10**item[0][1] for item in violin_parts["cmedians"].get_segments()])
            #violin_parts["cmaxes"].set_color("black")
            #violin_parts["cmins"].set_color("black")
            #violin_parts["cbars"].set_color("black")
        #    violin_parts["cmedians"].set_alpha(0.25)
            #violin_parts["cmaxes"].set_alpha(0.25)
            #violin_parts["cmins"].set_alpha(0.25)
            #violin_parts["cbars"].set_alpha(0.25)
            if h == 1:
                axx[h][i].set_xlabel(ratio_labs[i], fontfamily = "sans-serif",
                                   font = "Arial", fontweight = "bold",
                                fontsize = 14)  
            else:
                axx[h][i].spines["bottom"].set_visible(False)
            axx[h][i].set_xticks([])
            axx[h][i].set_xticklabels([])
    patches = [Patch(color = colours[i], label = labels[i],
                     alpha = 0.5) for i in range(len(labels))]
    axx[1][3].legend(handles=patches, bbox_to_anchor = (1,0.5),
                      prop = {"family" : "Arial",
                              "weight" : "bold",
                              "size" : 12})
    axx[0][3].set_ylim(-1,4)
    #axx[0][3].text(0.75, 3.5, "Median Ratio", ha = "left", va = "bottom",
    #               fontfamily = "sans-serif", font = "Arial", fontweight = "bold",
    #               fontsize = 10)
    axx[0][3].text(0.75, 0.5, "Expected Ratio", ha = "left", va = "bottom",
                   fontfamily = "sans-serif", font = "Arial", fontweight = "bold",
                   fontsize = 10, color = "red", alpha = 0.5)
    figg.text(0.04, 0.5, 'Ratio of Mean Abundances', va='center', rotation='vertical',
                   fontfamily = "sans-serif", font = "Arial", fontweight = "bold",
                   fontsize = 16)
    axx[1][0].set_ylabel("All Possible Ratios",
                   fontfamily = "sans-serif", font = "Arial", fontweight = "bold",
                   fontsize = 12)
    axx[0][0].set_ylabel("No Missing Values",
                   fontfamily = "sans-serif", font = "Arial", fontweight = "bold",
                   fontsize = 12)
    plt.savefig("figures/ratio_suppression.pdf")
    plt.show()
    print("\tGenerating Supporting Figure 1: Localisation Probability Histogram\n")
    fig14, ax14 = plt.subplots(figsize = (12,12))
    for key, value in sty_split:
        class1 = len([row[7] for row in value[1:] if row[7]>0.75])
        if key == "Y":
            ax14.hist([row[7] for row in value[1:] if row[7] ==row[7]], 
                      bins = 50,
                      edgecolor = "black", alpha = 1,
                      label = f"{key:>15}           $n={len(value[1:]):^4}$\t  {class1:>4}(${class1/len(value[1:])*100:>7.1f}\%$)")
        elif key == "S":
            ax14.hist([row[7] for row in value[1:] if row[7] ==row[7]], 
                      bins = 50,
                      edgecolor = "black", alpha = 1,
                      label = f"{key:>15}           $n={len(value[1:]):^4}$       {class1:>4}(${class1/len(value[1:])*100:>7.1f}\%$)")
        else:
            ax14.hist([row[7] for row in value[1:] if row[7] ==row[7]], 
                      bins = 50,
                      edgecolor = "black", alpha = 1,
                      label = f"{key:>15}           $n={len(value[1:]):^4}$\t   {class1:>4}(${class1/len(value[1:])*100:>7.1f}\%$)")
    patch = Patch(color = "white", label = "Phosphosite\tAll sites\tClass I Sites")
    hands, newlabs = ax14.get_legend_handles_labels()
    hands = [patch] + hands
    newlabs = [f"{'Phosphosite':^15} {'All sites':^10}       {'Class I Sites':^15}"] + newlabs
    ax14.set_yticks(ax14.get_yticks()[:-1])
    ax14.set_yticklabels([str(int(item)) for item in ax14.get_yticks()], fontfamily = "sans-serif",
                                    font = "Arial", fontweight = "bold", fontsize = 16)
    ax14.legend(handles = hands, labels = newlabs, prop = {"family" : "Arial",
                                                 "weight" : "bold",
                                                 "size" : 16}, loc = "upper left")
    ax14.plot([0.75,0.75], [0,4000], color = "black", linestyle = ":", alpha = 0.5)
    ax14.set_xticks([0,0.2,0.4,0.6,0.8,1])
    ax14.set_xticklabels(["0","0.2","0.4","0.6","0.8","1"], fontfamily = "sans-serif",
                                    font = "Arial", fontweight = "bold", fontsize = 16)
    ax14.set_ylim(-5,3600)
    ax14.text(0.74, 1800, "Localization probability $= 0.75$", fontfamily = "sans-serif",
                                    font = "Arial", fontsize = 18, rotation = "vertical",
              va = "center", ha = "center")
    ax14.spines["top"].set_visible(False)
    ax14.spines["right"].set_visible(False)
    ax14.set_ylabel("$\#$ of Phosphorylation Sites", fontfamily = "sans-serif",
                    font = "Arial", fontweight = "bold", fontsize = 22)
    ax14.set_xlabel("Localization Probability", fontfamily = "sans-serif",
                    font = "Arial", fontweight = "bold", fontsize = 22)
    plt.savefig("figures/localisation_prob_histogram.pdf")
    plt.show()
    print("\tGenerating Supporting Figure 6: Control$+\Phi$SDM Pairwise Correlation\n")
    fig2, ax2 = plt.subplots(3,3, figsize = (12,12), gridspec_kw = {"hspace" : 0.25,
                                                                    "wspace" : 0.25})
    ax2[0][0] = scatter_gausskde(csten_x_r1,csten_x_r2, ax2[0][0])
    ax2[0][1] = scatter_gausskde(csten_x_r1,csten_x_r3, ax2[0][1])
    ax2[0][2] = scatter_gausskde(csten_x_r2,csten_x_r3, ax2[0][2])
    ax2[0][0].legend(loc="upper left")
    ax2[0][1].legend(loc="upper left")
    ax2[0][2].legend(loc="upper left")
    ax2[1][0] = scatter_gausskde(csthree_x_r1,csthree_x_r2, ax2[1][0])
    ax2[1][1] = scatter_gausskde(csthree_x_r1,csthree_x_r3, ax2[1][1])
    ax2[1][2] = scatter_gausskde(csthree_x_r2,csthree_x_r3, ax2[1][2])
    ax2[1][0].legend(loc="upper left")
    ax2[1][1].legend(loc="upper left")
    ax2[1][2].legend(loc="upper left")
    ax2[2][0] = scatter_gausskde(csone_x_r1,csone_x_r2, ax2[2][0])
    ax2[2][1] = scatter_gausskde(csone_x_r1,csone_x_r3, ax2[2][1])
    ax2[2][2] = scatter_gausskde(csone_x_r2,csone_x_r3, ax2[2][2])
    ax2[2][0].legend(loc="upper left")
    ax2[2][1].legend(loc="upper left")
    ax2[2][2].legend(loc="upper left")
    for i in range(3):
        for j in range(3):
            ax2[i][j].set_ylabel(ylabels[j], **textdict)
            ax2[i][j].set_xlabel(xlabels[j], **textdict)
    for ax, row in zip(ax2[:,0], rows):
        ax.annotate(row, xy=(0, 0.5), xytext=(-ax.yaxis.labelpad - 10, 0),
                    xycoords=ax.yaxis.label, textcoords='offset points',
                    ha='center', va='center', rotation = 90,
                    fontfamily = "sans-serif", font = "Arial", fontsize = 20,
                    fontweight = "bold")
    fig2.suptitle("\n\nControl$+\Phi$SDM", fontfamily = "sans-serif", font = "Arial",
                  fontsize = 24, fontweight = "bold")
    plt.savefig("figures/control_sdm_corrplot.pdf")
    plt.show()
    print("\tGenerating Figure 3C: CV Percentage Boxplot\n")
    fig1, ax1 = plt.subplots(1,4,figsize = (12,6), gridspec_kw = {'wspace' : 0.05})#, sharey=True)
    positions = [1,1.55,2.1,2.65]
    bp1 = ax1[0].boxplot([[item[-7] for item in experiments["Control$+\Phi$SDM"][1:] if item[-7] == item[-7]],
                      [item[-7] for item in experiments["Control"][1:] if item[-7] == item[-7]],
                      [item[-7] for item in experiments["BOOST$+\Phi$SDM"][1:] if item[-7] == item[-7]],
                      [item[-7] for item in experiments["BOOST"][1:] if item[-7] == item[-7]]],
                      notch = True, showcaps = False,
                      patch_artist = True,
                      positions = positions,
                      widths = [0.4,0.4,0.4,0.4],
                      sym = ".")
    bp1_meds = [item.get_ydata()[0] for item in bp1["medians"]]
    for i in range(len(positions)):
        ax1[0].text(positions[i], 105, f"{bp1_meds[i]:.0f}", va = "center", 
                    ha = "center", **{"fontfamily" : "sans-serif",
                                 "font" : "Arial",
                                 "fontsize" : "18"})
    ax1[0].set_xticks([])
    #ax1[0].set_ylim(-5,100)
    ax1[0].set_yticks(ax1[0].get_yticks())
    ax1[0].set_yticklabels([str(item) for item in ax1[0].get_yticks()], **{"fontfamily" : "sans-serif",
                                 "font" : "Arial",
                                 "fontsize" : "18",
                                 "fontweight" : "bold"})
    ax1[0].spines["top"].set_visible(False)
    ax1[0].spines["right"].set_visible(False)
    bp2 = ax1[1].boxplot([[item[-8] for item in experiments["Control$+\Phi$SDM"][1:] if item[-8] == item[-8]],
                      [item[-8] for item in experiments["Control"][1:] if item[-8] == item[-8]],
                      [item[-8] for item in experiments["BOOST$+\Phi$SDM"][1:] if item[-8] == item[-8]],
                      [item[-8] for item in experiments["BOOST"][1:] if item[-8] == item[-8]]],
                      notch = True, showcaps = False,
                      patch_artist = True,
                      positions = positions,
                      widths = [0.4,0.4,0.4,0.4],
                      sym = ".")
    ax1[1].set_yticks([])
    ax1[1].set_xticks([])
    bp2_meds = [item.get_ydata()[0] for item in bp2["medians"]]
    for i in range(len(positions)):
        ax1[1].text(positions[i], 105, f"{bp2_meds[i]:.0f}", va = "center", 
                    ha = "center", **{"fontfamily" : "sans-serif",
                                 "font" : "Arial",
                                 "fontsize" : "18"})
    ax1[1].spines["top"].set_visible(False)
    ax1[1].spines["right"].set_visible(False)
    ax1[1].spines["left"].set_visible(False)
    bp3 = ax1[2].boxplot([[item[-9] for item in experiments["Control$+\Phi$SDM"][1:] if item[-9] == item[-9]],
                      [item[-9] for item in experiments["Control"][1:] if item[-9] == item[-9]],
                      [item[-9] for item in experiments["BOOST$+\Phi$SDM"][1:] if item[-9] == item[-9]],
                      [item[-9] for item in experiments["BOOST"][1:] if item[-9] == item[-9]]],
                      notch = True, showcaps = False,
                      patch_artist = True,
                      positions = positions,
                      widths = [0.4,0.4,0.4,0.4],
                      sym = ".")
    ax1[2].set_yticks([])
    ax1[2].set_xticks([])
    bp3_meds = [item.get_ydata()[0] for item in bp3["medians"]]
    for i in range(len(positions)):
        ax1[2].text(positions[i], 105, f"{bp3_meds[i]:.0f}", va = "center", 
                    ha = "center", **{"fontfamily" : "sans-serif",
                                 "font" : "Arial",
                                 "fontsize" : "18"})
    ax1[2].spines["top"].set_visible(False)
    ax1[2].spines["right"].set_visible(False)
    ax1[2].spines["left"].set_visible(False)
    for i in range(len(bp1["boxes"])):
        bp1["boxes"][i].set(facecolor=colours[i])
        bp1["medians"][i].set(color="black")
        bp1["fliers"][i].set(color=colours[i])
        bp2["boxes"][i].set(facecolor=colours[i])
        bp2["medians"][i].set(color="black")
        bp2["fliers"][i].set(color=colours[i])
        bp3["boxes"][i].set(facecolor=colours[i])
        bp3["medians"][i].set(color="black")
        bp3["fliers"][i].set(color=colours[i])
    ax1[0].set_ylim(-5,100)
    ax1[1].set_ylim(-5,100)
    ax1[2].set_ylim(-5,100)
    ax1[0].set_xlabel("0.1 mg", **{"fontfamily" : "sans-serif",
                                 "font" : "Arial",
                                 "fontsize" : "22",
                                 "fontweight" : "bold"})
    ax1[1].set_xlabel("0.3 mg", **{"fontfamily" : "sans-serif",
                                 "font" : "Arial",
                                 "fontsize" : "22",
                                 "fontweight" : "bold"})
    ax1[2].set_xlabel("1.0 mg", **{"fontfamily" : "sans-serif",
                                 "font" : "Arial",
                                 "fontsize" : "22",
                                 "fontweight" : "bold"})

    ax1[0].set_ylabel("CV (%)", **{"fontfamily" : "sans-serif",
                                 "font" : "Arial",
                                 "fontsize" : "22",
                                 "fontweight" : "bold"})
    ax1[3].spines["top"].set_visible(False)
    ax1[3].spines["right"].set_visible(False)
    ax1[3].spines["left"].set_visible(False)
    ax1[3].spines["bottom"].set_visible(False)
    ax1[3].set_xticks([])
    ax1[3].set_yticks([])
    patches = [Patch(color = colours[i], label = labels[i]) for i in range(len(labels))]
    ax1[3].legend(handles=patches, loc = "center left", prop = {"family" : "Arial",
                                                             "weight" : "bold",
                                                                "size" : 16})
    ax1[3].set_ylim(-5,100)
    ax1[3].text(0.1,105,"Median CV (%)", **{"fontfamily" : "sans-serif",
                                 "font" : "Arial",
                                 "fontsize" : "20",
                                 "fontweight" : "bold",
                                        "ha" : "left",
                                        "va" : "center"})
    plt.subplots_adjust(hspace = 0)
    plt.savefig("figures/cv_plot.pdf")
    plt.show()
    print("\tGenerating Supporting Figure 5: Control Pairwise Correlation\n")
    fig3, ax3 = plt.subplots(3,3,figsize=(12,12), gridspec_kw = {"hspace" : 0.25, 
                                                                 "wspace" : 0.25})
    ax3[0][0] = scatter_gausskde(cten_x_r1,cten_x_r2, ax3[0][0])
    ax3[0][1] = scatter_gausskde(cten_x_r1,cten_x_r3, ax3[0][1])
    ax3[0][2] = scatter_gausskde(cten_x_r2,cten_x_r3, ax3[0][2])
    ax3[0][0].legend(loc="upper left")
    ax3[0][1].legend(loc="upper left")
    ax3[0][2].legend(loc="upper left")
    ax3[1][0] = scatter_gausskde(cthree_x_r1,cthree_x_r2, ax3[1][0])
    ax3[1][1] = scatter_gausskde(cthree_x_r1,cthree_x_r3, ax3[1][1])
    ax3[1][2] = scatter_gausskde(cthree_x_r2,cthree_x_r3, ax3[1][2])
    ax3[1][0].legend(loc="upper left")
    ax3[1][1].legend(loc="upper left")
    ax3[1][2].legend(loc="upper left")
    ax3[2][0] = scatter_gausskde(cone_x_r1,cone_x_r2, ax3[2][0])
    ax3[2][1] = scatter_gausskde(cone_x_r1,cone_x_r3, ax3[2][1])
    ax3[2][2] = scatter_gausskde(cone_x_r2,cone_x_r3, ax3[2][2])
    ax3[2][0].legend(loc="upper left")
    ax3[2][1].legend(loc="upper left")
    ax3[2][2].legend(loc="upper left")
    for i in range(3):
        for j in range(3):
            ax3[i][j].set_ylabel(ylabels[j], **textdict)
            ax3[i][j].set_xlabel(xlabels[j], **textdict)
    for ax, row in zip(ax3[:,0], rows):
        ax.annotate(row, xy=(0, 0.5), xytext=(-ax.yaxis.labelpad - 10, 0),
                    xycoords=ax.yaxis.label, textcoords='offset points',
                    ha='center', va='center', rotation = 90,
                    fontfamily = "sans-serif", font = "Arial", fontsize = 20,
                    fontweight = "bold")
    fig3.suptitle("\n\nControl", fontfamily = "sans-serif", font = "Arial",
                  fontsize = 24, fontweight = "bold")
    plt.savefig("figures/control_corrplot.pdf")
    plt.show()
    print("\tGenerating Supporting Figure 4: BOOST$+\Phi$SDM Pairwise Correlation\n")
    fig4, ax4 = plt.subplots(3,3,figsize=(12,12), gridspec_kw = {"hspace" : 0.25, 
                                                                 "wspace" : 0.25})
    ax4[0][0] = scatter_gausskde(bsten_x_r1,bsten_x_r2, ax4[0][0])
    ax4[0][1] = scatter_gausskde(bsten_x_r1,bsten_x_r3, ax4[0][1])
    ax4[0][2] = scatter_gausskde(bsten_x_r2,bsten_x_r3, ax4[0][2])
    ax4[0][0].legend(loc="upper left")
    ax4[0][1].legend(loc="upper left")
    ax4[0][2].legend(loc="upper left")
    ax4[1][0] = scatter_gausskde(bsthree_x_r1,bsthree_x_r2, ax4[1][0])
    ax4[1][1] = scatter_gausskde(bsthree_x_r1,bsthree_x_r3, ax4[1][1])
    ax4[1][2] = scatter_gausskde(bsthree_x_r2,bsthree_x_r3, ax4[1][2])
    ax4[1][0].legend(loc="upper left")
    ax4[1][1].legend(loc="upper left")
    ax4[1][2].legend(loc="upper left")
    ax4[2][0] = scatter_gausskde(bsone_x_r1,bsone_x_r2, ax4[2][0])
    ax4[2][1] = scatter_gausskde(bsone_x_r1,bsone_x_r3, ax4[2][1])
    ax4[2][2] = scatter_gausskde(bsone_x_r2,bsone_x_r3, ax4[2][2])
    ax4[2][0].legend(loc="upper left")
    ax4[2][1].legend(loc="upper left")
    ax4[2][2].legend(loc="upper left")
    for i in range(3):
        for j in range(3):
            ax4[i][j].set_ylabel(ylabels[j], **textdict)
            ax4[i][j].set_xlabel(xlabels[j], **textdict)
    for ax, row in zip(ax4[:,0], rows):
        ax.annotate(row, xy=(0, 0.5), xytext=(-ax.yaxis.labelpad - 10, 0),
                    xycoords=ax.yaxis.label, textcoords='offset points',
                    ha='center', va='center', rotation = 90,
                    fontfamily = "sans-serif",font = "Arial", fontsize = 20,
                    fontweight = "bold")
    fig4.suptitle("\n\nBOOST$+\Phi$SDM", fontfamily = "sans-serif", font = "Arial",
                  fontsize = 24, fontweight = "bold")
    plt.savefig("figures/boost_sdm_corrplot.pdf")
    plt.show()
    print("\tGenerating Supporting Figure 3: BOOST Pairwise Correlation\n")
    fig5, ax5 = plt.subplots(3,3,figsize=(12,12), gridspec_kw = {"hspace" : 0.25, 
                                                                 "wspace" : 0.25})
    ax5[0][0] = scatter_gausskde(bten_x_r1,bten_x_r2, ax5[0][0])
    ax5[0][1] = scatter_gausskde(bten_x_r1,bten_x_r3, ax5[0][1])
    ax5[0][2] = scatter_gausskde(bten_x_r2,bten_x_r3, ax5[0][2])
    ax5[0][0].legend(loc="upper left")
    ax5[0][1].legend(loc="upper left")
    ax5[0][2].legend(loc="upper left")
    ax5[1][0] = scatter_gausskde(bthree_x_r1,bthree_x_r2, ax5[1][0])
    ax5[1][1] = scatter_gausskde(bthree_x_r1,bthree_x_r3, ax5[1][1])
    ax5[1][2] = scatter_gausskde(bthree_x_r2,bthree_x_r3, ax5[1][2])
    ax5[1][0].legend(loc="upper left")
    ax5[1][1].legend(loc="upper left")
    ax5[1][2].legend(loc="upper left")
    ax5[2][0] = scatter_gausskde(bone_x_r1,bone_x_r2, ax5[2][0])
    ax5[2][1] = scatter_gausskde(bone_x_r1,bone_x_r3, ax5[2][1])
    ax5[2][2] = scatter_gausskde(bone_x_r2,bone_x_r3, ax5[2][2])
    ax5[2][0].legend(loc="upper left")
    ax5[2][1].legend(loc="upper left")
    ax5[2][2].legend(loc="upper left")
    for i in range(3):
        for j in range(3):
            ax5[i][j].set_ylabel(ylabels[j], **textdict)
            ax5[i][j].set_xlabel(xlabels[j], **textdict)
    for ax, row in zip(ax5[:,0], rows):
        ax.annotate(row, xy=(0, 0.5), xytext=(-ax.yaxis.labelpad - 10, 0),
                    xycoords=ax.yaxis.label, textcoords='offset points',
                    ha='center', va='center', rotation = 90,
                    fontfamily = "sans-serif", font = "Arial", fontsize = 20,
                    fontweight = "bold")
    fig5.suptitle("\n\nBOOST", fontfamily = "sans-serif", font = "Arial",
                  fontsize = 24, fontweight = "bold")
    plt.savefig("figures/boost_corrplot.pdf")
    plt.show()
    print("\tGenerating Supporting Figure 2: Reporter Intensity Boxplots\n")
    n = 2 # of double rows
    m = 2 # of cols (note that the second column is purely for labels)
    t = 0.9 # Top space
    b = 0.1 # bottom space
    msp = 0.1 # Minor space
    sp = 0.5  # Major space
    offs=(1+msp)*(t-b)/(2*n+n*msp+(n-1)*sp) # grid offset
    hspace = sp+msp+1 #height space per grid
    # Make the GridSpec objects for the sets of subplots.
    gso = GridSpec(n,m, bottom=b+offs, top=t, hspace=hspace, wspace = 0.01)
    gse = GridSpec(n,m, bottom=b, top=t-offs, hspace=hspace, wspace = 0.01)
    # Make figure 9 and a list to hold the axes
    fig9 = plt.figure(figsize = (24,16))
    ax9 = []
    # Add the axes to the list. Because of how this is set up, the axes
    # have a strange indexing:
    for i in range(n*m):
        ax9.append([fig9.add_subplot(gso[i]),fig9.add_subplot(gse[i])])
    # 0,0 is the upper left
    # 0,1 is right below 0,0
    # 1,0 is the upper right
    # 1,1 is right below 1,0
    # 2,0 is the middle left
    # 2,1 is right below 2,0
    # 3,0 is the middle right
    # 3,1 is right below 3,0
    # Because I want all of the intensity boxplots to be in the left column, I need to
    # plot to 0,0 (Control + SDM), 0,1 (Control), 2,0 (BOOST + SDM) and 2,1 (BOOST)
    # Make the x positions for the boxplots using the same spacing as the CV plot
    i = 1
    x_pos = []
    for j in range(11):
        x_pos.append(i)
        i+=0.85
    # Loop over the number of rows
    for i in range(4):
        # If i is 0 or 2, I need to plot Control + SDM (sigsnals[0]) or BOOST + SDM (sigsnals[2])
        if i == 0 or i == 2:
            ax9[i][0].boxplot(sigsnals[i], notch = False, showcaps = False,
                              patch_artist = True,
                              positions = x_pos,
                              widths = [0.4 for _ in range(11)],
                              sym = ".",
                              boxprops=dict(facecolor=colours[i]),
                              medianprops=dict(color="black"))
            for j in range(11):
                ax9[i][0].text(x_pos[j], 8.5, f"{meds_list[i][j]:.0f}", fontfamily = "sans-serif",
                               font = "Arial", fontweight = "bold", ha = "center", va = "center",
                               fontsize = 16)
            ax9[i][0].set_ylim(0,9)
            ax9[i][0].set_yticks([2,4,6,8])
            ax9[i][0].set_yticklabels(["2","4","6","8"], fontfamily = "sans-serif", font = "Arial",
                                      fontsize = 18, fontweight = "bold")
            ax9[i][0].spines["right"].set_visible(False)
            ax9[i][0].spines["top"].set_visible(False)
            if i != 3 and i != 1:
                ax9[i][0].spines["bottom"].set_visible(False)
                ax9[i][0].set_xticks([])
            #
            ax9[i+1][1].spines["top"].set_visible(False)
            ax9[i+1][1].spines["right"].set_visible(False)
            ax9[i+1][1].spines["left"].set_visible(False)
            ax9[i+1][1].spines["bottom"].set_visible(False)
            ax9[i+1][1].set_xticks([])
            ax9[i+1][1].set_yticks([])
        # If i is 1, then we need to plot sigsnals[i] (Control) to 0,1
        elif i == 1:
            ax9[0][1].boxplot(sigsnals[i], notch = False, showcaps = False,
                              patch_artist = True,
                              positions = x_pos,
                              widths = [0.4 for _ in range(11)],
                              sym = ".",# label = labels[i],
                              boxprops=dict(facecolor=colours[i]),
                              medianprops=dict(color="black"))
            for j in range(11):
                ax9[0][1].text(x_pos[j], 8.5, f"{meds_list[i][j]:.0f}", fontfamily = "sans-serif",
                               font = "Arial", fontweight = "bold", ha = "center", va = "center",
                               fontsize = 16)
            ax9[0][1].set_ylim(0,9)
            ax9[0][1].set_yticks([2,4,6,8])
            ax9[0][1].set_yticklabels(["2","4","6","8"], fontfamily = "sans-serif", font = "Arial",
                                      fontsize = 18, fontweight = "bold")
            ax9[0][1].spines["right"].set_visible(False)
            ax9[0][1].spines["top"].set_visible(False)
            ax9[0][1].set_xticks(x_pos)
            ax9[0][1].set_xticklabels(control_labs, rotation = 90, fontfamily = "sans-serif", font = "Arial",
                                     fontweight = "bold", fontsize = 16)
            #
            ax9[i][0].spines["top"].set_visible(False)
            ax9[i][0].spines["right"].set_visible(False)
            ax9[i][0].spines["left"].set_visible(False)
            ax9[i][0].spines["bottom"].set_visible(False)
            ax9[i][0].set_ylim(0,9)
            ax9[i][0].text(0, 8.5, "Median Intensity", fontfamily = "sans-serif", font = "Arial",
                           fontsize = 16, fontweight = "bold", ha = "left", va = "center")
            ax9[i][0].set_xticks([])
            ax9[i][0].set_yticks([])
        # If i is 3, then we need to plot sigsnals[i] (BOOST) to 0,1
        elif i == 3:
            ax9[2][1].boxplot(sigsnals[i], notch = False, showcaps = False,
                                  patch_artist = True,
                                  positions = x_pos,
                                  widths = [0.4 for _ in range(11)],
                                  sym = ".",# label = labels[i],
                                  boxprops=dict(facecolor=colours[i]),
                                  medianprops=dict(color="black"))
            for j in range(11):
                ax9[2][1].text(x_pos[j], 8.5, f"{meds_list[i][j]:.0f}", fontfamily = "sans-serif",
                                   font = "Arial", fontweight = "bold", ha = "center", 
                               va = "center", fontsize = 16)
            ax9[2][1].set_ylim(0,9)
            ax9[2][1].set_yticks([2,4,6,8])
            ax9[2][1].set_yticklabels(["2","4","6","8"], fontfamily = "sans-serif", font = "Arial",
                                         fontsize = 18, fontweight = "bold")
            ax9[2][1].spines["right"].set_visible(False)
            ax9[2][1].spines["top"].set_visible(False)
            ax9[2][1].set_xticks(x_pos)
            ax9[2][1].set_xticklabels(boost_labs, rotation = 90, fontfamily = "sans-serif", font = "Arial",
                                         fontweight = "bold", fontsize = 16)
            ax9[i][0].spines["top"].set_visible(False)
            ax9[i][0].spines["right"].set_visible(False)
            ax9[i][0].spines["left"].set_visible(False)
            ax9[i][0].spines["bottom"].set_visible(False)
            ax9[i][0].set_xticks([])
            ax9[i][0].set_yticks([])
    fig9.text(0.04, 0.25, '\n\n\n$\log_{10}$(Reporter Intensity)', 
              va='center', rotation='vertical', fontfamily = "sans-serif",
              font = "Arial", fontsize = 20)
    fig9.text(0.04, 0.75, '\n\n\n$\log_{10}$(Reporter Intensity)', 
              va='center', rotation='vertical', fontfamily = "sans-serif",
              font = "Arial", fontsize = 20)
    # Add the legend stuffs. The patches were already made in the CV plot, so just add them
    ax9[1][1].legend(handles=patches, bbox_to_anchor = (0,0),
                     loc = "upper left", prop = {"family" : "Arial",
                                                 "weight" : "bold",
                                                 "size" : 16})
    plt.savefig("figures/report_intensity_boxplot.pdf")
    plt.show()
    print("\tGenerating Supporting Figure 10: BOOST Factor CDFs\n")
    fig12, ax12 = plt.subplots(2,2, figsize = (12,12),
                               gridspec_kw = {"wspace" : 0,
                                              "hspace" : 0.05})
    for i in range(2):
        for j in range(2):
            k = 0
            for cdf_d in cdf_data[i][j]:
                cdf([item for item in cdf_d if item == item], 50,
                        ax12[i][j], cdf_colours[k],
                        label = f"{cdf_labs[k]}$n={len([item for item in cdf_d if item == item])}$",
                    plot_90 = True)
                k+=1
            ax12[i][j].set_xlim(0.1,10000)
            ax12[i][j].set_ylim(-0.1,1.1)
            ax12[i][j].legend(loc = "center right", prop = {"family" : "Arial",
                                                             "weight" : "bold",
                                                                "size" : 16})
            ax12[i][j].spines["top"].set_visible(False)
            ax12[i][j].spines["right"].set_visible(False)
            if i == 0:
                #ax12[i][j].spines["bottom"].set_visible(False)
                ax12[i][j].set_xticks([1,10,100,1000,10000])
                ax12[i][j].tick_params(axis = "x", which = "minor", length = 6, width = 2)
                ax12[i][j].tick_params(axis = "x", which = "major", length = 12, width = 2)
                ax12[i][j].set_xticklabels(["","","","", ""], fontfamily = "sans-serif", font = "Arial", fontweight = "bold",
                            fontsize = 18, ha = "right", va = "top", rotation=45)
                ax12[i][j].plot([5,5],[-0.9,.9], color = "black",
                            alpha = 0.5, linestyle = ":")
                ax12[i][j].text(5.5,0.1,r"$5\times$",
                                rotation= 270, fontfamily = "sans-serif",
                                font = "Arial", fontsize = 16)
                ax12[i][j].plot([10,10],[-0.9,.9], color = "black",
                                alpha = 0.5, linestyle = ":")
                ax12[i][j].text(10.8,0.1,r"$10\times$",
                                rotation = 270, fontfamily = "sans-serif",
                                font = "Arial", fontsize = 16)
                ax12[i][j].plot([20,20],[-0.9,.9], color = "black",
                                alpha = 0.5, linestyle = ":")
                ax12[i][j].text(21,0.1,r"$20\times$",
                                rotation = 270, fontfamily = "sans-serif",
                                font = "Arial", fontsize = 16)
                #ax12[i][j].plot([0.1,10000], [-0.09,-0.09], color = "black", linestyle = "-")
            else:
                ax12[i][j].set_xticks([1,10,100,1000,10000])
                ax12[i][j].tick_params(axis = "x", which = "minor", length = 6, width = 2)
                ax12[i][j].tick_params(axis = "x", which = "major", length = 12, width = 2)
                ax12[i][j].set_xticklabels(["1","10","100","1000", "10000"], fontfamily = "sans-serif", font = "Arial", fontweight = "bold",
                            fontsize = 18, ha = "right", va = "top", rotation=45)
                ax12[i][j].plot([20,20],[-0.9,.9], color = "black",
                                alpha = 0.5, linestyle = ":")
                ax12[i][j].text(21,0.1,r"$20\times$",
                                rotation = 270, fontfamily = "sans-serif",
                                font = "Arial", fontsize = 16)
                ax12[i][j].plot([40,40],[-0.9,.9], color = "black",
                                alpha = 0.5, linestyle = ":")
                ax12[i][j].text(41,0.1,r"$40\times$",
                                rotation = 270, fontfamily = "sans-serif",
                                font = "Arial", fontsize = 16)
                ax12[i][j].plot([80,80],[-0.9,.9], color = "black",
                                alpha = 0.5, linestyle = ":")
                ax12[i][j].text(82,0.1,r"$80\times$",
                                rotation = 270, fontfamily = "sans-serif",
                                font = "Arial", fontsize = 16)
            if j == 1:
                ax12[i][j].spines["left"].set_visible(False)
                ax12[i][j].set_yticks([])
            else:
                ax12[i][j].text(0.2,0.92, "$90\%$", fontfamily = "sans-serif",
                                font = "Arial", fontsize = 16)
                ax12[i][j].set_yticks([0,0.2,0.4,0.6,0.8,1])
                ax12[i][j].set_yticklabels(["0","0.2","0.4","0.6","0.8","1"],
                           fontfamily = "sans-serif", font = "Arial", fontweight = "bold",
                            fontsize = 18, ha = "right", va = "center")
            ax12[i][j].plot([0,10000],[0.9,0.9], color = "black", linestyle = ":",
                            alpha = 0.5)
    ax12[0][0].set_title("BOOST", fontfamily = "sans-serif", font = "Arial",
                         fontweight = "bold", fontsize = 20,bbox=dict(facecolor='none', edgecolor='black'))
    ax12[0][1].set_title("BOOST$+\Phi$SDM", fontfamily = "sans-serif", font = "Arial",
                         fontweight = "bold", fontsize = 20, bbox=dict(facecolor='none', edgecolor='black'))
    ax12[0][1].yaxis.set_label_position("right")
    ax12[0][1].set_ylabel("pTyr peptides with q$<$0.05", fontfamily = "sans-serif", font = "Arial",
                         fontweight = "bold", fontsize = 20, bbox=dict(facecolor='none', edgecolor='black'),
                          va = "bottom", labelpad = 6, rotation = 270)
    ax12[1][1].yaxis.set_label_position("right")
    ax12[1][1].set_ylabel("All possible ratios", fontfamily = "sans-serif", font = "Arial",
                         fontweight = "bold", fontsize = 20, bbox=dict(facecolor='none', edgecolor='black'),
                          va = "bottom", labelpad = 6, rotation = 270)
    fig12.text(0, 0.5, '\n\nProportion of Unique pTyr peptides', 
              va='center', rotation='vertical', fontfamily = "sans-serif",
              font = "Arial", fontweight = "bold", fontsize = 22)
    fig12.text(0.45, 0.02, 'BOOST Factor', 
              va='center', fontfamily = "sans-serif",
              font = "Arial", fontweight = "bold", fontsize = 22)
    plt.savefig("figures/boostfactor_cdf.pdf")
    plt.show()
    print("\tGenerating Supporting Figure 8: BOOST-Control Overlap Volcanoes\n")
    fig15, ax15 = plt.subplots(3,4,figsize = (12,12), gridspec_kw = {"hspace" : 0.001,
                                                                     "wspace" : 0.001})
    # Four columns, so this will be second index
    for i in range(3):
        # Three rows, so this will be first index
        for j in range(4):
            ax15[i][j].scatter(boost_control_rs[j][i], 
                               boost_control_qs[j][i],
                               color = overlap_bc_colours[j],
                               s=20, edgecolor = "black", linewidth = 0.2)
            ax15[i][j].plot([-1.5,2],[-log10(0.05),-log10(0.05)], 
                           color = "black", linestyle = ":")
            ax15[i][j].set_xlim(-1.5,2)
            ax15[i][j].set_ylim(-0.5,4)
            ax15[i][j].spines["top"].set_visible(False)
            #ax15[i][j].spines["right"].set_visible(False)
            if i != 2:
                ax15[i][j].spines["bottom"].set_visible(False)
                ax15[i][j].set_xticks([])
            else:
                ax15[i][j].set_xticks([-1,0,1])
                ax15[i][j].set_xticklabels(["-1","0","1"],fontfamily = "sans-serif",
                                         font = "Arial", fontweight = "bold", fontsize = 18)
            if j != 0:
                #ax15[i][j].spines["left"].set_visible(False)
                ax15[i][j].set_yticks([])
            else:
                ax15[i][j].set_yticks([0,1,2,3,4])
                ax15[i][j].set_yticklabels(["0","1","2","3","4"], fontfamily = "sans-serif",
                                         font = "Arial", fontweight = "bold", fontsize = 18)
            
            if i == 0:
                ax15[i][j].spines["top"].set_visible(True)
                ax15[i][j].plot([log10(10),log10(10)],[-0.5,4], 
                               color = "black", linestyle = "dashed", alpha = 0.5)
                ax15[i][j].plot([log10(20),log10(20)],[-0.5,4], color = "grey",alpha = 0.1)
                ax15[i][j].plot([log10(5),log10(5)],[-0.5,4], color = "grey", alpha = 0.1)
                ax15[i][j].fill_between([log10(5),log10(20)],-0.5,4,color="grey",alpha=0.1)
                ax15[i][j].text(0,2.5,
                               f"{boost_control_qcounts[j][i]}\n$({(boost_control_qcounts[j][i]/len(boost_control_qs[j][i])*100):.0f}\%)$",
                               fontfamily = "sans-serif", font = "Arial", fontweight = "bold",
                               va = "center", ha = "right", fontsize = 16)
                ax15[i][j].text(0,1,
                               f"{len(boost_control_qs[j][i])-boost_control_qcounts[j][i]}\n$({((len(boost_control_qs[j][i])-boost_control_qcounts[j][i])/len(boost_control_qs[j][i])*100):.0f}\%)$",
                               fontfamily = "sans-serif", font = "Arial", fontweight = "bold",
                               va = "top", ha = "right", fontsize = 16)
            elif i == 1:
                ax15[i][j].plot([log10(3 + 1/3),log10(3 + 1/3)],[-0.5,4], 
                               color = "black", linestyle = "dashed", alpha = 0.5)
                ax15[i][j].plot([log10((3 + 1/3)/2),log10((3 + 1/3)/2)],[-0.5,4], color = "grey",alpha = 0.1)
                ax15[i][j].plot([log10((3 + 1/3)*2), log10((3 + 1/3)*2)],[-0.5,4], color = "grey", alpha = 0.1)
                ax15[i][j].fill_between([log10((3 + 1/3)/2),log10((3 + 1/3)*2)],-0.5,4,color="grey",alpha=0.1)
                ax15[i][j].text(0,2.5,
                               f"{boost_control_qcounts[j][i]}\n$({(boost_control_qcounts[j][i]/len(boost_control_qs[j][i])*100):.0f}\%)$",
                               fontfamily = "sans-serif", font = "Arial", fontweight = "bold",
                               va = "center", ha = "right", fontsize = 16)
                ax15[i][j].text(0,1,
                               f"{len(boost_control_qs[j][i])-boost_control_qcounts[j][i]}\n$({((len(boost_control_qs[j][i])-boost_control_qcounts[j][i])/len(boost_control_qs[j][i])*100):.0f}\%)$",
                               fontfamily = "sans-serif", font = "Arial", fontweight = "bold",
                               va = "top", ha = "right", fontsize = 16)
            else:
                ax15[i][j].plot([log10(3),log10(3)],[-0.5,4], 
                               color = "black", linestyle = "dashed", alpha = 0.5)
                ax15[i][j].plot([log10(3/2),log10(3/2)],[-0.5,4], color = "grey",alpha = 0.1)
                ax15[i][j].plot([log10(3*2), log10(3*2)],[-0.5,4], color = "grey", alpha = 0.1)
                ax15[i][j].fill_between([log10(3/2),log10(3*2)],-0.5,4,color="grey",alpha=0.1)
                ax15[i][j].text(0,2.5,
                               f"{boost_control_qcounts[j][i]}\n$({(boost_control_qcounts[j][i]/len(boost_control_qs[j][i])*100):.0f}\%)$",
                               fontfamily = "sans-serif", font = "Arial", fontweight = "bold",
                               va = "center", ha = "right", fontsize = 16)
                ax15[i][j].text(0,1,
                               f"{len(boost_control_qs[j][i])-boost_control_qcounts[j][i]}\n$({((len(boost_control_qs[j][i])-boost_control_qcounts[j][i])/len(boost_control_qs[j][i])*100):.0f}\%)$",
                               fontfamily = "sans-serif", font = "Arial", fontweight = "bold",
                               va = "top", ha = "right", fontsize = 16)
    ax15[0][0].set_title("BOOST\nGained", fontfamily = "sans-serif", font = "Arial",
                         fontweight = "bold", fontsize = 20, 
                         bbox=dict(facecolor='none', edgecolor='black'),
                         pad = 10)
    ax15[0][1].set_title("Overlap\n(BOOST)", fontfamily = "sans-serif", font = "Arial",
                         fontweight = "bold", fontsize = 20, 
                         bbox=dict(facecolor='none', edgecolor='black'),
                         pad = 10)
    ax15[0][2].set_title("Overlap\n(Control)", fontfamily = "sans-serif", font = "Arial",
                         fontweight = "bold", fontsize = 20, 
                         bbox=dict(facecolor='none', edgecolor='black'),
                         pad = 10)
    ax15[0][3].set_title("Control\nOnly", fontfamily = "sans-serif", font = "Arial",
                         fontweight = "bold", fontsize = 20, 
                         bbox=dict(facecolor='none', edgecolor='black'),
                         pad = 10)
    ax15[0][3].yaxis.set_label_position("right")
    ax15[0][3].set_ylabel("1.0 mg : 0.1 mg", fontfamily = "sans-serif", font = "Arial",
                         fontweight = "bold", fontsize = 22, bbox=dict(facecolor='none', edgecolor='black'),
                          va = "bottom", labelpad = 6, rotation = 270)
    ax15[1][3].yaxis.set_label_position("right")
    ax15[1][3].set_ylabel("1.0 mg : 0.3 mg", fontfamily = "sans-serif", font = "Arial",
                         fontweight = "bold", fontsize = 22, bbox=dict(facecolor='none', edgecolor='black'),
                          va = "bottom", labelpad = 6, rotation = 270)
    ax15[2][3].yaxis.set_label_position("right")
    ax15[2][3].set_ylabel("0.3 mg : 0.1 mg", fontfamily = "sans-serif", font = "Arial",
                         fontweight = "bold", fontsize = 22, bbox=dict(facecolor='none', edgecolor='black'),
                          va = "bottom", labelpad = 6, rotation = 270)
    fig15.text(0, 0.5, '\n\n\n$-\log_{10}(q)$', 
              va='center', rotation='vertical', fontfamily = "sans-serif",
              font = "Arial", fontweight = "bold", fontsize = 20)
    fig15.text(0.45, 0.07, '$\log_{10}$(Intensity Ratio)', 
              va='center', fontfamily = "sans-serif",
              font = "Arial", fontsize = 20)
    plt.savefig("figures/overlap_q_volcano_nosdm.pdf")
    plt.show()
    print("\tGenerating Supporting Figure 9: BOOST$+\Phi$SDM-Control$+\Phi$SDM Overlap Volcanoes\n")
    fig16, ax16 = plt.subplots(3,4,figsize = (12,12), gridspec_kw = {"hspace" : 0.001,
                                                                     "wspace" : 0.001})
    # Four columns, so this will be second index
    for i in range(3):
        # Three rows, so this will be first index
        for j in range(4):
            ax16[i][j].scatter(boostsdm_controlsdm_rs[j][i], 
                               boostsdm_controlsdm_qs[j][i],
                               color = overlap_bscs_colours[j],
                               s=20, edgecolor = "black", linewidth = 0.2)
            ax16[i][j].plot([-1.5,2],[-log10(0.05),-log10(0.05)], 
                           color = "black", linestyle = ":")
            ax16[i][j].set_xlim(-1.5,2)
            ax16[i][j].set_ylim(-0.5,4)
            ax16[i][j].spines["top"].set_visible(False)
            #ax15[i][j].spines["right"].set_visible(False)
            if i != 2:
                ax16[i][j].spines["bottom"].set_visible(False)
                ax16[i][j].set_xticks([])
            else:
                ax16[i][j].set_xticks([-1,0,1])
                ax16[i][j].set_xticklabels(["-1","0","1"],fontfamily = "sans-serif",
                                         font = "Arial", fontweight = "bold", fontsize = 18)
            if j != 0:
                #ax15[i][j].spines["left"].set_visible(False)
                ax16[i][j].set_yticks([])
            else:
                ax16[i][j].set_yticks([0,1,2,3,4])
                ax16[i][j].set_yticklabels(["0","1","2","3","4"], fontfamily = "sans-serif",
                                         font = "Arial", fontweight = "bold", fontsize = 18)
            
            if i == 0:
                ax16[i][j].spines["top"].set_visible(True)
                ax16[i][j].plot([log10(10),log10(10)],[-0.5,4], 
                               color = "black", linestyle = "dashed", alpha = 0.5)
                ax16[i][j].plot([log10(20),log10(20)],[-0.5,4], color = "grey",alpha = 0.1)
                ax16[i][j].plot([log10(5),log10(5)],[-0.5,4], color = "grey", alpha = 0.1)
                ax16[i][j].fill_between([log10(5),log10(20)],-0.5,4,color="grey",alpha=0.1)
                ax16[i][j].text(0,2.5,
                               f"{boostsdm_controlsdm_qcounts[j][i]}\n$({(boostsdm_controlsdm_qcounts[j][i]/len(boostsdm_controlsdm_qs[j][i])*100):.0f}\%)$",
                               fontfamily = "sans-serif", font = "Arial", fontweight = "bold",
                               va = "center", ha = "right", fontsize = 16)
                ax16[i][j].text(0,1,
                               f"{len(boostsdm_controlsdm_qs[j][i])-boostsdm_controlsdm_qcounts[j][i]}\n$({((len(boostsdm_controlsdm_qs[j][i])-boostsdm_controlsdm_qcounts[j][i])/len(boostsdm_controlsdm_qs[j][i])*100):.0f}\%)$",
                               fontfamily = "sans-serif", font = "Arial", fontweight = "bold",
                               va = "top", ha = "right", fontsize = 16)
            elif i == 1:
                ax16[i][j].plot([log10(3 + 1/3),log10(3 + 1/3)],[-0.5,4], 
                               color = "black", linestyle = "dashed", alpha = 0.5)
                ax16[i][j].plot([log10((3 + 1/3)/2),log10((3 + 1/3)/2)],[-0.5,4], color = "grey",alpha = 0.1)
                ax16[i][j].plot([log10((3 + 1/3)*2), log10((3 + 1/3)*2)],[-0.5,4], color = "grey", alpha = 0.1)
                ax16[i][j].fill_between([log10((3 + 1/3)/2),log10((3 + 1/3)*2)],-0.5,4,color="grey",alpha=0.1)
                ax16[i][j].text(0,2.5,
                               f"{boostsdm_controlsdm_qcounts[j][i]}\n$({(boostsdm_controlsdm_qcounts[j][i]/len(boostsdm_controlsdm_qs[j][i])*100):.0f}\%)$",
                               fontfamily = "sans-serif", font = "Arial", fontweight = "bold",
                               va = "center", ha = "right", fontsize = 16)
                ax16[i][j].text(0,1,
                               f"{len(boostsdm_controlsdm_qs[j][i])-boostsdm_controlsdm_qcounts[j][i]}\n$({((len(boostsdm_controlsdm_qs[j][i])-boostsdm_controlsdm_qcounts[j][i])/len(boostsdm_controlsdm_qs[j][i])*100):.0f}\%)$",
                               fontfamily = "sans-serif", font = "Arial", fontweight = "bold",
                               va = "top", ha = "right", fontsize = 16)
            else:
                ax16[i][j].plot([log10(3),log10(3)],[-0.5,4], 
                               color = "black", linestyle = "dashed", alpha = 0.5)
                ax16[i][j].plot([log10(3/2),log10(3/2)],[-0.5,4], color = "grey",alpha = 0.1)
                ax16[i][j].plot([log10(3*2), log10(3*2)],[-0.5,4], color = "grey", alpha = 0.1)
                ax16[i][j].fill_between([log10(3/2),log10(3*2)],-0.5,4,color="grey",alpha=0.1)
                ax16[i][j].text(0,2.5,
                               f"{boostsdm_controlsdm_qcounts[j][i]}\n$({(boostsdm_controlsdm_qcounts[j][i]/len(boostsdm_controlsdm_qs[j][i])*100):.0f}\%)$",
                               fontfamily = "sans-serif", font = "Arial", fontweight = "bold",
                               va = "center", ha = "right", fontsize = 16)
                ax16[i][j].text(0,1,
                               f"{len(boostsdm_controlsdm_qs[j][i])-boostsdm_controlsdm_qcounts[j][i]}\n$({((len(boostsdm_controlsdm_qs[j][i])-boostsdm_controlsdm_qcounts[j][i])/len(boostsdm_controlsdm_qs[j][i])*100):.0f}\%)$",
                               fontfamily = "sans-serif", font = "Arial", fontweight = "bold",
                               va = "top", ha = "right", fontsize = 16)
    ax16[0][0].set_title("BOOST$+\Phi$SDM\nGained", fontfamily = "sans-serif", font = "Arial",
                         fontweight = "bold", fontsize = 20, 
                         bbox=dict(facecolor='none', edgecolor='black'),
                         pad = 10)
    ax16[0][1].set_title("Overlap\n(BOOST$+\Phi$SDM)", fontfamily = "sans-serif", font = "Arial",
                         fontweight = "bold", fontsize = 20, 
                         bbox=dict(facecolor='none', edgecolor='black'),
                         pad = 10)
    ax16[0][2].set_title("Overlap\n(Control$+\Phi$SDM)", fontfamily = "sans-serif", font = "Arial",
                         fontweight = "bold", fontsize = 20, 
                         bbox=dict(facecolor='none', edgecolor='black'),
                         pad = 10)
    ax16[0][3].set_title("Control$+\Phi$SDM\nOnly", fontfamily = "sans-serif", font = "Arial",
                         fontweight = "bold", fontsize = 20, 
                         bbox=dict(facecolor='none', edgecolor='black'),
                         pad = 10)
    ax16[0][3].yaxis.set_label_position("right")
    ax16[0][3].set_ylabel("1.0 mg : 0.1 mg", fontfamily = "sans-serif", font = "Arial",
                         fontweight = "bold", fontsize = 22, bbox=dict(facecolor='none', edgecolor='black'),
                          va = "bottom", labelpad = 6, rotation = 270)
    ax16[1][3].yaxis.set_label_position("right")
    ax16[1][3].set_ylabel("1.0 mg : 0.3 mg", fontfamily = "sans-serif", font = "Arial",
                         fontweight = "bold", fontsize = 22, bbox=dict(facecolor='none', edgecolor='black'),
                          va = "bottom", labelpad = 6, rotation = 270)
    ax16[2][3].yaxis.set_label_position("right")
    ax16[2][3].set_ylabel("0.3 mg : 0.1 mg", fontfamily = "sans-serif", font = "Arial",
                         fontweight = "bold", fontsize = 22, bbox=dict(facecolor='none', edgecolor='black'),
                          va = "bottom", labelpad = 6, rotation = 270)
    fig16.text(0, 0.5, '\n\n\n$-\log_{10}(q)$', 
              va='center', rotation='vertical', fontfamily = "sans-serif",
              font = "Arial", fontweight = "bold", fontsize = 20)
    fig16.text(0.45, 0.07, '$\log_{10}$(Intensity Ratio)', 
              va='center', fontfamily = "sans-serif",
              font = "Arial", fontsize = 20)
    plt.savefig("figures/overlap_q_volcano_sdm.pdf")
    plt.show()
    print("\tGenerating Supporting Figures 8 & 9, Figure 4: Set Overlap\n")
    fig10, ax10 = plt.subplots(2,2,figsize = (12,12))
    boost_v_control = venn2([set(boost_peps_f), set(control_peps_f)], 
                              set_labels = ["",""],
                              ax = ax10[0][0], set_colors = [colours[3], colours[1]])
    ax10[0][0].set_title("Overlap of BOOST and Control\npTyr Peptides",
                         fontfamily = "sans-serif", font = "Arial",
                         fontweight = "bold", fontsize = 16)
    for text in boost_v_control.subset_labels:
        text.set_fontfamily("sans-serif")
        text.set_font("Arial")
        text.set_fontweight("bold")
        text.set_fontsize(12)
    bsdm_v_csdm = venn2([set(boostsdm_peps_f), set(controlsdm_peps_f)], 
                          set_labels = ["",""],
                          ax = ax10[0][1], set_colors = [colours[2], colours[0]])
    ax10[0][1].set_title("Overlap of BOOST$+\Phi$SDM and Control$+\Phi$SDM\npTyr Peptides",
                         fontfamily = "sans-serif", font = "Arial",
                         fontweight = "bold", fontsize = 16)
    for text in bsdm_v_csdm.subset_labels:
        text.set_fontfamily("sans-serif")
        text.set_font("Arial")
        text.set_fontweight("bold")
        text.set_fontsize(12)
    boost_v_bsdm = venn2([set(boost_peps_f), set(boostsdm_peps_f)], 
                          set_labels = ["",""],
                          ax = ax10[1][0], set_colors = [colours[3], colours[2]])
    ax10[1][0].set_title("Overlap of BOOST and BOOST$+\Phi$SDM\npTyr Peptides",
                         fontfamily = "sans-serif", font = "Arial",
                         fontweight = "bold", fontsize = 16)
    for text in boost_v_bsdm.subset_labels:
        text.set_fontfamily("sans-serif")
        text.set_font("Arial")
        text.set_fontweight("bold")
        text.set_fontsize(12)
    control_v_csdm = venn2([set(control_peps_f), set(controlsdm_peps_f)], 
                      set_labels = ["",""],
                      ax = ax10[1][1], set_colors = [colours[1], colours[0]])
    ax10[1][1].set_title("Overlap of Control and Control$+\Phi$SDM\npTyr Peptides",
                         fontfamily = "sans-serif", font = "Arial",
                         fontweight = "bold", fontsize = 16)
    for text in control_v_csdm.subset_labels:
        text.set_fontfamily("sans-serif")
        text.set_font("Arial")
        text.set_fontweight("bold")
        text.set_fontsize(12)
    plt.savefig("figures/set_overlap.pdf")
    plt.show()
    print("\tGenerating Figure 3B: q-value Volcano Plot\n")
    fig6, ax6 = plt.subplots(4,4,figsize = (10,12), gridspec_kw = {"hspace" : 0,
                                                               "wspace" : 0.01})
    handles = []
    leg_labs = []
    for i in range(4):
        for j in range(3,4):
            ax6[i][j].spines["top"].set_visible(False)
            ax6[i][j].spines["bottom"].set_visible(False)
            ax6[i][j].spines["left"].set_visible(False)
            ax6[i][j].spines["right"].set_visible(False)
            ax6[i][j].set_xticks([])
            ax6[i][j].set_yticks([])
    for i in range(len(qr_data)):
        for j in range(len(qr_data[i])):
            if j == 2:
                ax6[i][j].scatter(*qr_data[i][j], color = colours[i],
                                  label = labels[i],
                               s=20, edgecolor = "black", linewidth = 0.2)
            else:
                ax6[i][j].scatter(*qr_data[i][j], color = colours[i],
                               s=20, edgecolor = "black", linewidth = 0.2)
            
            if i == 0 and j == 0:
                ax6[i][j].plot([-1.5,2],[-log10(0.05),-log10(0.05)], 
                           color = "black", linestyle = ":", label = "$q=0.05$")
            else:
                ax6[i][j].plot([-1.5,2],[-log10(0.05),-log10(0.05)], 
                           color = "black", linestyle = ":")
            if j == 0:
                ax6[i][j].plot([log10(10),log10(10)],[-0.5,4], 
                               color = "black", linestyle = "dashed", alpha = 0.5)
                ax6[i][j].plot([log10(20),log10(20)],[-0.5,4], color = "grey",alpha = 0.1)
                ax6[i][j].plot([log10(5),log10(5)],[-0.5,4], color = "grey", alpha = 0.1)
                ax6[i][j].fill_between([log10(5),log10(20)],-0.5,4,color="grey",alpha=0.1)
                ax6[i][j].text(0,2.5,
                               f"{qcounts[i][j]}\n$({(qcounts[i][j]/len(qr_data[i][j][1])*100):.0f}\%)$",
                               fontfamily = "sans-serif", font = "Arial", fontweight = "bold",
                               va = "center", ha = "right", fontsize = 14)
                ax6[i][j].text(0,1,
                               f"{len(qr_data[i][j][1])-qcounts[i][j]}\n$({((len(qr_data[i][j][1])-qcounts[i][j])/len(qr_data[i][j][1])*100):.0f}\%)$",
                               fontfamily = "sans-serif", font = "Arial", fontweight = "bold",
                               va = "top", ha = "right", fontsize = 14)
            elif j == 1:
                ax6[i][j].plot([log10(3 + 1/3),log10(3 + 1/3)],[-0.5,4], 
                               color = "black", linestyle = "dashed", alpha = 0.5)
                ax6[i][j].plot([log10((3 + 1/3)/2),log10((3 + 1/3)/2)],[-0.5,4], color = "grey",alpha = 0.1)
                ax6[i][j].plot([log10((3 + 1/3)*2), log10((3 + 1/3)*2)],[-0.5,4], color = "grey", alpha = 0.1)
                ax6[i][j].fill_between([log10((3 + 1/3)/2),log10((3 + 1/3)*2)],-0.5,4,color="grey",alpha=0.1)
                ax6[i][j].text(-0.5,2.5,
                               f"{qcounts[i][j]}\n$({(qcounts[i][j]/len(qr_data[i][j][1])*100):.0f}\%)$",
                               fontfamily = "sans-serif", font = "Arial", fontweight = "bold",
                               va = "center", ha = "right", fontsize = 14)
                ax6[i][j].text(-0.5,1,
                               f"{len(qr_data[i][j][1])-qcounts[i][j]}\n$({((len(qr_data[i][j][1])-qcounts[i][j])/len(qr_data[i][j][1])*100):.0f}\%)$",
                               fontfamily = "sans-serif", font = "Arial", fontweight = "bold",
                               va = "top", ha = "right", fontsize = 14)
            else:
                ax6[i][j].plot([log10(3),log10(3)],[-0.5,4], 
                               color = "black", linestyle = "dashed", alpha = 0.5)
                ax6[i][j].plot([log10(3/2),log10(3/2)],[-0.5,4], color = "grey",alpha = 0.1)
                ax6[i][j].plot([log10(3*2), log10(3*2)],[-0.5,4], color = "grey", alpha = 0.1)
                ax6[i][j].fill_between([log10(3/2),log10(3*2)],-0.5,4,color="grey",alpha=0.1)
                ax6[i][j].text(-0.5,2.5,
                               f"{qcounts[i][j]}\n$({(qcounts[i][j]/len(qr_data[i][j][1])*100):.0f}\%)$",
                               fontfamily = "sans-serif", font = "Arial", fontweight = "bold",
                               va = "center", ha = "right", fontsize = 14)
                ax6[i][j].text(-0.5,1,
                               f"{len(qr_data[i][j][1])-qcounts[i][j]}\n$({((len(qr_data[i][j][1])-qcounts[i][j])/len(qr_data[i][j][1])*100):.0f}\%)$",
                               fontfamily = "sans-serif", font = "Arial", fontweight = "bold",
                               va = "top", ha = "right", fontsize = 14)
            ax6[i][j].set_ylim(-0.5,4)
            ax6[i][j].set_xlim(-1.5,2)
            ax6[i][j].spines["right"].set_visible(False)
            ax6[i][j].spines["top"].set_visible(False)
            if i != 3:
                ax6[i][j].spines["bottom"].set_visible(False)
                ax6[i][j].set_xticks([])
            else:
                ax6[i][j].set_xticks([-1,0,1])
                ax6[i][j].set_xticklabels(["-1","0","1"], fontfamily = "sans-serif",
                                         font = "Arial", fontweight = "bold", fontsize = 16)
            if j != 0:
                ax6[i][j].spines["left"].set_visible(False)
                ax6[i][j].set_yticks([])
            else:
                ax6[i][j].set_yticks([0,1,2,3,4])
                ax6[i][j].set_yticklabels(["0", "1", "2", "3", "4"],
                                          fontfamily = "sans-serif", font= "Arial",
                                          fontweight = "bold", fontsize = 16)
    labs = [axx.get_legend_handles_labels() for axx in fig6.axes]
    handles, labs = [sum(lol, []) for lol in zip(*labs)]
    fig6.text(0.43, 0.07, '$\log_{10}$(Intensity Ratio)', ha='center', fontfamily = "sans-serif",
              font = "Arial", fontsize = 18)
    fig6.text(0.04, 0.5, '\n$-\log_{10}$($q$)', va='center', rotation='vertical', fontfamily = "sans-serif",
              font = "Arial", fontsize = 18)
    ax6[2][3].legend(handles, labs, loc="upper left", prop = {"family" : "Arial",
                                                             "weight" : "bold",
                                                                "size" : 16})
    ax6[0][0].set_title("1.0 mg : 0.1 mg", va = "bottom", ha = "center", fontfamily = "sans-serif",
              font = "Arial", fontsize = 18, fontweight = "bold",
                       bbox=dict(facecolor='none', edgecolor='black'))
    ax6[0][1].set_title("1.0 mg : 0.3 mg", va = "bottom", ha = "center", fontfamily = "sans-serif",
              font = "Arial", fontsize = 18, fontweight = "bold",
                       bbox=dict(facecolor='none', edgecolor='black'))
    ax6[0][2].set_title("0.3 mg : 0.1 mg", va = "bottom", ha = "center", fontfamily = "sans-serif",
              font = "Arial", fontsize = 18, fontweight = "bold",
                       bbox=dict(facecolor='none', edgecolor='black'))
    plt.savefig("figures/q_volcano.pdf")
    plt.show()
    print("\tGenerating Figure 2A: Reporter Ion Count\n")
    fig7, ax7 = plt.subplots(2,2,figsize = (12,8), gridspec_kw = {"hspace" : 1,
                                                                 "wspace" :0.01})

    cooolor = 0
    for i in range(2):
        for j in range(2):
            ax7[i][j].bar([k for k in range(11)], 
                          report_counts[i][j], 
                          color = colours[cooolor], edgecolor = "black")
            for k in range(11):
                ax7[i][j].text(k, report_counts[i][j][k] + 10, 
                               f"{report_counts[i][j][k]}", va = "bottom", ha = "center",
                               fontfamily = "sans-serif", font = "Arial", 
                               fontweight = "bold", fontsize = 12)#, rotation = 45)
            ax7[i][j].set_xticks([k for k in range(11)])
            cooolor+=1
            if i ==0:
                ax7[i][j].set_ylim(0,400)
                ax7[i][j].set_xticklabels(control_labs, rotation = 90,
                                         fontfamily = "sans-serif", font = "Arial",
                                          fontweight = "bold", fontsize = 16)
            else:
                ax7[i][j].set_ylim(0,7000)
                ax7[i][j].set_xticklabels(boost_labs, rotation = 90,
                                         fontfamily = "sans-serif", font = "Arial",
                                          fontweight = "bold", fontsize = 16)
            ax7[i][j].spines["top"].set_visible(False)
            ax7[i][j].spines["right"].set_visible(False)
            if j == 1:
                ax7[i][j].spines["left"].set_visible(False)
                ax7[i][j].set_yticks([])
            ax7[i][j].set_title(plot_heads[i][j], fontfamily = "sans-serif", font = "Arial",
                                fontsize = 22, fontweight = "bold",
                                bbox=dict(facecolor='none', edgecolor='black'))
            ax7[i][j].set_yticks(ax7[i][j].get_yticks())
            ax7[i][j].set_yticklabels([str(int(d)) for d in ax7[i][j].get_yticks()], fontfamily = "sans-serif",
                                     font = "Arial", fontweight = "bold", fontsize = 16)
    fig7.text(0.04, 0.5, f'$\#$ of unique pTyr peptides\nwith quantified reporters', ha='center', fontfamily = "sans-serif",
              font = "Arial", fontsize = 20, rotation = 90,
              va = "center", fontweight = "bold")#, ha = "center")
    plt.subplots_adjust(bottom = 0.15)
    plt.savefig("figures/reporter_barchart.pdf")
    plt.show()
    print("\tGenerating Figure 2B: Missing Value Barchart\n")
    fig8, ax8 = plt.subplots(2,2,figsize = (12,8), gridspec_kw = {"hspace" : 1,
                                                                 "wspace" :0.01})
    percs = False
    cooolor = 0
    for i in range(2):
        for j in range(2):
            if percs:
                ax8[i][j].bar([k for k in range(11)], 
                              [100 for _ in range(11)], 
                              color = colours[cooolor], alpha = 0.5)
                ax8[i][j].bar([k for k in range(11)], 
                              report_percs[i][j], 
                              color = colours[cooolor], edgecolor = "black")
            else:
                ax8[i][j].bar([k for k in range(11)], 
                              miss_counts[i][j], 
                              color = colours[cooolor], edgecolor = "black")
            for k in range(11):
                if percs:
                    ax8[i][j].text(k, 100 + 3, 
                                   f"{report_percs[i][j][k]:.1f}", va = "bottom", ha = "center",
                                   fontfamily = "sans-serif", font = "Arial", fontweight = "bold")
                else:
                    if i == 0 and k != 10:
                        ax8[i][j].text(k, 303, 
                                   f"{report_percs[i][j][k]:.1f}", va = "bottom", ha = "center",
                                   fontfamily = "sans-serif", font = "Arial", fontweight = "bold")
                    elif i == 0 and k == 10:
                        ax8[i][j].text(k, 303, 
                                   f"{report_percs[i][j][k]:.1f}%", va = "bottom", ha = "center",
                                   fontfamily = "sans-serif", font = "Arial", fontweight = "bold")
                    elif i == 1 and k != 10:
                        ax8[i][j].text(k, 6003, 
                                   f"{report_percs[i][j][k]:.1f}", va = "bottom", ha = "center",
                                   fontfamily = "sans-serif", font = "Arial", fontweight = "bold")
                    else:
                        ax8[i][j].text(k, 6003, 
                                   f"{report_percs[i][j][k]:.1f}%", va = "bottom", ha = "center",
                                   fontfamily = "sans-serif", font = "Arial", fontweight = "bold")
            ax8[i][j].set_xticks([k for k in range(11)])
            cooolor+=1
            if i ==0:
                if percs:
                    ax8[i][j].set_ylim(0,100)
                else:
                    ax8[i][j].set_ylim(0,300)
                ax8[i][j].set_xticklabels(control_labs, rotation = 90,
                                         fontfamily = "sans-serif", font = "Arial",
                                          fontweight = "bold", fontsize = 16)
            else:
                if percs:
                    ax8[i][j].set_ylim(0,100)
                else:
                    ax8[i][j].set_ylim(0,6000)
                ax8[i][j].set_xticklabels(boost_labs, rotation = 90,
                                         fontfamily = "sans-serif", font = "Arial",
                                          fontweight = "bold", fontsize = 16)
            ax8[i][j].spines["top"].set_visible(False)
            ax8[i][j].spines["right"].set_visible(False)
            if j == 1:
                ax8[i][j].spines["left"].set_visible(False)
                ax8[i][j].set_yticks([])
            ax8[i][j].set_title(plot_heads[i][j], fontfamily = "sans-serif", font = "Arial",
                                fontsize = 20, fontweight = "bold",
                                bbox=dict(facecolor='none', edgecolor='black'))
            ax8[i][j].set_yticks(ax8[i][j].get_yticks())
            ax8[i][j].set_yticklabels([str(int(d)) for d in ax8[i][j].get_yticks()], fontfamily = "sans-serif",
                                     font = "Arial", fontweight = "bold", fontsize = 16)
            if percs:
                ax8[i][j].set_ylim(0,120)
            elif i == 0:
                ax8[i][j].set_ylim(0,360)
            else:
                ax8[i][j].set_ylim(0,7200)
    if percs:
        fig8.text(0.04, 0.5, f'$\%$ of unique pTyr peptides\nwith missing values', ha='center', fontfamily = "sans-serif",
                  font = "Arial", fontsize =20, rotation = 90,
                  va = "center", fontweight = "bold")#, ha = "center")
    else:
        fig8.text(0.04, 0.5, f'$\#$ of unique pTyr peptides\nwith missing values', ha='center', fontfamily = "sans-serif",
                  font = "Arial", fontsize = 20, rotation = 90,
                  va = "center", fontweight = "bold")#, ha = "center")
    plt.subplots_adjust(bottom = 0.15)
    if percs:
        plt.savefig("figures/reporter_nans_barchart_percentages.pdf")
    else:
        plt.savefig("figures/reporter_nans_barchart.pdf")
    plt.show()
    print("\tGenerating Figure 4C: BOOST Factor Histograms\n")
    fig11, ax11 = plt.subplots(2,3, figsize = (16,12),
                               gridspec_kw={"wspace" : 0.1,
                                            "hspace" : 0.05})
    histcolours = [[colours[3], colours[2]],
                   ["mediumaquamarine", "paleturquoise"]]
    groups = [["BOOST", "BOOST$+\Phi$SDM"],
              ["Overlap (BOOST)", "Overlap (BOOST$+\Phi$SDM)"]]
    overunder = [[get_above_below(facts[i][j]) for j in range(2)] for i in range(2)]
    use_bins = None
    for i in range(2):
        for j in range(2):
            hist, bins = np.histogram([item for item in facts[i][j] if item==item],
                                      bins = 50)
            logbins = np.logspace(log10(bins[0]),log10(bins[-1]),len(bins))
            ax11[i][j].hist(facts[i][j], bins = logbins,
                           color = histcolours[i][j], label = groups[i][j], edgecolor = "black")
            ax11[i][j].set_xscale("log")
            ax11[i][j].set_xlim((0.1, 100000.0))
            ax11[i][j].set_ylim(0,200)
            ax11[i][j].spines["top"].set_visible(False)
            ax11[i][j].spines["right"].set_visible(False)
            if i == 0 and j == 0:
                xtix = ax11[i][j].get_xticks()
                xlim = ax11[i][j].get_xlim()
                xtixlabs = [""] + [str(item) for item in xtix[1:3]] + [str(int(item)) for item in xtix[3:-1]] + [""]
            if j == 0:
                ax11[i][j].set_yticks(ax11[i][j].get_yticks()[:-1])
                ax11[i][j].set_yticklabels([str(int(item)) for item in ax11[i][j].get_yticks()],
                           fontfamily = "sans-serif", font = "Arial", fontweight = "bold",
                            fontsize = 18, ha = "right", va = "center")
            else:
                ax11[i][j].set_yticks([])
                ax11[i][j].spines["left"].set_visible(False)
            if i == 1:
                ax11[i][j].set_xticks(xtix[1:])
                ax11[i][j].tick_params(axis = "x", which = "minor", length = 6, width = 2)
                ax11[i][j].tick_params(axis = "x", which = "major", length = 12, width = 2)
                ax11[i][j].set_xticklabels(xtixlabs[1:], fontfamily = "sans-serif", font = "Arial", fontweight = "bold",
                            fontsize = 18, ha = "right", va = "top", rotation=45)
            else:
                ax11[i][j].set_ylim(-0.2,200)
                ax11[i][j].set_xticks(xtix[1:])
                ax11[i][j].tick_params(axis = "x", which = "minor", length = 6, width = 2)
                ax11[i][j].tick_params(axis = "x", which = "major", length = 12, width = 2)
                ax11[i][j].set_xticklabels(["" for _ in range(len(xtix[1:]))])
                ax11[i][j].spines["bottom"].set_visible(False)
            ax11[i][j].set_xlim((0.01, 100000.0))
            ax11[i][j].plot([0.1,0.1], [0,200], color = "grey", linestyle = (0, (5, 10)), alpha = 0.5)
            ax11[i][j].plot([1,1], [0,200], color = "grey", linestyle = ":", alpha = 1)
            ax11[i][j].plot([10,10], [0,200], color = "grey", linestyle = (0, (5, 10)), alpha = 0.5)
            ax11[i][j].plot([100,100], [0,200], color = "grey", linestyle = (0, (5, 10)), alpha = 0.5)
            ax11[i][j].plot([1000,1000], [0,200], color = "grey", linestyle = (0, (5, 10)), alpha = 0.5)
            ax11[i][j].plot([10000,10000], [0,200], color = "grey", linestyle = (0, (5, 10)), alpha = 0.5)
            ax11[i][j].plot([100000,100000], [0,200], color = "grey", linestyle = (0, (5, 10)), alpha = 0.5)
            ax11[i][j].text(0.7, 155, overunder[i][j][0], fontfamily = "sans-serif", font = "Arial", fontweight = "bold",
                            fontsize = 16, ha = "right", bbox=dict(facecolor='white',alpha = 1, edgecolor = "none"))
            ax11[i][j].text(1.8, 155, overunder[i][j][1], fontfamily = "sans-serif", font = "Arial", fontweight = "bold",
                            fontsize = 16, ha = "left", bbox=dict(facecolor='white',alpha = 1, edgecolor = "none"))
            ax11[i][j].text(10000, 100, f"Median$ = {meds[i][j]:.2f}$",
                           fontfamily = "sans-serif", font = "Arial", fontweight = "bold",
                            fontsize = 16, ha = "center",
                            bbox=dict(facecolor='white',alpha = 1, edgecolor = "none"))
    for i in range(2):
        for j in range(2,3):
            ax11[i][j].set_xticks([])
            ax11[i][j].set_yticks([])
            ax11[i][j].spines["top"].set_visible(False)
            ax11[i][j].spines["bottom"].set_visible(False)
            ax11[i][j].spines["right"].set_visible(False)
            ax11[i][j].spines["left"].set_visible(False)
    labs = [axx.get_legend_handles_labels() for axx in fig11.axes]
    handles, labs = [sum(lol, []) for lol in zip(*labs)]
    ax11[1][2].legend(handles, labs, loc="center", bbox_to_anchor = (0.5,1),
                      prop = {"family" : "Arial",
                              "weight" : "bold",
                              "size" : 16})
    fig11.text(0, 0.5, '\n\n# of Unique pTyr peptides', 
              va='center', rotation='vertical', fontfamily = "sans-serif",
              font = "Arial", fontweight = "bold", fontsize = 20)
    fig11.text(0.35, 0.02, 'BOOST Factor', 
              va='center', fontfamily = "sans-serif",
              font = "Arial", fontweight = "bold", fontsize = 20)
    plt.savefig("figures/boostfactor_hists.pdf")
    plt.show()
    print("\tGenerating Supporting Figures 4-6: Replicate Intensities vs the Means\n")
    group_labs = ["Control$+\Phi$SDM",
                  "Control",
                  "BOOST$+\Phi$SDM",
                  "BOOST"]

    cs_report_signal = [[experiments["Control$+\Phi$SDM"][j][i] for j in range(1,len(experiments["Control$+\Phi$SDM"]))]
                        for i in range(11)]  # For columns 0-10, which are the reporter channels
    # Output is a list of 11 lists, each with the data from one channel

    c_report_signal = [[experiments["Control"][j][i] for j in range(1,len(experiments["Control"]))]
                       for i in range(11)]

    bs_report_signal = [[experiments["BOOST$+\Phi$SDM"][j][i] for j in range(1,len(experiments["BOOST$+\Phi$SDM"]))]
                       for i in range(11)]

    b_report_signal = [[experiments["BOOST"][j][i] for j in range(1,len(experiments["BOOST"]))]
                       for i in range(11)]

    cs_report_signal = gh.transpose(*cs_report_signal)
    c_report_signal = gh.transpose(*c_report_signal)
    bs_report_signal = gh.transpose(*bs_report_signal)
    b_report_signal = gh.transpose(*b_report_signal)

    # grab reps and what not, no filtering by number of values
    cs_report_1_123 = [[log10(item[x]) for x in ten_x] for item in cs_report_signal if all([item[x] == item[x] for x in ten_x])]
    cs_report_1_mean = [sh.mean(item) for item in cs_report_1_123]
    c_report_1_123 = [[log10(item[x]) for x in ten_x] for item in c_report_signal if all([item[x] == item[x] for x in ten_x])]
    c_report_1_mean = [sh.mean(item) for item in c_report_1_123]
    bs_report_1_123 = [[log10(item[x]) for x in ten_x] for item in bs_report_signal if all([item[x] == item[x] for x in ten_x])]
    bs_report_1_mean = [sh.mean(item) for item in bs_report_1_123]
    b_report_1_123 = [[log10(item[x]) for x in ten_x] for item in b_report_signal if all([item[x] == item[x] for x in ten_x])]
    b_report_1_mean = [sh.mean(item) for item in b_report_1_123]

    ten_report_matrix = [[[row[i] for i in range(3)] for row in cs_report_1_123],
                         [[row[i] for i in range(3)] for row in c_report_1_123],
                         [[row[i] for i in range(3)] for row in bs_report_1_123],
                         [[row[i] for i in range(3)] for row in b_report_1_123]]
    ten_mean_lists = [cs_report_1_mean, c_report_1_mean, bs_report_1_mean,b_report_1_mean]

    cs_report_03_123 = [[log10(item[x]) for x in three_x] for item in cs_report_signal if all([item[x] == item[x] for x in three_x])]
    cs_report_03_mean = [sh.mean(item) for item in cs_report_03_123]
    c_report_03_123 = [[log10(item[x]) for x in three_x] for item in c_report_signal if all([item[x] == item[x] for x in three_x])]
    c_report_03_mean = [sh.mean(item) for item in c_report_03_123]
    bs_report_03_123 = [[log10(item[x]) for x in three_x] for item in bs_report_signal if all([item[x] == item[x] for x in three_x])]
    bs_report_03_mean = [sh.mean(item) for item in bs_report_03_123]
    b_report_03_123 = [[log10(item[x]) for x in three_x] for item in b_report_signal if all([item[x] == item[x] for x in three_x])]
    b_report_03_mean = [sh.mean(item) for item in b_report_03_123]

    three_report_matrix = [[[row[i] for i in range(3)] for row in cs_report_03_123],
                         [[row[i] for i in range(3)] for row in c_report_03_123],
                         [[row[i] for i in range(3)] for row in bs_report_03_123],
                         [[row[i] for i in range(3)] for row in b_report_03_123]]
    three_mean_lists = [cs_report_03_mean, c_report_03_mean, bs_report_03_mean,b_report_03_mean]


    cs_report_01_123 = [[log10(item[x]) for x in one_x] for item in cs_report_signal if all([item[x] == item[x] for x in one_x])]
    cs_report_01_mean = [sh.mean(item) for item in cs_report_01_123]
    c_report_01_123 = [[log10(item[x]) for x in one_x] for item in c_report_signal if all([item[x] == item[x] for x in one_x])]
    c_report_01_mean = [sh.mean(item) for item in c_report_01_123]
    bs_report_01_123 = [[log10(item[x]) for x in one_x] for item in bs_report_signal if all([item[x] == item[x] for x in one_x])]
    bs_report_01_mean = [sh.mean(item) for item in bs_report_01_123]
    b_report_01_123 = [[log10(item[x]) for x in one_x] for item in b_report_signal if all([item[x] == item[x] for x in one_x])]
    b_report_01_mean = [sh.mean(item) for item in b_report_01_123]

    one_report_matrix = [[[row[i] for i in range(3)] for row in cs_report_01_123],
                         [[row[i] for i in range(3)] for row in c_report_01_123],
                         [[row[i] for i in range(3)] for row in bs_report_01_123],
                         [[row[i] for i in range(3)] for row in b_report_01_123]]
    one_mean_lists = [cs_report_01_mean, c_report_01_mean,
                      bs_report_01_mean,b_report_01_mean]

    art_labels = ["$\log_{10}($R1$)$","$\log_{10}($R2$)$","$\log_{10}($R3$)$"]

    figgg, axxx = plt.subplots(4,3,figsize = (15,15))

    for i in range(4):
        for j in range(3):
            scatter_gausskde(ten_mean_lists[i],
                             [row[j] for row in ten_report_matrix[i]], axxx[i][j], colour = colours[i],
                             scatterargs = {"s" : 20,
                                                   "alpha" : 0.25,
                                                   "marker" : "o",
                                             "edgecolor" : "black"},
                             xlim = [0,8],
                             ylim = [0,8])
            axxx[i][j].legend(loc="upper left")
            axxx[i][j].set_xlabel("$\log_{10}($Geometric Mean$)$", fontfamily = "sans-serif",
                                  font = "Arial", fontsize = 10)
            axxx[i][j].set_ylabel(art_labels[j], fontfamily = "sans-serif",
                                  font = "Arial", fontsize = 10)
            above = sum([1 for k in range(len(ten_mean_lists[i])) if [row[j] for row in ten_report_matrix[i]][k] > ten_mean_lists[i][k] and [row[j] for row in ten_report_matrix[i]][k] == [row[j] for row in ten_report_matrix[i]][k]])
            below = len([row[j] for row in ten_report_matrix[i] if row[j] == row[j]]) - above
            #axxx[i][j].text(3,1,f"Reporter $> \mu$: $n={above}$", ha = "left", fontfamily = "sans-serif",
            #                      font = "Arial", fontsize = 10)
            #axxx[i][j].text(3,.5,f"Reporter $< \mu$: $n={below}$", ha = "left", fontfamily = "sans-serif",
            #                      font = "Arial", fontsize = 10)
            if j==2:
                axxx2= axxx[i][j].twinx()
                axxx2.set_ylabel(group_labs[i], rotation = 270, fontfamily = "sans-serif",
                                 font = "Arial", fontweight = "bold", fontsize = 18,
                                 bbox=dict(facecolor='none', edgecolor='black'), labelpad = 6,
                                va = "bottom")
                axxx2.set_yticks([])
                axxx2.set_yticklabels([])
            #axxx[i][j].scatter(ten_mean_lists[i], [row[j] for row in ten_report_matrix[i]],
            #                   color = colours[i], s=20, edgecolor = "black")
            
    figgg.suptitle("\n\n\nReplicates vs the Mean for 1.0 mg Protein Input",
                   fontfamily = "sans-serif", font = "Arial", fontweight = "bold",
                   fontsize = 18)
    plt.savefig("figures/reps_vs_mean_1mg.pdf")
    plt.show()

    figgg, axxx = plt.subplots(4,3,figsize = (15,15))

    for i in range(4):
        for j in range(3):
            scatter_gausskde(three_mean_lists[i],
                             [row[j] for row in three_report_matrix[i]], axxx[i][j], colour = colours[i],
                             scatterargs = {"s" : 20,
                                                   "alpha" : 0.25,
                                                   "marker" : "o",
                                             "edgecolor" : "black"},
                             xlim = [0,8],
                             ylim = [0,8])
            axxx[i][j].legend(loc="upper left")
            axxx[i][j].set_xlabel("$\log_{10}($Geometric Mean$)$", fontfamily = "sans-serif",
                                  font = "Arial", fontsize = 10)
            axxx[i][j].set_ylabel(art_labels[j], fontfamily = "sans-serif",
                                  font = "Arial", fontsize = 10)
            above = sum([1 for k in range(len(three_mean_lists[i])) if [row[j] for row in three_report_matrix[i]][k] > three_mean_lists[i][k] and [row[j] for row in three_report_matrix[i]][k] == [row[j] for row in three_report_matrix[i]][k]])
            below = len([row[j] for row in three_report_matrix[i] if row[j] == row[j]]) - above
            #axxx[i][j].text(3,1,f"Reporter $> \mu$: $n={above}$", ha = "left", fontfamily = "sans-serif",
            #                      font = "Arial", fontsize = 10)
            #axxx[i][j].text(3,.5,f"Reporter $< \mu$: $n={below}$", ha = "left", fontfamily = "sans-serif",
            #                      font = "Arial", fontsize = 10)
            if j==2:
                axxx2= axxx[i][j].twinx()
                axxx2.set_ylabel(group_labs[i], rotation = 270, fontfamily = "sans-serif",
                                 font = "Arial", fontweight = "bold", fontsize = 18,
                                 bbox=dict(facecolor='none', edgecolor='black'), labelpad = 6,
                                va = "bottom")
                axxx2.set_yticks([])
                axxx2.set_yticklabels([])
            #axxx[i][j].scatter(ten_mean_lists[i], [row[j] for row in ten_report_matrix[i]],
            #                   color = colours[i], s=20, edgecolor = "black")
            
    figgg.suptitle("\n\n\nReplicates vs the Mean for 0.3 mg Protein Input",
                   fontfamily = "sans-serif", font = "Arial", fontweight = "bold",
                   fontsize = 18)
    plt.savefig("figures/reps_vs_mean_03mg.pdf")
    plt.show()

    figgg, axxx = plt.subplots(4,3,figsize = (15,15))

    for i in range(4):
        for j in range(3):
            scatter_gausskde(one_mean_lists[i],
                             [row[j] for row in one_report_matrix[i]], axxx[i][j], colour = colours[i],
                             scatterargs = {"s" : 20,
                                                   "alpha" : 0.25,
                                                   "marker" : "o",
                                             "edgecolor" : "black"},
                             xlim = [0,8],
                             ylim = [0,8])
            axxx[i][j].legend(loc="upper left")
            axxx[i][j].set_xlabel("$\log_{10}($Geometric Mean$)$", fontfamily = "sans-serif",
                                  font = "Arial", fontsize = 10)
            axxx[i][j].set_ylabel(art_labels[j], fontfamily = "sans-serif",
                                  font = "Arial", fontsize = 10)
            above = sum([1 for k in range(len(one_mean_lists[i])) if [row[j] for row in one_report_matrix[i]][k] > one_mean_lists[i][k] and [row[j] for row in one_report_matrix[i]][k] == [row[j] for row in one_report_matrix[i]][k]])
            below = len([row[j] for row in one_report_matrix[i] if row[j] == row[j]]) - above
            #axxx[i][j].text(3,1,f"Reporter $> \mu$: $n={above}$", ha = "left", fontfamily = "sans-serif",
            #                     font = "Arial", fontsize = 10)
            #axxx[i][j].text(3,.5,f"Reporter $< \mu$: $n={below}$", ha = "left", fontfamily = "sans-serif",
            #                      font = "Arial", fontsize = 10)
            if j==2:
                axxx2= axxx[i][j].twinx()
                axxx2.set_ylabel(group_labs[i], rotation = 270, fontfamily = "sans-serif",
                                 font = "Arial", fontweight = "bold", fontsize = 18,
                                 bbox=dict(facecolor='none', edgecolor='black'), labelpad = 6,
                                va = "bottom")
                axxx2.set_yticks([])
                axxx2.set_yticklabels([])
            #axxx[i][j].scatter(ten_mean_lists[i], [row[j] for row in ten_report_matrix[i]],
            #                   color = colours[i], s=20, edgecolor = "black")
            
    figgg.suptitle("\n\n\nReplicates vs the Mean for 0.1 mg Protein Input",
                   fontfamily = "sans-serif", font = "Arial", fontweight = "bold",
                   fontsize = 18)
    plt.savefig("figures/reps_vs_mean_01mg.pdf")
    plt.show()
    print("Complete! Check the './figures' and './curated_results' folders for the output!\n")
    
#
#
#########################################################################################################
#
#  

if __name__ == "__main__":
    main()

#
#
#########################################################################################################
```
