## Supporting Folder 2 Data Analysis for "Mouse primary T cell phosphotyrosine proteomics enabled by BOOST": PTPN6_sites.pdf

| | |
|---|---|
| Y61 | Y61 |
| Y64 | Y64 |
| Y213 | Y213 |
| Y214 | Y214 |
| Y276 | Y276 |
| Y301 | Y301 |
| Y306 | Y306 |
| Y374 | Y374 |
| Y377 | Y377 |
| Y536 | Y536 |
| Y541 | Y541 |
| Y564 | Y564 |
