## Supporting Folder 2 Data Analysis for "Mouse primary T cell phosphotyrosine proteomics enabled by BOOST": PTPRC_sites.pdf

| | |
|---|---|
| Y631 | Y640 |
| Y672 | Y681 |
| Y678 | Y687 |
| Y680 | Y689 |
| Y711 | Y720 |
| Y754 | Y763 |
| Y781 | Y790 |
| Y852 | Y861 |
| Y871 | Y880 |
| Y937 | F946 |
| Y969 | Y978 |
| F971 | Y980 |
