## Supporting Folder 2 Data Analysis for "Mouse primary T cell phosphotyrosine proteomics enabled by BOOST": total.pdf

| Protein | Mouse | Human | Protein | Mouse | Human | Protein | Mouse | Human | Protein | Mouse | Human |
|---|---|---|---|---|---|---|---|---|---|---|---|
| Akt2 | Y122 | Y122 | GADS | Y45 | Y45 | p85$\alpha$ | Y470 | Y470 | TCR$\zeta$ | Y72 | Y72 |
| CARD11 | Y489 | Y489 | | Y218 | Y222 | | Y688 | Y688 | | Y83 | Y83 |
| Cbl-b | Y363 | Y363 | Grb2 | Y209 | Y209 | p85$\beta$ | Y458 | Y464 | | Y111 | Y111 |
| | Y763 | Y763 | GSK3$\beta$ | Y216 | Y216 | PAK1 | Y142 | Y142 | | Y123 | Y123 |
| CD28 | Y189 | Y191 | Itk | Y40 | Y40 | | Y153 | Y153 | | Y142 | Y142 |
| | Y204 | Y206 | | Y126 | Y120 | | Y474 | Y474 | | Y153 | Y153 |
| | Y207 | Y209 | | Y226 | Y220 | PAK2 | Y130 | Y130 | Tec | Y205 | Y206 |
| CD3$\delta$ | Y149 | Y149 | | Y243 | Y237 | | Y139 | Y139 | | Y227 | Y228 |
| | Y160 | Y160 | | Y517 | Y512 | | Y453 | Y453 | | Y280 | Y281 |
| CD3$\varepsilon$ | Y170 | Y188 | Jnk1 | Y185 | Y185 | PAK6 | Y366 | Y365 | | Y518 | Y519 |
| | Y181 | Y199 | Jnk2 | Y185 | Y185 | PD-1 | Y225 | Y223 | Vav1 | Y110 | Y110 |
| CD3$\gamma$ | Y160 | Y160 | Jnk3 | Y223 | Y223 | PKC$\theta$ | Y28 | Y28 | | Y192 | Y192 |
| | Y171 | Y171 | LAT | Y46 | Y45 | | Y545 | Y545 | | Y541 | Y541 |
| CD45 | Y631 | Y640 | | Y195 | Y220 | PLC$\gamma$1 | Y210 | Y210 | | Y791 | Y791 |
| | Y672 | Y681 | Lck | Y192 | Y192 | | Y472 | Y472 | Vav2 | Y142 | Y142 |
| | Y678 | Y687 | | Y394 | Y394 | | Y771 | Y771 | Vav3 | Y141 | Y141 |
| | Y680 | Y689 | | Y414 | Y414 | | Y775 | Y775 | | Y217 | Y217 |
| | Y711 | Y720 | | Y470 | Y470 | | Y783 | Y783 | Zap70 | Y69 | Y69 |
| | Y754 | Y763 | | Y505 | Y505 | | Y1003 | Y1003 | | Y87 | Y87 |
| | Y781 | Y790 | NCK1 | Y13 | Y13 | | Y1253 | Y1253 | | Y164 | Y164 |
| | Y852 | Y861 | | Y55 | Y55 | RHOA | Y66 | Y66 | | Y178 | Y178 |
| | Y871 | Y880 | | Y105 | Y105 | SHP-1 | Y61 | Y61 | | Y198 | Y198 |
| | Y937 | F946 | NCK2 | Y110 | Y110 | | Y64 | Y64 | | Y209 | Y209 |
| | Y969 | Y978 | NF$\kappa$B-p105 | Y238 | Y240 | | Y213 | Y213 | | Y211 | Y211 |
| | F971 | Y980 | NFAT1 | Y754 | Y752 | | Y214 | Y214 | | Y221 | Y221 |
| CDC42 | Y64 | Y64 | NFAT4 | Y86 | Y86 | | Y276 | Y276 | | Y248 | Y248 |
| CDK4 | Y17 | Y17 | | Y150 | Y150 | | Y301 | Y301 | | Y290 | Y292 |
| CTLA-4 | Y201 | Y201 | p110$\alpha$ | Y317 | Y317 | | Y306 | Y306 | | Y314 | Y315 |
| DLG1 | Y399 | Y399 | p110$\delta$ | Y523 | Y524 | | Y374 | Y374 | | Y396 | Y397 |
| | Y761 | Y760 | p38$\alpha$ | Y182 | Y182 | | Y377 | Y377 | | Y491 | Y492 |
| | Y785 | Y784 | p38$\beta$ | Y182 | Y182 | | Y536 | Y536 | | Y492 | Y493 |
| Erk1 | Y205 | Y204 | p38$\gamma$ | Y185 | Y185 | | Y541 | Y541 | | Y505 | Y506 |
| Erk2 | Y185 | Y187 | p55$\gamma$ | Y202 | Y202 | | Y564 | Y564 | | Y596 | Y597 |
| Fyn | Y28 | Y28 | p85$\alpha$ | Y76 | Y76 | SLP76 | Y173 | Y173 | | Y597 | Y598 |
| | Y214 | Y214 | | Y416 | Y416 | | Y483 | Y483 | | | |
| | Y420 | Y420 | | Y452 | Y452 | TAK1 | Y558 | Y585 | | | |
| | Y440 | Y440 | | Y467 | Y467 | TCR$\zeta$ | N64 | Y64 | | | |
