## Supporting Folder 2 Data Analysis for "Mouse primary T cell phosphotyrosine proteomics enabled by BOOST": ZAP70_sites.pdf

| | |
|---|---|
| Y69 | Y69 |
| Y87 | Y87 |
| Y164 | Y164 |
| Y178 | Y178 |
| Y198 | Y198 |
| Y209 | Y209 |
| Y211 | Y211 |
| Y221 | Y221 |
| Y248 | Y248 |
| Y290 | Y292 |
| Y314 | Y315 |
| Y396 | Y397 |
| Y491 | Y492 |
| Y492 | Y493 |
| Y505 | Y506 |
| Y596 | Y597 |
| Y597 | Y598 |
