## Supporting Folder 2 Data Analysis for "Mouse primary T cell phosphotyrosine proteomics enabled by BOOST": argcheck_helpers.py

```
"""
=================================================================================================
Kenneth P. Callahan

9 July 2021
                  
=================================================================================================
Python >= 3.8.5

archeck_helpers.py

This module contains functions that help with argument checking. Currently, they are mainly
type-checking and path checking functions.

As I need to check more things, I will put the functions here.
   
=================================================================================================
Dependencies:

There are no non-base Python dependencies. Currently, only sys, os, and glob are used.

=================================================================================================
"""
print(f"Loading the module: helpers.argcheck_helpers\n")
######################################################################################################
#
#     Imporatables

# os and sys are used for operating system level
# operations
import sys
import os

# glob is used to iterate over the files in directories
import glob

#
#
######################################################################################################
#
#     Checking Functions

def check_delim(argument,
                default = "\t"):
    """
    =================================================================================================
    check_delim(argument, default)
    
    This function is used to check file delimiters. Most files I see are tab delimited, hence
    the default being tab.
                  
    =================================================================================================
    Arguments:
    
    argument  ->  A string holding a (potential) delimiter
    defualt   ->  A string determining the default delimiter
    
    =================================================================================================
    Returns: The delimiter, after checking that it is a valid delimiter.
    
    =================================================================================================
    """
    # Assert thhat the argument is in the list of accepted arguments.
    assert argument in [None,':', '|', ';', '-', '\\', '/', '\\t', ',', "\t", "'\t'",
                        "':'", "'|'", "';'", "'-'", "'\\'", "'/'", "'\\t'" ,"','"], "The delimiter given is not accepted."
    # If the argument is None, then the argument is unset. Thus,
    # set it to the preset value of \t
    if argument == None:
        return default
    # If the argument is a properly formatted string, then just return the argument
    elif argument in [':', '|', ';', '-', '\\', '/', ',']:
        return argument
    # If the argument is an improperly formatted string (which happens when you pass
    # single-quote strings into the windows command line), then strip the single
    # quotes and return the properly formatted string
    elif argument in ["':'", "'|'", "';'", "'-'", "'\\'", "'/'" ,"','"]:
        return argument.strip("'")
    # Or if the argument is some form of the tab character, return the tab
    # character
    elif argument in ['\\t',"'\\t'", "\t","'\t'" ]:
        return "\t"
    # If something goes horribly wrong, simply return the tab character.
    else:
        return default

def check_existence(argument,
                    error = "An error occurred."):
    """
    =================================================================================================
    check_existence(argument, error)
    
    This function is meant to check the existence of a given path.
    
    =================================================================================================
    Arguments:
    
    argument  ->  A string containing a potential path
    error     ->  A string containing an error message, presented to the user if the path
                  does not exist.
    
    =================================================================================================
    Returns: The full path of the argument, or an error if the path does not exist.
    
    =================================================================================================
    """
    # Ensure that the argument exists, otherwise raise an error
    assert os.path.exists(argument), error
    # and if the argument exists, get the full path to the argument
    argument = os.path.abspath(argument)
    # and return the argument.
    return argument

def check_extension(argument,
                    desired_exts = ["txt", "py", "R"],
                    error = "I'm trapped in my computer. HELPPPPPP",
                    exists = True):
    """
    =================================================================================================
    check_extension(argument, desired_exts, error)
    
    This function is used to check whether a given filename has a proper extension.
    
    =================================================================================================
    Arguments:
    
    argument      ->  A string containing a file name (and possibly the path to the file)
    desired_exts  ->  A list of strings containing the allowable extensions, no periods are required.
    error         ->  A string containing an error message.
    exists        ->  A boolean that determines whether to check the path for existence
    
    =================================================================================================
    Returns:  The full path of the argument, or an error if the path does not exist or the extension
              is invalid.
    
    =================================================================================================
    """
    # Split the arugment on the period, and take the last element
    arg = argument.split(".")[-1]
    # Then assure that this is in the desired extensions.
    assert arg in desired_exts, error
    # If the exists is True
    if exists:
        # then check the argument for existence
        argument = check_existence(argument)
        # and return the argument
        return argument
    # OTherwise
    else:
        # Simply return the argument
        return argument

def check_dir(argument,
              create = True,
              error = "The directory does not exist, yet you have elected not to create it.\nProceed with caution."):
    """
    =================================================================================================
    check_dir(argument, create, error)
    
    This function is used to check for the existence of the argument, and whether or not to create
    the argument in the event the directory does not exist.
    
    =================================================================================================
    Arguments:
    
    argument  ->  A string contating the path to a directory
    create    ->  A boolean determining whether or not to create the directory, in the event it
                  does not exist.
    error     ->  A string containing an error message.
    
    =================================================================================================
    Returns: The argument after checking whether or not it exists and creating it if it did not.
    
    =================================================================================================
    """
    # If the path exists
    if os.path.exists(argument):
        # Return the absolute path to the argument
        return os.path.abspath(argument)
    # And if we create the argument
    elif create:
        # then create the directory to the argument
        os.makedirs(argument)
        # and return the absolute path to the argument.
        return os.path.abspath(argument)
    # If neither of these things happen,
    else:
        # print the error and return the argument.
        print(error)
        return argument
    
def check_filename(argument,
                   extension = "txt",
                   default = "im_a_file",
                   path = None):
    """
    =================================================================================================
    check_filename(argument, extension, default, path)
    
    This function is meant to check a filename for validity, given the kwargs.
    
    =================================================================================================
    Arguments:
    
    argument    ->  A string containing a file name (with no path information)
    extensions  ->  A string containing the desired extension
    default     ->  A string containing the default name, in the event something goes wrong
    path        ->  A path that the file should be appended to.
    
    =================================================================================================
    Returns: The filename with the extension, or the filename with the extension and the path.
    
    =================================================================================================
    """
    # If no path information is provided
    if path == None:
        # and the argument is None
        if argument == None:
            # Then just use the default file and extension
            return f"{default}.{extension}"
        # If the extension is already in the argument
        elif extension in argument:
            # Then just return the argument
            return argument
        # Otherwise, the extension is not in the argument
        else:
            # So split the argument and get the zeroeth value
            # (this works if no extension is present)
            arg = argument.split('.')[0]
            # and return arg with the extension
            return f"{arg}.{extension}"
    # If path information is provided
    elif path != None:
        # Then check the path for existence, and create
        # it if it does not
        path = check_dir(path,
                         create = True)
        # If the argument is not povided
        if argument == None:
            # Then just join the path with the default and the extension
            return os.path.join(path,f"{default}.{extension}")
        # If the extension is already in the argument
        elif extension in argument:
            # Then return the path joined to the argument
            return os.path.join(path,argument)
        # Otherwise
        else:
            # So split the argument and get the zeroeth value
            # (this works if no extension is present)
            arg = argument.split('.')[0]
            # and return the path joined to the argument and the extension.
            return os.path.join(path,f"{argument}.{extension}")


def check_type(argument,
               ob_type,
               error = "I'm a banana! I'm a banana! Look at me move YAYA"):
    """
    =================================================================================================
    check_type(argument, ob_type, error)
    
    This function is meant to check the type of an argument.
    
    =================================================================================================
    Arguments:
    
    argument  ->  Literally anything you want to check, as long as you know the expected type(s)
    ob_type   ->  An expected type, or a list of expected types
    error     ->  A string containing an error message, in the event something goes wrong.
    
    =================================================================================================
    Returns: The input argument after checking the validity of the type.
    
    =================================================================================================
    """
    # If multiple object types are provided
    if type(ob_type) in [list, tuple]:
        # Then see whether the argument type is in that list
        assert type(argument) in ob_type, error
    # Otherwise
    else:
        # See if the argument type is the preferred type
        assert type(argument) == ob_type, error
    # If all checks out, return the argument.
    return argument
    
#
#
######################################################################################################
```
