## Supporting Folder 2 Data Analysis for "Mouse primary T cell phosphotyrosine proteomics enabled by BOOST": general_helpers.py

```
"""
=================================================================================================
Kenneth P. Callahan

9 July 2021
                  
=================================================================================================
Python >= 3.8.5

general_helpers.py

This module contains functions that I use frequently, and that are generally helpful. They do
not belong to any specific script, and I tend to use them in many scripts.

I still need to add Type-checking to many of the functions here, but I will avoid that for
now. Currently, we need to be cautious about what the types of inputs are.

=================================================================================================
Dependencies:

os
sys
glob
copy

All of these modules are base Python modules. This file does not require any non-base modules.

=================================================================================================
"""
print(f"Loading the module: helpers.general_helpers\n")
######################################################################################################
#
#     Imporatables

# os and sys are used for operating system level
# operations
import os
import sys

# glob is used to iterate over the files in directories
import glob

# copy is used to create copies of input variables, as
# not to modify the original argument
import copy

#
#
######################################################################################################
#
# 

def remove_chars(string, remove = ["(", ")", "."] + [str(i) for i in range(10)]):
    newstr = ""
    for char in string:
        if char not in remove:
            newstr = f"{newstr}{char}"
    return newstr

#
#
######################################################################################################
#
#     Random Functions that I can't decide where they belong

def read_file(filename,
              delim = "\t"):
    """
    =================================================================================================
    read_file(filename, delim)
    
    This function is meant to read the lines of a file, split the lines on the delimiter, and
    return a list of lines
    
    =================================================================================================
    Arguments:
    
    filename  ->  A string containing the name of a file, and the path to the file if required.
    delim     ->  A string containing the character that splits elements of the file.
    
    =================================================================================================
    Returns: A list of lists
    
    =================================================================================================
    """
    # Open the file
    with open(filename, "r") as f:
        # and stip the lines, and split the lines on the delimiter
        lines = [line.rstrip("\n").split(delim) for line in f]
        # Close the file
        f.close()
    # and return the lines of the file
    return lines

def write_outfile(lines_list,
                  filename,
                  writestyle = "x"):
    """
    =================================================================================================
    write_outfile(lines_list, filename, writestyle)
    
    This function is meant to take a list of strings and a filename, then write the strings to the
    file.
    
    =================================================================================================
    Arguments:
    
    lines_list  ->  A list of strings, which will become the liens of the file
    filename    ->  A string containing the name of the output file, and the path if necessary
    writestyle  ->  A string determining how the file should be written. Look at the python
                    function 'open()' for more details.
    
    =================================================================================================
    Returns: None
    
    =================================================================================================
    """
    # This is a basic scheme for writing files. Open the file with writestyle,
    # use the writelines method of files to write the lines, and clsoe the file.
    with open(filename, writestyle) as f:
        f.writelines(lines_list)
        f.close()
    return None
        
def transpose(*args,
              errors = True):
    """
    =================================================================================================
    transpose(*args, errors)
    
    Given an arbitarary number of lists, flip the row and column space of those lists.
    
    =================================================================================================
    Arguments:
    
    *args   ->  An arbitrary number of lists
    errors  ->  A boolean, determines whether or not to check arguments
    
    =================================================================================================
    Returns: A list of lists, where the rows of the list are the columns of the input list.
    
    =================================================================================================
    Given a matrix A_{m,n}, return the matrix A^{T}_{n,m}, where
    the row space and the column space are exchanged:

    Example:

    a = [[1,2,3],[1,2,3]]

    print(transpose(a))

    -> [[1,1],[2,2],[3,3]]

    NOTE: This function did not end up in the final script, but I left
    it just in case it is needed in the future.
    """
    # Make sure the input arguments are all lists/tuples. We need iterables
    # for the list comprehension later to work.
    if errors:
        for item in args:
            assert type(item) in [list, tuple], "The inputs should either be lists or tuples of the same length"
    # Make sure that the size of all input lists are equivalent
    lengths = [len(item) for item in args]
    lengths = [lengths[0] == item for item in lengths]
    #
    if errors:
        assert all(lengths), "The input lists/tuples do not have the same dimensions..."
    # Use the zip() function to return all column-tuples
    a_transpose = list(zip(*args))
    # And turn the tuples into lists, as we often want them to be mutable
    return [list(item) for item in a_transpose]

def remove_dict_key(a_dictionary,
                    val = None,
                    dkey = None):
    """
    =================================================================================================
    remove_dict_key(a_dictionary, val, dkey)
    
    This function is meant to take either a value or a key that is in the input dictionary, and
    create a new dictionary either without that key, or without the key associated with the value.
    
    =================================================================================================
    Arguments:
    
    a_dictionary  ->  A dictionary
    val           ->  A value that should be in the dictionary, that you would like to remove
    dkey          ->  A key that should be in the dictionary, that you would like to remove
    
    NOTE: Only one of the two arguments <val> and <dkey> can be set at one time.
    
    =================================================================================================
    Returns: a dictionary with one key/value pair removed.
    
    =================================================================================================
    """
    # Check to make sure that only one of the two kwargs is set
    assert val != None or dkey != None, "Either the val or dkey argument must be set..."
    # Initialize the new_dict variable to hold the ouptput
    new_dict = {}
    # Loop over the keys and values of the input dictioanry
    for key, value in a_dictionary.items():
        # If the val is set, and the current value is
        # the same as the val
        if val != None and val == value:
            # Then pass, as this must be removed from the dictionary
            pass
        # Or if the dkey is set and the current key is
        # the same as the dkey
        elif dkey != None and dkey == key:
            # Then pass, this must be removed from the dictioary
            pass
        # Otherwise,
        else:
            # Add the key/value pair to the dictionary
            new_dict[key] = value
    # and return the dictionary once the filtering is complete
    return new_dict

def remove_all_keys(a_dictionary,
                    vals = [],
                    dkeys = []):
    """
    =================================================================================================
    remove_all_keys(a_dictionary, vals, dkeys)
    
    This function wraps <remove_dict_key> for all vals and all dkeys provided as input. For more
    information, refer to remove_dict_key()
    
    =================================================================================================
    Arguments:
    
    a_dictionary  ->  A dictionary
    vals          ->  A list of values that should be in the dictionary, that you want to remove
    dkeys         ->  A list of keys that should be in the dictioanry, that you want to remove
    
    =================================================================================================
    Returns: a new dictionary that contains all elements of the input dictionary, without the
             elements specified from vals and dkeys
    
    =================================================================================================
    """
    # Make a copy of the input dictionary, as we do not want to modify it
    newdict = copy.copy(a_dictionary)
    # Loop over the items in vals
    for item in vals:
        # and run remove_dict_key on each of the values
        newdict = remove_dict_key(newdict,
                                  val = item)
    # Loop over the items in dkeys
    for item in dkeys:
        # and run remove_dict_key on each of the keys
        newdict = remove_dict_key(newdict,
                                       dkey = item)
    # At the end, return the dictionary which is filtered.
    return newdict

#
#
######################################################################################################
#
#       Counting Functions

def count(a_list):
    """
    =================================================================================================
    count(a_list)
    
    This function takes a list as input and counts the occurrences of each element of the list.
    
    =================================================================================================
    Arguments:
    
    a_list  ->  A list with any type of items in it. The items in the dictionary will become the
                keys of the count dictionary.
    
    =================================================================================================
    Returns: A dictionary where the keys are the elements of a_list and the values are the number
             of occurrences of that element in the list.
    
    =================================================================================================
    """
    # Initialize the counts dictionary
    counts = {}
    # and loop over the elements of a_lsit
    for item in a_list:
        # if the string of the item is not already a key in the list
        if str(item) not in list(counts.keys()):
            # then initialize a key in the counts dictionary with value 1
            counts[str(item)] = 1
        # Otherwise, the string of the item is already a key
        else:
            # so increase the count by one
            counts[str(item)] += 1
    # and return the coutns dictionary after counting is complete.
    return counts

def merge_dicts(*dicts,
                filler = 0):
    """
    =================================================================================================
    merge_dicts(*dicts, filler)
    
    This function takes an arbitrary number of dictionaries and merges them together. If a key is not
    found in one of the dictionaries, the filler value is introduced.
    
    =================================================================================================
    Arguments:
    
    dicts   ->  An arbitrary number of dictionaries
    filler  ->  The value used as filler when one dictionary does not have the same key as another.
    
    =================================================================================================
    Returns: A dictionary of lists, where the keys are the keys from all dictionaries used as inputs
             and the values are lists of values, with filler values where values were not identified.
    
    =================================================================================================
    """
    # Use dictionary comprehension to initialize the new_dict
    new_dict = {key : [value] for key, value in dicts[0].items()}
    # Loop over the number of dictionaries, minus the zeroeth dictionary.
    for i in range(1,len(dicts)):
        # Loop over the keys and values in the ith dictionary
        for key, value in dicts[i].items():
            # If the current key is not already a key in the dictionary
            if str(key) not in new_dict:
                # Then make that a key, with value as a list containing i
                # filler values and the current dict value
                new_dict[key] = [filler for _ in range(i)] + [value]
            # If the key is already in the dict and there are the same
            # number of values as the index
            elif str(key) in new_dict and len(new_dict[key]) == i:
                # Then simply add the value to the dictionary, as
                # there are enough values in the current list
                new_dict[key].append(value)
            # Otherwise, we need to include a filler value along
            # with the value
            else:
                # so upate the lsit with a filer and the value
                new_dict[key] = [*new_dict[key], filler, value]
        # Once each dictionary is completeed,
        for key, value in new_dict.items():
            # loop over the newdict and see which items were not
            # updated
            if len(value) < i+1:
                # and add filler to the newdict, to ensure they stay
                # the same size as each list.
                new_dict[key] = new_dict[key] + [filler]
    # Once the merger is complete, return the newdict
    return new_dict

def count_lists(*lists):
    """
    =================================================================================================
    count_lists(*lists)
    
    =================================================================================================
    Arguments:
    
    lists  ->  An arbitrary number of lists
    
    =================================================================================================
    Returns: A list of dictionaries representing the counts for each input list.
    
    =================================================================================================
    """
    return [count(a_list) for a_list in lists]

#
#
######################################################################################################
#
#       Generators

def bipartite_pairs(iterable_1,
                    iterable_2, ret_type = tuple):
    """
    =================================================================================================
    bipartite_pairs(iterable_1, iterable_2, ret_type)
    
    This function is meant to make lists/tuples decribing the edges between two groups of nodes. For
    example,
    
    iterable_1 = [1,--\----/--[a,
                  2,---\--/----b,
                  3]----\/-----c] = iterable_2
                  
         output -> [[1,a],[1,b],[1,c],
                    [2,a],[2,b],[2,c],
                    [3,a],[3,b],[3,c]]
                  
    =================================================================================================
    Arguments:
    
    iterable_1  ->  An iterable
    iterable_2  ->  An iterable
    ret_type    ->  An iterable type, that the output will be returned as
    
    =================================================================================================
    Yields: A list of tuples(or lists) that define the edges between two groups.
    
    =================================================================================================
    """
    # Loop over the set of nodes in group 1
    for node1 in iterable_1:
        # Initialize a new group list
        new_group = []
        # Loop over the nodes in group 2
        for node2 in iterable_2:
            # and add the pair to the new group list.
            new_group.append(ret_type((node1,node2)))
        # Once this group of nodes has been made, yield this group.
        yield new_group

def unique_combinations(an_iterable):
    """
    =================================================================================================
    unique_combinations(an_iterable)
    
    This function is meant to take an iterable, and yeilds the unique elements of the iterable.
    The name comes from the use in filtering out forwards and backwards combinations.
    
    =================================================================================================
    Arguments:
    
    an_iterable  ->  An iterable object
    
    =================================================================================================
    Yields: Unique items from the iterable input
    
    =================================================================================================
    """
    # Initialize a list to hold the items that are seen.
    seen = []
    # Loop over the items in the iterable
    for item in an_iterable:
        # If the item is not already seen
        if item not in seen:
            # Then add the item to the seen list
            seen.append(item)
            #if type(item) in [list,tuple]:
            #    seen.append(*list(make_pairs(item, dupes = True)))
            # and yield the item
            yield item
            
def make_pairs(an_iterable,
               dupes = False,
               reverse = False):
    """
    =================================================================================================
    make_pairs(an_iterable, dupes, reverse)
    
    This function is meant to take an iterable and yield all mathematical combinations of pairs in
    the iterable.
    
    =================================================================================================
    Arguments:
    
    an_iterable  ->  An iterable
    dupes        ->  A boolean determining how to handle duplicates
    reverse      ->  A boolean determining whether to take the forward or backwards pairs
    
    =================================================================================================
    Yields: A two-list, containing pairs from the input list.
    
    =================================================================================================
    """
    # Initialize a list for the seen two-lists
    seen = []
    # Loop over the items in the iterable
    for item1 in an_iterable:
        # Loop over the items in the iterable again
        for item2 in an_iterable:
            # If the two items are equivalent, and the dupes argument
            # is True, and the current combination is not already seen.
            if item1 == item2 and dupes and [item1, item2] not in seen:
                # Then add the combination to the seen list
                seen.append([item1,item2])
                # and yield the current combination
                yield [item1, item2]
            # Or if the two items are not equivalent and
            # the current combination is not already seen
            elif item1 != item2 and [item1, item2] not in seen:
                # Then add both the forward and backwards combination
                # to the list
                seen.append([item1,item2])
                seen.append([item2,item1])
                # and if the reverse argument is True
                if reverse:
                    # Then yield the reverse combination
                    yield [item2, item1]
                # Otherwise
                else:
                    # yield the forward combination
                    yield [item1,item2]

#
#
######################################################################################################
#
#      Functions relating to lists

def unpack_list(a_list):
    """
    =================================================================================================
    unpack_list(a_list)
    
    This is a recursive function which takes a list of lists of... and returns a single list
    containing all elements of the input lists.
    
    =================================================================================================
    Arguments:
    
    a_list  ->  A list of an arbitrary number of sublists
    
    =================================================================================================
    Returns: A list containing all elements of all sublists.
    
    =================================================================================================
    """
    # Initialize the output list
    outlist = []
    # loop over the elements in the input list
    for element in a_list:
        # If the element is a list or a tuple
        if type(element) in [list, tuple]:
            # then use unpack_list() to unpack the
            # element and add that value to theoutlist
            outlist += unpack_list(element)
        # Otherwise,
        else:
            # Add the element to the list
            outlist.append(element)
    # and return the output list once the loop finishes
    return outlist

def remove_list_indices(a_list,
                        del_indices,
                        opposite = True):
    """
    =================================================================================================
    remove_list_indices(a_list, del_indices, opposite)
    
    This function is meant to remove elements from a list, based on the deletion indices and the
    value of opposite.
    
    =================================================================================================
    Arguments:
    
    a_list       ->  A list
    del_indices  ->  A list of integers that define either which indices to delete, or which to keep
    opposite     ->  A boolean, determines whether to delete all other indices or all indices in
                     del_indices
    
    =================================================================================================
    Returns: A list filtered by the commands specified above
    
    =================================================================================================
    Example:

    a = [1,2,3,4]
    del_indices = [0,2]

    print(remove_list_indices(a,del_indices))

    -> [2,4]

    print(remove_list_indices(a,del_indices, opposite = False))

    -> [1,3]
    =================================================================================================
    """
    # Make sure the user inputs the proper typed objects.
    # Opposite must be a boolean
    assert opposite in [True, False], "opposite argument must be a boolean"
    # del_indices must be an iterable, specifically a list/tuple
    assert type(del_indices) in [list, tuple], "Deletion indices should be given as a list or tuple of integers"
    # All of the values in del_indices must be integers
    assert all([type(ind) == int for ind in del_indices]), "Deletion indices should be given as a list or tuple of integers"
    # a_list must be an interable, specifically a list/tuple
    assert type(a_list) in [list, tuple], "The argument 'a_list' should be a list or a tuple."
    # The a_list must include all indices required for deletion.
    assert sorted(del_indices)[-1] <= len(a_list), "The deletion indices given expand beyond the size of the list."

    # If the argument opposite is True
    if opposite:
        # Then return a list where all elements NOT IN del_indices are kept
        return [a_list[i] for i in range(len(a_list)) if i not in del_indices]
    # If the argument opposite is False
    else:
        # Then return a list where all elements IN del_indices are kept
        return [a_list[i] for i in range(len(a_list)) if i in del_indices]

def list_to_str(a_list,
                delimiter = "\t",
                newline = True):

    """
    =================================================================================================
    list_to_str(a_list, delimiter, newline)
    
    Given a list, the delimiter (default '\t'), and whether to add a trailing newline
    character, take a list and convert the list into a string where each element is
    separated by the chosen delimiter.
    
    =================================================================================================
    Arguments:
    
    a_list     ->  A list
    delimiter  ->  A string to separate the elements of the list in a string
    newline    ->  A boolean that determines whether to add a newline character to the output string
    
    =================================================================================================
    Returns: A string containing the elements of a_list, separated by delimtmer, with or without
             a newline character
    
    =================================================================================================
    Example:

    a = [1,2,3]
    print(list_to_str(a))

    -> '1\t2\t3\n'
    =================================================================================================
    """
    # Make sure the user inputs the proper typed objects.
    # newline argument needs to be a boolean
    assert newline in [True, False], "newline argument must be a boolean"
    # a_list argument needs to be an iterable, specifically a list/tuple
    assert type(a_list) in [list, tuple], "The argument 'a_list' should be a list or a tuple."
    # These are the delimiter characters that I am currently able to work with
    #assert delimiter in [':', '|', ';', '-', '\\',
    #                     '/', ',', '\t', '\n', "",
    #                     " vs "], f"The delimiter provided is not an accepted delimiter."

    # Initialize the new string with the first element of the list.
    # This avoids having to slice the strings later
    newstr = f"{a_list[0]}"
    # If the list only has one element and the user elects to use
    # a trailing newline character
    if len(a_list) == 1 and newline:
        # Then simply return the newstr variable with an added
        # newline character
        return f"{newstr}\n"
    # If the list has only one element and the user does not elect to
    # use a trailing newline character
    elif len(a_list) == 1 and not newline:
        # Then simply return the newstr variable
        return f"{newstr}"

    # If the list has more then one element, then loop over all elements
    # (excluding the first one since that is already in the string)
    for i in range(1,len(a_list)):
        # and add those new elements to the newstring with the given
        # delimiter separating the elements.
        newstr = f"{newstr}{delimiter}{a_list[i]}"
    # If the user elects to use a trailing newline character
    if newline:
        # Then add the trailing newline character and return the string
        return f"{newstr}\n"
    # If the user does not elect to use a trailing newline character
    else:
        # Then simply return the newstr variable
        return newstr

def filter_matrix(a_matrix, header, value, keep, head_loc = 0, compare = "none"):
    """
    """
    # Get the header
    headers = a_matrix[head_loc]
    new_matrix = [item for item in a_matrix if item != headers]
    head_pos = headers.index(header)
    if keep and compare == "none":
        return [headers] + list(filter(lambda x: x[head_pos] == value,new_matrix))
    elif keep and compare == ">":
        return [headers] + list(filter(lambda x: x[head_pos] > value,new_matrix))
    elif keep and compare == ">=":
        return [headers] + list(filter(lambda x: x[head_pos] >= value,new_matrix))
    elif keep and compare == "<":
        return [headers] + list(filter(lambda x: x[head_pos] < value,new_matrix))
    elif keep and compare == "<=":
        return [headers] + list(filter(lambda x: x[head_pos] <= value,new_matrix))
    elif keep and compare == "in":
        return [headers] + list(filter(lambda x: x[head_pos] in value,new_matrix))
    elif keep and compare == "not in":
        return [headers] + list(filter(lambda x: x[head_pos] not in value,new_matrix))
    elif not keep and compare == ">":
        return [headers] + list(filter(lambda x: x[head_pos] <= value,new_matrix))
    elif not keep and compare == ">=":
        return [headers] + list(filter(lambda x: x[head_pos] < value,new_matrix))
    elif not keep and compare == "<":
        return [headers] + list(filter(lambda x: x[head_pos] >= value,new_matrix))
    elif not keep and compare == "<=":
        return [headers] + list(filter(lambda x: x[head_pos] > value,new_matrix))
    elif not keep and compare == "in":
        return [headers] + list(filter(lambda x: x[head_pos] not in value,new_matrix))
    elif not keep and compare == "not in":
        return [headers] + list(filter(lambda x: x[head_pos] in value,new_matrix))
    else:
        return [headers] + list(filter(lambda x: x[head_pos] != value,new_matrix))

def select_cols(a_matrix, head_dict, head_loc = 0):
    """
    """
    headers = a_matrix[head_loc]
    head_inds = [headers.index(head) for head in list(head_dict.keys())]
    matrix = transpose(*a_matrix)
    new_matrix = []
    for i in head_inds:
        new_matrix.append([head_dict[headers[i]]] + [row for row in matrix[i] if row != headers[i]])
    return transpose(*new_matrix)

def replace_value(a_list, value, newvalue, transform = None):
    """
    """
    new_list = []
    for item in a_list:
        if item == value:
            new_list.append(newvalue)
        else:
            new_list.append(item)
    return new_list

def transform_values(a_list, transform = float):
    new_list = []
    for item in a_list:
        try:
            new_list.append(float(item))
        except:
            new_list.append(item)
    return new_list

def grab_col(a_matrix, head, head_loc = 0):
    head_ind = a_matrix[head_loc].index(head)
    return [head] + [row[head_ind] for row in a_matrix if row[head_ind] != head]
    
def add_col(a_matrix, newhead, newval = [],
            head_loc = 0, newhead_pos = 0):
    if newval != []:
        assert len(newval) == len(a_matrix)-1, f"The new column should have length {len(a_matrix)-1}"
    else:
        newval = ["" for _ in range(len(a_matrix)-1)]
    if newhead_pos == -1:
        heads = a_matrix[head_loc] + [newhead]
    else:
        heads = a_matrix[head_loc][:newhead_pos+1] + [newhead] + a_matrix[head_loc][newhead_pos:]
    newmatrix = [heads]
    nohead = [row for row in a_matrix if a_matrix.index(row) != head_loc]
    for i in range(len(nohead)):
        if newhead_pos == -1:
            newrow = nohead[i] + [newval[i]]
        else:
            newrow = nohead[i][:newhead_pos+1] + [newval[i]] + nohead[i][newhead_pos:]
        newmatrix.append(newrow)
    return newmatrix

#
#
######################################################################################################
#
#  Functions relating to dictionaries

def bin_by_col(a_matrix, id_col, head_row = 0):
    """
    Given a matrix and the id column index, return a number of
    matrices where the id column is a single value
    """
    headers = a_matrix[head_row]
    parsed = {}
    i = 0
    for item in a_matrix:
        if item == headers:
            continue
        elif item[id_col] not in parsed.keys():
            parsed[item[id_col]] = [item]
        else:
            parsed[item[id_col]].append(item)
        i+=1
    return {key : [headers] + value for key, value in parsed.items()}

#
#
######################################################################################################
#
#      Path based stuff


def get_file_list(directory,
                  filename,
                  true_file = True):
    """
    =================================================================================================
    get_file_list(directory, filename, true_file)
    
    Given a directory, a filename, and a true_file boolean, recursively find all files in the input
    directory with the given file name (or string)
    
    =================================================================================================
    Arguments:
    
    directory  ->  A string containing the directory path in which to find files
    filename   ->  A string containing the name of the file to look for (or the string to look for
                   in a file)
    true_file  ->  A boolean determining whether the filename is a True file name, or whether 
                   it is a substring that all desired files have.
    
    =================================================================================================
    Returns: A list of all files with the given filename in the input directory
    
    =================================================================================================
    """
    # First, add a star to the directory path, so we may use
    # glob to get a list of all paths in the directory
    dirpaths = glob.glob(os.path.join(directory, "*"))
    # If the filename is an element of the paths found using glob
    if os.path.join(directory,filename) in dirpaths and true_file:
        # Then get the absolute path of the file and return it
        file = os.path.abspath(os.path.join(directory,filename))
        return file
    # Or, if we are looking for a substring and not a file name,
    # check to see if any of the paths in dirpaths contain the substring
    elif not true_file and any([True for path in dirpaths if filename in path]):
        # If so, get a list of all paths with that substring 
        files = [path for path in dirpaths if filename in path]
        # and return that list of file paths
        return files
    # Otherwise, there are no paths with the filename
    else:
        # So get a list of all paths that are directories
        dirpaths = [path for path in dirpaths if os.path.isdir(path)]
        # and use get_file_list() recursively to find files with
        # the given file name.
        files = [get_file_list(path,
                               filename = filename,
                               true_file = true_file) for path in dirpaths]
        # At the end, unpack the lsit of files
        return unpack_list(files)
    
    
#
#
######################################################################################################
```
