## Supporting Folder 2 Data Analysis for "Mouse primary T cell phosphotyrosine proteomics enabled by BOOST": mpl_plotting_helpers.py

```
"""
=================================================================================================
Kenneth P. Callahan

17 July 2021
                  
=================================================================================================
Python >= 3.8.5

mpl_plotting_helpers.py

This is a module meant to help with various types of plots. While Python has a ton of support
for plotting in MatPlotLib, some of the things aren't exactly intuitive. Many of these functions
are meant to help with plotting, wrapping MatPlotLib functions and performing formatting.

Currently type checking is not being done.

=================================================================================================
Dependencies:

   PACKAGES        VERSION
    matplotlib  ->  3.3.2
    numpy       ->  1.19.2

If the printed versions ever change, then change the versions here. These are the ones
in my current Python installation.
=================================================================================================
"""
print(f"Loading the module: helpers.mpl_plotting_helpers\n")
############################################################################################################
#
#     Importables

# Import base matplotlib to show the version.
import matplotlib

# Pyplot has the majority of the matplotlib plotting functions
# and classes, including figures, axes, etc.
import matplotlib.pyplot as plt
#    cm is used for colourmaps
import matplotlib.cm as cm
#    ticker is used for placing labels on heatmaps
import matplotlib.ticker as ticker

from matplotlib import gridspec
from matplotlib.patches import Patch

# Numpy is used for creating arrays that matplotlib is able to handle
import numpy as np

# Copy is used for making copies of objects, rather than manipulating
# the original form of an object.
import copy

# Random is used for making pseudorandom selections from a list of items.
import random

# General helpers has a number of functions I use frequently in
# my scripts. They are all placed in that module purely for
# convenience and generalizability.
from . import general_helpers as gh

# The Pandas Helper file has scripts that help manage Pandas
# DataFrames, and perform various actions on lists of dataframes
from . import pandas_helpers as ph

from . import stats_helpers as sh

print(f"matplotlib    {matplotlib.__version__}")
print(f"numpy         {np.__version__}\n")

plt.rcParams.update(plt.rcParamsDefault)
plt.rcParams['font.family'] = "sans-serif"

#
#
############################################################################################################
#
#     Global Variables

# This dictionary has some predefined sets of colours that can be used for
# barplot plotting. These colours should all be base matplotlib colours,
# so I can't imagine they'll be depricated soon.
colours = {"blues"  : ["steelblue", "cyan", "blue", "darkblue", "dodgerblue",
                       "lightblue", "deepskyblue"],
           "pinks"  : [ "mediumvioletred", "darkmagenta", "deeppink","violet", "magenta",
                       "pink", "lavenderblush"],
           "reds"   : ["darkred", "firebrick", "indianred", "red", "tomato", "salmon",
                      "lightcoral", "darksalmon", "mistyrose"],
           "purples": ["indigo", "mediumpurple", "purple",
                       "darkviolet", "mediumorchid", "plum", "thistle"],
           "greens" : ["darkolivegreen", "olivedrab", "green", "forestgreen",
                       "limegreen", "springgreen", "lawngreen"],
           "oranges": ["darkorange", "orange", "goldenrod", "gold", "yellow",
                       "khaki", "lightyellow"],
           "browns" : ["brown","saddlebrown", "sienna", "chocolate", "peru",
                      "sandybrown", "burlywood"],
           "monos"  : ["black", "dimgrey", "grey", "darkgrey",
                       "silver", "lightgrey", "gainsboro"],
           "cc"     : ["darkturquoise", "cyan", "thistle", "fuchsia", "violet"],
           "default": ["blue", "red", "green", "purple", "pink"],
           "all"    : ["darkblue", "steelblue", "blue", "dodgerblue", "deepskyblue",
                      "aqua", "lightblue", "cornflowerblue","darkmagenta", "mediumvioletred", 
                       "deeppink", "violet", "magenta", "pink", "lavenderblush",
                       "darkred", "firebrick", "indianred", "red", "tomato", "salmon",
                      "lightcoral", "darksalmon", "mistyrose",
                       "indigo", "rebeccapurple", "mediumpurple", "purple",
                       "darkviolet", "mediumorchid", "plum", "thistle",
                       "darkolivegreen", "olivedrab", "green", "forestgreen",
                       "limegreen", "springgreen", "lawngreen", "palegreen",
                       "darkorange", "orange", "goldenrod", "gold", "yellow",
                       "khaki", "lightyellow",
                       "brown","saddlebrown", "sienna", "chocolate", "peru",
                      "sandybrown", "burlywood", "wheat",
                       "black", "dimgrey", "grey", "darkgrey",
                       "silver", "lightgrey", "gainsboro", "white"]}

tab_colours = ["tab:blue", "tab:orange", "tab:green", "tab:red",
               "tab:purple", "tab:brown", "tab:pink", "tab:gray", "tab:olive", "tab:cyan"]

#
#
############################################################################################################
#
#     Functions: Bar Plots

def bars_ind(n_items,
             n_groups,
             sep_dist = 0.4,
             transpos = True):
    """
    =================================================================================================
    bars_ind(n_items, n_groups, sep_dist, transpos)
    
    This funciton is used to determine the center positions of all bars used for a bar plot, as
    well as the width of each bar.
    
    =================================================================================================
    Arguments:
    
    n_items   ->  An integer representing the number of categories to be plotted
    n_groups  ->  An integer representing the number of groups per category to be plotted
    sep_dist  ->  A float representing the distance between categories in the plot
    transpos  ->  A boolean determining whether to transpose the output list.
    
    =================================================================================================
    Returns: A list of lists determining the position of bars for each group, and the width
             of all bars in the plot.
    
    =================================================================================================
    """
    # Determine the width of a bar, by subtracting the separation distance from 1
    # and dividing by the number of groups.
    width = (1-sep_dist) / n_groups
    # If there is only one group,
    if n_groups == 1:
        # Then return a list with one sublist, centered
        # on the integers.
        return [[i for i in range(len(n_groups))]]
    # Initialize the output list. This will hold the lists returned.
    output = []
    # Next, loop over the number of categories
    for i in range(n_items):
        # Get the left most position of the interval
        interval = i + sep_dist/2 - 0.5
        # Add equidistant points from the interval to the output
        # Up to the number of groups per category and add that
        # to the outputs
        output.append([interval + j * width for j in range(1,n_groups+1)])
    # If the output must be transposed
    if transpos:
        # Then return the transposed output list and width
        return gh.transpose(*output), width
    # Otherwise,
    else:
        # Return the list itself and the width
        return output, width
    
def handle_colours(colour_type,
                   n_groups,
                   choice = "centered"):
    """
    =================================================================================================
    handle_colours(colout_type, n_groups, choice)
    
    This function is meant to get a list colours that will define the bars on the barplot. It uses
    the colours dictionary defined globally and the keys as the colour_type.
    
    =================================================================================================
    Arguments:
    
    colour_type  ->  A string that is in the keys of the colours dictionary. If a string not in
                     the colours dictionary is provided, the function will use either
                     'default' or 'all', depending on the size of n_groups.
    n_groups     ->  An integer representing the number of groups within a category.
    choice       ->  A string representing how to choose the colours from a list.
                     "centered" "random"
    
    =================================================================================================
    Returns: A list of colour strings of size n_groups.
    
    =================================================================================================
    """
    # Use the global colours dictionary for colour determination.
    global colours
    # First, check to see if the colour_type given is in the keys of the dictionary.
    if f"{colour_type}" in list(colours.keys()):
        # Next, check to see whether there are enough colours to support choice.
        #
        # If there are enough colours to support choices
        if n_groups <= len(colours[f'{colour_type}']):
            # Then check the choice string.
            #
            # If the choice string is random
            if choice == "random":
                # Then choose a random sample from the colours list, of size n_groups
                return random.sample(colours[f"{colour_type}"], n_groups)
            # If the choice string is centered
            elif choice == "centered":
                # Then get the center of the list. For even lists, this
                # will be the right-center.
                ind_cent = len(colours[f"{colour_type}"]) // 2
                # If the number of groups and the number of colours are equivalent
                if n_groups == len(colours[f"{colour_type}"]):
                    # Then return the list of colours
                    return colours[f"{colour_type}"]
                # If there is an even number of groups
                elif n_groups %2 == 0:
                    # Then get a slice object that is evenly spaced about
                    # the center of the list.
                    x = slice(ind_cent - n_groups//2, ind_cent + n_groups //2)
                    # Then attempt to return the colours list, sliced by x
                    try:
                        return colours[f'{colour_type}'][x]
                    # And if that fails, then return a random sample from the list
                    except:
                        return random.sample(colours[f'{colour_type}'], n_groups)
                # If there is an odd number of groups
                else:
                    # Then get a slice object that is left centered.
                    x = slice(ind_cent - 1 - n_groups //2, ind_cent - n_groups//2 + 2)
                    # and try to return the colours sliced at that point
                    try:
                        return colours[f'{colour_type}'][x]
                    # Otherwise return a random sample of the colours from that list.
                    except:
                        return random.sample(colours[f'{colour_type}'], n_groups)
            # If something other than centered or random is chosen
            else:
                # Then return a random sample from the colour
                return random.sample(colours[f"{colour_type}"])
        # If there are not enough colours in the chosen list,
        else:
            # Then return a random sample from all
            return random.sample(colours['all'], n_groups)
    # If the colour type is not in the keys and the number of groups is
    # less than or equal to the default list
    elif colour_type not in list(colours.keys()) and n_groups <= len(colours["default"]):
        # Then return a slice of the default list
        return colours["default"][:n_groups]
    # Otherwise, all things in the world are wrong
    else:
        # So just return a random sample from the all list of size n_groups.
        return random.sample(colours['all'], n_groups)

def bars(xvals,
         yval_matrix,
         col_labels = None,
         separation = 0.3,
         colour_type = "all",
         colour_choice = "centered",
         img_type = "pdf",
         img_name = "pokedex_completion",
         show = True,
         subplot_args = {"figsize" : (24,12)},
         set_kwargs = {"xlabel" : "Ash Ketchum",
                       "ylabel" : "Pokemon Caught",
                       "title"  : "I wanna be, the very best, like NO ONE EVER WAS"}):
    """
    =================================================================================================
    bars(*args, **kwargs)
    
    This function is meant to wrap axes.bars() and provide some of the formatting options for
    barplots
    
    =================================================================================================
    Arguments:
    
    xvals          ->  A list of strings that determine the categories of the bar chart. These values
                       and the values in each sublist of yval_matrix should be index paired.
    yval_matrix    ->  A list of lists of values, where each sublist represents the values for a
                       a specific group.
    col_labels     ->  A list of labels for the bars. These values should be index paired with the
                       yval_matrix list.
    separation     ->  A float (0<separation<1) that determines how much space is between two
                       categories
    colour_type    ->  A string that represents the colour group to use for the bars
    colour_choice  ->  A string that represents how to choose the colours from the colour list.
    img_type       ->  A string representing the file extension for the image.
    img_name       ->  A string representing the name of the file.
    show           ->  A boolean that determines whether or not to show the plot.
    subplot_args   ->  A dictionary of keyword arguments to be passed into plt.subplots()
    set_kwargs     ->  A dictionary of keyword arguments to be passed into ax.set()
    
    =================================================================================================
    Returns: None, but a figure is saved.
    
    =================================================================================================
    """
    # Set the global font size parameter to 20.
    plt.rcParams["font.size"] = 20
    # Get the indices list (of lists) and the width of the bars
    indices, width = bars_ind(len(xvals),
                              len(yval_matrix),
                              sep_dist = separation)
    # Get the colour list that wil be used for barplots
    color = handle_colours(colour_type,
                           len(yval_matrix),
                           choice = colour_choice)
    # With the indices and colours list, we are set to start plotting.
    # Create a figure and axes object using plt.subplots().
    fig, ax = plt.subplots(**subplot_args)
    # Then, loop over the number of indices lists created by
    # bars_ind()
    for i in range(len(indices)):
        # If column labels are provided, then we assume they are
        # index paired with the lists in yval_matrix and thus
        # the indices list.
        if col_labels != None:
            # So plot a bar to the axes for this combination,
            # with a colour.
            ax.bar(indices[i], yval_matrix[i],
                   width = width, label = col_labels[i],
                   edgecolor = "black", color = color[i])
        # If no labels are provided,
        else:
            # Then plot a bar all the same, but without a label.
            ax.bar(indices[i], yval_matrix[i],
                   width = width, label = col_labels[i], color = color[i])
    # Once all of the bars are plotted, plot a legend.
    ax.legend()
    # If the number of indices lists is even
    if len(indices) % 2 == 0:
        # Then calculate the position of the labels based on an even assumption.
        ax.set_xticks(indices[len(indices)//2 + 1])
        ax.set_xticklabels(xvals,rotation = 45, ha = 'right', rotation_mode = "anchor")
    # If the number of indices lists is odd
    elif len(indices) % 2 == 1:
        # Then calculate the position of the labels based on an odd assumption.
        ax.set_xticks([item + width/2 for item in indices[len(indices)//2]])
        ax.set_xticklabels(xvals,rotation = 45, ha = 'right', rotation_mode = "anchor")
    # If set_kwargs is not en empty dictioanry
    if set_kwargs != {}:
        # Then run ax.set{} using that dictionary. This will fail if the
        # arguments are not arguments in the set() method.
        ax.set(**set_kwargs)
    # Save the figure using the image name and image type
    if show:
        plt.savefig(f"{img_name}.{img_type}", bbox_inches = "tight")
    # If show is True
    return ax

#
#
############################################################################################################
#
#      Functions: Heatmaps

def get_range(a_list):
    """
    =================================================================================================
    get_range(a_list)
    
    This is meant to find the maximal span of a list of values.
    
    =================================================================================================
    Arguments:
    
    a_list  ->  A list of floats/ints.  [1,2,-3]
    
    =================================================================================================
    Returns: a tuple of the values that are either at the end/beginning of the list. (-3,3)

    =================================================================================================
    """
    # Make sure the input list is correctly formatted
    assert type(a_list) == list, "The input a_list should be of type list..."
    # First, unpack the list of lists. This makes one list with all values from
    # the lists within the input list.
    #print(a_list)
    unpacked = gh.unpack_list(a_list)
    #print(unpacked)
    # Next, float the items in the unpacked list. This will fail if any
    # strings that are not floatable are in the list.
    unpacked = [float(item) for item in unpacked if float(item) == float(item)]
    # Then we can get the max and min of the list.
    maxi = max(unpacked)
    mini = min(unpacked)
    # If the max value is greater than or equal to the minimum value
    if abs(maxi) >= abs(mini):
        # Then the bound is the int of max plus 1
        bound = int(abs(maxi)) + 1
        # We can then return the bounds, plus and minus bound
        return (-bound, bound)
    # If the min value is greater than the max value
    elif abs(maxi) < abs(mini):
        # Then the bound is the int of the absolute value
        # of mini plus 1
        bound = int(abs(mini)) + 1
        # We can then return the bounds, plus and minus bound
        return (-bound, bound)
    # If something goes wrong,
    else:
        # Then raise an error
        raise ValueError("This is an unexpected outcome...")

def infer_aspect_ratio(xlabels,
                       ylabels):
    """
    =================================================================================================
    infer_aspect_ratio(xlabels, ylabels)
    
    This function is meant to determine which aspect ratio to use when making a heatmap.
    
    =================================================================================================
    Arguments:
    
    xlabels  ->  a list of labels for the x axis of a heatmap
    ylabels  ->  a list of labels for the y axis of a heatmap
    
    =================================================================================================
    Returns: the string "auto" or "equal"
    
    =================================================================================================
    """
    # Check the types of the arguments
    assert type(xlabels) == list, "The xlabels should be of type <list>"
    assert type(ylabels) == list, "The ylabels should be of type <list>"
    # Get the length of the xlabels and the ylabels
    x = len(xlabels)
    y = len(ylabels)
    # Based on the lengths of the x and y labels, choose an aspect ratio
    #
    # If there are less than or equal to five y labels
    if y <= 5:
        # Then just set the aspect ratio to equal
        return "equal"
    # Or if there are more than 5 y labels and less than 5 xlabels
    elif y > 5 and x <= 5:
        # Then set the aspect ratio to auto
        return "auto"
    # Or if both x and y labels are longer than 5
    elif y >5 and x > 5:
        # Then set the aspect ratio to auto.
        return "auto"

def apply_sigstar(q_value,
                  char = "$*$"):
    """
    =================================================================================================
    apply_sigstar(q_value, char)
    
    =================================================================================================
    Arguments:
    
    q_value  ->  A q value (or some other significance value)
    char     ->  How to represent things with significance
    
    =================================================================================================
    Returns: A string which represents whether the value is < 0.001, 0.01, 0.05
    
    =================================================================================================
    """
    # Try to float the input value, and raise an error
    # if it does not work.
    try:
        q = float(q_value)
    except:
        raise ValueError(f"The input value is not floatable: {q_value}")
    # If the input value is not a number
    if q != q:
        # Then return an empty string
        return ""
    # Or if the input value is less than 0.001
    elif q <= 0.001:
        # Then return the char variable three times
        return fr" {char}{char}{char}"
    # Or if the input value is less than 0.01
    elif q <= 0.01:
        # Then return the char variable twice
        return fr" {char}{char}"
    # Or if the input value is less than 0.05
    elif q <= 0.05:
        # Then return the char variable once
        return fr" {char}"
    # Otherwise
    else:
        # Return an empty string
        return "n.s."

def make_sigstars(q_list, char = fr"$*$"):
    """
    =================================================================================================
    make_sigstars(q_list)
    
    =================================================================================================
    Argument:
    
    q_list  ->  A list of q values (or p values)
    
    =================================================================================================
    Returns: A list of significance stars that represent each value in the input list.
    
    =================================================================================================
    """
    # Use list comprehension and apply_sigstar to make a list of lists of significacne
    # stars for each value in the input lists of lists
    return [[apply_sigstar(q) for q in col] for col in q_list]

def plot_sigstars(axes,
                  q_list, 
                  text_dict = {}):
    """
    =================================================================================================
    plot_sigstars(axes, q_list)
    
    =================================================================================================
    Arguments:
    
    axes   ->  A matplotlib axes object where the significance stars will be plotted
    q_list ->  The list of q_values that represent significance
    
    =================================================================================================
    Returns: None
    
    =================================================================================================
    """
    # Get the significance stars using make_sigstars(q_list).
    # q_list is checked inside make_sigstars, so no need to check it.
    stars = make_sigstars(q_list)
    # Loop over the number of columns in sigstars
    for i in range(len(stars)):
        # Loop over the number of rows in sigstars
        for j in range(len(stars[0])):
            # Add text to the axes object at row j, column i.
            axes.text(j,i,stars[i][j], ha = "center",
                           va = "center", color = "black", **text_dict)
    return None

def get_sig_indices(sig_stars_list):
    """
    =================================================================================================
    get_sig_indices(sig_stars_list)
    
    =================================================================================================
    Arguments:
    
    sig_stars_list  ->  The list of lists of significance stars for the 
    
    =================================================================================================
    Returns: The indices of lists where significance stars exist.
    
    =================================================================================================
    """
    # Use list comprehension to get the indeices where there
    # is one or more star characters
    return [i for i in range(len(sig_stars_list)) if "*" in sig_stars_list[i] or "**" in sig_stars_list[i] or "***" in sig_stars_list[i]]

def sig_filtering(data_list,
                  yticks,
                  significance):
    """
    =================================================================================================
    sig_filtering(data_list, yticks, significance)
    
    =================================================================================================
    Arguments:
    
    data_list     ->  A list of data points to be filtered
    yticks        ->  A list of ytick vlaues to be filtered
    significance  ->  A list of significance values to be filtered
    
    =================================================================================================
    Returns: The filtered input lists
    
    =================================================================================================
    """
    # Use the get_sig_indices() function to get the indices of
    # lists with significanct values
    keep_indices = get_sig_indices(make_sigstars(significance))
    # If the list is empty
    if keep_indices == []:
        # Then return a list of empty lists
        return [[],[],[]]
    # Otherwise, use the remove_list_indices() function from general
    # helpers to filter out the lists without significant values
    # from each input list.
    else:
        # Filter the data list
        filt_dlist = gh.remove_list_indices(data_list,
                                            keep_indices,
                                            opposite = False)
        # Filter the yticks list
        filt_ylist = gh.remove_list_indices(yticks,
                                            keep_indices,
                                            opposite = False)
        # Filter the significance list
        filt_slist = gh.remove_list_indices(significance,
                                            keep_indices,
                                            opposite = False)
        # Return the filtered lists at the end.
        return filt_dlist, filt_ylist, filt_slist
            
def heatmap(data_list,
            xticks = None,
            yticks = None,
            cmap = "bwr",
            bad_color = "grey",
            clb_label = "Normalized\nEnrichment\nScore",
            heat_title = "I'm a heatmap!!",
            remove_spines = False,
            significance = None,
            sig_filter = False,
            save = True,
            img_type = 'pdf',
            img_name = "im_a_heatmap",
            subplot_args = {'figsize' : (12,12)},
            colorbar_args = {"shrink" : 0.5,
                             "pad"    : 0.08},
            textdict = {"fontfamily" : "sans-serif",
                        "font" : "Arial",
                        "fontweight" : "bold"}):
    """
    =================================================================================================
    heatmap(data_list, **kwargs)
    
    This function is meant to craete a heatmap from the input data list using matplotlibs imshow()
    
    =================================================================================================
    Arguments:
    
    data_list       ->  A list of lists containing the values for a heatmap
    
    xticks          ->  A list of strings which should label the x axis
    yticks          ->  A list of strings which should label the y axis
    cmap            ->  A string for a matplotlib.cm colormap
    bad_color       ->  A string for the color to make bad values (nan)
    clb_label       ->  A string to label the colorbar
    heat_title      ->  A string to label the heatmap
    significance    ->  A list of significance values
    sig_filter      ->  A boolean that determines whether to filter based on significance
                        and plot the filtered heatmap
    save            ->  A boolean for whether or not to save the plot
    img_type        ->  A string for the files extension
    img_name        ->  A string with the name for the image
    subplot_args    ->  A dictionary of arguments passed to plt.subplots()
    colorbar_args   ->  A dictionary of arguments to be passed to plt.colorbar()
    text_dict       ->  A dictionary of arguments to help format text
    
    =================================================================================================
    Returns: None, but a heatmap will be saved.
    
    =================================================================================================
    """
    # We aren't going to check the arguments here, since many of them
    # will be checked in other functions
    #
    plt.rcParams["font.size"] = 12
    # Make data_list into an array to be used in ax.imshow()
    data_arr = np.array(data_list, dtype = float)
    # Find the range of the values used in the heatmap.
    minval, maxval = get_range(data_list)
    # Changing the properties of global heatmap values are depricated.
    # Thus we have to copy the colourmap desired and change the copy.
    use_cmap = copy.copy(cm.get_cmap(cmap))
    use_cmap.set_bad(bad_color)
    # Use the subplots command to make a figure and axes
    # and pass in the subplots_args dictioanry
    fig, ax = plt.subplots(1,1,**subplot_args)
    # Make the heatmap using imshow() passing in
    # the data_array and the colormap. The aspect ratio
    # is controlled by infer_aspect_ratio()
    heat = ax.imshow(data_arr,
                     cmap = use_cmap,
                     alpha = 0.75,
                     vmin = minval,
                     vmax = maxval,
                     aspect = infer_aspect_ratio(xticks,yticks),
                     interpolation = "nearest")
    if remove_spines:
        ax.spines["right"].set_visible(False)
        ax.spines["left"].set_visible(False)
        ax.spines["top"].set_visible(False)
        ax.spines["bottom"].set_visible(False)
    # Set the title of the plot
    ax.set_title(heat_title, fontsize = 16, **text_dict)
    # Make the colorbar, passing in the colorbar_args dictionary
    clb = plt.colorbar(heat,
                       **colorbar_args)
    # And set the label of the colorbar.
    clb.ax.set_title(clb_label, fontsize = 12, **text_dict)
    # Next, set the ticks of the axis in the correct places.
    ax.set_xticks([i for i in range(len(data_arr[0]))])
    ax.set_yticks([i for i in range(len(data_arr))])
    # and if xticks are given
    if xticks != None:
        # Then label the x axis with the xticks
        ax.set_xticklabels(xticks, rotation = 45, ha = 'right', rotation_mode = "anchor", **text_dict)
    # and if yticks are given
    if yticks != None:
        # Then label the y axis with the given labels
        ax.set_yticklabels(yticks, ha = "right", va = "center", **text_dict)
    # These next commands are meant to help with placement of the xticks and yticks.
    # I admit I do not fully understand what they do.
    tick_spacing = 1
    ax.xaxis.set_major_locator(ticker.MultipleLocator(tick_spacing))
    ax.yaxis.set_major_locator(ticker.MultipleLocator(tick_spacing))
    # If significance values were given
    if significance != None:
        # Then run the plot_sigstars function passing in the
        # ax object and the significacne list.
        plot_sigstars(ax,
                      significance,
                      text_dict)
    # If the user wants to show the plot
    if save:
        # Then run plt.show() to print it
        plt.savefig(f"{img_name}.{img_type}", bbox_inches = 'tight')
    # This next set of code manages the filtering and replotting event.
    #
    # If the user elects to filter and significance values are given
    if significance != None and sig_filter:
        # Then run sig_filtering() and save the results.
        filt_dlist, filt_ylist, filt_slist = sig_filtering(data_list,
                                                           yticks,
                                                           significance)
        # If the lists are empty
        if filt_dlist == []:
            # Then return None, as there is no point in filtering.
            return ax
        # If the filtered lists are not empty,
        else:
            axes = [ax]
            # Then plot the filtered heatmap
            new_ax = plot_heatmap(filt_dlist,
                         xticks = xticks,
                         yticks = filt_ylist,
                         cmap = cmap,
                         bad_color = bad_color,
                         clb_label = clb_label,
                         heat_title = f"{heat_title}",
                         sig_filter = False,
                         significance = filt_slist,
                         img_type = img_type,
                         img_name = f"{img_name}_filtered",
                         subplot_args = subplot_args,
                         colorbar_args = colorbar_args,
                         text_dict = text_dict)
            axes.append(new_ax)
            return axes
    return ax

#
#
############################################################################################################
#
#

# Make dotplots
def add_errorbar(mpl_axes, x_pos, y_pos, 
                 std, color = "grey", x_offset = 0.05, 
                 transparency = 0.75):
    """
    Adds error bars to MatPlotLib axes object. Modifies the mpl_axes input, returns None.

    mpl_axes -> matplotlib axes object
    x_pos    -> position of the center of the error bars on the x axis
    y_pos    -> position of the center of the error bars on the y axis
    std      -> standard deviation of the data (or vertical offset for error bars)
    color    -> The color of the error bars (Default: "grey")
    x_offset -> Size of the middle of the error bars (Default: 0.05)
    """
    # Plot the vertical middle bar
    mpl_axes.plot([x_pos, x_pos], [y_pos + std, y_pos - std], color = color, alpha = transparency)
    # Plot the horizontal middle bar
    mpl_axes.plot([x_pos - x_offset, x_pos + x_offset], [y_pos, y_pos], color = color, alpha = transparency)
    # Plot the horizontal top bar
    mpl_axes.plot([x_pos - 0.75*x_offset, x_pos + 0.75*x_offset], [y_pos + std, y_pos + std], color = color, alpha = transparency)
    # Plot the horizontal bottom bar
    mpl_axes.plot([x_pos - 0.75*x_offset, x_pos + 0.75*x_offset], [y_pos - std, y_pos - std], color = color, alpha = transparency)
    return None

def get_centers(data_matrix, width = 7):
    """
    """
    assert width <= 10, "Width must be between 0 and 10"
    # Assume the data are labelled
    data = [[item[0], sorted(item[1], reverse = True)] for item in data_matrix]
    n = len(data)
    x_coords = []
    # Loop over the index of each list
    for i in range(n):
        m = len(data[i][1])
        xi_coords = []
        # Loop over the number of times width splits the list
        # Integer division rounds down, so for m in [0,width],
        # m // width = 0. The +1 tells us to loop once for that group
        for j in range(m//width + 1):
            xi_coords += [(i+1) - 0.2 + 0.05*(k) for k in range(width)]
        x_coords.append(xi_coords[:m])
    return x_coords, data

def get_data_info(data_matrix, width = 7):
    bins = len(data_matrix)
    centers = [i - 0.2 + 0.05 * (width/2) for i in range(1, bins+1)]
    xs, data = get_centers(data_matrix, width = width)
    means = [sh.mean(item[1]) for item in data]
    sds = [sh.standard_deviation(item[1]) for item in data_matrix]
    sems = [sh.sem(item[1]) for item in data]
    return {"centers" : centers,
            "xs"      : xs,
            "means"   : means,
            "sds"     : sds,
            "sems"    : sems}, data

def make_ymax(mpl_axes,
              labelled_groups):
    """
    """
    current = list(mpl_axes.get_yticks())
    diff = abs( max(current) - min(current))
    if 1 <= len(labelled_groups) < 5:
        return max(current) + 0.3*diff
    elif 5 <= len(labelled_groups) < 10:
        return max(current) + 0.45*diff
    else:
        return max(current) + 0.6*diff

def make_ymin(mpl_axes,
              labelled_groups):
    """
    """
    current = list(mpl_axes.get_yticks())
    diff = abs( max(current) - min(current))
    if 1 <= len(labelled_groups) < 5:
        return min(current) - 0.1*diff
    elif 5 <= len(labelled_groups) < 10:
        return min(current) - 0.15*diff
    else:
        return min(current) - 0.2*diff

def make_ytick_labels(current_ticks, n, numstring = ""):
    """
    """
    new_ticks = []
    for item in current_ticks:
        if int(item) == item:
            new_ticks.append(f"{int(item)}{numstring}")
        else:
            new_ticks.append(f"{item:.1f}{numstring}")
    return new_ticks

def update_yticks(mpl_axes, fontdict = {"fontfamily" : "sans-serif",
                                         "font" : "Arial",
                                         "ha" : "right",
                                         "fontweight" : "bold",
                                         "fontsize" : "12"}):
    """
    """
    current = mpl_axes.get_yticks()
    diff = current[-1] - current[0]
    if 0 < diff < 1000:
        diff = current[1]-current[0]
        mpl_axes.set_yticks(current)
        mpl_axes.set_yticklabels(make_ytick_labels(current, len(current)),
                                     **fontdict)
    elif 1000 <= diff <= 999999:
        diff = current[1]-current[0]
        mpl_axes.set_yticks([i*(diff) for i in range(len(current))])
        mpl_axes.set_yticklabels(make_ytick_labels([c/1000 for c in current], len(current), "K"),
                                     **fontdict)
    elif 1000000 <= diff:
        diff = current[1]-current[0]
        mpl_axes.set_yticks([i*(diff) for i in range(len(current))])
        mpl_axes.set_yticklabels(make_ytick_labels([c/1000000 for c in current], len(current), "M"),
                                     **fontdict)
    return None

def update_xticks(mpl_axes,
                  xpos,
                  xlabels,
                  fontdict = {"fontfamily" : "sans-serif",
                               "font" : "Arial",
                               "ha" : "center",
                               "fontweight" : "bold",
                               "fontsize" : "12"}):
    """
    """
    mpl_axes.set_xticks(xpos)
    mpl_axes.set_xticklabels(xlabels, **fontdict)
    return None

def add_relative_axis(mpl_axes,
                      rel_data,
                      data_matrix,
                      info_dict,
                      fontdict = {"fontfamily" : "sans-serif",
                               "font" : "Arial",
                               "ha" : "left",
                               "fontweight" : "bold",
                               "fontsize" : "12"},
                      ylabel = "Fold Change",
                      remove_top_spine = True):
    """
    """
    n = len(data_matrix)
    rel_index = [i for i in range(n) if rel_data in data_matrix[i]]
    data_scaled = [(item[0], [value / info_dict["means"][rel_index[0]] 
                              for value in item[1]])
                   for item in data_matrix]
    new_y = mpl_axes.twinx()
    if remove_top_spine:
        new_y.spines["top"].set_visible(False)
    ymax = make_ymax(mpl_axes,data_scaled)
    current_lims = mpl_axes.get_ylim()
    new_y.set_ylim(current_lims[0] / info_dict["means"][rel_index[0]],
                   current_lims[1] / info_dict["means"][rel_index[0]])
    for i in range(n):
        new_y.scatter(info_dict["xs"][i], data_scaled[i][1], alpha = 0)
    update_yticks(new_y, fontdict = fontdict)
    current_lims = mpl_axes.get_ylim()
    new_y.set_ylim(current_lims[0] / info_dict["means"][rel_index[0]],
                   current_lims[1] / info_dict["means"][rel_index[0]])
    new_y.set_ylabel(ylabel,
                     font = "Arial",
                     fontsize = 14,
                     fontweight = "bold",
                     rotation = 270, va = "baseline")
    return None

def update_ylims(ymin, ymax, mpl_axes, labelled_groups):
    if ymin == ymax:
        ymin = make_ymin(mpl_axes, labelled_groups)
        ymax = make_ymax(mpl_axes, labelled_groups)
        mpl_axes.set_ylim(ymin, ymax)
    elif ymin == None and ymax != None:
        ymin = make_ymin(mpl_axes, labelled_groups)
        mpl_axes.set_ylim(ymin, ymax)
    elif ymin != None and ymax == None:
        ymax = make_ymax(mpl_axes, labelled_groups)
        mpl_axes.set_ylim(ymin, ymax)
    else:
        mpl_axes.set_ylim(ymin, ymax)
    return None

def update_ylims(axes, ymin, ymax):
    current = list(axes.get_ylim())
    if ymin == ymax and ymin == None:
        axes.set_ylim(*current)
        return
    elif ymin == None and ymax != None:
        axes.set_ylim(current[0], ymax)
        return
    elif ymin != None and ymax == None:
        axes.set_ylim(ymin, current[1])
        return
    else:
        axes.set_ylim(ymin, ymax)
        return

def update_xlims(mpl_axes_1, mpl_axes_2):
    xvals = list(mpl_axes_2.get_xlim())
    mpl_axes_1.set_xlim(*xvals)
    return

def plot_id_lines(mpl_axes_1, # Comparison axis
                  mpl_axes_2, # Dotplot_axis
                  labelled_data,
                  centers,  # Center value for each group
                  heights,  # list of highest comparison for the groups
                  colours = [],  # list of colours, one per group
                  plot_kwargs = {"alpha" : 0.5,
                                 "linestyle" : ":"}): 
    ax2_max = max(list(mpl_axes_2.get_ylim()))
    ax1_min = min(list(mpl_axes_1.get_ylim()))
    n = len(centers)
    if colours == []:
        colours = ["grey" for _ in range(n)]
    for i in range(n):
        mpl_axes_2.plot([centers[i], centers[i]],
                        [max(labelled_data[i][1]), ax2_max], colours[i], **plot_kwargs)
        mpl_axes_1.plot([centers[i], centers[i]],
                        [ax1_min, heights[i]], colours[i], **plot_kwargs)
    return None

# Functions to handle comparisons. Comparisons should be a dictionary of groups & pvalues
# from statistical tests in stats_helpers

# These functions are to format the comparisons such that we can
#    a) Filter them on significance if desired/required
#    b) Get the indices of the groups fromt he data
#    c) Get all of the x positions (centers) using the indices

def format_comps(comp_dict):
    # Return a list of [group 1, group 2] and [pval]
    groups = list(zip(comp_dict["Group 1"], comp_dict["Group 2"]))
    ps = comp_dict["pvalue"]
    combined = [[sorted(groups[i]), ps[i]] for i in range(len(ps))]
    return sorted(combined, key = lambda x: x[0][0])

def match_comp_to_data(comp_list, labelled_data):
    # Return the index for each group in the list
    inds = []
    labels = [item[0] for item in labelled_data]
    for item in comp_list:
        inds.append([sorted([labels.index(item[0][0]), labels.index(item[0][1])]), item[1]])
    inds = sorted(inds, key = lambda x: x[0])
    return inds

def make_xvals(comp_dict,
               labelled_data,
               centers,
               filter_by_alpha = False,
               alpha = 0.05):
    comp_list = format_comps(comp_dict)
    if filter_by_alpha:
        comp_list = [item for item in comp_list if item[1] < alpha]
        if len(comp_list) >= 19:
            return "toomany", "comps", "toplot"
        filtered = True
    elif len(comp_list) >= 19:
        print(f"Too many comparisons provided. Filtering by significance : {alpha}")
        comp_list = [item for item in comp_list if item[1] < alpha]
        if len(comp_list) >= 19:
            return "toomany", "comps", "toplot"
        filtered = True
    else:
        filtered = False
    data_inds = match_comp_to_data(comp_list, labelled_data)
    return [[centers[ind[0][0]], centers[ind[0][1]]] for ind in data_inds], [item[1] for item in data_inds], filtered

def make_sigstrings(value_list, sig_dict = {0.05 : "$*$",
                                            0.01 : "$**$",
                                            0.001 : "$**$$*$"}):
    sig_thresholds = list(sig_dict.keys())
    strings = []
    for val in value_list:
        sig = "n.s."
        for thresh in sig_thresholds:
            if val < thresh:
                sig = sig_dict[thresh]
        strings.append(sig)
    return strings

def generate_legend_string(p_or_q,
                           sig_dict,
                           omit = False):
    """
    """
    # Values in the dict are the representation
    # Keys in the dict are the cutoff
    items = list(sig_dict.items())
    #
    if omit:
        string = f"${p_or_q}\geq{items[0][0]}$ omitted\n"
    else:
        string = f"{'n.s.':<9}: ${p_or_q}\geq{items[0][0]}$"
    for item in items:
        centering = len(item[1]) + 5
        string = f"{string}\n{item[1]:<10}: ${p_or_q}<{item[0]}$"
    return string
    


def plot_comparisons(mpl_axes_1,                   # Comparison axes
                     mpl_axes_2,                   # Dotplot axes
                     labelled_data,                # 
                     comp_dict = {},
                     filter_by_alpha = False,
                     alpha = 0.05,
                     centers = [],
                     colours = [],
                     sig_dict = {0.05 : "$*$",
                                 0.01 : "$**$",
                                 0.001 : "$**$$*$"},
                     textdict = {"fontfamily" : "sans-serif",
                                 "font" : "Arial",
                                 "ha" : "left",
                                 "va" : "center",
                                 "fontweight" : "bold"},
                     return_height = True,
                     p_or_q = "p"):
    # Max comparisons hard set to 19, so make the y values for bars and text
    ys = [[i/19+0.02, i/19+0.02] for i in range(20)]
    text_y = [(ys[i][0] + ys[i+1][0])/2 for i in range(19)]
    # Handle anovas
    if comp_dict["id"][0] == "ANOVA":
        comp_dict["Group 1"] = [labelled_data[0][0]]
        comp_dict["Group 2"] = [labelled_data[-1][0]]
    # Need the xvalues for the bars and comparisons
    xs, pvals, filtered = make_xvals(comp_dict, labelled_data, 
                           centers, filter_by_alpha = filter_by_alpha,
                           alpha = alpha)
    if xs == "toomany":
        return None, None
    text_x = [sh.mean(item) for item in xs]
    # Make the significance strings
    pvals = make_sigstrings(pvals)
    # Keep track of the heights
    heights = {}
    maxtext = 0
    # Now we should be able to plot comparisons
    for i in range(len(xs)):
        mpl_axes_1.plot(xs[i], ys[i], color = "black", alpha = 0.5)
        mpl_axes_1.text(text_x[i], text_y[i], pvals[i], **textdict)
        # Update heights
        heights[xs[i][0]] = ys[i][0]
        heights[xs[i][1]] = ys[i][1]
        maxtext = text_y[i]
    # Turn the heights into a list
    if comp_dict["id"][0] == "ANOVA":
        heights = [ys[0][0] for _ in range(len(labelled_groups))]
    else:
        heights = sorted([[key, value] for key, value in heights.items()], key = lambda x: x[0])
        heights = [item[1] for item in heights]
    plot_id_lines(mpl_axes_1, mpl_axes_2, labelled_data, centers, heights, colours = colours)
    if filtered:
        string = generate_legend_string(p_or_q, sig_dict, omit = True)
        #string = f"Test : {comp_dict['id'][0]}\n\n${p_or_q}\geq0.05$ omitted\n$*$     : $p < 0.05$\n$**$   : $p<0.01$\n$**$$*$ : $p<0.001$"
    else:
        string = generate_legend_string(p_or_q, sig_dict)
        #string = f"Test : {comp_dict['id'][0]}\n\nn.s.   : $p\geq 0.05$\n$*$       : $p < 0.05$\n$**$    : $p<0.01$\n$**$$*$   : $p<0.001$"
    handles, labels = mpl_axes_1.get_legend_handles_labels()
    handles.append(Patch(color="none", label = string))
    if return_height:
        return maxtext, handles
    else:
        return None, handles

def dotplot(labelled_groups,
            foldchange_axis = False,
            foldchange_group = None,
            comparisons = {},        # Args for plot_comparisons
            filename = "dotplot.pdf",
            save_file = True,
            colours = [],
            title = "Dotplot",
            xlabel = "",
            ylabel = "Abundance",
            ymin = None,
            ymax = None,
            errorbar = "sem",
            filter_by_alpha = False,
            alpha = 0.05, 
            sig_dict = {0.05 : "$*$",
                        0.01 : "$*$$*$",
                        0.001 : "$*$$*$$*$"},
            p_or_q = "p"):
    assert errorbar.lower() in ["sem", "sd"], "The accepted errorbar settings are Standard Error of the Mean (sem) or Standard Deviation (sd)"
    global tab_colours
    if type(colours) == str:
        colours = handle_colours(colours, len(labelled_groups), choice = "random")
        print(f"Colours chosen: {colours}")
    #
    info_dict, groups = get_data_info(labelled_groups)
    #
    fig, ax = plt.subplots(2,1,sharex = True, figsize = (2*len(labelled_groups), 10))
    #
    nrow = 2
    ncol = 1
    #
    gs = gridspec.GridSpec(nrow,ncol,
         wspace=0.0, hspace=0.0, 
         top=1.-0.5/(nrow+1), bottom=0.5/(nrow+1), 
         left=0.5/(ncol+1), right=1-0.5/(ncol+1)) 
    #
    ax1 = plt.subplot(gs[0])
    ax1.set_ylim(0,1)
    ax1.axis("off")
    #
    ax2 = plt.subplot(gs[1])
    ax2.spines["right"].set_visible(False)
    #
    for i in range(len(groups)):
        if colours == []:
            ax2.scatter(info_dict["xs"][i], groups[i][1], edgecolors = "black")
        elif colours != []:
            ax2.scatter(info_dict["xs"][i], groups[i][1], edgecolors = "black", color = colours[i])
        if errorbar.lower() == "sem":
            add_errorbar(ax2, info_dict["centers"][i], info_dict["means"][i], info_dict["sems"][i])
        else:
            add_errorbar(ax2, info_dict["centers"][i], info_dict["means"][i], info_dict["sds"][i])
    # Axes related updates
    update_ylims(ax2, ymin, ymax)
    update_xlims(ax1,ax2)
    update_yticks(ax2)
    update_xticks(ax2, info_dict["centers"], [item[0] for item in groups])
    if foldchange_axis and foldchange_group != None:
        add_relative_axis(ax2, foldchange_group, groups, info_dict)
    ax2.set_xlabel(xlabel, **{"fontfamily" : "sans-serif",
                               "font" : "Arial",
                               "ha" : "center",
                               "fontweight" : "bold",
                               "fontsize" : "14"})
    ax2.set_ylabel(ylabel, **{"fontfamily" : "sans-serif",
                               "font" : "Arial",
                               "ha" : "center",
                               "fontweight" : "bold",
                               "fontsize" : "14"})
    # If comparisons are provided, plot them unless there are too many.
    if comparisons != {}:
        if colours == []:
            title_height, handles = plot_comparisons(ax1, ax2, groups, comparisons,
                                                     filter_by_alpha = filter_by_alpha, alpha = alpha,
                                                     centers = info_dict["centers"],
                                                     colours = tab_colours, sig_dict = sig_dict,
                                                     p_or_q = p_or_q)
        else:
            title_height, handles = plot_comparisons(ax1, ax2, groups, comparisons,
                                                     filter_by_alpha = filter_by_alpha, alpha = alpha,
                                                     centers = info_dict["centers"],
                                                     colours = colours, sig_dict = sig_dict,
                                                     p_or_q = p_or_q)
        if title_height != handles:
            ax1.text(sh.mean(list(ax1.get_xlim())), title_height + 0.1,
                     title, **{"fontfamily" : "sans-serif",
                               "font" : "Arial",
                               "ha" : "center",
                               "fontweight" : "bold",
                               "fontsize" : "16"})
            ax1.legend(handles = handles, loc = "center right", 
                       bbox_to_anchor = (0,0.5),
                       frameon = False)
        else:
            ax2.set_title("title", **{"fontfamily" : "sans-serif",
                                  "font" : "Arial",
                                  "ha" : "center",
                                  "fontweight" : "bold",
                                  "fontsize" : "16"})
    else:
        ax2.set_title(title, **{"fontfamily" : "sans-serif",
                                  "font" : "Arial",
                                  "ha" : "center",
                                  "fontweight" : "bold",
                                  "fontsize" : "16"})
    ax2.spines["top"].set_visible(False)
    if save_file:
        plt.savefig(filename)
    return ax


#
#
############################################################################################################
#
#         Exploiting matplotlib.axes.Axes.table()

from matplotlib.transforms import Bbox

def make_mpl_table(*lists,
                   fig = None,
                   ax = None, 
                   colours = [], 
                   columns = True,
                   colLabels = [],
                   rowLabels = [],
                   colLabels_colours = [],
                   rowLabels_colours = [],
                   tableargs = {"loc" : (0,0),
                                "edges" : "open"},
                   fontfamily = "sans-serif",
                   font = "Arial",
                   fontsize = "14",
                   fontweight = "bold"):
    """
    """
    if ax == None and fig == None:
        x = float(input("Please input the X component of figure size (float):\t" ))
        y = float(input("Please input the Y component of figure size (float):\t" ))
        fig, ax = plt.subplots(figsize = (x,y))
    elif ax == None or fig == None:
        assert False, "Both a figure and axes object should be passed in, or neither."
    if columns:
        lists = gh.transpose(*lists)
    if colLabels != []:
        assert len(colLabels) == len(lists[0]), "Not enough column labels provided"
    else:
        colLabels = ["" for _ in range(len(lists[0]))]
    if rowLabels != []:
        assert len(rowLabels) == len(lists), "Not enough row labels provided"
    else:
        rowLabels = ["" for _ in range(len(lists))]
    if colLabels_colours != []:
        assert len(colLabels_colours) == len(colLabels), "The number of colours provided for column labels\ndoes not match the number of labels"
    else:
        colLabel_colours = ["black" for _ in range(len(colLabels))]
    if rowLabels_colours != []:
        assert len(rowLabels_colours) == len(rowLabels), "The number of colours provided for row labels\ndoes not match the number of labels"
    else:
        rowLabel_colours = ["black" for _ in range(len(rowLabels))]
    if colours != [] and columns:
        colours = gh.transpose(*colours)
        assert all([len(row) == len(lists[0]) for row in colours]), "Too few colours provided for each column"
        assert len(colours) == len(lists), "Too few rows of colours provided"
    elif colours != [] and not columns:
        assert all([len(row) == len(lists[0]) for row in colours]), "Too few colours provided for each column"
        assert len(colours) == len(lists), "Too few rows of colours provided"
    else:
        colours = [["black" for _ in range(len(lists[0])+1)] for i in range(len(lists)+1)]
    tab = ax.table(lists, rowLabels = rowLabels, colLabels = colLabels, **tableargs)
    r = fig.canvas.get_renderer()
    cellwidths = [[0 for _ in range(len(lists[0])+1)] for i in range(len(lists)+1)]
    cellheights = [[0 for _ in range(len(lists[0])+1)] for i in range(len(lists)+1)]
    for cell in tab._cells:
        text = tab._cells[cell].get_text()
        text.set_fontfamily(fontfamily)
        text.set_font(font)
        text.set_fontweight(fontweight)
        text.set_fontsize(fontsize)
        text.set_ha("center")
        coords = text.get_window_extent(renderer=r)
        coords = Bbox(ax.transData.inverted().transform(coords))
        width = coords.width
        height = coords.height
        cellwidths[cell[0]][cell[1]+1] = width
        cellheights[cell[0]][cell[1]+1] = height
    cellwidths = [max(col) for col in gh.transpose(*cellwidths)]
    cellheights = [max(col) for col in cellheights]
    for cell in tab._cells:
        if cell[0] == 0:
            text = tab._cells[cell].get_text()
            text.set_color(colLabel_colours[cell[1]])
            tab._cells[cell].set_width(cellwidths[cell[1]+1])
            tab._cells[cell].set_height(cellheights[cell[0]])
        elif cell[1] == -1:
            text = tab._cells[cell].get_text()
            text.set_color(rowLabel_colours[cell[0]-1])
            tab._cells[cell].set_width(cellwidths[cell[1]+1])
            tab._cells[cell].set_height(cellheights[cell[0]])
        else:
            text = tab._cells[cell].get_text()
            text.set_color(colours[cell[0]-1][cell[1]])
            tab._cells[cell].set_width(cellwidths[cell[1]+1])
            tab._cells[cell].set_height(cellheights[cell[0]])
    return fig, ax, tab

#
#
############################################################################################################
```
