## Supporting Folder 2 Data Analysis for "Mouse primary T cell phosphotyrosine proteomics enabled by BOOST": pandas_helpers.py

```
"""
=================================================================================================
Kenneth P. Callahan

9 July 2021
                  
=================================================================================================
Python >= 3.8.5

pandas_helpers.py

Functions to help manipulate pandas dataframes (and other pandas objects). This is meant to be
imported into other scripts.

=================================================================================================
Dependencies:

 PACKAGE      VERSION
  numpy   ->  1.20.1
  pandas  ->  1.2.3
  
Kenny's Dependencies:

  homebrew_stats  ->  Statistics, should be in the same folder as this script
  general_helpers ->  General functions, should be in the same folder as this script
=================================================================================================
"""
print(f"Loading the module: helpers.pandas_helpers\n")
######################################################################################################
#
#     Importables

# This module is used for unix-like string matching,
# namely WildCard notation (*string*).
import fnmatch
import os

# DataFrames, the main focus of this file, are
# pandas objects.
import pandas as pd

# Numpy is used for applying logarithms to columns
# of pandas DataFrames.
import numpy as np

# Homebrew stats is my self curated statistics module. I had
# some trouble using the formal statistics modules like
# SciPy, so I coded some statistics. There also is no
# standard module for applying the Storey method for
# q-value estimation, so I coded that as well.
from . import stats_helpers as hs

# General helpers has a number of functions I use frequently in
# my scripts. They are all placed in that module purely for
# convenience and generalizability.
from . import general_helpers as gh

# Argcheck helpers has functions that I use to check the
# validity of arguments in functions, since Python is
# weakly typed.
from . import argcheck_helpers as ah

print(f"pandas        {pd.__version__}")
print(f"numpy         {np.__version__}\n")

#
#
######################################################################################################
#
#     Dealing with column headers

def df_make_headers(strs_list,
                    delim = "_vs_",
                    reverse = False):
    """
    =================================================================================================
    df_make_headers(strs_list, delim, reverse)
    
    This function is meant to take a list of lists of strings, and return a list of strings separated
    by the delim argument. In use, these strings act as column headers in a Pandas DataFrame.
                  
    =================================================================================================
    Arguments:
    
    strs_list  ->  A list of lists of strings, where each sublist contains the information required
                   to label one column of a Pandas DataFrame
    delim      ->  A string which will separate each sublist in strs_list. Try not to use spaces
    reverse    ->
    
    =================================================================================================
    Returns:
    
    =================================================================================================
    """
    # First, check the input types to make sure they're good
    # strs_list should be a lsit
    strs_list = ah.check_type(strs_list, [list, type(gh.unique_combinations([1,2,3]))],
                              error = "The strs_list argument is not a list...")
    # and each element of strs list should be a list
    for i in range(len(strs_list)):
        strs_list[i] = ah.check_type(strs_list[i], list,
                                     error = f"Sublist {i} of strs_list is not a list...")
        # and each element of strs_list[i] should be a string
        for j in range(len(strs_list[i])):
            strs_list[i][j] = ah.check_type(strs_list[i][j], str,
                                            error = f"Element {j} of sublist {i} is not a string...")
    delim = ah.check_type(delim, str,
                          error = f"The delim argument is not a string...")
    reverse = ah.check_type(reverse, bool,
                            error = "The reverse argument is not a boolean")
    # Once all of the inputs are checked,
    # determine whether to reverse the order of each sublist.
    # If reverse is true
    if reverse:
        # Then the new headers should use the reverse order
        # of the input sublists. So use list comprehension to
        # reverse the list order, and use list_to_str to convert
        # the reversed list into a string. Do this for each
        # sublist in the input list.
        new_heads = [gh.list_to_str([item[-i] for i in range(1,len(item)+1)], delimiter = f"{delim}", newline = False) for item in strs_list]
    # If reverse is False
    else:
        # Then use the sublists verbatum to make strings,
        # and use list comprehension to convert each sublist
        # into a string.
        new_heads = [gh.list_to_str(item, delimiter = f"{delim}", newline = False) for item in strs_list]
    # Once this is complete, return new_heads.
    return new_heads

def df_pair_heads_and_indices(group_lists,
                              headers):
    """
    =================================================================================================
    df_pair_heads_and_indices(group_lists, headers)
    
    This function is meant to take a list of lists of group headers and a list of formatted headers,
    and determine which elements should go together (by index in group_lists)
    
    Example:   group_lists = [["t1", "t2"],         headers = ["t2_vs_t1",
                              ["t1", "t3"],                    "t3_vs_t1",
                              ["t2", "t3"]]                    "t3_vs_t2"]
                              
               out_dict = {"t2_vs_t1" : [0], "t3_vs_t1" : [1], "t3_vs_t2" : [2]}
               
                  
    =================================================================================================
    Arguments:
    
    group_lists  ->  A list of lists of strings containing groups that are compared
    headers      ->  A list of strings describing all combinations group comparisons
    
    =================================================================================================
    Returns: A dictionary of lists of integers where keys are the headers and values are lists of
             indices that have the substrings for the key
    
    =================================================================================================
    """
    # Check that the argument types are correct
    # The group_lists should be a list of lists of strings
    group_lists = ah.check_type(group_lists, list, error = "The group_lists argument is not a list...")
    for i in range(len(group_lists)):
        group_lists[i] = ah.check_type(group_lists[i], list,
                                       error = f"Sublist {i} of group_lists is not a list...")
        for j in range(len(group_lists[i])):
            group_lists[i][j] = ah.check_type(group_lists[i][j], str,
                                              error = f"Element {j} of sublist {i} is not a string...")
    # The headers should be a list of strings
    headers = ah.check_type(headers, list,
                            error = "The headers argument is not a list...")
    headers = [ah.check_type(headers[i], str,
                             error = f"Element {i} of headers is not a string...") for i in range(len(headers))]
    # Once the arguments are checked, we can
    # proceed with the function.
    # Initialize the output dictionary
    out_dict = {}
    # and create header : list pairs for each header given.
    for head in headers:
        out_dict[head] = []
    # Initialize an index to keep track of the loop
    index = 0
    # Loop over each element in the group_lists
    for element in group_lists:
        # Loop over the headers in the headers list
        for head in headers:
            # If each element in the group is also
            # in the header
            if all([group in head for group in element]):
                # Then add the current index to the output
                # dictionary under the current header key.
                out_dict[head].append(index)
        # Once we've checked the headers,
        # increase the index by one and continue
        index+=1
    # Once the output dictionary has been updated,
    # return it.
    return out_dict

def df_reformat(headers,
                field_df,
                index_dictionary,
                identity = "peptide",
                field = 'pvalue'):
    """
    =================================================================================================
    df_reformat(headers, value_df, index_dictionary, identity, field)
    
    This function is meant to 
                  
    =================================================================================================
    Arguments:
    
    headers           ->  A list of strings describing the comparisons being used
    field_df          ->  A DataFrame containing a header with the field string as a column header 
    index_dictionary  ->  A dictionary with keys as comparisons being used and values as indices
                          (rows) where the comparison was used
    identity          ->  A string containing the header that identifies each row in the DataFrame
    field             ->  A string containing the column that we care about, the 'x-value'
    
    =================================================================================================
    Returns: A new DataFrame where the columns are defined by <headers>, the rows come from
             the <field> column of the input DataFrame, and the position of the values in the
             rows are defined by the index dictionary.
    
    =================================================================================================
    """
    # Check the input argument types:
    # The headers should be a list of strings
    headers = ah.check_type(headers, list,
                            error = "The headers argument is not a list...")
    headers = [ah.check_type(headers[i], str,
                             error = f"Element {i} of headers is not a string...") for i in range(len(headers))]
    # The field_df should be a dataframe
    field_df = ah.check_type(field_df, pd.DataFrame,
                             error = f"The field_df argument is not a DataFrame...")
    # The index_dictionary should be a dictionary
    index_dictionary = ah.check_type(index_dictionary, dict,
                                     error = f"The index_dictionary argument is not a dictionary...")
    # The identity and field arguments should be strings
    identity = ah.check_type(identity, str,
                             error = f"The identity argument should be a string...")
    field = ah.check_type(field, str,
                          error = f"The field argument should be a string...")
    # Once all the types have been checked, proceed with the program
    # Make a new DataFrame by taking all rows with indices
    # associated with the zeroeth header.
    new_df = field_df.iloc[index_dictionary[headers[0]]]
    # Then make the identity column the new index. All joining
    # will happen according to the identity column.
    new_df.set_index(identity, inplace = True)
    # Rename the field-column of the new DataFrame. The
    # rename method will filter all unspecified columns, so
    # this should result in a one-column DataFrame with field values
    new_df.rename(columns = {field:headers[0]}, inplace = True)
    # With the new dataframe initialized, we need to add the
    # remaining columns. Those are specified in the index_dictionary
    # So loop over the headers
    for head in headers:
        # and if the current headers is not already a
        # column header in the DataFrame
        if head not in list(new_df.columns.to_numpy()):
            # Parse the dictionary based on row indices in index_dict
            head_parse = field_df.iloc[index_dictionary[head]]
            # Change the index to the identity column
            head_parse.set_index(identity, inplace = True)
            # Rename the field column to the header, filtering
            # the other columns out.
            head_parse.rename(columns = {field:head}, inplace= True)
            # Join the new_df and head_parse DataFrame. When how=outer,
            # this is the equivalent of union in sets.
            new_df = new_df.join(head_parse, how = 'outer')
        # If the current header is in the columns
        else:
            # Just continue
            continue
    # Once the new_df has been sufficiently updated,
    # return the new_df
    return new_df

#
#
######################################################################################################
#
#     Dealing with Separating and Combining

def df_combine(*dataframes,
               keep_order = True,
               remake_index = False):
    """
    =================================================================================================
    df_combine(*dataframes, keep_order, remake_index)
    
    This function is meant to combine multiple dataframes by index.
                  
    =================================================================================================
    Arguments:
    
    dataframes    ->  An arbitrary number of pandas DataFrames
    keep_order    ->  A boolean, whether or not to keep the current order of DataFrame columns
    remake_index  ->  False or a string. If a string is given, then use that column as the
                      index when combining DataFrames
    
    =================================================================================================
    Returns: A new dataframe, which is the combination of all input DataFrames
    
    =================================================================================================
    """
    # Check the input arguments for validity
    # *dataframes should contain dataframes
    dataframes = [ah.check_type(dataframes[i], pd.DataFrame,
                                error = f"Element {i} of dataframes is not a DataFrame...") for i in range(len(dataframes))]
    # keep_order should be a boolean
    keep_order = ah.check_type(keep_order, bool,
                               error = f"The keep_order argument is not a boolean...")
    # The remake index argument should either be False
    # or a string
    remake_index = ah.check_type(remake_index, [bool, str],
                                 error = "The remake_index argument should be False or of type string...")
    # Now that all of the arguments are checked, continute.
    # First, if remake_index is a string
    if type(remake_index) == str:
        # Then loop over the dataframes
        for df in dataframes:
            # and set the indices to the remake_index value
            df.set_index(remake_index, inplace = True)
    # If keep_order is True
    if keep_order:
        # Then get a list of the columns of the zeroeth dataframe
        og_col_order = list(dataframes[0].columns.values)
        # and loop over the dataframes in the list
        for df in dataframes:
            # and update the columns for all columns not in
            # the current columns list.
            og_col_order = og_col_order + [col for col in list(df.columns.values) if col not in og_col_order]
    # Next, sort the dataframes by length, in reverse order.
    dataframes = sorted(dataframes, key=lambda x:len(x), reverse = True)
    # Initialize the new_df variable
    new_df = None
    # and loop over the dataframes
    for df in dataframes:
        # If new_df has not been set yet
        if type(new_df) == type(None):
            # Then set new_df to be the first
            # (and current) dataframe
            new_df = df
        # If the new_df has already ben set
        else:
            # Then get a list of the unique columns from the current dataframe
            unique = [item for item in list(df.columns.values) if item not in list(new_df.columns.values)]
            # and update the new_df with the the columns of the current
            # dataframe that are unique.
            new_df = new_df.join(df[unique], how = "outer")
    # Once new_df has been updated,
    # if keep_order is True
    if keep_order:
        # Then change the order of the columns to reflect the
        # original dataframe column order.
        new_df = new_df.reindex(columns = og_col_order)
    # If the index of the dataframes was changed during the process of combination
    if type(remake_index) == str:
        # Then reset the index to numbers
        new_df.reset_index(inplace = True)
        # and rename the new column as the remake_index string.
        new_df = new_df.rename(columns = {"index":remake_index})
    # Once all transformations have ceased,
    # return the new dataframe
    return new_df

def make_filtstrs(parsed_strs):
    """
    =================================================================================================
    make_filtstrs(parsed_strs)
    
    This is meant to make a number of wildcard like strings from a list of lists of strings.
                  
    =================================================================================================
    Arguments:
    
    parsed_strs  ->  A list of lists of strings, where the strings represent the pairs of
                     experiment types and conditions.
    
    =================================================================================================
    Returns: a list of the strings in wildcard notation. These will be used for filtering
             with fnmatch.
    
    =================================================================================================
    """
    # Check the input arguments:
    parsed_strs = ah.check_type(parsed_strs, list,
                                error = f"The parsed_strs argument is not a list...")
    for i in range(len(parsed_strs)):
        parsed_strs[i] = ah.check_type(parsed_strs[i], list,
                                       error = f"Sublist {i} of parsed_strs is not a list...")
        for j in range(len(parsed_strs[i])):
            parsed_strs[i][j] = ah.check_type(parsed_strs[i][j], str,
                                              error = f"Element {j} of sublist {i} is not a string...")
    # Once parsed_strs was checked, continue.
    # Initialize a list to hold the wildcard strings
    filterers = []
    # Loop over the tuples in parsed_strs
    for tup in parsed_strs:
        # Add a star to the newstr variable
        newstr = "*"
        # Loop over the strings in tup
        for string in tup:
            # and update newstr with the string and a star
            newstr += f"{string}*"
        # Once all strings have been added, add the newstr
        # to the filterers list
        filterers.append(newstr)
    # Once all of the strings have been made, return the
    # filterers list.
    return filterers

def df_parser(main_df,
              parsed_strs,
              id_col = None,
              set_index = False):
    """
    =================================================================================================
    df_parser(main_df, parsed_strs, id_col, set_index)
    
    This function is meant to take a DataFrame and split the DataFrame based on the strings in
    parsed_strs
    
    =================================================================================================
    Arguments:
    
    main_df      ->  A dataframe to be split into new dataframes
    parsed_strs  ->  A list of lists of strings that should be present in the column headers
    id_col       ->  A string representing the column of the DataFrame that is the identity
    set_index    ->  A boolean which determines whether the index needs to be set
    
    =================================================================================================
    Returns: a list of DataFrames, and strings used to filter the columns of the original
             DataFrame
    
    =================================================================================================
    """
    # Check some of the input arguments
    main_df = ah.check_type(main_df, pd.DataFrame,
                            error = "The main_df argument is not a pandas DataFrame...")
    # id_col should either be a string or None or list
    id_col = ah.check_type(id_col, [type(None), str, list],
                           error = "The id_col is neither a string, nor None, nor a list...")
    # The set_index should be a boolean
    set_index = ah.check_type(set_index, bool,
                              error = "The set_index argument is not a boolean...")
    # Make the filtering strings using Unix-like wildcard syntax
    # parsed_strs will be checked in make_filtstrs()
    filterers = make_filtstrs(parsed_strs)
    # Initialize a list to hold DataFrames after parsing
    dfs = []
    # Loop over the unix-like wildcard strings
    for f_string in filterers:
        # fnmatch filter returns a list of all column headers
        # with the wildcard string(s) in them
        cols = fnmatch.filter(list(main_df.columns.values), f_string)
        # If the number of columns found is zero,
        if len(cols) == 0:
            # then append an empty dataframe to the list
            dfs.append(pd.DataFrame(np.empty(shape=(len(main_df),0), dtype=np.float64)))
        # If the id_col was not set
        elif id_col == None:
            # and the number of columns is 1
            if len(cols) == 1:
                # Then append a dataframe of the one column
                # to the dfs list
                dfs.append(main_df[cols[0]])
            # If there is more than one column
            else:
                # Then add the dataframe with all of those
                # columns to the list
                dfs.append(main_df[cols])
        # If the id_col was set to a string
        elif type(id_col) == str:
            # Then add the dataframe with the id column and
            # all other columns to the list
            dfs.append(main_df[[id_col, *cols]])
        # If the id_column is neither None nor str type,
        else:
            # Then add the dataframe with the id_cols
            # and the filtered columns.
            dfs.append(main_df[[*id_col, *cols]])
    # Once the main_df has been filtered, check for any empty dataframes
    # by looping over the dataframes
    for i in range(len(dfs)):
        # and seeing whether or not the size of the dataframe is zero
        if bool(dfs[i].to_numpy().size == 0):
            raise ValueError("An empty DataFrame is present. It may be due to misordered parsed_strs tuples.")
    # If set_index is True and the id_col is a string
    if set_index and type(id_col) == str:
        # Then set the index of every dataframe to the id_col
        dfs = [df.set_index(id_col) for df in dfs]
    # At the end, return the dataframe list
    # and the strings used for filtering columns
    return dfs, filterers

def df_parse_fieldvalues(a_dataframe,
                         groups = ["Group 1", "Group 2"],
                         identity = "peptide",
                         reverse = True,
                         field = 'pvalue'):
    """
    =================================================================================================
    df_parse_fieldvalues(a_dataframe, groups, identity, reverse, field)
    
    This function is meant to take a pandas DataFrame, parse it, and reformat the parsed dataframes
    by group.
    
    =================================================================================================
    Arguments:
    
    a_dataframe  ->  A pandas DataFrame that has columns labels in <group>, <identity> and <field>
    groups       ->  A list of strings that identifys which groups are being compared.
    identity     ->  A string describing which column acts as the identity for rows in the dataframe
    reverse      ->  A boolean that determines whether comparison labels are in forward or reverse
                     order
    field        ->  A string describing which column will be used for the row values in the output
                     DataFrame
    
    =================================================================================================
    Returns: A pandas DataFrame where the columns are the comparisons between the groups in <groups>
             and the values are the values from the input DataFrame corresponding to the index of
             the compared group.
    
    =================================================================================================
    """
    # Check the input arguments
    a_dataframe = ah.check_type(a_dataframe, pd.DataFrame,
                                error = "The a_dataframe argument is not a DataFrame...")
    # groups will be checked in unique_combinations
    # The identity argument should be a string
    identity = ah.check_type(identity, str,
                             error = "The identity argument is not a string...")
    # The field argument should be a string
    field = ah.check_type(field, str,
                          error = "The field argument is not a string...")
    # Create a minimal dataframe by keeping only the identity,
    # the groups, and the field of interest.
    df_minimal = a_dataframe[[identity, *groups, field]]
    # Use unique combinations to to find all unique combinations of
    # the items in group_values. First, convert the values into a list
    group_values = [list(item) for item in df_minimal[groups].to_numpy()]
    # Then, run unique_combinations on the list
    uniq_combs = gh.unique_combinations(group_values)
    uniq_combs = gh.unique_combinations(gh.unpack_list(list(uniq_combs)))
    uniq_combs = sorted(list(uniq_combs))
    uniq_combs = gh.make_pairs(uniq_combs)
    # Unpack the list, then use make pairs with reverse = True
    # then deal with the consequences later
    
    
    # Use df_make_headers() to make headers from the unique
    # combinations of values in groups
    new_heads = df_make_headers(list(uniq_combs),
                                reverse = reverse)
    # Then, create a dataframe using only the field value and the identity.
    df_vals = a_dataframe[[identity, field]]
    # Next, use df_pair_heads_and_indices() to get a dictionary of
    # heads and indices for those headers
    paired = df_pair_heads_and_indices(group_values,new_heads)
    # Finally, use df_reformat() to create a new dataframe, using
    # the new headers, dataframe, and paired_indices
    return df_reformat(new_heads,
                       df_vals,
                       paired,
                       identity = identity,
                       field = field)

#
#
######################################################################################################
#
#     Dealing with Filtering Rows

def df_filter_nans(*dataframes, 
                   threshold = 3,
                   subset = None,
                   not_included = [None]):
    
    """
    =================================================================================================
    df_filter_nans(dataframes, threshold, subset, not_included)
    
    This function is meant to take an arbitrary number of dataframes and filter out the rows
    with >= threshold nan values
    
    =================================================================================================
    Arguments:
    
    dataframes    ->  An arbitrary number of DataFrames
    threshold     ->  An integer, the minimum number of values before filtering the row
    subset        ->  A string present in some of the headers, these are the headers to focus
                      filtering on. Can also be a list
    not_included  ->  A list of strings, which are headers to exclude from filtering.
    
    =================================================================================================
    Returns: A list of the dataframes after nan filtering.
    
    =================================================================================================
    """
    # Check the input arguments
    dataframes = list(dataframes)
    for i in range(len(dataframes)):
        dataframes[i] = ah.check_type(dataframes[i], pd.DataFrame,
                                      error = f"Element {i} of dataframes is not a DataFrame")
    # The threshold should be an integer
    threshold = ah.check_type(threshold, int,
                              error = "The threshold given is not an integer...")
    # The subset should be a string
    subset = ah.check_type(subset, [str,list, type(None)],
                           error = "The subset argument is not a string nor a list...")
    # The not_included should be a list of strings/None
    not_included = ah.check_type(not_included, list,
                                 error = "The not_included argument is not a list...")
    not_included = [ah.check_type(not_included[i], [str, type(None)],
                                  error = f"Element {i} of not_included is neither a string nor NoneType...") for i in range(len(not_included))]
    # Once argument checking is complete, continue.
    # Initialize the list of filtered dataframe
    filtered = []
    # Loop over the dataframes in the dataframes list.
    for df in dataframes:
        # Loop over the columns of the dataframe
        for col in list(df.columns.values):
            # If the column is not in the not_included list
            if col not in not_included:
                # Then we want to change the type of the values to floats,
                # and ignore the column if errors arise
                df[col] = df[[col]].astype(float, errors = "ignore")
        # If the subset value is set, and it is a string
        if subset != None and type(subset) == str:
            # Then get a list of columns that contain the subset
            new_sub = [item for item in list(df.columns.values) if subset in item]
            # and use the dropna() method of dataframes to filter out the
            # values in the current df, using only the subset rows.
            filtered.append(df.dropna(thresh = threshold, subset = new_sub))
        # If the subset value is set and it is a list
        elif subset != None and type(subset) == list:
            # Then use the dropna() method of datafranes to filter out
            # the values in the current df using the subset rows
            filtered.append(df.dropna(thresh = threshold, subset = subset))
        # Otherwise, the subset value has not been set,
        else:
            # So we can simply filter the rows using dropna()
            filtered.append(df.dropna(thresh = threshold))
    # Once all of the rows have been filtered in each dataframe,
    # return the list of filtered dataframes
    return filtered

#
#
######################################################################################################
#
#     Dealing with Statistics-Related  stuff

def df_row_mean(dataframes):
    """
    =================================================================================================
    df_row_mean(dataframes)
    
    This function is meant to take the row mean of each DataFrame in the list of inputs. The column
    'mean' will be added to the columns of the DataFrame
    
    =================================================================================================
    Arguments:
    
    dataframes  ->  A list of Pandas DataFrames
    
    =================================================================================================
    Returns: A list of pandas DataFrames, where the column 'mean' has been added to each DataFrame
    
    =================================================================================================
    """
    # Check the input arguments
    dataframes = ah.check_type(dataframes, list,
                               error = "The dataframes argument is not a list of DataFrames...")
    dataframes = [ah.check_type(dataframes[i], pd.DataFrame,
                                error = f"Element {i} of dataframes is not a DataFrame...") for i in range(len(dataframes))]
    # Loop over the DataFrames in dataframes
    for df in dataframes:
        # Make a new column in the DataFrame and add
        # the mean of each row.
        df['mean'] = df.mean(axis = 1)
    # At the end of the loop, return the dataframes list.
    return dataframes

def df_foldchange(dataframe1,
                  dataframe2,
                  label = "Hello World!"):
    """
    =================================================================================================
    df_foldchange(dataframe1, dataframe2, label)
    
    This function is meant to calculate the foldchange between two pandas DataFrames. It does so on
    index, so I suggest you convert the index column into some identity before applying this function.
                  
    =================================================================================================
    Arguments:
    
    dataframe1  ->  A pandas DataFrame, with the same indices as DataFrame2
    dataframe2  ->  A pandas DataFrame, with the same indices as DataFrame1
    label       ->  A string to replace the label of the new foldchange dataframe.
    
    =================================================================================================
    Returns: A new dataframe, containing the ratio of of the means from dataframe1/dataframe2
    
    =================================================================================================
    """
    # Check the input arguments
    dataframe1 = ah.check_type(dataframe1, pd.DataFrame,
                               error = "The dataframe1 argument is not a DataFrame")
    dataframe2 = ah.check_type(dataframe2, pd.DataFrame,
                               error = "The dataframe2 argument is not a DataFrame")
    label = ah.check_type(label, str,
                          error = "The label argument is not a string")
    # Onnce the input values are checked, continue.
    # First, determine whether the input dataframes have mean columns
    if "mean" not in list(dataframe1.columns.values):
        # If dataframe1 has no mean column, create one
        dataframe1 = df_row_mean([dataframe1])[0]
    if "mean" not in list(dataframe2.columns.values):
        # If dataframe2 has no mean column, create one
        dataframe2 = df_row_mean([dataframe2])[0]
    # Make a new dataframe as the ratio of the mean columns
    # This operation divides the elements of df1 by df2 using
    # the index values of the dataframe.
    new_df = pd.DataFrame(dataframe1['mean']/dataframe2['mean'])
    # and rename the column using the label
    new_df.rename(columns = {"mean": label}, inplace = True)
    # Once this is complete, return the new_df
    return new_df

def df_pairwise_foldchange(dataframes,
                           labels,
                           reverse_comp = True):
    """
    =================================================================================================
    df_pairwise_foldchange(dataframes, labels)
    
    This function wraps together the df_foldchange function, as well as properly applying labels.
    
    =================================================================================================
    Arguments:
    
    dataframes    ->  A list of pandas DataFrames
    labels        ->  A list of strings, which will label the columns of the output dataframe
    reverse_comp  ->  A boolean that determines the order of the comparisons, and thus the
                      order of the foldchanges.
    
    =================================================================================================
    Returns: A DataFrame of ratios of means between compared groups.
    
    =================================================================================================
    """
    # First, check the arguments
    dataframes = ah.check_type(dataframes, list,
                               error = "The dataframes is not a list of dataframes...")
    dataframes = [ah.check_type(dataframes[i], pd.DataFrame,
                                error = f"Element {i} of the dataframes argument is not a dataframe") for i in range(len(dataframes))]
    labels = ah.check_type(labels, list,
                           error = "The argument labels is not a list...")
    labels = [ah.check_type(labels[i], str,
                            error = f"Element {i} of the labels is not a string...") for i in range(len(labels))]
    reverse_comp = ah.check_type(reverse_comp, bool,
                                 error = f"The reverse_comp argument is not a boolean...")
    # Once the labels are checked, continue.
    # First, add the mean column to each of the dataframes.
    dataframes = df_row_mean(dataframes)
    # Then use the make_pairs() function to get a list
    # of all comparison index pairs, in reverse order.
    comparisons = list(gh.make_pairs(range(len(dataframes)),
                                     reverse=reverse_comp))
    # Initialize the new_df by getting the first foldchange list
    new_df = df_foldchange(dataframes[comparisons[0][0]],
                           dataframes[comparisons[0][1]],
                           label = labels[0])
    # Loop over the number of remaining comparisons
    for i in range(1,len(comparisons)):
        # and get a new foldchange dataframe with the current comparison
        new_fold = df_foldchange(dataframes[comparisons[i][0]],
                                 dataframes[comparisons[i][1]],
                                 label = labels[i])
        # Then join that dataframe with new_df
        new_df = new_df.join(new_fold, how = "outer")
    # At the end, return the new_df
    return new_df

def df_foldchange_signs(fc_df):
    """
    =================================================================================================
    df_foldchange_signs(fc_df):
    
    This function is meant to get the sign of the log transformed foldchange of a dataframe.
                  
    =================================================================================================
    Arguments:
    
    fc_df  ->  A pandas DataFrame of fold changes between to groups.
    
    =================================================================================================
    Returns: A DataFrame of 1/-1 that represent the sign of the log of the fold change in the
             input DataFrames.
    
    =================================================================================================
    """
    # Check the argument
    fc_df = ah.check_type(fc_df, pd.DataFrame,
                          error = "The fc_df argument is not a DataFrame...")
    # Apply the transformation to the dataframe and return the result
    return fc_df.apply(lambda x: np.log2(x)/abs(np.log2(x)))

def df_bin_row(index,
               condition_dfs,
               filterers,
               group_labels = None,
               comparisons = 'all'):
    """
    =================================================================================================
    df_bin_row(index, condition_dfs, filterers, group_labels, comparisons)
    
    This function is meant to take a row from a set of DataFrames and determine how the comparisons
    between the values in those rows should be handled. The results from this function are used
    in df_ttests(), and df_ttests() is the only place this function should be used.
    
    =================================================================================================
    Arguments:
    
    index          ->  An integer (or string) that determines the current row of the DataFrames
    condition_dfs  ->  A list of DataFrames that represent a specific condition
    filterers      ->  A list of Unix like wildcard strings that are used for filtering.
    group_labels   ->  Either None or a list of strings defining the groups being compared
    comparisons    ->  A string determining how to order the items for comparisons.
                       'all' 'highest' 'lowest'
                       
    NOTE: condition_dfs, filterers and group_labels lists should be index paired.
    
    ======================================================================#
    all_data = {"raw":  {key : {"data" : value} for key, value in update_with_rowflanks(data_dict).items()},
                "log":  {key : {"data" : value} for key, value in update_with_rowflanks(data_dict,
                                                                  rowflanks_kwargs = {"id_column" : "peptide",
                                                                  "assign_score" : "ascore",
                                                                  "flanks_given" : ["flank1", "flank2", "flank3"],
                                                                  "dupe_method" : "highest",
                                                                  "log_trans" : 2}).items()}}
    # and write the raw GCT file
    write_field_gct_files(args[2],
                          all_data,
                          field = 'data')
    # Then transform the AllRowFlanks objects into dataframes
    all_data = {"raw" : {key : {"data" : value} for key, value in update_rowflanks_df(all_data['raw']).items()},
                "log":  {key : {"data" : value} for key, value in update_rowflanks_df(all_data['log']).items()}}
    # and perform statistics on those dataframes
    stats_dict = {'raw' : { key : transform_stats(value["data"],
                                                  experiment_metadata[1],
                                                  parsing_strings,
                                                  fields = field_strings,
                                                  ttest_settings = df_t_settings) for key, value in all_data['raw'].items()},
                  "log" : { key : transform_stats(value["data"],
                                                  experiment_metadata[1],
                                                  parsing_strings,
                                                  fields = field_strings,
                                                  ttest_settings = df_t_settings) for key, value in all_data['log'].items()}}
    # Finally, write the Pvalue and Qvalue GCT files
    write_field_gct_files(args[2],
                          stats_dict,
                          field = None)
    return None
===========================
    Returns: A tuple of lists, where the zeroeth list is the data, and the first list is the
             combinations to be omitted
    
    =================================================================================================
    """
    # I'm not going to check the input arguments here, since this is only
    # used inside of the df_ttests() function.
    #
    # Initialize the list to hold the data from the rows.
    current_items = []
    # Loop over the number of DataFrames in the condition_dfs list
    for i in range(len(condition_dfs)):
        # If no group labels were provided
        if group_labels == None:
            # Then get and filter the headers using the filterers strings
            filt_heads = fnmatch.filter(list(condition_dfs[i]),filterers[i])
            # and attempt to get a numpy array of the row defined
            # by index.
            try:
                current_items.append(condition_dfs[i][filt_heads].loc[[index]].astype(float).to_numpy())
            # If that fails, then simply pass.
            except:
                pass
        # If group labels were provided
        elif group_labels != None:
            # Then get and filter the headers using the filterers strings
            filt_heads = fnmatch.filter(list(condition_dfs[i]),filterers[i])
            # And try to add both the group labels and the
            # numpy array of the row to the list
            try:
                current_items.append((group_labels[i],
                                      condition_dfs[i][filt_heads].loc[[index]].astype(float).to_numpy()))
            # If this fails, then simply pass.
            except:
                pass
    # If there is only one current_items or the list is empty
    if current_items == [] or len(current_items) == 1:
        # Return two empty lists
        return [],[]
    # Once we have some rows of the DataFrames, we need to
    # manipulate them such that Statistics can be performed.
    # First, we will filter out nans
    # Loop over the number of items in current_items
    for j in range(len(current_items)):
        # If group labels were procided, then there will be tuples
        # in the current_items list
        if group_labels != None:
            # Thus, keep the zeroeth element the same, and filter out
            # nan values from the numpy array in the first item position.
            current_items[j] = current_items[j][0], current_items[j][1][~np.isnan(current_items[j][1])]
        # Otherwise, the list only contains numpy arrays
        else:
            # So we can simply filter out nan values from the numpy array
            current_items[j] = current_items[j][~np.isnan(current_items[j])]
    # Once the nan values have been filtered, we can order the lists
    # to abide by the comparisons variable
    # If the comparisons variable is 'all'
    if comparisons in ["all","a"]:
        # Then no sorting is required, and no comparisons
        # are to be omitted from the statistics.
        return current_items, []
    # If the comparisons are 'lowest', then we only want comparisons
    # between all elements to the element with the lowest mean.
    elif comparisons in ["lowest mean", "l", "low", "lowest"]:
        # If group labels are provided
        if group_labels != None:
            # Then the numpy array is in the first slot of each tuple.
            # So sort the list based on the means of those elements
            # (list sorting is automatically lowest to highest)
            current_items = sorted(current_items, key=lambda x: x[1].mean())
            # And remove all items that have two or less items.
            current_items = [item for item in current_items if len(item[1]) >2]
        # OR, if group labels are not provided
        else:
            # Then filter out the arrays with less than two values or less
            current_items = [item for item in current_items if len(item) >2]
            # and sort the list on the mean of the numpy arrays
            current_items = sorted(current_items, key=lambda x: x.mean())
        # If two or less items are remaining after sorting/filtering
        #print(len(current_items))
        if len(current_items) <= 2:
            # then no combinations should be omitted
            omit_combs = []
        # Otherwise, we need to determine which combinations to omit.
        else:
            # Get all combinations of the elements of current_items that
            # do not include the zeroeth element, as it is the lowest mean
            holder = gh.make_pairs(current_items)
            omit_combs = [item for item in holder if current_items[0] not in item]
            holder = gh.make_pairs(current_items, reverse = True)
            omit_combs += [item for item in holder if current_items[0] not in item]
        # Onve the conditional statements have been determined,
        # return the current_items and the omit_combs lists.
        #print(current_items)
        return current_items, omit_combs
    # If the comparisons are 'highest', then we only want to compare
    # things to the list with the highest mean
    elif comparisons in ["highest mean", "h", "high", "highest"]:
        # If group labels were provided
        if group_labels != None:
            # Then the numpy array is in the first slot of each tuple.
            # So sort the list based on the means of those elements
            # (list sorting is automatically lowest to highest, so reverse)
            current_items = sorted(current_items, key=lambda x: x[1].mean(), reverse = True)
            # And remove all items that have two or less items.
            current_items = [item for item in current_items if len(item[1]) >2]
        # If group labels were not provided
        else:
            # Sort the list on the mean of the numpy arrays, highest to lowest
            current_items = sorted(current_items, key=lambda x:x.mean(), reverse = True)
            # Then filter out the arrays with less than two values or less
            current_items = [item for item in current_items if len(item) >2]
        # If two or less items remain after filtering,
        if len(current_items) <= 2:
            # Then there are no combinations to omit
            omit_combs = []
        # Otherwise, we need to determine which combinations to omit
        # during statistics
        else:
            # Get all combinations of the elements of current_items that
            # do not include the zeroeth element, as it is the highest mean
            holder = gh.make_pairs(current_items)
            omit_combs = [item for item in holder if current_items[0] not in item]
            holder = gh.make_pairs(current_items, reverse = True)
            omit_combs += [item for item in holder if current_items[0] not in item]
        # Once the conditional states have been determined,
        # return the current_items and the omit_combs lists.
        return current_items, omit_combs
        
def df_ttests(main_df,
              parsed_strs,
              identity_column = None,
              comparisons = "all",
              group_labels = None,
              filt_thresh = 3,
              pw_ttest_optargs = {},
              qvalue = False,
              storey_pi0 = 1):
    
    """
    =================================================================================================
    df_ttests(main_df, parsed_strs, comparisons, group_labels, filt_thresh, pw_ttest_optargs, qvalue)
    
    This function is meant to take a dataframe, parse it by conditions, and perform statstical tests
    between rows with the same indices (or the index specified by identity column)
    
    =================================================================================================
    Arguments:
    
    main_df           ->  A pandas DataFrame containing the data for which to perform statistics
    parsed_strs       ->  A list of lists of strings, which define how to parse the columns of the
                          DataFrame
    identity_column   ->  A string which defines a column that uniquely identifies rows
    comparisons       ->  A string defining how to perform comparisons.
                          "all", "highest", "lowest"
    group_labels      ->  A list of strings which are used to label the comparisons
    filt_thresh       ->  An integer, the minimum number of real data points to consider for comparisons
    pw_ttest_optargs  ->  A dictionary that defines the arguments of the ttest
    qvalue            ->  A boolean which determines whether or not to apply Storey's False Discovery
                          Rate estimation on all P-values calculated.
    storey_pi0        ->  The value used for pi0 in the Storey FDR algorithm. The default value of
                          1 is equivalent to the Benjamini-Hochberg method of FDR estimation. For
                          the pure Storey method, set this value to None.
    
    =================================================================================================
    Returns: A Pandas DataFrame containing the results of T-Tests, and possibly FDR estimation
    
    =================================================================================================
    """
    # This list contains all comparison types that are permitted
    valid_comparisons = ["all",
                         "lowest mean",
                         "highest mean",
                         "a",
                         "l",
                         "h",
                         "low",
                         "high",
                         "lowest",
                         "highest"]
    
    # First thing to do is make sure the user either passed parsed dataframes
    # or gave an iterable of strings to look for
    # And make sure the rest of the arguments are somewhat valid
    assert comparisons.lower() in valid_comparisons, f"Your comparisons value is not accepted. Try: {valid_comparisons}"
    assert type(identity_column) in [str, type(None)], "The idenitity column should be unset (None) or type str"
    assert type(pw_ttest_optargs) == dict, "The pairwise T-test optional arguments should be in a dictionary of key = arg_name, value = arg_value"
    assert qvalue in [True, False], "The qvalue argument should be a boolean"
    # First, parse the input DataFrame using the parsed_strs provided,
    # settingt he index to the identity column.
    condition_dfs, filt_strs = df_parser(main_df, 
                                         parsed_strs = parsed_strs, 
                                         id_col = identity_column)
    # Next, filter out rows that do not contain enough data points
    condition_dfs = df_filter_nans(*condition_dfs, 
                                   threshold = filt_thresh)
    # If group labels were provided
    if group_labels != None:
        # Then check to make sure there is one label per dataframe
        assert len(condition_dfs) == len(group_labels), f"You did not provide the correct number of group labels. Expecting {len(condition_dfs)}, but got {len(group_labels)}"
    # Set the total_ttests value, which will hold the results of
    # the statistical tests being conducted.
    total_ttests = None
    # Loop over the indices in the main dataframe
    for index in list(main_df.index.values):
        # Save the current comparison and the omittion labels
        # using df_bin_row
        current_comp, omit = df_bin_row(index,
                                        condition_dfs,
                                        filt_strs,
                                        group_labels = group_labels,
                                        comparisons = comparisons.lower())
        #print(omit)
        # If this list is empty,
        if current_comp == [] or len(current_comp) == 1:
            # Then continue, we do not want to try statistics with nothing
            continue
        # If the lsit was not empty, then perform pairwise T-tests
        ttest_results = hs.pairwise_t(*current_comp, omit_comb = omit, **pw_ttest_optargs)
        # If the identity column was set
        if identity_column != None:
            # Get the string identifying each row in the main dataframe
            name = main_df[f"{identity_column}"].loc[[index]].to_numpy()[0]
            # And repeat that string for the number of tests completed
            newcol = [name for _ in range(len(ttest_results))]
            # Then add that list as a column in ttest_results
            ttest_results[f"{identity_column}"] = newcol
        # Now that we've done T-tests and identified them
        # See whether this is the first test being done.
        # If so,
        if type(total_ttests) == type(None):
            # Then assign this result to the total_ttests variable
            total_ttests = ttest_results
        # Otherwise, this is not the first test beign done
        else:
            # So add this result to the total_ttests variables, ignoring the index
            total_ttests = pd.concat([total_ttests,ttest_results], ignore_index=True)
    # Once all T-tests and their results are completed and recorded
    # we can determine whether or not to apply FDR estimations.
    # If qvalue is False
    if qvalue == False:
        # Then simply return the T-test results
        return total_ttests
    # Otherwise, qvalue is True, and we should apply FDR estimations.
    else:
        # This step requires information regarding group labels and identity columns.
        # If both identity columns and group labels are set
        if identity_column != None and group_labels != None:
            # Then apply the Storey FDR algorithm using the pvalue column
            # the group labels, the id_column, and carry over the T-test metadata.
            return hs.storey(total_ttests[['pvalue']], 
                             groups = total_ttests[[f"{identity_column}", "Group 1", "Group 2"]],
                             test = total_ttests["Test"],
                             pi0 = storey_pi0)
        # If the identity column is not set, but the group labels are set
        elif identity_column == None and group_labels != None:
            # Then apply Storey FDR estimation using the pvalue column,
            # the groups, and carry over the T-test metadata.
            return hs.storey(total_ttests[['pvalue']], 
                             groups = total_ttests[["Group 1", "Group 2"]],
                             test = total_ttests["Test"],
                             pi0 = storey_pi0)
        # If the identity column was provided but no group lables are set
        elif identity_column != None and group_labels == None:
            # Them apply Storey FDR estimation using the pvalue column,
            # the identity column, and carry over the T-test metadata
            return hs.storey(total_ttests[['pvalue']], 
                             groups = total_ttests[[f"{identity_column}"]],
                             test = total_ttests["Test"],
                             pi0 = storey_pi0)
        # If neither the ID column onr the group labels are provided,
        elif identity_column == None and group_labels == None:
            # Then apply the Storey algorithm using only the P-value column
            # and carrying over the T-test metadata.
            return hs.storey(total_ttests[['pvalue']], 
                             test = total_ttests["Test"],
                             pi0 = storey_pi0)

def df_ttest_on_dict(data_dict,
                     condition_strs,
                     parsing_strs,
                     fields = ["pvalue", "qvalue"],
                     ttest_settings = {"identity_column"  : "peptide",
                                       "comparisons"      : "all",
                                       "pw_ttest_optargs" : {"t_type" : "welch"},
                                       "qvalue"           : True,
                                       "storey_pi0"       : 1},
                     write = True,
                     path = "",
                     filename = "a_file",
                     parse = True):
    """
    =================================================================================================
    df_ttest_on_dict(data_dict, condition_strs, parsing_strs, **kwargs)
    
    =================================================================================================
    Arguments:
    
    data_dict       ->  A dictionary containing key/value pairs where the values are DataFrames
    condition_strs  ->  A list of strings describing the labels for groups
    parsing_strs    ->  A list of lists of strings, where each sublist describes how to parse the
                        DataFrame
    fields          ->  A list of strings describing the statistics field to parse. Should only
                        use P-values and Q-values
    ttest_settings  ->  A dictionary that defines the arguments of the ttest
    write           ->  A boolean value that determines whether or not to write the statistics
                        DataFrame to a file.
    path            ->  A string describing the path to the output file directory. Not used if
                        the write boolean is False
    filename        ->  A string describing the name of the output file. Not used if the write
                        boolean is False
    
    =================================================================================================
    Returns: A dictionary of the outputs from statistical tests on the dictionaries.
    
    =================================================================================================
    """
    # Initialize the newdict dictionary, which will hold the
    # outputs from statistical tests
    newdict = {}
    # Loop over the keys and values in the input data dictionary
    for key, value in data_dict.items():
        # Initialize a dictionary for each key in the data_dict
        newdict[key] = {}
        # Set saved to zero, as this will
        # hold the parsing string for this dataframe
        saved = 0
        # Loop over the number of parsing string lists
        for i in range(len(parsing_strs)):
            # If the current key is in the parsing string
            if key in parsing_strs[i][0]:
                # Set saved to the index
                saved = i
                # and break the loop
                break
        # Next, perform T-tests using the df_ttests wrapper
        stats_df = df_ttests(value,
                             parsed_strs = parsing_strs[saved],
                             group_labels = condition_strs,
                             **ttest_settings)
        # If the write boolean is True
        if write:
            # and the output filepath does not exist
            if not os.path.exists(os.path.join(path,key)):
                # First make the path
                os.makedirs(os.path.join(path,key))
            # And write the statistics dataframe to the file, tab separated.
            stats_df.to_csv(os.path.join(path, key, f"{filename}_{key}.txt"), sep = "\t")
        
        if parse:
            # Loop over the field strings in fields
            for field in fields:
                # And add an element to the newdict[key] dictionary with the
                # field key and the value as the flipped dataframe.
                newdict[key][f"{field}"] = df_parse_fieldvalues(stats_df,
                                                            field = field,
                                                            identity = ttest_settings["identity_column"])
        else:
            newdict[key]["stats"] = stats_df
    # At the end, return the newdict.
    return newdict


#
#
######################################################################################################
#
#     Dealing with Object Type Transformations

def df_to_dict(*args, 
               label_loc = 0, 
               to_type = float,
               keep_headers = False):
    """
    =================================================================================================
    df_to_dict(*args, label_loc, to_type, keep_headers)
                  
    =================================================================================================
    Arguments:
    
    args          ->  An arbitrary number of pandas DataFrames
    label_loc     ->  An integer that determines where the location of the label
    to_type       ->  The type to change the values to
    keep_headers  ->  A boolean determining whether or not to keep the headers of the dataframes
    
    =================================================================================================
    Returns: A list of dictionaries, where each dictionary contains the elements from a Pandas
             DataFrame
    
    =================================================================================================
    """
    # Check the input argumetns
    assert type(label_loc) == int, "label_loc shoud be an integer"
    assert to_type in [float, int, str], "The to_type specified is invalid. Takes: float/int/str"
    # Initialize the list to hold dictionaries returned
    returns = []
    # Loop over the input dataframes
    for arg in args:
        # And check that each item is, indeed a DataFrame
        assert type(arg) == type(pd.DataFrame()), "Inputs should be dataframes"
        # For each dataframe, initialize a dictionary. If we
        # want to keep the headers,
        if keep_headers == True:
            # Initialize the dictionary using the headers 
            newdict = {"headers" : list(arg.column.values)}
        # Otherwise
        else:
            # just intiialize an empty dictioanry
            newdict = {}
        # Loop over the lists in the numpified dataframe
        for sub_list in [list(item) for item in arg.to_numpy()]: 
            # Get the key for the dictionary
            label = sub_list.pop(label_loc)
            # Add the list to the dictionary, transforming the values based on
            # the given type
            newdict[label] = [to_type(value) for value in sub_list ] 
        # Add the dictionary to the returns list
        returns.append(newdict)
    # Return the returns list
    return returns

def df_to_lists(a_dataframe):
    """
    =================================================================================================
    df_to_lists(a_dataframe)
                  
    =================================================================================================
    Arguments:
    
    a_dataframe  ->  A pandas DataFrame
    
    =================================================================================================
    Returns: The contents of the pandas DataFrame as a list of lists
    
    =================================================================================================
    """
    # Get the columns of the dataframe as a list
    cols = list(a_dataframe.columns.values)
    # Get the rows as a DataFrame as a list of lists
    rows = [list(row) for row in a_dataframe.to_numpy()]
    # Return the column headers and the rows
    return [cols, *rows]

#
#
######################################################################################################
```
