## Supporting Folder 2 Data Analysis for "Mouse primary T cell phosphotyrosine proteomics enabled by BOOST": stats_helpers.py

```
"""
=================================================================================================
Kenneth P. Callahan

9 July 2021
                  
=================================================================================================
Python >= 3.8.5

homebrew_stats.py

This module is meant to help with general statistical functions. Currently, there is only
a small number of statistics options supported, but I suspect this will grow in the future.

Currently supported:

    FDR Estimation (Storey)
    T-tests

=================================================================================================
Dependencies:

 PACKAGE     VERSION
  Pandas  ->  1.2.3
  Numpy   ->  1.20.1
  SciPy   ->  1.7.2

=================================================================================================
"""
import os
print(f"Loading the module: helpers.{os.path.basename(__file__)}\n")
#####################################################################################################################
#
#   Importables

import fnmatch                                        # Unix-like string searching

import pandas as pd                                   # General use for data
import numpy as np                                    # General use for data

from math import sqrt
import copy

import scipy
from scipy.stats import f, t
from scipy.stats import studentized_range as q
 
from scipy.interpolate import splrep, splev           # Used for Storey Q-value estimation, fitting cubic spline
from scipy.interpolate import UnivariateSpline

from . import general_helpers as gh

print(f"numpy        {np.__version__}")
print(f"scipy         {scipy.__version__}")
print(f"pandas        {pd.__version__}\n")

#
#
#####################################################################################################################
#
#     Miscellaneous Functions

def filter_nans(data,
                threshold = 3, 
                threshold_type = "data"):
    """
    =================================================================================================
    filter_nans(data, threshold, threshold_type)
    
    This function is meant to filter out the nan values from a list, based on the input arguments.
                  
    =================================================================================================
    Arguments:
    
    data            ->  A list (or iterable) of data points. The points are assumed to be numbers.
    threshold       ->  An integer describing the minimum value requirement.
    threshold_type  ->  A string describing how the threshold integer will be applied. 
                        "on_data" "on_nan"
    
    =================================================================================================
    Returns: The filtered list, or an empty list if the threshold requirements were not met.
    
    =================================================================================================
    """
    # Make sure the user gave a valid thresholding option
    assert threshold_type.lower() in ["data", 
                                      "on_data", 
                                      "on data", 
                                      "nan", 
                                      "on_nan", 
                                      "on nan"], "Threshold is either relative to NaN or data."
    assert type(data) == list, "The data should be in a list"
    # Filter NaNs, as they do not equal themselves
    filtered = [val for val in data if val == val]
    # Keep data if there are at least <threshold> data points
    if threshold_type.lower() in ["data", "on_data", "on data"]:
        if len(filtered) >= threshold:
            return filtered
        else:
            return []
    # Keep data if there are no more than <threshold> nans
    elif threshold_type.lower() in ["nan", "on_nan", "on nan"]:
        if len(data) - len(filtered) <= threshold:
            return filtered
        else:
            return []

def filter_nan_dict(data,
                    threshold = 3, 
                    threshold_type = "data"):
    """
    =================================================================================================
    filter_nan_dict(data, threshold, thershold_type)
    
    This function is meant to filter out nan values from the list-values in a dictionary. This
    function uses filter_nans() to filter lists.
                  
    =================================================================================================
    Arguments:
    
    data            ->  A dictionary of lists of data points. The points are assumed to be numbers.
    threshold       ->  An integer describing the minimum value requirement.
    threshold_type  ->  A string describing how the threshold integer will be applied. 
                        "on_data" "on_nan"
    
    =================================================================================================
    Returns: A dictionary where all values have been filtered from the list-values.
    
    =================================================================================================
    """
    # Make sure a dictionary is given as input
    assert type(data) == dict, "The data should be in a dictionary"
    # Initialize the new dictionary
    filtered_dict = {}
    # Loop over the keys/values in the dictionary
    for key, value in data.items():
        # Filter the nans
        filt_list = filter_nans(value)
        # If the list is not empty
        if filt_list != []:
            # then add it to the dictionary
            filtered_dict[key] = filt_list
        # IF the list is empty, it will not be added to the dictionary
    # Return the filtered dictionary.
    return filtered_dict

def count_list(dataset): 
    '''
    Given a dataset, count the occurence of
    each data point and return them as a dictionary. 
    '''
    # First we create a dictionary to hold the counts. 
    # We then loop over the elements of the dataset
    # and attempt to check the dictionary keys for them.
    # If the element has appeard, we add one to the count
    # and if the element is not in the dictionary, we 
    # add a key to the dictionary with that elements name 
    # and initialize it to 1 (since we have seen it once). 
    # At the end, return the dictionary. 
    dic = {}                          # Create the empty dictionary
    for element in dataset:           # Loop over the elemnts in the dataset
        try:                          # Attempt
            dic[str(element)] +=  1   # Look for the key of the element, add one to count
        except:                       # Otherwise 
            dic[str(element)] = 1     # Add a key to the dicitonary with value 1
    return dic                        # Return the dictionary


#
#
#######################################################################################################
#
#    General Statistical Functions

def mean(dataset, filter_nans = True, threshold = 1):
    '''
    Given a dataset, return the average value.
    '''
    if filter_nans:
        data = [d for d in dataset if d == d]
    else:
        data = [d for d in dataset]
    if len(data) >= threshold:
        return sum(data) / len(data)
    else:
        return float("nan")

def median(dataset, filter_nans = True):
    '''
    Given a dataset, return the median value.
    '''
    # The calculation for median depends on whether the 
    # number of datapoints in the set is even or odd. 
    # If the number of datapoints is even, then we need to 
    # find the average of the two middle numbers. If the 
    # number of datapoints is odd, then we simply need
    # to find the middle one. They also need to be sorted.
    if filter_nans:
        data = sorted([d for d in dataset if d == d])
    else:
        data = sorted(dataset)
    if len(data) == 0:
        return float("nan")
    elif len(data) == 1:
        return data[0]
    elif len(data) % 2 == 0:                            # if the dataset is even
        index = len(data) // 2                        # get the middle data point
        med = (data[index] + data[index -1]) / 2      # average the middle two points
        return med                                    # return this value
    elif len(data) % 2 == 1:                          # if the dataset is odd
        index = len(data) // 2                        # get the middle point
        return data[index]                            # return the middle point

def grand_mean(*data):
    all_data = gh.unpack_list(data)
    return sum(all_data)/len(all_data)

def demean(data, grand = False):
    if not grand:
        return [d - mean(data) for d in data]
    else:
        return [mean(d) - grand_mean(data) for d in data]

def variance(dataset, correction = 1, threshold = 2): 
    '''
    Given a dataset, calculate the variance of the 
    parent population of the dataset. 
    '''
    # Calculate the data without the mean, square
    # all of the elements, then return the sum of 
    # those squares divided by the number of 
    # datapoints minus 1. 
    meanless = demean(dataset)          # Remove the mean from the data
    if len(meanless) < threshold:
        return float("nan")
    squares = [x**2 for x in meanless]       # Square all of the meanless datapoints
    return sum(squares) / (len(dataset) - correction) # return the sum of the squares divided by n-1

def standard_deviation(data, correction = 1, threshold = 2):
    return sqrt(variance(data, correction = correction, threshold = threshold))

def sem(data, correction = 1):
    if len(data) < 2:
        return float("nan")
    return standard_deviation(data, correction=correction) / sqrt(len(data))
    
def sum_of_squares(vector):
    return sum([v**2 for v in vector])

def var_within(*data, correction = 1):
    variances = [variance(d, correction = correction) for d in data]
    return mean(variances)

def var_between(*data, correction = 1):
    means = [mean(d) for d in data]
    return len(data[0]) * variance(means, correction = correction)

def total_variation(*data):
    return sum_of_squares(demean(data, grand=True))

def sos_between(*data):
    demeaned = demean(data, grand = True)
    lens = [len(data[i]) for i in range(len(data))]
    return sum([lens[i]*demeaned[i]**2 for i in range(len(data))])

def ms_between(*data):
    return sos_between(*data) / (len(data)-1)

def sos_within(*data, correction = 1):
    deg_frees = [len(d) - 1 for d in data]
    var = [variance(d, correction = correction) for d in data]
    return sum([deg_frees[i] * var[i] for i in range(len(data))])

def ms_within(*data, correction = 1):
    deg_free = sum([len(d) for d in data]) - len(data)
    return sos_within(*data, correction = correction)/deg_free

def mode(dataset):
    '''
    Given a dataset, returns the most frequent value
    and how many times that value appears
    '''
    # First, we count all of the elements and arrange them in 
    # a dictionary. Then, we create a sorted list of tuples 
    # from the key value pairs. Initialize 'pair', to hold
    # the key value pair of the highest value, and an empty
    # list to hold any of the pairs that tie. We then loop
    # over the sorted lists keys and values, and look for the 
    # highest counts. We return the highest count from the 
    # dictionary, or the highest counts if there were any ties.
    counted = count_list(dataset)      # Count the elements of the dataset
    sort = sorted(counted.items())     # Sort the numbers and occurences
    pair = 'hold', 0                   # Initialize the pair
    ties = []                          # Initialize the tie list
    for key, value in sort:            # Loop over key, value in sorted dictionary
        if value > pair[1]:            # If the value is greater than the pair
            pair = key, value          # Re assign the pair to the current one
            ties = []                  # Reset the tie list
        elif value == pair[1]:         # If the value is equal to the current value
            ties.append((key, value))  # Append the new key, value pair to the list
    ties.append(pair)                  # After, append the pair to the list
    svar = sorted(ties)                # Sort the list of ties
    if len(ties) > 1:                  # If there are any ties, 
        return svar                    # Return the sorted list of ties 
    elif len(ties) == 1:               # If there are no ties
        return pair                    # Return the highest value

def quantile(dataset, percentage):
    '''
    Given a dataset and a pecentage, the function returns
    the value under which the given percentage of the data
    lies
    '''
    # First, sort the dataset, then find the index at the 
    # given percentage of the list. Then, return the 
    # value of the dataset at that index. 
    dataset = sorted(dataset)               # Sort the dataset
    index = int(percentage * len(dataset))  # Get the index of element percentage 
    return dataset[index]                   # return the element at the index  

def interquantile_range(dataset, per_1, per_2):
    '''
    Given a dataset and two percentages that define 
    a range of the dataset, find the range of the 
    elements between those elements. 
    '''
    dataset = sorted(dataset)
    return quantile(dataset, per_2) - quantile(dataset, per_1)  

def data_range(dataset):
    '''
    Given a dataset, return the range of the elements.
    '''
    dataset = sorted(dataset)
    return dataset[-1] - dataset[1]

def dot_product(data_1, data_2):
    
    '''
    Given two datasets of equal length, return the 
    dot product. 
    '''
    # First, we make sure that the lists are the same size,
    # Then we loop over the length of the lists, and sum the 
    # product of the corresponding elements of each list. 
    # Then, that sum is returned. 
    assert len(data_1) == len(data_2), "These lists are not the same length"
    sum_total = 0                            # Initialize the sum
    for i in range(len(data_1)):             # Loop over the size of the list
        sum_total += data_1[i] * data_2[i]   # Add to the sum the product of the datapoints in 1 and 2
    return sum_total                         # Return the sum

def covariance(data_1, data_2):
    
    '''
    Given two datasets, calculate the covariance between them
    '''
    
    n = len(data_1)
    return dot_product(demean(data_1),demean(data_2)) / (n-1)

def correlation(data_1, data_2):
    '''
    Given two datasets, calculate the correlation between them. 
    '''
    return covariance(data_1, data_2) / (standard_deviation(data_1) * standard_deviation(data_2))

def vector_sum(vectors):
    '''
    Given a set of vectors, return a vector which contains the 
    sum of the ith elements from each vector in index i
    '''
    for i in range(len(vectors)-1):
        assert len(vectors[i]) == len(vectors[i+1]), 'Vectors are not the same length'
    return [sum(vector[i] for vector in vectors)
           for i in range(len(vectors[0]))]

assert vector_sum([[1,2],[2,3],[3,4]]) == [6,9]

def scalar_multiply(scalar, vector):
    '''
    Given a scalar and a vector (list), return a vector
    where each component is multiplied by the scalar. 
    '''
    return [scalar * var for var in vector]

assert scalar_multiply(3, [1,2,3,4]) == [3,6,9,12]

def vector_subtract(vectors):
    '''
    Given a set of vectors, return the difference between
    the vectors, in index order. 
    
    This will look like:
    vectors[0] - vectors[1] - ... - vectors[n] = result
    '''
    for i in range(len(vectors)-1):
        assert len(vectors[i]) == len(vectors[i+1]), 'Vectors are not the same length'
    pass_count = 0
    result = vectors[0]
    for column in vectors:
        if column == result and pass_count == 0:
            pass_count += 1
            pass
        else:
            for i in range(len(result)):
                result[i] += -column[i]
            pass_count += 1
    return result      

assert vector_subtract([[1,2,3], [3,4,5]]) == [-2,-2,-2]

def vector_mean(vectors):
    '''
    Given a list of lists (which correspond to vectors, where
    each element of the vector represents a different variable)
    return the vector mean of the vectors (add each the vectors
    component-wise, divide each sum by the number of vectors)
    '''
    n = len(vectors)
    return scalar_multiply(1/n, vector_sum(vectors))

assert vector_mean([[1,2],[2,3],[3,4]]) == [2,3]

def scale(data_1):
    '''
    Given a set of datapoint sets, return the mean of each
    dataset and the standard deviation for each set. 
    
    Data points should be give as 
    x = [[x1_1, x2_1,..., xn_1],...,[x1_n, x2_n,..., xn_n]]
    if the data are not in this format, but are in the format
    x = [[x1_1, x1_2,..., x1_n],...,[xn_1, xn_2,..., xn_n]]
    apply the function reformat_starts(*args) to the data. 
    '''
    # First, we make sure that all of the data points
    # given are the same length, then save the size of
    # the datasets as n. We then calculate the vector 
    # mean of the data, as well as the standard deviations
    # of each data type. Then the means and SDs are returned
    for q in range(len(data_1) -1):
        assert len(data_1[q]) == len(data_1[q+1]), 'Data lists are different sizes'
    n = len(data_1[0])
    means = vector_mean(data_1)
    s_deviations = [standard_deviation([vector[i] 
                                        for vector in data_1])
                   for i in range(n)]
    return means, s_deviations

t_vectors = [[-3, -1, 1], [-1, 0, 1], [1, 1, 1]]
t_means, t_stdevs = scale(t_vectors)
assert t_means == [-1, 0, 1]
assert t_stdevs == [2, 1, 0]

def rescale(data_1):
    '''
    Given a set of data sets, return a list of the
    data rescaled, based on the means and the 
    standard deviations of the data. 
    '''
    # First, we calculate the mean and standard deviations
    # of the data, and save the size of the datasets as 
    # n. We then copy each of the vectors to the list
    # rescaled. Next, we loop over the vectors in rescaled,
    # and loop over the size of the datasets, and 
    # scale each term in v[i] based on the mean and SD
    means, s_deviations = scale(data_1)
    n = len(data_1[0])
    rescaled = [v[:] for v in data_1]
    for v in rescaled:
        for i in range(n):
            if s_deviations[i] > 0:
                v[i] = (v[i] - means[i]) / s_deviations[i]
    return rescaled

t2_means, t2_stdevs = scale(rescale(t_vectors))
assert t2_means == [0, 0, 1]
assert t2_stdevs == [1, 1, 0]

def unscaled(scaled_data_1, data_1, coefficients = False):
    '''
    Given a set of scaled datapoints, the original datapoints,
    and a truthy value for whether we are unscaling coefficients
    of regression, return the unscaled data points. 
    '''
    # This is basically 'rescale' in reverse, with the
    # condition of if we are unscaling coefficients. If 
    # we are unscaling coefficients, we subtract from the 
    # alpha term (v[0]) all elements in the form 
    # v[j] * mean[j] / s_deviations[j] (as described in 
    # Data Science from Scratch, 2nd Edition in Chapter 
    # 16, Logistic Regression). OTherwise, all coefficient 
    # are divided by the standard deviation term of the 
    # corresponding data. 
    n = len(data_1[0])
    means, s_deviations = scale(data_1)
    unscaled = [v[:] for v in scaled_data_1]
    for v in unscaled:
        for i in range(n):
            if coefficients == False:
                v[i] = v[i]*s_deviations[i] + means[i]
            elif coefficients == True:
                if i == 0:
                    for j in range(1,n):
                        if s_deviations[j] > 0:
                            v[0] = v[0] - (v[j]*means[j])/s_deviations[j]
                        else:
                            v[0] = v[0] - v[j]
                elif i != 0: 
                    if s_deviations[i] > 0: 
                        v[i] = v[i] / s_deviations[i]
                    else:
                        pass
    return unscaled

assert unscaled(rescale(t_vectors), t_vectors) == t_vectors

# Simple Linear Regression

def prediction(alpha, beta, variable):
    
    '''
    Given a constant term (alpha), a coefficient (beta) and 
    a variable value (variable), predict the output.
    '''
    
    return alpha + beta * variable

def error(alpha, beta, variable, output):
    
    '''
    Given a constant term (alpha), a coefficient (beta) and 
    a variable value (variable) and the outcome of the 
    experiment (output), calculate the error of the prediction
    '''
    
    return prediction(alpha, beta, variable) - output

def sum_of_squerrors(alpha, beta, variable, output):
    
    '''
    Given a constant term (alpha), a coefficient (beta) and 
    a list of variable values (variable) and a list of the 
    outcomes of the experiment (output), return the sum of
    the squared errors.
    '''
    
    return sum([error(alpha, beta, variable[i], output[i])**2
                for i in range(len(variable))])

def r_squared(alpha, beta, data_1, data_2):
    
    '''
    Given an estimate for a constant term (alpha) and a coefficient
    (beta), as well as the datasets used to make those estimates
    (data_1 and data_2) for a least squared fit model, calculate
    the r squared value for how good of a fit the line is for the
    data. 
    '''
    
    return 1.0 - (sum_of_squerrors(alpha, beta, data_1, data_2)/
                 sum_of_squares(demean(data_2)))

def least_squares_fit(data_1, data_2):
    
    '''
    Given sets of data, calculate the estimate of a constant
    term (alpha) and a coefficient (beta) for the least 
    squares fit model
    
    y = alpha + beta * x
    '''
    
    beta = correlation(data_1, data_2) * standard_deviation(data_2) / standard_deviation(data_1)
    alpha = mean(data_2) - beta * mean(data_1)
    r2 = r_squared(alpha, beta, data_1, data_2)
    
    return alpha, beta, r2


#
#
#####################################################################################################################
#
#   Q-Value Estimation Algorithms

#### Storey

def storey_check_groups(groups):
    
    """
    =================================================================================================
    storey_check_groups(groups)
    
    This function is meant to check the groups argument input into the function storey()
    
    =================================================================================================
    Arguments:
    
    groups  ->  Either a list, tuple, pandas DataFrame, or a numpy array that describes the groups
                in Storey FDR estimation.
    
    =================================================================================================
    Returns: A list of lists that describe the groups used in Storey FDR estimation
    
    =================================================================================================
    """
    # If the input groups are a pandas DataFrame
    if type(groups) == type(pd.DataFrame()):
        # Then convert the groups into a transposed numpy array
        groups = groups.to_numpy().transpose()
        # and use list comprehension to reformat the groups into
        # a list of pairs.
        groups = [[groups[j][i] for j in range(len(groups))] 
                  for i in range(len(groups[0])) ]
    # If the input groups are a lsit
    elif type(groups) == list:
        # Then loop over the number of lists
        for i in range(len(groups)):
            # If the type of the input groups are not
            # a list, tuple or array
            if type(groups[i]) not in [list, tuple, type(np.array([]))]:
                # Then list the element
                groups[i] = list(groups[i])
            # Otherwise
            else:
                # Just keep the list the same
                groups[i] = groups[i]
    # If the groups were given as a tuple
    elif type(groups) == tuple:
        # Then turn the groups into a lsit
        groups = list(groups)
        # and loop over the number of items in the groups
        for i in range(len(groups)):
            # and if the element is not a list, tuple, array,
            if type(groups[i]) not in [list, tuple, type(np.array([]))]:
                # then list the element and save it
                groups[i] = list(groups[i])
            # Otherwsie,
            else:
                # Keep the element the same
                groups[i] = groups[i]
    # If the input is a numpy array
    elif type(groups) == type(np.array([])):
        # Then use list comprehension to format the groups list.
        # Assumes the groups have been transposed in this instance
        groups = [[groups[j][i] for j in range(len(groups))] for i in range(len(groups[0]))]
    # At then end, return the groups list.
    return groups

def storey_check_test(test):
    
    """
    =================================================================================================
    storey_check_test(test)
    
    This function is meant to check and format the T-test type from the inputs. The function works
    almost exactly like storey_check_groups()
    
    =================================================================================================
    Arguments:
    
    test  ->  A list, array, tuple, or array describing the T-test types used for each P-value.
    
    =================================================================================================
    Returns: A properly formatted list
    
    =================================================================================================
    """
    
    # If the groups are dataframes, make the input into a list  of two-lists
    if type(test) == type(pd.DataFrame()) or type(test) == type(pd.Series([])):
        # If the input is a series or DataFrame object
        # then attempt to list it
        try:
            test = list(test)
        except:
            raise ValueError("The test dataframe is incorrectly formatted. Try:  df_name['Test']")
    # If the input type is a list
    elif type(test) == list:
        # then iterate through each element of the list
        for i in range(len(test)):
            # and if any elements are not strings, then string them
            if type(test[i]) != str:
                test[i] = str(test[i])
            else:
                test[i] = test[i]
    # If the input type is a tuple
    elif type(test) == tuple:
        # then list the test
        test = list(test)
        # and loop over the elements of test
        for i in range(len(test)):
            # If any elements are not strings, then string them
            if type(test[i]) != str:
                test[i] = str(test[i])
            else:
                test[i] = test[i]
    # If the input is a numpy araray
    elif type(test) == type(np.array([])):
        # then use list comprehension to str all elements of the array
        test = [str(test[i]) for i in range(len(test))]
    # And at the end, return the test, reformatted
    return test

def storey_check_args(pvals,
                      groups,
                      test):
    """
    =================================================================================================
    storey_check_args(pvals, groups, test)
    
    This function is meant to check the arguments passed into the storey() function, and ensures that
    FDR estimation may proceed without conflict.
    
    =================================================================================================
    Arguments:
    
    pvals   ->  A list, numpy array, dataframe, tuple of P-values
    groups  ->  A list, numpy array, dataframe, tuple of group labels, index matched to the pvalues
    test    ->  A list, numpy array, dataframe, tuple of T-tests used for calculating P-values,
                index matched to the pvals argument.
    
    =================================================================================================
    Returns: The pvalues, g_checker boolean (group checker) and the t_checker boolean (test checker)
    
    =================================================================================================
    """
    # First, type-check the inputs 
    assert type(pvals) in [list, 
                           type(np.array([] ,dtype = float)),
                           type(pd.Series()),
                           type(pd.DataFrame())], "The p-values should be given as a list or a numpy array"
    assert type(groups) in [type(None),
                            list, 
                            tuple,
                            type(np.array([])),
                            type(pd.Series()),
                            type(pd.DataFrame())], "The p-values should be given as a list, tuple, numpy array, series or dataframe."
    # Then, if the pvals were a series or DataFrame object
    if type(pvals) == type(pd.Series()) or type(pvals) == type(pd.DataFrame()):
        # Turn them into numpy arrays and transpose them
        pvals = pvals.to_numpy().transpose()
        # Then, if the length of pvals is not 1, then raise an error
        if len(pvals) != 1:
            raise ValueError("The DataFrame or Series input has more than one dimension...")
        # Otherwise, pvals are the zeroeth element of the array
        else:
            pvals = pvals[0]
    # Next, check the groups. If somethign other than NoneType
    # was provided
    if type(groups) in [list, 
                        tuple,
                        type(np.array([])),
                        type(pd.Series()),
                        type(pd.Series()),
                        type(pd.DataFrame())]:
        # Then set g_checker to True, so we will check groups
        g_checker = True
    # Otherwise, set g_checker to False, as we do not need to check groups
    else:
        g_checker = False
    # If the test is a proper typed object
    if type(test) in [list, 
                      tuple,
                      type(np.array([])),
                      type(pd.Series()),
                      type(pd.Series()),
                      type(pd.DataFrame())]:
        # Then set t_checker to True, as we need to check the test
        t_checker = True
    # Otherwise, set t_checker to False, as we do not need to check test
    else:
        t_checker = False
    # and return pvals, g_chekcer and t_checker
    return pvals, g_checker, t_checker

def storey_make_id_dict(pvals, 
                        groups,
                        test,
                        g_checker,
                        t_checker):
    
    """
    =================================================================================================
    storey_make_id_dict(pvals, groups, test, g_checker, t_checker)
    
    This function is meant to take all relevant arguments to storey() and perform checking
    operations on each of those inputs.
    
    =================================================================================================
    Arguments:
    
    For more information on pvals, groups, test, refer to storey_check_args().
    
    g_checker  ->  A boolean, determines whether a group is in need of checking
    t_checker  ->  A boolean, determines whether a test is in need of checking
    
    =================================================================================================
    Returns: A dictionary based on the pvals argument, and the groups and test arguments checked.
    
    =================================================================================================
    """
    # Initialize the idenities dictionary
    identities = {}
    # Then proceed with making the identity dictionary
    # If groups are given and tests are not given
    if g_checker != False and t_checker == False:
        # Make sure the groups are in the correct format.
        # Otherwise, terminate gracefully.
        groups = storey_check_groups(groups)
        # If there are not enough group labels given for all the pvals,
        if len(groups) != len(pvals): 
            # Just proceed without group labels
            print("Each p-value should have a corresponding label/group tuple, proceeding without labels")
            # And make a dict of lists with key = pval, value = [position]
            for i in range(len(pvals)):
                identities[f"{i}?{pvals[i]}"] = [i] 
        # Otherwise, use the labels as the keys of the dictionary
        else:
            # make a dict of lists with key = pval, value = [position, label]
            for i in range(len(pvals)):
                identities[f"{i}?{pvals[i]}"] = [i, *groups[i]]
    # If no groups were provided but tests were provieded
    elif g_checker == False and t_checker != False:
        # Make sure the tests are in the right format. Otherwise, terminate gracefully
        test = storey_check_test(test)
        # If there are not enough tests given for all the pvals,
        if len(test) != len(pvals): 
            # Just proceed without labels
            print("Each p-value should have a corresponding test, proceeding without test identifier")
            # And make a dict of lists with key = pval, value = [position]
            for i in range(len(pvals)):
                identities[f"{i}?{pvals[i]}"] = [i]
        # Otherwise, use the tests as the keys of the dictionary
        else:
            # make a dict of lists with key = pval, value = [position, label]
            for i in range(len(pvals)):
                identities[f"{i}?{pvals[i]}"] = [i, test[i]]
    # If both tests and groups are provided
    elif g_checker != False and t_checker != False:
        # Make sure they're in the right format. Otherwise, terminate gracefully
        groups = storey_check_groups(groups)
        test = storey_check_test(test)
        # If there are not enough labels given for all the pvals,
        if len(groups) != len(pvals) and len(test) != len(pvals): 
            # Just proceed without labels
            print("Each p-value should have a corresponding label/group tuple and test, proceeding without labels and test identifiers")
            # And make a dict of lists with key = pval, value = [position]
            for i in range(len(pvals)):
                identities[f"{i}?{pvals[i]}"] = [i]
        # Otherwise, use the labels as the keys of the dictionary
        elif len(groups) != len(pvals) and len(test) == len(pvals):
            print("Each p-value should have a corresponding test, proceeding without test identifiers")
            for i in range(len(pvals)):
                identities[f"{i}?{pvals[i]}"] = [i, *groups[i]]
        elif len(groups) == len(pvals) and len(test) != len(pvals):
            print("Each p-value should have a corresponding label/group tuple, proceeding without labels")
            for i in range(len(pvals)):
                identities[f"{i}?{pvals[i]}"] = [i, test[i]]
        else:
            # make a dict of lists with key = pval, value = [position, label]
            for i in range(len(pvals)):
                identities[f"{i}?{pvals[i]}"] = [i, *groups[i], test[i]]
    # If no labels are given, then just make the identities dictionary
    else:
        # by looping over the pvals
        for i in range(len(pvals)):
            # and making keys as index/pval and value index.
            identities[f"{i}?{pvals[i]}"] = [i]
    # Once checking is over, return the identities dictionary, groups and test
    return identities, groups, test

def storey_reorder_qs(qs, 
                      pvals, 
                      og_pvals):
    """
    =================================================================================================
    storey_reorder_qs(qs, pvals, og_pvals)
    
    This function is used in storey(), and is meant to take the list of qvalues, the list of pvalues,
    and the original list of pvalues, and reorder the qvalue list in correspondence with the original
    pvalue list.
                  
    =================================================================================================
    Arguments:
    
    qs        ->  A list of q values created using the pvals list
    pvals     ->  A list of p values used to estimate the q values
    og_pvals  ->  A list of p values in their original order
    
    =================================================================================================
    Returns: A list of q values which ahve been reordered to match the order of og_pvals
    
    =================================================================================================
    """
    # Initialize the list of seen pvalues and new qvalues
    seen = []
    newqs = []
    # Loop over original order of pvalues
    for i in range(len(og_pvals)):
        # If the current pvalue is already seen
        if og_pvals[i] in seen:
            # Then find the the index of this particular pvalue. It will
            # find the first instance of this pvalue in the pvals list.
            ind = pvals.index(og_pvals[i])
            # Then, see how many of these pvalues have been identified
            num = seen.count(og_pvals[i])
            # and qvalue corresponding to this pvalue is the index of
            # the pvals list up to the current pval + the number seen,
            # plus the number of elements before that list.
            ind = pvals[ind+num:].index(og_pvals[i]) + len(pvals[:ind+num])
        # If the current pvalue is not seen
        else:
            # find the index of the og_pval[i] in pvals
            ind = pvals.index(og_pvals[i])
        # move qs[ind] to newqs
        newqs.append(qs[ind])
        # Add this value to the seen list
        seen.append(og_pvals[i])
    # Once the loop is complete, the q value will be reordered based
    # on the original pvalue order.
    return newqs
    
    

def pi_0(ps, lam):
    """
    =================================================================================================
    pi_0(ps, lam)
    
    This function is used to calculate the value of pi_0 in the Storey FDR estimation algorithm
                  
    =================================================================================================
    Arguments:
    
    ps   ->  A list of pvalues
    lam  ->  A critical value of lambda
    
    =================================================================================================
    Returns: The sum of all p values greater than lambda divided by the number of p values times the
             difference and 1 and lambda.
    
    =================================================================================================
    """
    # hat(pi_0) = num(p_j >0) / len(p)/(1-lambda)
    # This just uses numpy functions to do that for us
    return np.sum(ps>lam) / (ps.shape[0]*(1-lam))

def storey(pvals, 
           pi0 = None, 
           groups = None,
           test = None):
    """
    =================================================================================================
    storey(pvals, pi0, groups, test)
    
    This function performs Storey False Discovery Rate Estimation, as described in the publication
    Statistical Significance for Genomewide Studies (Storey, Tibshirani 2003)
    https://www.pnas.org/content/pnas/100/16/9440.full.pdf
    
    =================================================================================================
    Arguments:
    
    pvals   ->  A list of pvalues. Can be unordered
    pi0     ->  A value for pi0. This does not need to be set.
    groups  ->  A list, tuple, dataframe, or numpy array describing comparison groups
    test    ->  A list, tuple, dataframe or numpy array describing the test used for Pvalue generation
    
    =================================================================================================
    Returns: A DataFrame describing the P-values, the Q-values, and whatever metadata was provided.
    
    =================================================================================================
    """
    # First, get a list of the group names if groups are provided.
    group_names = None
    if type(groups) == type(pd.DataFrame([])):
        group_names = list(groups.columns.values)
    # and check the arguments provided to the function.
    og_pvals, g_checker, t_checker  = storey_check_args(pvals, groups, test)
    #
    ##################################################
    # 1.) The pvalues should be sorted. Make a dict to keep track of the order and info
    # Use the storey_make_id_dict() function to organize the arguments
    identities, groups, test = storey_make_id_dict(og_pvals,
                                                   groups,
                                                   test,
                                                   g_checker,
                                                   t_checker)
    # Once the original sequence is preserved, sort the pvals
    pvals = sorted([float(val) for val in og_pvals])
    og_pvals = [float(val) for val in og_pvals]
    # IF a list is given, make it a numpy array
    if type(pvals) == list:
        pvals = np.array(pvals, dtype = float)
    #
    ##################################################
    # 2,3,4.) Calculate the hat(pi_0) for the lambdas and Fit the cubic spline or just set pi0s value
    # Make a range of lambda values and make an empty array
    # to hold the hat(pi_0)s
    # The lambdas are meant to be a range of values used to fit the cubic spline
    lambdas = np.arange(0.01,0.97,0.01)
    # If pi0 is not set, and the pvalues are less than 100 in length
    if len(pvals) < 100 and pi0 is None:
        # Set the hat_pi0 to 1, as it will be clsoe to 1 anyway
        hat_pi0 = 1.0
    # If pi0 is set, then set it to the input value
    elif pi0 != None:
        hat_pi0 = pi0
    # Otherwise, determine an appriopriate value.
    else:
        # Make the pi0s array
        pi0s = np.empty(lambdas.shape[0])
        # and for each lambda
        for i, cur_l in enumerate(lambdas):
            # calculate pi0 and assign it to the pi0s array
            pi0s[i] = pi_0(pvals, cur_l)
        #spline = UnivariateSpline(x=lambdas, y=pi0s, k=3)
        #hat_pi0 = spline(1)
        # Then use the scipy function splrep and splrev to
        # fit the cubic spline and calculate hat_pi0
        tck = splrep(lambdas, pi0s, k=3)
        hat_pi0 = splev(lambdas[-1], tck)
        # If the value of hat_pi0 is greater than 1
        if hat_pi0 > 1:
            # Just set it to 1, it cannot be greater than 1
            hat_pi0 = 1
    # If the value of hat_pi0 is 1
    if hat_pi0 == 1:
        # Then this algorithm is equivalent to BH
        method = "Benj-Hoch"
    # If hat_pi0 is less than 1
    else:
        # Then this is the true Storey algorithm
        method = "Storey"
    #
    ##################################################
    # 5.) Make the calculation q(p_m) = min_{t\geq p_{i}} * \dfrac{(hat_pi0 * m * t)}{num(p_j \leq t)} = hat_pi0 * p_m
    # Initialize the qs list, by multiplying hat_pi0 to
    # the highest pvalue
    qs = [hat_pi0 * pvals[-1]]
    #
    ##################################################
    # 6.) Calculate min( \dfrac{hat_pi0 * m * p_i}{i}, q(p_{i+1}))
    # Then use the calculation above to determine what the next qvalue should be
    # Loop backwards over length of pvals minus 1 to zero, since we have qm already
    for i in range(len(pvals)-1, 0, -1):
        # append the minimum of the previous q and the next prediction.
        # Note pvals[i-1], since i is mathematically counted (1,2,...,m)
        qs.append(min(hat_pi0*len(pvals)*pvals[i-1]/i, qs[-1]))
    # This list is reversed compared to the pvals list, so reverse it
    qs = [qs[-i] for i in range(1,len(qs)+1)]
    #
    ##################################################
    # 7.) The q value estimate for the ith most significant feature is q(p_i)
    # So we just need to format the output. Since the identities has the information
    # to format the qvals list,
    # align the pvalues and qvalues
    qs = storey_reorder_qs(qs, list(pvals), og_pvals)
    # Now, we need only to format the output properly.
    # if groups were provided, but test was not provided
    if g_checker != False and t_checker == False:
        # and if group_names == None,
        if group_names == None:
            # Then use the generic group names as the labels
            g_nums = [f"Group {i+1}" for i in range(len(groups[0])) ]
        # otherwise
        else:
            # Use the group names found in the beginning
            g_nums = group_names
        # and format the output as a list of lists of columns names.
        output = [["index", *g_nums, "Test", "pvalue", "method", "qvalue"]]
    # Or if the groups are not provcided and the test is provided
    elif g_checker == False and t_checker != False:
        # format the output headers as follows
        output = [["index", "Test", "pvalue", "method", "qvalue"]]
    # Or if both groups and test are provided
    elif g_checker != False and t_checker != False:
        # then determine whether group names were provided
        if group_names == None:
            # and if not, use generic group names
            g_nums = [f"Group {i+1}" for i in range(len(groups[0]))]
        # Otherwise,
        else:
            # use the group names identified eearlier
            g_nums = group_names
        # and make the output headers as follows
        output = [["index", *g_nums, "Test", "pvalue", "method", "qvalue"]]
    # IF no group names and no test was provided
    else:
        # Make the output headers minimal.
        output = [["index", "pvalue", "method", "qvalue"]]
    # Loop over the identities dictionary
    for pval, value in identities.items():
        # and add lists to the output, containing the value, the pvalue, Storey, and the qvalue
        output.append([*value, float(pval.split("?")[1]), f"{method}", qs[value[0]]])
    # Turn the outputs list into a dataframe
    returner = pd.DataFrame(np.array(output[1:]), columns = output[0])
    # remvoe the index column
    del returner["index"]
    # and return the output dataframe.
    return returner
###
#
#
#####################################################################################################################
#
#   Statistical Test Functions, DEFUNCT

def pairwise_t(*data_list, 
               comp_labels = True, 
               omit_comb = [],
               t_type = "d",
               nan_policy = "omit"):
    
    """ 
    =================================================================================================
    pairwise_t(*data_list, comp_labels, omit_comb, t_type, nan_policy)
    
    This function is meant to perform T-tests between the the input data.
                  
    =================================================================================================
    Arguments:
    
    data_list    ->  A list of lists, where each list is a set of data.
    comp_labels  ->  A boolean describing whether or not to include comparison labels.
                        If True, then the data_list is expected to be tuples, where element
                        zero is a label, and element one is a list of data.
    omit_comb    ->  A list of lists that describe combinations to omit from comparisons.
    t_type       ->  A string describing the type of comparison to be used.
    nan_policy   ->  A string describing how to handle nan values.
    
    =================================================================================================
    Returns: A Pandas DataFrame containing the results of the T-tests.
    
    =================================================================================================
    """
    # Check to make sure the inputs are correct
    assert comp_labels in [True, False], "comp_labels argument must be a boolean"
    assert type(omit_comb) == list, "Combinations to be ommitted should be a list of 2-tuples/2-lists"
    # Next, check the omit_comb list. If it is not empty
    if omit_comb != []:
        # Initialize the seen list
        seen = []
        # and loop over each combination to omit
        for comb in omit_comb:
            # and check each combination for validity
            if len(comb) == 1:
                assert type(comb) in [list, tuple] and len(comb) == 2, "Combinations to be ommitted should be a list of 2-tuples/2-lists"
            # Once the combination is checked, add it to the seen list
            seen.append(list(comb))
    # If the omit_comb list is empty
    else:
        # Then initialize an empty seen list.
        seen = []
    # After checking the omittion combinations, initialize
    # the statistics dictioanry
    stats = []
    # And loop over the given data lists
    for data1 in data_list:
        # And loop over the data lists again
        for data2 in data_list:
            # If the current combination has not been seen
            # and the lists are not the same list
            # Then also check whether or not comparison labels are expect. If not
            if [data1,data2] not in seen and [data2,data1] not in seen and data1 != data2 and comp_labels == False:
                # And add the forward and reverse lsits to the seen list
                seen.append([data1,data2])
                seen.append([data2,data1])
                # Then run a T-Test on the data and add it
                # to the stats list
                stats.append(list(ttest(data1,
                                        data2,
                                        t_type = t_type,
                                        nan_policy = nan_policy)))
            # If comparison labels are provided
            elif [data1,data2] not in seen and [data2,data1] not in seen and data1 != data2 and comp_labels == True:
                # And add the forward and reverse lsits to the seen list
                seen.append([data1,data2])
                seen.append([data2,data1])
                # Then run a T-Test on the data and add it
                # to the stats list with the labels
                stats.append([data1[0], 
                             data2[0],
                             *list(ttest(data1[1],
                                         data2[1],
                                         t_type = t_type,
                                        nan_policy = nan_policy))])
    # If comparison labels are expected,
    if comp_labels:
        #  then add the Group columns to the output DataFrame
        return pd.DataFrame(np.array(stats), columns = ["Group 1", "Group 2", "T-statistic", "DoF", "pvalue", "Test"])
    # If comparison labels are not expected
    else:
        # Then make the simple column headers for the DataFrame
        return pd.DataFrame(np.array(stats), columns = ["T-statistic", "DoF", "pvalue", "Test"])
    
####################################################################################################
#
#        Statistics Objects

#
#
############
#
#   StatFormatter base class. Used for representing the objects and writing to files.
    
class StatFormatter():
    
    def __init__(self):
        self.output = [{}]
    
    def _get_printinfo(self, a_dictionary):
        if "TukeyHSD" in a_dictionary["id"]:
            important = ["id", "Group 1", "Group 2", "q stat", "pvalue", "DF", "alpha","Reject", "Post hoc"]
            return {key : value for key, value in a_dictionary.items() if key in important}
        elif "ANOVA" in a_dictionary["id"]:
            important = ["id", "df1", "df2", "f stat", "pvalue", "alpha", "Reject"]
            return {key : value for key, value in a_dictionary.items() if key in important}
        elif "TTest" in a_dictionary["id"]:
            important = ["Test", "Group 1", "Group 2", "t stat", "pvalue", "tails", "DF", "alpha"]
            return {key : value for key, value in a_dictionary.items() if key in important}
        elif "FisherLSD" in a_dictionary["id"]:
            important = ["id", "Test", "Group 1", "Group 2", "t stat", "pvalue", "tails", "DF", "alpha", "Post hoc"]
            return {key : value for key, value in a_dictionary.items() if key in important}
        elif "Holm-Sidak" in a_dictionary["id"]:
            important = ["id", "Test", "Group 1", "Group 2", "pvalue", "alpha", "Reject", "Post hoc"]
            return {key : value for key, value in a_dictionary.items() if key in important}
        else:
            return {}
    
    def _print_table(self,a_dictionary):
        filtered = self._get_printinfo(a_dictionary)
        headers = list(filtered.keys())
        values = list(filtered.values())
        max_len = max([len(item) for item in values])
        newstr = ""
        for head in headers:
            newstr = f"{newstr}|{head:^10}"
        newstr = f"{newstr}|\n"
        head_sep = gh.list_to_str(["-" for _ in range(len(newstr))], delimiter = "", newline = True)
        newstr = f"{newstr}{head_sep}"
        for i in range(max_len):
            for sublist in values:
                try:
                    sublist[i]
                except:
                    newstr = f"{newstr}|{'':^10}"
                    continue
                if type(sublist[i]) == int:
                    newstr = f"{newstr}|{sublist[i]:^10}"
                elif type(sublist[i]) == float:
                    newstr = f"{newstr}|{sublist[i]:^10.4f}"
                elif type(sublist[i]) == str:
                    newstr = f"{newstr}|{sublist[i]:^10}"
                elif type(sublist[i]) == bool:
                    newstr = f"{newstr}|{str(sublist[i]):^10}"
            newstr = f"{newstr}|\n"
        return newstr
    
    def __str__(self):
        newstr = ""
        for item in self.output:
            newstr = f"{newstr}\n\n{self._print_table(item)}"
        return newstr
    
    def __repr__(self):
        return self.__str__()
    
    def fill_values(self, inplace = True):
        lens = [max([len(item) for item in list(a_dict.values())]) for a_dict in self.output]
        index = 0
        new_output = []
        for a_dict in self.output:
            newdict = {}
            for key, value in a_dict.items():
                if len(value) < lens[index]:
                    newdict[key] = value + ["" for _ in range(lens[index] - len(value))]
                else:
                    newdict[key] = value
            new_output.append(newdict)
        if inplace:
            self.output = new_output
        else:
            return new_output
                    
    
    def to_dfs(self):
        dfs = []
        for a_dict in self.output:
            max_len = max([len(item) for item in list(a_dict.values())])
            newdict = {}
            for key, value in a_dict.items():
                if len(value) == max_len:
                    newdict[key] = value
                else:
                    newdict[key] = [*value, *["" for _ in range(max_len - len(value))]]
            dfs.append(pd.DataFrame(newdict))
        return dfs
    
    def _make_outlists(self):
        lists = []
        for a_dict in self.output:
            max_len = max([len(item) for item in list(a_dict.values())])
            headlist = list(a_dict.keys())
            newlist = []
            for key, value in a_dict.items():
                if len(value) == max_len:
                    newlist.append(value)
                else:
                    newlist.append([*value, *["" for _ in range(max_len - len(value))]])
            lists.append([headlist] + gh.transpose(*newlist))
        return lists
    
    def write_output(self, filename = "output", file_type = "csv"):
        assert file_type.lower() in ["csv", "txt", "xlsx", "xls"], f"Invalid file type provided: {file_type}"
        if file_type.lower() in ["xls", "xlsx"]:
            output_file = f"{filename}.xlsx"
            dfs = self.to_dfs()
            i = 0
            with pd.ExcelWriter(output_file) as writer:
                for df in dfs:
                    df.to_excel(writer, engine = "openpyxl",sheet_name = f"sheet_{i}", index = False)
                    i+=1
            return f"File {output_file}.xlsx has been written."
        elif file_type.lower() == "csv":
            output_file = f"{filename}.csv"
            organised = self._make_outlists()
            outlines = []
            for line_list in organised:
                for line in line_list:
                    outlines.append(gh.list_to_str(line, ",", newline = True))
                outlines.append("\n\n")
            gh.write_outfile(outlines, output_file)
            return f"File {output_file}.csv has been written."
        elif file_type.lower() == "txt":
            output_file = f"{filename}.txt"
            organised = self._make_outlists()
            outlines = []
            for line_list in organised:
                for line in line_list:
                    outlines.append(gh.list_to_str(line, "\t", newline = True))
                outlines.append("\n\n")
            gh.write_outfile(outlines, output_file)
            return f"File {output_file} has been written."
        else:
            raise ValueError(f"The file type you requested is not supported:{file_type}")
            
    def _fix_group_ids(self, list1, list2, current_id = 1):
        """
        RECURSIVE: probably slow for high numbers of repeats.
        """
        list1 = [str(item) for item in list1]
        list2 = [str(item) for item in list2]
        comp = [item for item in list1 if item in list2]
        if comp != []:
            if "_?_" in list2[0]:
                self._fix_group_ids(list1, list2, current_id = current_id + 1)
            else:
                return list1 + [f"{item}_?_{current_id}" for item in list2]
        else:
            return list1 + list2
    
    def _merge_dicts(self, dict1, dict2, 
                     check_groups = True):
        """
        """
        keys = list(dict1.keys())
        newdict = {}
        for key in keys:
            if "Group" in key and check_groups:
                newdict[key] = self._fix_group_ids(dict1[key], dict2[key])
            elif "id" == key:
                newdict[key] = dict1[key]
            else:
                newdict[key] = dict1[key] + dict2[key]
        return newdict
    
    def _merge_outputs(self, sf_object,
                       check_groups = True):
        """
        """
        new_output = []
        ids = []
        for dict1 in self.output:
            missed = 0
            for dict2 in sf_object.output:
                if dict1["id"] != dict2["id"]:
                    missed += 1
                else:
                    ids.append(dict1["id"])
                    new_output.append(self._merge_dicts(dict1, dict2,
                                                        check_groups = check_groups))
        for item in sf_object.output:
            if item["id"] not in ids:
                new_output.append(item)
        for item in self.output:
            if item["id"] not in ids:
                new_output.append(item)
        return new_output
    
    def combine(self, sf_object, inplace = True,
                check_groups = True):
        # If the objects have the same type, then we can combine
        # them as one.
        if type(sf_object) == type(self):
            new_output = self._merge_outputs(sf_object,
                                             check_groups = check_groups)
            if inplace:
                self.output = new_output
            else:
                return new_output
        else:
            new_output = []
            id1 = [item["id"] for item in self.output]
            id2 = [item["id"] for item in sf_object.output]
            comp = [item for item in id1 if item in id2]
            if inplace:
                if comp != []:
                    self.output = self._merge_outputs(sf_object,
                                                      check_groups = check_groups)
                else:
                    self.output += sf_object.output
            else:
                if comp != []:
                    new_output = self._merge_outputs(sf_object,
                                                      check_groups = check_groups)
                else:
                    new_output = self.output + sf_object.output
                return new_output
    
    def grab_results(self, test_id):
        """
        """
        for item in self.output:
            if test_id in item["id"]:
                return item
        return None

#
#
#######################
#
#   T tests

class TTest(StatFormatter):
    
    def __init__(self, data1, data2, 
                 test_type = "d", tails = 2, 
                 nan_policy = "omit", alpha = 0.05,
                 labels = True, df = None, msew = None,
                 threshold = 2):
        if not labels:
            assert all([type(d) in [int, float] for d in data1]), "There are non-numbers in the data..." 
            assert all([type(d) in [int, float] for d in data2]), "There are non-numbers in the data..."
        else:
            assert all([type(d) in [int, float] for d in data1[1]]), "There are non-numbers in the data..." 
            assert all([type(d) in [int, float] for d in data2[1]]), "There are non-numbers in the data..."
        assert test_type in ["w", "welch", "s", "student", "d", "determine"], "The test_type given is not valid. Try 'w', 's' or 'd'"
        assert nan_policy.lower() in ["omit", "propogate"], "The nan policy is not valid. Try 'omit' or 'propogate'."
        assert type(alpha) == float and alpha <= 1 and alpha >= 0, "The alpha value is not valid."
        assert tails in [2,1,-1], "The tails are not valid. Try 2, 1 or -1."
        
        assert any([df == None and msew == None, df != None and msew != None]), "Mean squared error within and the passed in Degrees of freedom must both be None or numbers..."
        if df != None and msew != None:
            assert type(df) in [int, float], "The Degrees of freedom (df) argument must be a number..."
            assert type(msew) in [int, float], "The Degrees of freedom (df) argument must be a number..."
        
        if not labels:
            self.d1 = copy.copy(["1", data1])
            self.d2 = copy.copy(["2", data2])
        else:
            self.d1 = copy.copy(data1)
            self.d2 = copy.copy(data2)
        if test_type.lower() in ["d", "determine"]:
            self.tt = "determine"
        elif test_type.lower() in ["s", "student"]:
            self.tt = "student"
        else:
            self.tt = "welch"
        self.tails = tails
        self.alpha = alpha
        
        if nan_policy.lower() == "propogate":
            if any([d != d for d in data1]) or any([d != d for d in data2]):
                self.output = [{"id"  : ["TTest"],
                               "Test" : ["nan"],
                               "Group 1"   : [self.d1[0]],
                               "Group 2"   : [self.d2[0]],
                               "Size 1"    : [len(self.d1[1])],
                               "Size 2"    : [len(self.d2[1])],
                               "muDiff" : [float("nan")],
                               "t stat" : [float("nan")],
                               "pvalue" : [float("nan")],
                               "tails"  : [self.tails],
                               "DF" : [float("nan")],
                               "alpha" : [self.alpha]}]
                return None
        else:
            self._correct_nans_ttest()
        
        if len(self.d1[1]) < threshold or len(self.d2[1]) < threshold:
            self.output = [{"id"  : ["TTest"],
                           "Test" : ["nan"],
                           "Group 1"   : [self.d1[0]],
                           "Group 2"   : [self.d2[0]],
                           "Size 1"    : [len(self.d1[1])],
                           "Size 2"    : [len(self.d2[1])],
                           "muDiff" : [float("nan")],
                           "t stat" : [float("nan")],
                           "pvalue" : [float("nan")],
                           "tails"  : [self.tails],
                           "DF" : [float("nan")],
                           "alpha" : [self.alpha]}]
            self.fill_values()
            return
        
        if df == None:
            self.df = self.t_degree_freedom()
        else:
            self.df = df
        
        if df != None and msew != None:
            return self.ttest(msew = msew)
        
        if self.tt in ["d","determine"]:
            var1 = variance(self.d1[1])
            var2 = variance(self.d2[1])
            if 1/2 <= var1/var2 <= 2:
                self.tt = "student"
                self.output = [self.t_test()]
            else:
                self.tt = "welch"
                self.output = [self.t_test()]
                
        else:
            self.output = [self.t_test()]
        self.fill_values()
    
    def _correct_nans_ttest(self):
        self.d1[1] = [d for d in self.d1[1] if d == d]
        self.d2[1] = [d for d in self.d2[1] if d == d]
        return None
            
    def t_crit(self):
        """
        """
        if self.tails == 2:
            return t.ppf(1 - (self.alpha/2), df = self.df)
        elif self.tails == 1:
            return t.ppf(1 - self.alpha, df = self.df)
        elif self.tails == -1:
            return t.ppf(self.alpha, df = self.df)
        
    def _pooled_var(self, msew = None):
        """
        """
        var1 = variance(self.d1[1])
        var2 = variance(self.d2[1])
        if msew != None:
            return sqrt(msew * (1/len(self.d1[1]) + 1/len(self.d2[1])))
        if self.tt in ["s", "student"]:
            pooled_num = (len(self.d1[1]) - 1) * var1 + (len(self.d2[1]) - 1) * var2
            pooled_den = len(self.d1[1]) + len(self.d2[1]) - 2
            return sqrt(pooled_num/pooled_den)
        else:
            return sqrt((var1 / len(self.d1[1])) + (var2 / len(self.d2[1])))
    
    def t_degree_freedom(self):
        """
        """
        if self.tt in ["s", "student"]:
            return len(self.d1[1]) + len(self.d2[1]) - 2
        else:
            var1 = variance(self.d1[1])
            var2 = variance(self.d2[1])
            numer = ((var1 / len(self.d1[1])) + (var2 / len(self.d2[1])))**2
            denom = ((var1 / len(self.d1[1]))**2 / (len(self.d1[1]) - 1)) + ( (var2 / len(self.d2[1]))**2 / (len(self.d2[1]) - 1) )
            return numer / denom
    
    def t_statistic(self,
                    msew = None):
        """
        """ 
        numer = mean(self.d1[1]) - mean(self.d2[1])
        poolvar = self._pooled_var(msew = msew)
        if msew != None:
            return numer / poolvar
        elif self.tt in ["s", "student"]:
            denom = poolvar * sqrt(1/len(self.d1[1]) + 1/len(self.d2[1]))
        else:
            denom = poolvar
        return numer/denom
    
    def t_pval(self, t_stat):
        if self.tails == 2:
            return 2 * (1 - t.cdf(abs(t_stat), df = self.df))
        elif self.tails == 1:
            return 1 - t.cdf(abs(t_stat), df = self.df)
        elif self.tails == -1:
            return t.cdf(t_stat, df = self.df)
        else:
            raise ValueError("Something catastrophic has occurred...")
    
    def t_test(self,
               msew = None):
        """
        """
        if msew == None:
            t_stat = self.t_statistic()
            pval = self.t_pval(t_stat)
            return {"id"          : ["TTest"],
                    "Test"        : [self.tt],
                    "Group 1"     : [self.d1[0]],
                    "Group 2"     : [self.d2[0]],
                    "Size 1"      : [len(self.d1[1])],
                    "Size 2"      : [len(self.d2[1])],
                    "muDiff"      : [mean(self.d1[1]) - mean(self.d2[1])],
                    "t stat"      : [t_stat],
                    "pvalue"      : [float(pval)],
                    "tails"       : [self.tails],
                    "DF"          : [self.df],
                    "alpha"       : [self.alpha]}
        else:
            t_stat = self.t_statistic(msew = msew)
            pval = self.t_pval(t_stat)
            return {"id"          : ["FisherLSD"],
                    "Test"        : ["Protected"],
                    "Group 1"     : [self.d1[0]],
                    "Group 2"     : [self.d2[0]],
                    "Size 1"      : [len(self.d1[1])],
                    "Size 2"      : [len(self.d2[1])],
                    "muDiff"      : [mean(self.d1[1]) - mean(self.d2[1])],
                    "t stat"      : [t_stat],
                    "pvalue"      : [float(pval)],
                    "tails"       : [self.tails],
                    "DF"          : [self.df],
                    "alpha"       : [self.alpha]}

class PairwiseT(TTest):
    
    def __init__(self):
        return None

#
#
############
#
#    One Way ANOVA
    
class ANOVA(StatFormatter):
    
    def __init__(self,
                 *data,
                 way = 1,
                 alpha = 0.05,
                 nan_policy = "omit"):
        """
        """
        assert way in [1,2], "The argument way only takes values 1 or 2."
        assert nan_policy in ["omit", "propogate"], "The nan policy is not valid. Try 'omit' or 'propogate'."
        self.d = data
        if nan_policy == "propogate":
            if any([any([d != d for d in data_list]) for data_list in data]):
                self.output = [{"id"     : ["ANOVA"],
                               "way"     : [way],
                               "df1"     : [float("nan")],
                               "df2"     : [float("nan")],
                               "f stat"  : [float("nan")],
                               "f_crit"  : [float("nan")],
                               "pvalue"  : [float("nan")],
                                "alpha"  : [alpha],
                                "Reject"    : float("nan")}]
                return None
        else:
            self.cor_data = self._correct_nans_anova()
        df1 = len(self.cor_data) - 1
        df2 = sum([len(d) for d in self.cor_data]) - len(self.cor_data)
        if way == 1:
            self.output = [{"id"    : ["ANOVA"],
                            "way"   : [way],
                           "df1"    : [df1],
                           "df2"    : [df2],
                           "f stat" : [self.f_statistic(*self.cor_data)],
                           "f_crit" : [self.f_critical(alpha, df1, df2)],
                           "pvalue" : [float(1 - f.cdf(self.f_statistic(*self.cor_data), df1, df2))],
                           "alpha"  : [alpha]}]
            self.output[0]["Reject"] = [self.output[0]["pvalue"] < self.output[0]["alpha"]]
            self.fill_values()
        else:
            raise ValueError("Two way ANOVAs are not yet programmed...")
    
    def _correct_nans_anova(self):
        return [[d for d in data_list if d == d] for data_list in self.d]
            
    def f_statistic(self, *data, correction = 1):
        """
        """
        lens = [len(d) for d in data]
        if all([l == len(data[0]) for l in lens]):
            return var_between(*data, correction = correction) / var_within(*data, correction=correction)
        else:
            return ms_between(*data) / ms_within(*data, correction = correction)
    
    def f_critical(self, alpha, df1, df2):
        """
        """
        return float(f.ppf(1-alpha, df1, df2))

#
#
############
#
# Tukey Honest Significant Difference Test (Protected)

class TukeyHSD(ANOVA):
    
    def __init__(self,
                 *data,
                 alpha = 0.05,
                 labels = True,
                 nan_policy = "omit",
                 override = False):
        """
        """
        # Run the ANOVA
        if labels:
            super().__init__(*[d[1] for d in data], way = 1, alpha = alpha, nan_policy = nan_policy)
            self.cor_data = [[data[i][0], self.cor_data[i]] for i in range(len(data))]
        else:
            super().__init__(*data, way = 1, alpha = alpha, nan_policy = nan_policy)
        #
        if nan_policy.lower() == "propogate":
            if any([any([d != d for d in data_list]) for data_list in data]):
                self.output = {"id"  : ["TukeyHSD"],
                               "Test" : [float("nan")],
                               "Group 1" : [float("nan")],
                                "Group 2" : [float("nan")],
                                "muDiff"   : [float("nan")],
                                "Lower B" : [float("nan")],
                                "Upper B" : [float("nan")],
                                "q stat" : [float("nan")],
                                "pvalue" : [float("nan")],
                                "alpha" : [float("nan")],
                                "Reject" : [float("nan")],
                                "q crit" : [float("nan")],
                                "DF" : [float("nan")],
                                "Group N" : [float("nan")]}
            return None
        self.override = override
        # If a significant difference in the means is not identified
        if self.output[0]["pvalue"][0] >= alpha and not override:
            # Then save the output with a message for post hoc tests
            self.output.append({"id": ["TukeyHSD"],
                                "Test" : ["Protected"],
                                "Post hoc" : ["Not", "permitted."]})
        elif self.output[0]["pvalue"][0] >= alpha and override:
            self.output.append(self.tukey_hsd(*self.cor_data, alpha = alpha, labels = labels))
            self.output[-1]["Test"] = ["Not", "protected"]
        # Otherwise
        else:
            # Save the results 
            self.output.append(self.tukey_hsd(*self.cor_data, alpha = alpha, labels = labels))
        self.fill_values()
        
    def q_statistic(self,
                    data1,
                    data2,
                    msw):
        standard_error = sqrt(msw/2*((1/len(data1) + 1/len(data2))))
        return abs(mean(data1) - mean(data2))/standard_error

    def q_conf(self,
               data1,
               data2,
               q_crit,
               msw):
        standard_error = sqrt(msw/2*((1/len(data1) + 1/len(data2))))
        mean_diff = abs(mean(data1) - mean(data2))
        return (mean_diff - q_crit * standard_error, mean_diff + q_crit * standard_error)
    
    def tukey_hsd(self,
                  *user_data,
                  alpha = 0.05,
                  labels = False):
        """
        """
        if not labels:
            data = [[i, user_data[i]] for i in range(len(user_data))]
        else:
            data = copy.copy(user_data)
        df = sum([len(d[1]) for d in data]) - len(data)
        k = len(data)
        tukeyout = {"id"      : ["TukeyHSD"],
                    "Group 1" : [],
                    "Group 2" : [],
                    "muDiff"   : [],
                    "Lower B" : [],
                    "Upper B" : [],
                    "q stat" : [],
                    "pvalue" : [],
                    "alpha" : [alpha],
                    "Reject"   : [],
                    "q crit" : [float(q.ppf(1-alpha, k, df))],
                    "DF" : [df],
                    "Group N" : [k]}
        msw = ms_within(*[d[1] for d in data])
        paired = gh.make_pairs(data, dupes = False, reverse = False)
        for pair in paired:
            tukeyout["Group 1"].append(pair[0][0])
            tukeyout["Group 2"].append(pair[1][0])
            tukeyout["muDiff"].append(mean(pair[0][1]) - mean(pair[1][1]))
            tukeyout["Lower B"].append(self.q_conf(pair[0][1],
                                                       pair[1][1],
                                                       tukeyout["q crit"][0],
                                                       msw)[0])
            tukeyout["Upper B"].append(float(self.q_conf(pair[0][1],
                                                       pair[1][1],
                                                       tukeyout["q crit"][0],
                                                       msw)[1]))
            tukeyout["q stat"].append(float(self.q_statistic(pair[0][1],
                                                            pair[1][1],
                                                            msw)))
            tukeyout["pvalue"].append(float(1 - q.cdf(self.q_statistic(pair[0][1],
                                                                 pair[1][1],
                                                                 msw), k, df)))
            tukeyout["Reject"].append(tukeyout["pvalue"][-1] < alpha)
        if self.override:
            tukeyout["Test"] = ["Unprotected"]
        else:
            tukeyout["Test"] = ["Protected"]
        return tukeyout
    
#
#
############
#
#    Fisher LSD test (Protected)

class FisherLSD(TTest,ANOVA):
    
    def __init__(self, *data, labels = True, 
                 override = False, alpha = 0.05):
        # Override -> unprotected
        # perform pairwise T tests with DF from the ANOVA and MSEw for the data
        if labels:
            # self.cor_data and self.alpha should be assigned here
            ANOVA.__init__(self, *[d[1] for d in data], way = 1, alpha = alpha)
            self.cor_data = [[data[i][0], self.cor_data[i]] for i in range(len(data))]
        else:
            ANOVA.__init__(self, data, way = 1, alpha = alpha)
            self.cor_data = [[i +1, self.cor_data[i]] for i in range(len(self.cor_data))]
        self.df = sum([len(d[1]) for d in self.cor_data]) - len(self.cor_data)
        self.tails = 2
        self.alpha = alpha
        if not override and self.output[0]["pvalue"][0] >= alpha:
            self.output.append({"id" : ["FisherLSD"],
                                "Test": ["Protected"],
                                "Post hoc" : ["Not", "permitted."]})
        elif override and self.output[0]["pvalue"][0] >= alpha:
            self.output.append(self.fisher_lsd())
            self.output[-1]["Test"] = ["Not", "protected"]
            del self.d1
            del self.d2
        else:
            self.output.append(self.fisher_lsd())
            del self.d1
            del self.d2
        return None
    
    def lsd_statistic(self):
        return TTest.t_crit()
    
    def fisher_lsd(self, labels = True):
        # make the pairs
        comparisons = gh.make_pairs(self.cor_data, dupes = False, reverse = False)
        #
        if labels:
            msew = ms_within(*[d[1] for d in self.cor_data])
        else:
            msew = ms_within(*self.cor_data)
        #
        tests = []
        #
        for comp in comparisons:
            self.d1, self.d2 = comp
            tests.append(TTest.t_test(self, msew = msew))
        tests = gh.merge_dicts(*tests, filler = "")
        tests = {key : gh.unpack_list(value) for key, value in tests.items()}
        # Reformate some of the output by slicing lists
        tests["id"] = tests["id"][:1]
        tests["Test"] = tests["Test"][:1]
        tests["tails"] = tests["tails"][:1]
        tests["DF"] = tests["DF"][:1]
        tests["alpha"] = tests["alpha"][:1]
        return tests

#
#
###########
#
#    Holm-Sidak FWER Correction (Protected)

class HolmSidak(FisherLSD):
    
    def __init__(self, *data, labels = True, 
                 override = False, alpha = 0.05):
        # Run FisherLSD.__init__() to get Pvalues and what not
        super().__init__(*data, labels = labels, override = override,
                         alpha = alpha)
        # Now self.output should have ANOVA and FisherLSD
        fisher_ind = self._find_fisher()
        if not override and "Post hoc" in list(self.output[fisher_ind].keys()):
            self.output.append({"id" : ["Holm-Sidak"],
                                "Test" : ["Protected"],
                                "Post hoc" : ["Not", "permitted."]})
        else:
            self.output.append(self.holm_sidak())
        return None
    
    def _find_fisher(self):
        return max([self.output.index(item) for item in self.output if "FisherLSD" in item["id"]])
    
    def _adjust_pval(self, pval, n, k):
        # Formula for calculating an adjusted Pvalue
        return 1 - (1 - pval)**(k - n + 1)
    
    def _adjust_pvals(self, pvals):
        # K is the number of comparisons, also the number of pvalues
        k = len(pvals)
        # index the pvals
        sort_pvals = list(zip([i for i in range(k)],pvals))
        sort_pvals = sorted(sort_pvals, key = lambda x: x[1])
        # Adjust first value
        adjusted = [[sort_pvals[0][0], self._adjust_pval(sort_pvals[0][1], 1, k)]]
        # Loop over the remaining values and compare
        for n in range(1, len(sort_pvals)):
            adjusted.append([sort_pvals[n][0], max([adjusted[n-1][1], self._adjust_pval(sort_pvals[n][1], n+1, k)])])
        adjusted = sorted(adjusted, key = lambda x: x[0])
        return [adjusted[i][1] for i in range(k)]
    
    def holm_sidak(self):
        fisher_ind = self._find_fisher()
        new_dict = copy.copy(self.output[fisher_ind])
        new_dict["pvalue"] = self._adjust_pvals(new_dict["pvalue"])
        new_dict["id"] = ["Holm-Sidak"]
        new_dict["Reject"] = [new_dict["pvalue"][i] < self.alpha for i in range(len(new_dict["pvalue"]))]
        return new_dict

#
#
#####################################################################################################################
```
