## Supporting Folder 2 Data Analysis for "Mouse primary T cell phosphotyrosine proteomics enabled by BOOST": western_helpers.py

```
"""
Kenny Callahan

File for plotting LI-COR Odyssey CLx Western data.
"""
##################################################################################################
#
#

import pandas as pd

from . import stats_helpers as sh

from . import mpl_plotting_helpers as mph

#
#
##################################################################################################
#
#

def get_signal(file_df, channel, signal_column = "Signal", channel_column = "Channel"):
    """
    """
    assert type(channel) == int, "The channel should be a number (700 or 800)."
    assert type(file_df) == type(pd.DataFrame([])), "The data should be in a DataFrame."
    channel_filtered = file_df[file_df[channel_column] == channel]
    return list(channel_filtered[signal_column].astype(float))

def licor_correction(exp_signal, control_signal):
    """
    """
    # Make sure all of the inputs are numbers
    assert all([type(item) == float or type(item) == int for item in exp_signal]), "Signal should be provided as a float."
    assert all([type(item) == float or type(item) == int for item in control_signal]), "Signal should be provided as a float."
    assert len(exp_signal) == len(control_signal), "The exp_signal and control_signal should be index paired lists of the same size."
    # Make the correction factors using the max signal
    corr_facts = [signal / max(control_signal) for signal in control_signal]
    # Correct the experimental signals using the controls
    return [exp_signal[i]/corr_facts[i] for i in range(len(exp_signal))]

def bin_data(signal_list, bins, labels = [], group_sizes = []):
    """
    """
    assert all([type(item) == float or type(item) == int for item in signal_list]), "Signal should be provided as a float."
    assert type(bins) == int, "The number of groups (bins) should be an integer."
    assert type(labels) in [list, tuple], "The labels should be in a list or tuple."
    if group_sizes != []:
        assert all([type(item) == int for item in group_sizes]), "The group sizes should be integers."
        assert len(group_sizes) == bins, "The number of given group sizes does not match the number of groups."
        groups = []
        count = 0
        for item in group_sizes:
            groups.append(signal_list[count:count+item])
            count += item
    else:
        # Assume an even number of things in each bin
        n_members = len(signal_list)//bins
        groups = [signal_list[n_members*i: n_members*(i+1)] for i in range(bins)]
    if labels != []:
        assert all([type(item) == str for item in labels]), "The labels should be strings"
        assert len(labels) == bins, "Each group should have a corresponding label"
        return [(labels[i], groups[i]) for i in range(bins)]
    else:
        return [(f"{i}", groups[i]) for i in range(bins)]
    
def perform_statistics(labelled_groups,
                       test = "Holm-Sidak",
                       alpha = 0.05):
    tests = {"TTest" : sh.TTest,
             "TukeyHSD" : sh.TukeyHSD,
             "Holm-Sidak" : sh.HolmSidak}
    assert test in ["TTest", "TukeyHSD", "Holm-Sidak", "none"], f'The test provided is not valid. Try :{"TTest"} (For two tests), {"TukeyHSD"}, or {"Holm-Sidak"}'
    if test == "none" or len(labelled_groups) == 1:
        return {}
    if len(labelled_groups) < 3 and test != "TTest":
        print(f"You chose {test} for hypothesis testing, but provided {len(labelled_groups)} groups.\nProceeding with a TTest instead.")
        test = "TTest"
    elif len(labelled_groups) > 2 and test == "TTest":
        print(f"You chose {test} for hypothesis testing, but provided {len(labelled_groups)} groups.\nProceeding with Holm-Sidak multiple comparisons instead.")
        test = "Holm-Sidak"
    output = tests[test](*labelled_groups, alpha = alpha)
    if test != "TTest" and output.grab_results("ANOVA")["pvalue"][0] >= alpha:
        return output, "ANOVA"
    else:
        return output, test

#
#
###################################################################################################
#
#

def western_quant(file1,                  # File with LI-COR intensity data
                  file2 = None,           # File with control LI-COR intensity data
                  exp_channel = 800,      # Channel with experimental data
                  control_channel = 700,  # Channel with control data
                  bins = 4,               # Number of groups represented in the data
                  labels = [],            # Labels for each group, same order as File
                  group_sizes = [],        # Sizes of the groups
                  filename = "dotplot.pdf",# Name of the output file
                  title = "Dotplot",       # Title for the plot
                  ylabel = "Abundance",    # Label for the Y axis
                  colours = [],            # List of colours for each group. Must be >= bins
                  errorbar = "sem",        # Value for error bars: SEM or SD currently supported
                  foldchange_axis = False, # Boolean for foldchange axis.
                  foldchange_group = None, # Group to compare other groups to
                  test = "Holm-Sidak",     # Statistical test to use
                  perform_stats = True,
                  stats_outfile_type = "xlsx",
                  alpha = 0.05,            # Significance level to use
                  sig_dict = {0.05 : "$*$",# Dictionary of values for how to represent significance
                        0.01 : "$*$$*$",
                        0.001 : "$*$$*$$*$"},
                  p_or_q = "p"):           # Significance letter (p value or q value)
    """
    """
    
    file = pd.read_excel(file1)
    if file2 == None:
        exp_signal = get_signal(file,
                                exp_channel)
        control_signal = get_signal(file,
                                    control_channel)
        corr_signal = licor_correction(exp_signal,
                                       control_signal)
        groups = bin_data(corr_signal,
                          bins = bins,
                          labels = labels,
                          group_sizes = group_sizes)
        if type(colours) == str:
            colours = mph.handle_colours(colours, len(groups))
        elif type(colours) == list and colours != []:
            if len(colours) < len(groups):
                print("Not enough colours were provided. Proceeding with a random selection from all colours.")
                colours = mph.handle_colours("all", len(groups), "random")
                print(f"Colours chosen: {colours}\n")
        if perform_stats:
            comp_dict, test = perform_statistics(groups, test = test, alpha = alpha)
            new_filename = file1.split(".")[0]
            new_filename = f"{new_filename}_statistics"
            comp_dict.write_output(new_filename, stats_outfile_type)
            comp_dict = comp_dict.grab_results(test)
        else:
            comp_dict = {}
        ax = mph.dotplot(groups,
                         filename = filename,
                         comparisons = comp_dict,
                         colours = colours,
                         title = title,
                         ylabel = ylabel,
                         foldchange_axis = foldchange_axis,
                         foldchange_group = foldchange_group,
                         ymin = 0,
                         errorbar = errorbar)
        return ax
    else:
        file_2 = pd.read_excel(file2)
        exp_signal = get_signal(file,
                                exp_channel)
        control_signal = get_signal(file_2,
                                    control_channel)
        corr_signal = licor_correction(exp_signal,
                                       control_signal)
        groups = bin_data(corr_signal,
                          bins = bins,
                          labels = labels,
                          group_sizes = group_sizes)
        if type(colours) == str:
            colours = mph.handle_colours(colours, len(groups))
        elif type(colours) == list and colours != []:
            if len(colours) < len(groups):
                print("Not enough colours were provided. Proceeding with a random selection from all colours.")
                colours = mph.handle_colours("all", len(groups), "random")
                print(f"Colours chosen: {colours}\n")
        
        if perform_stats:
            comp_dict, test = perform_statistics(groups, test = test, alpha = alpha)
            new_filename = file1.split(".")[0]
            new_filename = f"{new_filename}_statistics"
            comp_dict.write_output(new_filename, stats_outfile_type)
            comp_dict = comp_dict.grab_results(test)
        else:
            comp_dict = {}
        ax = mph.dotplot(groups,
                         filename = filename,
                         comparisons = comp_dict,
                         colours = colours,
                         title = title,
                         ylabel = ylabel,
                         foldchange_axis = foldchange_axis,
                         foldchange_group = foldchange_group,
                         ymin = 0,
                         errorbar = errorbar)
        return ax
    
#
#
###################################################################################################
```
