## Supporting Folder 2 Data Analysis for "Mouse primary T cell phosphotyrosine proteomics enabled by BOOST": mouse_jurkat_site_comparison.py

```
"""
============================================================================================================================================================
Mouse primary T cell phosphotyrosine proteomics enabled by BOOST
Xien Yu Chua(1), Kenneth P. Callahan(2), Alijah A. Griffith(2), Tobias Hildebrandt(1), Guoping Fu(3), Mengzhou Hu(1), Renren Wen(3), Arthur R. Salomon(2,*)
(1)Department of Molecular Pharmocology, Physiology & Biotechnology, Brown University, Providence, RI, 02912
(2)Department of Molecular Biology, Cell Biology & Biochemistry, Brown University, Providence, RI, 02912
(3)Blood Research Institute, Blood Center of Wisconsin, Milwaukee, WI, 53226
* Corresponding Author,
============================================================================================================================================================
Kenneth P. Callahan
19 February 2022
Salomon Laboratory
Post-MaxQuant Data Analysis

Python 3.8.10

Dependencies:

helpers: A folder of common functions created by Kenneth P. Callahan for data analysis
         Put this folder in the same directory as this file and include '__init__.py' in this
         this directory as well

NumPy Version 1.19.2
SciPy Version 1.7.2
Pandas Version 1.2.3
matplotlib Version 3.3.2
matplotlib-venn Version 0.11.6

============================================================================================================================================================
This script was used to compare unique pTyr sites between our data and the previously published
data from Jurkat T cells (Chua et al. 2020) and create two figures:

- Supporting Figure 12: KEGG TCR Site Comparison
- Figure 5: Mouse-Jurkat Site overlap

The data from Chua et al. 2020 is publically available (see their publication for details).
============================================================================================================================================================

Chua, X. Y.; Mensah, T.; Aballo, T.; Mackintosh, S. G.; Edmondson, R. D.; Salomon, A. R. Tandem mass tag 
approach utilizing pervanadate BOOST channels delivers deeper quantitative characterization of the 
tyrosine phosphoproteome. Molecular & Cellular Proteomics 2020, 19, 730– 743, 
DOI: 10.1074/mcp.TIR119.001865

============================================================================================================================================================
"""

###################################################################################################
#
#  Importables

from helpers import stats_helpers as sh
from helpers import mpl_plotting_helpers as mph
from helpers import general_helpers as gh

from matplotlib_venn import venn2, venn2_circles
import matplotlib.pyplot as plt

from data_analysis import keep_first, make_loc_probs, add_sequence_window, compare_seqs_to_database, get_other_species_sites

#
#
###################################################################################################
#
#  Colours, labels, headers to rename, etc.

rename_cols = {"Reporter intensity corrected 1" : "0.1 mg R1",
               "Reporter intensity corrected 2" : "0.1 mg R2",
               "Reporter intensity corrected 3" : "0.1 mg R3",
               "Reporter intensity corrected 4" : "0.3 mg R1",
               "Reporter intensity corrected 5" : "0.3 mg R2",
               "Reporter intensity corrected 6" : "0.3 mg R3",
               "Reporter intensity corrected 7" : "1.0 mg R1",
               "Reporter intensity corrected 8" : "1.0 mg R2",
               "Reporter intensity corrected 9" : "1.0 mg R3",
               "Reporter intensity corrected 10" : "PV BOOST R1",
               "Reporter intensity corrected 11" : "PV BOOST R2",
               "Phospho (STY) Probabilities" : "Localisation Probability (Sequence)",
               "Gene names" : "Gene names",
               "Protein names" : "Protein names",
               "id" : "id",
               "Phospho (STY) site IDs" : "Phospho (STY) site IDs",
               "Experiment" : "Experiment",
               "Modified sequence" : "Modified sequence",
               "Charge" : "Charge"}

keep_boost_cols = {"Gene names" : "Gene names",
               "Protein names" : "Protein names",
               "Modified sequence" : "Modified sequence",
               "Localisation Probability (Sequence)" : "Localisation Probability (Sequence)",
               "Modified Site" : "Modified Site",
               "PSP Modified Site (Equivalent in MOUSE)" : "PSP Modified Site (Equivalent in MOUSE)",
               "Highest (Y) Localisation Probability" : "Highest (Y) Localisation Probability",
               "All (Y) Localisation Probabilities":"All (Y) Localisation Probabilities",
               "Sequence Window" : "Sequence Window",
               "Flanking Sequence" : "Flanking Sequence",
               "PSP Flanking Sequence (Equivalent in MOUSE)" : "PSP Flanking Sequence (Equivalent in MOUSE)",
               "Charge" : "Charge",
               "Experiment" : "Experiment",
               "0.1 mg R1" : "0.1 mg R1",
               "0.1 mg R2" : "0.1 mg R2",
               "0.1 mg R3" : "0.1 mg R3",
               "0.3 mg R1" : "0.3 mg R1",
               "0.3 mg R2" : "0.3 mg R2",
               "0.3 mg R3" : "0.3 mg R3",
               "1.0 mg R1" : "1.0 mg R1",
               "1.0 mg R2" : "1.0 mg R2",
               "1.0 mg R3" : "1.0 mg R3",
               "PV BOOST R1" : "PV BOOST R1",
               "PV BOOST R2" : "PV BOOST R2",
               "Median"   : "Row Median",
               "Missing values"  : "Missing Values",
               "Unique ID" : "Unique ID",
               "PSP Site Group ID" : "PSP Site Group ID",
               "id" : "evidence.txt ID",
               "Phospho (STY) site IDs" : "Phospho (STY)Sites.txt ID"}

kegg_tcr_g_to_p = {"AKT2" : "Akt2",
                     "CARD11" : "CARD11",
                     "CBLB" : "Cbl-b",
                     "CD247" : r"TCR$\zeta$",
                     "CD28" : "CD28",
                     "CD3D" : r"CD3$\delta$",
                     "CD3E" : r"CD3$\epsilon$",
                     "CD3G" : r"CD3$\gamma$",
                     "CDC42" : "CDC42",
                     "CDK4" : "CDK4",
                     "CTLA4" : "CTLA-4",
                     "DLG1" : "DLG1",
                     "FYN" : "Fyn",
                     "GRAP2" : "GADS",
                     "GRB2" : "Grb2",
                     "GSK3B" : r"GSK3$\beta$",
                     "ITK" : "Itk",
                     "LAT" : "LAT",
                     "LCK" : "Lck",
                     "LCP2" : "SLP76",
                     "MAP3K7" : "TAK1",
                     "MAPK1" : "Erk2",
                     "MAPK3" : "Erk1",
                     "MAPK8" : "Jnk1",
                     "MAPK9" : "Jnk2",
                     "MAPK10" : "Jnk3",
                     "MAPK11" : r"p38$\beta$",
                     "MAPK12" : r"p38$\gamma$",
                     "MAPK14" : r"p38$\alpha$",
                     "NCK1" : "NCK1",
                     "NCK2" : "NCK2",
                     "NFATC2" : "NFAT1",
                     "NFATC3" : "NFAT4",
                     "NFKB1" : r"NF$\kappa$B-p105",
                     "PAK1" : "PAK1",
                     "PAK2" : "PAK2",
                     "PAK6" : "PAK6",
                     "PDCD1" : "PD-1",
                     "PIK3CA" : r"p110$\alpha$",
                     "PIK3CD" : r"p110$\delta$",
                     "PIK3R1" : r"p85$\alpha$", 
                     "PIK3R2" : r"p85$\beta$",
                     "PIK3R3" : r"p55$\gamma$",
                     "PLCG1" : r"PLC$\gamma$1",
                     "PRKCQ" : r"PKC$\theta$",
                     "PTPN6" : "SHP-1",
                     "PTPRC" : "CD45",
                     "RHOA" : "RHOA",
                     "TEC" : "Tec",
                     "VAV1" : "Vav1",
                     "VAV2" : "Vav2",
                     "VAV3" : "Vav3",
                     "ZAP70" : "Zap70"
                     }

kegg_tcr_p_to_g = {value : key for key, value in kegg_tcr_g_to_p.items()}

heads = ["gene", "h site", "h flank", "m site", "m flank"]

venn_colours = ["red","green"]

#
#
###################################################################################################
#
#  Functions

def pair(list_1, list_2):
    """
    Pair two lists based on elements
    """
    both, unique_1, unique_2 = [], [], []
    for element in list_1:
        if element in list_2:
            both.append(element)
        else:
            unique_1.append(element)
    for element in list_2:
        if element not in list_1:
            unique_2.append(element)
    return [both, unique_1, unique_2]

def unique(a_list):
    seen = []
    for item in a_list:
        if item not in seen:
            seen.append(item)
    return seen
    
def remove_p(site):
    if "-" in site:
        return site.split("-")[0]
    else:
        return site

def make_site_colours(combined_site_list,
                      both, unique_1, unique_2,
                      color1 = "red",
                      color2 = "green",
                      unseen = "grey",
                      gene = False,
                      genecolor = "purple"):
    colours = []
    if not gene:
        for item in combined_site_list:
            if item in both:
                colours.append([color1, color2])
            elif item in unique_1:
                colours.append([color1,unseen])
            elif item in unique_2:
                colours.append([unseen,color2])
    else:
        for item in combined_site_list:
            if item in both:
                colours.append([genecolor,color1, color2])
            elif item in unique_1:
                colours.append([genecolor,color1,unseen])
            elif item in unique_2:
                colours.append([genecolor,unseen,color2])
    return colours

def replace_repeats(a_list):
    seen = []
    for item in a_list:
        if item not in seen:
            seen.append(item)
        else:
            seen.append("")
    return seen


#
#
###################################################################################################
#
#  main()

def main():
    print("STAGE 1: Reading and Managing Data")
    print("\tReading 'chua_2020_tandem/evidence.txt'")
    xien_ev = gh.read_file("chua_2020_tandem/evidence.txt")
    print(f"\t\tTotal number of rows: {len(xien_ev)-1}\n")
    print("\tManaging the data")
    print(f"\t\tFiltering reverse and potential contaminants")
    xien_ev = gh.filter_matrix(gh.filter_matrix(xien_ev, "Reverse", "+", False),
                             "Potential contaminant", "+", False)
    print(f"\t\tDone. Rows remaining: {len(xien_ev)-1}")
    print("\t\tGrabbing desired columns...")
    xien_ev = gh.select_cols(xien_ev, rename_cols)
    print("\t\tDone. Columns remaining:")
    for item in xien_ev[0]:
        print(f"\t\t\t{item}")
    print("\t\tSelecting only the PV BOOST experiment (#2)")
    xien_ev = [xien_ev[0]] + [row for row in xien_ev[1:] if row[-3] == "2"]
    print(f"\t\tDone. Rows remaining:{len(xien_ev)-1}")
    mod_seq_col = xien_ev[0].index("Modified sequence")
    print(f"\t\tTransforming data to floats")
    xien_ev = [gh.transform_values(row) for row in xien_ev]
    for i in range(len(xien_ev)-1):
        xien_ev[i+1][-3] = int(xien_ev[i+1][-3])
        xien_ev[i+1][-1] = int(xien_ev[i+1][-1])
    print("\t\tChanging zeroes to nans")
    xien_ev = [gh.replace_value(item, 0, float("nan")) for item in xien_ev]
    print("\t\tAdding Unique ID: <Experiment>_<Modified sequence>_<Charge>")
    xien_ev = [item + [gh.list_to_str(item[-3:], delimiter = "", newline = False)] for item in xien_ev]
    xien_ev[0][-1] = "Unique ID"
    print("\t\tCounting missing values")
    xien_ev = [xien_ev[0] + [""]] + [item + [sum([1 for _ in item[:9] if _ != _])] for item in xien_ev[1:]]
    xien_ev[0][-1] = "Missing values"
    print("\t\tCalculating row median")
    xien_ev = [xien_ev[0] + [""]] + [item + [sh.median(item[:11])] for item in xien_ev[1:]]
    xien_ev[0][-1] = "Median"
    print("\t\tSorting by Unique ID, missing values, and median intensity (descending).")
    xien_ev = [xien_ev[0]] + sorted(xien_ev[1:], key = lambda x: (x[-3],x[-2],-x[-1]))
    print("\t\tRemoving duplicates...")
    xien_ev = [xien_ev[0]] + keep_first(xien_ev[1:], -3)
    print(f"\t\t{len(xien_ev)-1} remaining\n")
    xien_ev = [item + ["pY" in item[mod_seq_col]] for item in xien_ev]
    xien_ev[0][-1] = "ispY"
    for item in xien_ev[1:]:
        if item[-7] == 2:
            item[-7] = "PV"
        else:
            item[-7] = "no PV"
    experiments = gh.bin_by_col(xien_ev[1:], xien_ev[0].index("Experiment"))
    experiments = {key : [xien_ev[0]] + value for key, value in experiments.items()}
    pv_exp = experiments["PV"]
    print("\t\tGrabbing Unique IDs")
    ev_uniqueid = pv_exp[0].index("Unique ID")
    print("\tGrabbing information from Xien's BOOST experiment (Supp Tab 3 and Supp Tab 5)")
    print("\t\tReading 'chua_2020_tandem/chua_2020_tandem_st3_st5.txt'")
    xienmcp = gh.read_file("chua_2020_tandem/chua_2020_tandem_st3_st5.txt")
    print("\t\tGrabbing Unique IDs from Xien's Supplementary Tables")
    xien_uniqueid = xienmcp[0].index("uniqueID")
    uniqueids = gh.transform_values([row[xien_uniqueid] for row in xienmcp[1:]], transform = float)
    print("\t\tRetaining data found in Xien's Supporting Tables 3 and 5")
    pv_filt = [pv_exp[0]] + [row for row in pv_exp[1:] if row[ev_uniqueid] in uniqueids]
    print("\tPreparing flanking sequences for Jurkat Data")
    print("\t\tReading 'chua_2020_tandem/Phospho (STY)Sites.txt'")
    sty = gh.read_file("chua_2020_tandem/Phospho (STY)Sites.txt")
    sty = [gh.transform_values(row) for row in sty]
    sty = [gh.replace_value(row, "", float("nan")) for row in sty]
    print("\t\tReading 'database/phosphositeplus_ptm_data.txt'")
    psp = gh.read_file("database/phosphositeplus_ptm_data.txt")[3:]
    psp_database = gh.bin_by_col(psp, psp[0].index("ORGANISM"))
    # Grab the index of the localisation probability column
    loc_col = pv_exp[0].index("Localisation Probability (Sequence)")
    # Aggregate the data we want for the new sheets
    new_sheets = [pv_filt]
    print("\t\tAdding localisation probabilities column")
    new_sheets = [make_loc_probs(matrix, "Localisation Probability (Sequence)") for matrix in new_sheets]
    print("\t\tDone\n")
    print("\t\tAdding sequence windows and flanking sequences")
    new_sheets[0] = add_sequence_window(new_sheets[0], sty)
    print("\t\tDone\n")
    print("\t\tAdding site numbers using PSP database")
    new_sheets = compare_seqs_to_database(psp_database["human"], new_sheets)
    print("\t\tDone\n")
    print("\t\tAdding Mouse equivalent sites")
    new_sheets = get_other_species_sites(psp_database["mouse"], new_sheets,
                                         organism = "mouse")
    print("\t\tDone\n")
    print("\t\tPreparing to write the Jurkat data")
    newsheets = [gh.select_cols(new_sheets[0], keep_boost_cols)]
    newsheets = [[gh.transform_values(row) for row in sheet] for sheet in newsheets]
    newsheets = [[matrix[0]] + sorted(matrix[1:], key = lambda x: (x[0],
                                                                   x[4],
                                                                   -x[6])) for matrix in newsheets]
    newsheets = [[gh.list_to_str(row,
                                  delimiter = "\t",
                                  newline = True) for row in matrix]
                  for matrix in newsheets]
    names = ["chua_2020_tandem/chua_2020_tandem_after_flanking.txt"]
    i=0
    for sheet in newsheets:
        print(f"t\t\tWriting {names[i]}\n")
        gh.write_outfile(sheet, filename = names[i], writestyle = "w")
        i+=1
    newsheets = [[string.rstrip("\n").split("\t") for string in matrix] for matrix in newsheets]
    print("\tGrabbing the Mouse BOOST Data")
    print("\t\tReading 'curated_results/boost_gained_peptides.txt'")
    bgp = gh.read_file("curated_results/boost_gained_peptides.txt", delim = "\t")
    print("\t\tReading 'curated_results/boost_control_peptides.txt'")
    bcp = gh.read_file("curated_results/boost_control_peptides.txt", delim = "\t")
    print("\t\tGrabbing the flanking sequences from Mouse and Jurakts")
    # Merge the BOOST mouse data
    mouse_boost = bgp + bcp[1:]
    # Grab all the flanking sequences
    mouse_flank_ind = mouse_boost[0].index("Flanking Sequence")
    mouse_flanks = [row[mouse_flank_ind].lower() for row in mouse_boost[1:]]
    xien_flank_ind = newsheets[0][0].index("Flanking Sequence")
    xien_flanks = [row[xien_flank_ind].lower() for row in newsheets[0][1:]]
    print("\t\tReading 'database/PSdataTRH.txt', a manually curated comparison of phosphosites")
    print("\t\tusing the PhosphoSitePlus website (www.phosphosite.org)")
    kegg_tcr = gh.read_file("database/PSdataTRH.txt", delim = "\t")
    # Make a site id for each of the sites
    kegg_tcr = [row + [gh.list_to_str(row, delimiter = ";", newline = False)] for row in kegg_tcr]
    # Lowercase columns 2 and 4
    for row in kegg_tcr:
        row[2] = row[2].lower()
        row[4] = row[4].lower()
    print("\t\tFinding overlap between Jurkats and Mouse KEGG TCR Sites")
    # Find the Human sites
    human = [row for row in kegg_tcr if row[2] in xien_flanks]
    # Find the Mouse sites
    mouse = [row for row in kegg_tcr if row[4] in mouse_flanks]
    compared = pair(mouse,human)
    compared = [unique(comp) for comp in compared]
    for item in compared:
        print(len(item))
    compared = [[[remove_p(item) for item in row]for row in a_list]for a_list in compared]
    print("STAGE 2: Plotting Overlap Venn Diagram")
    fig, ax = plt.subplots(figsize = (3,3))
                         # Mouse           # Human          # Both
    v = venn2(subsets = (len(compared[1]), len(compared[2]), len(compared[0])),
              set_labels = ("",""),
              set_colors = venn_colours)
                             # Mouse           # Human          # Both
    venn2_circles(subsets = (len(compared[1]), len(compared[2]), len(compared[0])),
                  linewidth = 1)
    for text in v.subset_labels:
        text.set_fontfamily("sans-serif")
        text.set_font("Arial")
        text.set_fontweight("bold")
        text.set_fontsize(16)
    plt.savefig("figures/human_mouse_overlap.pdf")
    plt.show()
    print("\tDone\n")
    print("STAGE 3: Writing the Unique Site Tables")
    heads = ["Protein", "h site", "h flank", "m site", "m flank"]
    comp_with_heads = [heads] + sorted(compared[0] + compared[1] + compared[2], key=lambda x: (x[0],int(x[1][1:])))
    comp_no_heads = comp_with_heads[1:]
    compared_prot = [[[kegg_tcr_g_to_p[row[0]]] + row[1:] for row in comp] for comp in compared]
    print("\tAggregating Comparison data")
    comp_no_heads = [[kegg_tcr_g_to_p[row[0]]] + row[1:] for row in comp_no_heads]
    comp_no_heads = sorted(comp_no_heads, key = lambda x: x[0].lower())
    print("\tBinning data into 37 groups")
    cols = [comp_no_heads[i*37:(i+1)*37] for i in range(4)]
    for i in range(3):
        cols[-1].append(["","","","",""])
    cols = [[[row[0],row[3],row[1]] for row in sect] for sect in cols]
    newcols = []
    for i in range(37):
        newrow = []
        for j in range(len(cols)):
            newrow += cols[j][i]
        newcols.append(newrow)
    newcols = gh.transpose(*newcols)
    for i in range(4):
        newcols[i*3] = replace_repeats(newcols[i*3])
    print("\tAssigning colours to each site and gene")
    ncolours = make_site_colours(comp_no_heads, compared_prot[0],compared_prot[1],compared_prot[2], gene = True)
    print(len(ncolours))
    colours = [ncolours[i*37:(i+1)*37] for i in range(4)]
    for i in range(3):
        colours[-1].append(["grey","grey","grey"])
    newcolours = []
    for i in range(37):
        newrowc = []
        for j in range(len(colours)):
            newrowc += colours[j][i]
        newcolours.append(newrowc)
    newcolours = gh.transpose(*newcolours)
    print("\tMaking the table")
    fig1,ax1,tab = mph.make_mpl_table(*newcols,colours=newcolours,
                                       colLabels = ["Protein", "Mouse", "Human","Protein", "Mouse", "Human","Protein", "Mouse", "Human","Protein", "Mouse", "Human"])#,"Gene", "Mouse", "Human"])#,"Gene", "Mouse", "Human","Gene", "Mouse", "Human"])
    ax1.spines["top"].set_visible(False)
    ax1.spines["bottom"].set_visible(False)
    ax1.spines["left"].set_visible(False)
    ax1.spines["right"].set_visible(False)
    ax1.set_xticks([])
    ax1.set_yticks([])
    plt.savefig("figures/sites/total.pdf")
    plt.show()
    print("\t\tDone\n")
    print("\tMaking Gene-specific tables")
    sites_by_gene = gh.bin_by_col(comp_with_heads,0)
    sites_by_gene = {key: value[1:] for key, value in sites_by_gene.items() if key != "gene"}
    sites_by_gene = {key : [[remove_p(item) for item in row]for row in value] for key, value in sites_by_gene.items()}
    for key, value in sites_by_gene.items():
        newval = sorted(value, key = lambda x: int(x[1][1:]))
        mousesites = [item[3] for item in newval]
        humansites = [item[1] for item in newval]
                                            #both       #mouse      #human 
        gene_spec_colours = make_site_colours(newval, compared[0],compared[1],compared[2])
        fig2, ax2 = plt.subplots()
        fig2,ax2, tab = mph.make_mpl_table(mousesites,
                                           humansites,
                                           colours = gh.transpose(*gene_spec_colours),
                                            fig = fig2, ax = ax2)
        ax2.spines["top"].set_visible(False)
        ax2.spines["bottom"].set_visible(False)
        ax2.spines["left"].set_visible(False)
        ax2.spines["right"].set_visible(False)
        ax2.set_xticks([])
        ax2.set_yticks([])
        fig1.tight_layout()
        print(f"\t\t\tSaving file 'figures/sites/{key}_sites.pdf'")
        plt.savefig(f"figures/sites/{key}_sites.pdf")
        plt.close()
    print("\tDone\n")
    return "Comparison complete!!!"

#
#
###################################################################################################
#
#

if __name__ == "__main__":
    a = main()
    print(a)

#
#
###################################################################################################
```
