## Supporting Folder 2 Data Analysis for "Mouse primary T cell phosphotyrosine proteomics enabled by BOOST": experimental_design.pdf

(A)

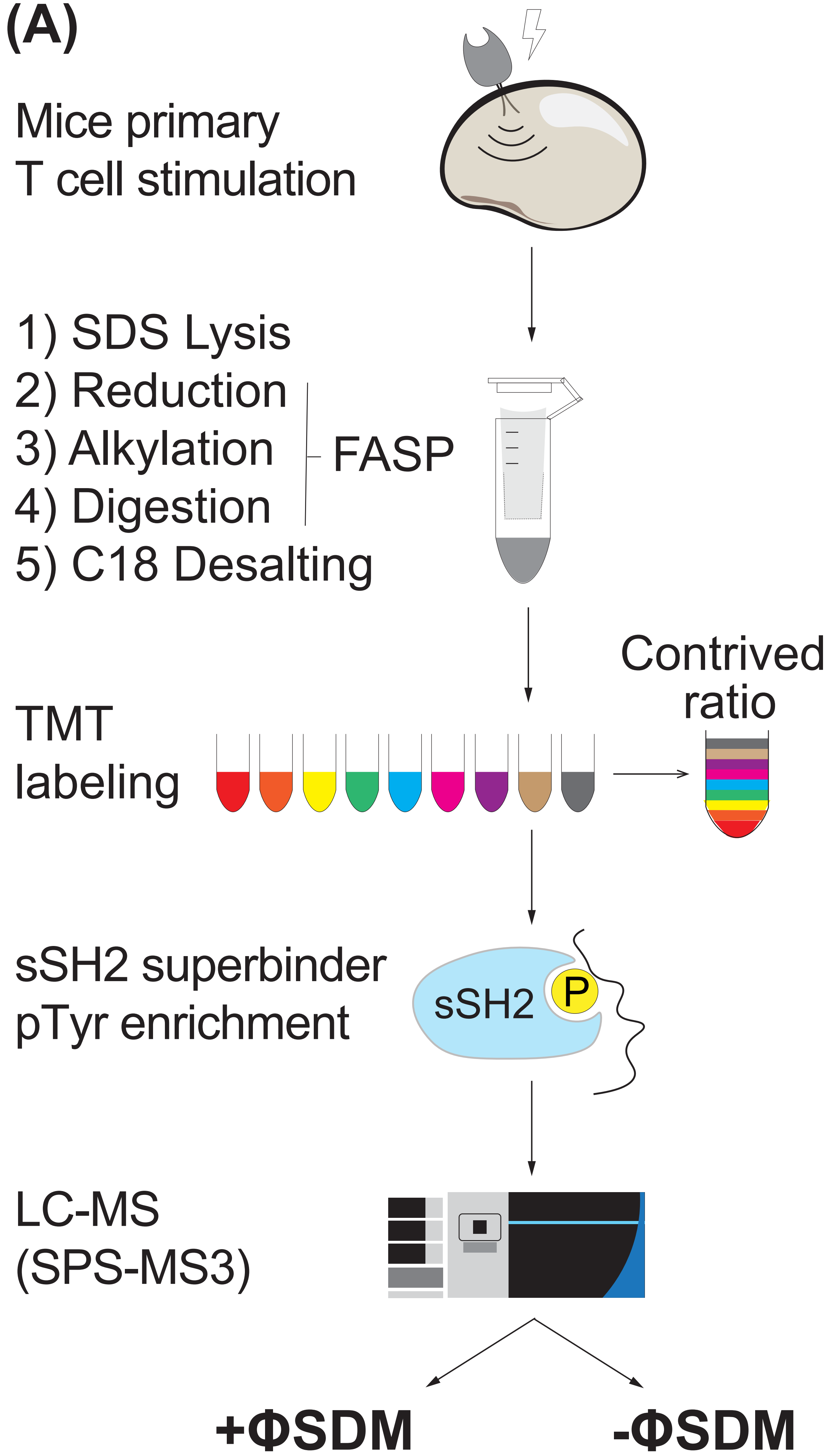


(B)

| TMT | Control +ΦSDM | Control | BOOST +ΦSDM | BOOST |
|---|---|---|---|---|
| 126 | 1 mg control | 1 mg control | 1 mg PV | 1 mg PV |
| 127N | 1.0 mg Rep.1 | 1.0 mg Rep.1 | 1.0 mg Rep.1 | 1.0 mg Rep.1 |
| 127C | Blank | Blank | Blank | Blank |
| 128N | 1.0 mg Rep.2 | 1.0 mg Rep.2 | 1.0 mg Rep.2 | 1.0 mg Rep.2 |
| 128C | 0.3 mg Rep.1 | 0.3 mg Rep.1 | 0.3 mg Rep.1 | 0.3 mg Rep.1 |
| 129N | 1.0 mg Rep.3 | 1.0 mg Rep.3 | 1.0 mg Rep.3 | 1.0 mg Rep.3 |
| 129C | 0.3 mg Rep.2 | 0.3 mg Rep.2 | 0.3 mg Rep.2 | 0.3 mg Rep.2 |
| 130N | 0.1 mg Rep.1 | 0.1 mg Rep.1 | 0.1 mg Rep.1 | 0.1 mg Rep.1 |
| 130C | 0.3 mg Rep.3 | 0.3 mg Rep.3 | 0.3 mg Rep.3 | 0.3 mg Rep.3 |
| 131N | 0.1 mg Rep.2 | 0.1 mg Rep.2 | 0.1 mg Rep.2 | 0.1 mg Rep.2 |
| 131C | 0.1 mg Rep.3 | 0.1 mg Rep.3 | 0.1 mg Rep.3 | 0.1 mg Rep.3 |
