## Supporting Folder 2 Data Analysis for "Mouse primary T cell phosphotyrosine proteomics enabled by BOOST": boost_factor_cdfs.pdf

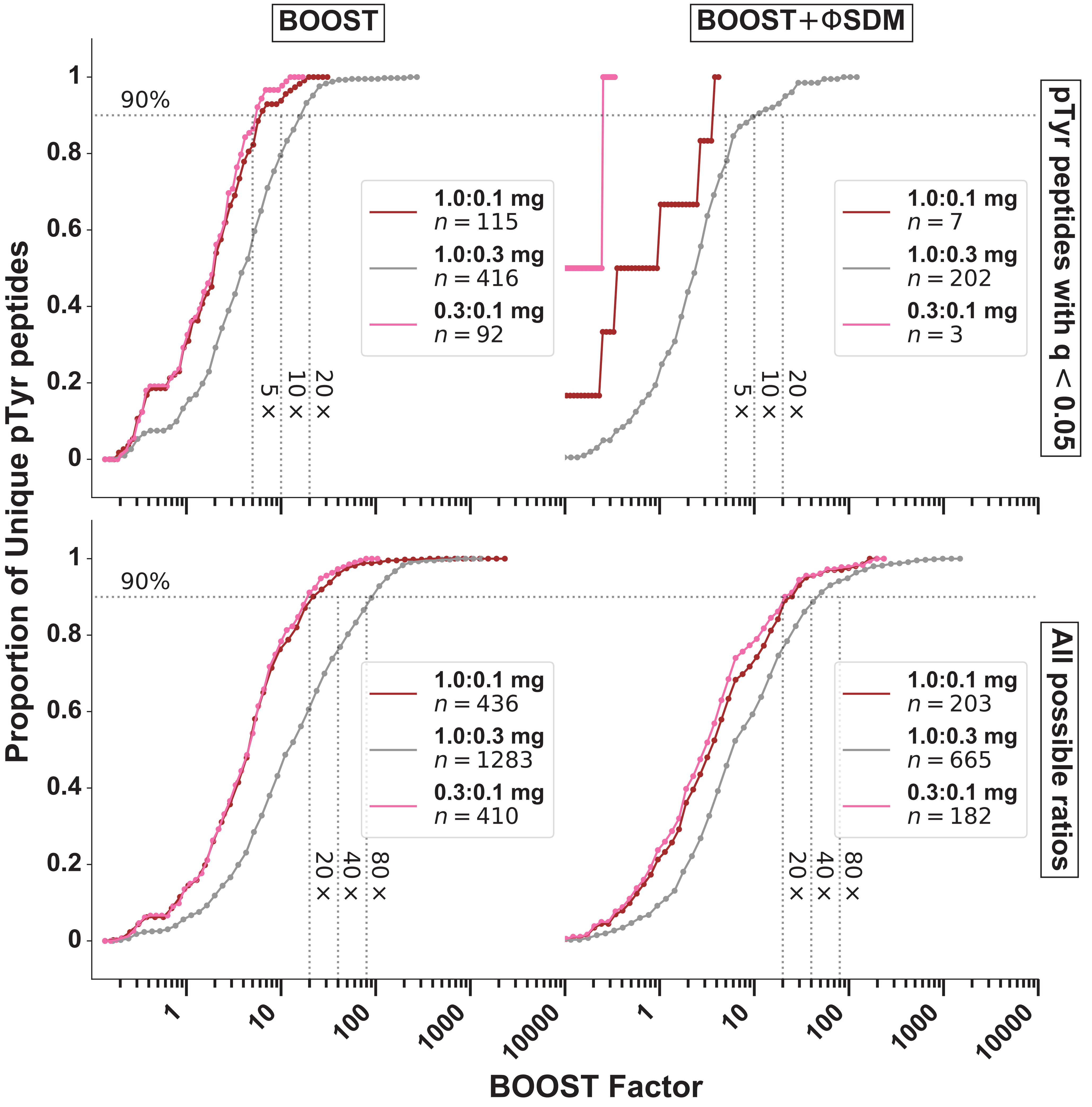


$$\text{BOOST Factor} = \frac{\text{Total reporter ion current}_{\text{BOOST}}}{\text{Total reporter ion current}_{\text{Experimental}}} \approx \frac{1\text{mg PV intensity}}{\text{Sum}(1\text{mg}_{\text{Rep1-3}} + 0.3\text{mg}_{\text{Rep1-3}} + 0.1\text{mg}_{\text{Rep1-3}}\ \text{intensity})}$$
