## Supplementary figures and images for "Mouse primary T cell phosphotyrosine proteomics enabled by BOOST"

### accuracy_and_precision.pdf

(A)

| | 1.0 mg | 0.3 mg | 0.1 mg |
|---|---|---|---|
| Control+ΦSDM | 0.977 | 0.941 | 0.56 |
| Control | 0.983 | 0.955 | 0.863 |
| BOOST+ΦSDM | 0.879 | 0.753 | 0.527 |
| BOOST | 0.894 | 0.869 | 0.775 |

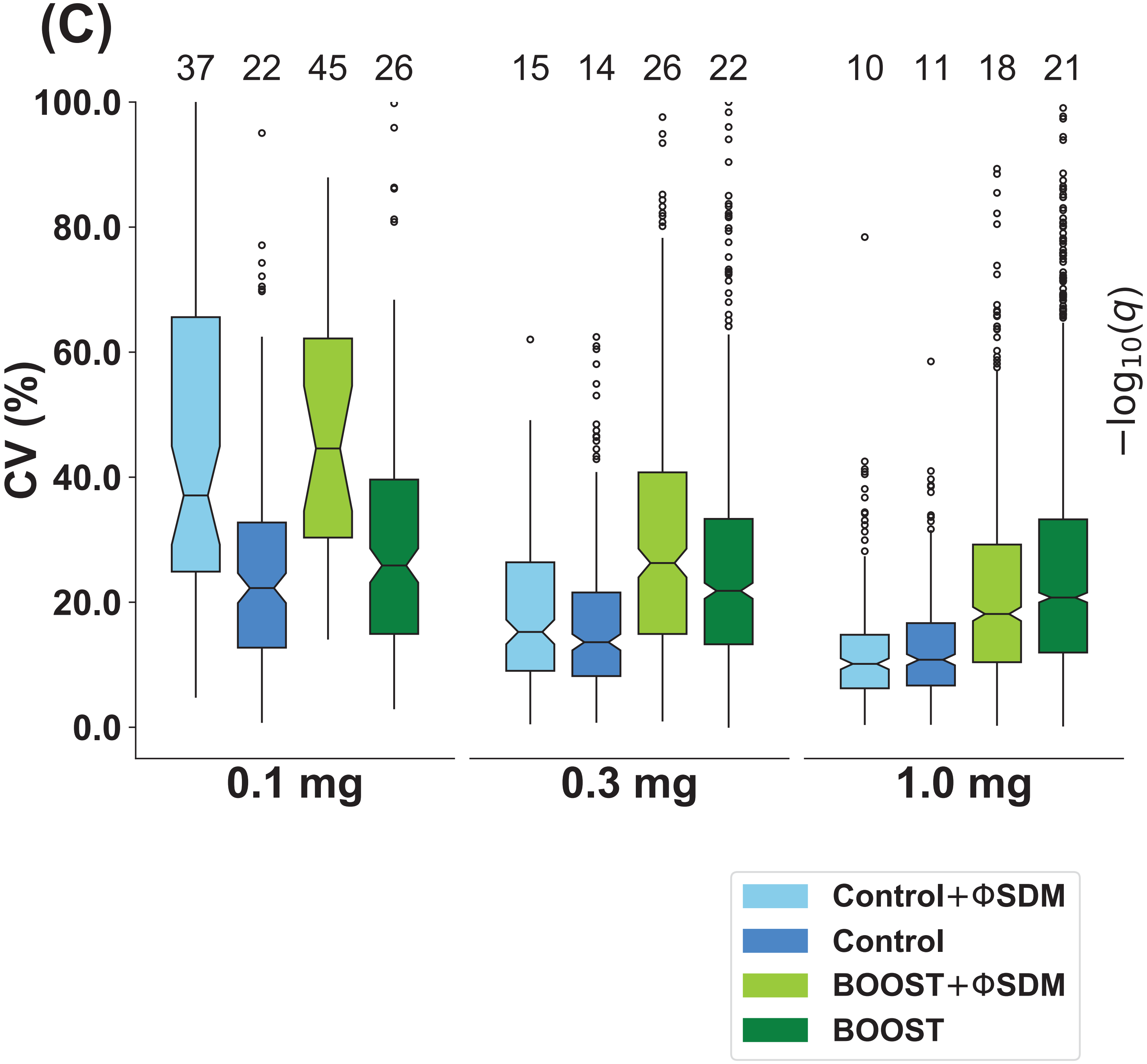


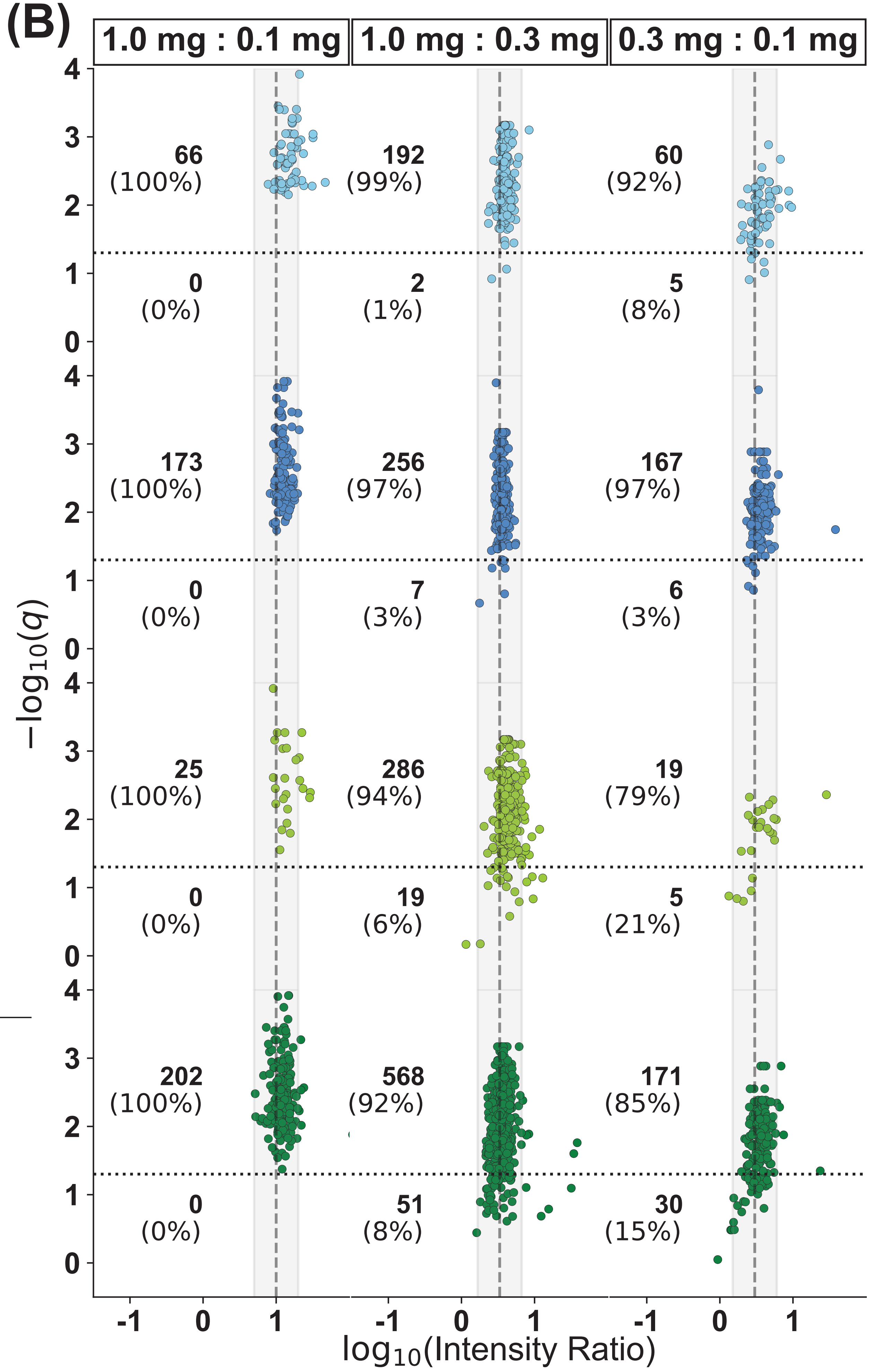

### biologically_interesting.pdf

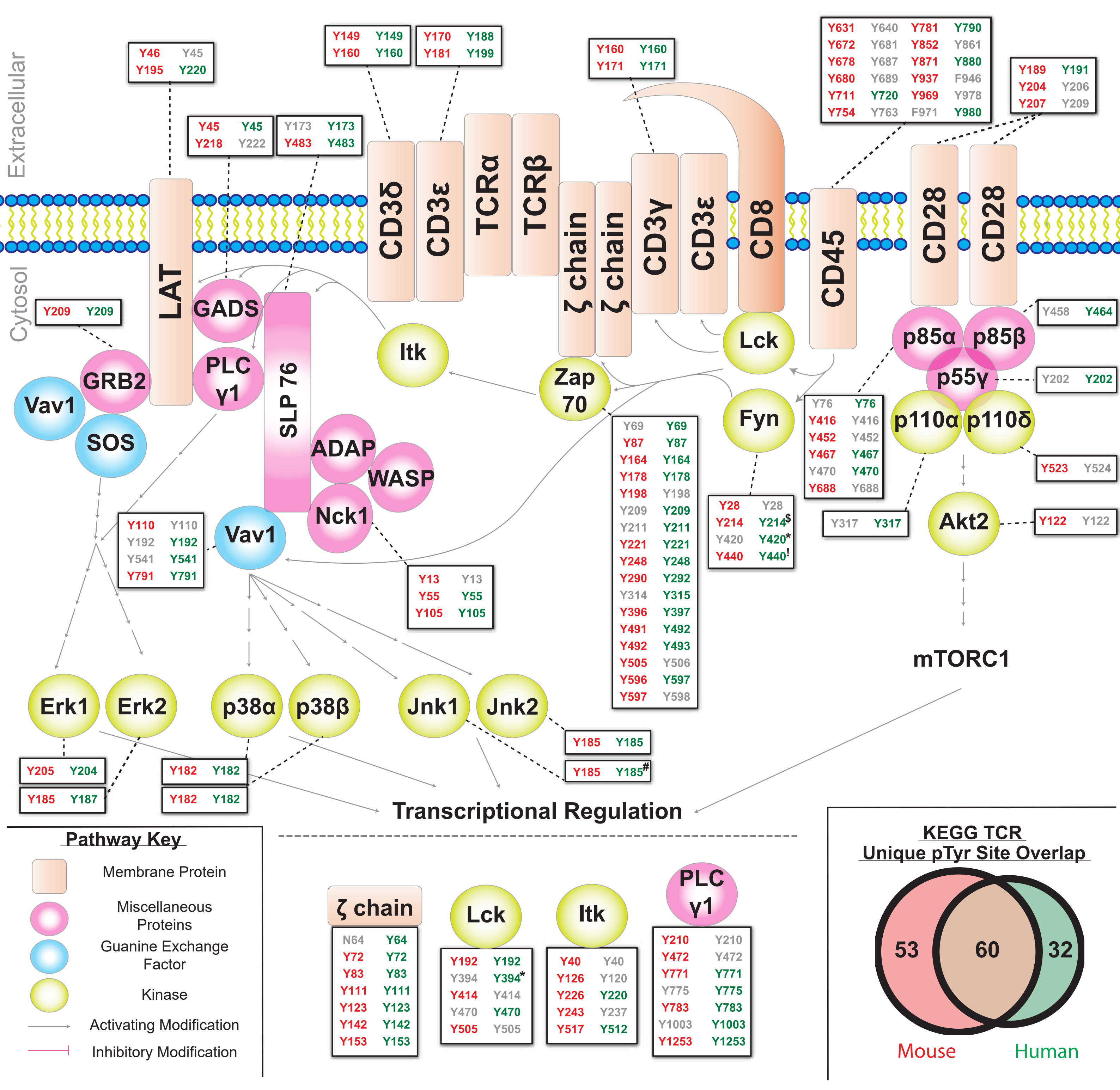

### boost_control_gained_qvolcanoes.pdf

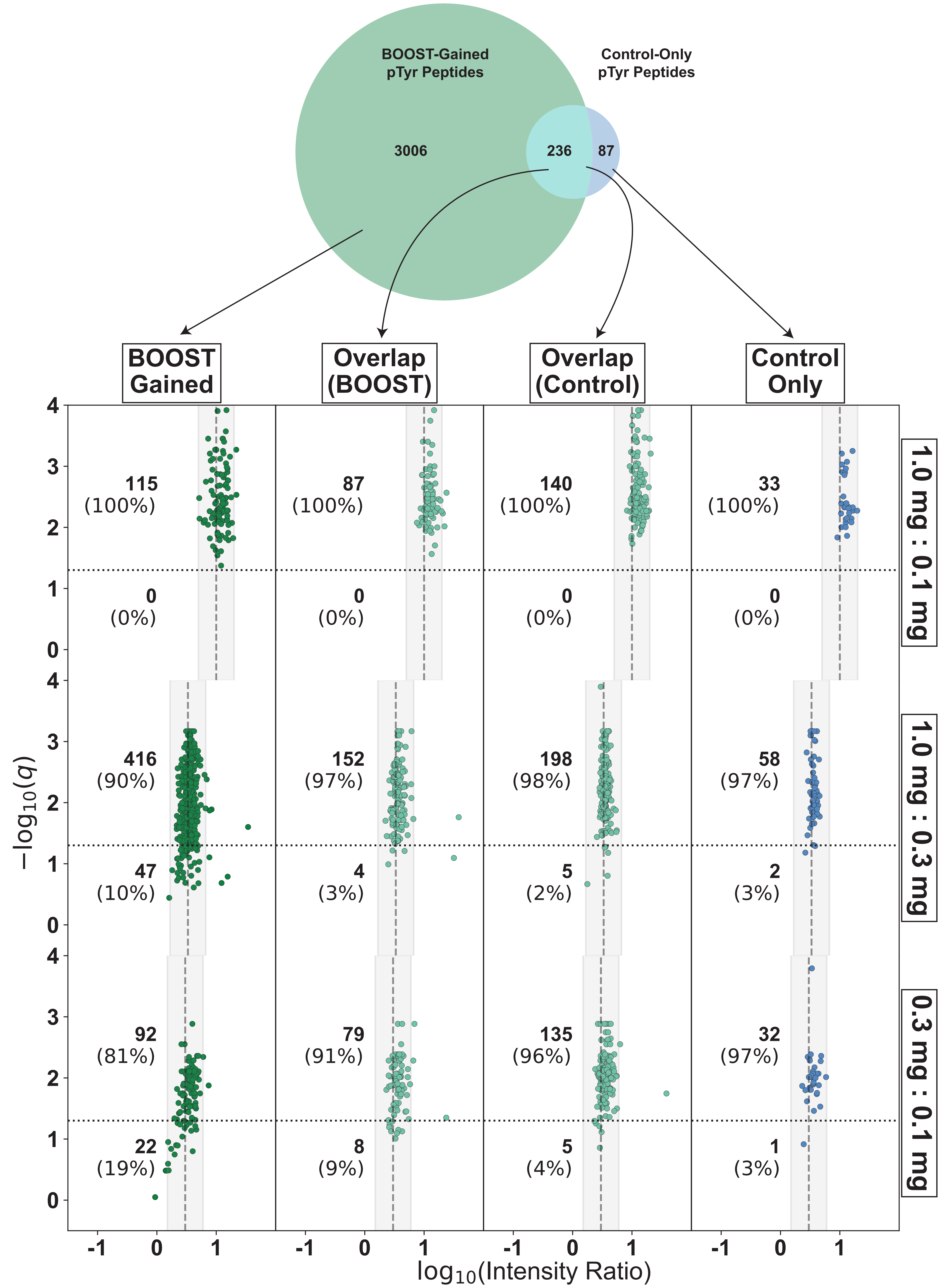

### boost_corrplot.pdf

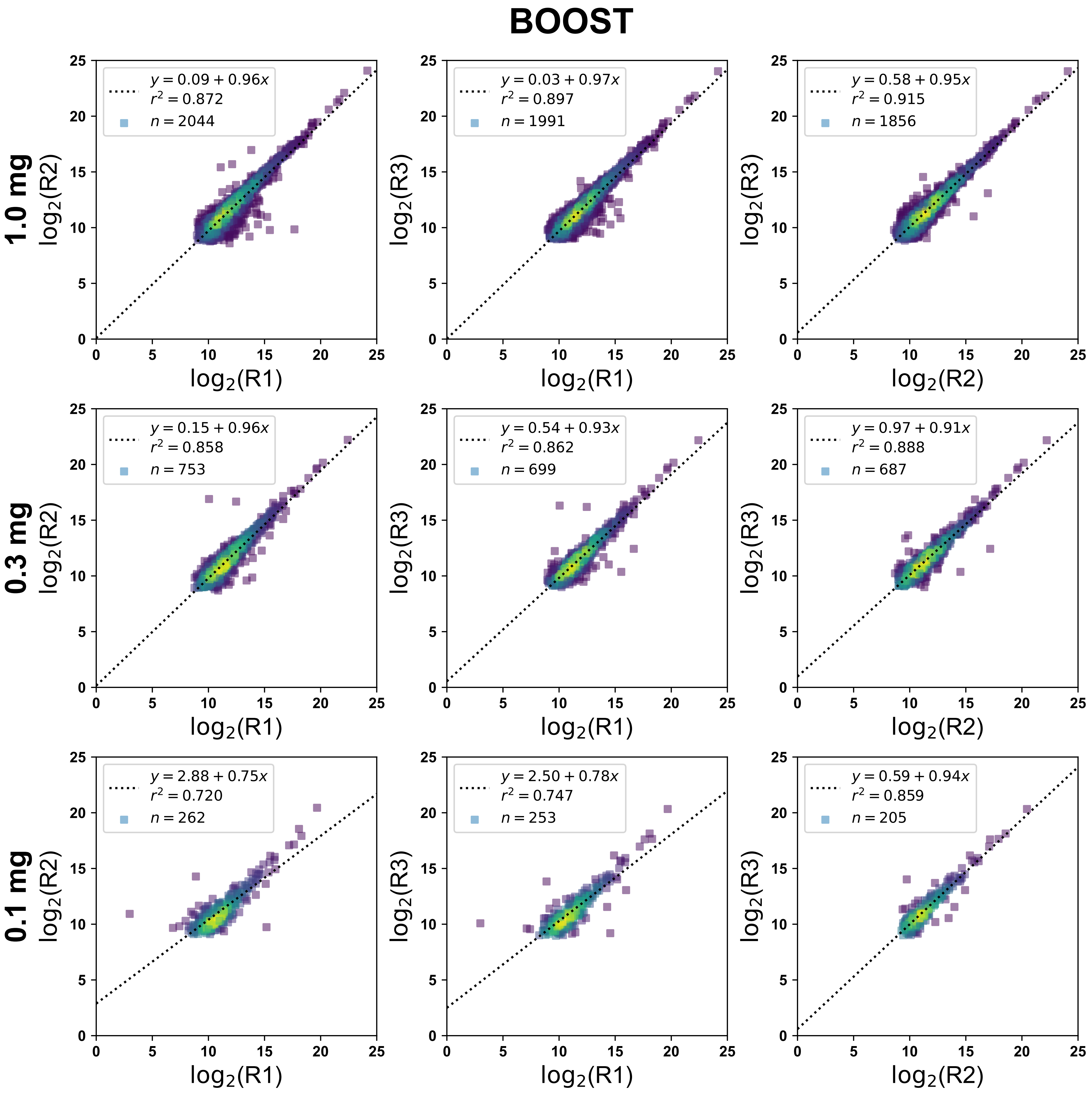

### boost_factor.pdf

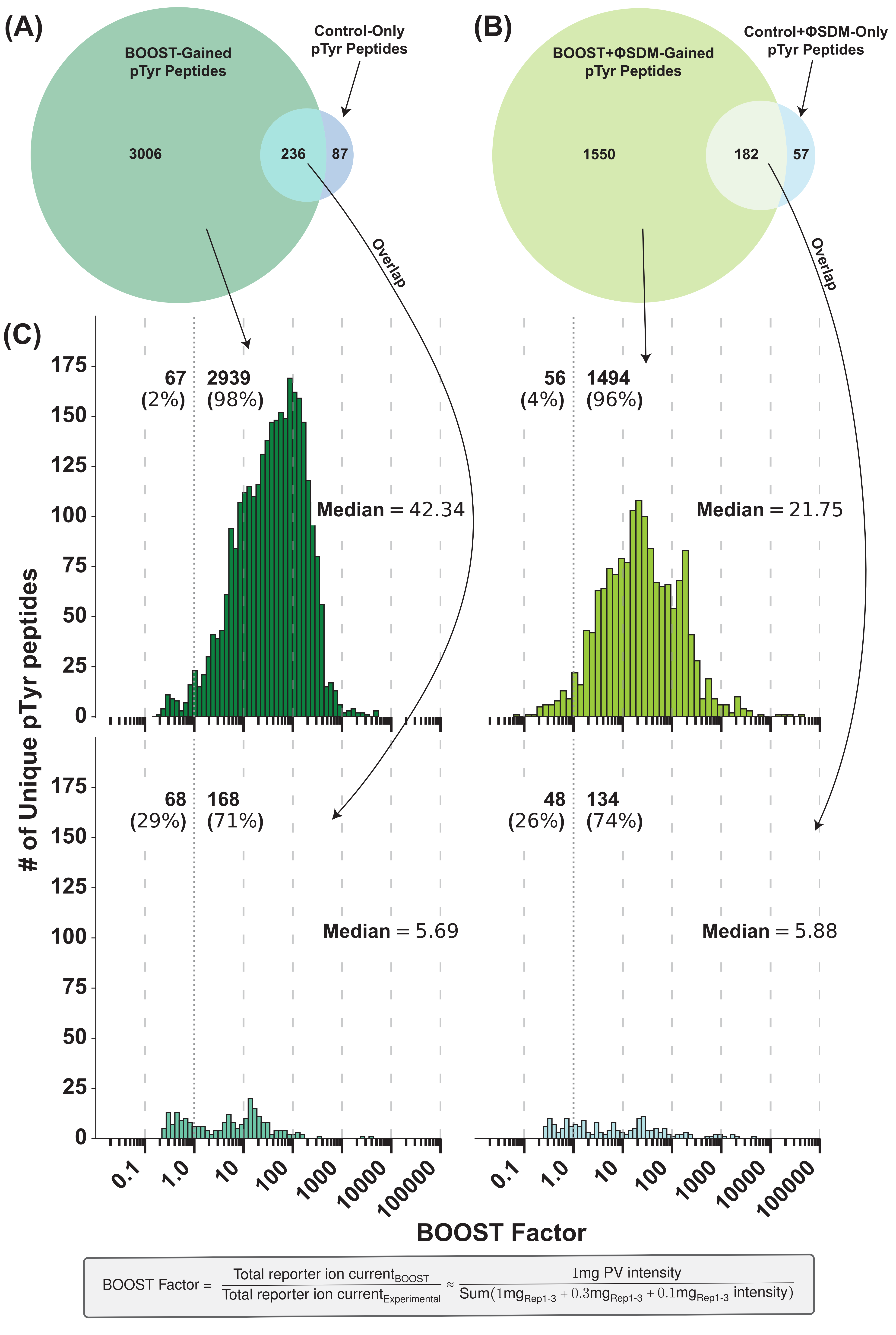

### boost_replicate_reprod.pdf

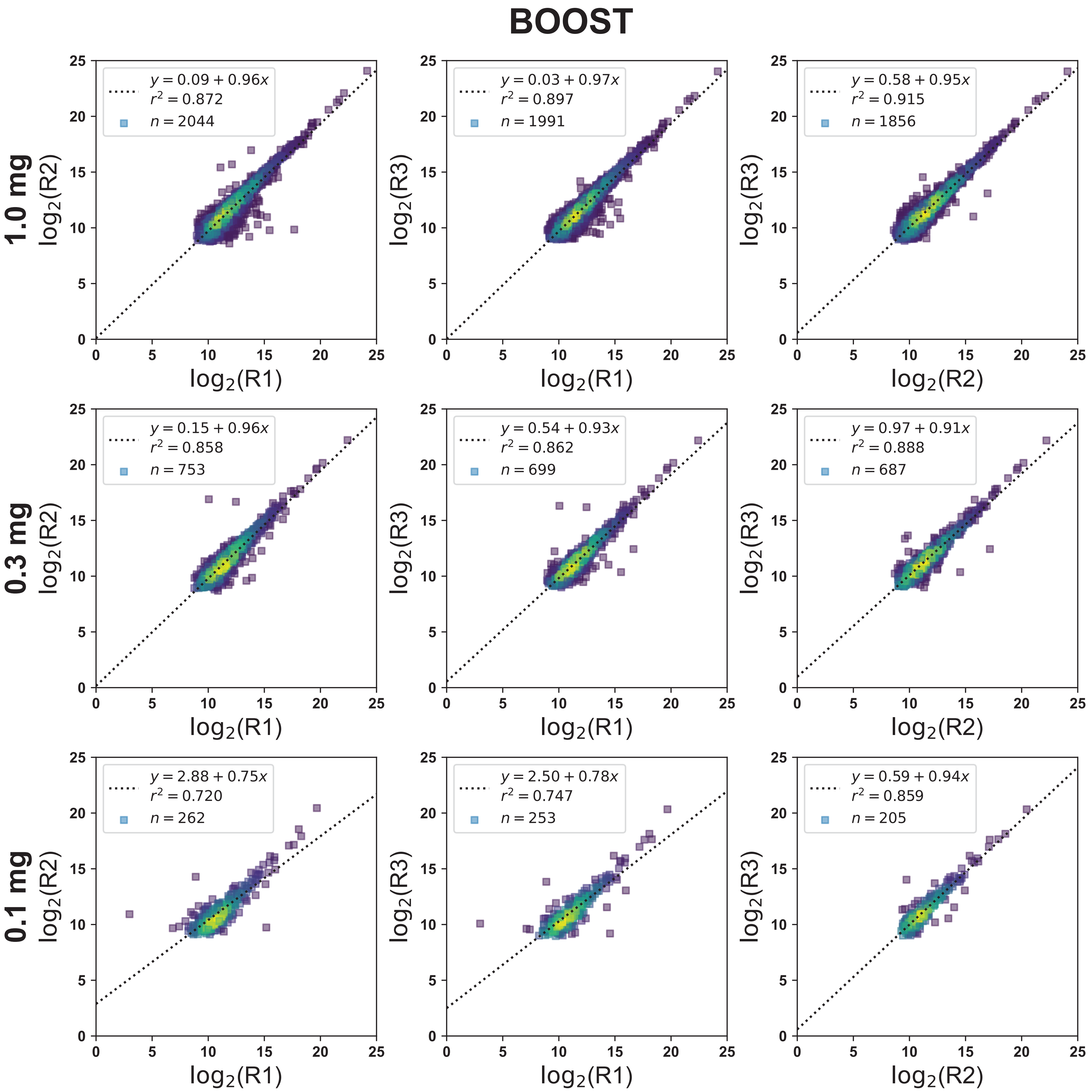

### boost_sdm_corrplot.pdf

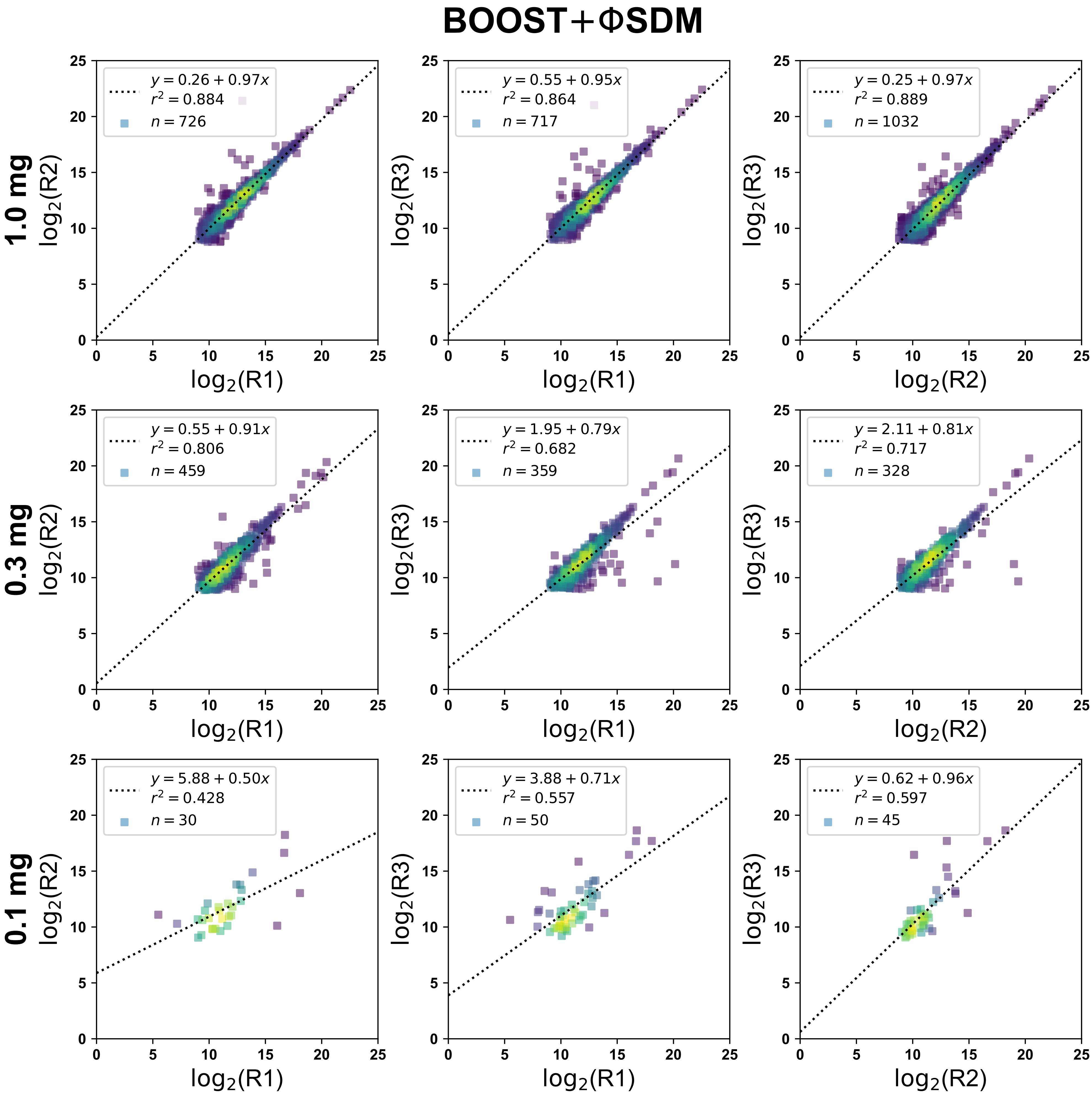

### boostfactor_cdf.pdf

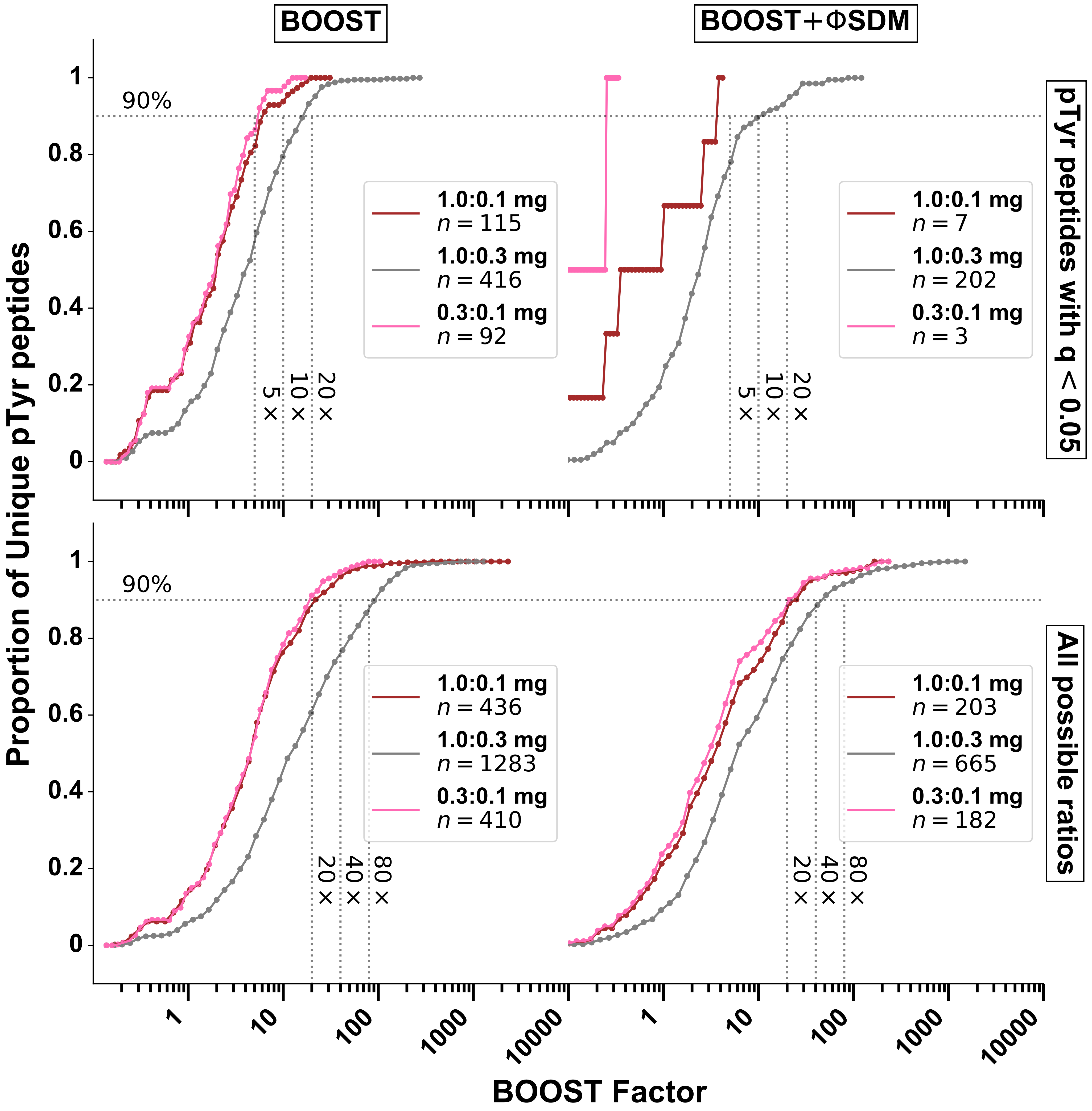

### boostfactor_hists.pdf

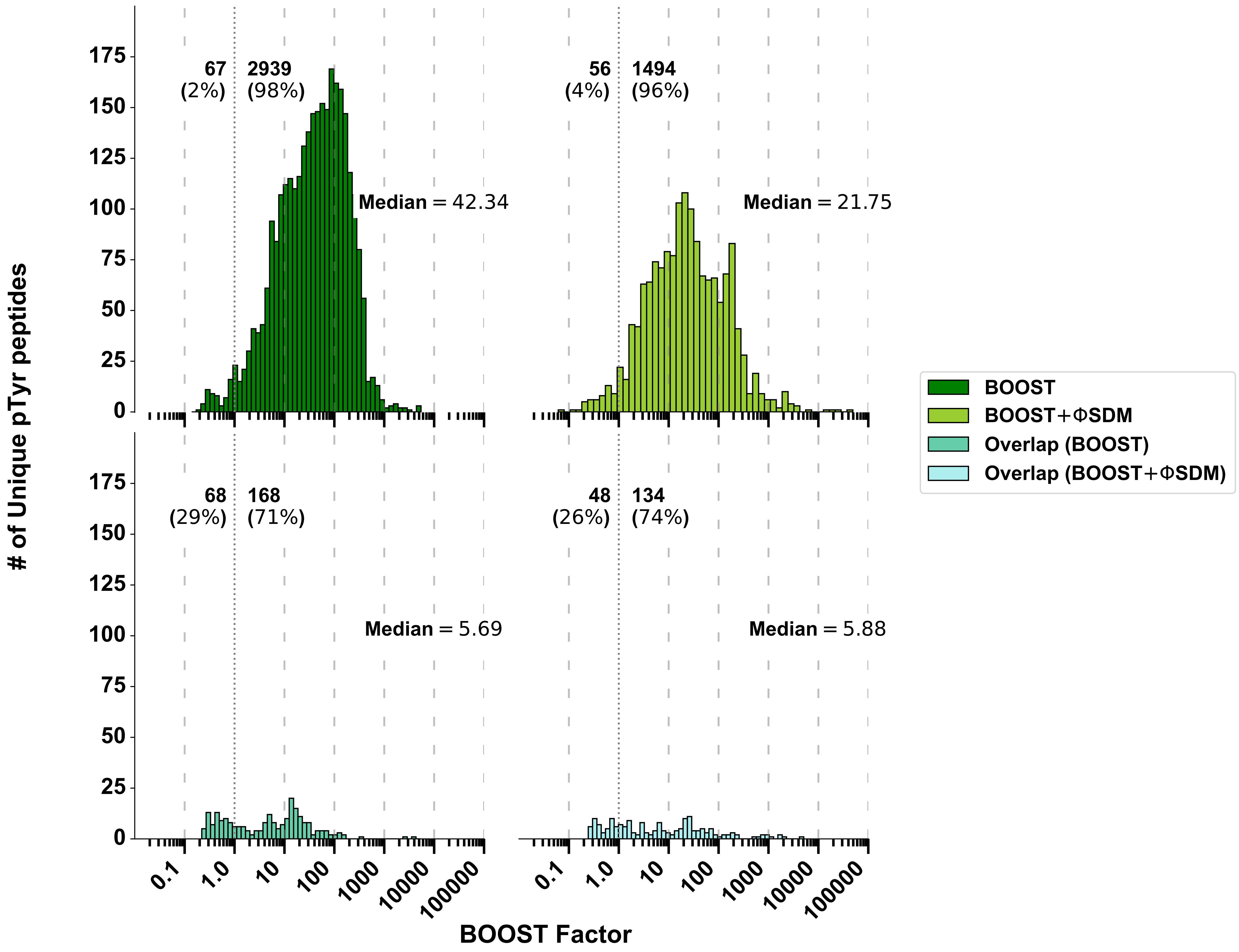

### boostsdm_controlsdm_gained_qvolcanoes.pdf

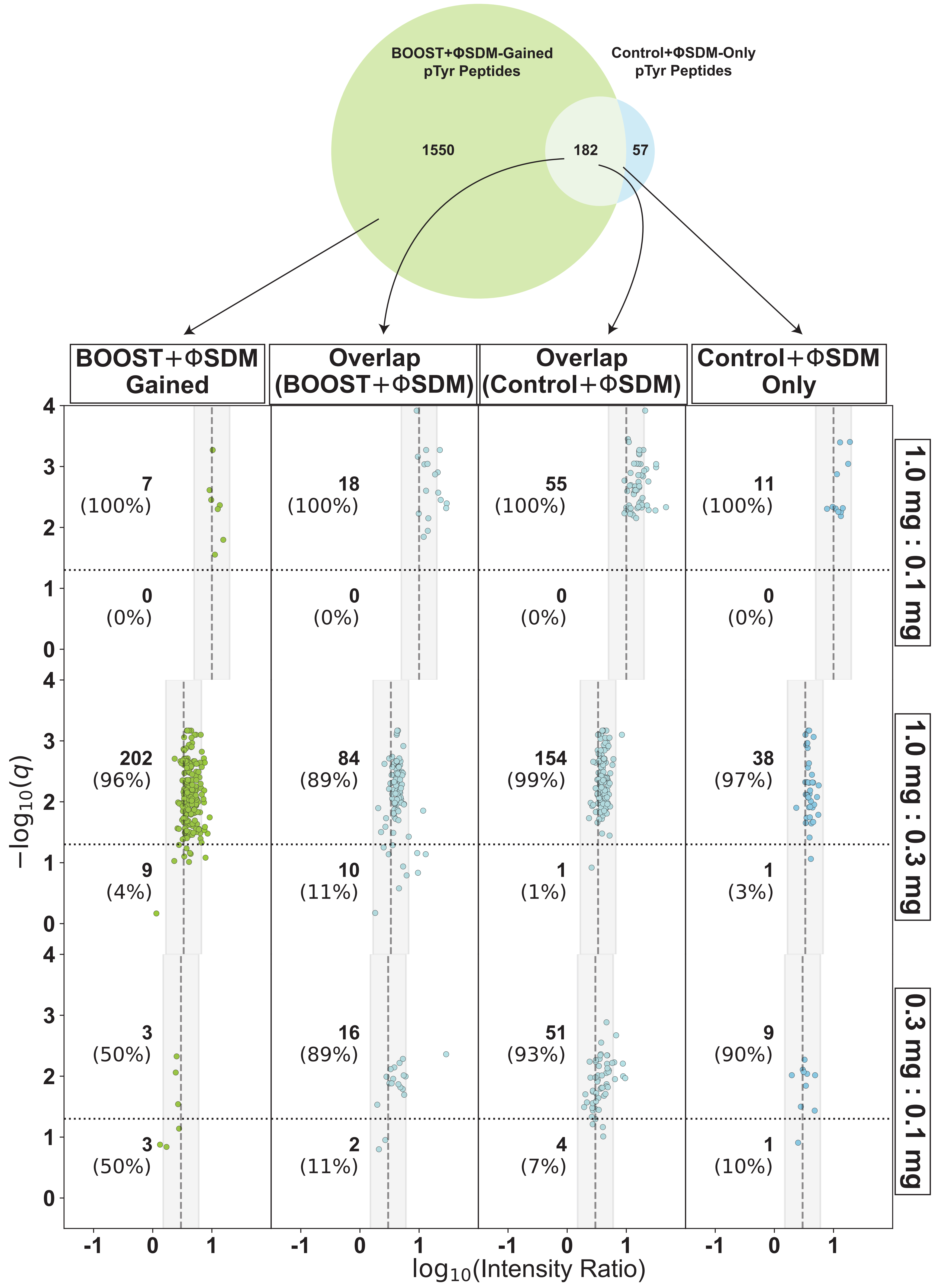

### boostsdm_replicate_reprod.pdf

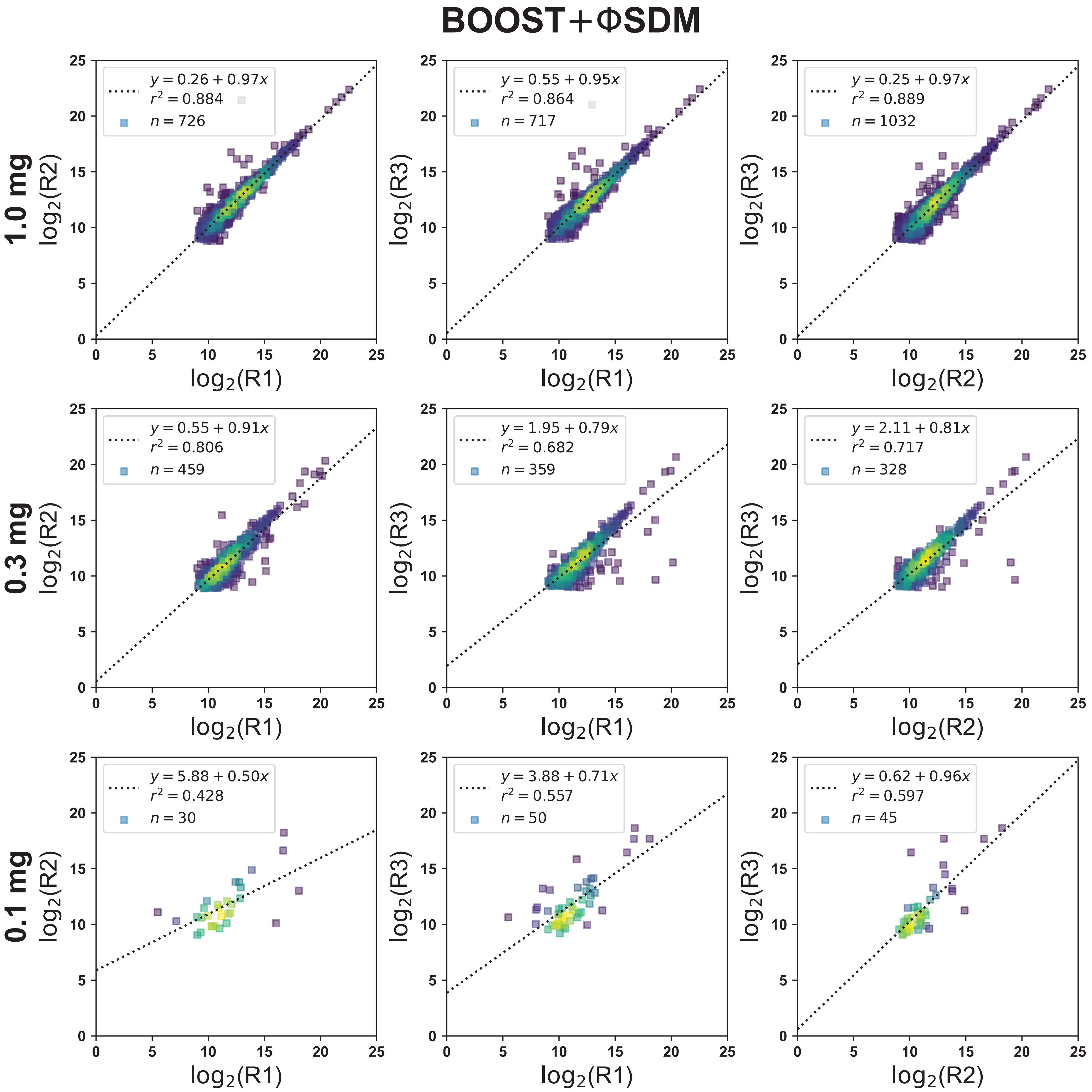

### control_corrplot.pdf

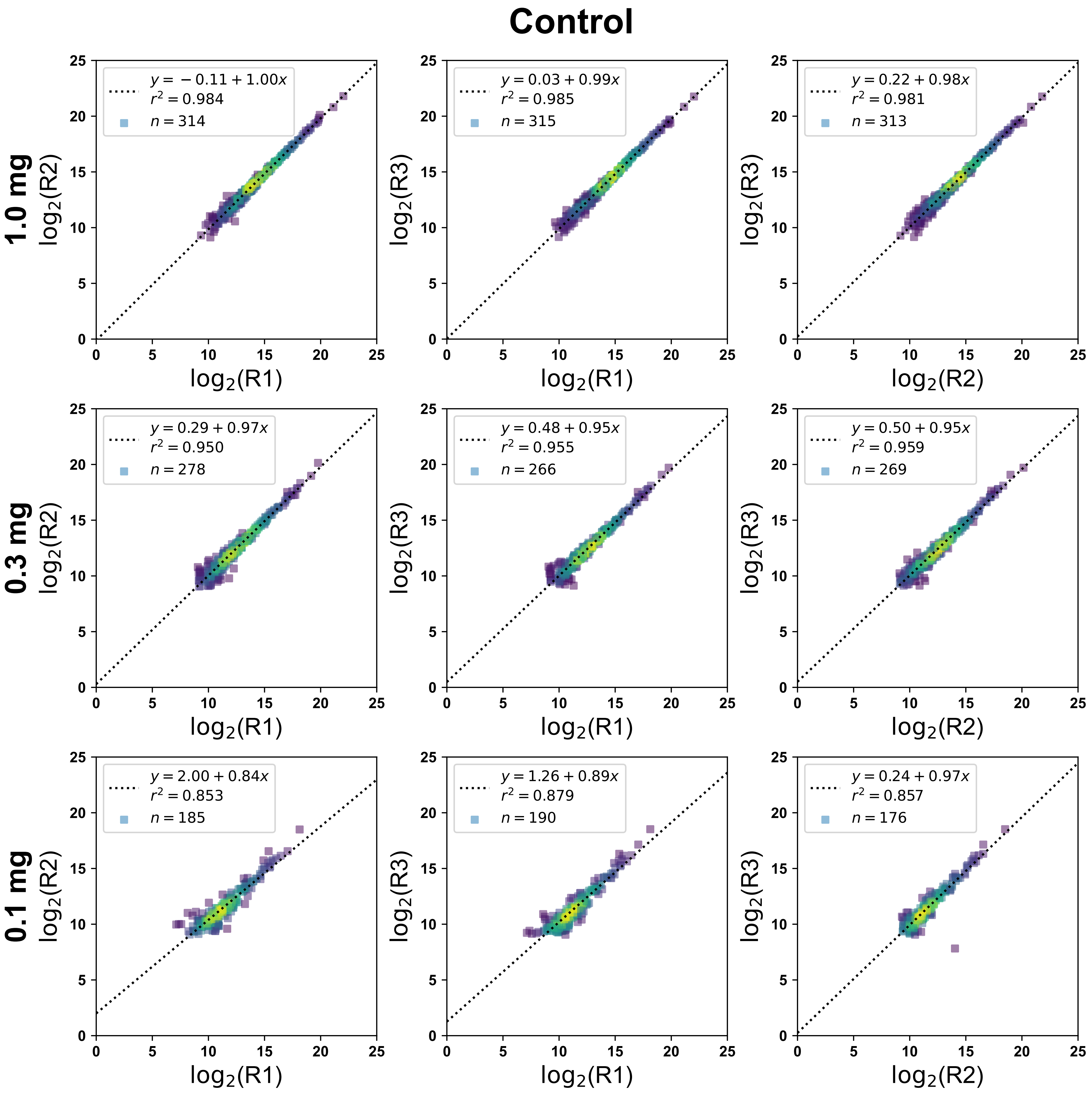

### control_replicate_reprod.pdf

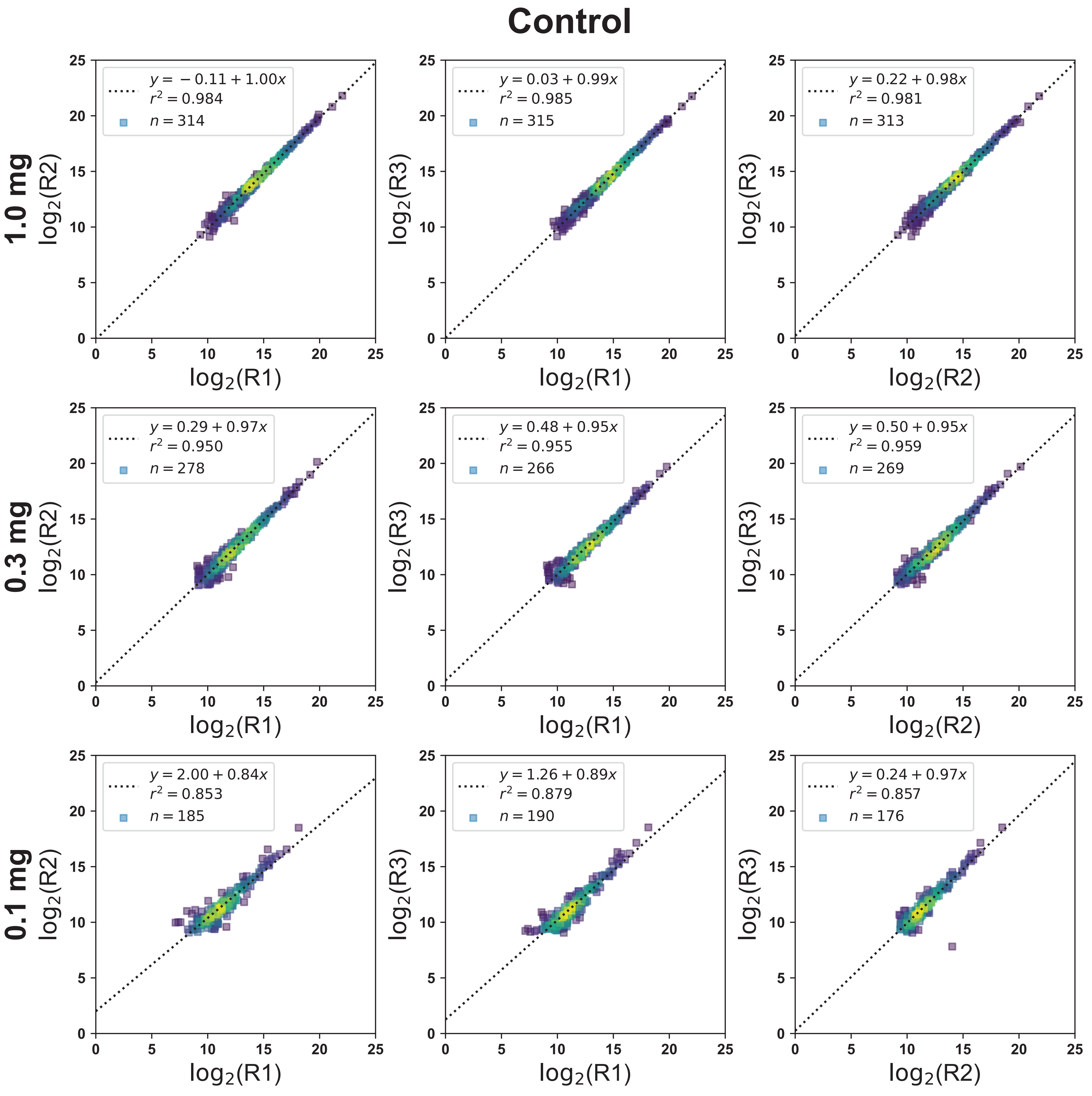

### control_sdm_corrplot.pdf

**Control+ΦSDM**

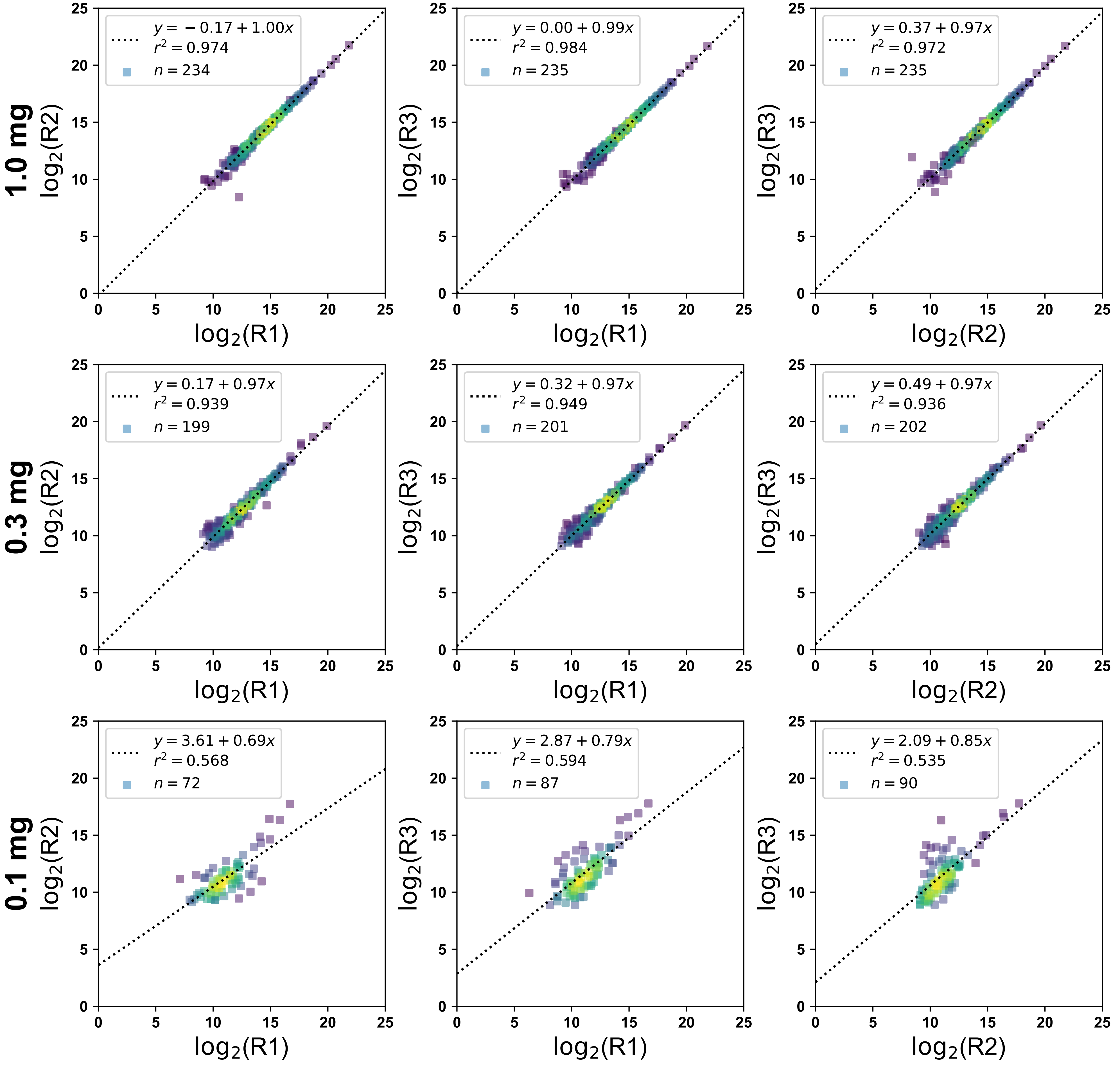

### controlsdm_replicate_reprod.pdf

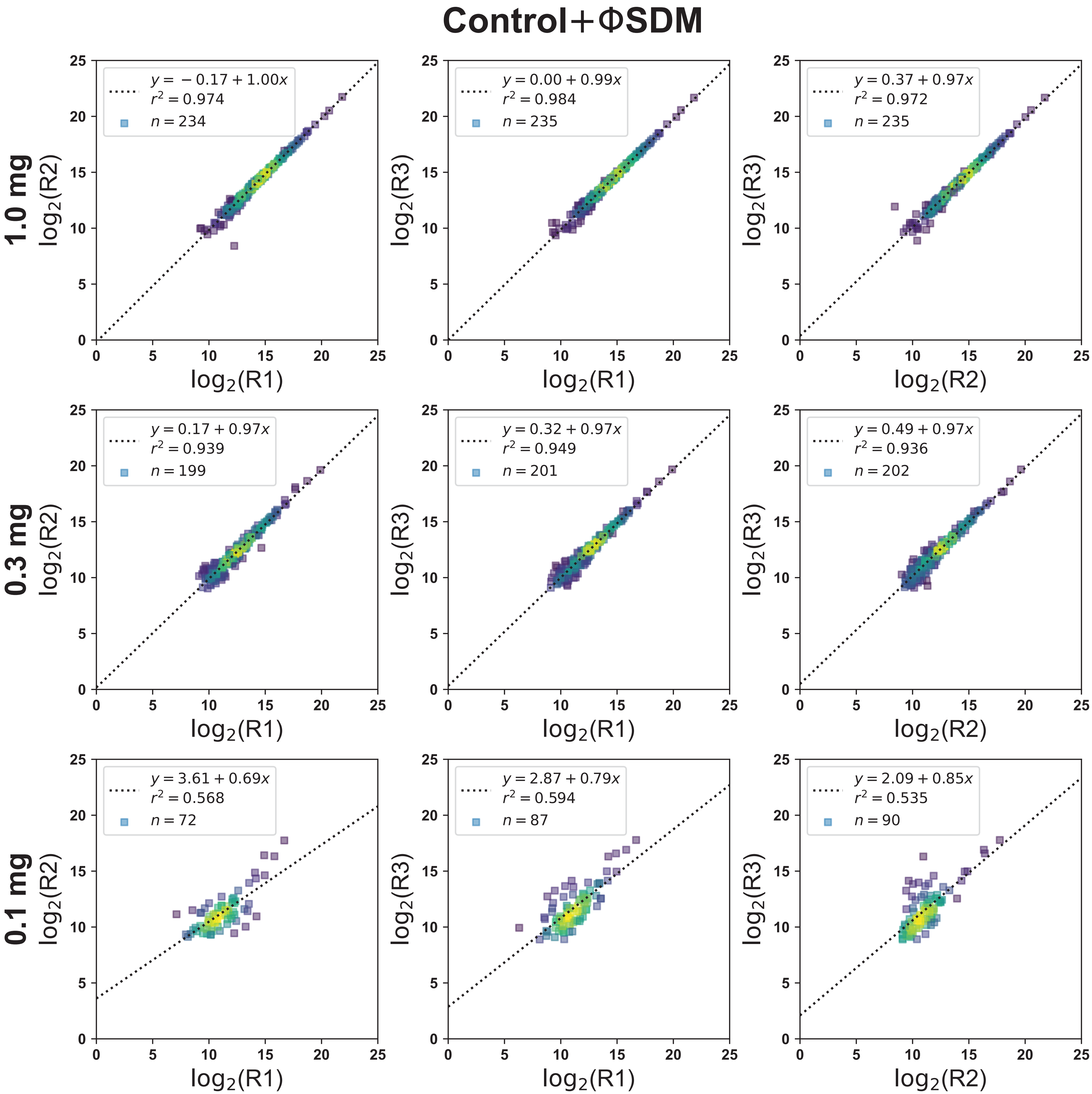

### cv_plot.pdf

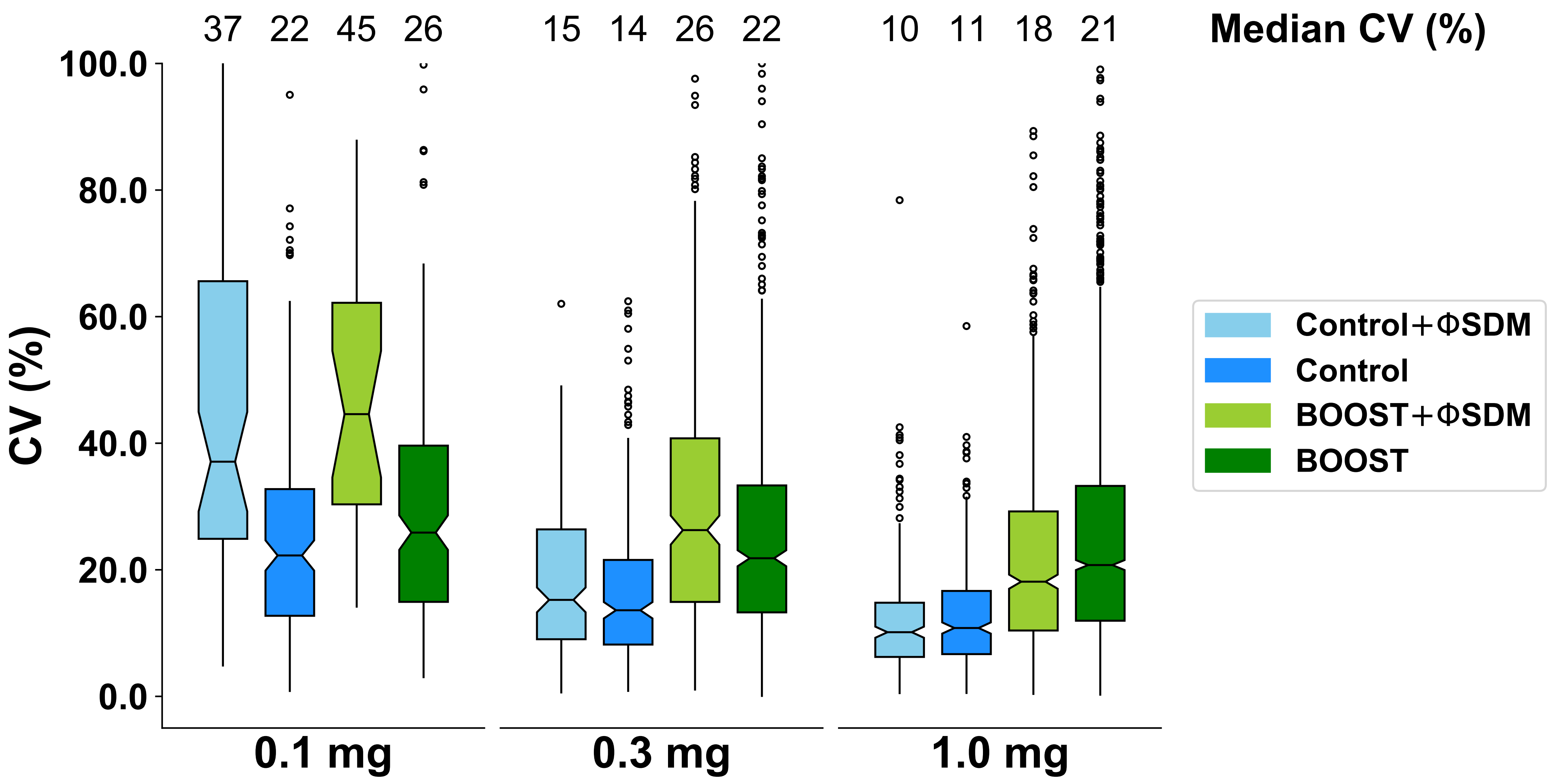

### depth_nans.pdf

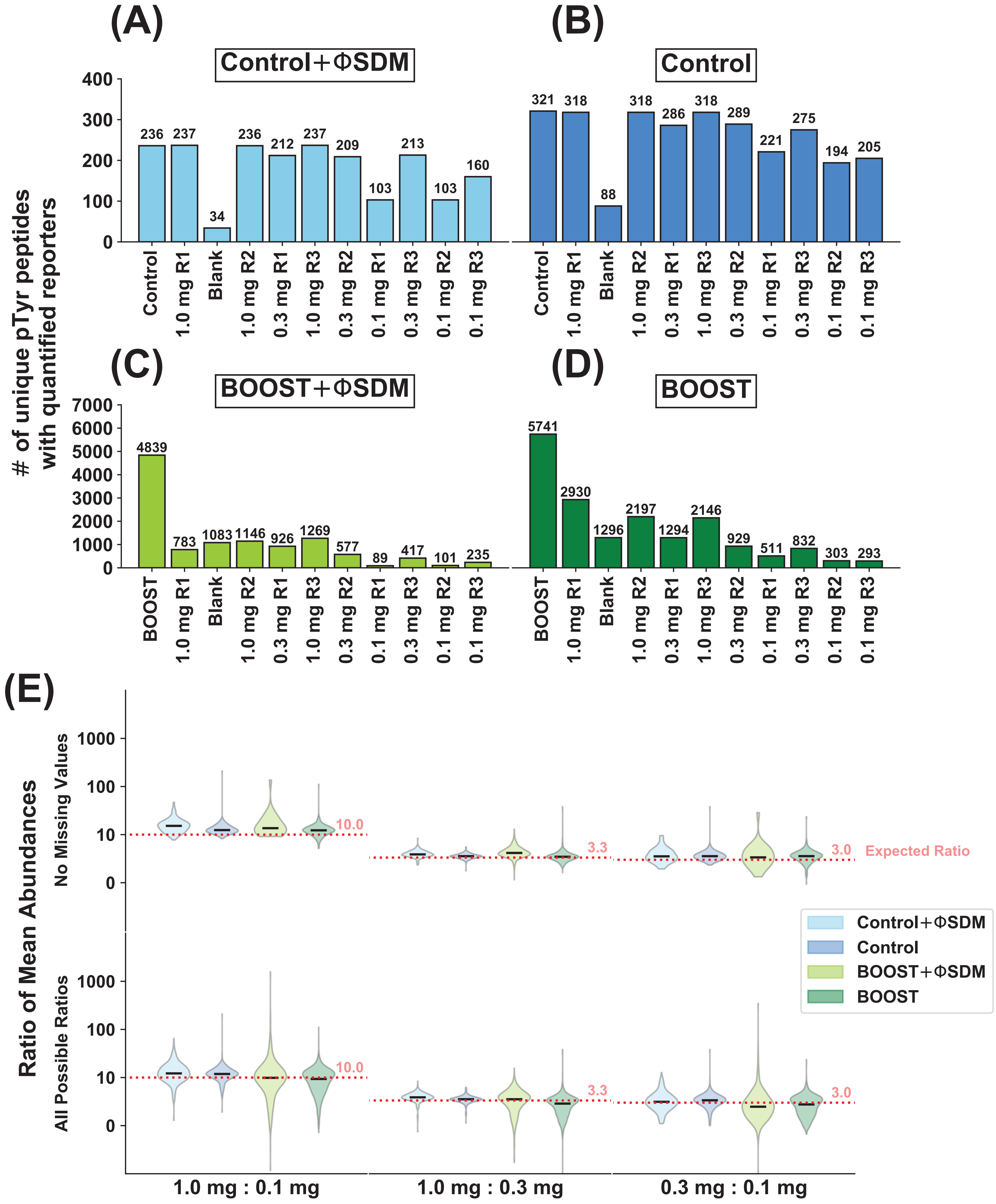

### human_mouse_overlap.pdf

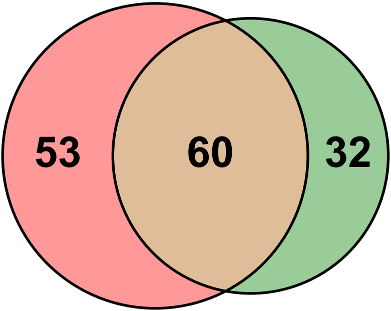

### intensity_plots.pdf

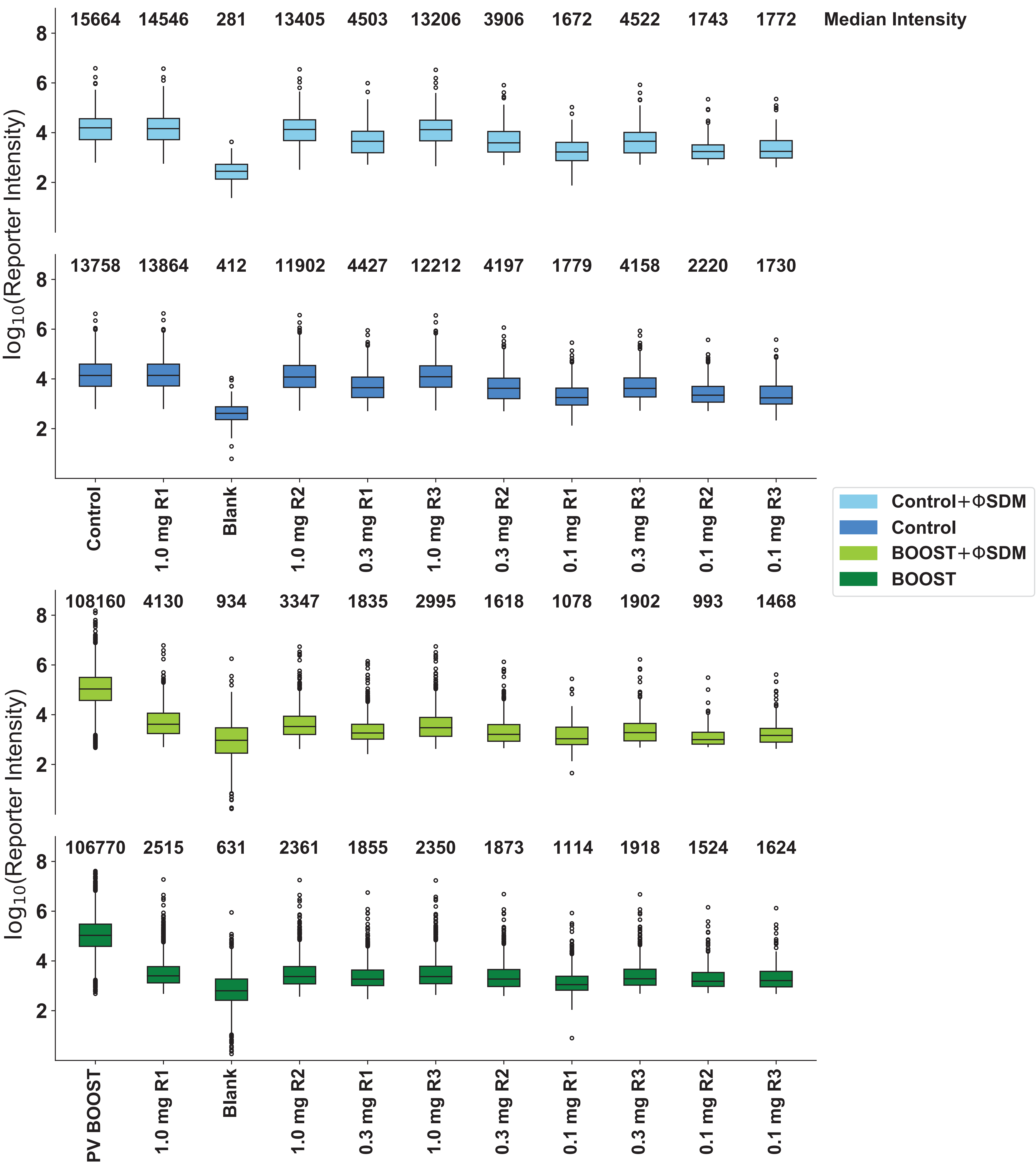

### localisation_prob_histogram.pdf

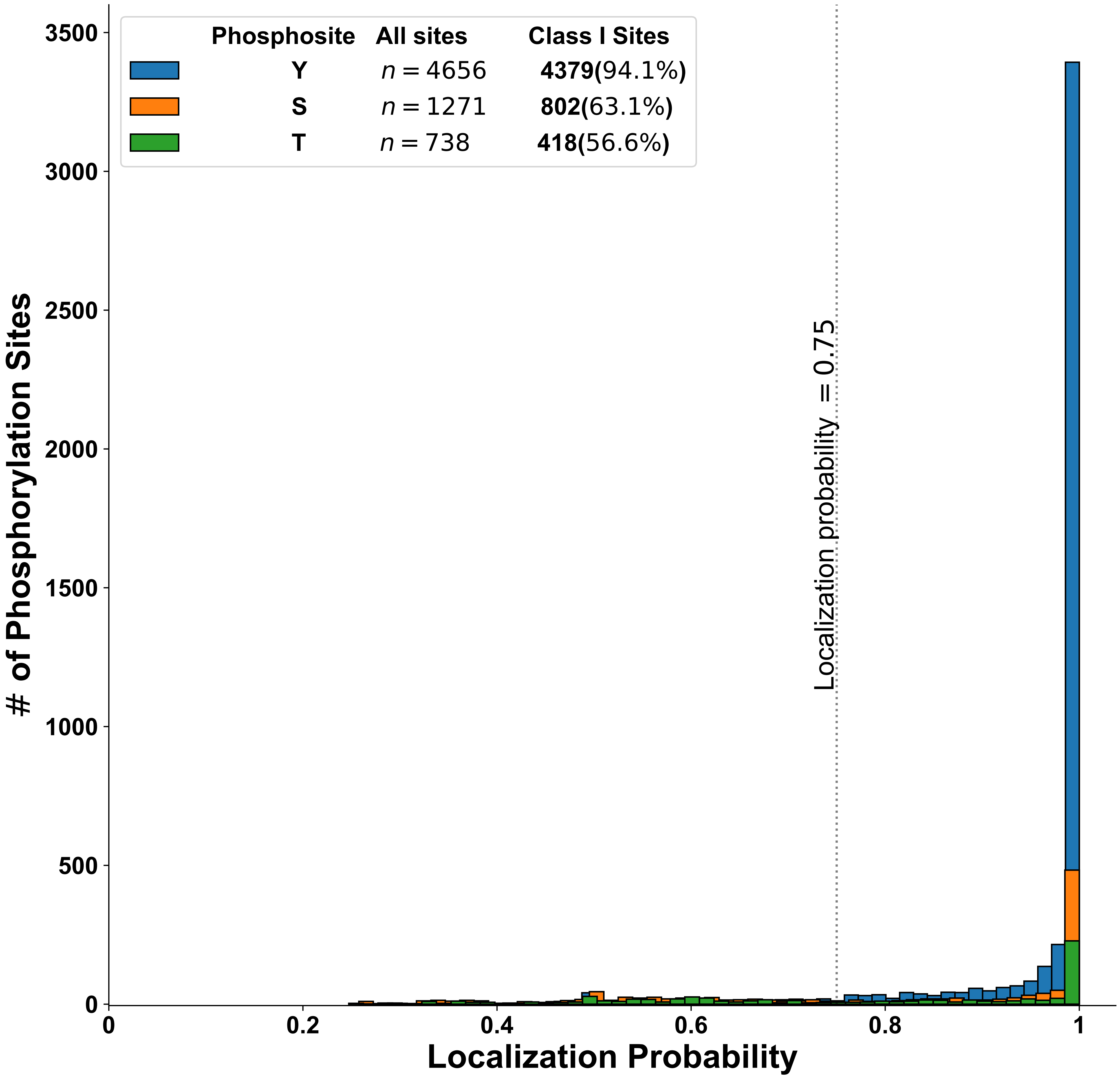

### locprob.pdf

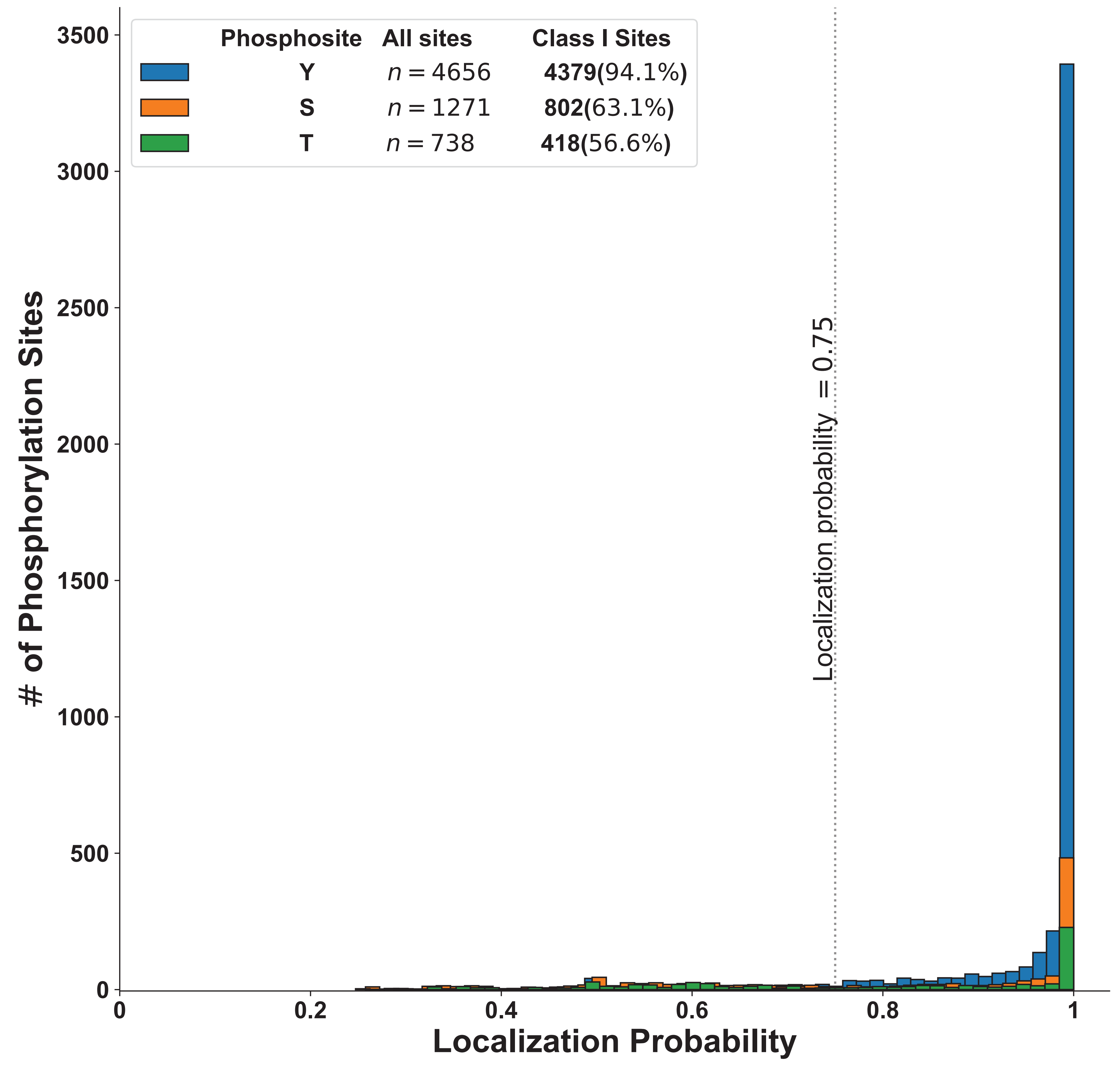

### mg01_repmean.pdf

#### Replicates vs the Mean for 0.1 mg Protein Input

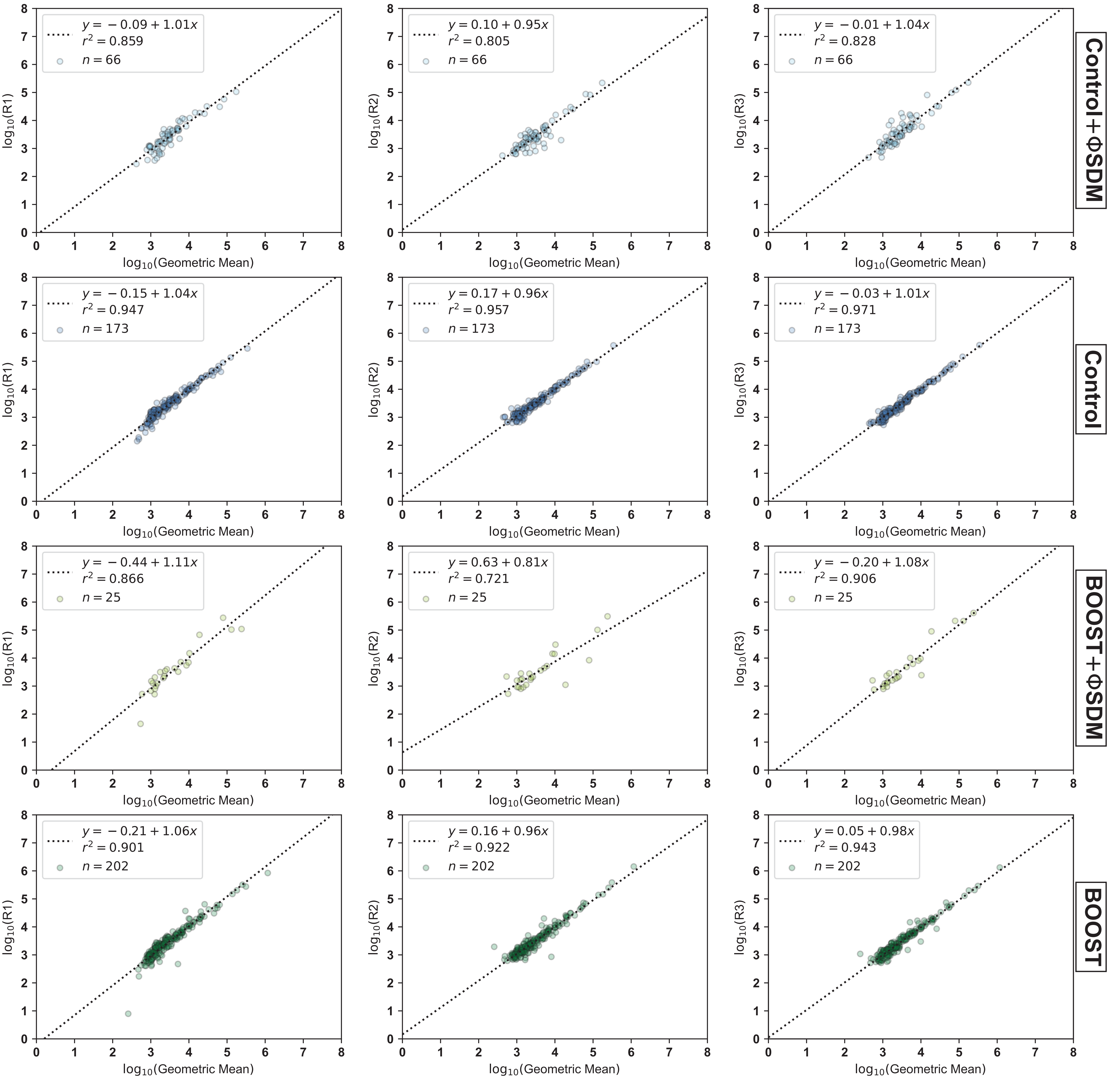

### mg1_repmean.pdf

#### Replicates vs the Mean for 1.0 mg Protein Input

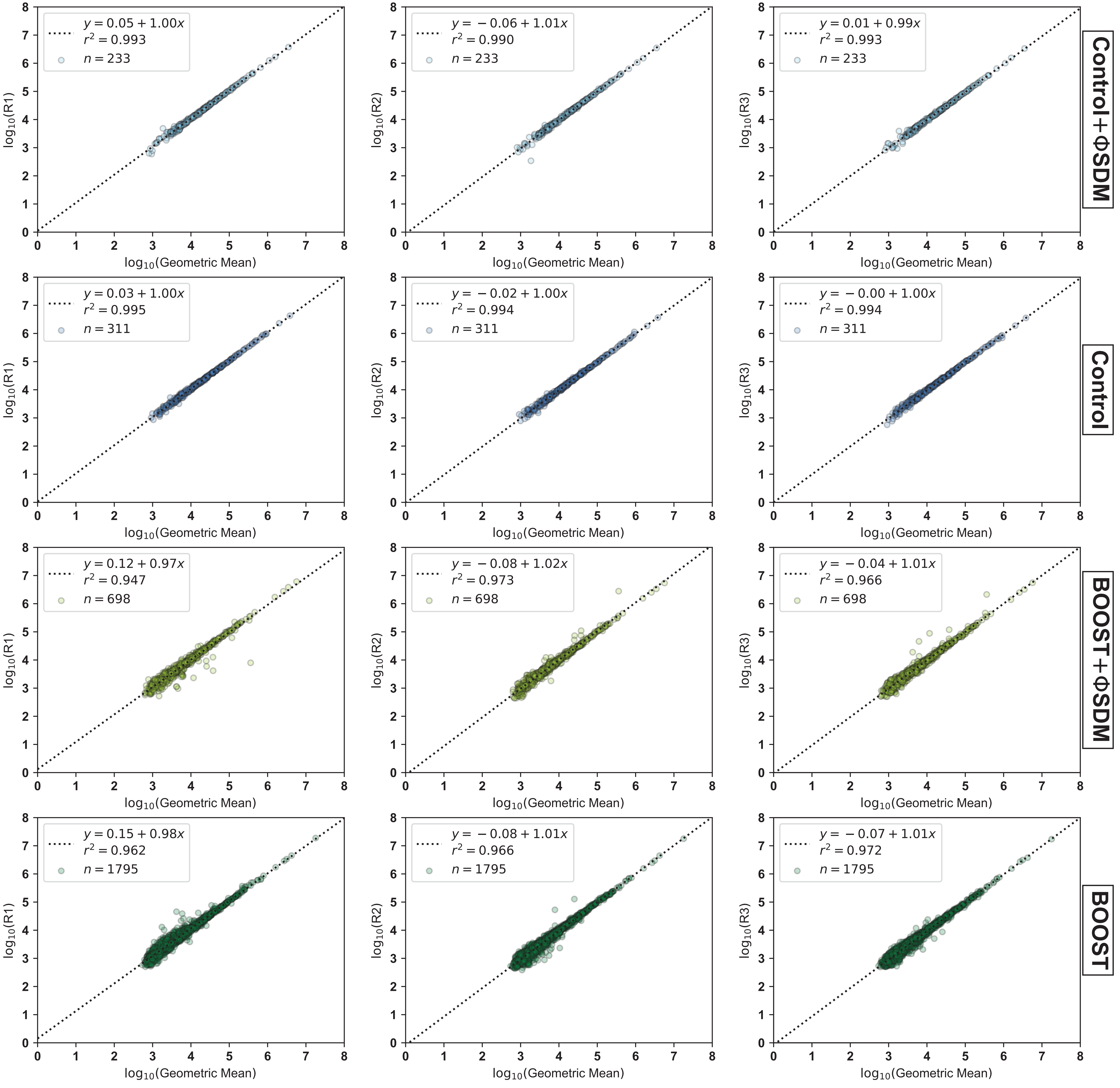

### mg03_repmean.pdf

#### Replicates vs the Mean for 0.3 mg Protein Input

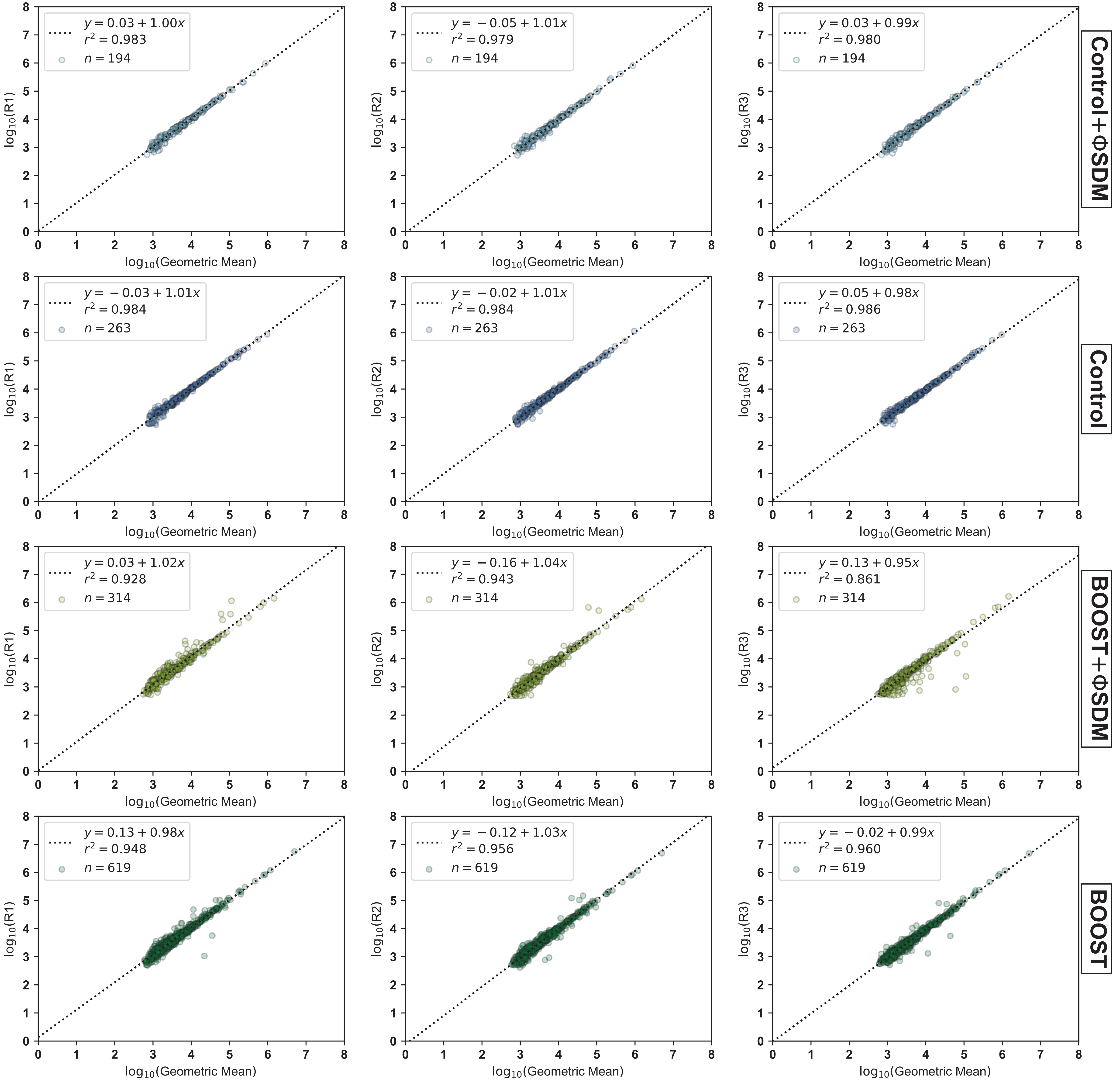

### mouse-1.jpg

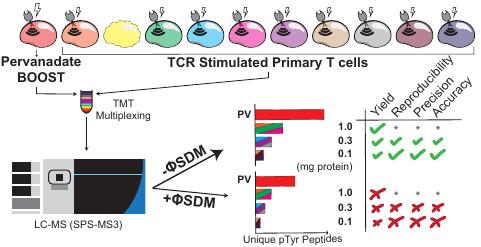

### mouse_turbotmt_gtoc.pdf

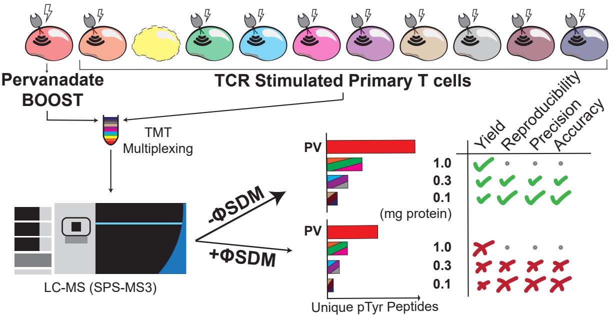

### nancounts.pdf

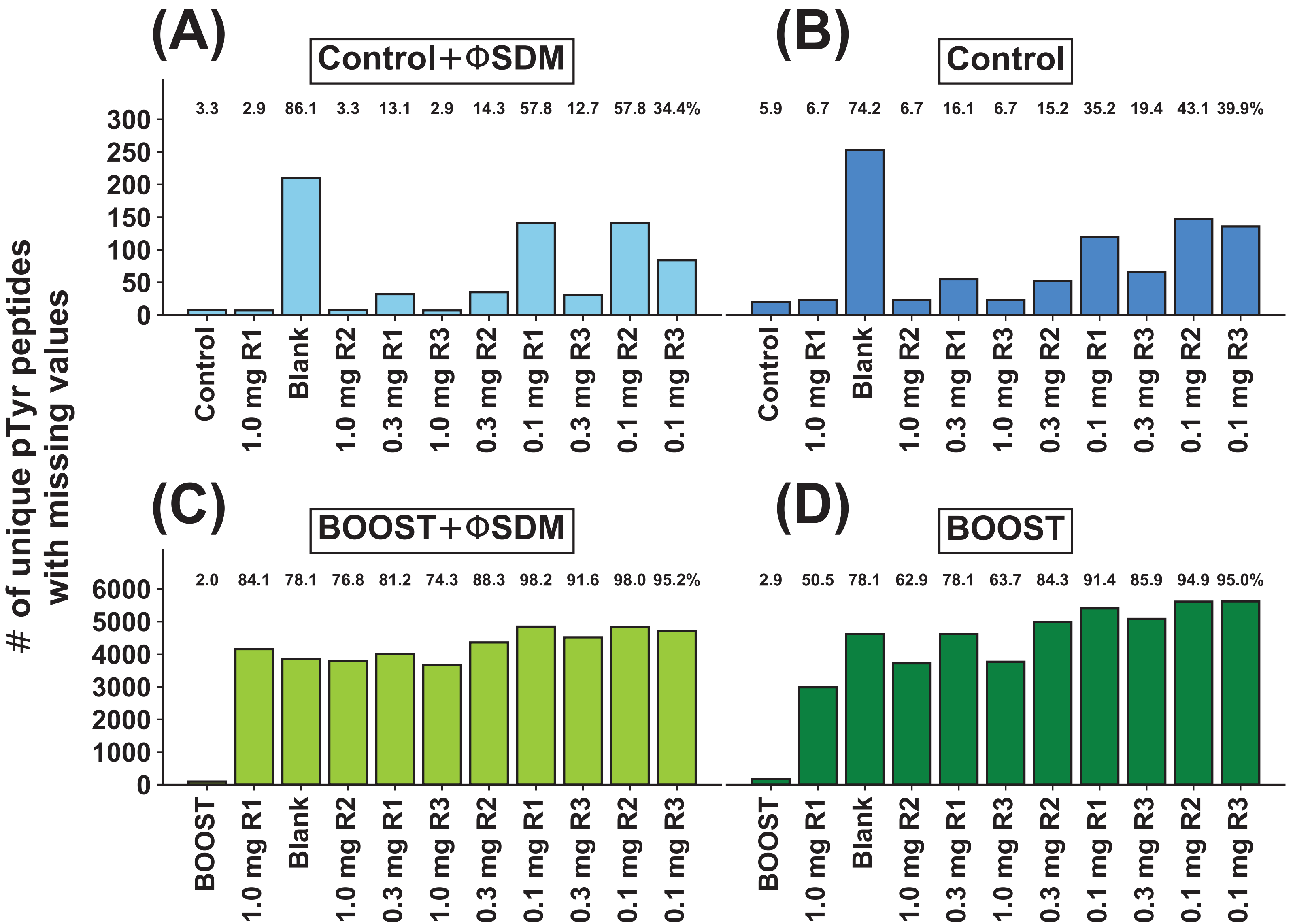

### overlap_q_volcano_nosdm.pdf

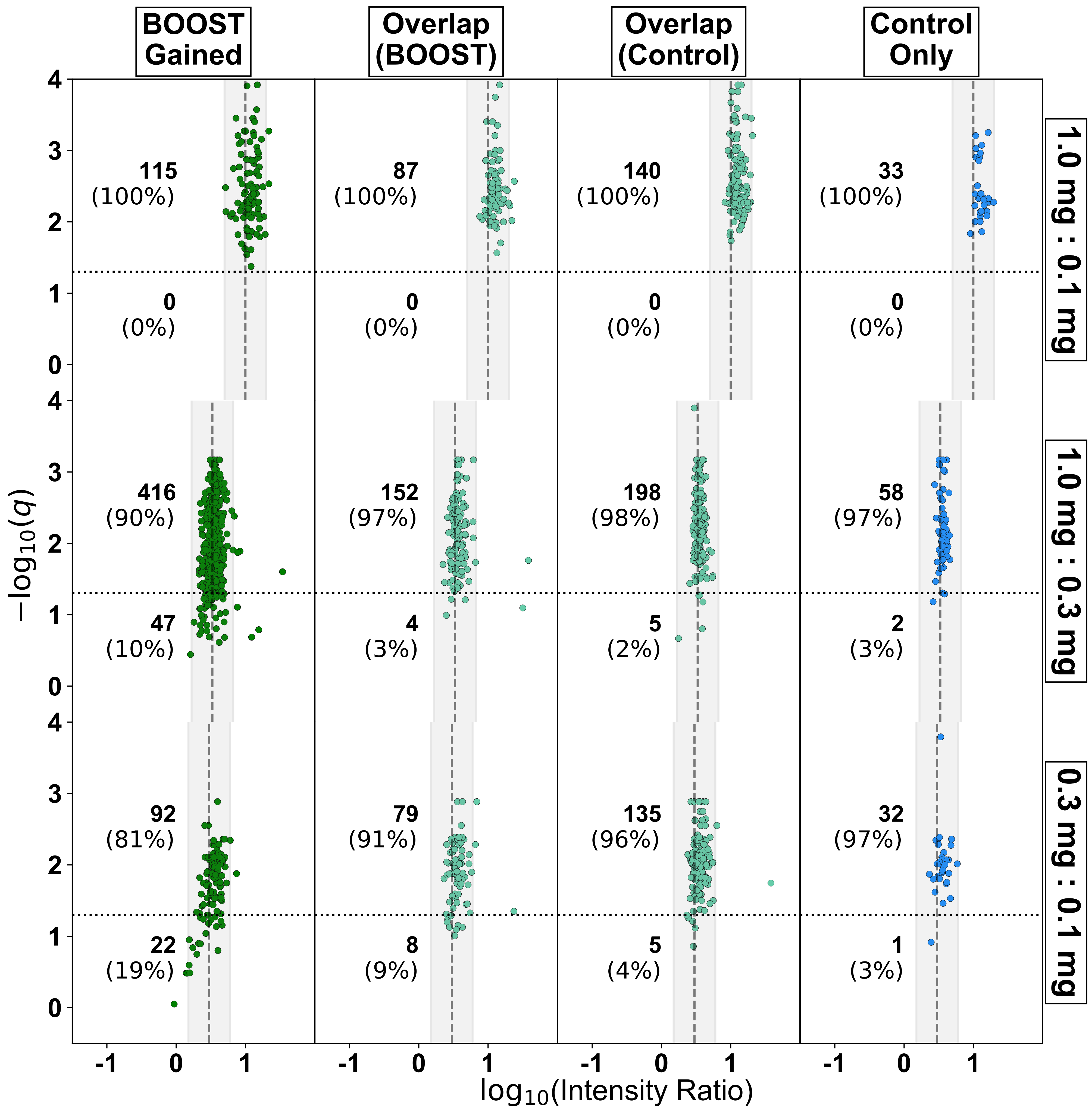

### overlap_q_volcano_sdm.pdf

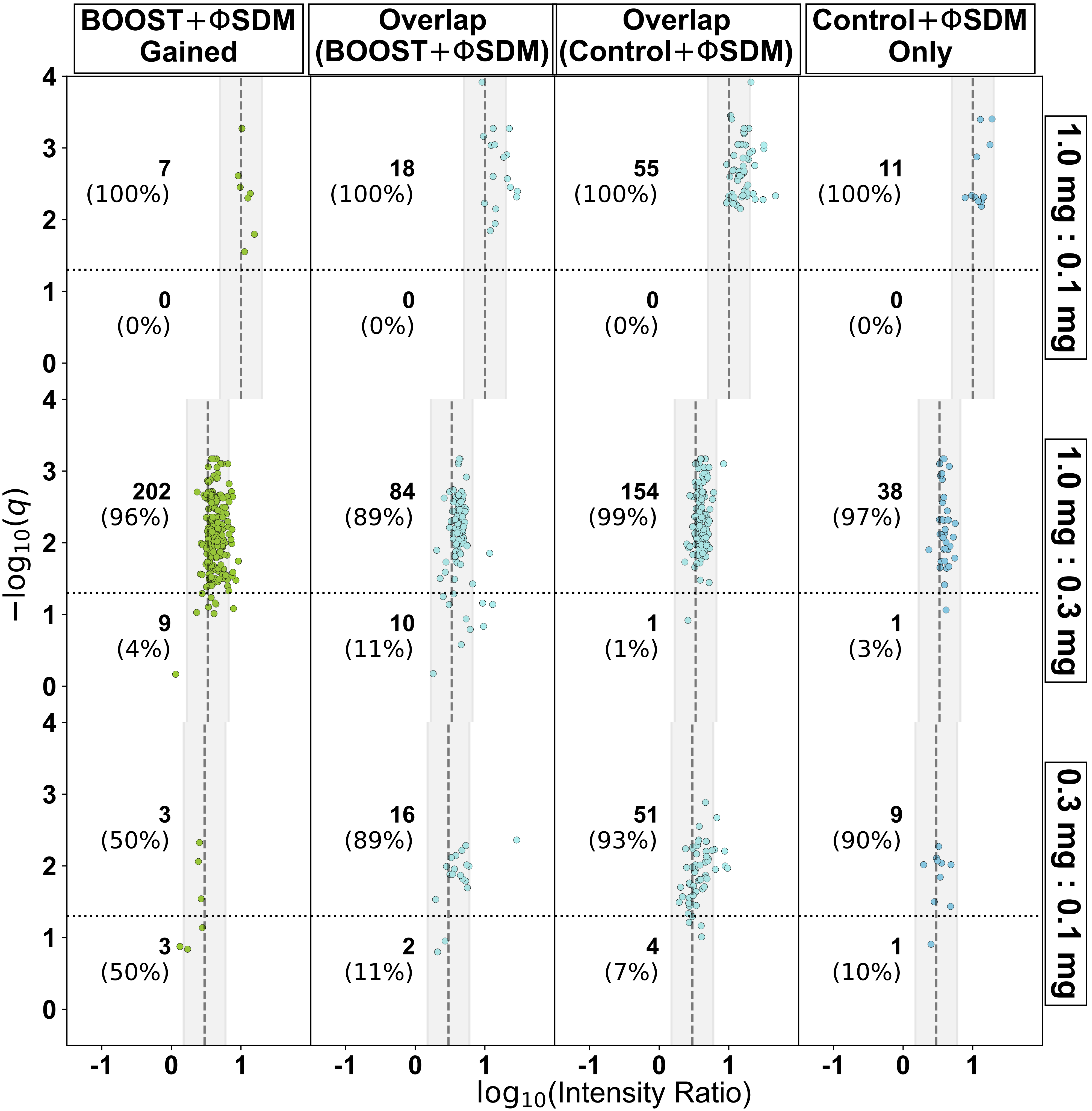

### q_volcano.pdf

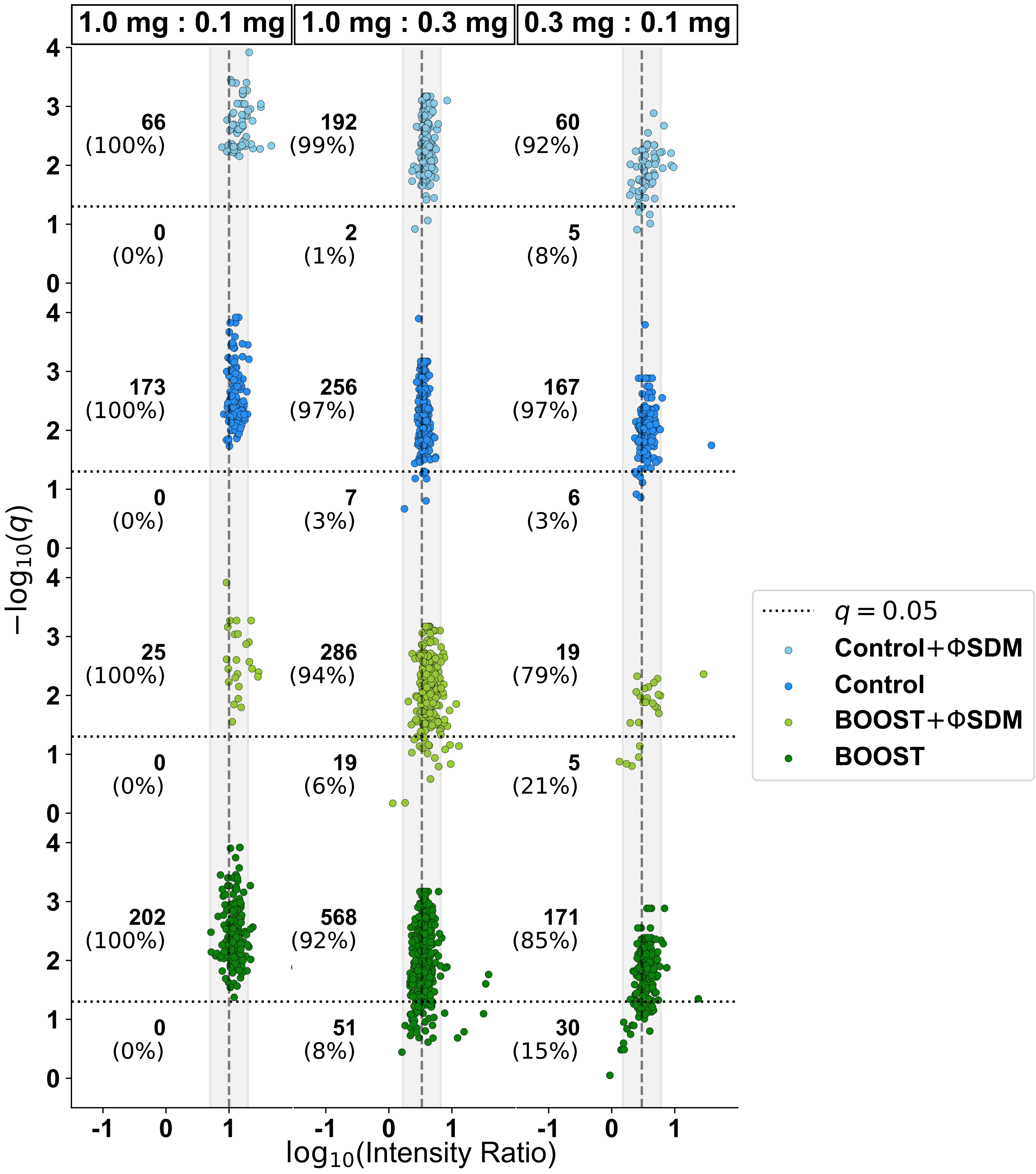

### ratio_suppression.pdf

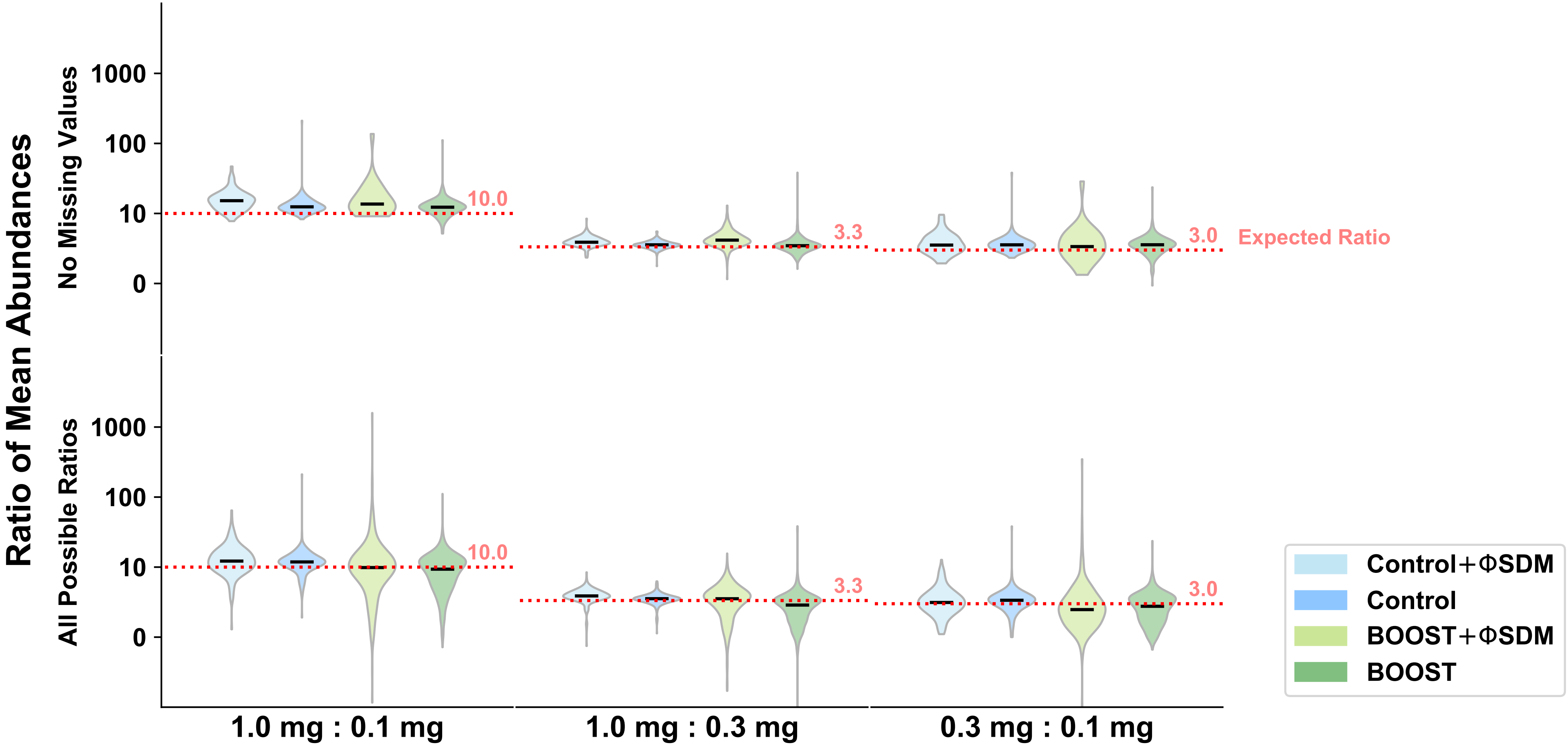

### report_intensity_boxplot.pdf

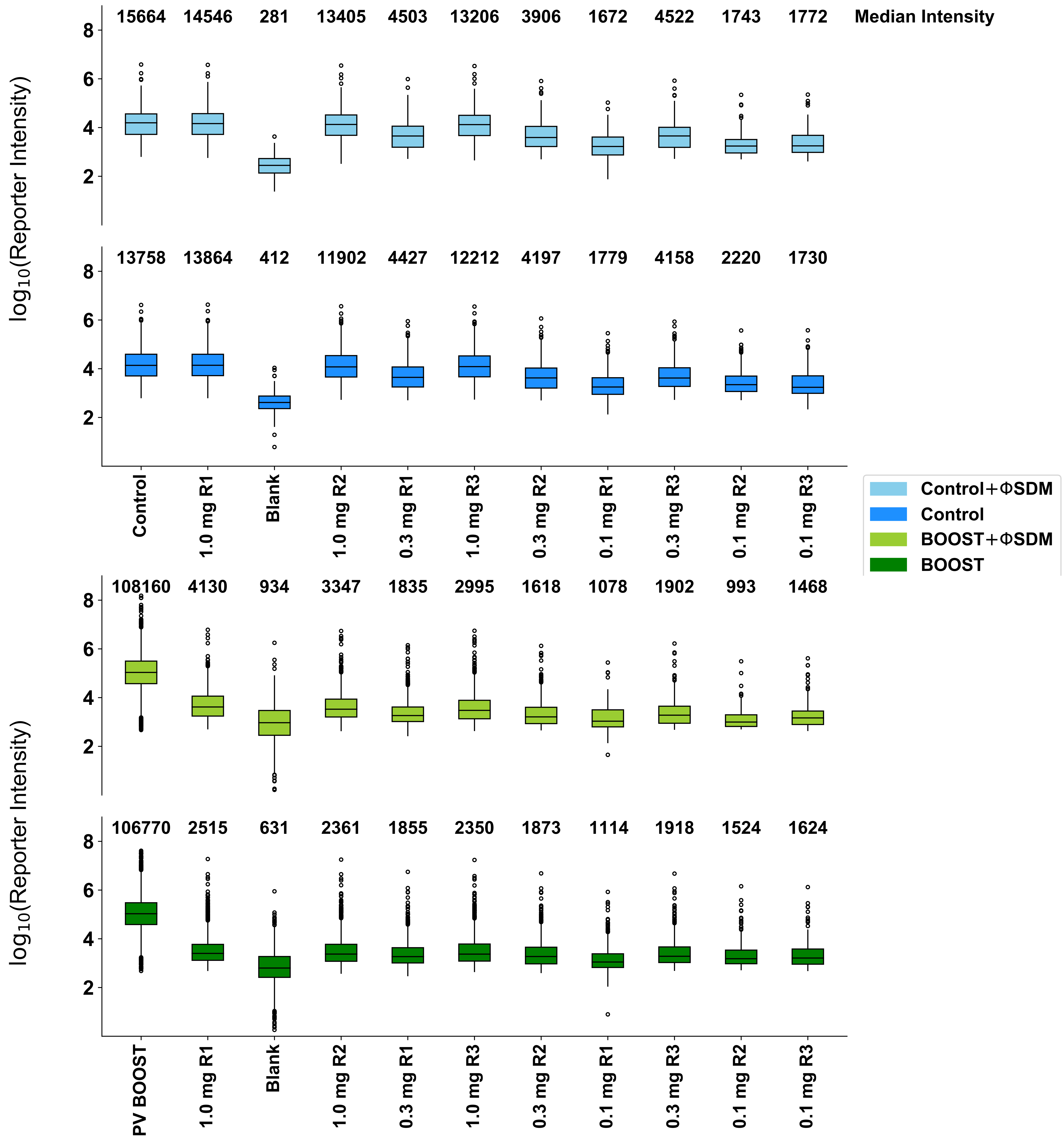

### reporter_barchart.pdf

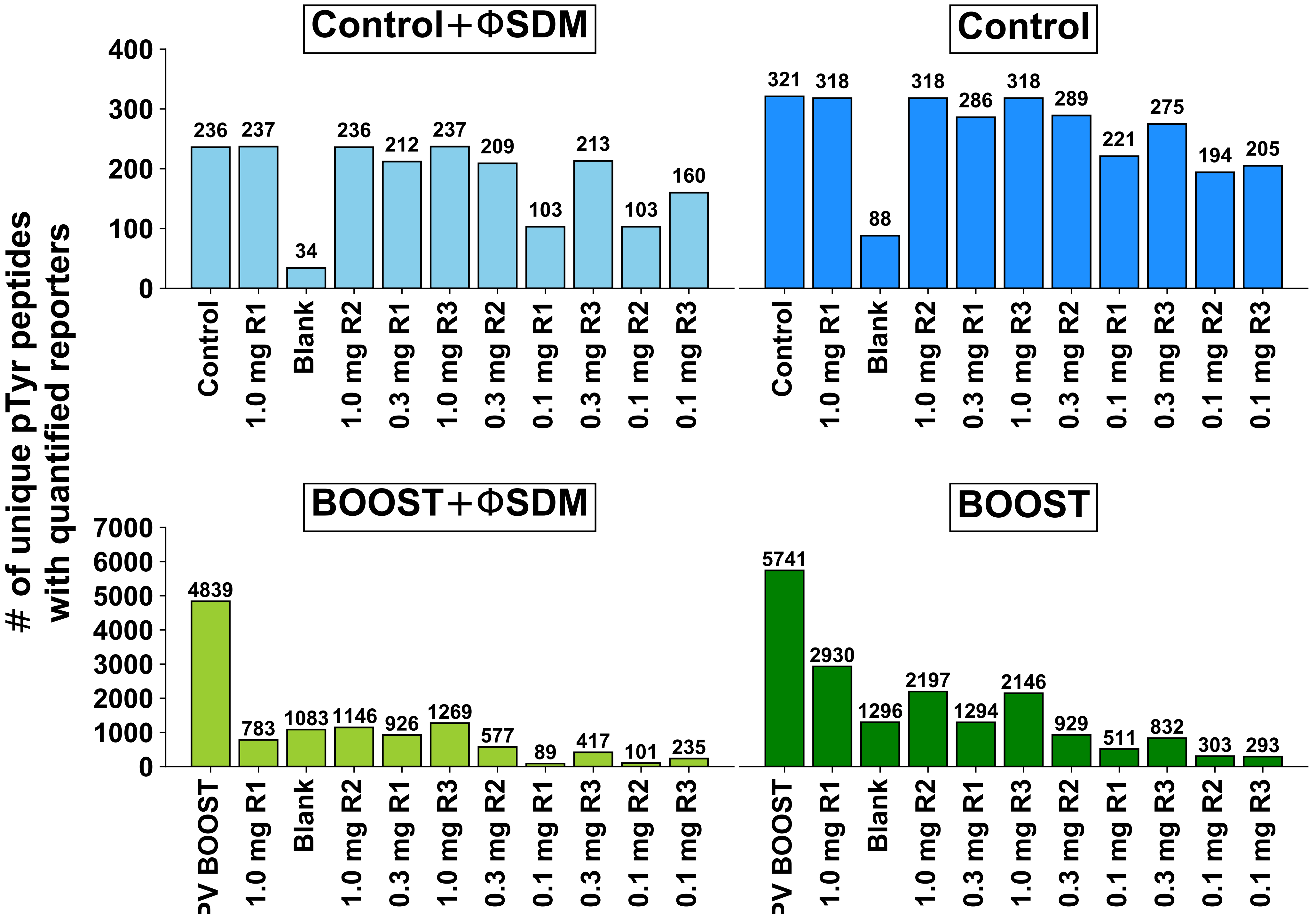

### reporter_nans_barchart.pdf

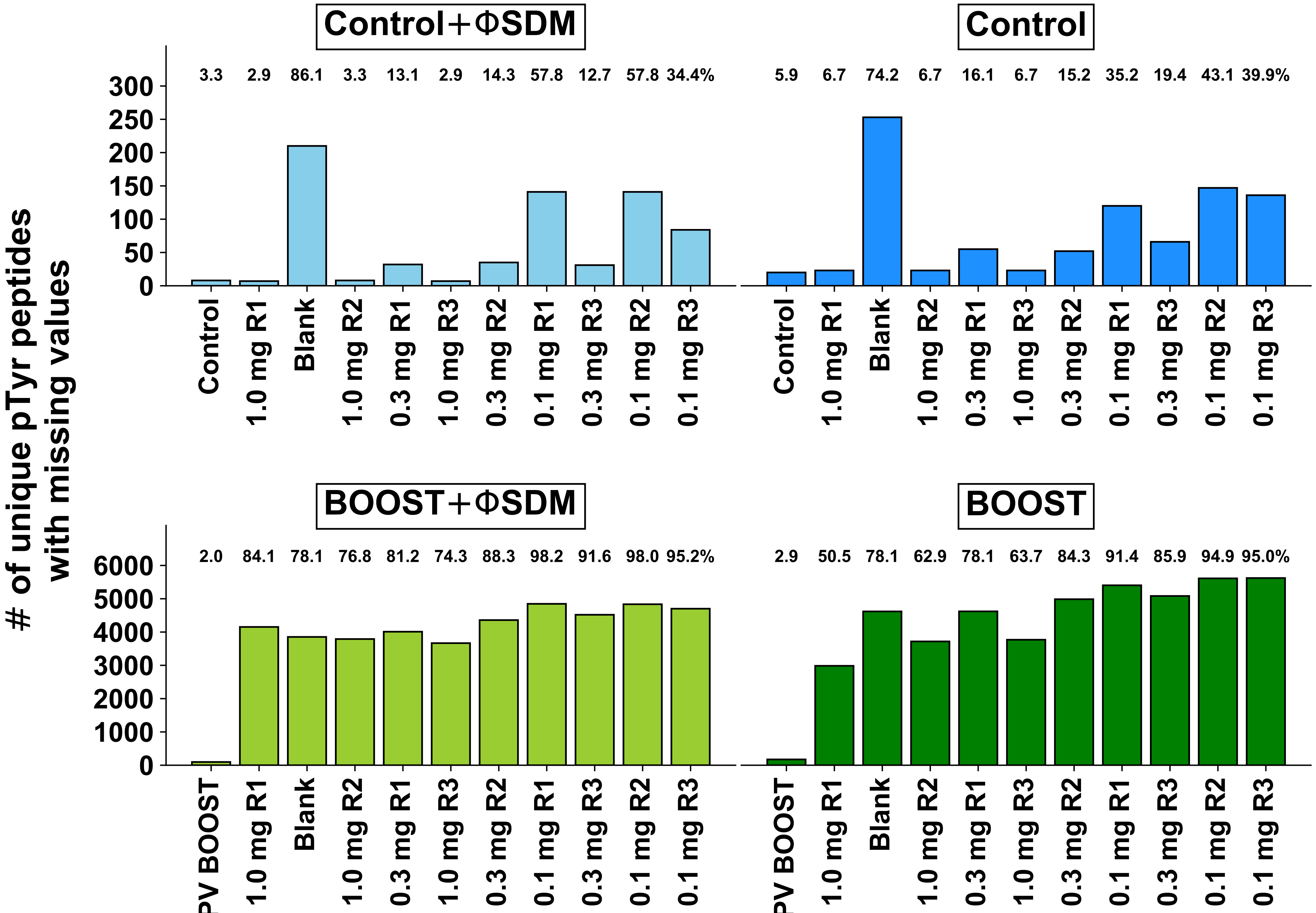

### reps_vs_mean_01mg.pdf

#### Replicates vs the Mean for 0.1 mg Protein Input

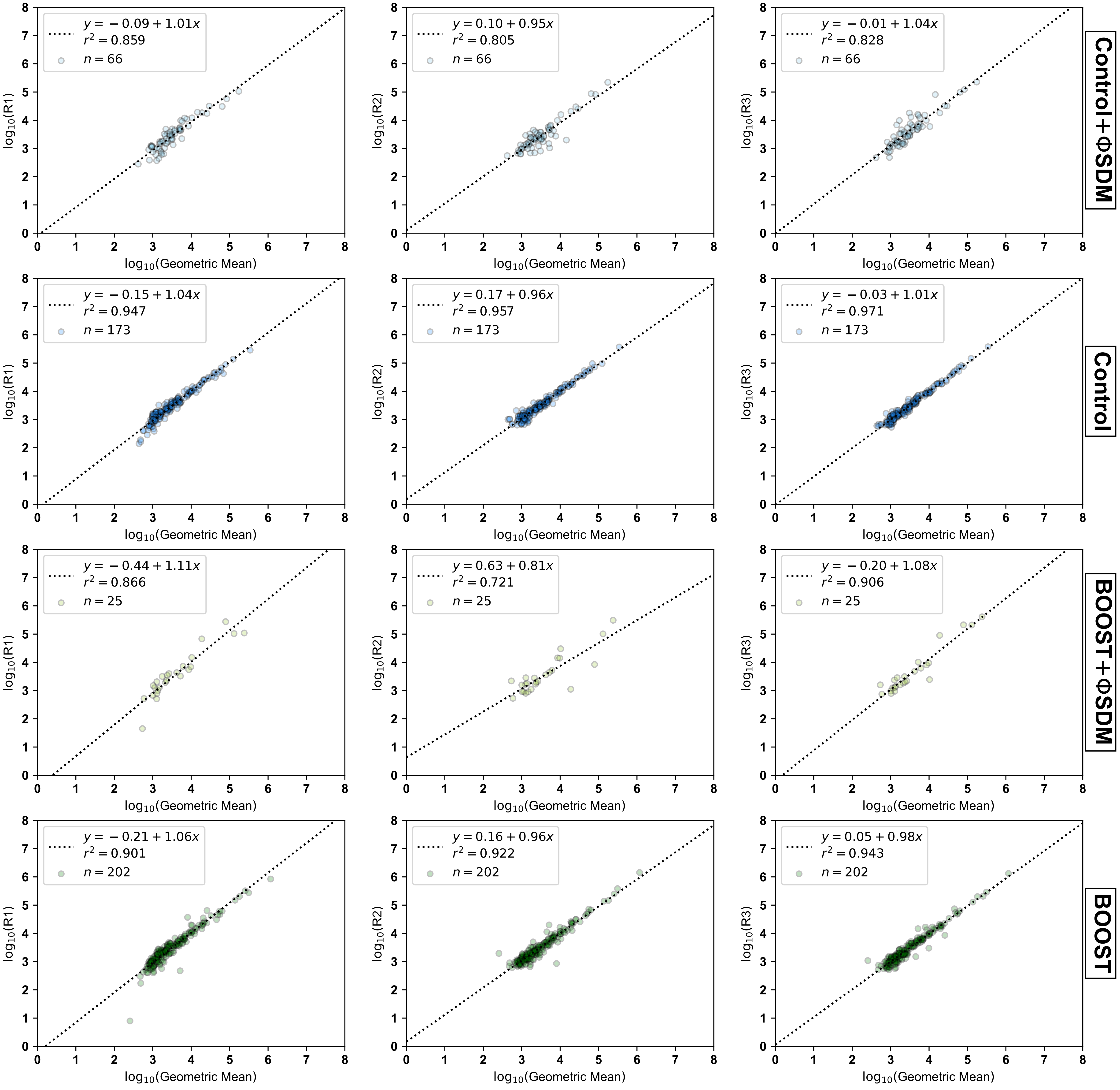

### reps_vs_mean_1mg.pdf

##### Replicates vs the Mean for 1.0 mg Protein Input

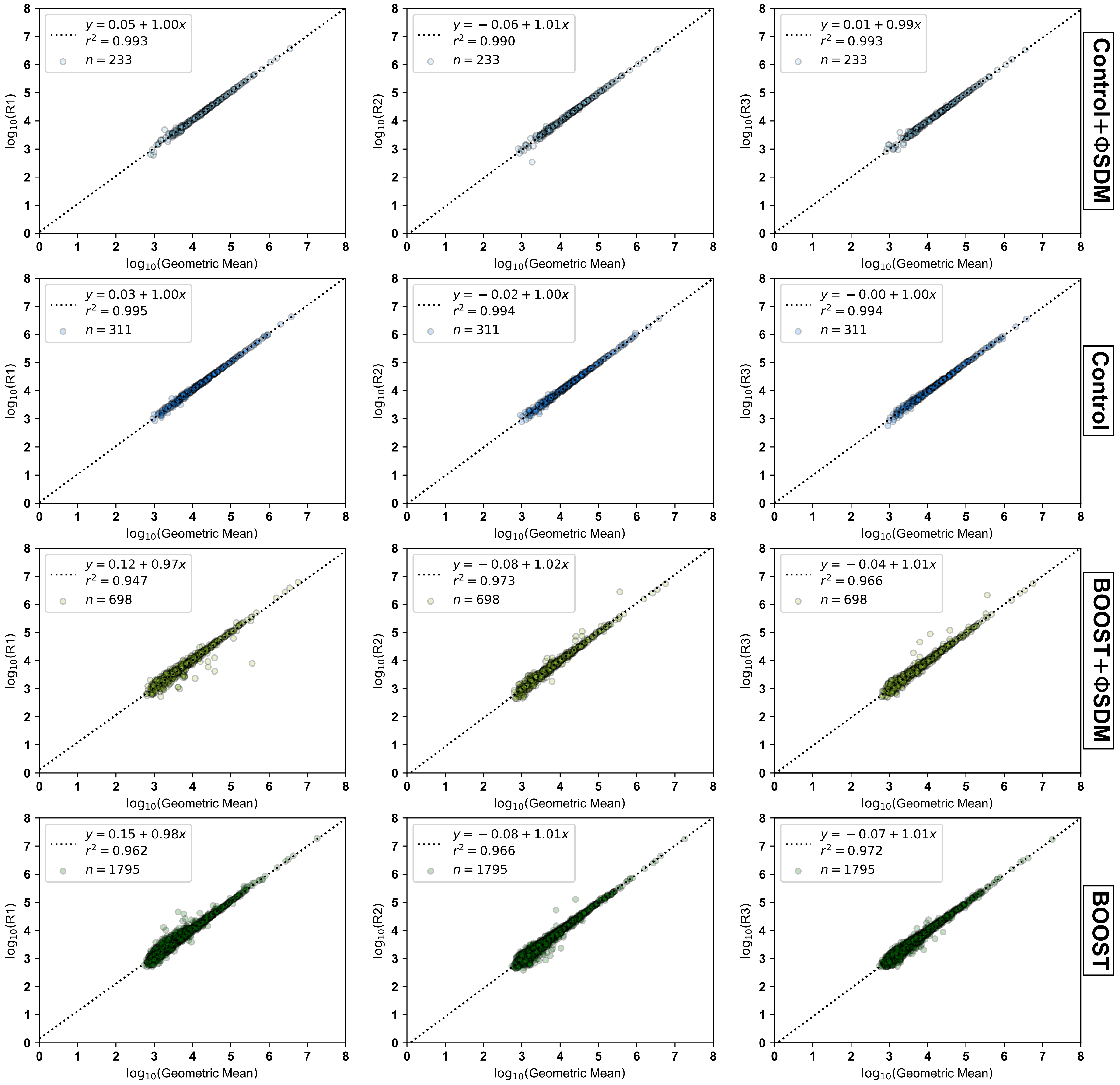

### reps_vs_mean_03mg.pdf

##### Replicates vs the Mean for 0.3 mg Protein Input

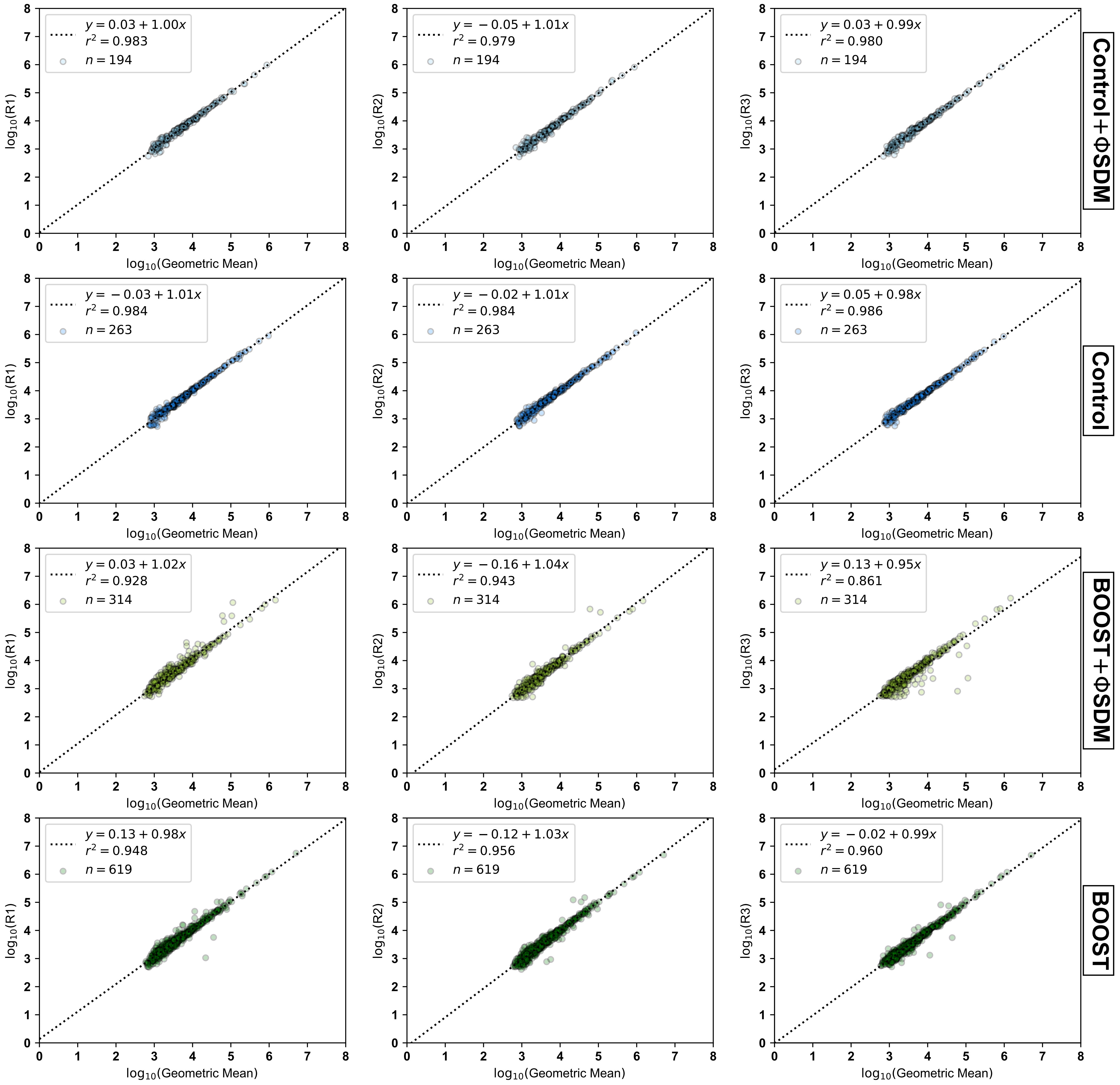

### scans.pdf

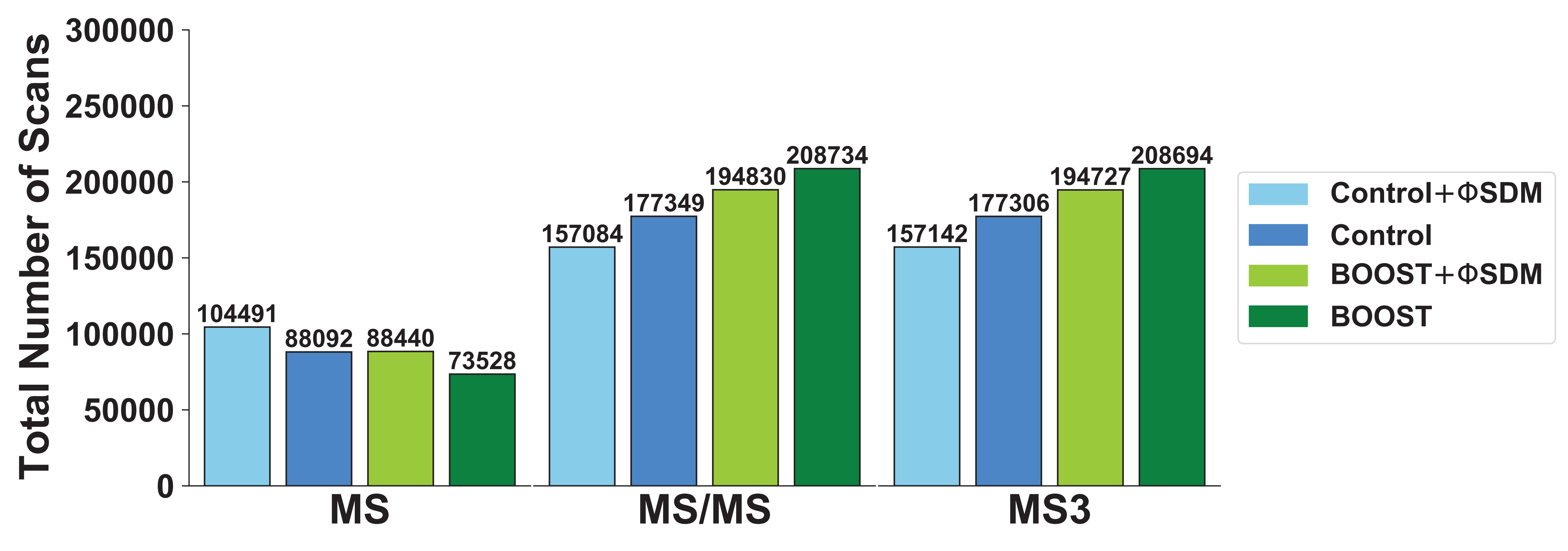

### set_overlap.pdf

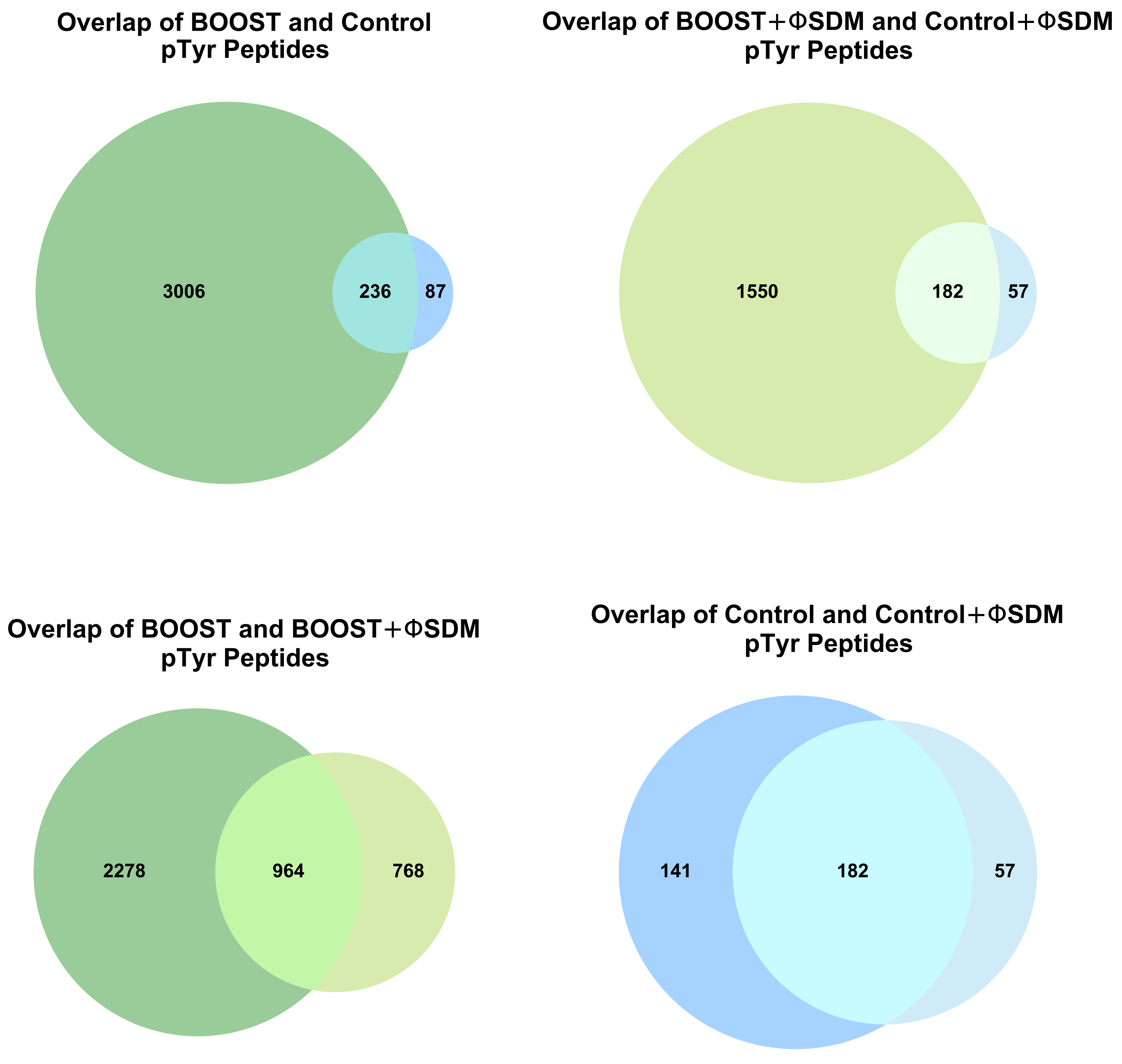

### set_overlap_2.pdf

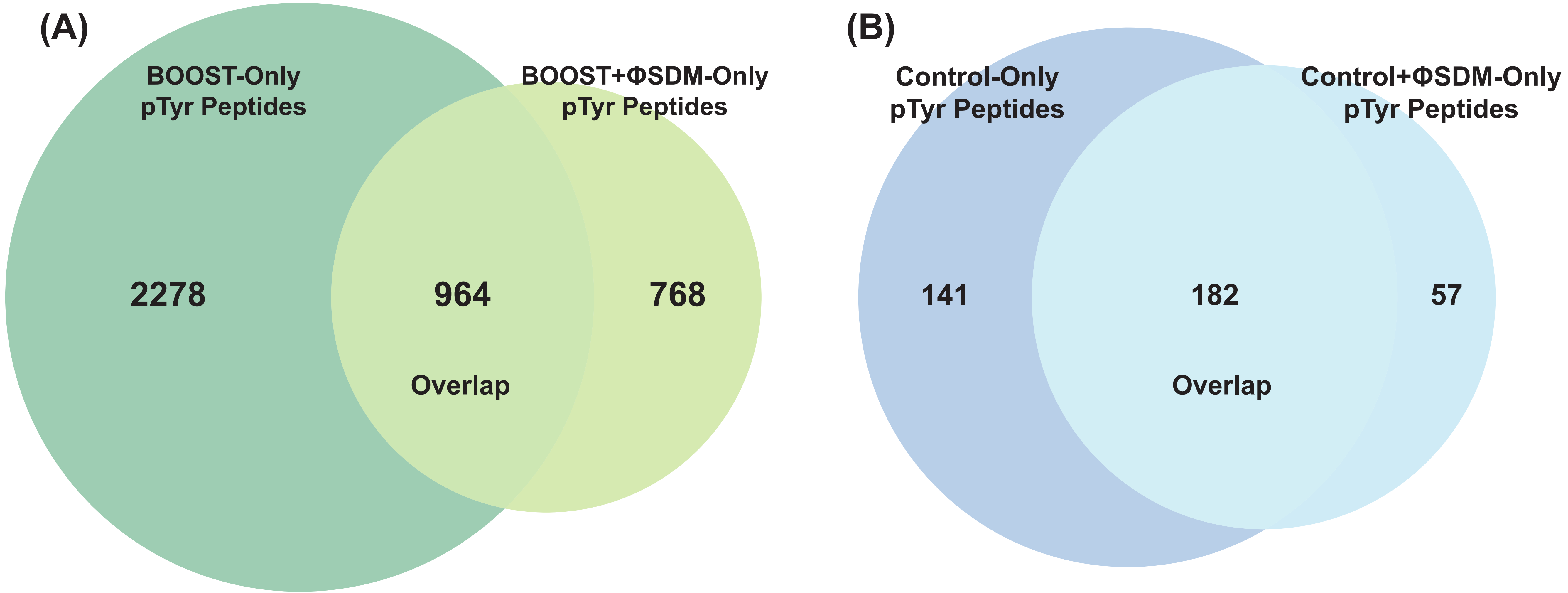
