## Supporting Information for "Mouse primary T cell phosphotyrosine proteomics enabled by BOOST"

### Table Of Contents


| | |
|---|---|
| **Supporting Table 1:** | (.XLSX) All unique peptides observed in the BOOST experiment without ΦSDM |
| **Supporting Table 2:** | (.XLSX) All unique peptides observed in the BOOST and 1.0 mg Control experiments without ΦSDM |
| **Supporting Table 3:** | (.XLSX) All unique peptides observed in the 1.0 mg Control experiment without ΦSDM |
| **Supporting Table 4:** | (.XLSX) All unique peptides observed in the BOOST experiment with ΦSDM |
| **Supporting Table 5:** | (.XLSX) All unique peptides observed in the BOOST and 1.0 mg Control experiments with ΦSDM |
| **Supporting Table 6:** | (.XLSX) All unique peptides observed in the 1.0 mg Control experiment with ΦSDM |
| **Supporting Folder 1** | (.ZIP) All tables output from MaxQuant |
| **Supporting Folder 2** | (.ZIP) All Python3 code used for data analysis |
| **Supporting Figure 1:** | Histogram of all PSMs containing a phosphorylated amino acid in all conditions binned by localization probability |
| **Supporting Figure 2:** | Missing value bar charts for each experiment |
| **Supporting Figure 3:** | $\log_{10}$ transformed reporter intensity box-and-whisker plots |
| **Supporting Figure 4:** | Comparison of replicate intensities to the mean of each pTyr peptide PSM in the 1.0 mg condition. |
| **Supporting Figure 5:** | Comparison of replicate intensities to the mean of each pTyr peptide PSM in the 0.3 mg condition. |
| **Supporting Figure 6:** | Comparison of replicate intensities to the mean of each pTyr peptide PSM in the 0.1 mg condition. |
| **Supporting Figure 7:** | Pairwise replicate comparisons of unique PSMs identified in BOOST when ΦSDM is disabled |
| **Supporting Figure 8:** | Pairwise replicate comparisons of unique PSMs identified in BOOST when ΦSDM is enabled |
| **Supporting Figure 9:** | Pairwise replicate comparisons of unique PSMs identified in 1.0 mg Control when ΦSDM is disabled |
| **Supporting Figure 10:** | Pairwise replicate comparisons of unique peptides identified in 1.0 mg Control when ΦSDM is enabled |

| | |
|---|---|
| **Supporting Figure 11:** | Venn Diagram and Volcano Plots for BOOST and 1.0 mg Control when ΦSDM is disabled |
| **Supporting Figure 12:** | Venn Diagram and Volcano Plots for BOOST and 1.0 mg Control when ΦSDM is enabled |
| **Supporting Figure 13:** | Venn Diagrams for BOOST conditions (with and without ΦSDM) and 1.0 mg Control (with and without ΦSDM) |
| **Supporting Figure 14:** | Cumulative distributions of unique pTyr peptides from BOOST experiments (with and without ΦSDM) |
| **Supporting Figure 15:** | Overlap between Mouse-BOOST and Jurkat-BOOST experiments |

Supporting Table 1: All unique peptide PSMs observed exclusively in the BOOST experiment with ΦSDM disabled and reporter ions quantified in at least one experimental channel. Included are MaxQuant outputs such as the gene name(s), protein name(s), modified sequence, modified sequence with phosphorylation site localization probabilities, sequence windows, and corrected TMT reporter ion intensities, as well as calculations used in the manuscript (mean intensity values, standard deviations, CV percentages, $p$-values, $q$-values, BOOST factors) and WikiPathways[1] Annotations for each unique peptide.

Supporting Table 2: All unique peptide PSMs observed in both the BOOST experiment and the 1.0 mg Control experiment with ΦSDM disabled and reporter ions quantified in at least one experimental channel. Included are MaxQuant outputs such as the gene name(s), protein name(s), modified sequence, modified sequence with phosphorylation site localization probabilities, sequence windows, and corrected TMT reporter ion intensities, as well as calculations used in the manuscript (mean intensity values, standard deviations, CV percentages, $p$-values, $q$-values, BOOST factors) and WikiPathways[1] Annotations for each unique peptide.

Supporting Table 3: All unique peptide PSMs observed exclusively in the 1.0 mg Control experiment with ΦSDM disabled and reporter ions quantified in at least one experimental channel. Included are MaxQuant outputs such as the gene name(s), protein name(s), modified sequence, modified sequence with phosphorylation site localization probabilities, sequence windows, and corrected TMT reporter ion intensities, as well as calculations used in the manuscript (mean intensity values, standard deviations, CV percentages, $p$-values, $q$-values) and WikiPathways[1] Annotations for each unique peptide.

Supporting Table 4: All unique peptide PSMs observed exclusively in the BOOST experiment with ΦSDM enabled and reporter ions quantified in at least one experimental channel. Included are MaxQuant outputs such as the gene name(s), protein name(s), modified sequence, modified sequence with phosphorylation site localization probabilities, sequence windows, and corrected TMT reporter ion intensities, as well as calculations used in the manuscript (mean intensity values, standard deviations, CV percentages, $p$-values, $q$-values, BOOST factors) and WikiPathways[1] Annotations for each unique peptide.

Supporting Table 5: All unique peptide PSMs observed in both the BOOST experiment and the 1.0 mg Control experiment with ΦSDM enabled and reporter ions quantified in at least one experimental channel. Included are MaxQuant outputs such as the gene name(s), protein name(s), modified sequence, modified sequence with phosphorylation site localization probabilities, sequence windows, and corrected TMT reporter ion intensities, as well as calculations used in the manuscript (mean intensity values, standard deviations, CV percentages, *p*-values, *q*-values, BOOST factors) and WikiPathways[1] Annotations for each unique peptide.

Supporting Table 6: All unique peptide PSMs observed exclusively in the 1.0 mg Control experiment with ΦSDM enabled and reporter ions quantified in at least one experimental channel. Included are MaxQuant outputs such as the gene name(s), protein name(s), modified sequence, modified sequence with phosphorylation site localization probabilities, sequence windows, and corrected TMT reporter ion intensities, as well as PhosphoSitePlus® site annotations, calculations used in the manuscript (mean intensity values, standard deviations, CV percentages, *p*-values, *q*-values) and WikiPathways[1] Annotations for each unique peptide.

Supporting Folder 1: All tables generated by MaxQuant as text files. These include “summary.txt” (a summary of parameters, information, .raw files, and statistics used for peak detection), “evidence.txt” (all information about unique peptides quantified from .raw files), “peptides.txt” (information about the peptides identified from .raw files), “modificationSpecificPeptides.txt” (information about posttranlational modifications to the peptides), “Oxidation (M)Sites.txt” (information about oxidized peptides), “Phospho (STY)Sites.txt” (information about phosphorylated peptides), “proteinGroups.txt” (information about estimated protein abundance from the .raw files), “allPeptides.txt” (all information for each unique peptide identified in each .raw file), “msScans.txt” (information about the scans observed on the mass spectrometer), “mzRange.txt”, “msmsScans.txt” (information about the MS/MS scans for each .raw file), and “msms.txt” (information about the MS/MS spectra for each peptide identified in each .raw file).

Supporting Folder 2: All Python3 code used to analyze the MaxQuant output files and databases referenced. These include “data_analysis.py” (script used to generate plots), “helpers/” (Python3 files used to assist in data analysis), “database/” (all external databases used in analysis), and “maxquant_results” (the “evidence.txt” and “Phospho (STY)Sites.txt” files from Supporting Folder 2), as well as the output folders “figures/” (all figures generated by data_analysis.py) and “curated_results/” (all .txt output files from Python3 analysis, which are aggregated and formatted in Supporting Tables 1-6).

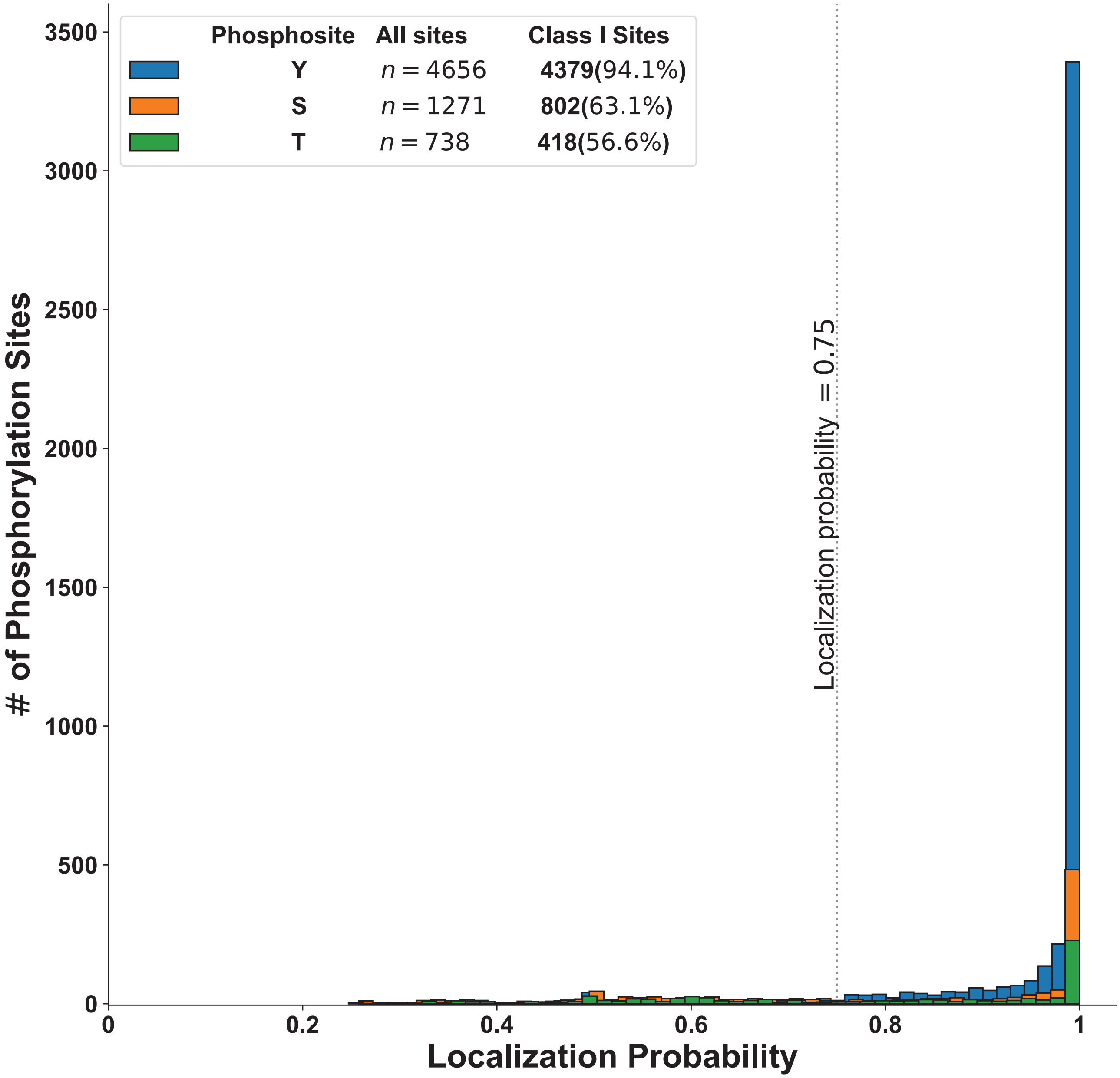


Supporting Figure 1: A histogram with depicting all PSMs from all experiments containing at least one phosphorylated serine (S), threonine (T), or tyrosine (Y) amino acid as a function of localization probability ($n_{\text{bins}} = 75$). The total number and number of Class I (localization probability $> 0.75$) phosphorylation sites for each amino acid are noted in the Figure Legend.

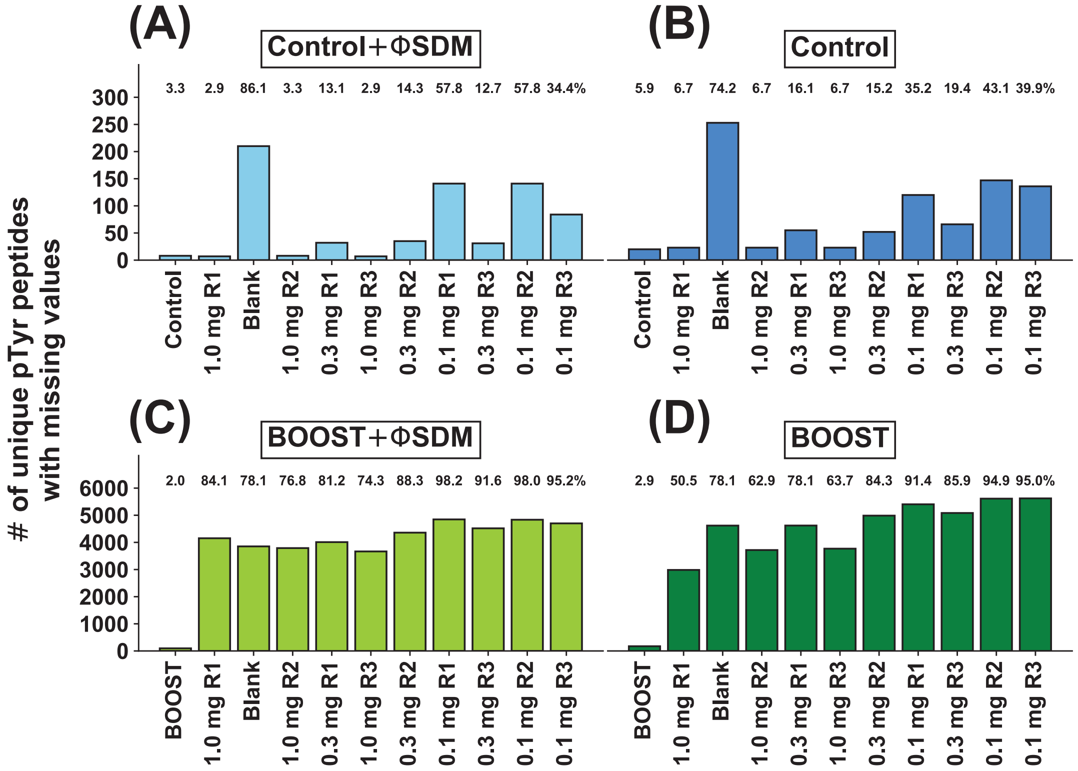


Supporting Figure 2: Disabling ΦSDM reduces the proportion of missing values in BOOST and 1.0 mg Control experiments. The number of missing values in each TMT channel for the (A) 1.0 mg Control experiment with ΦSDM enabled, (B) 1.0 mg Control experiment with ΦSDM disabled, (C) BOOST experiment with ΦSDM enabled, (D) BOOST experiment with ΦSDM disabled. The percentage of missing values in each TMT channel is indicated above each bar.

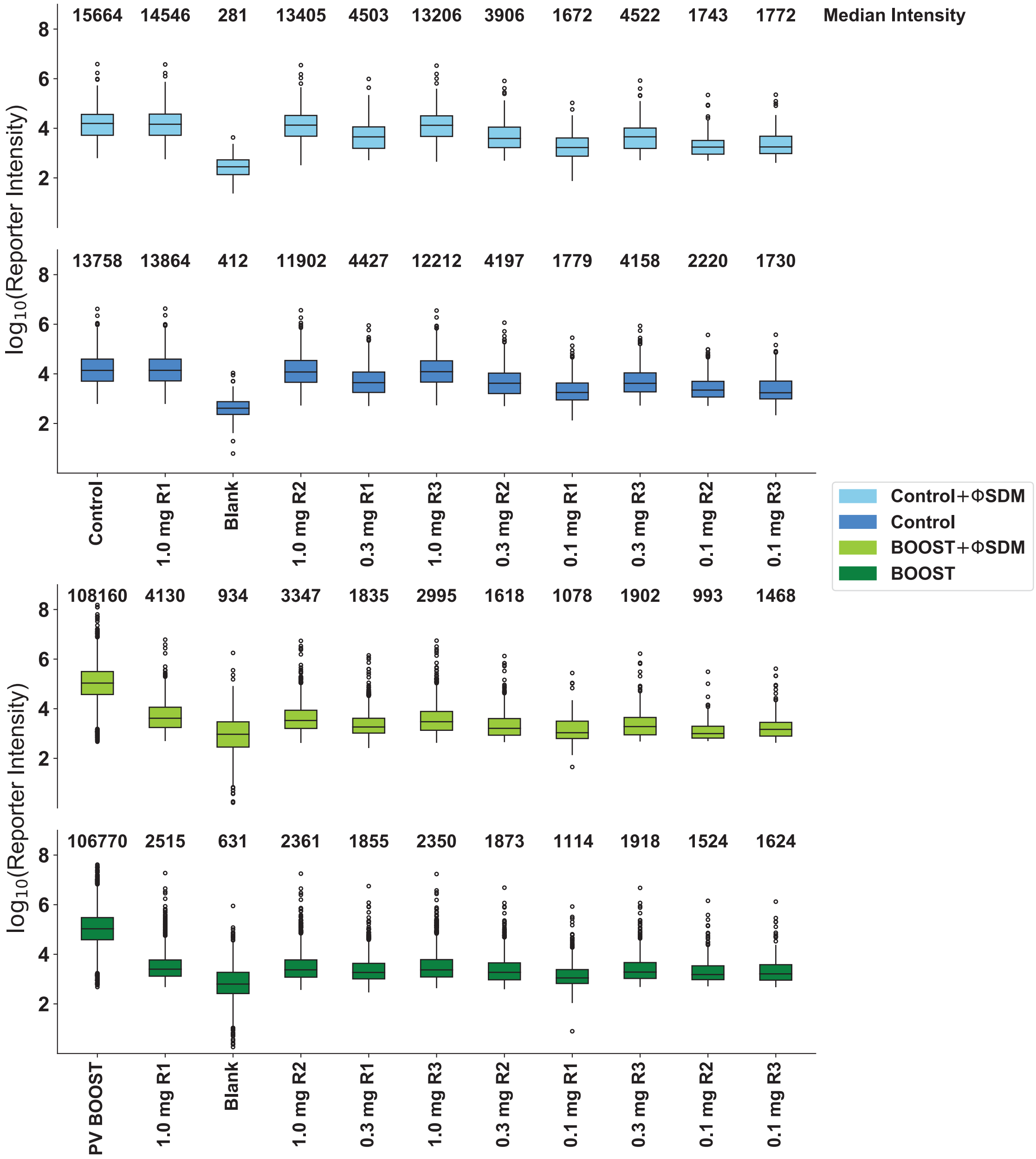


Supporting Figure 3: Box-and-whisker plots showing the $\log_{10}$ transformed reporter intensities for each TMT mix and each condition. Non-transformed, median intensities are displayed above each box-and-whisker plot.

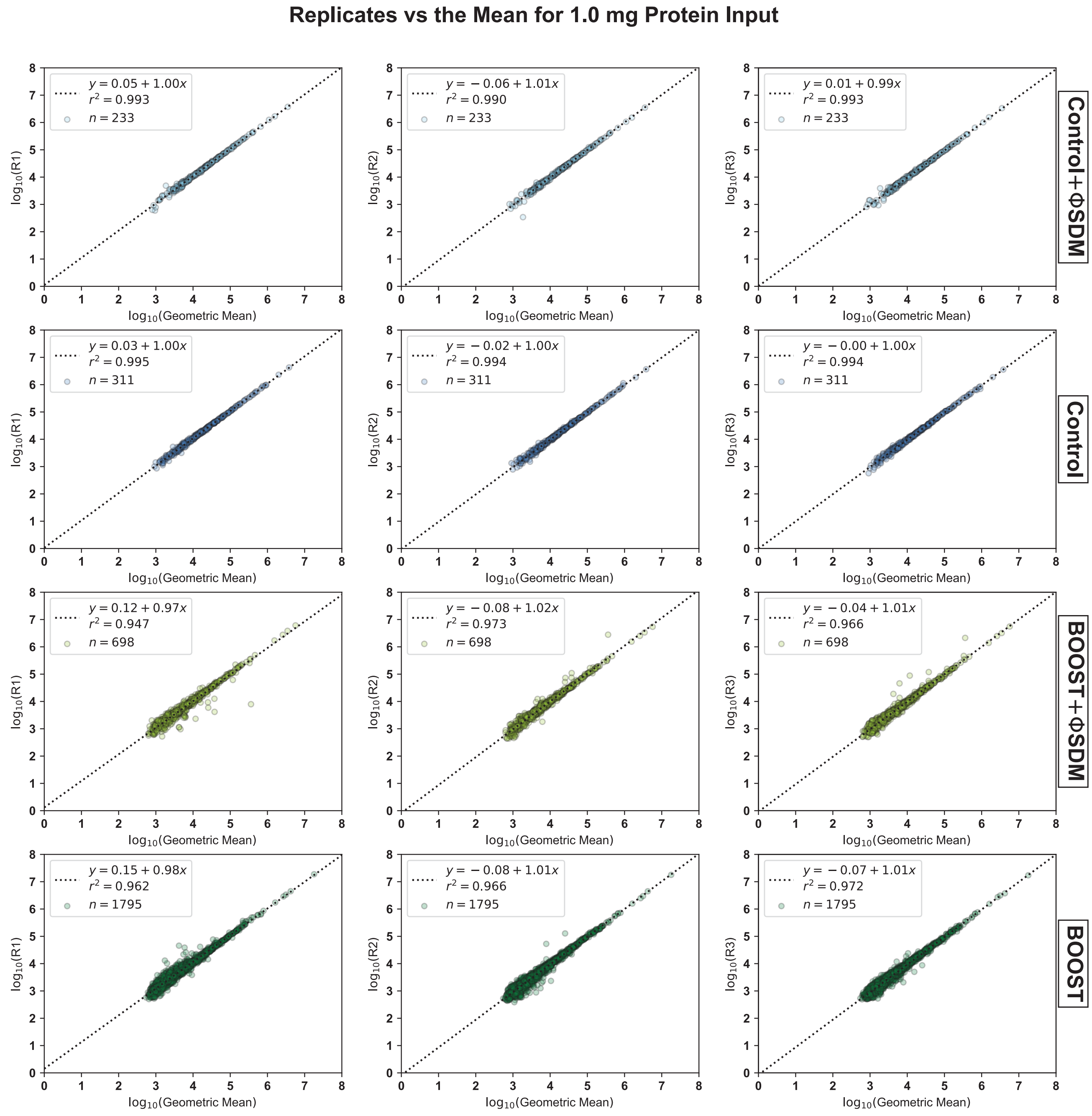


Supporting Figure 4: Comparison between the reporter ion intensity values and the mean value for a given pTyr peptide PSM with no missing values in the 1.0 mg condition of each experiment. The $\log_{10}$(Geometric Mean) is on the $x$-axis, while each replicate intensity value is on the $y$-axis. The legend shows the line of best fit as determined by simple linear regression,[2] and the experiment is noted on the right side of each row.

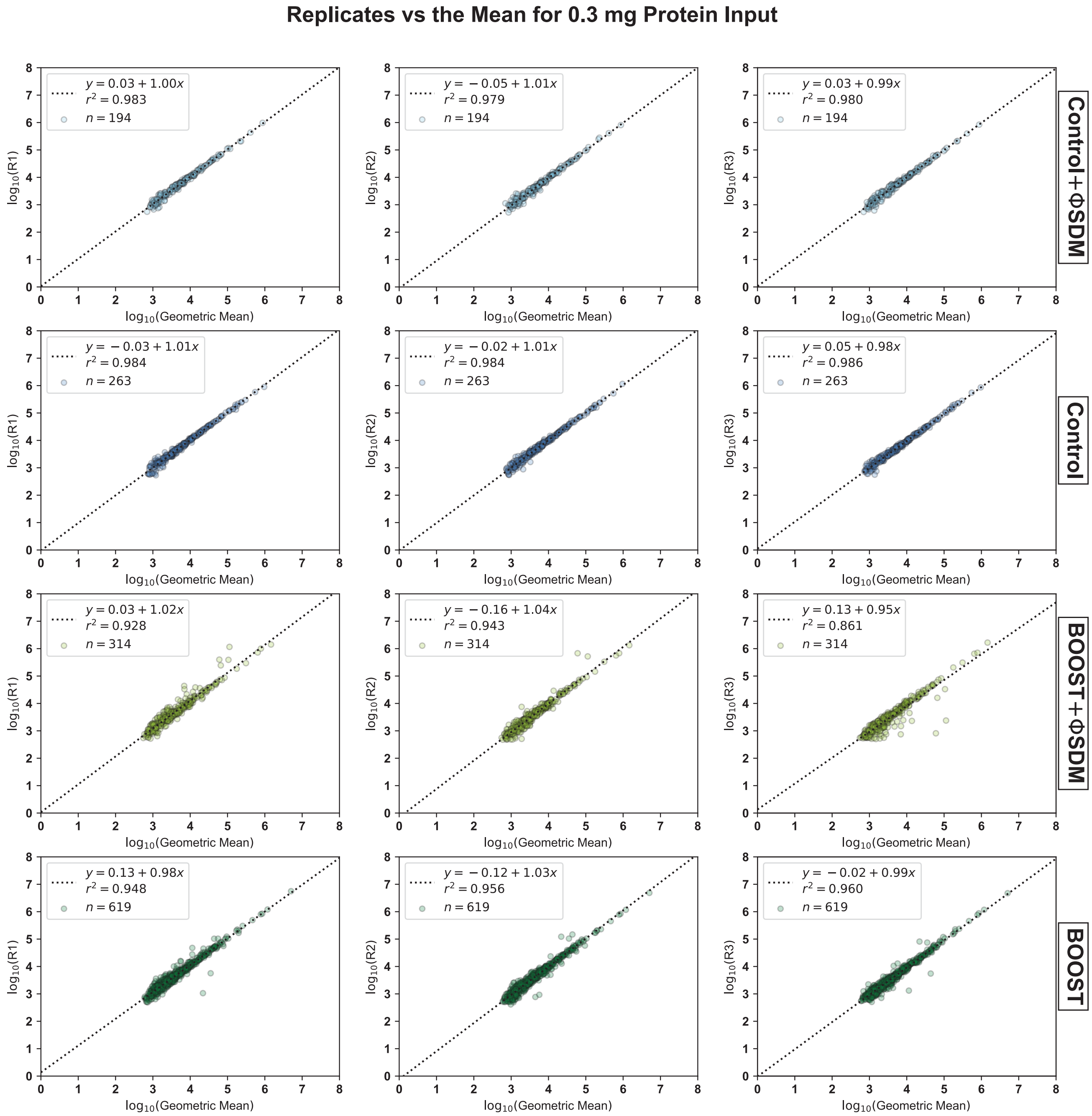


Supporting Figure 5: Comparison between the reporter ion intensity values and the mean value for a given pTyr peptide PSM with no missing values in the 0.3 mg condition of each experiment. The $\log_{10}$(Geometric Mean) is on the $x$-axis, while each replicate intensity value is on the $y$-axis. The legend shows the line of best fit as determined by simple linear regression,[2] and the experiment is noted on the right side of each row.

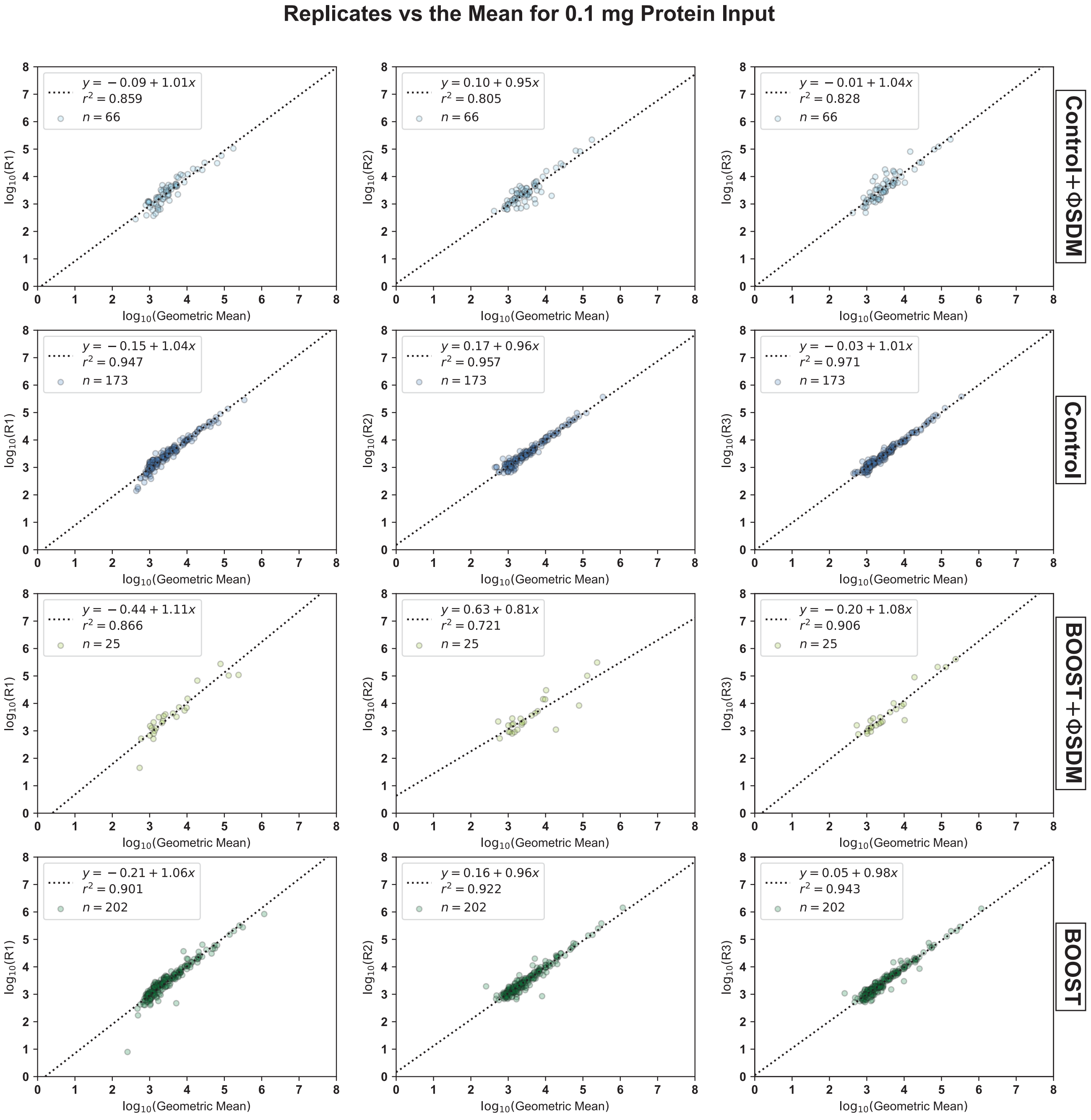


Supporting Figure 6: Comparison between the reporter ion intensity values and the mean value for a given pTyr peptide PSM with no missing values in the 0.1 mg condition of each experiment. The $\log_{10}$(Geometric Mean) is on the $x$-axis, while each replicate intensity value is on the $y$-axis. The legend shows the line of best fit as determined by simple linear regression,[2] and the experiment is noted on the right side of each row.

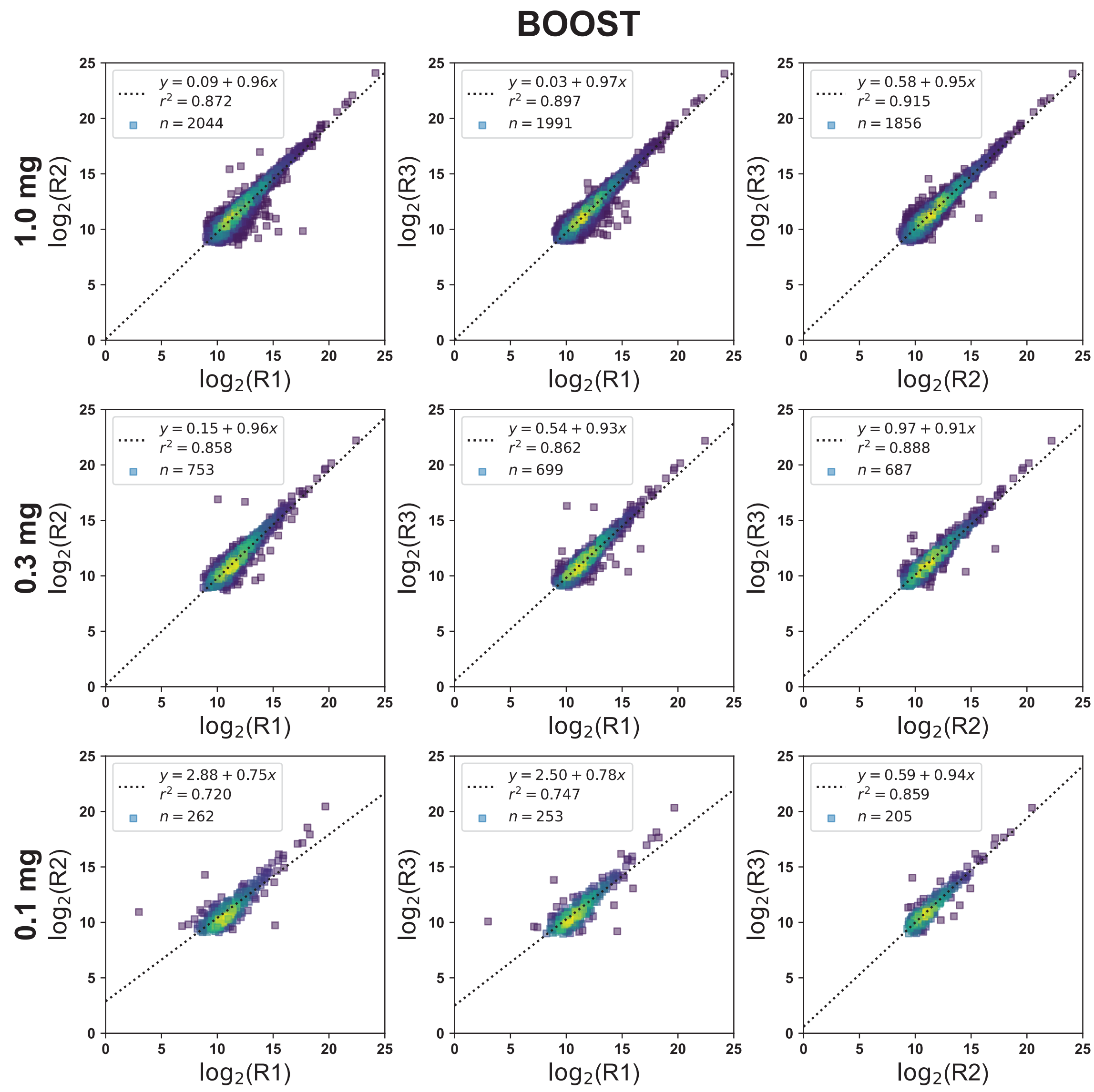


Supporting Figure 7: Replicate reproducibility is stable when ΦSDM is disabled for low protein input camples in the pervanadate BOOST condition. Evaluation of replicate reproducibility in the BOOST experiment (with ΦSDM disabled) using pairwise comparisons of $\log_2$ transformed abundances for pTyr peptide PSMs with the same charge state and assigned sequence. For each comparison, we show the line of best fit as determined by simple least-squares regression,[2] the $r^2$ value as an estimate of the quality of the fitted line, and the total number of points ($n$) in each comparison.

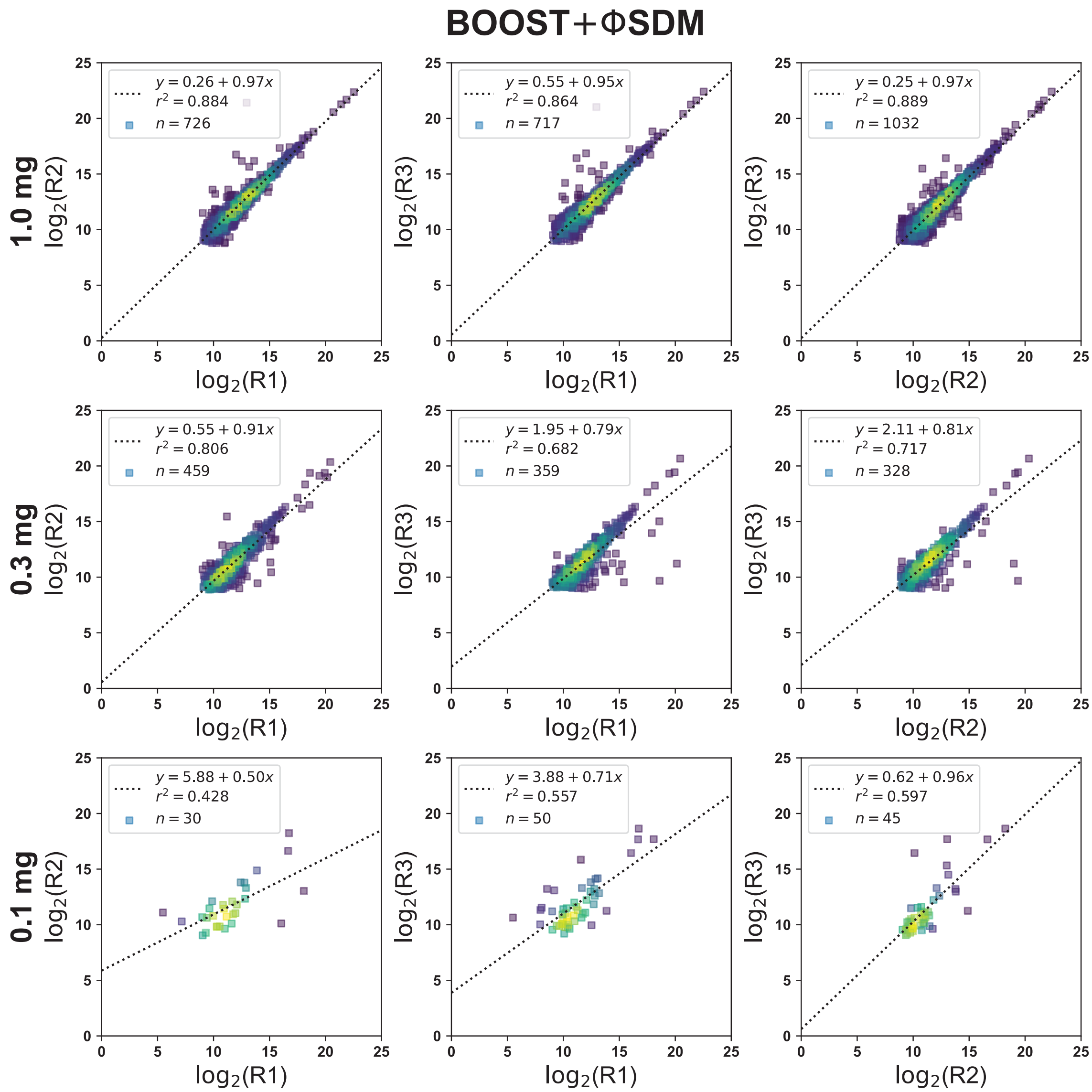


Supporting Figure 8: Replicate reproducibility is severely degraded when ΦSDM is enabled for low protein input samples in the pervanadate BOOST condition. Evaluation of replicate reproducibility in the BOOST experiment (with ΦSDM enabled) using pairwise comparisons of $\log_2$ transformed abundances for pTyr peptide PSMs with the same charge state and assigned sequence. For each comparison, we show the line of best fit as determined by simple least-squares regression,[2] the $r^2$ value as an estimate of the quality of the fitted line, and the total number of points ($n$) in each comparison.

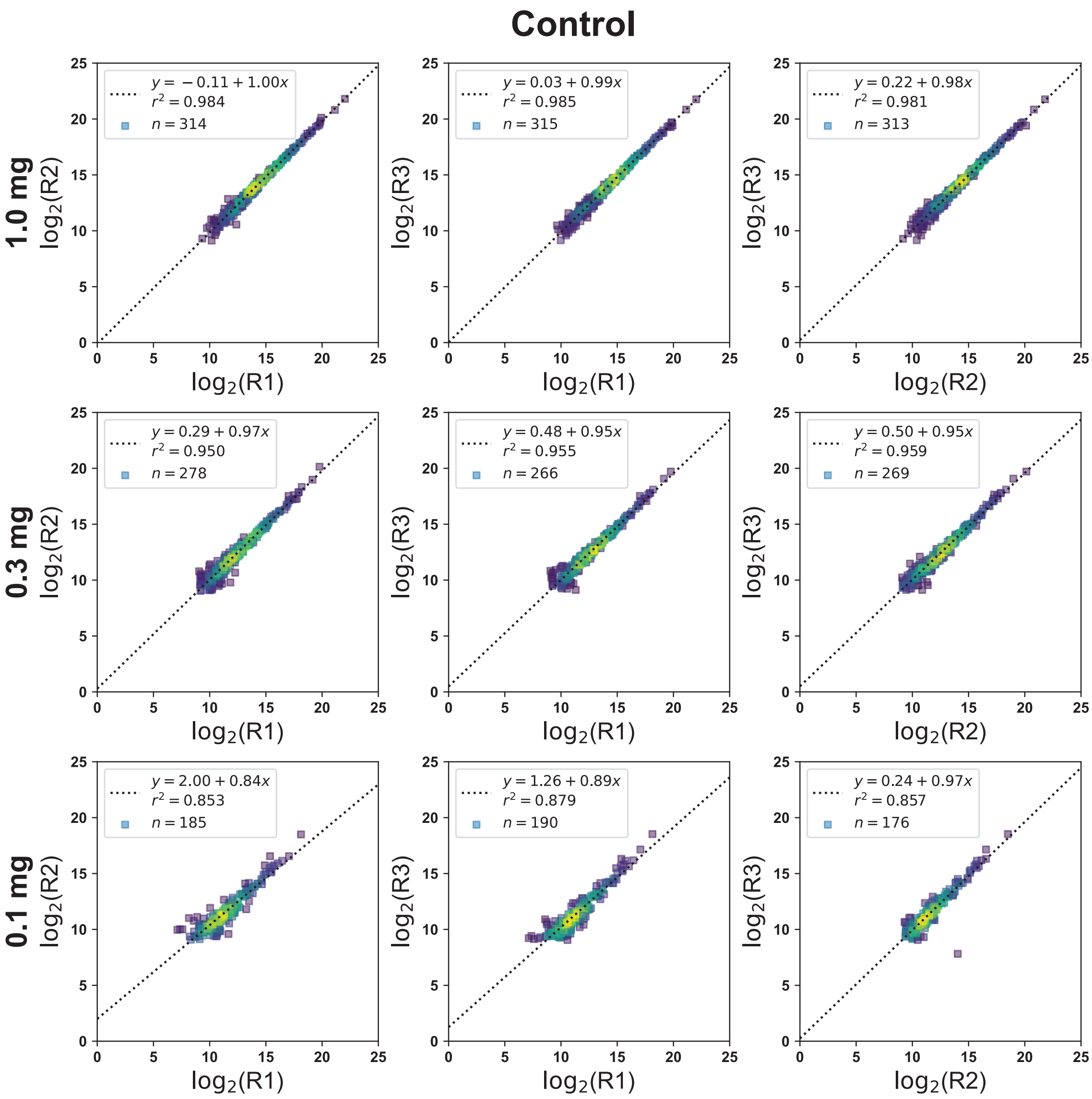


Supporting Figure 9: Replicate reproducibility is stable when ΦSDM is disabled for low protein input camples in the 1.0 mg Control condition. Evaluation of replicate reproducibility in the 1.0 mg Control experiment (with ΦSDM disabled) using pairwise comparisons of $\log_2$ transformed abundances for pTyr peptide PSMs with the same charge state and assigned sequence. For each comparison, we show the line of best fit as determined by simple least-squares regression,[2] the $r^2$ value as an estimate of the quality of the fitted line, and the total number of points ($n$) in each comparison.

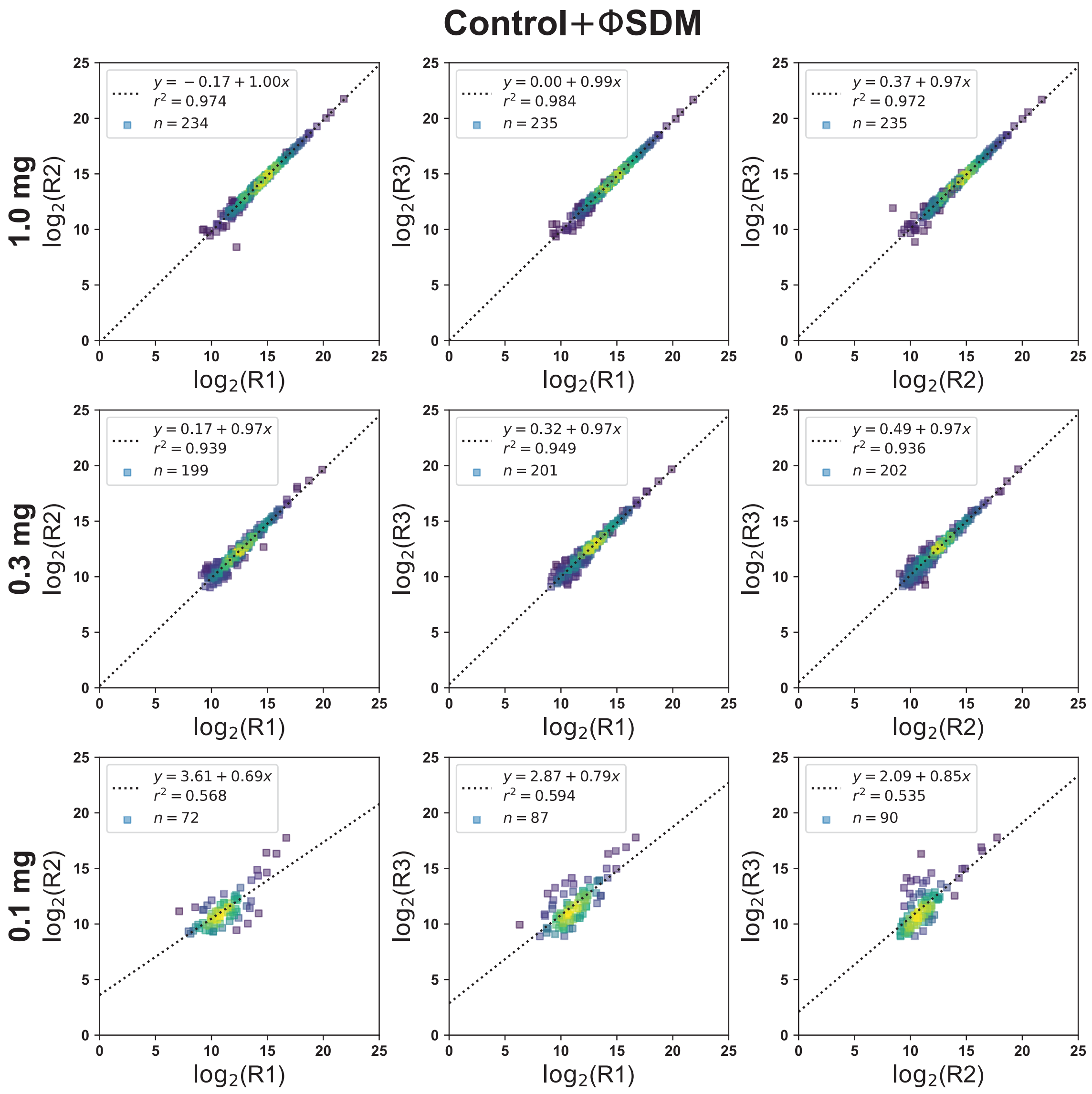


Supporting Figure 10: Replicate reproducibility is degraded when ΦSDM is enabled for low protein input samples in the 1.0 mg Contro conditionl. Evaluation of replicate reproducibility in the 1.0 mg experiment (with ΦSDM enabled) using pairwise comparisons of $\log_2$ transformed abundances for pTyr peptide PSMs with the same charge state and assigned sequence. For each comparison, we show the line of best fit as determined by simple least-squares regression,[2] the $r^2$ value as an estimate of the quality of the fitted line, and the total number of points ($n$) in each comparison.

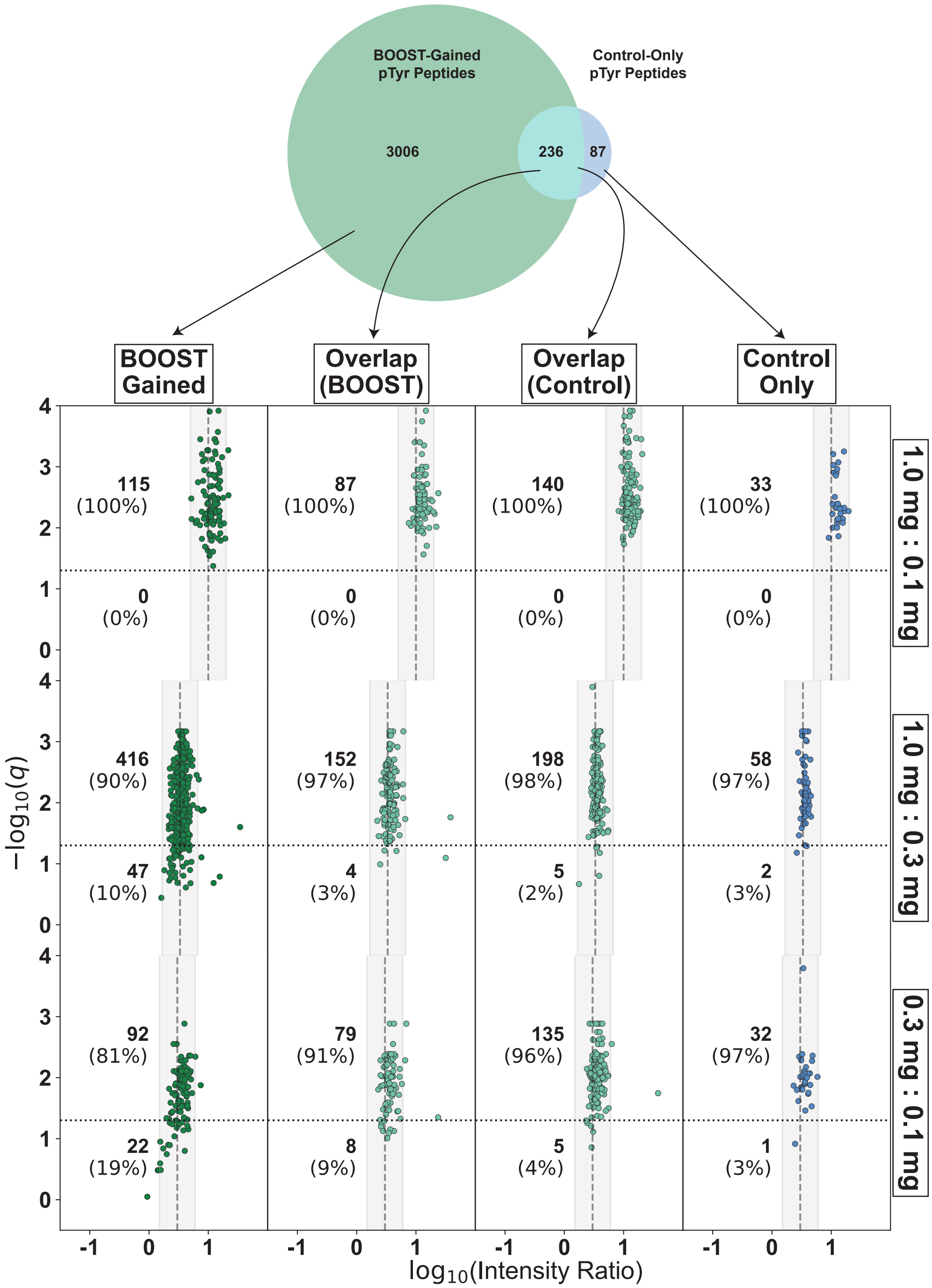


Supporting Figure 11: With ΦSDM disabled, the pervanadate BOOST channel dramatically increases the number of unique pTyr peptide PSMs observed as compared to a 1.0 mg Control channel. A Venn diagram showing the overlap of unique pTyr peptide PSMs between the BOOST and 1.0 mg Control experiments (with ΦSDM disabled). Volcano plots show $-\log_{10}(q\text{-value})$ as a function of $\log_{10}$(Intensity Ratio) for unique pTyr peptide PSMs from groups show in the Venn diagram. For the overlapping section, volcano plots were created using data from both the BOOST experiment and the control experiment acquired with ΦSDM disabled.

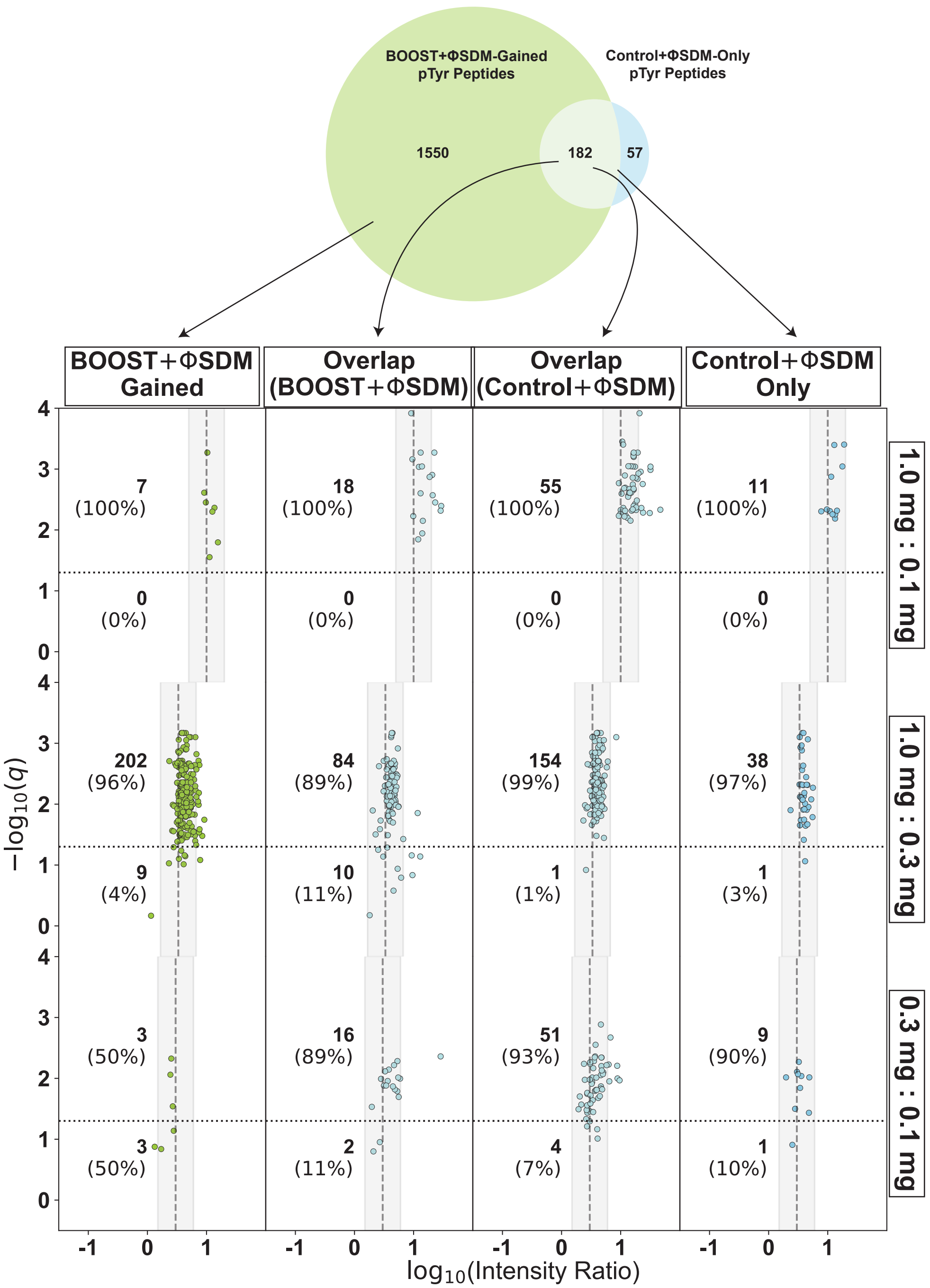


Supporting Figure 12: The pervanadate BOOST channel increases the number of unique pTyr peptide PSMs observed when ΦSDM is enabled, although few peptide PSMs are observed in low abundance samples. A Venn diagram showing the overlap of unique pTyr peptide PSMs between the BOOST+ΦSDM and 1.0 mg Control+ΦSDMl experiments. Volcano plots show $-\log_{10}(q\text{-value})$ as a function of $\log_{10}(\text{Intensity Ratio})$ for unique pTyr peptide PSMs from groups show in the Venn diagram. For the overlapping section, volcano plots were created using data from both the BOOST experiment and the control experiment acquired with ΦSDM enabled.

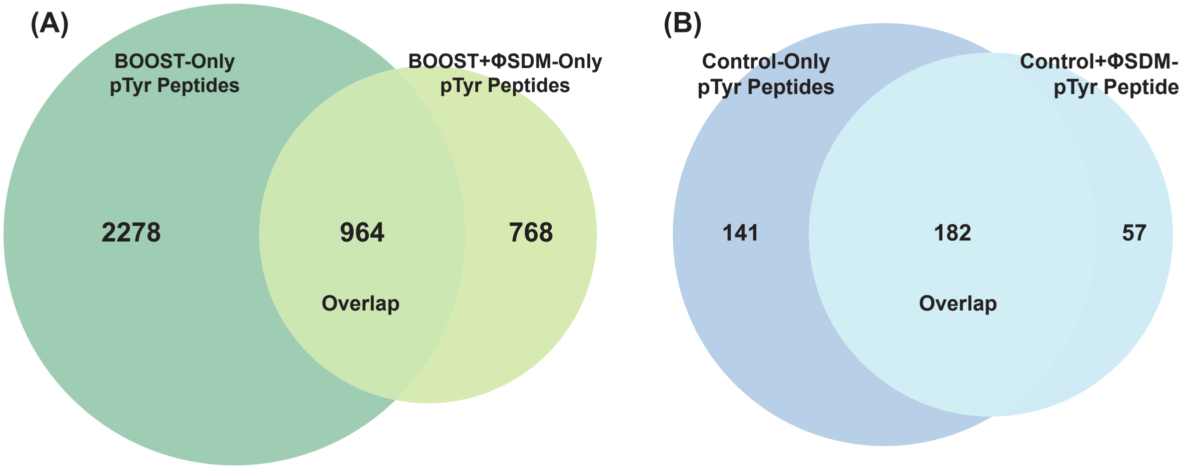


Supporting Figure 13: Enabling ΦSDM results in lower yield in both pervanadate BOOST and 1.0 mg Control conditions. Venn diagrams showing the number of unique pTyr peptide PSMs observed when ΦSDM is enabled or disabled using (A) pervanadate BOOST samples, and (B) 1.0 mg Control samples.

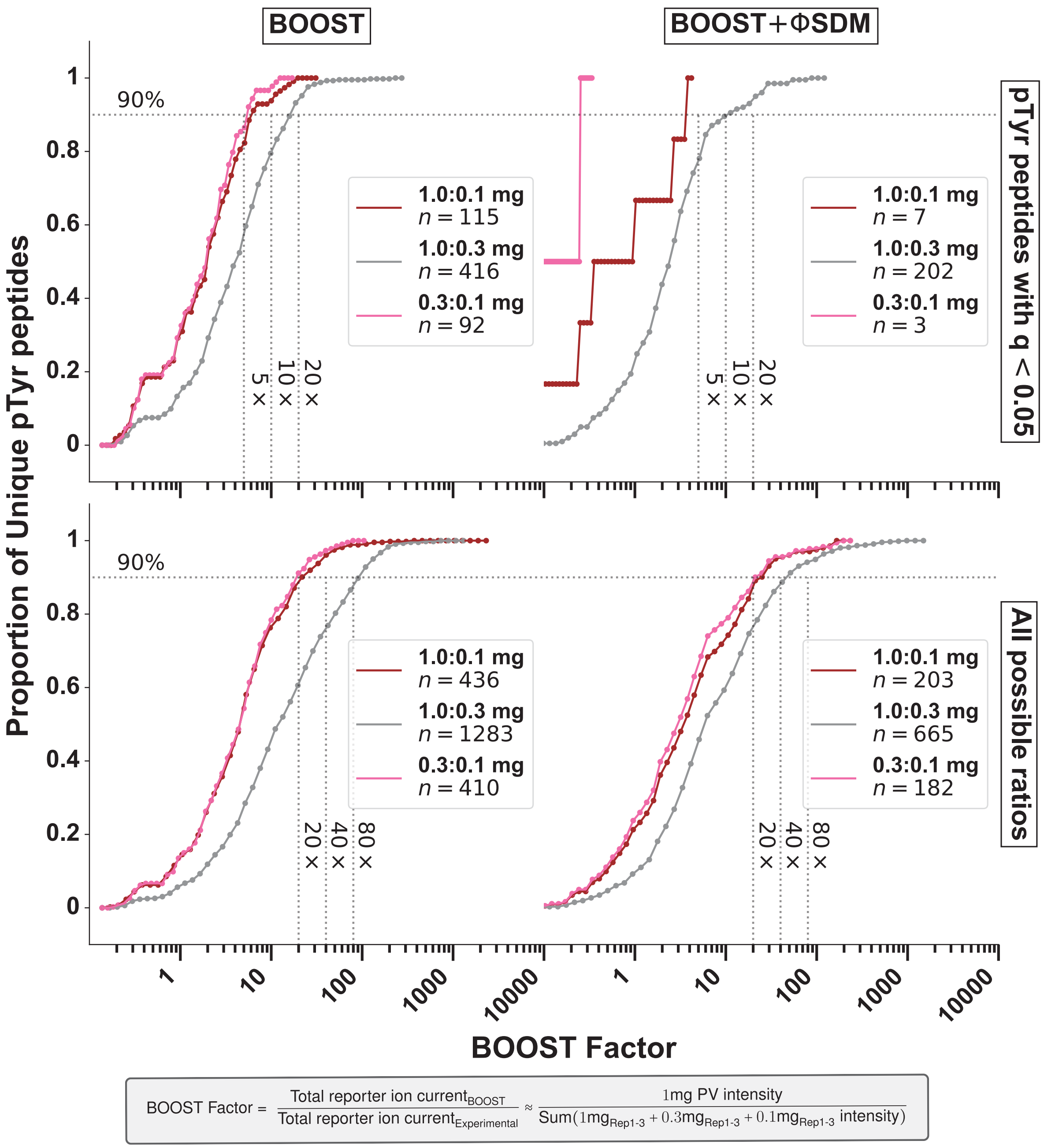


Supporting Figure 14: Enabling ΦSDM decreases quantitation depth, particularly in low abundance samples. Cumulative distribution of BOOST factors for unique pTyr peptides identified in the pervanadate BOOST experiments with ΦSDM disabled or with ΦSDM enabled for pTyr peptides with a statistically significant ratio ($q < 0.05$) or for all calculable ratios. For each cumulative distribution, the range of BOOST factors are split into 50 bins of equal size on a $\log_{10}$ scale.

| Protein | Mouse | Human | Protein | Mouse | Human | Protein | Mouse | Human | Protein | Mouse | Human |
|---|---|---|---|---|---|---|---|---|---|---|---|
| Akt2 | Y122 | Y122 | GADS | Y45 | Y45 | p85α | Y470 | Y470 | TCRζ | Y72 | Y72 |
| CARD11 | Y489 | Y489 | | Y218 | Y222 | | Y688 | Y688 | | Y83 | Y83 |
| Cbl-b | Y363 | Y363^ | Grb2 | Y209 | Y209 | p85β | Y458 | Y464 | | Y111 | Y111 |
| | Y763 | Y763 | GSK3β | Y216 | Y216& | PAK1 | Y142 | Y142 | | Y123 | Y123 |
| CD28 | Y189 | Y191 | Itk | Y40 | Y40 | | Y153 | Y153 | | Y142 | Y142 |
| | Y204 | Y206 | | Y126 | Y120 | | Y474 | Y474+ | | Y153 | Y153 |
| | Y207 | Y209 | | Y226 | Y220 | PAK2 | Y130 | Y130 | Tec | Y205 | Y206 |
| CD3δ | Y149 | Y149 | | Y243 | Y237 | | Y139 | Y139 | | Y227 | Y228 |
| | Y160 | Y160 | | Y517 | Y512 | | Y453 | Y453+ | | Y280 | Y281 |
| CD3ε | Y170 | Y188 | Jnk1 | Y185 | Y185# | PAK6 | Y366 | Y365 | | Y518 | Y519 |
| | Y181 | Y199 | Jnk2 | Y185 | Y185 | PD-1 | Y225 | Y223 | Vav1 | Y110 | Y110 |
| CD3γ | Y160 | Y160 | Jnk3 | Y223 | Y223# | PKCθ | Y28 | Y28 | | Y192 | Y192 |
| | Y171 | Y171 | LAT | Y46 | Y45 | | Y545 | Y545 | | Y541 | Y541 |
| CD45 | Y631 | Y640 | | Y195 | Y220 | PLCγ1 | Y210 | Y210 | | Y791 | Y791 |
| | Y672 | Y681 | Lck | Y192 | Y192 | | Y472 | Y472 | Vav2 | Y142 | Y142 |
| | Y678 | Y687 | | Y394 | Y394* | | Y771 | Y771 | Vav3 | Y141 | Y141 |
| | Y680 | Y689 | | Y414 | Y414 | | Y775 | Y775 | | Y217 | Y217 |
| | Y711 | Y720 | | Y470 | Y470 | | Y783 | Y783 | Zap70 | Y69 | Y69 |
| | Y754 | Y763 | | Y505 | Y505 | | Y1003 | Y1003 | | Y87 | Y87 |
| | Y781 | Y790 | NCK1 | Y13 | Y13 | | Y1253 | Y1253 | | Y164 | Y164 |
| | Y852 | Y861 | | Y55 | Y55 | RHOA | Y66 | Y66% | | Y178 | Y178 |
| | Y871 | Y880 | | Y105 | Y105 | SHP-1 | Y61 | Y61 | | Y198 | Y198 |
| | Y937 | F946 | NCK2 | Y110 | Y110 | | Y64 | Y64 | | Y209 | Y209 |
| | Y969 | Y978 | NFκB-p105 | Y238 | Y240 | | Y213 | Y213 | | Y211 | Y211 |
| | F971 | Y980 | NFAT1 | Y754 | Y752 | | Y214 | Y214 | | Y221 | Y221 |
| CDC42 | Y64 | Y64% | NFAT4 | Y86 | Y86 | | Y276 | Y276 | | Y248 | Y248 |
| CDK4 | Y17 | Y17 | | Y150 | Y150 | | Y301 | Y301 | | Y290 | Y292 |
| CTLA-4 | Y201 | Y201 | p110α | Y317 | Y317 | | Y306 | Y306 | | Y314 | Y315 |
| DLG1 | Y399 | Y399 | p110δ | Y523 | Y524 | | Y374 | Y374 | | Y396 | Y397 |
| | Y761 | Y760 | p38α | Y182 | Y182 | | Y377 | Y377 | | Y491 | Y492 |
| | Y785 | Y784 | p38β | Y182 | Y182 | | Y536 | Y536 | | Y492 | Y493 |
| Erk1 | Y205 | Y204 | p38γ | Y185 | Y185 | | Y541 | Y541 | | Y505 | Y506 |
| Erk2 | Y185 | Y187 | p55γ | Y202 | Y202 | | Y564 | Y564 | | Y596 | Y597 |
| Fyn | Y28 | Y28 | p85α | Y76 | Y76 | SLP76 | Y173 | Y173 | | Y597 | Y598 |
| | Y214 | Y214$ | | Y416 | Y416 | | Y483 | Y483 | | | |
| | Y420 | Y420* | | Y452 | Y452 | TAK1 | Y558 | Y585 | | | |
| | Y440 | Y440! | | Y467 | Y467 | TCRζ | N64 | Y64 | | | |

Supporting Figure 15: Comparison of pTyr sites identified using BOOST in primary T cells from mice and pTyr sites identified using BOOST in Jukat T cells. Flanking sequences (phosphorylation site ±7 amino acids) and phosphorylation sites for each protein in the Kyoto Encyclopedia of Genes and Genomes T cell receptor signaling pathway were manually curated from PhosphoSitePlus®[3] for humans and mice. Flanking sequences for each peptide in the Mouse-BOOST and Jurkat-BOOST datasets were compared with the manually curated KEGG TCR/PhosphoSitePlus flanking sequences and filtered for unique sites. Gene names are colored purple. Phosphotyrosine sites identified in mice are colored red. Phosphotyrosine sites identified in Jurkat T cells are colored green. Sites that were not identified either mice or Jurkat T cells are colored grey. PSMs that can be assigned to multiple proteins: ˆCbl-B;Cbl | %CDC42/RHOA | $Fyn/Yes1 | *Fyn/Yes1/Src/Lck | !Fyn/Yes1/Src | &GSK3β/GSK3α | #Jnk1/Jnk3 | +PAK1/PAK2.
