## Supporting Folder 1 MaxQuant Results for "Mouse primary T cell phosphotyrosine proteomics enabled by BOOST": tables.pdf

### Summary

The summary file contains summary information for all the raw files processed with a single MaxQuant run. The summary information consists of some MaxQuant parameters, information of the raw file contents, and statistics on the peak detection. Based on this file a quick overview can be gathered on the quality of the data in the raw file.

The last row in this file contains the summary information for each column on each of the processed files.

| Name | Separator | Description |
|---|---|---|
| Raw file | | The raw file processed. |
| Experiment | | Experiment name assigned to this LC-MS run in the experimental design. |
| Enzyme | | The protease used to digest the protein sample. |
| Enzyme mode | | The protease used to digest the protein sample. |
| Enzyme first search | | The protease used for the first search. |
| Enzyme mode first search | | The protease used for the first search. |
| Use enzyme first search | | Marked with '+' when a different protease setup was used for the first search. |
| Variable modifications | | The variable modification(s) used during the identification of peptides. |
| Fixed modifications | | The fixed modification(s) used during the identification of peptides. |
| Multi modifications | | The multi modification(s) used during the identification of peptides. |
| Variable modifications first search | | The variable modification(s) used during the first search. |
| Use variable modifications first search | | Marked with '+' when different variable modifications were used for the first search. |
| Requantify | | The number of labels used. |
| Multiplicity | | The number of labels used. |
| Max. missed cleavages | | The maximum allowed number of missed cleavages. |
| Labels0 | | The labels used in the labeling experiment. Allowed values for X: 0=light; 1=medium; 2=heavy label partner. |
| LC-MS run type | | The type of LC-MS run. Usually it will be 'Standard' which refers to a conventional shotgun proteomics run with data-dependent MS/MS. |
| Time-dependent recalibration | | When marked with '+', time-dependent recalibration was applied to improve the data quality. |
| MS | | The number of MS spectra recorded in this raw file. |
| MS/MS | | The number of MS/MS spectra recorded in this raw file. |
| MS3 | | The number of MS3 spectra recorded in this raw file. |
| MS/MS Submitted | | The number of tandem MS spectra submitted for analysis. |
| MS/MS Submitted (SIL) | | The number of tandem MS spectra submitted for analysis, where the precursor ion was detected as part of a labeling cluster. |
| MS/MS Submitted (ISO) | | The number of tandem MS spectra submitted for analysis, where the precursor ion was detected as an isotopic pattern. |
| MS/MS Submitted (PEAK) | | The number of tandem MS spectra submitted for analysis, where the precursor ion was detected as a single peak. |
| MS/MS Identified | | The total number of identified tandem MS spectra. |
| MS/MS Identified (SIL) | | The total number of identified tandem MS spectra, where the precursor ion was detected as part of a labeling cluster. |
| MS/MS Identified (ISO) | | The total number of identified tandem MS spectra, where the precursor ion was detected as an isotopic pattern. |
| MS/MS Identified (PEAK) | | The total number of identified tandem MS spectra, where the precursor ion was detected as a single peak. |
| MS/MS Identified [%] | | The percentage of identified tandem MS spectra. |
| MS/MS Identified (SIL) [%] | | The percentage of identified tandem MS spectra, where the precursor ion was detected as part of a labeling cluster. |
| MS/MS Identified (ISO) [%] | | The percentage of identified tandem MS spectra, where the precursor ion was detected as an isotopic pattern. |
| MS/MS Identified (PEAK) [%] | | The percentage of identified tandem MS spectra, where the precursor ion was detected as a single peak. |
| Peptide Sequences Identified | | The total number of unique peptide amino acid sequences identified from the recorded tandem mass spectra. |
| Peaks | | The total number of peaks detected in the full scans. |
| Peaks Sequenced | | The total number of peaks sequenced by tandem MS. |

| | | |
|---|---|---|
| Peaks Sequenced [%] | | The percentage of peaks sequenced by tandem MS. |
| Peaks Repeatedly Sequenced | | The total number of peaks repeatedly sequenced (i.e. 1 or more times) by tandem MS. |
| Peaks Repeatedly Sequenced [%] | | The percentage of peaks repeatedly sequenced (i.e. 1 or more times) by tandem MS. |
| Isotope Patterns | | The total number of detected isotope patterns. |
| Isotope Patterns Sequenced | | The total number of isotope patterns sequenced by tandem MS. |
| Isotope Patterns Sequenced (z>1) | | The total number of isotope patterns sequenced by tandem MS with a charge state of 2 or more. |
| Isotope Patterns Sequenced [%] | | The percentage of isotope patterns sequenced by tandem MS. |
| Isotope Patterns Sequenced (z>1) [%] | | The percentage of isotope patterns sequenced by tandem MS with a charge state of 2 or more. |
| Isotope Patterns Repeatedly Sequenced | | The total number of isotope patterns repeatedly sequenced (i.e. 1 or more times) by tandem MS. |
| Isotope Patterns Repeatedly Sequenced [%] | | The percentage of isotope patterns repeatedly sequenced (i.e. 1 or more times) by tandem MS. |
| Recalibrated | | When marked with '+', the masses taken from the raw file were recalibrated. |
| Av. Absolute Mass Deviation [ppm] | | The average absolute mass deviation found comparing to the identification mass in parts per million. |
| Mass Standard Deviation [ppm] | | The standard deviation of the mass deviation found comparing to the identification mass in parts per million. |
| Av. Absolute Mass Deviation [mDa] | | The average absolute mass deviation found comparing to the identification mass in milli-Dalton. |
| Mass Standard Deviation [mDa] | | The standard deviation of the mass deviation found comparing to the identification mass in milli-Dalton. |

### Evidence

The evidence file combines all the information about the identified peptides and normally is the only file required for processing the results. Additional information about the peptides, modifications, proteins, etc. can be found in the other files by unique identifier linkage.

| **Name** | **Separator** | **Description** |
|---|---|---|
| Sequence | | The identified AA sequence of the peptide. |
| Length | | The length of the sequence stored in the column 'Sequence'. |
| Modifications | | Post-translational modifications contained within the identified peptide sequence. |
| Modified sequence | | Sequence representation including the post-translational modifications (abbreviation of the modification in brackets before the modified AA). The sequence is always surrounded by underscore characters ('_'). |
| Oxidation (M) Probabilities | | Sequence representation of the peptide including PTM positioning probabilities ([0..1], where 1 is best match) for 'Oxidation (M)'. |
| Phospho (STY) Probabilities | | Sequence representation of the peptide including PTM positioning probabilities ([0..1], where 1 is best match) for 'Phospho (STY)'. |
| Oxidation (M) Score Diffs | | Sequence representation for each of the possible PTM positions in each possible configuration, the difference is calculated between the identification score with the PTM added to that position and the best scoring identification where no PTM is added to that position. When this value is negative, it is unlikely that the particular modification is located at this position. |
| Phospho (STY) Score Diffs | | Sequence representation for each of the possible PTM positions in each possible configuration, the difference is calculated between the identification score with the PTM added to that position and the best scoring identification where no PTM is added to that position. When this value is negative, it is unlikely that the particular modification is located at this position. |
| Acetyl (Protein N-term) | | The number of occurrences of the modification 'Acetyl (Protein N-term)'. |
| Oxidation (M) | | The number of occurrences of the modification 'Oxidation (M)'. |
| Phospho (STY) | | The number of occurrences of the modification 'Phospho (STY)'. |
| Missed cleavages | | Number of missed enzymatic cleavages. |
| Proteins | | The identifiers of the proteins this particular peptide is associated with. |
| Leading proteins | | The identifiers of the proteins in the proteinGroups file, with this protein as best match, this particular peptide is associated with. When multiple matches are found here, the best scoring protein can be found in the 'Leading Razor Protein' column. |
| Leading razor protein | | The identifier of the best scoring protein, from the proteinGroups file this, this peptide is associated to. |
| Gene names | | Names of genes this peptide is associated with. |
| Protein names | | Names of proteins this peptide is associated with. |
| Type |  | The type of the feature. 'MSMS' for an MS/MS spectrum without an MS1 isotope pattern assigned. 'ISO-MSMS' MS1 isotope cluster identified by MS/MS. 'MULTI-MSMS' MS1 labeling cluster identified by MS/MS. 'MULTI-SECPEP' MS1 labeling cluster identified by MS/MS as second peptide. 'MULTI-MATCH' MS1 labeling cluster identified by matching between runs. In case of label-free data there is no difference between 'MULTI' and 'ISO'. |
| Raw file | | The name of the RAW-file the mass spectral data was derived from. |
| Experiment | | |
| MS/MS m/z | | The m/z used for fragmentation (not necessarily the mono-isotopic m/z). |
| Charge | | The charge-state of the precursor ion. |
| m/z | | The recalibrated mass-over-charge value of the precursor ion. |
| Mass | | The predicted monoisotopic mass of the identified peptide sequence. |
| Uncalibrated - Calibrated m/z [ppm] | | The difference between the uncalibrated and recalibrated mass-over-charge value of the precursor ion measured in parts-per-million. This gives an indication of the mass drift in the original data, which was automatically corrected by MaxQuant. |

| | | |
|---|---|---|
| Uncalibrated - Calibrated m/z [Da] | | The difference between the uncalibrated and recalibrated mass-over-charge value of the precursor ion measured in parts-per-million. This gives an indication of the mass drift in the original data, which was automatically corrected by MaxQuant. |
| Mass error [ppm] | | Mass error of the recalibrated mass-over-charge value of the precursor ion in comparison to the predicted monoisotopic mass of the identified peptide sequence in parts per million. |
| Mass error [Da] | | Mass error of the recalibrated mass-over-charge value of the precursor ion in comparison to the predicted monoisotopic mass of the identified peptide sequence in milli-Dalton. |
| Uncalibrated mass error [ppm] | | Mass error of the uncalibrated mass-over-charge value of the precursor ion in comparison to the predicted monoisotopic mass of the identified peptide sequence.<br><br>Note: This column can contain missing values (denoted as NaN). |
| Uncalibrated mass error [Da] | | Mass error of the uncalibrated mass-over-charge value of the precursor ion in comparison to the predicted monoisotopic mass of the identified peptide sequence.<br><br>Note: This column can contain missing values (denoted as NaN). |
| Max intensity m/z 0 | | Summed up eXtracted Ion Current (XIC) of all isotopic clusters associated with the identified AA sequence. In case of a labeled experiment this is the total intensity of all the isotopic patterns in the label cluster. |
| Retention time | | The uncalibrated retention time in minutes in the elution profile of the precursor ion. |
| Retention length |  | The total retention time length of the peak (last time point first time point). |
| Calibrated retention time | | The recalibrated retention time in minutes in the elution profile of the precursor ion. |
| Calibrated retention time start | | The recalibrated retention start in minutes in the elution profile of the precursor ion. |
| Calibrated retention time finish | | The recalibrated retention finish in minutes in the elution profile of the precursor ion. |
| Retention time calibration | | The difference in minutes between the uncalibrated and recalibrated retention time. This gives an indication of the retention time drift in the original data, which was automatically corrected by MaxQuant.<br><br>Note: This column can contain missing values (NaN). |
| Match time difference | | When the option match between runs is used in MaxQuant, this value indicates the time difference between the feature from the raw file it was taken from and the feature from the raw file it was matched to. |
| Match m/z difference | | When the option match between runs is used in MaxQuant, this value indicates the m/z difference between the feature from the raw file it was taken from and the feature from the raw file it was matched to. |
| Match q-value | | This is the q-value for features that have been identified by 'matching between runs'. |
| Match score | | The andromeda score of the MS/MS identification that is the source of this identification by 'matching between runs'. |
| Number of data points | | The number of data points (peak centroids) collected for this peptide feature. |
| Number of scans | | The number of MS scans that the 3d peaks of this peptide feature are overlapping with. |
| Number of isotopic peaks | | The number of isotopic peaks contained in this peptide feature. |
| PIF | | Short for Parent Ion Fraction; indicates the fraction the target peak makes up of the total intensity in the inclusion window. |
| Fraction of total spectrum | | The percentage the ion intensity makes up of the total intensity of the whole spectrum. |
| Base peak fraction | | The percentage the parent ion intensity in comparison to the highest peak in the MS spectrum. |
| PEP | | Posterior Error Probability of the identification. This value essentially operates as a p-value, where smaller is more significant. |
| MS/MS count | | The number of sequencing events for this sequence, which matches the number of identifiers stored in the column MS/MS IDs. |
| MS/MS scan number | | The RAW-file derived scan number of the MS/MS with the highest peptide identification score (the highest score is stored in the column 'Score'). |
| Score | | Andromeda score for the best associated MS/MS spectrum. |
| Delta score | | Score difference to the second best identified peptide. |
| Combinatorics | | Number of possible distributions of the modifications over the peptide sequence. |

| | | |
|---|---|---|
| Intensity | | Summed up eXtracted Ion Current (XIC) of all isotopic clusters associated with the identified AA sequence. In case of a labeled experiment this is the total intensity of all the isotopic patterns in the label cluster. |
| Reporter intensity corrected 1 | | |
| Reporter intensity corrected 2 | | |
| Reporter intensity corrected 3 | | |
| Reporter intensity corrected 4 | | |
| Reporter intensity corrected 5 | | |
| Reporter intensity corrected 6 | | |
| Reporter intensity corrected 7 | | |
| Reporter intensity corrected 8 | | |
| Reporter intensity corrected 9 | | |
| Reporter intensity corrected 10 | | |
| Reporter intensity corrected 11 | | |
| Reporter intensity 1 | | |
| Reporter intensity 2 | | |
| Reporter intensity 3 | | |
| Reporter intensity 4 | | |
| Reporter intensity 5 | | |
| Reporter intensity 6 | | |
| Reporter intensity 7 | | |
| Reporter intensity 8 | | |
| Reporter intensity 9 | | |
| Reporter intensity 10 | | |
| Reporter intensity 11 | | |
| Reporter intensity count 1 | | |
| Reporter intensity count 2 | | |
| Reporter intensity count 3 | | |
| Reporter intensity count 4 | | |
| Reporter intensity count 5 | | |
| Reporter intensity count 6 | | |
| Reporter intensity count 7 | | |
| Reporter intensity count 8 | | |
| Reporter intensity count 9 | | |
| Reporter intensity count 10 | | |
| Reporter intensity count 11 | | |
| Reverse | | When marked with '+', this particular peptide was found to be part of a protein derived from the reversed part of the decoy database. These should be removed for further data analysis. |
| Potential contaminant | | When marked with '+', this particular peptide was found to be part of a commonly occurring contaminant. These should be removed for further data analysis. |
| id | | A unique (consecutive) identifier for each row in the evidence table, which is used to cross-link the information in this file with the information stored in the other files. |
| Protein group IDs | | The identifier of the protein-group this redundant peptide sequence is associated with, which can be used to look up the extended protein information in the file 'proteinGroups.txt'. As a single peptide can be linked to multiple proteins (e.g. in the case of razor-proteins), multiple ids can be stored here separated by a semicolon. As a protein can be identified by multiple peptides, the same id can be found in different rows. |
| Peptide ID | | The identifier of the non-redundant peptide sequence. |
| Mod. peptide ID | | Identifier of the associated modification summary stored in the file 'modificationSpecificPeptides.txt'. |
| MS/MS IDs | | Identifier(s) of the associated MS/MS summary(s) stored in the file 'msms.txt'. |
| Best MS/MS | | Identifier(s) of the best MS/MS associated spectrum stored in the file 'msms.txt'. |
| Oxidation (M) site IDs | | Identifier(s) of the modification summary stored in the file 'Oxidation (M)Sites.txt'. |
| Phospho (STY) site IDs | | Identifier(s) of the modification summary stored in the file 'Phospho (STY)Sites.txt'. |
| Taxonomy IDs | | Taxonomy identifiers. |

### Peptides

The peptides table contains information on the identified peptides in the processed raw-files.

| Name | Separator | Description |
|---|---|---|
| Sequence | | The amino acid sequence of the identified peptide. |
| N-term cleavage window | | Sequence window from -15 to 15 around the N-terminal cleavage site of this peptide. |
| C-term cleavage window | | Sequence window from -15 to 15 around the C-terminal cleavage site of this peptide. |
| Amino acid before | | The amino acid in the protein sequence before the peptide. |
| First amino acid | | The amino acid in the first position of the peptide sequence. |
| Second amino acid | | The amino acid in the first position of the peptide sequence. |
| Second last amino acid | | The amino acid in the last position of the peptide sequence. |
| Last amino acid | | The amino acid in the last position of the peptide sequence. |
| Amino acid after | | The amino acid in the protein sequence after the peptide. |
| A Count | | The number of instances of the 'A' amino acid contained within the sequence. |
| R Count | | The number of instances of the 'R' amino acid contained within the sequence. |
| N Count | | The number of instances of the 'N' amino acid contained within the sequence. |
| D Count | | The number of instances of the 'D' amino acid contained within the sequence. |
| C Count | | The number of instances of the 'C' amino acid contained within the sequence. |
| Q Count | | The number of instances of the 'Q' amino acid contained within the sequence. |
| E Count | | The number of instances of the 'E' amino acid contained within the sequence. |
| G Count | | The number of instances of the 'G' amino acid contained within the sequence. |
| H Count | | The number of instances of the 'H' amino acid contained within the sequence. |
| I Count | | The number of instances of the 'I' amino acid contained within the sequence. |
| L Count | | The number of instances of the 'L' amino acid contained within the sequence. |
| K Count | | The number of instances of the 'K' amino acid contained within the sequence. |
| M Count | | The number of instances of the 'M' amino acid contained within the sequence. |
| F Count | | The number of instances of the 'F' amino acid contained within the sequence. |
| P Count | | The number of instances of the 'P' amino acid contained within the sequence. |
| S Count | | The number of instances of the 'S' amino acid contained within the sequence. |
| T Count | | The number of instances of the 'T' amino acid contained within the sequence. |
| W Count | | The number of instances of the 'W' amino acid contained within the sequence. |
| Y Count | | The number of instances of the 'Y' amino acid contained within the sequence. |
| V Count | | The number of instances of the 'V' amino acid contained within the sequence. |
| U Count | | The number of instances of the 'U' amino acid contained within the sequence. |
| O Count | | The number of instances of the 'O' amino acid contained within the sequence. |
| Length | | The length of the sequence stored in the column "Sequence". |
| Missed cleavages | | Number of missed enzymatic cleavages. |
| Mass | | Monoisotopic mass of the peptide. |
| Proteins | | Identifiers of proteins this peptide is associated with. |
| Leading razor protein | | Identifier of the leading protein in the protein group which uses this peptide for quantification. (Either unique or razor.) |
| Start position | | Position of the first amino acid of this peptide in the protein sequence. (one-based) |
| End position | | Position of the last amino acid of this peptide in the protein sequence. (one-based) |

| | | |
|---|---|---|
| Gene names | | Names of genes this peptide is associated with. |
| Protein names | | Names of proteins this peptide is associated with. |
| Unique (Groups) | | When marked with '+', this particular peptide is unique to a single protein group in the proteinGroups file. |
| Unique (Proteins) | | When marked with '+', this particular peptide is unique to a single protein sequence in the fasta file(s). |
| Charges | | All charge states that have been observed. |
| PEP | | Posterior Error Probability of the identification. This value essentially operates as a p-value, where smaller is more significant. |
| Score | | Highest Andromeda score for the associated MS/MS spectra. |
| Experiment 1 | | Number of evidence entries for this 'Experiment'. |
| Experiment 2 | | Number of evidence entries for this 'Experiment'. |
| Experiment 3 | | Number of evidence entries for this 'Experiment'. |
| Experiment 4 | | Number of evidence entries for this 'Experiment'. |
| Reporter intensity corrected 1 | | |
| Reporter intensity corrected 2 | | |
| Reporter intensity corrected 3 | | |
| Reporter intensity corrected 4 | | |
| Reporter intensity corrected 5 | | |
| Reporter intensity corrected 6 | | |
| Reporter intensity corrected 7 | | |
| Reporter intensity corrected 8 | | |
| Reporter intensity corrected 9 | | |
| Reporter intensity corrected 10 | | |
| Reporter intensity corrected 11 | | |
| Reporter intensity 1 | | |
| Reporter intensity 2 | | |
| Reporter intensity 3 | | |
| Reporter intensity 4 | | |
| Reporter intensity 5 | | |
| Reporter intensity 6 | | |
| Reporter intensity 7 | | |
| Reporter intensity 8 | | |
| Reporter intensity 9 | | |
| Reporter intensity 10 | | |
| Reporter intensity 11 | | |
| Reporter intensity count 1 | | |
| Reporter intensity count 2 | | |
| Reporter intensity count 3 | | |
| Reporter intensity count 4 | | |
| Reporter intensity count 5 | | |
| Reporter intensity count 6 | | |
| Reporter intensity count 7 | | |
| Reporter intensity count 8 | | |
| Reporter intensity count 9 | | |
| Reporter intensity count 10 | | |
| Reporter intensity count 11 | | |
| Reporter intensity corrected 1 1 | | |
| Reporter intensity corrected 2 1 | | |
| Reporter intensity corrected 3 1 | | |
| Reporter intensity corrected 4 1 | | |
| Reporter intensity corrected 5 1 | | |
| Reporter intensity corrected 6 1 | | |
| Reporter intensity corrected 7 1 | | |
| Reporter intensity corrected 8 1 | | |
| Reporter intensity corrected 9 1 | | |
| Reporter intensity corrected 10 1 | | |
| Reporter intensity corrected 11 1 | | |
| Reporter intensity 1 1 | | |
| Reporter intensity 2 1 | | |
| Reporter intensity 3 1 | | |
| Reporter intensity 4 1 | | |
| Reporter intensity 5 1 | | |
| Reporter intensity 6 1 | | |

| |
|---|
| Reporter intensity 7 1 |
| Reporter intensity 8 1 |
| Reporter intensity 9 1 |
| Reporter intensity 10 1 |
| Reporter intensity 11 1 |
| Reporter intensity count 1 1 |
| Reporter intensity count 2 1 |
| Reporter intensity count 3 1 |
| Reporter intensity count 4 1 |
| Reporter intensity count 5 1 |
| Reporter intensity count 6 1 |
| Reporter intensity count 7 1 |
| Reporter intensity count 8 1 |
| Reporter intensity count 9 1 |
| Reporter intensity count 10 1 |
| Reporter intensity count 11 1 |
| Reporter intensity corrected 1 2 |
| Reporter intensity corrected 2 2 |
| Reporter intensity corrected 3 2 |
| Reporter intensity corrected 4 2 |
| Reporter intensity corrected 5 2 |
| Reporter intensity corrected 6 2 |
| Reporter intensity corrected 7 2 |
| Reporter intensity corrected 8 2 |
| Reporter intensity corrected 9 2 |
| Reporter intensity corrected 10 2 |
| Reporter intensity corrected 11 2 |
| Reporter intensity 1 2 |
| Reporter intensity 2 2 |
| Reporter intensity 3 2 |
| Reporter intensity 4 2 |
| Reporter intensity 5 2 |
| Reporter intensity 6 2 |
| Reporter intensity 7 2 |
| Reporter intensity 8 2 |
| Reporter intensity 9 2 |
| Reporter intensity 10 2 |
| Reporter intensity 11 2 |
| Reporter intensity count 1 2 |
| Reporter intensity count 2 2 |
| Reporter intensity count 3 2 |
| Reporter intensity count 4 2 |
| Reporter intensity count 5 2 |
| Reporter intensity count 6 2 |
| Reporter intensity count 7 2 |
| Reporter intensity count 8 2 |
| Reporter intensity count 9 2 |
| Reporter intensity count 10 2 |
| Reporter intensity count 11 2 |
| Reporter intensity corrected 1 3 |
| Reporter intensity corrected 2 3 |
| Reporter intensity corrected 3 3 |
| Reporter intensity corrected 4 3 |
| Reporter intensity corrected 5 3 |
| Reporter intensity corrected 6 3 |
| Reporter intensity corrected 7 3 |
| Reporter intensity corrected 8 3 |
| Reporter intensity corrected 9 3 |
| Reporter intensity corrected 10 3 |
| Reporter intensity corrected 11 3 |
| Reporter intensity 1 3 |
| Reporter intensity 2 3 |
| Reporter intensity 3 3 |
| Reporter intensity 4 3 |

| | | |
|---|---|---|
| Reporter intensity 5 3 | | |
| Reporter intensity 6 3 | | |
| Reporter intensity 7 3 | | |
| Reporter intensity 8 3 | | |
| Reporter intensity 9 3 | | |
| Reporter intensity 10 3 | | |
| Reporter intensity 11 3 | | |
| Reporter intensity count 1 3 | | |
| Reporter intensity count 2 3 | | |
| Reporter intensity count 3 3 | | |
| Reporter intensity count 4 3 | | |
| Reporter intensity count 5 3 | | |
| Reporter intensity count 6 3 | | |
| Reporter intensity count 7 3 | | |
| Reporter intensity count 8 3 | | |
| Reporter intensity count 9 3 | | |
| Reporter intensity count 10 3 | | |
| Reporter intensity count 11 3 | | |
| Reporter intensity corrected 1 4 | | |
| Reporter intensity corrected 2 4 | | |
| Reporter intensity corrected 3 4 | | |
| Reporter intensity corrected 4 4 | | |
| Reporter intensity corrected 5 4 | | |
| Reporter intensity corrected 6 4 | | |
| Reporter intensity corrected 7 4 | | |
| Reporter intensity corrected 8 4 | | |
| Reporter intensity corrected 9 4 | | |
| Reporter intensity corrected 10 4 | | |
| Reporter intensity corrected 11 4 | | |
| Reporter intensity 1 4 | | |
| Reporter intensity 2 4 | | |
| Reporter intensity 3 4 | | |
| Reporter intensity 4 4 | | |
| Reporter intensity 5 4 | | |
| Reporter intensity 6 4 | | |
| Reporter intensity 7 4 | | |
| Reporter intensity 8 4 | | |
| Reporter intensity 9 4 | | |
| Reporter intensity 10 4 | | |
| Reporter intensity 11 4 | | |
| Reporter intensity count 1 4 | | |
| Reporter intensity count 2 4 | | |
| Reporter intensity count 3 4 | | |
| Reporter intensity count 4 4 | | |
| Reporter intensity count 5 4 | | |
| Reporter intensity count 6 4 | | |
| Reporter intensity count 7 4 | | |
| Reporter intensity count 8 4 | | |
| Reporter intensity count 9 4 | | |
| Reporter intensity count 10 4 | | |
| Reporter intensity count 11 4 | | |
| Intensity | | Summed up eXtracted Ion Current (XIC) of all isotopic clusters associated with the identified AA sequence. In case of a labeled experiment this is the total intensity of all the isotopic patterns in the label cluster. |
| Intensity 1 | | Summed up eXtracted Ion Current (XIC) of all isotopic clusters associated with the identified AA sequence. In case of a labeled experiment this is the total intensity of all the isotopic patterns in the label cluster. |
| Intensity 2 | | Summed up eXtracted Ion Current (XIC) of all isotopic clusters associated with the identified AA sequence. In case of a labeled experiment this is the total intensity of all the isotopic patterns in the label cluster. |
| Intensity 3 | | Summed up eXtracted Ion Current (XIC) of all isotopic clusters associated with the identified AA sequence. In case of a labeled experiment this is the total intensity of all the isotopic patterns in the label cluster. |

| | | |
|---|---|---|
| Intensity 4 | | Summed up eXtracted Ion Current (XIC) of all isotopic clusters associated with the identified AA sequence. In case of a labeled experiment this is the total intensity of all the isotopic patterns in the label cluster. |
| Reverse | | When marked with '+', this particular peptide was found to be part of a protein derived from the reversed part of the decoy database. These should be removed for further data analysis. |
| Potential contaminant | | When marked with '+', this particular peptide was found to be part of a commonly occurring contaminant. These should be removed for further data analysis. |
| id | | A unique (consecutive) identifier for each row in the peptides table, which is used to cross-link the information in this table with the information stored in the other tables. |
| Protein group IDs | | The identifiers of the protein groups this peptide was linked to, referenced against the proteinGroups table. |
| Mod. peptide IDs | | Identifier(s) for peptide sequence(s), associated with the peptide, referenced against the corresponding modified peptides table. |
| Evidence IDs | | Identifier(s) for analyzed peptide evidence associated with the protein group referenced against the evidence table. |
| MS/MS IDs | | The identifiers of the MS/MS scans identifying this peptide, referenced against the msms table. |
| Best MS/MS | | The identifier of the best (in terms of quality) MS/MS scan identifying this peptide, referenced against the msms table. |
| Oxidation (M) site IDs | | Identifier(s) for site(s) associated with the protein group, which show(s) evidence of the modification, referenced against the appropriate modification site file. |
| Phospho (STY) site IDs | | Identifier(s) for site(s) associated with the protein group, which show(s) evidence of the modification, referenced against the appropriate modification site file. |
| Taxonomy IDs | | Taxonomy identifiers. |
| MS/MS Count | | |

### Modification-specific peptides

| Name | Separator | Description |
|---|---|---|
| Sequence | | The identified AA sequence of the peptide. |
| Modifications | | Post-translational modifications contained within the sequence. When no modifications exist, this is set to 'unmodified'. |
| Mass | | Charge corrected mass of the precursor ion. |
| Mass Fractional Part | | The values after the decimal point (ie value - floor(value)). |
| Protein Groups | | IDs of the protein groups to which this peptide belongs. |
| Proteins | | The identifiers of the proteins this particular peptide is associated with. |
| Gene Names | | Names of genes this peptide is associated with. |
| Protein Names | | Names of proteins this peptide is associated with. |
| Unique (Groups) | | When marked with '+', this particular peptide is unique to a single protein group in the proteinGroups file. |
| Unique (Proteins) | | When marked with '+', this particular peptide is unique to a single protein sequence in the fasta file(s). |
| Acetyl (Protein N-term) | | Number of Acetyl (Protein N-term) on this peptide. |
| Oxidation (M) | | Number of Oxidation (M) on this peptide. |
| Phospho (STY) | | Number of Phospho (STY) on this peptide. |
| Missed cleavages | | Number of missed enzymatic cleavages. |
| Experiment 1 | | Number of evidence entries for this 'Experiment'. |
| Experiment 2 | | Number of evidence entries for this 'Experiment'. |
| Experiment 3 | | Number of evidence entries for this 'Experiment'. |
| Experiment 4 | | Number of evidence entries for this 'Experiment'. |
| Retention time | | Retention time in minutes averaged over the evidence entries belonging to this modification-specific peptide. |
| Calibrated retention time | | Calibrated retention time averaged over the evidence entries belonging to this modification-specific peptide. Obviously this only makes sense if retention time recalibration has been performed which is the case when matching between run is selected. |
| Charges | | All charge states that have been observed. |
| PEP | | Posterior Error Probability of the identification. This value essentially operates as a p-value, where smaller is more significant. |
| MS/MS scan number | | The RAW-file derived scan number of the MS/MS with the highest peptide identification score (the highest score is stored in the column 'Score'). |
| Raw file | | The name of the RAW-file the mass spectral data was derived from. |
| Score | | Andromeda score for the best identified among the associated MS/MS spectra. |
| Delta score | | Score difference to the second best identified peptide. |
| Reverse | | When marked with '+', this particular peptide was found to be part of a protein derived from the reversed part of the decoy database. These should be removed for further data analysis. |
| Potential contaminant | | When marked with '+', this particular peptide was found to be part of a commonly occurring contaminant. These should be removed for further data analysis. |
| Reporter intensity corrected 1 | | |
| Reporter intensity corrected 2 | | |
| Reporter intensity corrected 3 | | |
| Reporter intensity corrected 4 | | |
| Reporter intensity corrected 5 | | |
| Reporter intensity corrected 6 | | |
| Reporter intensity corrected 7 | | |
| Reporter intensity corrected 8 | | |
| Reporter intensity corrected 9 | | |
| Reporter intensity corrected 10 | | |
| Reporter intensity corrected 11 | | |
| Reporter intensity 1 | | |
| Reporter intensity 2 | | |
| Reporter intensity 3 | | |
| Reporter intensity 4 | | |
| Reporter intensity 5 | | |
| Reporter intensity 6 | | |

| |
|---|
| Reporter intensity 7 |
| Reporter intensity 8 |
| Reporter intensity 9 |
| Reporter intensity 10 |
| Reporter intensity 11 |
| Reporter intensity count 1 |
| Reporter intensity count 2 |
| Reporter intensity count 3 |
| Reporter intensity count 4 |
| Reporter intensity count 5 |
| Reporter intensity count 6 |
| Reporter intensity count 7 |
| Reporter intensity count 8 |
| Reporter intensity count 9 |
| Reporter intensity count 10 |
| Reporter intensity count 11 |
| Reporter intensity corrected 1 1 |
| Reporter intensity corrected 2 1 |
| Reporter intensity corrected 3 1 |
| Reporter intensity corrected 4 1 |
| Reporter intensity corrected 5 1 |
| Reporter intensity corrected 6 1 |
| Reporter intensity corrected 7 1 |
| Reporter intensity corrected 8 1 |
| Reporter intensity corrected 9 1 |
| Reporter intensity corrected 10 1 |
| Reporter intensity corrected 11 1 |
| Reporter intensity 1 1 |
| Reporter intensity 2 1 |
| Reporter intensity 3 1 |
| Reporter intensity 4 1 |
| Reporter intensity 5 1 |
| Reporter intensity 6 1 |
| Reporter intensity 7 1 |
| Reporter intensity 8 1 |
| Reporter intensity 9 1 |
| Reporter intensity 10 1 |
| Reporter intensity 11 1 |
| Reporter intensity count 1 1 |
| Reporter intensity count 2 1 |
| Reporter intensity count 3 1 |
| Reporter intensity count 4 1 |
| Reporter intensity count 5 1 |
| Reporter intensity count 6 1 |
| Reporter intensity count 7 1 |
| Reporter intensity count 8 1 |
| Reporter intensity count 9 1 |
| Reporter intensity count 10 1 |
| Reporter intensity count 11 1 |
| Reporter intensity corrected 1 2 |
| Reporter intensity corrected 2 2 |
| Reporter intensity corrected 3 2 |
| Reporter intensity corrected 4 2 |
| Reporter intensity corrected 5 2 |
| Reporter intensity corrected 6 2 |
| Reporter intensity corrected 7 2 |
| Reporter intensity corrected 8 2 |
| Reporter intensity corrected 9 2 |
| Reporter intensity corrected 10 2 |
| Reporter intensity corrected 11 2 |
| Reporter intensity 1 2 |
| Reporter intensity 2 2 |
| Reporter intensity 3 2 |
| Reporter intensity 4 2 |

| |
|---|
| Reporter intensity 5 2 |
| Reporter intensity 6 2 |
| Reporter intensity 7 2 |
| Reporter intensity 8 2 |
| Reporter intensity 9 2 |
| Reporter intensity 10 2 |
| Reporter intensity 11 2 |
| Reporter intensity count 1 2 |
| Reporter intensity count 2 2 |
| Reporter intensity count 3 2 |
| Reporter intensity count 4 2 |
| Reporter intensity count 5 2 |
| Reporter intensity count 6 2 |
| Reporter intensity count 7 2 |
| Reporter intensity count 8 2 |
| Reporter intensity count 9 2 |
| Reporter intensity count 10 2 |
| Reporter intensity count 11 2 |
| Reporter intensity corrected 1 3 |
| Reporter intensity corrected 2 3 |
| Reporter intensity corrected 3 3 |
| Reporter intensity corrected 4 3 |
| Reporter intensity corrected 5 3 |
| Reporter intensity corrected 6 3 |
| Reporter intensity corrected 7 3 |
| Reporter intensity corrected 8 3 |
| Reporter intensity corrected 9 3 |
| Reporter intensity corrected 10 3 |
| Reporter intensity corrected 11 3 |
| Reporter intensity 1 3 |
| Reporter intensity 2 3 |
| Reporter intensity 3 3 |
| Reporter intensity 4 3 |
| Reporter intensity 5 3 |
| Reporter intensity 6 3 |
| Reporter intensity 7 3 |
| Reporter intensity 8 3 |
| Reporter intensity 9 3 |
| Reporter intensity 10 3 |
| Reporter intensity 11 3 |
| Reporter intensity count 1 3 |
| Reporter intensity count 2 3 |
| Reporter intensity count 3 3 |
| Reporter intensity count 4 3 |
| Reporter intensity count 5 3 |
| Reporter intensity count 6 3 |
| Reporter intensity count 7 3 |
| Reporter intensity count 8 3 |
| Reporter intensity count 9 3 |
| Reporter intensity count 10 3 |
| Reporter intensity count 11 3 |
| Reporter intensity corrected 1 4 |
| Reporter intensity corrected 2 4 |
| Reporter intensity corrected 3 4 |
| Reporter intensity corrected 4 4 |
| Reporter intensity corrected 5 4 |
| Reporter intensity corrected 6 4 |
| Reporter intensity corrected 7 4 |
| Reporter intensity corrected 8 4 |
| Reporter intensity corrected 9 4 |
| Reporter intensity corrected 10 4 |
| Reporter intensity corrected 11 4 |
| Reporter intensity 1 4 |
| Reporter intensity 2 4 |

| | | |
|---|---|---|
| Reporter intensity 3 4 | | |
| Reporter intensity 4 4 | | |
| Reporter intensity 5 4 | | |
| Reporter intensity 6 4 | | |
| Reporter intensity 7 4 | | |
| Reporter intensity 8 4 | | |
| Reporter intensity 9 4 | | |
| Reporter intensity 10 4 | | |
| Reporter intensity 11 4 | | |
| Reporter intensity count 1 4 | | |
| Reporter intensity count 2 4 | | |
| Reporter intensity count 3 4 | | |
| Reporter intensity count 4 4 | | |
| Reporter intensity count 5 4 | | |
| Reporter intensity count 6 4 | | |
| Reporter intensity count 7 4 | | |
| Reporter intensity count 8 4 | | |
| Reporter intensity count 9 4 | | |
| Reporter intensity count 10 4 | | |
| Reporter intensity count 11 4 | | |
| Intensity | | Summed up eXtracted Ion Current (XIC) of all isotopic clusters associated with the identified AA sequence. In case of a labeled experiment this is the total intensity of all the isotopic patterns in the label cluster. |
| Intensity 1 | | Summed up eXtracted Ion Current (XIC) of all isotopic clusters associated with the identified AA sequence. In case of a labeled experiment this is the total intensity of all the isotopic patterns in the label cluster. |
| Intensity 2 | | Summed up eXtracted Ion Current (XIC) of all isotopic clusters associated with the identified AA sequence. In case of a labeled experiment this is the total intensity of all the isotopic patterns in the label cluster. |
| Intensity 3 | | Summed up eXtracted Ion Current (XIC) of all isotopic clusters associated with the identified AA sequence. In case of a labeled experiment this is the total intensity of all the isotopic patterns in the label cluster. |
| Intensity 4 | | Summed up eXtracted Ion Current (XIC) of all isotopic clusters associated with the identified AA sequence. In case of a labeled experiment this is the total intensity of all the isotopic patterns in the label cluster. |
| id | | A unique (consecutive) identifier for each row in the peptides table, which is used to cross-link the information in this table with the information stored in the other tables. |
| Protein group IDs | | The identifiers of the protein groups this peptide was linked to, referenced against the proteinGroups table. |
| Peptide ID | | Identifier of the associated peptide sequence summary, which can be found in the file 'peptides.txt'. |
| Evidence IDs | | Identifier(s) for analyzed peptide evidence associated with the protein group referenced against the evidence table. |
| MS/MS IDs | | The identifiers of the MS/MS scans identifying this peptide, referenced against the msms table. |
| Best MS/MS | | The identifier of the best (in terms of quality) MS/MS scan identifying this peptide, referenced against the msms table. |
| Oxidation (M) site IDs | | Identifier(s) for site(s) associated with this peptide, which show(s) evidence of the modification, referenced against the appropriate modification site file. |
| Phospho (STY) site IDs | | Identifier(s) for site(s) associated with this peptide, which show(s) evidence of the modification, referenced against the appropriate modification site file. |
| MS/MS Count | | |
| Taxonomy IDs | | Taxonomy identifiers. |

### Oxidation (M)Sites

| **Name** | **Separator** | **Description** |
|---|---|---|
| Proteins | | Identifiers of proteins this site is associated with. |
| Positions within proteins | | For each protein identifier in the 'Proteins' column you find here the position of the site in the respective protein sequence. The index of the first amino acid in the sequence is 1. |
| Leading proteins | | |
| Protein | | Identifier of the protein this peptide is associated with. |
| Protein names | | Names of proteins this peptide is associated with. |
| Gene names | | Names of genes this peptide is associated with. |
| Fasta headers | | Descriptions of proteins this peptide is associated with. |
| Localization prob | | |
| Score diff | | |
| PEP | | The posterior error probability (PEP) of the best identified modified peptide containing this site. |
| Score | | The Andromeda score of the best identified modified peptide containing this site. |
| Delta score | | The Andromeda delta score of the best identified modified peptide containing this site. |
| Score for localization | | The Andromeda score of the MS/MS spectrum used for calculating the localization score for this site. |
| Localization prob 1 | | |
| Score diff 1 | | |
| PEP 1 | | |
| Score 1 | | |
| Localization prob 2 | | |
| Score diff 2 | | |
| PEP 2 | | |
| Score 2 | | |
| Localization prob 3 | | |
| Score diff 3 | | |
| PEP 3 | | |
| Score 3 | | |
| Localization prob 4 | | |
| Score diff 4 | | |
| PEP 4 | | |
| Score 4 | | |
| Diagnostic peak | | |
| Number of Oxidation (M) | | Different numbers of Oxidation (M) on peptides that this site is involved in. |
| Amino acid | | |
| Sequence window | | |
| Modification window | | |
| Peptide window coverage | | |
| Oxidation (M) Probabilities | | |
| Oxidation (M) Score diffs | | |
| Position in peptide | | |
| Charge | | Charge state of the precursor ion. |
| Mass error [ppm] | | Mass error of the recalibrated mass-over-charge value of the precursor ion in comparison to the predicted monoisotopic mass of the identified peptide sequence. |
| Reporter intensity corrected 1___1 | | |
| Reporter intensity corrected 1___2 | | |
| Reporter intensity corrected 1___3 | | |
| Reporter intensity corrected 2___1 | | |
| Reporter intensity corrected 2___2 | | |
| Reporter intensity corrected 2___3 | | |
| Reporter intensity corrected 3___1 | | |
| Reporter intensity corrected 3___2 | | |
| Reporter intensity corrected 3___3 | | |
| Reporter intensity corrected 4___1 | | |
| Reporter intensity corrected 4___2 | | |

| |
|---|
| Reporter intensity corrected 4___3 |
| Reporter intensity corrected 5___1 |
| Reporter intensity corrected 5___2 |
| Reporter intensity corrected 5___3 |
| Reporter intensity corrected 6___1 |
| Reporter intensity corrected 6___2 |
| Reporter intensity corrected 6___3 |
| Reporter intensity corrected 7___1 |
| Reporter intensity corrected 7___2 |
| Reporter intensity corrected 7___3 |
| Reporter intensity corrected 8___1 |
| Reporter intensity corrected 8___2 |
| Reporter intensity corrected 8___3 |
| Reporter intensity corrected 9___1 |
| Reporter intensity corrected 9___2 |
| Reporter intensity corrected 9___3 |
| Reporter intensity corrected 10___1 |
| Reporter intensity corrected 10___2 |
| Reporter intensity corrected 10___3 |
| Reporter intensity corrected 11___1 |
| Reporter intensity corrected 11___2 |
| Reporter intensity corrected 11___3 |
| Reporter intensity 1___1 |
| Reporter intensity 1___2 |
| Reporter intensity 1___3 |
| Reporter intensity 2___1 |
| Reporter intensity 2___2 |
| Reporter intensity 2___3 |
| Reporter intensity 3___1 |
| Reporter intensity 3___2 |
| Reporter intensity 3___3 |
| Reporter intensity 4___1 |
| Reporter intensity 4___2 |
| Reporter intensity 4___3 |
| Reporter intensity 5___1 |
| Reporter intensity 5___2 |
| Reporter intensity 5___3 |
| Reporter intensity 6___1 |
| Reporter intensity 6___2 |
| Reporter intensity 6___3 |
| Reporter intensity 7___1 |
| Reporter intensity 7___2 |
| Reporter intensity 7___3 |
| Reporter intensity 8___1 |
| Reporter intensity 8___2 |
| Reporter intensity 8___3 |
| Reporter intensity 9___1 |
| Reporter intensity 9___2 |
| Reporter intensity 9___3 |
| Reporter intensity 10___1 |
| Reporter intensity 10___2 |
| Reporter intensity 10___3 |
| Reporter intensity 11___1 |
| Reporter intensity 11___2 |
| Reporter intensity 11___3 |
| Reporter intensity count 1___1 |
| Reporter intensity count 1___2 |
| Reporter intensity count 1___3 |
| Reporter intensity count 2___1 |
| Reporter intensity count 2___2 |
| Reporter intensity count 2___3 |
| Reporter intensity count 3___1 |
| Reporter intensity count 3___2 |
| Reporter intensity count 3___3 |

| |
|---|
| Reporter intensity count 4___1 |
| Reporter intensity count 4___2 |
| Reporter intensity count 4___3 |
| Reporter intensity count 5___1 |
| Reporter intensity count 5___2 |
| Reporter intensity count 5___3 |
| Reporter intensity count 6___1 |
| Reporter intensity count 6___2 |
| Reporter intensity count 6___3 |
| Reporter intensity count 7___1 |
| Reporter intensity count 7___2 |
| Reporter intensity count 7___3 |
| Reporter intensity count 8___1 |
| Reporter intensity count 8___2 |
| Reporter intensity count 8___3 |
| Reporter intensity count 9___1 |
| Reporter intensity count 9___2 |
| Reporter intensity count 9___3 |
| Reporter intensity count 10___1 |
| Reporter intensity count 10___2 |
| Reporter intensity count 10___3 |
| Reporter intensity count 11___1 |
| Reporter intensity count 11___2 |
| Reporter intensity count 11___3 |
| Reporter intensity corrected 1 1___1 |
| Reporter intensity corrected 1 1___2 |
| Reporter intensity corrected 1 1___3 |
| Reporter intensity corrected 2 1___1 |
| Reporter intensity corrected 2 1___2 |
| Reporter intensity corrected 2 1___3 |
| Reporter intensity corrected 3 1___1 |
| Reporter intensity corrected 3 1___2 |
| Reporter intensity corrected 3 1___3 |
| Reporter intensity corrected 4 1___1 |
| Reporter intensity corrected 4 1___2 |
| Reporter intensity corrected 4 1___3 |
| Reporter intensity corrected 5 1___1 |
| Reporter intensity corrected 5 1___2 |
| Reporter intensity corrected 5 1___3 |
| Reporter intensity corrected 6 1___1 |
| Reporter intensity corrected 6 1___2 |
| Reporter intensity corrected 6 1___3 |
| Reporter intensity corrected 7 1___1 |
| Reporter intensity corrected 7 1___2 |
| Reporter intensity corrected 7 1___3 |
| Reporter intensity corrected 8 1___1 |
| Reporter intensity corrected 8 1___2 |
| Reporter intensity corrected 8 1___3 |

| |
|---|
| Reporter intensity corrected 9 1___1 |
| Reporter intensity corrected 9 1___2 |
| Reporter intensity corrected 9 1___3 |
| Reporter intensity corrected 10 1___1 |
| Reporter intensity corrected 10 1___2 |
| Reporter intensity corrected 10 1___3 |
| Reporter intensity corrected 11 1___1 |
| Reporter intensity corrected 11 1___2 |
| Reporter intensity corrected 11 1___3 |
| Reporter intensity corrected 1 2___1 |
| Reporter intensity corrected 1 2___2 |
| Reporter intensity corrected 1 2___3 |
| Reporter intensity corrected 2 2___1 |
| Reporter intensity corrected 2 2___2 |
| Reporter intensity corrected 2 2___3 |
| Reporter intensity corrected 3 2___1 |
| Reporter intensity corrected 3 2___2 |
| Reporter intensity corrected 3 2___3 |
| Reporter intensity corrected 4 2___1 |
| Reporter intensity corrected 4 2___2 |
| Reporter intensity corrected 4 2___3 |
| Reporter intensity corrected 5 2___1 |
| Reporter intensity corrected 5 2___2 |
| Reporter intensity corrected 5 2___3 |
| Reporter intensity corrected 6 2___1 |
| Reporter intensity corrected 6 2___2 |
| Reporter intensity corrected 6 2___3 |
| Reporter intensity corrected 7 2___1 |
| Reporter intensity corrected 7 2___2 |
| Reporter intensity corrected 7 2___3 |
| Reporter intensity corrected 8 2___1 |
| Reporter intensity corrected 8 2___2 |
| Reporter intensity corrected 8 2___3 |
| Reporter intensity corrected 9 2___1 |
| Reporter intensity corrected 9 2___2 |
| Reporter intensity corrected 9 2___3 |
| Reporter intensity corrected 10 2___1 |
| Reporter intensity corrected 10 2___2 |

| |
|---|
| Reporter intensity corrected 10 2___3 |
| Reporter intensity corrected 11 2___1 |
| Reporter intensity corrected 11 2___2 |
| Reporter intensity corrected 11 2___3 |
| Reporter intensity corrected 1 3___1 |
| Reporter intensity corrected 1 3___2 |
| Reporter intensity corrected 1 3___3 |
| Reporter intensity corrected 2 3___1 |
| Reporter intensity corrected 2 3___2 |
| Reporter intensity corrected 2 3___3 |
| Reporter intensity corrected 3 3___1 |
| Reporter intensity corrected 3 3___2 |
| Reporter intensity corrected 3 3___3 |
| Reporter intensity corrected 4 3___1 |
| Reporter intensity corrected 4 3___2 |
| Reporter intensity corrected 4 3___3 |
| Reporter intensity corrected 5 3___1 |
| Reporter intensity corrected 5 3___2 |
| Reporter intensity corrected 5 3___3 |
| Reporter intensity corrected 6 3___1 |
| Reporter intensity corrected 6 3___2 |
| Reporter intensity corrected 6 3___3 |
| Reporter intensity corrected 7 3___1 |
| Reporter intensity corrected 7 3___2 |
| Reporter intensity corrected 7 3___3 |
| Reporter intensity corrected 8 3___1 |
| Reporter intensity corrected 8 3___2 |
| Reporter intensity corrected 8 3___3 |
| Reporter intensity corrected 9 3___1 |
| Reporter intensity corrected 9 3___2 |
| Reporter intensity corrected 9 3___3 |
| Reporter intensity corrected 10 3___1 |
| Reporter intensity corrected 10 3___2 |
| Reporter intensity corrected 10 3___3 |
| Reporter intensity corrected 11 3___1 |
| Reporter intensity corrected 11 3___2 |
| Reporter intensity corrected 11 3___3 |
| Reporter intensity corrected 1 4___1 |

| |
|---|
| Reporter intensity corrected 1 4___2 |
| Reporter intensity corrected 1 4___3 |
| Reporter intensity corrected 2 4___1 |
| Reporter intensity corrected 2 4___2 |
| Reporter intensity corrected 2 4___3 |
| Reporter intensity corrected 3 4___1 |
| Reporter intensity corrected 3 4___2 |
| Reporter intensity corrected 3 4___3 |
| Reporter intensity corrected 4 4___1 |
| Reporter intensity corrected 4 4___2 |
| Reporter intensity corrected 4 4___3 |
| Reporter intensity corrected 5 4___1 |
| Reporter intensity corrected 5 4___2 |
| Reporter intensity corrected 5 4___3 |
| Reporter intensity corrected 6 4___1 |
| Reporter intensity corrected 6 4___2 |
| Reporter intensity corrected 6 4___3 |
| Reporter intensity corrected 7 4___1 |
| Reporter intensity corrected 7 4___2 |
| Reporter intensity corrected 7 4___3 |
| Reporter intensity corrected 8 4___1 |
| Reporter intensity corrected 8 4___2 |
| Reporter intensity corrected 8 4___3 |
| Reporter intensity corrected 9 4___1 |
| Reporter intensity corrected 9 4___2 |
| Reporter intensity corrected 9 4___3 |
| Reporter intensity corrected 10 4___1 |
| Reporter intensity corrected 10 4___2 |
| Reporter intensity corrected 10 4___3 |
| Reporter intensity corrected 11 4___1 |
| Reporter intensity corrected 11 4___2 |
| Reporter intensity corrected 11 4___3 |
| Reporter intensity 1 1___1 |
| Reporter intensity 1 1___2 |
| Reporter intensity 1 1___3 |
| Reporter intensity 2 1___1 |
| Reporter intensity 2 1___2 |
| Reporter intensity 2 1___3 |
| Reporter intensity 3 1___1 |
| Reporter intensity 3 1___2 |
| Reporter intensity 3 1___3 |
| Reporter intensity 4 1___1 |

| |
|---|
| Reporter intensity 4 1___2 |
| Reporter intensity 4 1___3 |
| Reporter intensity 5 1___1 |
| Reporter intensity 5 1___2 |
| Reporter intensity 5 1___3 |
| Reporter intensity 6 1___1 |
| Reporter intensity 6 1___2 |
| Reporter intensity 6 1___3 |
| Reporter intensity 7 1___1 |
| Reporter intensity 7 1___2 |
| Reporter intensity 7 1___3 |
| Reporter intensity 8 1___1 |
| Reporter intensity 8 1___2 |
| Reporter intensity 8 1___3 |
| Reporter intensity 9 1___1 |
| Reporter intensity 9 1___2 |
| Reporter intensity 9 1___3 |
| Reporter intensity 10 1___1 |
| Reporter intensity 10 1___2 |
| Reporter intensity 10 1___3 |
| Reporter intensity 11 1___1 |
| Reporter intensity 11 1___2 |
| Reporter intensity 11 1___3 |
| Reporter intensity 1 2___1 |
| Reporter intensity 1 2___2 |
| Reporter intensity 1 2___3 |
| Reporter intensity 2 2___1 |
| Reporter intensity 2 2___2 |
| Reporter intensity 2 2___3 |
| Reporter intensity 3 2___1 |
| Reporter intensity 3 2___2 |
| Reporter intensity 3 2___3 |
| Reporter intensity 4 2___1 |
| Reporter intensity 4 2___2 |
| Reporter intensity 4 2___3 |
| Reporter intensity 5 2___1 |
| Reporter intensity 5 2___2 |
| Reporter intensity 5 2___3 |
| Reporter intensity 6 2___1 |
| Reporter intensity 6 2___2 |
| Reporter intensity 6 2___3 |
| Reporter intensity 7 2___1 |
| Reporter intensity 7 2___2 |
| Reporter intensity 7 2___3 |
| Reporter intensity 8 2___1 |
| Reporter intensity 8 2___2 |
| Reporter intensity 8 2___3 |
| Reporter intensity 9 2___1 |
| Reporter intensity 9 2___2 |
| Reporter intensity 9 2___3 |
| Reporter intensity 10 2___1 |
| Reporter intensity 10 2___2 |
| Reporter intensity 10 2___3 |
| Reporter intensity 11 2___1 |
| Reporter intensity 11 2___2 |
| Reporter intensity 11 2___3 |
| Reporter intensity 1 3___1 |
| Reporter intensity 1 3___2 |
| Reporter intensity 1 3___3 |
| Reporter intensity 2 3___1 |
| Reporter intensity 2 3___2 |
| Reporter intensity 2 3___3 |
| Reporter intensity 3 3___1 |
| Reporter intensity 3 3___2 |

| |
|---|
| Reporter intensity 3 3___3 |
| Reporter intensity 4 3___1 |
| Reporter intensity 4 3___2 |
| Reporter intensity 4 3___3 |
| Reporter intensity 5 3___1 |
| Reporter intensity 5 3___2 |
| Reporter intensity 5 3___3 |
| Reporter intensity 6 3___1 |
| Reporter intensity 6 3___2 |
| Reporter intensity 6 3___3 |
| Reporter intensity 7 3___1 |
| Reporter intensity 7 3___2 |
| Reporter intensity 7 3___3 |
| Reporter intensity 8 3___1 |
| Reporter intensity 8 3___2 |
| Reporter intensity 8 3___3 |
| Reporter intensity 9 3___1 |
| Reporter intensity 9 3___2 |
| Reporter intensity 9 3___3 |
| Reporter intensity 10 3___1 |
| Reporter intensity 10 3___2 |
| Reporter intensity 10 3___3 |
| Reporter intensity 11 3___1 |
| Reporter intensity 11 3___2 |
| Reporter intensity 11 3___3 |
| Reporter intensity 1 4___1 |
| Reporter intensity 1 4___2 |
| Reporter intensity 1 4___3 |
| Reporter intensity 2 4___1 |
| Reporter intensity 2 4___2 |
| Reporter intensity 2 4___3 |
| Reporter intensity 3 4___1 |
| Reporter intensity 3 4___2 |
| Reporter intensity 3 4___3 |
| Reporter intensity 4 4___1 |
| Reporter intensity 4 4___2 |
| Reporter intensity 4 4___3 |
| Reporter intensity 5 4___1 |
| Reporter intensity 5 4___2 |
| Reporter intensity 5 4___3 |
| Reporter intensity 6 4___1 |
| Reporter intensity 6 4___2 |
| Reporter intensity 6 4___3 |
| Reporter intensity 7 4___1 |
| Reporter intensity 7 4___2 |
| Reporter intensity 7 4___3 |
| Reporter intensity 8 4___1 |
| Reporter intensity 8 4___2 |
| Reporter intensity 8 4___3 |
| Reporter intensity 9 4___1 |
| Reporter intensity 9 4___2 |
| Reporter intensity 9 4___3 |
| Reporter intensity 10 4___1 |
| Reporter intensity 10 4___2 |
| Reporter intensity 10 4___3 |
| Reporter intensity 11 4___1 |
| Reporter intensity 11 4___2 |
| Reporter intensity 11 4___3 |
| Reporter intensity count 1 1___1 |
| Reporter intensity count 1 1___2 |
| Reporter intensity count 1 1___3 |
| Reporter intensity count 2 1___1 |
| Reporter intensity count 2 1___2 |
| Reporter intensity count 2 1___3 |

| |
|---|
| Reporter intensity count 3 1___1 |
| Reporter intensity count 3 1___2 |
| Reporter intensity count 3 1___3 |
| Reporter intensity count 4 1___1 |
| Reporter intensity count 4 1___2 |
| Reporter intensity count 4 1___3 |
| Reporter intensity count 5 1___1 |
| Reporter intensity count 5 1___2 |
| Reporter intensity count 5 1___3 |
| Reporter intensity count 6 1___1 |
| Reporter intensity count 6 1___2 |
| Reporter intensity count 6 1___3 |
| Reporter intensity count 7 1___1 |
| Reporter intensity count 7 1___2 |
| Reporter intensity count 7 1___3 |
| Reporter intensity count 8 1___1 |
| Reporter intensity count 8 1___2 |
| Reporter intensity count 8 1___3 |
| Reporter intensity count 9 1___1 |
| Reporter intensity count 9 1___2 |
| Reporter intensity count 9 1___3 |
| Reporter intensity count 10 1___1 |
| Reporter intensity count 10 1___2 |
| Reporter intensity count 10 1___3 |
| Reporter intensity count 11 1___1 |
| Reporter intensity count 11 1___2 |
| Reporter intensity count 11 1___3 |
| Reporter intensity count 1 2___1 |
| Reporter intensity count 1 2___2 |
| Reporter intensity count 1 2___3 |
| Reporter intensity count 2 2___1 |
| Reporter intensity count 2 2___2 |
| Reporter intensity count 2 2___3 |
| Reporter intensity count 3 2___1 |
| Reporter intensity count 3 2___2 |
| Reporter intensity count 3 2___3 |
| Reporter intensity count 4 2___1 |
| Reporter intensity count 4 2___2 |
| Reporter intensity count 4 2___3 |
| Reporter intensity count 5 2___1 |
| Reporter intensity count 5 2___2 |
| Reporter intensity count 5 2___3 |
| Reporter intensity count 6 2___1 |
| Reporter intensity count 6 2___2 |
| Reporter intensity count 6 2___3 |
| Reporter intensity count 7 2___1 |
| Reporter intensity count 7 2___2 |
| Reporter intensity count 7 2___3 |
| Reporter intensity count 8 2___1 |
| Reporter intensity count 8 2___2 |
| Reporter intensity count 8 2___3 |
| Reporter intensity count 9 2___1 |
| Reporter intensity count 9 2___2 |
| Reporter intensity count 9 2___3 |
| Reporter intensity count 10 2___1 |
| Reporter intensity count 10 2___2 |
| Reporter intensity count 10 2___3 |
| Reporter intensity count 11 2___1 |
| Reporter intensity count 11 2___2 |
| Reporter intensity count 11 2___3 |
| Reporter intensity count 1 3___1 |
| Reporter intensity count 1 3___2 |
| Reporter intensity count 1 3___3 |
| Reporter intensity count 2 3___1 |

| |
|---|
| Reporter intensity count 2 3___2 |
| Reporter intensity count 2 3___3 |
| Reporter intensity count 3 3___1 |
| Reporter intensity count 3 3___2 |
| Reporter intensity count 3 3___3 |
| Reporter intensity count 4 3___1 |
| Reporter intensity count 4 3___2 |
| Reporter intensity count 4 3___3 |
| Reporter intensity count 5 3___1 |
| Reporter intensity count 5 3___2 |
| Reporter intensity count 5 3___3 |
| Reporter intensity count 6 3___1 |
| Reporter intensity count 6 3___2 |
| Reporter intensity count 6 3___3 |
| Reporter intensity count 7 3___1 |
| Reporter intensity count 7 3___2 |
| Reporter intensity count 7 3___3 |
| Reporter intensity count 8 3___1 |
| Reporter intensity count 8 3___2 |
| Reporter intensity count 8 3___3 |
| Reporter intensity count 9 3___1 |
| Reporter intensity count 9 3___2 |
| Reporter intensity count 9 3___3 |
| Reporter intensity count 10 3___1 |
| Reporter intensity count 10 3___2 |
| Reporter intensity count 10 3___3 |
| Reporter intensity count 11 3___1 |
| Reporter intensity count 11 3___2 |
| Reporter intensity count 11 3___3 |
| Reporter intensity count 1 4___1 |
| Reporter intensity count 1 4___2 |
| Reporter intensity count 1 4___3 |
| Reporter intensity count 2 4___1 |
| Reporter intensity count 2 4___2 |
| Reporter intensity count 2 4___3 |
| Reporter intensity count 3 4___1 |
| Reporter intensity count 3 4___2 |
| Reporter intensity count 3 4___3 |
| Reporter intensity count 4 4___1 |
| Reporter intensity count 4 4___2 |
| Reporter intensity count 4 4___3 |
| Reporter intensity count 5 4___1 |
| Reporter intensity count 5 4___2 |
| Reporter intensity count 5 4___3 |
| Reporter intensity count 6 4___1 |
| Reporter intensity count 6 4___2 |
| Reporter intensity count 6 4___3 |
| Reporter intensity count 7 4___1 |
| Reporter intensity count 7 4___2 |
| Reporter intensity count 7 4___3 |
| Reporter intensity count 8 4___1 |
| Reporter intensity count 8 4___2 |
| Reporter intensity count 8 4___3 |
| Reporter intensity count 9 4___1 |
| Reporter intensity count 9 4___2 |
| Reporter intensity count 9 4___3 |
| Reporter intensity count 10 4___1 |
| Reporter intensity count 10 4___2 |
| Reporter intensity count 10 4___3 |
| Reporter intensity count 11 4___1 |
| Reporter intensity count 11 4___2 |
| Reporter intensity count 11 4___3 |

| | | |
|---|---|---|
| Intensity | | Summed up eXtracted Ion Current (XIC) of all isotopic clusters associated with the identified AA sequence. In case of a labeled experiment this is the total intensity of all the isotopic patterns in the label cluster. |
| Intensity___1 | | Summed up eXtracted Ion Current (XIC) of all isotopic clusters associated with the identified AA sequence. In case of a labeled experiment this is the total intensity of all the isotopic patterns in the label cluster. |
| Intensity___2 | | Summed up eXtracted Ion Current (XIC) of all isotopic clusters associated with the identified AA sequence. In case of a labeled experiment this is the total intensity of all the isotopic patterns in the label cluster. |
| Intensity___3 | | Summed up eXtracted Ion Current (XIC) of all isotopic clusters associated with the identified AA sequence. In case of a labeled experiment this is the total intensity of all the isotopic patterns in the label cluster. |
| Ratio mod/base | | |
| Intensity 1 | | Summed up eXtracted Ion Current (XIC) of all isotopic clusters associated with the identified AA sequence. In case of a labeled experiment this is the total intensity of all the isotopic patterns in the label cluster. |
| Intensity 2 | | Summed up eXtracted Ion Current (XIC) of all isotopic clusters associated with the identified AA sequence. In case of a labeled experiment this is the total intensity of all the isotopic patterns in the label cluster. |
| Intensity 3 | | Summed up eXtracted Ion Current (XIC) of all isotopic clusters associated with the identified AA sequence. In case of a labeled experiment this is the total intensity of all the isotopic patterns in the label cluster. |
| Intensity 4 | | Summed up eXtracted Ion Current (XIC) of all isotopic clusters associated with the identified AA sequence. In case of a labeled experiment this is the total intensity of all the isotopic patterns in the label cluster. |
| Ratio mod/base 1 | | |
| Ratio mod/base 2 | | |
| Ratio mod/base 3 | | |
| Ratio mod/base 4 | | |
| Intensity 1___1 | | Summed up eXtracted Ion Current (XIC) of all isotopic clusters associated with the identified AA sequence. In case of a labeled experiment this is the total intensity of all the isotopic patterns in the label cluster. |
| Intensity 1___2 | | Summed up eXtracted Ion Current (XIC) of all isotopic clusters associated with the identified AA sequence. In case of a labeled experiment this is the total intensity of all the isotopic patterns in the label cluster. |
| Intensity 1___3 | | Summed up eXtracted Ion Current (XIC) of all isotopic clusters associated with the identified AA sequence. In case of a labeled experiment this is the total intensity of all the isotopic patterns in the label cluster. |
| Intensity 2___1 | | Summed up eXtracted Ion Current (XIC) of all isotopic clusters associated with the identified AA sequence. In case of a labeled experiment this is the total intensity of all the isotopic patterns in the label cluster. |
| Intensity 2___2 | | Summed up eXtracted Ion Current (XIC) of all isotopic clusters associated with the identified AA sequence. In case of a labeled experiment this is the total intensity of all the isotopic patterns in the label cluster. |
| Intensity 2___3 | | Summed up eXtracted Ion Current (XIC) of all isotopic clusters associated with the identified AA sequence. In case of a labeled experiment this is the total intensity of all the isotopic patterns in the label cluster. |
| Intensity 3___1 | | Summed up eXtracted Ion Current (XIC) of all isotopic clusters associated with the identified AA sequence. In case of a labeled experiment this is the total intensity of all the isotopic patterns in the label cluster. |
| Intensity 3___2 | | Summed up eXtracted Ion Current (XIC) of all isotopic clusters associated with the identified AA sequence. In case of a labeled experiment this is the total intensity of all the isotopic patterns in the label cluster. |
| Intensity 3___3 | | Summed up eXtracted Ion Current (XIC) of all isotopic clusters associated with the identified AA sequence. In case of a labeled experiment this is the total intensity of all the isotopic patterns in the label cluster. |
| Intensity 4___1 | | Summed up eXtracted Ion Current (XIC) of all isotopic clusters associated with the identified AA sequence. In case of a labeled experiment this is the total intensity of all the isotopic patterns in the label cluster. |
| Intensity 4___2 | | Summed up eXtracted Ion Current (XIC) of all isotopic clusters associated with the identified AA sequence. In case of a labeled experiment this is the total intensity of all the isotopic patterns in the label cluster. |

| | | |
|---|---|---|
| Intensity 4___3 | | Summed up eXtracted Ion Current (XIC) of all isotopic clusters associated with the identified AA sequence. In case of a labeled experiment this is the total intensity of all the isotopic patterns in the label cluster. |
| Reverse | | When marked with '+', this particular peptide was found to be part of a protein derived from the reversed part of the protein sequence database. These should be removed for further data analysis. |
| Potential contaminant | | When marked with '+', this particular peptide was found to be part of a commonly occurring contaminant. These should be removed for further data analysis. |
| id | | A unique (consecutive) identifier for each row in the site table, which is used to cross-link the information in this file with the information stored in the other files. |
| Protein group IDs | | The identifier of the protein-group this peptide sequence is associated with, which can be used to look up the extended protein information in the file 'proteinGroups.txt'.<br>As a single peptide can be linked to multiple proteins (e.g. in the case of razor-proteins), multiple id's can be stored here separated by a semicolon.<br>As a protein can be identified by multiple peptides, the same id can be found in different rows. |
| Positions | | The positions of the modifications in the protein amino acid sequence. |
| Position | | The position of the modification in the protein amino acid sequence. |
| Peptide IDs | | Identifier(s) of the associated peptide sequence(s) summary, which can be found in the file 'peptides.txt'. |
| Mod. peptide IDs | | Identifier(s) of the associated peptide sequence(s) summary, which can be found in the file 'modificationSpecificPeptides.txt'. |
| Evidence IDs | | Identifier(s) for analyzed peptide evidence associated with the protein group referenced against the evidence table. |
| MS/MS IDs | | The identifiers of the MS/MS scans identifying this peptide, referenced against the msms table. |
| Best localization evidence ID | | |
| Best localization MS/MS ID | | |
| Best localization raw file | | |
| Best localization scan number | | |
| Best score evidence ID | | |
| Best score MS/MS ID | | |
| Best score raw file | | |
| Best score scan number | | |
| Best PEP evidence ID | | |
| Best PEP MS/MS ID | | |
| Best PEP raw file | | |
| Best PEP scan number | | |

### Phospho (STY)Sites

| Name | Separator | Description |
|---|---|---|
| Proteins | | Identifiers of proteins this site is associated with. |
| Positions within proteins | | For each protein identifier in the 'Proteins' column you find here the position of the site in the respective protein sequence. The index of the first amino acid in the sequence is 1. |
| Leading proteins | | |
| Protein | | Identifier of the protein this peptide is associated with. |
| Protein names | | Names of proteins this peptide is associated with. |
| Gene names | | Names of genes this peptide is associated with. |
| Fasta headers | | Descriptions of proteins this peptide is associated with. |
| Localization prob | | |
| Score diff | | |
| PEP | | The posterior error probability (PEP) of the best identified modified peptide containing this site. |
| Score | | The Andromeda score of the best identified modified peptide containing this site. |
| Delta score | | The Andromeda delta score of the best identified modified peptide containing this site. |
| Score for localization | | The Andromeda score of the MS/MS spectrum used for calculating the localization score for this site. |
| Localization prob 1 | | |
| Score diff 1 | | |
| PEP 1 | | |
| Score 1 | | |
| Localization prob 2 | | |
| Score diff 2 | | |
| PEP 2 | | |
| Score 2 | | |
| Localization prob 3 | | |
| Score diff 3 | | |
| PEP 3 | | |
| Score 3 | | |
| Localization prob 4 | | |
| Score diff 4 | | |
| PEP 4 | | |
| Score 4 | | |
| Diagnostic peak | | |
| Number of Phospho (STY) | | Different numbers of Phospho (STY) on peptides that this site is involved in. |
| Amino acid | | |
| Sequence window | | |
| Modification window | | |
| Peptide window coverage | | |
| Phospho (STY) Probabilities | | |
| Phospho (STY) Score diffs | | |
| Position in peptide | | |
| Charge | | Charge state of the precursor ion. |
| Mass error [ppm] | | Mass error of the recalibrated mass-over-charge value of the precursor ion in comparison to the predicted monoisotopic mass of the identified peptide sequence. |
| Reporter intensity corrected 1___1 | | |
| Reporter intensity corrected 1___2 | | |
| Reporter intensity corrected 1___3 | | |
| Reporter intensity corrected 2___1 | | |
| Reporter intensity corrected 2___2 | | |
| Reporter intensity corrected 2___3 | | |
| Reporter intensity corrected 3___1 | | |
| Reporter intensity corrected 3___2 | | |
| Reporter intensity corrected 3___3 | | |
| Reporter intensity corrected 4___1 | | |
| Reporter intensity corrected 4___2 | | |

| |
|---|
| Reporter intensity corrected 4___3 |
| Reporter intensity corrected 5___1 |
| Reporter intensity corrected 5___2 |
| Reporter intensity corrected 5___3 |
| Reporter intensity corrected 6___1 |
| Reporter intensity corrected 6___2 |
| Reporter intensity corrected 6___3 |
| Reporter intensity corrected 7___1 |
| Reporter intensity corrected 7___2 |
| Reporter intensity corrected 7___3 |
| Reporter intensity corrected 8___1 |
| Reporter intensity corrected 8___2 |
| Reporter intensity corrected 8___3 |
| Reporter intensity corrected 9___1 |
| Reporter intensity corrected 9___2 |
| Reporter intensity corrected 9___3 |
| Reporter intensity corrected 10___1 |
| Reporter intensity corrected 10___2 |
| Reporter intensity corrected 10___3 |
| Reporter intensity corrected 11___1 |
| Reporter intensity corrected 11___2 |
| Reporter intensity corrected 11___3 |
| Reporter intensity 1___1 |
| Reporter intensity 1___2 |
| Reporter intensity 1___3 |
| Reporter intensity 2___1 |
| Reporter intensity 2___2 |
| Reporter intensity 2___3 |
| Reporter intensity 3___1 |
| Reporter intensity 3___2 |
| Reporter intensity 3___3 |
| Reporter intensity 4___1 |
| Reporter intensity 4___2 |
| Reporter intensity 4___3 |
| Reporter intensity 5___1 |
| Reporter intensity 5___2 |
| Reporter intensity 5___3 |
| Reporter intensity 6___1 |
| Reporter intensity 6___2 |
| Reporter intensity 6___3 |
| Reporter intensity 7___1 |
| Reporter intensity 7___2 |
| Reporter intensity 7___3 |
| Reporter intensity 8___1 |
| Reporter intensity 8___2 |
| Reporter intensity 8___3 |
| Reporter intensity 9___1 |
| Reporter intensity 9___2 |
| Reporter intensity 9___3 |
| Reporter intensity 10___1 |
| Reporter intensity 10___2 |
| Reporter intensity 10___3 |
| Reporter intensity 11___1 |
| Reporter intensity 11___2 |
| Reporter intensity 11___3 |
| Reporter intensity count 1___1 |
| Reporter intensity count 1___2 |
| Reporter intensity count 1___3 |
| Reporter intensity count 2___1 |
| Reporter intensity count 2___2 |
| Reporter intensity count 2___3 |
| Reporter intensity count 3___1 |
| Reporter intensity count 3___2 |
| Reporter intensity count 3___3 |

| |
|---|
| Reporter intensity count 4___1 |
| Reporter intensity count 4___2 |
| Reporter intensity count 4___3 |
| Reporter intensity count 5___1 |
| Reporter intensity count 5___2 |
| Reporter intensity count 5___3 |
| Reporter intensity count 6___1 |
| Reporter intensity count 6___2 |
| Reporter intensity count 6___3 |
| Reporter intensity count 7___1 |
| Reporter intensity count 7___2 |
| Reporter intensity count 7___3 |
| Reporter intensity count 8___1 |
| Reporter intensity count 8___2 |
| Reporter intensity count 8___3 |
| Reporter intensity count 9___1 |
| Reporter intensity count 9___2 |
| Reporter intensity count 9___3 |
| Reporter intensity count 10___1 |
| Reporter intensity count 10___2 |
| Reporter intensity count 10___3 |
| Reporter intensity count 11___1 |
| Reporter intensity count 11___2 |
| Reporter intensity count 11___3 |
| Reporter intensity corrected 1 1___1 |
| Reporter intensity corrected 1 1___2 |
| Reporter intensity corrected 1 1___3 |
| Reporter intensity corrected 2 1___1 |
| Reporter intensity corrected 2 1___2 |
| Reporter intensity corrected 2 1___3 |
| Reporter intensity corrected 3 1___1 |
| Reporter intensity corrected 3 1___2 |
| Reporter intensity corrected 3 1___3 |
| Reporter intensity corrected 4 1___1 |
| Reporter intensity corrected 4 1___2 |
| Reporter intensity corrected 4 1___3 |
| Reporter intensity corrected 5 1___1 |
| Reporter intensity corrected 5 1___2 |
| Reporter intensity corrected 5 1___3 |
| Reporter intensity corrected 6 1___1 |
| Reporter intensity corrected 6 1___2 |
| Reporter intensity corrected 6 1___3 |
| Reporter intensity corrected 7 1___1 |
| Reporter intensity corrected 7 1___2 |
| Reporter intensity corrected 7 1___3 |
| Reporter intensity corrected 8 1___1 |
| Reporter intensity corrected 8 1___2 |
| Reporter intensity corrected 8 1___3 |

| |
|---|
| Reporter intensity corrected 9 1___1 |
| Reporter intensity corrected 9 1___2 |
| Reporter intensity corrected 9 1___3 |
| Reporter intensity corrected 10 1___1 |
| Reporter intensity corrected 10 1___2 |
| Reporter intensity corrected 10 1___3 |
| Reporter intensity corrected 11 1___1 |
| Reporter intensity corrected 11 1___2 |
| Reporter intensity corrected 11 1___3 |
| Reporter intensity corrected 1 2___1 |
| Reporter intensity corrected 1 2___2 |
| Reporter intensity corrected 1 2___3 |
| Reporter intensity corrected 2 2___1 |
| Reporter intensity corrected 2 2___2 |
| Reporter intensity corrected 2 2___3 |
| Reporter intensity corrected 3 2___1 |
| Reporter intensity corrected 3 2___2 |
| Reporter intensity corrected 3 2___3 |
| Reporter intensity corrected 4 2___1 |
| Reporter intensity corrected 4 2___2 |
| Reporter intensity corrected 4 2___3 |
| Reporter intensity corrected 5 2___1 |
| Reporter intensity corrected 5 2___2 |
| Reporter intensity corrected 5 2___3 |
| Reporter intensity corrected 6 2___1 |
| Reporter intensity corrected 6 2___2 |
| Reporter intensity corrected 6 2___3 |
| Reporter intensity corrected 7 2___1 |
| Reporter intensity corrected 7 2___2 |
| Reporter intensity corrected 7 2___3 |
| Reporter intensity corrected 8 2___1 |
| Reporter intensity corrected 8 2___2 |
| Reporter intensity corrected 8 2___3 |
| Reporter intensity corrected 9 2___1 |
| Reporter intensity corrected 9 2___2 |
| Reporter intensity corrected 9 2___3 |
| Reporter intensity corrected 10 2___1 |
| Reporter intensity corrected 10 2___2 |

| |
|---|
| Reporter intensity corrected 10 2___3 |
| Reporter intensity corrected 11 2___1 |
| Reporter intensity corrected 11 2___2 |
| Reporter intensity corrected 11 2___3 |
| Reporter intensity corrected 1 3___1 |
| Reporter intensity corrected 1 3___2 |
| Reporter intensity corrected 1 3___3 |
| Reporter intensity corrected 2 3___1 |
| Reporter intensity corrected 2 3___2 |
| Reporter intensity corrected 2 3___3 |
| Reporter intensity corrected 3 3___1 |
| Reporter intensity corrected 3 3___2 |
| Reporter intensity corrected 3 3___3 |
| Reporter intensity corrected 4 3___1 |
| Reporter intensity corrected 4 3___2 |
| Reporter intensity corrected 4 3___3 |
| Reporter intensity corrected 5 3___1 |
| Reporter intensity corrected 5 3___2 |
| Reporter intensity corrected 5 3___3 |
| Reporter intensity corrected 6 3___1 |
| Reporter intensity corrected 6 3___2 |
| Reporter intensity corrected 6 3___3 |
| Reporter intensity corrected 7 3___1 |
| Reporter intensity corrected 7 3___2 |
| Reporter intensity corrected 7 3___3 |
| Reporter intensity corrected 8 3___1 |
| Reporter intensity corrected 8 3___2 |
| Reporter intensity corrected 8 3___3 |
| Reporter intensity corrected 9 3___1 |
| Reporter intensity corrected 9 3___2 |
| Reporter intensity corrected 9 3___3 |
| Reporter intensity corrected 10 3___1 |
| Reporter intensity corrected 10 3___2 |
| Reporter intensity corrected 10 3___3 |
| Reporter intensity corrected 11 3___1 |
| Reporter intensity corrected 11 3___2 |
| Reporter intensity corrected 11 3___3 |
| Reporter intensity corrected 1 4___1 |

| |
|---|
| Reporter intensity corrected 1 4___2 |
| Reporter intensity corrected 1 4___3 |
| Reporter intensity corrected 2 4___1 |
| Reporter intensity corrected 2 4___2 |
| Reporter intensity corrected 2 4___3 |
| Reporter intensity corrected 3 4___1 |
| Reporter intensity corrected 3 4___2 |
| Reporter intensity corrected 3 4___3 |
| Reporter intensity corrected 4 4___1 |
| Reporter intensity corrected 4 4___2 |
| Reporter intensity corrected 4 4___3 |
| Reporter intensity corrected 5 4___1 |
| Reporter intensity corrected 5 4___2 |
| Reporter intensity corrected 5 4___3 |
| Reporter intensity corrected 6 4___1 |
| Reporter intensity corrected 6 4___2 |
| Reporter intensity corrected 6 4___3 |
| Reporter intensity corrected 7 4___1 |
| Reporter intensity corrected 7 4___2 |
| Reporter intensity corrected 7 4___3 |
| Reporter intensity corrected 8 4___1 |
| Reporter intensity corrected 8 4___2 |
| Reporter intensity corrected 8 4___3 |
| Reporter intensity corrected 9 4___1 |
| Reporter intensity corrected 9 4___2 |
| Reporter intensity corrected 9 4___3 |
| Reporter intensity corrected 10 4___1 |
| Reporter intensity corrected 10 4___2 |
| Reporter intensity corrected 10 4___3 |
| Reporter intensity corrected 11 4___1 |
| Reporter intensity corrected 11 4___2 |
| Reporter intensity corrected 11 4___3 |
| Reporter intensity 1 1___1 |
| Reporter intensity 1 1___2 |
| Reporter intensity 1 1___3 |
| Reporter intensity 2 1___1 |
| Reporter intensity 2 1___2 |
| Reporter intensity 2 1___3 |
| Reporter intensity 3 1___1 |
| Reporter intensity 3 1___2 |
| Reporter intensity 3 1___3 |
| Reporter intensity 4 1___1 |

| |
|---|
| Reporter intensity 4 1___2 |
| Reporter intensity 4 1___3 |
| Reporter intensity 5 1___1 |
| Reporter intensity 5 1___2 |
| Reporter intensity 5 1___3 |
| Reporter intensity 6 1___1 |
| Reporter intensity 6 1___2 |
| Reporter intensity 6 1___3 |
| Reporter intensity 7 1___1 |
| Reporter intensity 7 1___2 |
| Reporter intensity 7 1___3 |
| Reporter intensity 8 1___1 |
| Reporter intensity 8 1___2 |
| Reporter intensity 8 1___3 |
| Reporter intensity 9 1___1 |
| Reporter intensity 9 1___2 |
| Reporter intensity 9 1___3 |
| Reporter intensity 10 1___1 |
| Reporter intensity 10 1___2 |
| Reporter intensity 10 1___3 |
| Reporter intensity 11 1___1 |
| Reporter intensity 11 1___2 |
| Reporter intensity 11 1___3 |
| Reporter intensity 1 2___1 |
| Reporter intensity 1 2___2 |
| Reporter intensity 1 2___3 |
| Reporter intensity 2 2___1 |
| Reporter intensity 2 2___2 |
| Reporter intensity 2 2___3 |
| Reporter intensity 3 2___1 |
| Reporter intensity 3 2___2 |
| Reporter intensity 3 2___3 |
| Reporter intensity 4 2___1 |
| Reporter intensity 4 2___2 |
| Reporter intensity 4 2___3 |
| Reporter intensity 5 2___1 |
| Reporter intensity 5 2___2 |
| Reporter intensity 5 2___3 |
| Reporter intensity 6 2___1 |
| Reporter intensity 6 2___2 |
| Reporter intensity 6 2___3 |
| Reporter intensity 7 2___1 |
| Reporter intensity 7 2___2 |
| Reporter intensity 7 2___3 |
| Reporter intensity 8 2___1 |
| Reporter intensity 8 2___2 |
| Reporter intensity 8 2___3 |
| Reporter intensity 9 2___1 |
| Reporter intensity 9 2___2 |
| Reporter intensity 9 2___3 |
| Reporter intensity 10 2___1 |
| Reporter intensity 10 2___2 |
| Reporter intensity 10 2___3 |
| Reporter intensity 11 2___1 |
| Reporter intensity 11 2___2 |
| Reporter intensity 11 2___3 |
| Reporter intensity 1 3___1 |
| Reporter intensity 1 3___2 |
| Reporter intensity 1 3___3 |
| Reporter intensity 2 3___1 |
| Reporter intensity 2 3___2 |
| Reporter intensity 2 3___3 |
| Reporter intensity 3 3___1 |
| Reporter intensity 3 3___2 |

| |
|---|
| Reporter intensity 3 3___3 |
| Reporter intensity 4 3___1 |
| Reporter intensity 4 3___2 |
| Reporter intensity 4 3___3 |
| Reporter intensity 5 3___1 |
| Reporter intensity 5 3___2 |
| Reporter intensity 5 3___3 |
| Reporter intensity 6 3___1 |
| Reporter intensity 6 3___2 |
| Reporter intensity 6 3___3 |
| Reporter intensity 7 3___1 |
| Reporter intensity 7 3___2 |
| Reporter intensity 7 3___3 |
| Reporter intensity 8 3___1 |
| Reporter intensity 8 3___2 |
| Reporter intensity 8 3___3 |
| Reporter intensity 9 3___1 |
| Reporter intensity 9 3___2 |
| Reporter intensity 9 3___3 |
| Reporter intensity 10 3___1 |
| Reporter intensity 10 3___2 |
| Reporter intensity 10 3___3 |
| Reporter intensity 11 3___1 |
| Reporter intensity 11 3___2 |
| Reporter intensity 11 3___3 |
| Reporter intensity 1 4___1 |
| Reporter intensity 1 4___2 |
| Reporter intensity 1 4___3 |
| Reporter intensity 2 4___1 |
| Reporter intensity 2 4___2 |
| Reporter intensity 2 4___3 |
| Reporter intensity 3 4___1 |
| Reporter intensity 3 4___2 |
| Reporter intensity 3 4___3 |
| Reporter intensity 4 4___1 |
| Reporter intensity 4 4___2 |
| Reporter intensity 4 4___3 |
| Reporter intensity 5 4___1 |
| Reporter intensity 5 4___2 |
| Reporter intensity 5 4___3 |
| Reporter intensity 6 4___1 |
| Reporter intensity 6 4___2 |
| Reporter intensity 6 4___3 |
| Reporter intensity 7 4___1 |
| Reporter intensity 7 4___2 |
| Reporter intensity 7 4___3 |
| Reporter intensity 8 4___1 |
| Reporter intensity 8 4___2 |
| Reporter intensity 8 4___3 |
| Reporter intensity 9 4___1 |
| Reporter intensity 9 4___2 |
| Reporter intensity 9 4___3 |
| Reporter intensity 10 4___1 |
| Reporter intensity 10 4___2 |
| Reporter intensity 10 4___3 |
| Reporter intensity 11 4___1 |
| Reporter intensity 11 4___2 |
| Reporter intensity 11 4___3 |
| Reporter intensity count 1 1___1 |
| Reporter intensity count 1 1___2 |
| Reporter intensity count 1 1___3 |
| Reporter intensity count 2 1___1 |
| Reporter intensity count 2 1___2 |
| Reporter intensity count 2 1___3 |

| |
|---|
| Reporter intensity count 3 1___1 |
| Reporter intensity count 3 1___2 |
| Reporter intensity count 3 1___3 |
| Reporter intensity count 4 1___1 |
| Reporter intensity count 4 1___2 |
| Reporter intensity count 4 1___3 |
| Reporter intensity count 5 1___1 |
| Reporter intensity count 5 1___2 |
| Reporter intensity count 5 1___3 |
| Reporter intensity count 6 1___1 |
| Reporter intensity count 6 1___2 |
| Reporter intensity count 6 1___3 |
| Reporter intensity count 7 1___1 |
| Reporter intensity count 7 1___2 |
| Reporter intensity count 7 1___3 |
| Reporter intensity count 8 1___1 |
| Reporter intensity count 8 1___2 |
| Reporter intensity count 8 1___3 |
| Reporter intensity count 9 1___1 |
| Reporter intensity count 9 1___2 |
| Reporter intensity count 9 1___3 |
| Reporter intensity count 10 1___1 |
| Reporter intensity count 10 1___2 |
| Reporter intensity count 10 1___3 |
| Reporter intensity count 11 1___1 |
| Reporter intensity count 11 1___2 |
| Reporter intensity count 11 1___3 |
| Reporter intensity count 1 2___1 |
| Reporter intensity count 1 2___2 |
| Reporter intensity count 1 2___3 |
| Reporter intensity count 2 2___1 |
| Reporter intensity count 2 2___2 |
| Reporter intensity count 2 2___3 |
| Reporter intensity count 3 2___1 |
| Reporter intensity count 3 2___2 |
| Reporter intensity count 3 2___3 |
| Reporter intensity count 4 2___1 |
| Reporter intensity count 4 2___2 |
| Reporter intensity count 4 2___3 |
| Reporter intensity count 5 2___1 |
| Reporter intensity count 5 2___2 |
| Reporter intensity count 5 2___3 |
| Reporter intensity count 6 2___1 |
| Reporter intensity count 6 2___2 |
| Reporter intensity count 6 2___3 |
| Reporter intensity count 7 2___1 |
| Reporter intensity count 7 2___2 |
| Reporter intensity count 7 2___3 |
| Reporter intensity count 8 2___1 |
| Reporter intensity count 8 2___2 |
| Reporter intensity count 8 2___3 |
| Reporter intensity count 9 2___1 |
| Reporter intensity count 9 2___2 |
| Reporter intensity count 9 2___3 |
| Reporter intensity count 10 2___1 |
| Reporter intensity count 10 2___2 |
| Reporter intensity count 10 2___3 |
| Reporter intensity count 11 2___1 |
| Reporter intensity count 11 2___2 |
| Reporter intensity count 11 2___3 |
| Reporter intensity count 1 3___1 |
| Reporter intensity count 1 3___2 |
| Reporter intensity count 1 3___3 |
| Reporter intensity count 2 3___1 |

| |
|---|
| Reporter intensity count 2 3___2 |
| Reporter intensity count 2 3___3 |
| Reporter intensity count 3 3___1 |
| Reporter intensity count 3 3___2 |
| Reporter intensity count 3 3___3 |
| Reporter intensity count 4 3___1 |
| Reporter intensity count 4 3___2 |
| Reporter intensity count 4 3___3 |
| Reporter intensity count 5 3___1 |
| Reporter intensity count 5 3___2 |
| Reporter intensity count 5 3___3 |
| Reporter intensity count 6 3___1 |
| Reporter intensity count 6 3___2 |
| Reporter intensity count 6 3___3 |
| Reporter intensity count 7 3___1 |
| Reporter intensity count 7 3___2 |
| Reporter intensity count 7 3___3 |
| Reporter intensity count 8 3___1 |
| Reporter intensity count 8 3___2 |
| Reporter intensity count 8 3___3 |
| Reporter intensity count 9 3___1 |
| Reporter intensity count 9 3___2 |
| Reporter intensity count 9 3___3 |
| Reporter intensity count 10 3___1 |
| Reporter intensity count 10 3___2 |
| Reporter intensity count 10 3___3 |
| Reporter intensity count 11 3___1 |
| Reporter intensity count 11 3___2 |
| Reporter intensity count 11 3___3 |
| Reporter intensity count 1 4___1 |
| Reporter intensity count 1 4___2 |
| Reporter intensity count 1 4___3 |
| Reporter intensity count 2 4___1 |
| Reporter intensity count 2 4___2 |
| Reporter intensity count 2 4___3 |
| Reporter intensity count 3 4___1 |
| Reporter intensity count 3 4___2 |
| Reporter intensity count 3 4___3 |
| Reporter intensity count 4 4___1 |
| Reporter intensity count 4 4___2 |
| Reporter intensity count 4 4___3 |
| Reporter intensity count 5 4___1 |
| Reporter intensity count 5 4___2 |
| Reporter intensity count 5 4___3 |
| Reporter intensity count 6 4___1 |
| Reporter intensity count 6 4___2 |
| Reporter intensity count 6 4___3 |
| Reporter intensity count 7 4___1 |
| Reporter intensity count 7 4___2 |
| Reporter intensity count 7 4___3 |
| Reporter intensity count 8 4___1 |
| Reporter intensity count 8 4___2 |
| Reporter intensity count 8 4___3 |
| Reporter intensity count 9 4___1 |
| Reporter intensity count 9 4___2 |
| Reporter intensity count 9 4___3 |
| Reporter intensity count 10 4___1 |
| Reporter intensity count 10 4___2 |
| Reporter intensity count 10 4___3 |
| Reporter intensity count 11 4___1 |
| Reporter intensity count 11 4___2 |
| Reporter intensity count 11 4___3 |

| | | |
|---|---|---|
| Intensity | | Summed up eXtracted Ion Current (XIC) of all isotopic clusters associated with the identified AA sequence. In case of a labeled experiment this is the total intensity of all the isotopic patterns in the label cluster. |
| Intensity___1 | | Summed up eXtracted Ion Current (XIC) of all isotopic clusters associated with the identified AA sequence. In case of a labeled experiment this is the total intensity of all the isotopic patterns in the label cluster. |
| Intensity___2 | | Summed up eXtracted Ion Current (XIC) of all isotopic clusters associated with the identified AA sequence. In case of a labeled experiment this is the total intensity of all the isotopic patterns in the label cluster. |
| Intensity___3 | | Summed up eXtracted Ion Current (XIC) of all isotopic clusters associated with the identified AA sequence. In case of a labeled experiment this is the total intensity of all the isotopic patterns in the label cluster. |
| Ratio mod/base | | |
| Intensity 1 | | Summed up eXtracted Ion Current (XIC) of all isotopic clusters associated with the identified AA sequence. In case of a labeled experiment this is the total intensity of all the isotopic patterns in the label cluster. |
| Intensity 2 | | Summed up eXtracted Ion Current (XIC) of all isotopic clusters associated with the identified AA sequence. In case of a labeled experiment this is the total intensity of all the isotopic patterns in the label cluster. |
| Intensity 3 | | Summed up eXtracted Ion Current (XIC) of all isotopic clusters associated with the identified AA sequence. In case of a labeled experiment this is the total intensity of all the isotopic patterns in the label cluster. |
| Intensity 4 | | Summed up eXtracted Ion Current (XIC) of all isotopic clusters associated with the identified AA sequence. In case of a labeled experiment this is the total intensity of all the isotopic patterns in the label cluster. |
| Ratio mod/base 1 | | |
| Ratio mod/base 2 | | |
| Ratio mod/base 3 | | |
| Ratio mod/base 4 | | |
| Intensity 1___1 | | Summed up eXtracted Ion Current (XIC) of all isotopic clusters associated with the identified AA sequence. In case of a labeled experiment this is the total intensity of all the isotopic patterns in the label cluster. |
| Intensity 1___2 | | Summed up eXtracted Ion Current (XIC) of all isotopic clusters associated with the identified AA sequence. In case of a labeled experiment this is the total intensity of all the isotopic patterns in the label cluster. |
| Intensity 1___3 | | Summed up eXtracted Ion Current (XIC) of all isotopic clusters associated with the identified AA sequence. In case of a labeled experiment this is the total intensity of all the isotopic patterns in the label cluster. |
| Intensity 2___1 | | Summed up eXtracted Ion Current (XIC) of all isotopic clusters associated with the identified AA sequence. In case of a labeled experiment this is the total intensity of all the isotopic patterns in the label cluster. |
| Intensity 2___2 | | Summed up eXtracted Ion Current (XIC) of all isotopic clusters associated with the identified AA sequence. In case of a labeled experiment this is the total intensity of all the isotopic patterns in the label cluster. |
| Intensity 2___3 | | Summed up eXtracted Ion Current (XIC) of all isotopic clusters associated with the identified AA sequence. In case of a labeled experiment this is the total intensity of all the isotopic patterns in the label cluster. |
| Intensity 3___1 | | Summed up eXtracted Ion Current (XIC) of all isotopic clusters associated with the identified AA sequence. In case of a labeled experiment this is the total intensity of all the isotopic patterns in the label cluster. |
| Intensity 3___2 | | Summed up eXtracted Ion Current (XIC) of all isotopic clusters associated with the identified AA sequence. In case of a labeled experiment this is the total intensity of all the isotopic patterns in the label cluster. |
| Intensity 3___3 | | Summed up eXtracted Ion Current (XIC) of all isotopic clusters associated with the identified AA sequence. In case of a labeled experiment this is the total intensity of all the isotopic patterns in the label cluster. |
| Intensity 4___1 | | Summed up eXtracted Ion Current (XIC) of all isotopic clusters associated with the identified AA sequence. In case of a labeled experiment this is the total intensity of all the isotopic patterns in the label cluster. |
| Intensity 4___2 | | Summed up eXtracted Ion Current (XIC) of all isotopic clusters associated with the identified AA sequence. In case of a labeled experiment this is the total intensity of all the isotopic patterns in the label cluster. |

| | | |
|---|---|---|
| Intensity 4___3 | | Summed up eXtracted Ion Current (XIC) of all isotopic clusters associated with the identified AA sequence. In case of a labeled experiment this is the total intensity of all the isotopic patterns in the label cluster. |
| Occupancy 1 | | |
| Occupancy ratio1 | | |
| Occupancy error scale 1 | | |
| Occupancy 2 | | |
| Occupancy ratio2 | | |
| Occupancy error scale 2 | | |
| Occupancy 3 | | |
| Occupancy ratio3 | | |
| Occupancy error scale 3 | | |
| Occupancy 4 | | |
| Occupancy ratio4 | | |
| Occupancy error scale 4 | | |
| Reverse | | When marked with '+', this particular peptide was found to be part of a protein derived from the reversed part of the protein sequence database. These should be removed for further data analysis. |
| Potential contaminant | | When marked with '+', this particular peptide was found to be part of a commonly occurring contaminant. These should be removed for further data analysis. |
| id | | A unique (consecutive) identifier for each row in the site table, which is used to cross-link the information in this file with the information stored in the other files. |
| Protein group IDs | | The identifier of the protein-group this peptide sequence is associated with, which can be used to look up the extended protein information in the file 'proteinGroups.txt'.<br>As a single peptide can be linked to multiple proteins (e.g. in the case of razor-proteins), multiple id’s can be stored here separated by a semicolon.<br>As a protein can be identified by multiple peptides, the same id can be found in different rows. |
| Positions | | The positions of the modifications in the protein amino acid sequence. |
| Position | | The position of the modification in the protein amino acid sequence. |
| Peptide IDs | | Identifier(s) of the associated peptide sequence(s) summary, which can be found in the file 'peptides.txt'. |
| Mod. peptide IDs | | Identifier(s) of the associated peptide sequence(s) summary, which can be found in the file 'modificationSpecificPeptides.txt'. |
| Evidence IDs | | Identifier(s) for analyzed peptide evidence associated with the protein group referenced against the evidence table. |
| MS/MS IDs | | The identifiers of the MS/MS scans identifying this peptide, referenced against the msms table. |
| Best localization evidence ID | | |
| Best localization MS/MS ID | | |
| Best localization raw file | | |
| Best localization scan number | | |
| Best score evidence ID | | |
| Best score MS/MS ID | | |
| Best score raw file | | |
| Best score scan number | | |
| Best PEP evidence ID | | |
| Best PEP MS/MS ID | | |
| Best PEP raw file | | |
| Best PEP scan number | | |

### Protein groups

The Protein Groups table contains information on the identified proteins in the processed raw-files. Each single row contains the group of proteins that could be reconstructed from a set of peptides.

| Name | Separator | Description |
|---|---|---|
| Protein IDs | | Identifiers of proteins contained in the protein group. They are sorted by number of identified peptides in descending order. |
| Majority protein IDs | | These are the IDs of those proteins that have at least half of the peptides that the leading protein has. |
| Peptide counts (all) | | Number of peptides associated with each protein in protein group, occurring in the order as the protein IDs occur in the 'Protein IDs' column. Here distinct peptide sequences are counted. Modified forms or different charges are counted as one peptide. |
| Peptide counts (razor+unique) | | Number of peptides associated with each protein in protein group, occurring in the order as the protein IDs occur in the 'Protein IDs' column. Here distinct peptide sequences are counted. Modified forms or different charges are counted as one peptide. |
| Peptide counts (unique) | | Number of peptides associated with each protein in protein group, occurring in the order as the protein IDs occur in the 'Protein IDs' column. Here distinct peptide sequences are counted. Modified forms or different charges are counted as one peptide. |
| Protein names | | Names of proteins contained within the group. |
| Gene names | | Names of the genes associated to the proteins contained within the group. |
| Fasta headers | | Fasta headers(s) of protein(s) contained within the group. |
| Number of proteins | | Number of proteins contained within the group. This corresponds to the number of entries in the column 'Protein IDs'. |
| Peptides | | The total number of peptide sequences associated with the protein group (i.e. for all the proteins in the group). |
| Razor + unique peptides | | The total number of razor + unique peptides associated with the protein group (i.e. these peptides are shared with another protein group). |
| Unique peptides | | The total number of unique peptides associated with the protein group (i.e. these peptides are not shared with another protein group). |
| Peptides 1 | | Number of peptides (distinct peptide sequences) in experiment 1 |
| Peptides 2 | | Number of peptides (distinct peptide sequences) in experiment 2 |
| Peptides 3 | | Number of peptides (distinct peptide sequences) in experiment 3 |
| Peptides 4 | | Number of peptides (distinct peptide sequences) in experiment 4 |
| Razor + unique peptides 1 | | Number of razor + unique peptides (distinct peptide sequences) in experiment 1 |
| Razor + unique peptides 2 | | Number of razor + unique peptides (distinct peptide sequences) in experiment 2 |
| Razor + unique peptides 3 | | Number of razor + unique peptides (distinct peptide sequences) in experiment 3 |
| Razor + unique peptides 4 | | Number of razor + unique peptides (distinct peptide sequences) in experiment 4 |
| Unique peptides 1 | | Number of unique peptides (distinct peptide sequences) in experiment 1 |
| Unique peptides 2 | | Number of unique peptides (distinct peptide sequences) in experiment 2 |
| Unique peptides 3 | | Number of unique peptides (distinct peptide sequences) in experiment 3 |
| Unique peptides 4 | | Number of unique peptides (distinct peptide sequences) in experiment 4 |
| Sequence coverage [%] | | Percentage of the sequence that is covered by the identified peptides of the best protein sequence contained in the group. |
| Unique + razor sequence coverage [%] | | Percentage of the sequence that is covered by the identified unique and razor peptides of the best protein sequence contained in the group. |
| Unique sequence coverage [%] | | Percentage of the sequence that is covered by the identified unique peptides of the best protein sequence contained in the group. |
| Mol. weight [kDa] | | Molecular weight of the leading protein sequence contained in the protein group. |

| | | |
|---|---|---|
| Sequence length | | The length of the leading protein sequence contained in the group. |
| Sequence lengths | | The length of all sequences of the proteins contained in the group. |
| Q-value | | This is the ratio of reverse to forward protein groups. |
| Score | | Protein score which is derived from peptide posterior error probabilities. |
| Reporter intensity corrected 1 1 | | |
| Reporter intensity corrected 2 1 | | |
| Reporter intensity corrected 3 1 | | |
| Reporter intensity corrected 4 1 | | |
| Reporter intensity corrected 5 1 | | |
| Reporter intensity corrected 6 1 | | |
| Reporter intensity corrected 7 1 | | |
| Reporter intensity corrected 8 1 | | |
| Reporter intensity corrected 9 1 | | |
| Reporter intensity corrected 10 1 | | |
| Reporter intensity corrected 11 1 | | |
| Reporter intensity corrected 1 2 | | |
| Reporter intensity corrected 2 2 | | |
| Reporter intensity corrected 3 2 | | |
| Reporter intensity corrected 4 2 | | |
| Reporter intensity corrected 5 2 | | |
| Reporter intensity corrected 6 2 | | |
| Reporter intensity corrected 7 2 | | |
| Reporter intensity corrected 8 2 | | |
| Reporter intensity corrected 9 2 | | |
| Reporter intensity corrected 10 2 | | |
| Reporter intensity corrected 11 2 | | |
| Reporter intensity corrected 1 3 | | |
| Reporter intensity corrected 2 3 | | |
| Reporter intensity corrected 3 3 | | |
| Reporter intensity corrected 4 3 | | |
| Reporter intensity corrected 5 3 | | |
| Reporter intensity corrected 6 3 | | |
| Reporter intensity corrected 7 3 | | |
| Reporter intensity corrected 8 3 | | |
| Reporter intensity corrected 9 3 | | |
| Reporter intensity corrected 10 3 | | |
| Reporter intensity corrected 11 3 | | |
| Reporter intensity corrected 1 4 | | |
| Reporter intensity corrected 2 4 | | |
| Reporter intensity corrected 3 4 | | |
| Reporter intensity corrected 4 4 | | |
| Reporter intensity corrected 5 4 | | |
| Reporter intensity corrected 6 4 | | |
| Reporter intensity corrected 7 4 | | |
| Reporter intensity corrected 8 4 | | |
| Reporter intensity corrected 9 4 | | |
| Reporter intensity corrected 10 4 | | |
| Reporter intensity corrected 11 4 | | |
| Reporter intensity 1 1 | | |
| Reporter intensity 2 1 | | |
| Reporter intensity 3 1 | | |
| Reporter intensity 4 1 | | |
| Reporter intensity 5 1 | | |
| Reporter intensity 6 1 | | |
| Reporter intensity 7 1 | | |
| Reporter intensity 8 1 | | |
| Reporter intensity 9 1 | | |
| Reporter intensity 10 1 | | |
| Reporter intensity 11 1 | | |
| Reporter intensity 1 2 | | |
| Reporter intensity 2 2 | | |
| Reporter intensity 3 2 | | |

| |
|---|
| Reporter intensity 4 2 |
| Reporter intensity 5 2 |
| Reporter intensity 6 2 |
| Reporter intensity 7 2 |
| Reporter intensity 8 2 |
| Reporter intensity 9 2 |
| Reporter intensity 10 2 |
| Reporter intensity 11 2 |
| Reporter intensity 1 3 |
| Reporter intensity 2 3 |
| Reporter intensity 3 3 |
| Reporter intensity 4 3 |
| Reporter intensity 5 3 |
| Reporter intensity 6 3 |
| Reporter intensity 7 3 |
| Reporter intensity 8 3 |
| Reporter intensity 9 3 |
| Reporter intensity 10 3 |
| Reporter intensity 11 3 |
| Reporter intensity 1 4 |
| Reporter intensity 2 4 |
| Reporter intensity 3 4 |
| Reporter intensity 4 4 |
| Reporter intensity 5 4 |
| Reporter intensity 6 4 |
| Reporter intensity 7 4 |
| Reporter intensity 8 4 |
| Reporter intensity 9 4 |
| Reporter intensity 10 4 |
| Reporter intensity 11 4 |
| Reporter intensity count 1 1 |
| Reporter intensity count 2 1 |
| Reporter intensity count 3 1 |
| Reporter intensity count 4 1 |
| Reporter intensity count 5 1 |
| Reporter intensity count 6 1 |
| Reporter intensity count 7 1 |
| Reporter intensity count 8 1 |
| Reporter intensity count 9 1 |
| Reporter intensity count 10 1 |
| Reporter intensity count 11 1 |
| Reporter intensity count 1 2 |
| Reporter intensity count 2 2 |
| Reporter intensity count 3 2 |
| Reporter intensity count 4 2 |
| Reporter intensity count 5 2 |
| Reporter intensity count 6 2 |
| Reporter intensity count 7 2 |
| Reporter intensity count 8 2 |
| Reporter intensity count 9 2 |
| Reporter intensity count 10 2 |
| Reporter intensity count 11 2 |
| Reporter intensity count 1 3 |
| Reporter intensity count 2 3 |
| Reporter intensity count 3 3 |
| Reporter intensity count 4 3 |
| Reporter intensity count 5 3 |
| Reporter intensity count 6 3 |
| Reporter intensity count 7 3 |
| Reporter intensity count 8 3 |
| Reporter intensity count 9 3 |
| Reporter intensity count 10 3 |
| Reporter intensity count 11 3 |
| Reporter intensity count 1 4 |

| | | |
|---|---|---|
| Reporter intensity count 2 4 | | |
| Reporter intensity count 3 4 | | |
| Reporter intensity count 4 4 | | |
| Reporter intensity count 5 4 | | |
| Reporter intensity count 6 4 | | |
| Reporter intensity count 7 4 | | |
| Reporter intensity count 8 4 | | |
| Reporter intensity count 9 4 | | |
| Reporter intensity count 10 4 | | |
| Reporter intensity count 11 4 | | |
| Sequence coverage 1 [%] | | Percentage of the sequence that is covered by the identified peptides in this sample of the longest protein sequence contained within the group. |
| Sequence coverage 2 [%] | | Percentage of the sequence that is covered by the identified peptides in this sample of the longest protein sequence contained within the group. |
| Sequence coverage 3 [%] | | Percentage of the sequence that is covered by the identified peptides in this sample of the longest protein sequence contained within the group. |
| Sequence coverage 4 [%] | | Percentage of the sequence that is covered by the identified peptides in this sample of the longest protein sequence contained within the group. |
| Intensity | | Summed up eXtracted Ion Current (XIC) of all isotopic clusters associated with the identified AA sequence. In case of a labeled experiment this is the total intensity of all the isotopic patterns in the label cluster. |
| Intensity 1 | | Summed up eXtracted Ion Current (XIC) of all isotopic clusters associated with the identified AA sequence. In case of a labeled experiment this is the total intensity of all the isotopic patterns in the label cluster. |
| Intensity 2 | | Summed up eXtracted Ion Current (XIC) of all isotopic clusters associated with the identified AA sequence. In case of a labeled experiment this is the total intensity of all the isotopic patterns in the label cluster. |
| Intensity 3 | | Summed up eXtracted Ion Current (XIC) of all isotopic clusters associated with the identified AA sequence. In case of a labeled experiment this is the total intensity of all the isotopic patterns in the label cluster. |
| Intensity 4 | | Summed up eXtracted Ion Current (XIC) of all isotopic clusters associated with the identified AA sequence. In case of a labeled experiment this is the total intensity of all the isotopic patterns in the label cluster. |
| MS/MS count | | |
| Only identified by site | | When marked with '+', this particular protein group was identified only by a modification site. |
| Reverse | | When marked with '+', this particular protein group contains no protein, made up of at least 50% of the peptides of the leading protein, with a peptide derived from the reversed part of the decoy database. These should be removed for further data analysis. The 50% rule is in place to prevent spurious protein hits to erroneously flag the protein group as reverse. |
| Potential contaminant | | When marked with '+', this particular protein group was found to be a commonly occurring contaminant. These should be removed for further data analysis. |
| id | | A unique (consecutive) identifier for each row in the proteinGroups table, which is used to cross-link the information in this file with the information stored in the other files. |
| Peptide IDs | | Identifier(s) of the associated peptide sequence(s) summary, which can be found in the file 'peptides.txt'. |
| Peptide is razor | | Indicates for each peptide ID if it is a razor or group unique peptide (true) or a non unique non razor peptide (false). |
| Mod. peptide IDs | | |
| Evidence IDs | | |
| MS/MS IDs | | |
| Best MS/MS | | The identifier of the best (in terms of quality) MS/MS scans identifying the peptides of this protein, referenced against the msms table. |
| Oxidation (M) site IDs | | Identifier(s) for site(s) associated with the protein group, which show(s) evidence of the modification, referenced against the appropriate modification site file. |
| Phospho (STY) site IDs | | Identifier(s) for site(s) associated with the protein group, which show(s) evidence of the modification, referenced against the appropriate modification site file. |
| Oxidation (M) site positions | | Positions of the sites in the leading protein of this group. |
| Phospho (STY) site positions | | Positions of the sites in the leading protein of this group. |
| Taxonomy IDs | | Taxonomy identifiers. |

### All peptides

| Name | Separator | Description |
|---|---|---|
| Raw file | | Name of the raw file the spectral data was extracted from. |
| Type | | The type of detection for the peptide. MULTI – A labeling multiplet was detected.<br>ISO – An isotope pattern was detected. |
| Charge | | The charge state of the peptide. |
| m/z | | The mass divided by the charge of the charged peptide. |
| Mass | | The mass of the neutral peptide ((m/z-proton) * charge). |
| Uncalibrated m/z | | m/z before re-calibrations have been applied. |
| Number of data points | | The number of data points (peak centroids) collected for this peptide feature. |
| Number of scans | | The number of MS scans that the 3d peaks of this peptide feature are overlapping with. |
| Number of isotopic peaks | | The number of isotopic peaks contained in this peptide feature. |
| PIF | | Short for Parent Ion Fraction; indicates the fraction the target peak makes up of the total intensity in the inclusion window. |
| Mass fractional part | | The values after the radix point (ie value - floor(value)). |
| Mass deficit | | Empirically derived deviation measure to the next nearest integer scaled to center around 0. Can be used to visually detect contaminants in a plot setting Mass against this value.<br><br>m*a+b – round(m*a+b)<br>m: the peptide mass<br>a: 0.99954<br>b: -0.04 |
| Mass precision [ppm] | | The precision of the mass detection of the peptide in parts-per-million. |
| Max intensity m/z 0 | | Summed up eXtracted Ion Current (XIC) of all isotopic clusters associated with the identified AA sequence. In case of a labeled experiment this is the total intensity of all the isotopic patterns in the label cluster. |
| Retention time | | The retention time of the peak detected for the peptide measured in minutes. |
| Retention length | | The total retention time width of the peak (last time point – first time point) in seconds. |
| Retention length (FWHM) | | The full width at half maximum value retention time width of the peak in seconds. |
| Min scan number | | The first scan number at which the peak was encountered. |
| Max scan number | | The last scan number at which the peak was encountered. |
| Identified | | When marked with '+' this particular MS/MS scan was identified as a peptide; when marked with '-' no identification was made. |
| Reverse | | When marked with '+' this particular MS/MS scan was assigned to a decoy sequence. |
| MS/MS IDs | | Unique identifier linking this identification to the MS/MS scans. |
| Sequence | | The identified AA sequence of the peptide. |
| Length | | The length of the sequence stored in the column "Sequence". |
| Modifications | | Post-translational modifications contained within the sequence. When no modifications exist, this is set to 'unmodified'.<br><br>Note: This column only set when this MS/MS spectrum has been identified. |
| Modified sequence | | Sequence representation of the peptide including location(s) of modified AAs.<br><br>Note: This column only set when this MS/MS spectrum has been identified. |
| Proteins | | Identifiers of proteins this peptide is associated with.<br><br>Note: This column only set when this MS/MS spectrum has been identified. |
| Score | | The score of the identification (higher is better). |
| PEP | | The posterior error probability of the identification (smaller is better). |
| Intensity | | Summed up eXtracted Ion Current (XIC) of all isotopic clusters associated with the identified AA sequence. In case of a labeled experiment this is the total intensity of all the isotopic patterns in the label cluster. |
| Intensities | | Elution profile. |
| Isotope pattern | | Isotope pattern. |
| MS/MS Count | | The number of MS/MS spectra recorded for the peptide. |

| | | |
|---|---|---|
| MSMS Scan Numbers | | The scan numbers where the MS/MS spectra were recorded. |
| MSMS Isotope Indices | | Indices of the isotopic peaks that the MS/MS spectra reside on. A value of 0 corresponds to the monoisotopic peak. |

### MS scans

The msScans table contains information about the full scans, which can be used to verify data quality and generated useful statistics about the interaction between the samples and LC.

| Name | Separator | Description |
|---|---|---|
| Raw file | | The name of the RAW-file the mass spectral data originates from. |
| Scan number | | The scan number (defined in the raw-file) at which the full scan was made. |
| Scan index | | The consecutive index of this full scan. |
| Retention time | | The retention time at which the full scan was made. |
| Cycle time | | The total time (full scan including the tandem MS scans) this full scan has taken up. |
| Ion injection time | | The total injection time that was required to capture the specified amount of ions. This value is limited by a maximum, which can be used to determine whether the time has maxed out (indicative of a bad acquisition). |
| Base peak intensity | | The intensity of the most intense ion in the spectrum. |
| Total ion current | | The total intensity acquired in the full scan. |
| MS/MS count | | The number of tandem MS scans that were made based on this full scan (e.g. a top 10 method selects the top 10 most intense ions in the scan and fragments those). |
| Mass calibration | | The applied mass correction in Th to the full scan. |
| Experiment | | |
| Peak length | | The average time between the start and the end of the peaks detected in the full scan. |
| Isotope pattern length | | The average time between the start and the end of the isotope patterns detected in the full scan. |
| Multiplet length | | The average time between the start and the end of the isotope patterns of the labeling multiplets detected in the full scan. |
| Peaks / s | | The average number of peaks detected per second of chromatography. |
| Single peaks / s | | The average number of single peaks detected per second of chromatography. |
| Isotope patterns / s | | The average number of isotope patterns detected per second of chromatography. |
| Single isotope patterns / s | | The average number of single isotope patterns detected per of second chromatography. |
| Multiplets / s | | The average number of labeling multiplets detected per of second chromatography. |
| Identified multiplets / s | | The percentage of labeling multiplets actually identified. |
| Multiplet identification rate [%] | | The percentage of the detected labeling multiplets that were identified. |
| Intens Comp Factor | | Taken from the Thermo RAW file. |
| CTCD Comp | | Taken from the Thermo RAW file. |
| RawOvFtT | | For Thermo Fisher only. TIC estimation done with the orbitrap cell. |
| AGC Fill | | Taken from the Thermo RAW file. |

### MZ range

| **Name** | **Separator** | **Description** |
|---|---|---|
| Raw file | | The name of the RAW-file the mass spectral data was derived from. |
| m/z | | The mass-over-charge value. |
| Peaks / Da | | The average number of peaks detected per Dalton. |
| Single peaks / Da | | The average number of single peaks detected per Dalton. |
| Isotope patterns / Da | | The average number of isotope patterns detected per Dalton. |
| Single isotope patterns / Da | | The average number of single isotope patterns detected per Dalton. |
| SILAC pairs / Da | | The average number of SILAC pairs detected per Dalton. |
| Identified SILAC pairs / Da | | The percentage of SILAC pairs actually identified. |
| SILAC identification rate [%] | | The percentage of the detected SILAC pairs that were identified. |

### MS/MS scans

| Name | Separator | Description |
|---|---|---|
| Raw file | | Name of the RAW file the spectral MS/MS data was extracted from. |
| Scan number | | RAW file derived scan number for the MS/MS spectrum. |
| Retention time | | Time point along the elution profile at which the MS/MS data was recorded. |
| Ion injection time | | The ion inject time for the MS/MS scan. This can be used to determine if this time equals to the maximum ion inject time, general indicative of a lower quality spectrum. |
| Total ion current | | The total ion current of the MS/MS scan. For Thermo data this value is calculated by summing all the intensity values found in the mass spectral data, which is different from the Xcalibur reported TIC (Xcalibur TIC is about 25% of the value reported here). |
| Collision energy | | The collision energy used for the fragmentation that resulted in this MS/MS scan. |
| Summations | | For time of flight instruments only. |
| Base peak intensity | | The intensity of the most intense ion in the spectrum. |
| Elapsed time | | The time the MS/MS scan took to complete. |
| Identified | | When marked with '+' this particular MS/MS scan was identified as a peptide; when marked with '-' no identification was made. |
| Matched | | When marked with '+' this particular MS/MS scan was retrieved by matching between runs. |
| Reverse | | When marked with '+' this particular MS/MS scan was assigned to a decoy sequence. |
| MS/MS IDs | | Unique identifier linking this identification to the MS/MS scans. |
| Sequence | | The identified AA sequence of the peptide. |
| Length | | The length of the sequence stored in the column “Sequence”. |
| Filtered peaks | | Number of peaks after the 'top X per 100 Da' filtering. |
| m/z | | Recalibrated m/z of the precursor ion. |
| Mass | | Charge corrected mass of the precursor ion. |
| Charge | | Charge state of the precursor ion. |
| Type | | The type of precursor ion as identified by MaxQuant. ISO – isotopic cluster.<br>PEAK – single peak.<br>MULTI – labeling cluster. |
| Fragmentation | | The type of fragmentation used to create the MS/MS spectrum.<br>CID – Collision Induced Dissociation.<br>HCD – High energy Collision induced Dissociation.<br>ETD – Electron Transfer Dissociation. |
| Mass analyzer | | The mass analyzer used to record the MS/MS spectrum. ITMS – Ion trap.<br>FTMS – Fourier transform ICR or orbitrap cell.<br>TOF – Time of flight. |
| Parent intensity fraction | | The percentage the parent ion intensity makes up of the total intensity in the selection window. |
| Fraction of total spectrum | | The percentage the parent ion intensity makes up of the total intensity of the whole MS spectrum. |
| Base peak fraction | | The percentage the parent ion intensity in comparison to the highest peak in he MS spectrum. |
| Precursor full scan number | | The full scan number where the precursor ion was selected for fragmentation. |
| Precursor intensity | | The intensity of the precursor ion at the scan number it was selected. |
| Precursor apex fraction | | The fraction the intensity of the precursor ion makes up of the peak (apex) intensity. |
| Precursor apex offset | | How many full scans the precursor ion is offset from the peak (apex) position. |
| Precursor apex offset time | | How much time the precursor ion is offset from the peak (apex) position. |
| Scan event number | | This number indicates which MS/MS scan this one is in the consecutive order of the MS/MS scans that are acquired after an MS scan. |
| Modifications | | Post-translational modifications contained within the sequence. When no modifications exist, this is set to ‘unmodified’.<br><br>Note: This column only set when this MS/MS spectrum has been identified. |

| | | |
|---|---|---|
| Modified sequence | | Sequence representation of the peptide including location(s) of modified AAs.<br><br>Note: This column only set when this MS/MS spectrum has been identified. |
| Proteins | | Identifiers of proteins this peptide is associated with.<br><br>Note: This column only set when this MS/MS spectrum has been identified. |
| Score | | The score of the identification (higher is better). |
| PEP | | The posterior error probability of the identification (smaller is better). |
| Experiment | | |
| Reporter intensity corrected 1 | | |
| Reporter intensity corrected 2 | | |
| Reporter intensity corrected 3 | | |
| Reporter intensity corrected 4 | | |
| Reporter intensity corrected 5 | | |
| Reporter intensity corrected 6 | | |
| Reporter intensity corrected 7 | | |
| Reporter intensity corrected 8 | | |
| Reporter intensity corrected 9 | | |
| Reporter intensity corrected 10 | | |
| Reporter intensity corrected 11 | | |
| Reporter intensity 1 | | |
| Reporter intensity 2 | | |
| Reporter intensity 3 | | |
| Reporter intensity 4 | | |
| Reporter intensity 5 | | |
| Reporter intensity 6 | | |
| Reporter intensity 7 | | |
| Reporter intensity 8 | | |
| Reporter intensity 9 | | |
| Reporter intensity 10 | | |
| Reporter intensity 11 | | |
| Reporter PIF | | |
| Reporter fraction | | |
| Intens Comp Factor | | Taken from the Thermo RAW file. |
| CTCD Comp | | Taken from the Thermo RAW file. |
| RawOvFtT | | For Thermo Fisher only. TIC estimation done with the orbitrap cell. |
| AGC Fill | | Taken from the Thermo RAW file. |
| Scan index | | Consecutive index of the MS/MS spectrum. |
| MS scan index | | Consecutive index of the MS spectrum prior to this MS/MS spectrum. |
| MS scan number | | Scan number of the MS spectrum prior to this MS/MS spectrum. |

# MS/MS

| Name | Separator | Description |
|---|---|---|
| Raw file | | The name of the RAW file the mass spectral data was read from. |
| Scan number | | The RAW-file derived scan number of the MS/MS spectrum. |
| Scan index | | The consecutive index of the MS/MS spectrum. |
| Sequence | | The identified AA sequence of the peptide. |
| Length | | The length of the sequence stored in the column "Sequence". |
| Missed cleavages | | Number of missed enzymatic cleavages. |
| Modifications | | Post-translational modifications contained within the identified peptide sequence. |
| Modified sequence | | Sequence representation including the post-translational modifications (abbreviation of the modification in brackets before the modified AA). The sequence is always surrounded by underscore characters ('_'). |
| Oxidation (M) Probabilities | | Sequence representation of the peptide including PTM positioning probabilities ([0..1], where 1 is best match) for 'Oxidation (M)'. |
| Phospho (STY) Probabilities | | Sequence representation of the peptide including PTM positioning probabilities ([0..1], where 1 is best match) for 'Phospho (STY)'. |
| Oxidation (M) Score diffs | | |
| Phospho (STY) Score diffs | | |
| Acetyl (Protein N-term) | | |
| Oxidation (M) | | |
| Phospho (STY) | | |
| Proteins | | The identifiers of the proteins the identified peptide is associated with. |
| Gene Names | | Names of genes the identified peptide is associated with. |
| Protein Names | | Descriptions of the proteins the identified peptide is associated with. |
| Charge | | The charge state of the precursor ion. |
| Fragmentation | | The type of fragmentation used to create the MS/MS spectrum.<br>CID – Collision Induced Dissociation.<br>HCD – High energy Collision induced Dissociation.<br>ETD – Electron Transfer Dissociation. |
| Mass analyzer | | The mass analyzer used to record the MS/MS spectrum. ITMS – Ion trap.<br>FTMS – Fourier transform ICR or orbitrap cell.<br>TOF – Time of flight. |
| Type | | The type of precursor ion as identified by MaxQuant. ISO – isotopic cluster.<br>PEAK – single peak.<br>MULTI – labeling cluster. |
| Scan event number | | |
| Isotope index | | |
| m/z | | The mass-over-charge of the precursor ion. |
| Mass | | The charge corrected mass of the precursor ion. |
| Mass error [ppm] | | Mass error of the recalibrated mass-over-charge value of the precursor ion in comparison to the predicted monoisotopic mass of the identified peptide sequence expressed in parts per million. |
| Mass error [Da] | | Mass error of the recalibrated mass-over-charge value of the precursor ion in comparison to the predicted monoisotopic mass of the identified peptide sequence expressed in atomic mass units. |
| Simple mass error [ppm] | | |
| Retention time | | The uncalibrated retention time in minutes where the MS/MS spectrum has been acquired. |
| PEP | | Posterior Error Probability of the identification. This value essentially operates as a p-value, where smaller is more significant. |
| Score | | Andromeda score for the best associated MS/MS spectrum. |
| Delta score | | Score difference to the second best identified peptide with a different amino acid sequence. |
| Score diff | | Score difference to the second best positioning of modifications identified peptide with the same amino acid sequence. |
| Localization prob | | |
| Combinatorics | | Number of possible distributions of the modifications over the peptide sequence. |

| | | |
|---|---|---|
| PIF | | Short for Parent Ion Fraction; indicates the fraction the target peak makes up of the total intensity in the inclusion window. |
| Fraction of total spectrum | | The percentage the parent ion intensity makes up of the total intensity of the whole spectrum. |
| Base peak fraction | | The percentage the parent ion intensity in comparison to the highest peak in he MS spectrum. |
| Precursor full scan number | | The full scan number where the precursor ion was selected for fragmentation. |
| Precursor Intensity | | The intensity of the precursor ion at the scan number it was selected. |
| Precursor apex fraction | | The fraction the intensity of the precursor ion makes up of the peak (apex) intensity. |
| Precursor apex offset | | How many full scans the precursor ion is offset from the peak (apex) position. |
| Precursor apex offset time | | How much time the precursor ion is offset from the peak (apex) position. |
| Diagnostic peak Phospho (STY) Y | | |
| Matches | | The species of the peaks in the fragmentation spectrum after TopN filtering. |
| Intensities | | The intensities of the peaks in the fragmentation spectrum after TopN filtering. |
| Mass deviations [Da] | | The mass deviation of each peak in the fragmentation spectrum in absolute mass units. |
| Mass deviations [ppm] | | The mass deviation of each peak in the fragmentation spectrum in parts per million. |
| Masses | | The masses-over-charge of the peaks in the fragmentation spectrum. |
| Number of matches | | The number of peaks matching to the predicted fragmentation spectrum. |
| Intensity coverage | | The fraction of intensity in the MS/MS spectrum that is annotated. |
| Peak coverage | | The fraction of peaks in the MS/MS spectrum that are annotated. |
| Neutral loss level | | How many neutral losses were applied to each fragment in the Andromeda scoring. |
| ETD identification type | | For ETD spectra several different combinations of ion series are scored. Here the highest scoring combination is indicated |
| Reverse | | When marked with '+', this particular peptide was found to be part of a protein derived from the reversed part of the decoy database. These should be removed for further data analysis. |
| All scores | | |
| All sequences | | |
| All modified sequences | | |
| Reporter intensity corrected 1 | | |
| Reporter intensity corrected 2 | | |
| Reporter intensity corrected 3 | | |
| Reporter intensity corrected 4 | | |
| Reporter intensity corrected 5 | | |
| Reporter intensity corrected 6 | | |
| Reporter intensity corrected 7 | | |
| Reporter intensity corrected 8 | | |
| Reporter intensity corrected 9 | | |
| Reporter intensity corrected 10 | | |
| Reporter intensity corrected 11 | | |
| Reporter intensity 1 | | |
| Reporter intensity 2 | | |
| Reporter intensity 3 | | |
| Reporter intensity 4 | | |
| Reporter intensity 5 | | |
| Reporter intensity 6 | | |
| Reporter intensity 7 | | |
| Reporter intensity 8 | | |
| Reporter intensity 9 | | |
| Reporter intensity 10 | | |
| Reporter intensity 11 | | |
| Reporter mass deviation [mDa] 1 | | |
| Reporter mass deviation [mDa] 2 | | |
| Reporter mass deviation [mDa] 3 | | |
| Reporter mass deviation [mDa] 4 | | |
| Reporter mass deviation [mDa] 5 | | |

| | | |
|---|---|---|
| Reporter mass deviation [mDa] 6 | | |
| Reporter mass deviation [mDa] 7 | | |
| Reporter mass deviation [mDa] 8 | | |
| Reporter mass deviation [mDa] 9 | | |
| Reporter mass deviation [mDa] 10 | | |
| Reporter mass deviation [mDa] 11 | | |
| Reporter PIF | | |
| Reporter fraction | | |
| id | | A unique (consecutive) identifier for each row in the msms table, which is used to cross-link the information in this file with the information stored in the other files. |
| Protein group IDs | | The identifier of the protein-group this redundant peptide sequence is associated with, which can be used to look up the extended protein information in the file 'proteinGroups.txt'.<br>As a single peptide can be linked to multiple proteins (e.g. in the case of razor-proteins), multiple id's can be stored here separated by a semicolon.<br>As a protein can be identified by multiple peptides, the same id can be found in different rows. |
| Peptide ID | | The identifier of the non-redundant peptide sequence. |
| Mod. peptide ID | | Identifier of the associated modification summary stored in the file 'modificationSpecificPeptides.txt'. |
| Evidence ID | | Identifier of the associated evidence stored in the file 'evidence.txt'. |
| Oxidation (M) site IDs | | Identifier of the associated entry stored in the file 'Oxidation (M)Sites.txt'. |
| Phospho (STY) site IDs | | Identifier of the associated entry stored in the file 'Phospho (STY)Sites.txt'. |
